# Genetic tool development in marine protists: Emerging model organisms for experimental cell biology

**DOI:** 10.1101/718239

**Authors:** Drahomíra Faktorová, R. Ellen R. Nisbet, José A. Fernández Robledo, Elena Casacuberta, Lisa Sudek, Andrew E. Allen, Manuel Ares, Cristina Aresté, Cecilia Balestreri, Adrian C. Barbrook, Patrick Beardslee, Sara Bender, David S. Booth, François-Yves Bouget, Chris Bowler, Susana A. Breglia, Colin Brownlee, Gertraud Burger, Heriberto Cerutti, Rachele Cesaroni, Miguel A. Chiurillo, Thomas Clemente, Duncan B. Coles, Jackie L. Collier, Elizabeth C. Cooney, Kathryn Coyne, Roberto Docampo, Christopher L. Dupont, Virginia Edgcomb, Elin Einarsson, Pía A. Elustondo, Fernan Federici, Veronica Freire-Beneitez, Nastasia J. Freyria, Kodai Fukuda, Paulo A. García, Peter R. Girguis, Fatma Gomaa, Sebastian G. Gornik, Jian Guo, Vladimír Hampl, Yutaka Hanawa, Esteban R. Haro-Contreras, Elisabeth Hehenberger, Andrea Highfield, Yoshihisa Hirakawa, Amanda Hopes, Christopher J. Howe, Ian Hu, Jorge Ibañez, Nicholas A.T. Irwin, Yuu Ishii, Natalia Ewa Janowicz, Adam C. Jones, Ambar Kachale, Konomi Fujimura-Kamada, Binnypreet Kaur, Jonathan Z. Kaye, Eleanna Kazana, Patrick J. Keeling, Nicole King, Lawrence A. Klobutcher, Noelia Lander, Imen Lassadi, Zhuhong Li, Senjie Lin, Jean-Claude Lozano, Fulei Luan, Shinichiro Maruyama, Tamara Matute, Cristina Miceli, Jun Minagawa, Mark Moosburner, Sebastián R. Najle, Deepak Nanjappa, Isabel C. Nimmo, Luke Noble, Anna M.G. Novák Vanclová, Mariusz Nowacki, Isaac Nuñez, Arnab Pain, Angela Piersanti, Sandra Pucciarelli, Jan Pyrih, Joshua S. Rest, Mariana Rius, Deborah Robertson, Albane Ruaud, Iñaki Ruiz-Trillo, Monika A. Sigg, Pamela A. Silver, Claudio H. Slamovits, G. Jason Smith, Brittany N. Sprecher, Rowena Stern, Estienne C. Swart, Anastasios D. Tsaousis, Lev Tsypin, Aaron Turkewitz, Jernej Turnšek, Matus Valach, Valérie Vergé, Peter von Dassow, Tobias von der Haar, Ross F. Waller, Lu Wang, Xiaoxue Wen, Glen Wheeler, April Woods, Huan Zhang, Thomas Mock, Alexandra Z. Worden, Julius Lukeš

## Abstract

Diverse microbial ecosystems underpin life in the sea. Among these microbes are many unicellular eukaryotes that span the diversity of the eukaryotic tree of life. However, genetic tractability has been limited to a few species, which do not represent eukaryotic diversity or environmentally relevant taxa. Here, we report on the development of genetic tools in a range of protists primarily from marine environments. We present evidence for foreign DNA delivery and expression in 13 species never before transformed and advancement of tools for 8 other species, as well as potential reasons for why transformation of yet another 17 species tested was not achieved. Our resource in genetic manipulation will provide insights into the ancestral eukaryotic lifeforms, general eukaryote cell biology, protein diversification and the evolution of cellular pathways.

## INTRODUCTION

The ocean represents the largest continuous planetary ecosystem, hosting an enormous variety of organisms, which include microscopic biota such as unicellular eukaryotes (protists). Despite their small size, protists play key roles in marine biogeochemical cycles and harbor tremendous evolutionary diversity^1,2^. Notwithstanding their importance for understanding the evolution of life on Earth and their role in marine food webs, as well as driving biogeochemical cycles to maintain habitability, little is known about their cell biology including reproduction, metabolism, and signalling^3^. Most of the biological knowledge available is based on comparison of proteins from cultured species to homologs in genetically tractable model taxa^4–7^. A major impediment to understanding the cell biology of these diverse eukaryotes is that protocols for genetic modification are only available for a small number of species^8,9^ that represent neither the most ecologically relevant protists nor the breadth of eukaryotic diversity.

The development of genetic tools requires reliable information about gene organisation and regulation of the emergent model species. Over the last decade, genome^4–6^ and transcriptome sequencing initiatives^7^ have resulted in nearly 120 million unigenes being identified in protists^10^, which facilitates the developments of genetic tools used for model species Insights from these studies enabled the phylogenetically-informed approach^7^ for selecting and developing key marine protists into model systems in the Environmental Model Systems (EMS) Project presented herein. Forty-one research groups took part in the EMS Project, a collaborative effort resulting in the development of genetic tools that significantly expand the number of eukaryotic lineages that can be manipulated, and which encompass multiple ecologically important marine protists.

Here, we summarize detailed methodological achievements and analyse results to provide a synthetic ‘Transformation Roadmap’ for creating new microeukaryotic model systems. Although the organisms reported here are diverse, the paths to overcome difficulties share similarities, highlighting the importance of building a well-connected community to overcome technical challenges and accelerate the development of genetic tools. The 13 emerging model species presented herein, and the collective set of genetic tools from the overall collaborative project, will not only extend our knowledge of marine cell biology, evolution and functional biodiversity, but also serve as platforms to advance protistan biotechnology.

## RESULTS

### Overview of taxa in the EMS Initiative

Taxa were selected from multiple eukaryotic supergroups^1,7^ to maximize the potential of cellular biology and to evaluate the numerous unigenes with unknown functions found in marine protists (**Fig. 1**). Prior to the EMS initiative, reproducible transformation of marine protists was limited to only a few species such as *Thalassiosira pseudonana, Phaeodactylum tricornutum*, and *Ostreococcus tauri* (**Suppl. Table 1**). The EMS initiative included 39 species, specifically, 6 archaeplastids, 2 haptophytes, 2 rhizarians, 9 stramenopiles, 12 alveolates, 4 discobans, and 4 opisthokonts (**Fig. 1**). Most of these taxa were isolated from coastal habitats, the focus area of several culture collections^7^. More than 50% of the selected species are considered photoautotrophs, with another 35% divided between heterotrophic osmotrophs and phagotrophs, the remainder being predatory mixotrophs. Almost 20% of the chosen species are symbionts and/or parasites of marine plants or animals, 5% are associated with detritus, and several are responsible for harmful algal blooms (**Suppl. Table 2**).

**Fig. 1.**
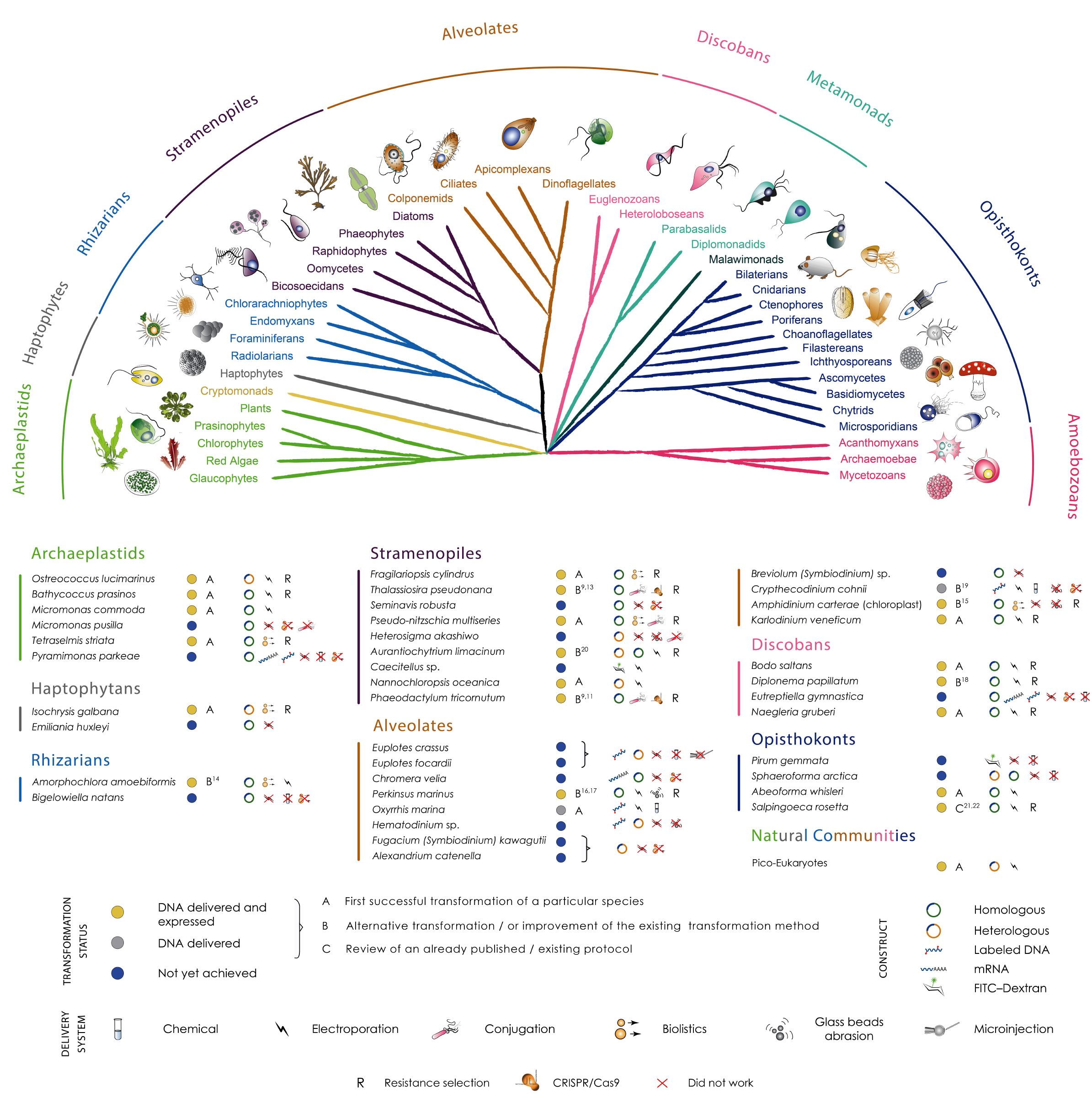
Phylogenetic relationships and transformation status of marine protists. A schematic view of the eukaryotic tree of life with effigies of main representatives. Colour-coordinated species we have attempted to genetically modify are listed below. Current transformability status is schematized in circles indicating: DNA delivered and shown to be expressed (yellow; for details see text and Table 1); DNA delivered, but no expression seen (grey); no successful transformation achieved despite efforts (blue). The details of transformation of species that belong to “DNA delivered” and “Not achieved yet” categories are described in the **Suppl. Table 5**.

While some transformation systems for protists have been developed in the past^8,9,11^, the challenge for this initiative was to develop genetic tools for species which not only require different cultivation conditions but are also phenotypically diverse. It should be noted that not all major lineages were explored. For example, amoebozoans did not feature in this aquatic-focused initiative, in part because they tend to be most important in soils, at least based on current knowledge, and manipulation systems exist for members of this eukaryotic supergroup, such as *Dictyostelium discoideum^12^*. The overall EMS initiative outcomes are summarized in Figure 1 and Table 1. We provide detailed protocols for 13 taxa, for which no transformation systems have been previously reported (Category A), and 8 taxa, for which existing protocols^9,11,13–19,20,21^ were advanced (Category B) (**Figs. 2, 3 and 4; Table 1; Suppl. Tables 1-5 and Online Methods**). We also review an already published EMS transformation protocol^22^ in 1 species (Category C), and we discuss unsuccessful transformation attempts for 17 additional taxa (**Fig. 1; Online Methods**). Finally, we synthesize our findings in a roadmap for the development of transformation systems in protists (**Fig. 5**).

**Table 1:**
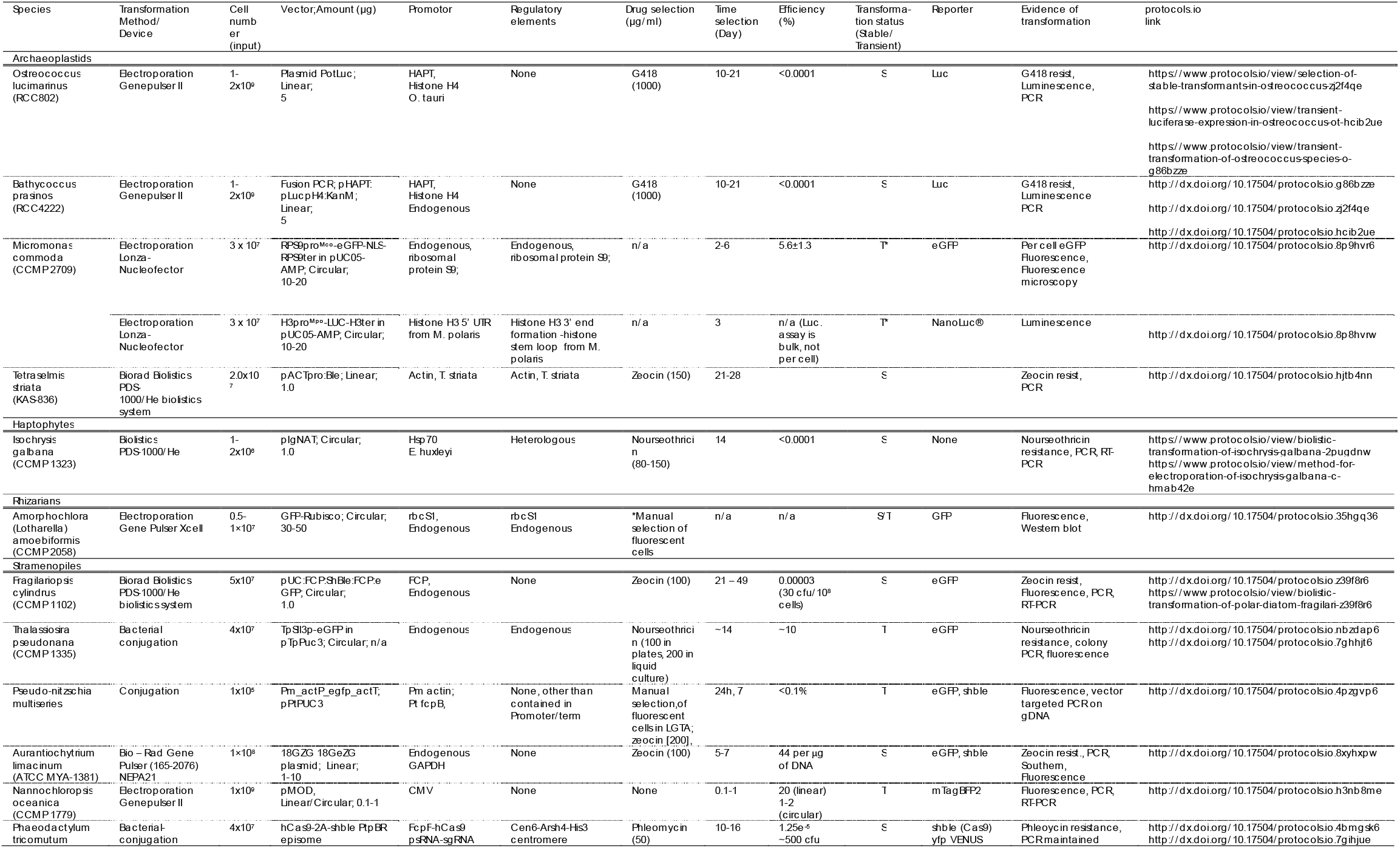
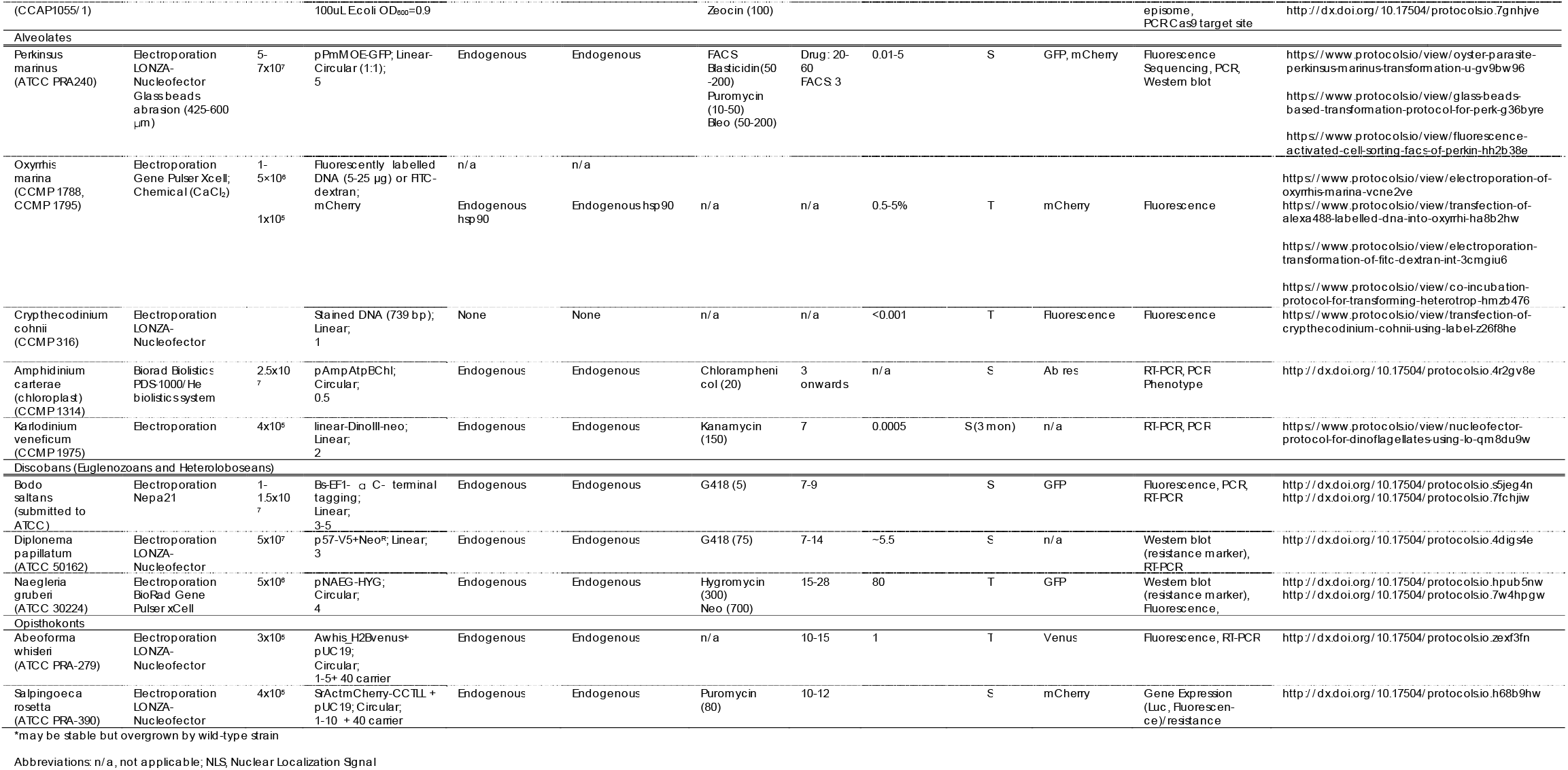
Parameters used for successful transformation as shown in Figs. 2, 3 and 4. For additional information see **Suppl. Tables 5, 7 and 8** and **Suppl. Notes 1** and protocols.io. For contacting laboratories working with particular species, see details given in **Suppl. Table 6**.

**Fig. 2.**
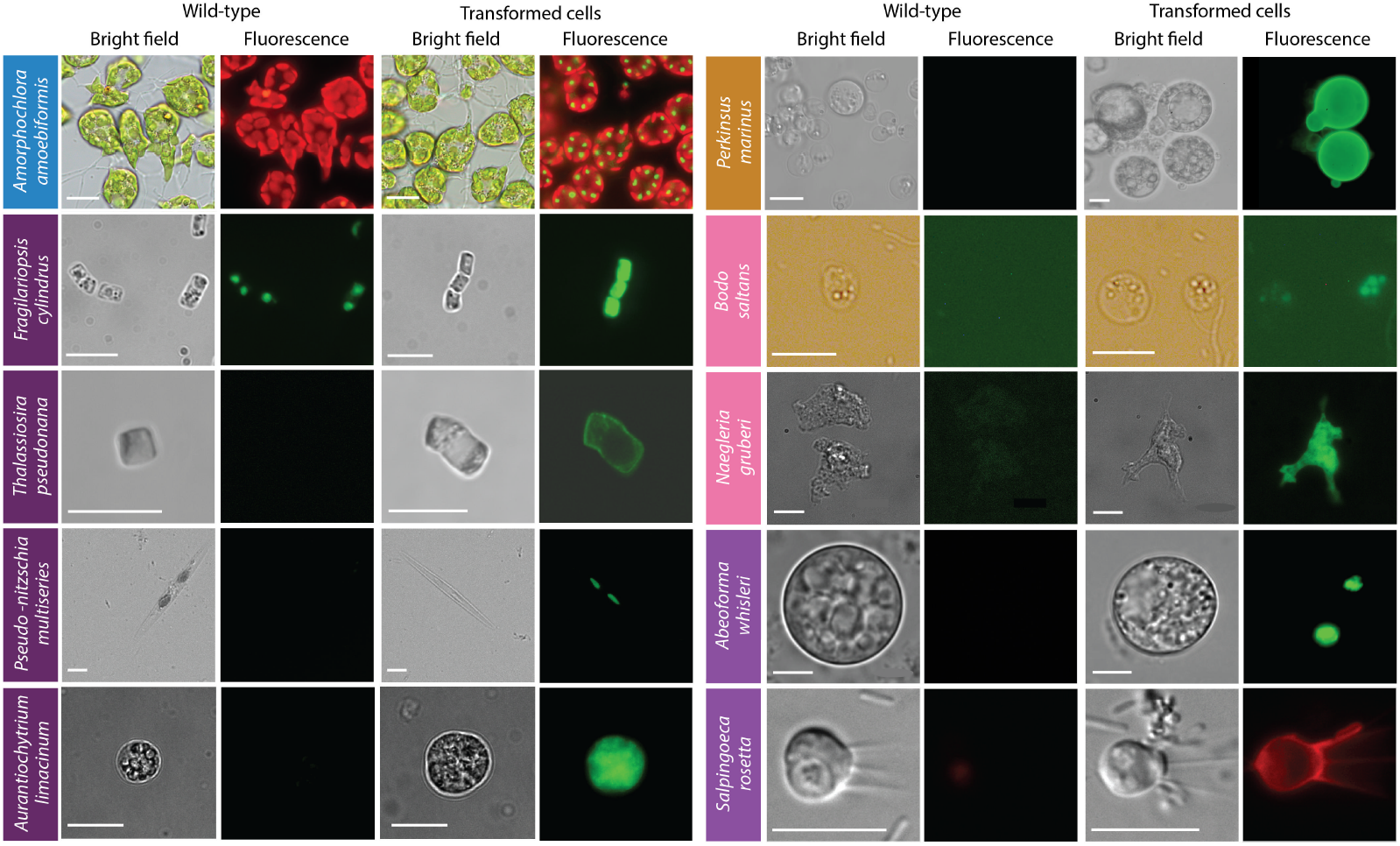
Epifluorescence micrographs of transformed marine protists. Representative images of transformants and wild-type cell lines of 10 selected protist species. Colored boxes behind species names refer to phylogenetic supergroup assignments given in Fig. 1. Representative data of at least two independent experiments are shown. The fluorescent images show the expression of individual fluorescent marker genes introduced via transformation for all organisms shown, except in the case of *A. amoebiformis*. For the latter, red depicts the natural autofluorescence of photosynthetic pigments in the cell, while the additional green spheres in the transformant fluorescence panel shows introduced GFP fluorescence (see **Suppl. Fig. 15C** for a trace of these different regions in the cell). Scale bars are as follows: 10 μm for *A. amoebiformis*, *T. pseudonana*, *A. limacinum*, *B. saltans*, *N. gruberi*, *A. whisleri*, and *S. rosetta*; 15 μm for *P. marinus*; 20 μm for *F. cylindrus*; 100 μm for *P. multiseries*.

**Fig. 3.**
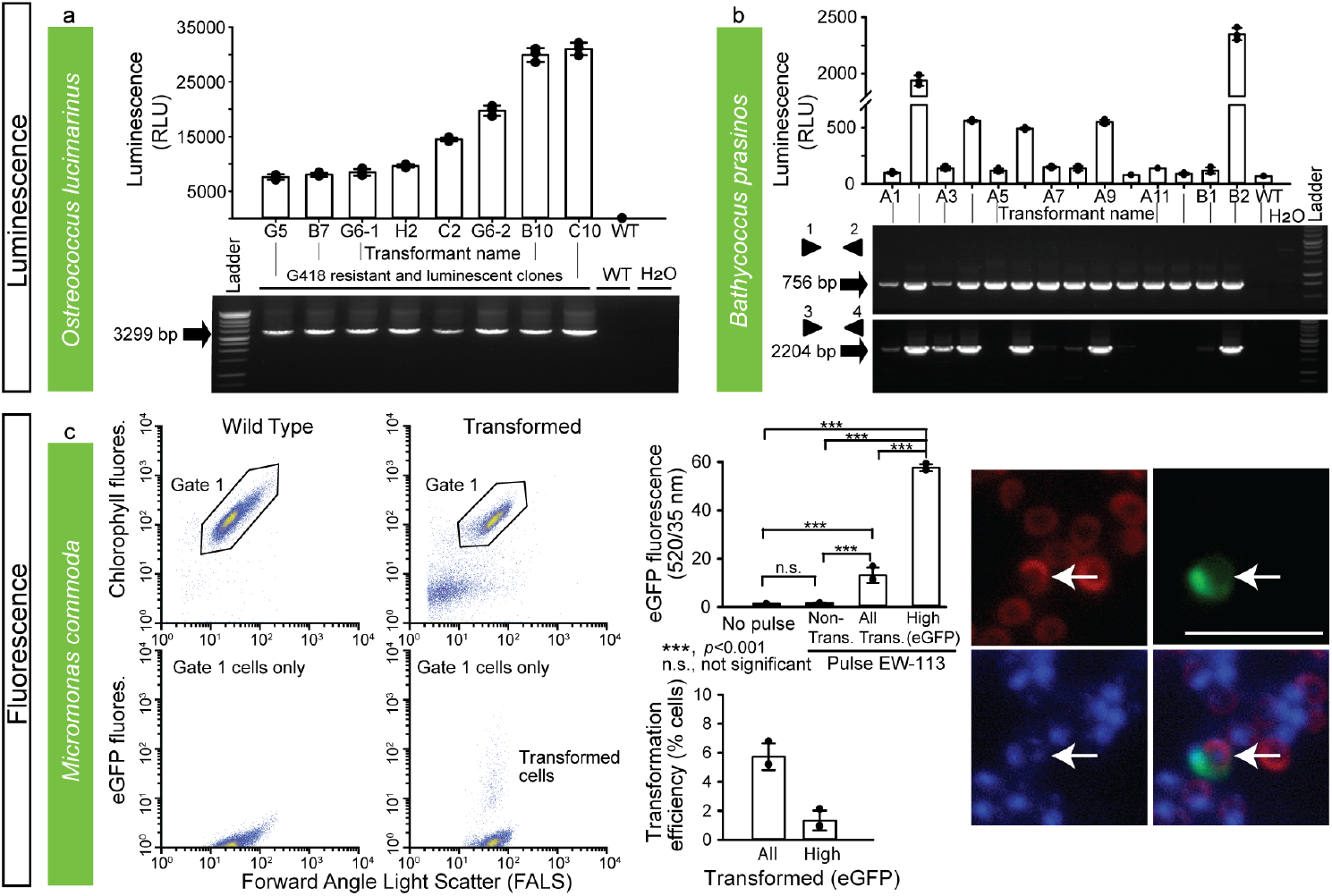
Various methods were used to demonstrate successful transformation in different archaeplastid species –. Luminescence and Fluorescence. Luminescence and Fluorescence (by FACS and epifluorescence microscopy) were used to verify expression of introduced constructs in three archaeplastids - *O. lucimarinus*, *B. prasinos*, and *M. commoda* (**a, b, c**). For the latter, red in the image depicts the natural autofluorescence of photosynthetic pigments in the cell, while green shows introduced eGFP fluorescence and blue shows the DAPI stained nucleus; the overlay shows colocalization of eGFP and nucleus signals. See Supplementary Fig. 15d for a trace of these different regions in the cell. NS, not significant; trans., transformed. Representative data of at least two independent experiments are shown. For detailed figure description see **Suppl. Notes 2**.

**Fig. 4.**
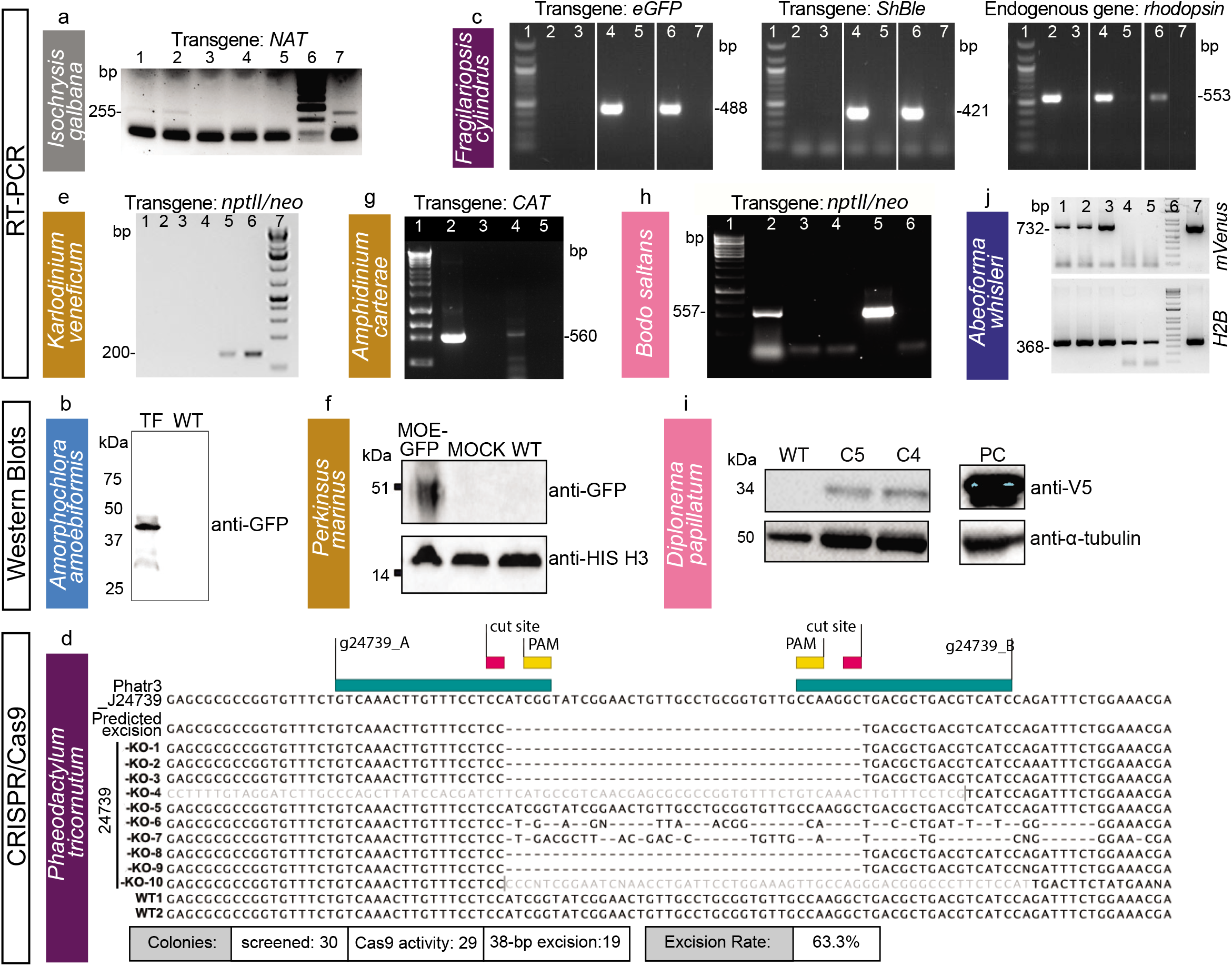
Various methods were used to demonstrate successful transformation in different species -. RT-PCR, western blot and sequencing. Western blot, RT-PCR or sequencing (in case of Cas9-induced excision by CRISPR) were used to verify expression of introduced constructs in one haptophyte – *I. galbana* (**a**), one rhizarian – *A. amoebiformis* (**b**), two stramenopiles – *F. cylindrus* and *P. tricornutum* (**c, d**), three alveolates – *K. veneficum, P. marinus* and *A. carterae* (**e, f, g**), two discobans – *B. saltans* and *D. papillatum* (**h, i**) and one opisthokont – *A. whisleri* (**j**). Note that *nptII/neo* is used synonymously with amino 3’-glycosyl phosphotransferase gene (*aph*(*3*’)) conferring resistance to kanamycin and neomycin. Representative data of at least two independent experiments are shown. For detailed figure description see **Suppl. Notes 2**.

**Fig. 5.**
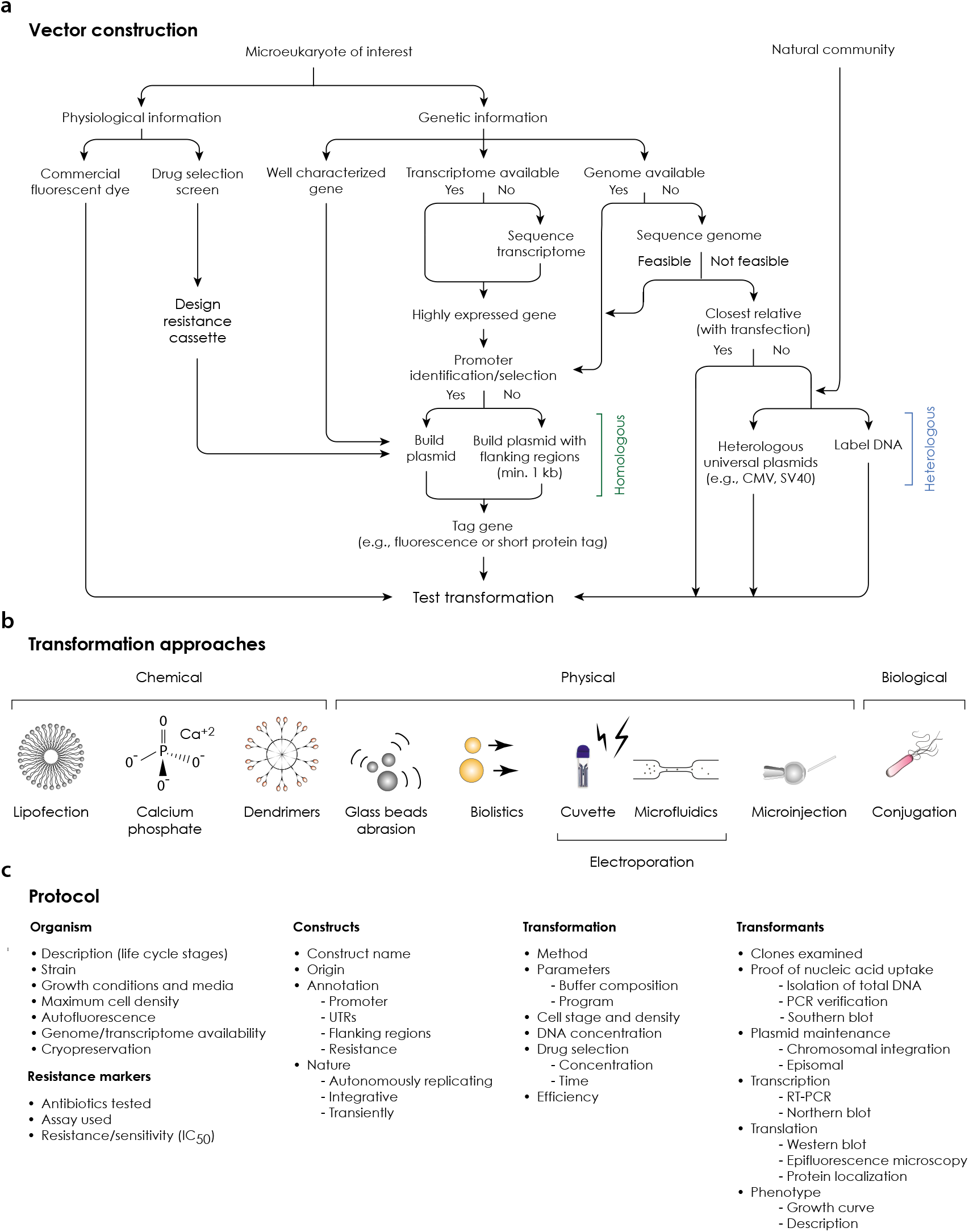
‘Transformation Roadmap’ for the creation of genetically tractable protists. (**a**) **Vector design and construction for microeukaryotes of interest and a natural community**. (**b**) **Transformation approaches**. Different symbols represent methods (e.g. chemical, physical or biological) for introducing DNA/RNA/protein into a living cell. (**c**) **Protocol**. Key methodological steps for successful transformation are listed in an abbreviated form (for particular examples, see Table 1 and text).

#### Archaeplastids

Prasinophytes are important marine green algae distributed from polar to tropical regions. They form a sister group to chlorophyte algae, and together, these two groups branch adjacent to land plants, collectively comprising the Viridiplantae, which are part of the Archaeplastida^1,23^ (**Fig. 1**). Genome sequences are available for the picoprasinophytes (<3 μm cell diameter) tested herein, specifically, *Micromonas commoda, M. pusilla*, *Ostreococcus lucimarinus* and *Bathycoccus prasinos*. As part of the EMS initiative, we report on the first genetic tools for *Bathycoccus*, a scaled, non-motile genus, and *Micromonas*, a motile, naked genus with larger genomes than *Bathycoccus* and *Ostreococcus*^22^. We also report on the first genetic tools for *Tetraselmis striata* and *Ostreococcus lucimarinus*. The latter was transformed based on an adapted homologous recombination system for *Ostreococcus tauri*^24,25^.

*O. lucimarinus* (RCC802) and *B. prasinos* (RCC4222) were transformed using protocols adapted from *O. tauri^24,25^*. Briefly, using electroporation for transfer of exogenous genes, *O. lucimarinus* was transformed using a DNA fragment encoding the *O. tauri* high-affinity phosphate transporter (*HAPT*) gene fused to a luciferase gene and a kanamycin selection marker (**Table 1; Suppl. Table 3**), which resulted in transient luciferase expression 24 h after electroporation (**Table 1; Fig. 3a**). After 2 weeks of growth in low-melting agarose plates containing G418 (1 mg/ml), 480 colonies were obtained, picked, and grown in artificial seawater with the antibiotic neomycin. Of these, 76 displayed luminescence ≥ 2.5 fold above background (80 Relative Luminescence Units, RLU), with widely variable levels (200 to 31020 RLU), likely reflecting either variations in the site of integration and/or the number of integrated genes (**Fig. 3a; Suppl. Fig. 1; Online Methods**).

The *O. tauri* construct did not work in *B. prasinos*, while the use of the *B. prasinos* histone *H4* and *HAPT* sequences in an otherwise identical construct and conditions was successful. Although luciferase expression was not detected 24 h after electroporation, 48 G418-resistant colonies were obtained 2 weeks later, 20 being luminescent when grown in liquid medium. Analysis of 14 resistant transformants revealed that the luciferase sequence was integrated into the genome of 5 luminescent clones, and one non-luminescent clone (**Fig. 3b; Online Methods**), suggesting that the chromatin context at integration sites in the latter was not favourable to luciferase expression.

Although transformation methods successful for *Bathycoccus* and *Ostreococcus* failed in *Micromonas*, Lonza nucleofection was successful with *M. commoda* (CCMP2709) (**Table 1; Fig. 3c**) using 2 different codon-optimized plasmids, one encoding the luciferase gene (NanoLuc, Promega) flanked by an exogenous promoter and terminator sequence from the 5’- and 3’-untranslated regions (UTRs) of histone *H3* in *Micromonas polaris* (CCMP2099), and the other encoding an enhanced fluorescent protein (*eGFP*) gene flanked by endogenous promoter and terminator sequences from ribosomal protein S9 (**Suppl. Table 5**). Sensitivities to antibiotics were established (**Suppl. Table 3**). Constructs did not include a selectable marker, as we aimed to introduce and express foreign DNA while developing conditions suitable for transfection that supported robust growth in this cell wall-lacking protist (**Table 1**). Transformants revealed a significantly higher level of eGFP fluorescence than wild type cells, with 1.3% of the population showing fluorescence per cell 45-fold higher than both the non-transformed portion of the culture and the wild type cells (**Fig. 3c; Online Methods**). Additionally, the RLU was 1500-fold higher than controls when using the luciferase-bearing construct, such that multiple experiments with both plasmids confirmed expression of exogenous genes in *M. commoda*.

*T. striata* (KAS-836) was transformed using microprojectile bombardment (**Suppl. Fig. 2a**). Two selectable marker genes were tested, consisting of a putative promoter and 5’ UTR sequences from the *T. striata* actin gene and either the coding sequences of the *Streptoalloteichus hindustanus* bleomycin gene (conferring resistance to zeocin) or the *Streptomyces hygroscopicus bar* gene (conferring resistance to glufosinate) (**Table 1; Suppl. Fig. 2a; Online Methods**). The terminator sequence was obtained from the *T. striata* glyceraldehyde-3-phosphate dehydrogenase gene. Linearized plasmids were coated on gold particles and introduced into *T. striata* cells by using the PDS-1000/He Particle Delivery System (BioRad). Transformants were successfully selected on half-strength f/2 at 50% salinity agar plates containing either 150 μg/ml zeocin or 150 μg/ml glufosinate.

#### Haptophytes (incertae sedis)

Haptophytes are a group of photosynthetic protists that are abundant in marine environments and include the major calcifying lineage, the coccolithophores. Genome sequences are available for *Emiliania huxleyi*^6^ and *Chrysochromulina tobin^26^*, and there is one report of nuclear transformation of a calcifying coccolithophore species^27^ but transformation of *E. huxleyi*, the most prominent coccolithophore, has not been achieved yet^27^. Here, as part of the EMS initiative, a stable nuclear transformation system was developed for *Isochrysis galbana*, a species that lacks coccoliths, but represents an important feedstock for shellfish aquaculture^28^.

*I. galbana* (CCMP1323) was transformed by biolistic bombardment with the pIgNAT vector, which contains nourseothricin *N*-acetyltransferase (*NAT*, for nourseothricin resistance) driven by the promoter and terminator of *Hsp70* from *E. huxleyi* (CCMP1516). Twenty four hours after bombardment, cells were transferred to liquid f/2 medium at 50% salinity containing 80 μg/ml nourseothricin (NTC) and left to grow for 2-3 weeks to select for transformants (**Table 1**). The presence of *NAT* in NTC-resistant cells was verified by PCR and RT-PCR (**Fig. 4a; Suppl. Fig. 2b; Online Methods**) and the sequence was verified. To confirm NTC resistance was a stable phenotype, cells were sub-cultured every 2-4 weeks at progressively higher NTC concentrations (up to 150 μg/ml) in the above-mentioned media. Cells remained resistant to NTC for approximately 6 months, as confirmed by PCR screening to identify the presence of the *NAT* gene.

#### Rhizarians

Rhizarians include diverse non-photosynthetic protists, as well as the photosynthetic chlorarachniophytes that acquired a plastid *via* secondary endosymbiosis of a green alga^4^. Uniquely, they represent an intermediate stage of the endosymbiotic process, since their plastids still harbor a relict nucleus (nucleomorph). Here, we report on an advanced transformation protocol for the chlorarachniophyte *Amorphochlora (Lotharella) amoebiformis* for which low-efficiency transient transformation has previously been achieved using particle bombardment^14^.

*A. amoebiformis* (CCMP2058) cells were resuspended in 100 μl of Gene Pulse Electroporation Buffer (BioRad) with 20-50 μg of the reporter plasmid encoding eGFP-RubisCO fusion protein under the control of the native *rbcS1* promoter and subjected to electroporation (**Table 1**). Cells were immediately transferred to fresh ESM medium and incubated for 24 h. Transformation efficiency was estimated by the fraction of cells expressing eGFP, resulting in 0.03-0.1% efficiency, as enumerated by microscopy, showing an efficiency up to 1000-fold higher than in the previous study^14^ (**Table 1**). Stable transformants were generated by manual isolation using a micropipette, and a transformed line has maintained eGFP fluorescence for at least 10 months without antibiotic selection (**Fig. 2; Fig. 4b; Online Methods**).

#### Stramenopiles

Stramenopiles are a diverse lineage harboring important photoautotrophic, mixotrophic (combining photosynthetic and phagotrophic nutrition) and heterotrophic taxa. As the most studied class in this lineage, diatoms (Bacillariophyceae) were early targets for the development of reverse genetics tool^11,29^. Diatoms are estimated to contribute approximately 20% of annual carbon fixation^30^ and, like several other algal lineages, are used in bioengineering applications and biofuels^31^. Although other cold-adapted eukaryotes have yet to be transformed, here we present a protocol for the Antarctic diatom *Fragilariopsis cylindrus*^32^. A first transformation protocol has also been developed for *Pseudo-nitzschia multiseries*, a toxin-producing diatom^33^. Here we also present work for non-diatom stramenopiles, including the first transformation protocol for the eustigmatophyte *Nannochloropsis oceanica*, and an alternative protocol for the labyrinthulomycete *Aurantiochytrium limacinum*^20^, both of which are used for biotechnological applications. Furthermore, we report on advances for CRISPR/Cas-driven gene knockouts in *Phaeodactylum tricornutum*^8,13^ and a more efficient bacterial conjugation system for *Thalassiosira pseudonana*^13^.

Microparticle bombardment was used on *F. cylindrus* (CCMP1102) which was grown, processed and maintained at 4 °C in 24 h light. Exponential phase cells were harvested onto a 1.2 μm membrane filter which was then placed on an 1.5% agar Aquil plate for bombardment with beads coated with a plasmid containing zeocin resistance and *eGFP*, both controlled by an endogenous fucoxanthin chlorophyll *a/c* binding protein (FCP) promoter and terminator (**Table 1; Suppl. Table 3; Online Methods**)^34^. Transformation was performed using 0.7 μm tungsten particles and the biolistic particle delivery system PDS1000/He (BioRad). Rupture discs for 1350 and 1550 pounds per square inch (psi) gave the highest colony numbers with efficiencies of 20.7 colony forming units (cfu)/10^8^ cells and 30 cfu/10^8^ cells, respectively. Following bombardment, the filter was turned upside down and left to recover for 24 h on the plate, then cells were rinsed from the plate/filter and spread across five 0.8% agar Aquil plates with 100 μg/ml zeocin. Colonies appeared 3 to 5 weeks later. PCR on genomic DNA showed that 100% and 60% of colonies screened positive for the bleomycin gene (*ShBle*) for zeocin resistance and the gene encoding eGFP, respectively. Confirmed by fluorescence-activated cell sorting (FACS) and microscopy, eGFP was localized to the cytosol and was distinguishable from plastid autofluorescence (**Fig. 2**). Additional confirmation by PCR and RT-PCR (**Fig. 4c**) revealed that the *ShBle* and *eGFP* genes were present in the genomes of transformants after multiple transfers (> 10) 2 years later, indicating long-term stability.

Bacterial conjugation methods were improved in *T. pseudonana* (CCMP1335) using the silaffin precursor *TpSil3p* (**Table 1; Online Methods**) as the target gene. *TpSil3p* was fused to *eGFP* flanked by an FCP promoter and terminator, cloned into a pTpPuc3 episomal backbone, and transformed into mobilization plasmid-containing EPI300 *E. coli* cells (Lucigen). The donor cells were grown in SOC medium at 37 °C until OD_600_ of 0.3–0.4, centrifuged and resuspended in 267 μl SOC medium. Next, 200 μl of donor cells were mixed with *T. pseudonana* cells, co-cultured on pre-dried 1% agar plates, dark incubated at 30 °C for 90 min, then at 18 °C in constant light for 4 h, followed by selection in 0.25% agar plates containing 100 μg/ml NTC. Colonies were observed after 2 weeks, inoculated into 300 μl L1 medium and supplemented with 200 μg/ml NTC to reduce the number of false positives. Positive transformants were identified by colony PCR screening (**Suppl. Fig. 3**) and epifluorescence microscopy (**Fig. 2**).

The diatom *Pseudo-nitzschia multiseries* (15093C) and other members of this genus form buoyant linear chains with overlapping cell tips during active growth, and were unconducive to punctate colony formation on agar, where their growth is generally poor. To address this challenge, a low-gelation-temperature agarose seawater medium (LGTA) was developed to facilitate growth, antibiotic selection, and cell recovery. *P. multiseries* exhibited growth inhibition at relatively low concentrations under NTC, formaldehyde, and zeocin (**Suppl. Table 3**). Biolistic transformation of two other *Pseudo-nitzschia* species had been demonstrated at low efficiency^35^. To complement this approach and explore potentially higher efficiency methods for transformation with diatom episomal plasmids, we modified the existing conjugation-based method^13^. The published conjugation protocol was modified to enhance *P. multiseries* post-conjugation viability by reducing SOC content. An episomal version of the Pm_actP_egfp_actT expression cassette was transfected into *E. coli* EPI300+pTAMOB and used for conjugation (**Table 1; Online Methods**). After 48 h in L1 medium, cells were plated in LGTA and eGFP-positive cells were observed 7 days later (**Fig. 2**). PCR revealed the presence of plasmids in all eGFP positive colonies (**Suppl. Fig. 4**). Similarly, conjugation with the episome pPtPUC3 (bleomycin selection marker)-containing bacterial donors was followed under zeocin selection (200 μg/ml). After 7 days, only viable cells (based on bright chlorophyll fluorescence) contained the episome, as confirmed by PCR. Propagation of transformants after the first medium transfer (under selection) has so far been unsuccessful.

Stable transformation of *A. limacinum* (ATCC MYA-1381) was achieved by knock-in of a resistance cassette composed of *ShBle* driven by 1.3 kb promoter and 1.0 kb terminator regions of the endogenous glyceraldehyde-3-phosphate dehydrogenase gene carried in a pUC19-based plasmid (18GZG) along with the native 18S rRNA gene, and by knock-in of a similar construct containing a *eGFP:ShBle* fusion (**Suppl. Fig. 5**). Approximately 1 x 10^8^ cells were electroporated, adapting the electroporation protocol used for *Schizochytrium*^36^. The highest transformation efficiency was achieved using 1 μg of linearized 18GZG plasmid with 2 pulses, resulting in a time constant of ~5 ms (**Table 1; Online Methods**). Expression of the fusion protein was confirmed by both the zeocin-resistance phenotype and the detection of eGFP (**Fig. 2**). Six 18GZG transformants derived from uncut and linearized plasmids were examined in detail. All maintained antibiotic resistance throughout 13 serial transfers, first in selective, and subsequently in non-selective media, and then again in selective medium. Integration of the plasmid into the genome was confirmed by PCR as well as by Southern blots using a digoxigenin-labeled *ShBle* gene probe, showing that 4 transformants had integrations by single homologous recombination, while in 2 transformants, additional copies of the antibiotic resistance cassette were integrated by non-homologous recombination elsewhere in the genome (**Suppl. Fig. 5**).

Electroporation of *N. oceanica* (CCMP1779) was optimized based on observation of cells treated with fluorescein-conjugated 2000 kDa dextran and subsequent survival (**Table 1; Online Methods**). A sorbitol concentration of 800 mM and electroporation at between 5 and 9 kV/cm resulted in highest cell recovery. These conditions were used during introduction of plasmids containing the gene for the blue fluorescent reporter *mTagBFP2* under the control of cytomegalovirus (CMV), the cauliflower Mosaic Virus 35S, or the VCP1 promoter previously described from *Nannochloropsis* sp.^37^. Transient expression of blue fluorescence (compared to cells electroporated simultaneously under the same conditions without plasmid) appeared within 2 h, lasted for at least 24 h, and disappeared by 48 h in subsets of cells electroporated with *mTagBFP2* under the control of CMV (**Suppl. Fig. 6**). The transient transformation was more effective when a linearized plasmid was used compared to a circular plasmid (**Table 1**). VCP1 did not induce blue fluorescence with a circular plasmid, while 35S gave inconsistent results with either circularized or linearized plasmids.

For *P. tricornutum* (CCAP1055/1), we adapted the CRISPR/Cas9 system^8^ for multiplexed targeted mutagenesis. Bacterial conjugation^13^ was used to deliver an episome that contained a Cas9 cassette and 2 single-guide RNA (sgRNA) expression cassettes designed to excise a 38 bp-long domain from the coding region of a nuclear-encoded, chloroplastic glutamate synthase (*Phatr3_J24739*) and introduce an in-frame stop codon after strand ligation (**Table 1; Online Methods**). The GoldenGate assembly was used to clone 2 expression cassettes carrying sgRNAs into a *P. tricornutum* episome that contained a *Cas9-2A-ShBle* expression cassette and the centromeric region CenArsHis (**Suppl. Fig. 7**). After their addition to a *P. tricornutum* culture, plates were incubated in a growth chamber under standard growth conditions for 2 days and transformed *P. tricornutum* colonies began to appear after 2 weeks. Only colonies maintaining *Cas9-2A-ShBle* sequence on the delivered episome were able to grow on selection plates because *Cas9* and *ShBle* were transcriptionally fused by the 2A peptide^38^ (**Suppl. Fig. 7**). Gel electrophoresis migration and sequencing of the genomic target loci confirmed the 38 bp-long excision and premature stop codon (**Fig. 4d**).

#### Alveolates

This species-rich and diverse group is comprised of ciliates, apicomplexans, and dinoflagellates (**Fig. 1**). As a link between apicomplexan parasites and dinoflagellate algae, perkinsids are key for understanding the evolution of parasitism, and also have potential biomedical applications^17^. Techniques currently exist for transformation of only a small number of ciliates, perkinsids and apicomplexans^39^. Here, we present a first transformation protocol for *Karlodinium veneficum* (CCMP1975), a phagotrophic mixotroph that produces fish-killing karlotoxins^40^. Experiments were also performed on *Oxyrrhis marina* (CCMP 1788/CCMP 1795), a basal-branching phagotroph that lacks photosynthetic plastids and *Crypthecodinium cohnii* (CCMP 316), a heterotroph used in food supplements. For both of these taxa first evidence of DNA delivery was achieved (**Table 1; Suppl. Results; Supp. Fig. 15; Online Methods**), a goal recently achieved for *C. cohnii* using electroporation^19^. Additionally, we report on improved transformation systems for *Perkinsus marinus* (PRA240) and *Amphidinium carterae* (CCMP1314) chloroplast, published recently as part of the EMS initiative^15^.

*K. veneficum* (CCMP1975) was transformed based on electroporation and cloning the selectable marker gene aminoglycoside 3’-phosphotransferase (*nptII/neo*; note that *nptII/neo* is used synonymously with amino 3’-glycosyl phosphotransferase gene conferring resistance to kanamycin, neomycin, paromomycin, ribostamycin, butirosin and gentamicin B) into the backbone of the dinoflagellate-specific expression vector DinoIII-*neo*^41^, which confers resistance to neomycin and kanamycin (**Table 1**). In brief, DinoIII-*neo* was linearized and electroporated using the Nucleofector optimization pulse codes, buffer SF/Solution I (Lonza), and 2 μg/μl of linearized DinoIII-*neo*. Electroporated cells were selected under 150 μg/ml kanamycin 3 days post-electroporation. Fresh seawater with kanamycin was added every 2 weeks to the cultures and new subcultures were inoculated monthly. After 3 months, DNA and RNA were isolated from the resistant cultures as previously reported^42^ and cDNA was synthesized using random hexamers. Out of 16 transformations, 2 cell lines (CA-137, DS-138) showed stable growth under kanamycin selection. CA-137 developed dense cultures after 3 months, and the resistance gene was detected in both DNA and RNA by nested PCR and RT-PCR, respectively (**Fig. 4e; Suppl. Fig. 8; Online Methods**).

We improved the transformation protocol^16,17^ of *P. marinus*, a pathogen of marine mollusks, fish, and amphibians^43^ (**Suppl. Table 5**). We co-expressed 2 genes and efficiently selected transient and stable transformants using FACS (**Table 1; Figs. 2** and **4f; Suppl. Fig. 9; Online Methods**). In addition, we established the integration profile of ectopic DNA once introduced into the *P. marinus* genome. We did not see evidence of integration through homologous recombination and observed a propensity for plasmid fragmentation and integration within transposable elements sites. An optimized alternative protocol for transformation using glass bead abrasion was also developed. Two versions of the previously published *Moe* gene promoter^16^ were tested. Whereas the 1.0 kb promoter version induced expression after 2 or 3 days, the truncated version (0.5 kb) took 7 days for expression to be detected. Resistance genes to zeocin, blasticidin and puromycin have all been shown to confer resistance to transformed *P. marinus*; however, selection regimes are still relatively slow and inefficient, indicating further room for improvement^17^.

We also report a new vector for the transformation of the *A. carterae* chloroplast, a photosynthetic dinoflagellate. *A. carterae*, like other dinoflagellates with a peridinin-containing chloroplast, contains a fragmented chloroplast genome made up of multiple plasmid-like minicircles^40^. The previous transformation protocols made use of this to introduce two vectors based on the psbA minicircle^15^. Here, we show that other minicircles are also suitable for use as vectors. We created a new artificial minicircle, using the atpB minicircle as a backbone, but replacing the *atpB* gene with a codon-optimized chloramphenicol acetyl transferase (**Table 1; Online Methods**). This circular vector was introduced by biolistics to *A. carterae* (**Suppl. Fig. 10a**). Following selection with chloramphenicol, we were able to detect transcription of the chloramphenicol acetyl transferase gene via RT-PCR (**Fig. 4g**). This result suggests that all 20 or so minicircles in the dinoflagellate chloroplast genome would be suitable for use as artificial minicircles, thus providing a large pool of potential vectors.

#### Discobans

This diverse group, recently split into Discoba and Metamonada^44^, includes heterotrophs, photoautotrophs, predatory mixotrophs, as well parasites. Discobans include parasitic kinetoplastids with clinical significance, such as *Trypanosoma brucei, T. cruzi* and *Leishmania* spp., for which efficient transformation protocols are available^45^. However, such protocols are missing for aquatic species. Here, we describe available transformation protocols for the kinetoplastid *Bodo saltans* and the heterolobosean *Naegleria gruberi*. The former was isolated from a lake, but identical 18S rRNA gene sequences have been reported from marine environments^46^. The latter is a freshwater protist that represents a model organism for closely related marine heterolobosean amoebas. Furthermore, we provide advanced methods that build on prior EMS results^18^ for the diplonemid *Diplonema papillatum*.

*B. saltans* (ATCC 30904) was transformed with a plasmid containing a cassette designed to fuse an endogenous *EF-1α* gene with *eGFP* for C-terminal tagging. This cassette includes downstream of *eGFP*, a *B. saltans* tubulin intergenic region followed by the selectable marker *nptII/neo* gene, conferring resistance to neomycin. *EF-1α* genes exist in tandem repeats. The homologous regions that flank the cassette were chosen as targets for inducing homology-directed repair, however, they target only one copy of the gene. As transcription in *B. saltans* is polycistronic^46^, insertion of the tubulin intergenic region into the plasmid is essential for polyadenylation of the *EF1-αGFP* fusion and *trans*-splicing of the *nptII/neo* gene (**Suppl. Table 5**). Selection of transfected cells began with 2 μg/ml of neomycin added 24 h after electroporation, and this concentration was gradually increased over 2 weeks to 5 μg/ml (**Table 1; Online Methods**). Cells were washed and subcultured into fresh selection medium every 4 days, and neomycin-resistant cells emerged 7-9 days post-electroporation. The eGFP signal was detected 2 days post-electroporation, albeit with low intensity. This may be due to the inefficient translation of *eGFP* since it has not been codon-optimized for *B. saltans* (**Fig. 2**). Genotyping analysis 9 months post-transfection confirmed the presence of the *nptII/neo* gene and at least partial plasmid sequence (**Fig. 4h; Suppl. Fig. 10b**). However, plasmid integration into the *B. saltans* genome through homologous recombination is still unconfirmed. This suggests either off-target plasmid integration or episomal maintenance.

For *N. gruberi* (ATCC 30224) two plasmids were designed. The first one carried the hygromycin B resistance gene (*hph*) with an actin promoter and terminator, along with an HA-tagged *eGFP* driven by the ubiquitin promoter and terminator. The second plasmid carried the *nptII/neo* gene instead. For each individual circular plasmid, 4 μg was electroporated (**Table 1; Online Methods**). About 48 h after electroporation, dead cells were removed from the suspension and viable cells were washed with PBS. Afterwards, 300 μg/ml of hygromycin B or 700 μg/ml of neomycin was added to the fresh media. One to 4 weeks later, resistant clones were recovered and expression of *eGFP* and/or hygromycin was confirmed by western blotting (**Suppl. Fig. 11**). Expression of *eGFP* was observed by epifluorescence microscopy (**Fig. 2; Suppl. Fig. 11**) with ~80% of transformants maintaining hygromycin B or neomycin resistance in addition to expressing *eGFP*.

*D. papillatum* (ATCC 50162) was transformed by electroporation using 3 μg a *SwaI*-linearised fragment (cut from p57-V5+NeoR plasmid) containing the V5-tagged *nptII/neo* gene flanked by partial regulatory sequences derived from the hexokinase gene of the kinetoplastid *Blastocrithidia* (strain p57) (**Table 1; Online Methods**) using a published protocol^18^. About 18 h after electroporation, 75 μg/ml G418 was added to the medium and after 2 weeks 7 neomycin-resistant clones were recovered. Transcription of *nptII/neo* was verified in 4 clones by RT-PCR (**Suppl. Fig. 12**) and the expression of the tagged *nptII/neo* protein was confirmed in 2 clones by western blotting using the α-V5 antibody (**Fig. 4i**).

### Opisthokonts

The opisthokont clade Holozoa includes animals and their closest unicellular relatives choanoflagellates, filastereans, ichthyosporeans, and corallochytreans. The establishment of genetic tools in non-metazoan holozoans promises to help illuminate the cellular and genetic foundations of animal multicellularity^47^. Genomic and transcriptomic data are available for multiple representatives characterized by diverse cell morphologies, some of which can even form multicellular structures^46^. Untill recently, only transient transformations had been achieved for some opistokonts such as the filasterean *Capsaspora owczarzaki*^48^, the ichthyosporean *Creolimax fragrantissima*^49^ and the choanoflagellate *Salpingoeca rosetta*^21^. Through the EMS initiative, we report on the first evidence for transient transformation of the ichthyosporean *Abeoforma whisleri*, isolated from the digestive tract of mussels, and review a recently published stable transformation protocol for *S. rosetta* achieved by using the selectable puromycin N-acetyl-transferase gene (**Fig. 2**)^22^.

All *A. whisleri* life stages are highly sensitive to a variety of methods for transformation. However, we developed a 4D-nucleofection-based protocol using 16-well strips, wherein PBS-washed cells were resuspended in 20 μl of buffer P3 (Lonza) containing 40 μg of carrier plasmid (empty pUC19) and 1-5 μg of the reporter plasmid (*A. whisleri* H2B fused to mVenus fluorescent protein, mVFP) (**Table 1; Online Methods**), and subjected to code EN-138 (Lonza). Immediately after the pulse, cells were recovered by adding 80 μl of marine broth (Gibco) prior to plating in 12-well culture plates previously filled with 1 ml marine broth. After 24 h, ~1% of the culture was transformed based on the fraction of cells expressing *mVFP* in the nucleus (**Figs. 2** and **4j**).

### Microbial eukaryotes in natural planktonic communities

Model organisms are typically selected based on criteria such as relative ease of isolation and asexual cultivation in the laboratory, however these attributes may not correlate with the capacity for uptake and expression of the exogenous DNA. We explored whether natural marine planktonic pico- and nanoeukaryote communities would take up DNA in a culture-independent setting. Microbial plankton from natural seawater was concentrated and electroporated with plasmids containing mTagBFP2 under the control of CMV or 35S promoters (**Suppl. Results; Online Methods**). In most trials, blue fluorescent cells were rare if detected at all (compared to control samples). However, in one natural community tested, a photosynthetic picoeukaryote population exhibited up to 50% of cells with transient expression of blue fluorescence when the CMV promoter was used (**Suppl. Fig. 13**). This suggests it might be possible to selectively culture eukaryotic microorganisms based on capacity to express exogenous DNA.

## Discussion

The collaborative effort by the EMS initiative facilitated identification and optimization of the steps required to create new protist model systems, which culminated in the synthetic ‘Transformation Roadmap’ (**Fig. 5**). Our genetic manipulation systems for aquatic (largely marine) protists will enable deeper insights into their cell biology, with potentially valuable outcomes for aquatic sciences, evolutionary studies, nanotechnology, biotechnology, medicine, and pharmacology. Successes and failures with selectable markers, transformation conditions, and reporters were qualitatively compared across species (**Suppl. Tables 3** and **4**; **Table 1; Figs. 2, 3** and **4; Online Methods**).

For some of the selected species, the first step was to identify cultivation conditions for robust growth in the laboratory to either generate high cell densities or large culture volumes for obtaining sufficient biomass required for a variety of molecular biology experiments. Unlike established microbial model species, cultivation of marine protists can be challenging especially under axenic conditions or for predatory taxa that require co-cultivation with their prey. Nevertheless, 13 out of 35 species were rendered axenic prior to the development of transformation protocols. For the remaining species, we were unable to remove bacteria and therefore had to make sure that transformation signals were coming from the targeted protist rather than contaminants (**Suppl. Table 2**). Subsequent steps included the identification of suitable antibiotics and their corresponding selectable markers (**Table 1; Suppl. Table 3**), conditions for introducing exogenous DNA (**Table 1; Suppl. Table 4**), and selection of promoter and terminator sequences for designing transformation vectors (**Table 1; Online Methods – Suppl. Table 5 and Suppl. Notes 1**).

As exemplified in the new model systems provided herein (**Table 1; Figs. 2, 3** and **4**), a variety of methods were used to test whether exogenous DNA was integrated into the genome or maintained as a plasmid, and whether the introduced genes were expressed. Approaches to show the former included inverse PCR, Southern blotting and whole genome sequencing, whereas approaches to demonstrate the latter included various combinations of PCR, RT-PCR, western blotting, epifluorescence microscopy, FACS, antibody-based methods, and/or growth assays in the presence of antibiotics to confirm transcription and translation of introduced selection and reporter genes (e.g. *eGFP, YFP, mCherry*). For fluorescent markers, it was first ensured that the wild type, or manipulated controls cells, had no signals conflicting with the marker (**Figs. 2** and **3c**), an important step because photosynthetic protists contain chlorophyll and other autofluorescent pigments. Overall transformation outcomes for each species were parsed into three groups according to the level of success or lack thereof (A = first transformation protocol for a given species; B = advanced protocol based on prior work; C = published protocol based on the EMS initiative) and are discussed according to their phylogenetic position (**Fig. 1**).

Our studies did not result in a universally applicable protocol because transformability and a range of other key conditions varied greatly across taxa and approaches, such as intrinsic features of the genome and differences in cellular structure and morphology. In general, electroporation proved to be the most common method for introducing exogenous DNA stably into cells. This approach was utilized for naked cells and protoplasts, yet frequently also worked, albeit with lower efficiency, on cells protected by cell walls. Linearized plasmids were most effective for delivery, and 5’ and 3’ UTRs-containing promotors of highly expressed endogenous genes provided the strongest expression of selective reporters and markers. If successful, teams usually continued with fluorescence-based methods. Furthermore, large amounts of carrier DNA usually facilitated successful initial transformations (e.g. *M. commoda, A. whisleri*) or improved existing protocols (*S. rosetta*^21^). We also provide the contact details of all co-authors who are assigned to particular species (**Suppl. Table 6**).

Some lineages were difficult to transform, especially dinoflagellates and coccolithophores. Here, even if DNA appeared to be delivered (**Suppl. Table 5**), expression of the transformed genes could not be confirmed. Examples include the dinoflagellates *C. cohnii* and *Symbiodinium microadriaticum*, and the coccolithophore *E. huxleyi*. Thus, at least these 3 species need concerted future efforts.

The combination of results presented herein together with previously published protocols from the EMS initiative^50^ significantly expands the segment of extant eukaryotic diversity amenable to reverse genetics approaches. Out of the 39 microbial eukaryotes selected for the initiative, exogenous DNA was delivered and expressed in more than 50% of them. The transformation systems enable us for the first time to shed light on the function of species-specific genes, which likely reflect key adaptations to specific niches in dynamic ocean habitats.

## ACKNOWLEDGEMENTS

We thank M. Salisbury and D. Lacono, C. Poirier, M. Hamilton, C. Eckmann, H. Igel, C. Yung and K. Hoadley for assistance; and V. K. Nagarajan, M. Accerbi, and P. J. Green who carried out *Agrobacterium* studies in *Heterosigma akashiwo*; and N. Kraeva, C. Bianchi and V. Yurchenko for the help with designing the p57-V5+NeoR construct. We are also grateful to the protocols.io team (L. Teytelman and A. Broellochs) for their support and thank four anonymous reviewers for their constructive criticisms of our manuscript. This collaborative effort was supported by the Gordon and Betty Moore Foundation EMS Program of the Marine Microbiology Initiative (grant numbers: GBMF4972 and 4972.01 to F.Y.B.; GBMF4970 and 4970.01 to M.A. & A.Z.W.; GBMF3788 to A.Z.W.; GBMF 4968 and 4968.01 to H.C.; GBMF4984 to V.H.; GBMF4974 and 4974.01 to C. Brownlee; GBMF4964 to Y. Hirakawa; GBMF4961 to T. Mock; GBMF4958 to P.S.; GBMF4957 to A.T.; GBMF4960 to G.J.S.; GBMF4979 to K.C.; GBMF4982 and 4982.01 to J.L.C.; GBMF4964 to P.J.K.; GBMF4981 to P.V.D.; GBMF5006 to A.E.A.; GBMF4986 to C.M.; GBMF4962 to J.A.F.R.; GBMF4980 and 4980.01 to S.L.; GBMF 4977 and 4977.01 to R.F.W.; GBMF4962.01 to C.H.S.; GBMF4985 to J.M.; GBMF4976 and 4976.01 to C.H.; GBMF4963 and 4963.01 to V.E.; GBMF5007 to C.L.D; GBMF4983 and 4983.01 to J.L; GBMF4975 and 4975.01 to A.T.; GBMF4973 and 4973.01 to I.R.T. and GBMF4965 to N.K.), by The Leverhulme Trust (RPG-2017-364) (to T. Mock and A. Hopes) and by ERD Funds (16_019/0000759) from the Czech Ministry of Education (to JL).

## AUTHOR CONTRIBUTIONS

The project was conceived and designed by A.C.J., J.Z.K., S.B., D.F., J.L., R.E.R.N., J.A.F.R., E.C., L.S., A.Z.W., T. Mock, A.E.A., F.Y.B, C. Brownlee, C. Bowler, H.C., T.C., J. L.C., K.C., C.L.D., V.E., V.H., Y. Hirakawa., C.J.H., P.J.K., N.K., S.L., C.M., J.M., I.R.T., P.A.S., C.H.S., G.J.S., A.D.T., P.V.D., A.T. and R.F.W. Data analysis was carried out by M.A.J., C.A., C. Balestreri, A.C.B., P.B., D.S.B., S.A.B., G.B., R.C., M.A.C., D.B.C., E.C.C., R.D., E.E., P.A.E., F.F., V.F.B., N.J.F., K.F., P.A.G., P.R.G., F.G., S.G.G., J.G., Y. Hanawa, E.R.H.C., E.H., A. Highfield, A. Hopes, I.H., J.I., N.A.T.I., Y.I., N.E.J., A.K., K. F.K., B.K., E.K., L.A.K., N.L., I.L., Z.L., J.C.L., F.L., S.M., T. Matute, M.M., S.R.N., D.N., I.C.N., L.N., A.M.G.N.V., M.N., I.N., A. Pain, A. Piersanti, S.P., J.P., J.S.R., M.R., D.R., A.R., M.A.S., E.C.S., B.N.S., R.S., T.V.H., L.T., J.T., M.V., V.V., L.W., X.W., G.W., A.W. and H.Z. The manuscript was written by D.F., R.E.R.N., J.A.F.R., E.C., L.S., T. Mock, A.Z.W. and J.L. with input from all authors.

## ONLINE METHODS

### Studied species and used transformation methods

For the full list of vector sequences and maps see **Suppl. Notes 1** and for detailed description of **Figs. 3 and 4** see **Suppl. Notes 2**. Antibiotic concentrations effective for selection of transformats can be found in **Suppl. Table 3**, the details of the transformation methods applied to this study in **Suppl. Table 4** and contact details for individual laboratories in **Suppl. Table 6**. Full list of protists (including details of culture collection) and links to the complete step-by steps transformation protocols and published vector sequences are listed in **Suppl. Table 5**. The protocols.io links listed in **Table 1** and **Suppl. Table 5** are summarized in **Suppl. Tables 7** and **8**.

### Life Sciences Reporting Summary

Further information is available in the Life Sciences Reporting Summary.

### Data availability

The data that support the findings of this study are available from the corresponding authors as well as the other authors upon request (for the contacts see **Suppl. Table 6**). Source data for **Fig. 3, Fig. 4** and **Suppl. Figs. 9B, C; 11A** and **12B, C** are available online.

