## Supplementary material for "Genetic tool development in marine protists: Emerging model organisms for experimental cell biology": Suppl. Fig.

**Suppl. Fig. 1.**

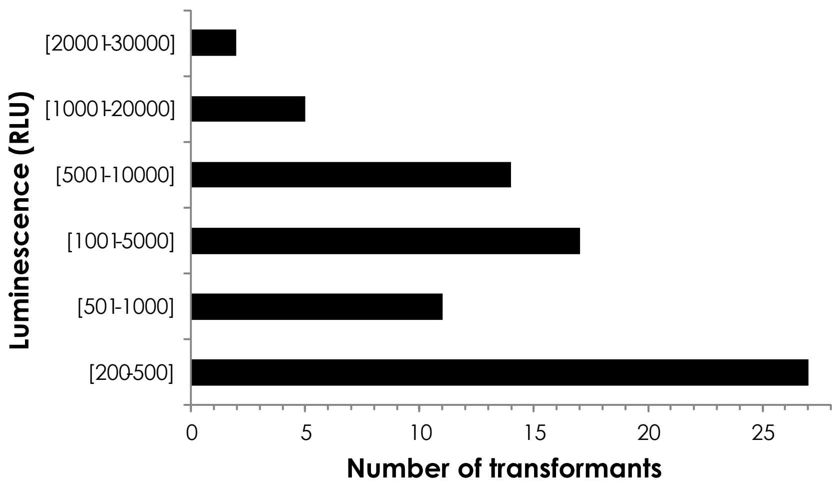


**Stable transformation of *Ostreococcus lucimarinus* RCC802**
Luminescence of 76 RCC802 transformed by HAPT:luc transgene exhibiting luminescence levels at least two fold above RCC802 WT control transformed without DNA (80 RLU). The range of luminescence (RLU) is shown on the *Y* axis. In total 480 transformants resistant to G418 were obtained in two independent electroporation experiments.

**Transient expression and stable transformation of selected Mamiellophyceae**

| Species (Transgene) | Transient luciferase expression (RLU/mg of protein) | Stable transformants resistant to G418 (clones/ µg of transgene) | Stable transformant s expressing luciferase[1] |
|---|---|---|---|
| *O tauri* RCC745 (pH4 :KanMX-pHAPT :Luc)[2] | 400 | 480 (100) | 80 |
| *O tauri* RCC 1110 (pH4 :KanMX-pHAPT :Luc)[2] | 600 | 32 (32) | 32 |
| *O lucimarinus* (pH4 :KanMX-pHAPT :Luc RCC 802)[2] | 650 | 480 (96) | 76 |
| *B. prasinos* RCC 4222 (pH4 :KanMX-pHAPT :Luc)[3] | Nd | 48 (20) | 14 |
| *B. prasinos* RCC 4222 (pH4 :KanMX-BCCT)[4] | Na | 384 (77)[5] | Na |

[1] Luminescence two fold above background
[2] *O. tauri* RCC745 Histone H4 and high affinity phosphate transporter (HAPT) promoters
[3] *B. prasinos* RC4222 Histone H4 and high affinity phosphate transporter (HAPT) promoters
[4] *B. prasinos* BCCT gene encoding putative DMSP transporter interrupted by pH4:KanMX selection marker to test homologous recombination
[5] No BBCT knock-out by homologous recombination was obtained

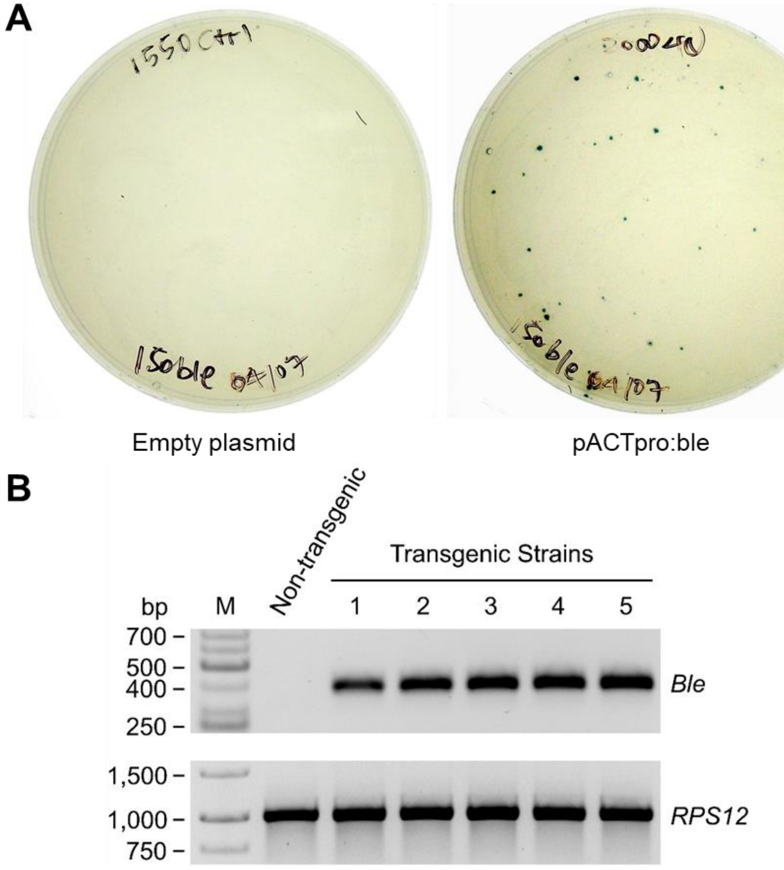


**Suppl. Fig. 2a | Transformation of *Tetraselmis striata*** (KAS-836), by microprojectile bombardment, with a plasmid (pACTpro:ble) containing a selectable marker consisting of the coding sequence of the *Streptoalloteichus hindustanus Ble* gene (conferring resistance to zeocin) under the control of regulatory sequences from the *T. striata* actin gene. **(A)** Transformants were selected on half-strength f/2 agar plates containing 150 µg/ml zeocin. **(B)** Colonies surviving on the selective plates (or non-transgenic control colonies) were examined by PCR amplification of the *Ble* coding sequence using total genomic DNA as the template. The promoter of the *T. striata* 40S ribosomal protein S12 gene was also amplified as a control for equivalent amount of input DNA in the PCR reactions. The panels show reverse images of ethidium bromide stained PCR products amplified from the non-transgenic recipient strain and from five putative transformants. The size of molecular weight markers (M) is indicated in base pairs (bp). GeneRuler DNA ladders (50 bp and 1 kb) from ThermoFisher Scientific. The representative image of three independent experiments is shown.

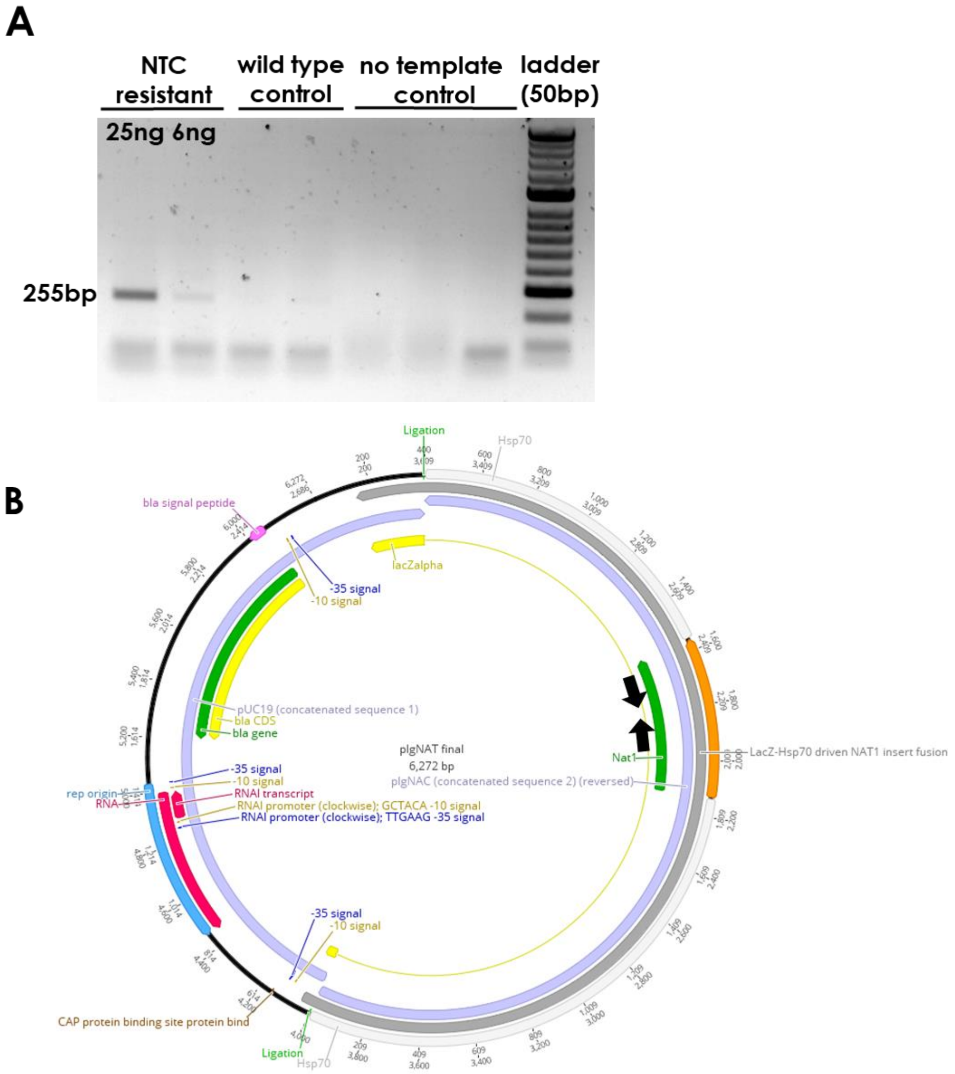


**Suppl. Fig. 2b | Transformation of *Isochrysis galbana* – verification by PCR.**
**(A)** Agarose gel displaying PCR amplification of the NAT gene from *I. galbana* genomic DNA, comparing a nourseothricin-resistant strain to untransformed wild type cells. 50 bp ladder (Bioline) was used. The PCR was performed twice with similar results.
**(B)** Schematic map of the plasmid pIgNAT used for transformation of *I. galbana*. The nouseothricin-resistance gene NAT is flanked by the promoter and terminator of the HSP70 gene from *E. huxleyi*. Black arrows show position of primers used in (A).

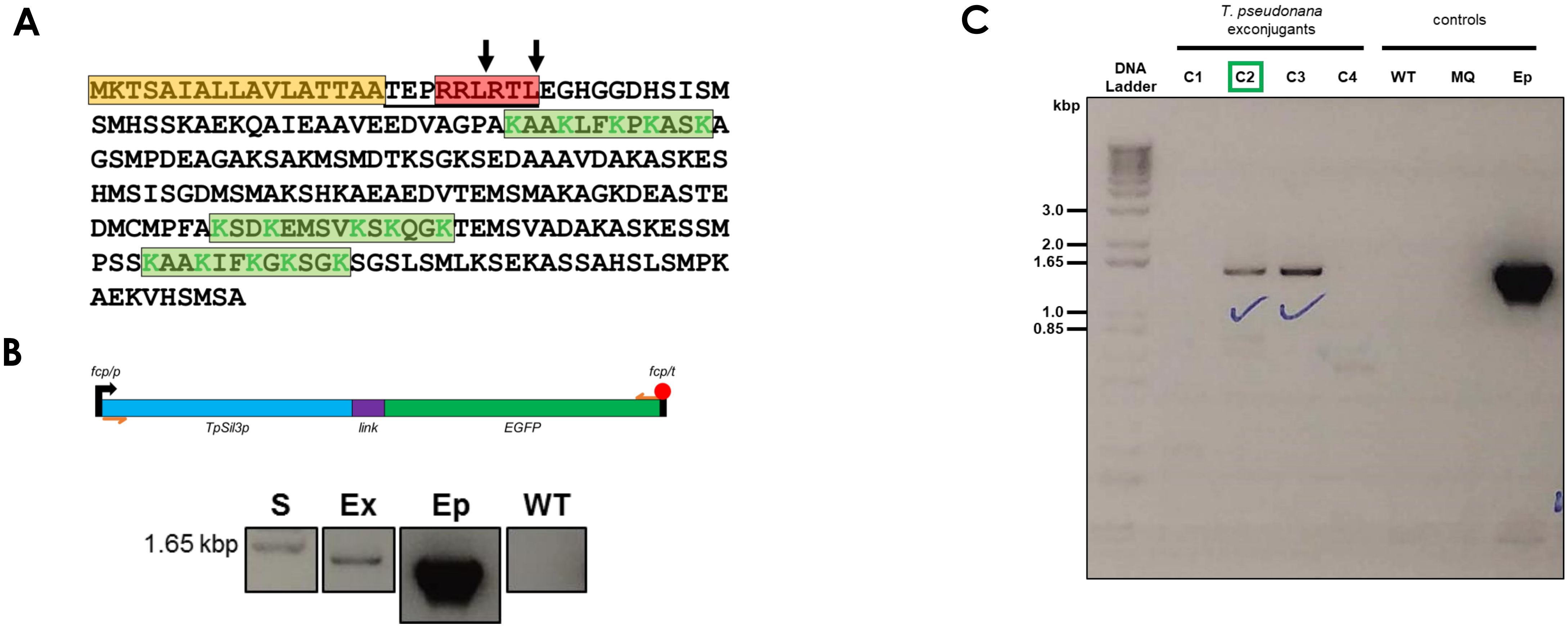


**Suppl. Fig. 3 | TpSil3p features and positive *T. pseudonana* exconjugant confirmation with colony PCR.**
**(A)** The amino acid sequence of TpSil3p, with signal the peptide shown in orange, and the pro-peptide (underlined) with consecutive RxL motifs (in red) - known cleavage sites in biosilicification proteins possibly recognized by signal-1 protease in endoplasmic reticulum or Golgi apparatus (arrows indicate putative cleavage sites); green: pentalysine clusters – 12 to 14 amino acid residues-long motifs with 5 non-consecutive lysines known to promote biosilica association.
**(B)** Top: expression cassette scheme. fcp/p: fcp promoter; TpSil3p: silaffin 3 precursor; *link*: KGSGSTSGSG linker; EGFP: enhanced GFP. The whole open reading frame was optimized for expression in *T. pseudonana* using IDT's Codon Optimization Tool. Depicted in orange are colony PCR primers. The expression cassette was inserted into pTpPuc3 32 bp downstream of the HIS3 gene. Bottom: colony PCR result. S: DNA standard; Ex: *T. pseudonana* exconjugant; Ep: positive control (episome used for conjugation); WT: wild type *T. pseudonana*. 0.5 µl cells were genotyped with Phire Plant Direct PCR Master Mix. Annealing temperature and extension time were set to 57.5 °C and 40 sec, respectively. Expected ~1.45 kb long amplicon was observed in the exconjugant strain corresponding to the full length of designed ORF. 1 Kb Plus DNA Ladder was used. Full gel is shown in (C).
**(C)** C1–C4: four different Nou200-resistant *T. pseudonana* cell lines (i.e. tested exconjugants); C1, C2, C3: three cell lines that exhibited the expected protein localization pattern as demonstrated by fluorescence microscopy; C2, C3: two colony PCR-positive cell lines; C2: cell line selected for the manuscript; WT: wild type *T. pseudonana* cells; MQ: Mili-Q control; Ep: pure episome control. The single experiment resulted in three cell lines exhibiting the protein localization pattern.

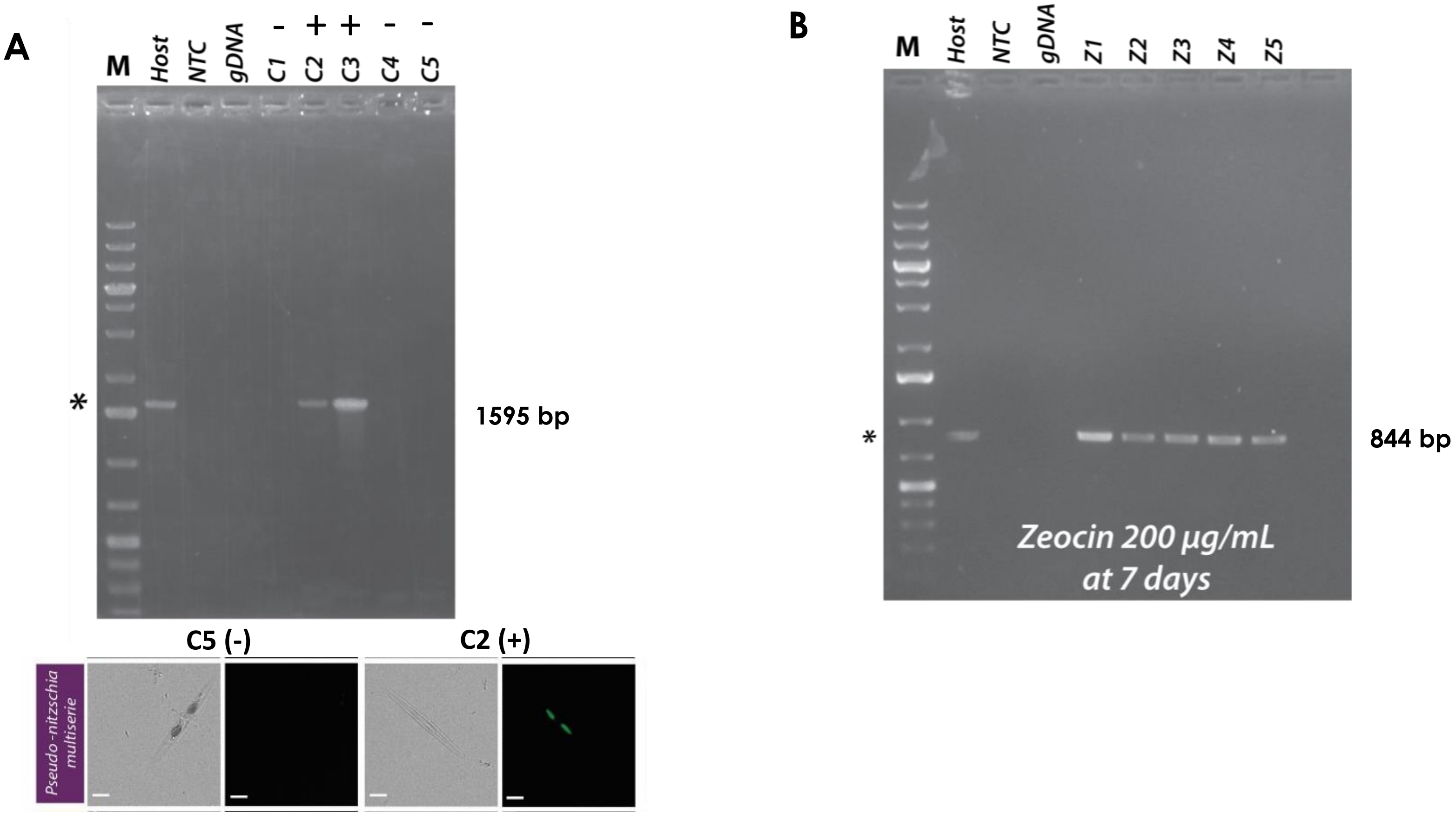


**Suppl. Fig. 4 | Transformation of *Pseudo-nitzschia multiseries*.**

**(A)** Evidence of successful bacterial conjugation-based transfection. The episomal version of the Pm_actP_egfp_actT expression cassette was transfected into the bacterial host EPI300+pTAMOB and used in our modified conjugation protocol. After 48 h recovery in liquid media cultures were plated in LGTA. After 7 d of outgrowth, wells were screened for presence of EGFP+ cells (green cells). PCR screens of + and – picks reveal plasmid maker only associated with EGFP+ phenotype.

**(B)** Evidence for maintenance of heterologous markers under selection. EPI300+pTAMOB+pPtPUC3 bacterial hosts were used in our modified conjugation protocol. Following recovery and 7 d of zeocin selection in LGTA, PCR screens of viable cells (bright chlorophyll fluorescence) picks confirmed presence of the *fcp* promoter and *shble* marker domain, consistent with successful transfection and expression of this heterologous construct. Predicted and observed product size for the screening PCR of pPmAGFPC10 is 1595 bp. PCR screening product for pPtPUC3 is 844 bp. O'GeneRuler 1kb Plus DNA Ladder was used as a marker. 'Host': bacterial donor DNA; 'NTC': no template control; 'gDNA': WT P. multiseries. Representative results from three independent experiments for each construct.

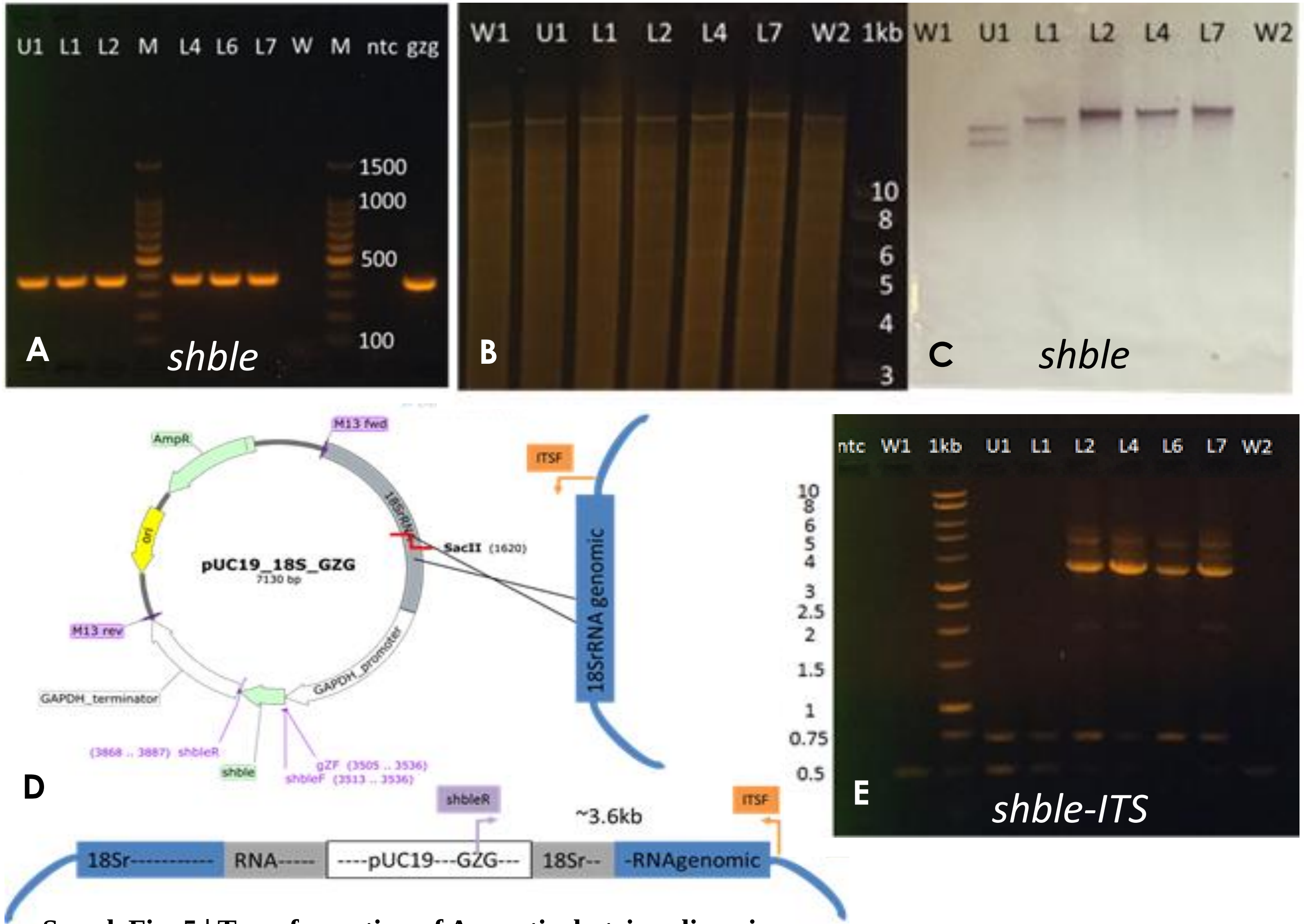


**Transformation conditions and efficiencies for *A. limacinum*.**

| Instrument | Condition | pUC19_18 GZG DNA | Initial Colonies | Stable Transform-ants | Efficiency (transformant/ µg DNA) |
|---|---|---|---|---|---|
| Bio-Rad Gene Pulser | 300 V, 1 pulse | 10 µg | 10 | 1 | 0.1 |
| | 450 V, 2 pulses | 1 µg | 54 | 44 | 44 |
| | | 2.5 µg | 62 | 52 | 20.8 |
| | | 5 µg | 50 | 39 | 7.8 |
| | | 10 µg | 75 | 68 | 6.8 |
| NEPA | 250 V, 4 ms, 2 pulses | 1 µg | 25 | 18 | 18 |
| | 275 V, 8 ms, 2 pulses | 1 µg | 24 | 13 | 13 |
| | 300 V, 12 ms, 2 pulses | 1 µg | 11 | 7 | 7 |

F

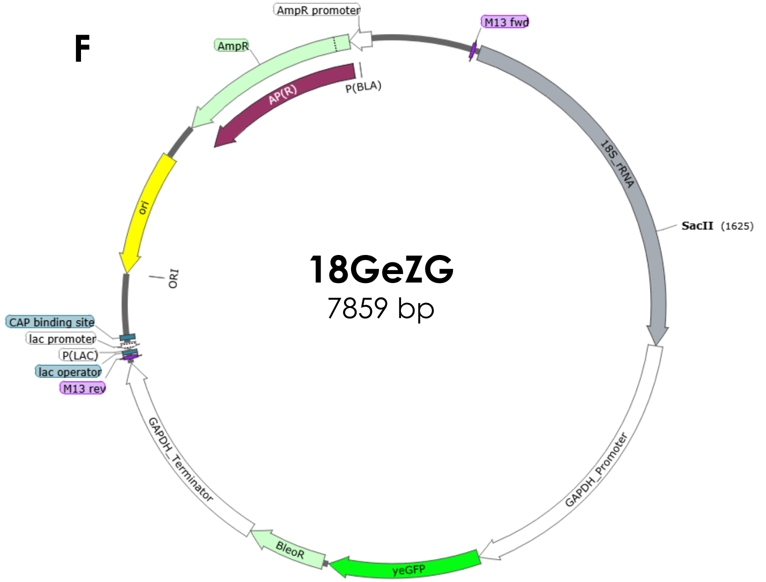


**Suppl. Fig. 5 | Transformation of Aurantiochytrium limacinum.**
(**A**) Genomic DNA PCR confirmation of *shble* (375 bp) in six 18GZG transformants (U1, L1, L2, L4, L6, L7). W is the wild-type, gzg is the plasmid control, ntc is the no template control, and M is a 100 bp marker. (**B**) Gel electrophoresis of *Xho*I-digested genomic DNA from five transformants and two wild types (W1, W2). (**C**) Southern blot using digoxigenin-labeled probe of *shble*. (**D**) Diagram of expected 18GZG plasmid integration into the nuclear genome. The *Sac*II restriction endonuclease site located in the 18S rRNA region was used for linearization. Plasmid primers are annotated in purple, and ITS specific primer in orange (ITSF). (**E**) Genomic DNA PCR of *shble*-ITS region (3.6 kb) in six putative transformants. L2, L4, L6 and L7 produced a band expected from single homologous recombination. (**F**) 18GeZG plasmid map, used in yeGFP::*shble* expression. *Sac*II restriction site was used for linearization. The transformation of 18GZG was done at least 11 independent times and always recovered zeocin-resistant transformants. A set of them was analysed from the same experiment in detail, as shown above.

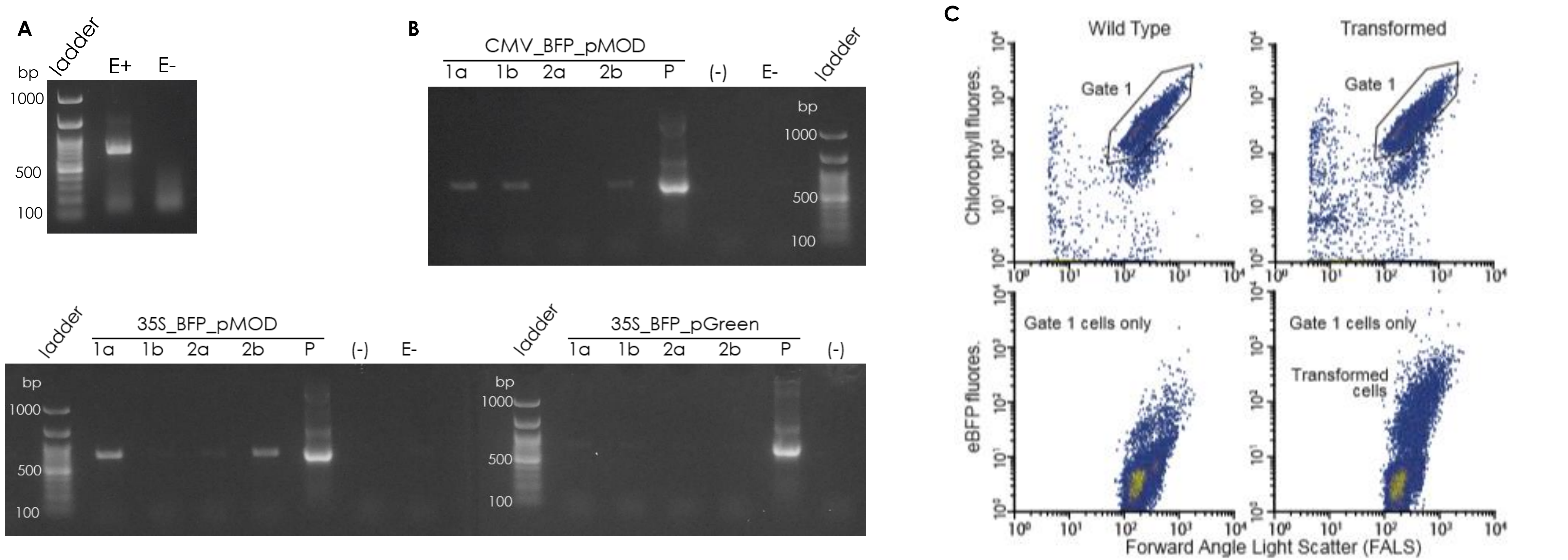


**Suppl. Fig. 6 | Transformation of *Nannochloropsis oceanica* and natural populations.**
(A) RT-PCR of sorted cells incubated with plasmid with (E+) and without (E-) electroporation. Cells were washed 3 times with 800 mM sorbitol, mixed with 10 ug/ml of linearized CMV-BFP plasmid in 800 mM sorbitol and then elecroporated using an exponential decay pulse protocol (electric field of 9 kV/cm and resistance of 500 W). Non-electroporated cells were kept as controls. After 24 h of recovery, flow cytometry was used to sort blue fluorescent cells from electroporated samples (E+) and chlorophyll cells from the non-electroporated ones. 2000 cells were sorted directly into the lysis buffer from the NucleoSpin RNA XS kit and stored at -80℃ until RNA extraction and RT-PCR with the same kit. The band in E+ is the expected product size (750 bp). **(B)** Transformation of natural marine plankton community. Cells were concentrated by tangential flow filtration and washed with 800 mM sorbitol and mixed with 2 mg/mL FITC-dextran (MW = 2000 kDa), 25 ug/mL plasmid and 250 ug/mL carrier DNA (salmon sperm) in 800 mM sorbitol. Plasmids had the mTagBFP2 gene under the control of either the CMV or the 35S promoter. Cells were electroporated in duplicates with an exponential decay pulse protocol (labelled 1a and 1b) (electric field of 5 kV/cm and resistance of 550 W) or a time constant protocol (labelled 2a and 2b) (electric field of 10 kV/cm and time constant of 8 ms). Negative control cells were prepared by incubating them with a mix of all 3 plasmids but without electroporation (E-). Electroporated blue fluorescent cells and non-electroporated chlorophyll cells (from negative control) were sorted (100 cells per reaction) using flow cytometry into 100 µl of TE Buffer prior to PCR. For each set of samples for each promoter, 1a and 1b correspond cells electroporated with the exponential decay pulse protocol, 2a and 2b to cells electroporated with the time constant protocol, P corresponds to the respective plasmid positive control, E- corresponds to cells incubated with the same plasmids but without electroporation, (-) is the negative control for the PCR mix, lad is the 100 bp ladder and corresponding DNA band molecular weights. **(C)** FACS analysis of *N. oceanica* control cells (no plasmid added but pulse applied) and cells subjected to a pulse with a plasmid, after which a subset of cells appeared to have been transformed at 24 h. Two samples electroporated with linearized CMV-BFP plasmid and two controls were performed on each of two days of independent trials using this protocol, with similar results obtained in the other three sample sets. In three independent trials, using linearized CMV-BFP plasmid instead of circular plasmid, percentages of blue-fluorescent cells were much lower (c. 1-2%) though still much higher than controls (not shown). The RT-PCR test and the PCR test were only performed on one trial of each type to provide independent evidence to support BFP expression indicated by flow cytometry.

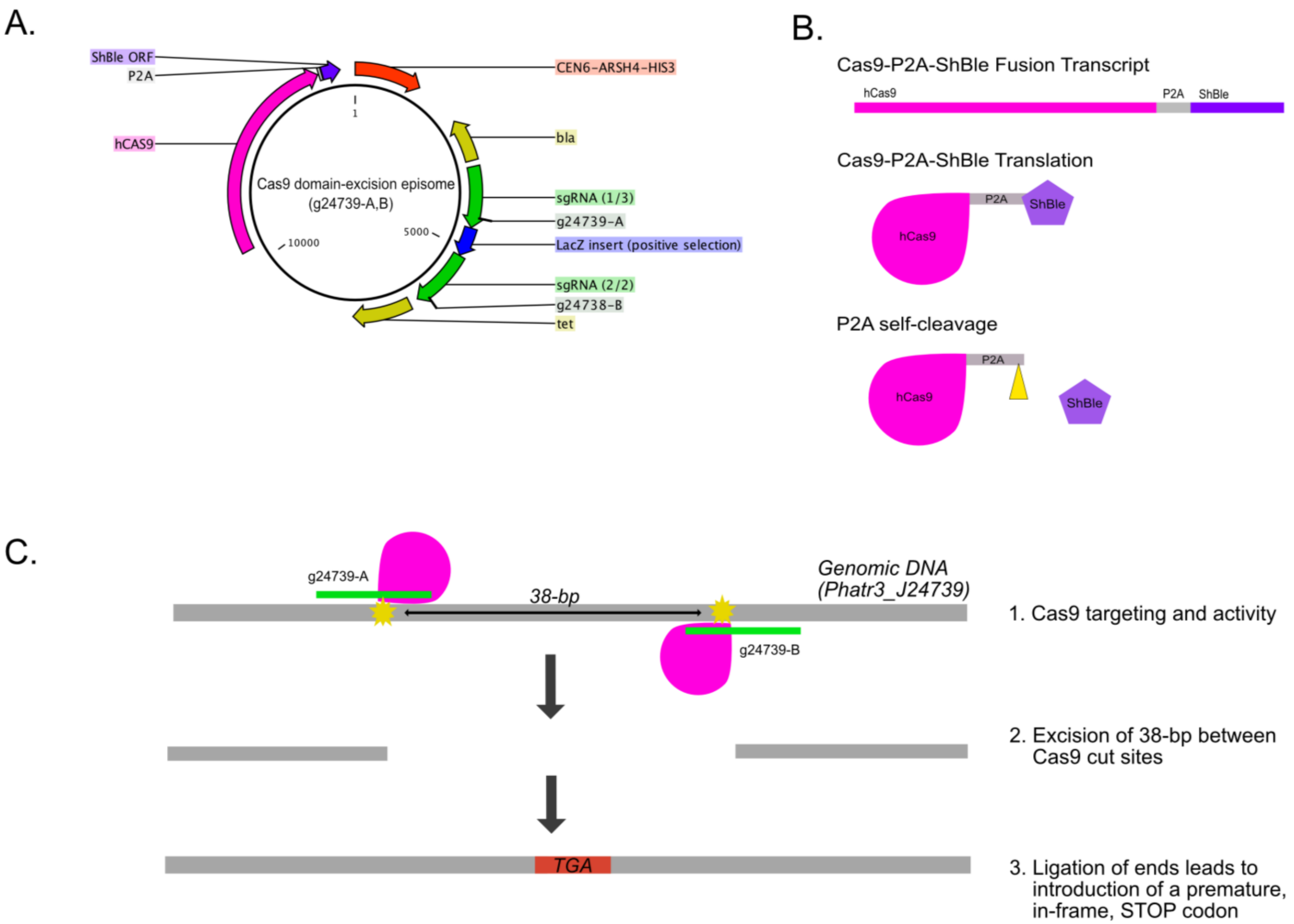


**Suppl. Fig. 7 | CRISPR-Cas9 methodology in *Phaeodactylum tricornutum* CCAP 1055/1.**
Single-gene excision mutagenesis was developed to deliver a selectable hCas9 and two sgRNAs on a conjugation episome. **(A)** Episomal plasmid map containing hCas9 and two sgRNAs. The hCas9 is transcribed with the phleomycin-resistant gene (shble) by the 2A peptide. The two sgRNAs are expressed separately within their own expression cassettes and are separated by a LacZ bacterial expression cassette. Selectable markers for bacterial selection (blasticidin and tetracyclin) and the *P. tricornutum* centromeric region, CEN6-ARSH4-HIS3, are also present. **(B)** The hCas9-2A-ShBle expression cassette permits transcription of the hCas9-2A-ShBle product followed by translation where 2A self-cleaves. **(C)** The two sgRNAs were designed to target the gene cGOGAt precisely 38-bp apart. After cutting of both target loci followed by ligation of the free ends, a premature TGA stop codon was induced downstream of the excised loci.

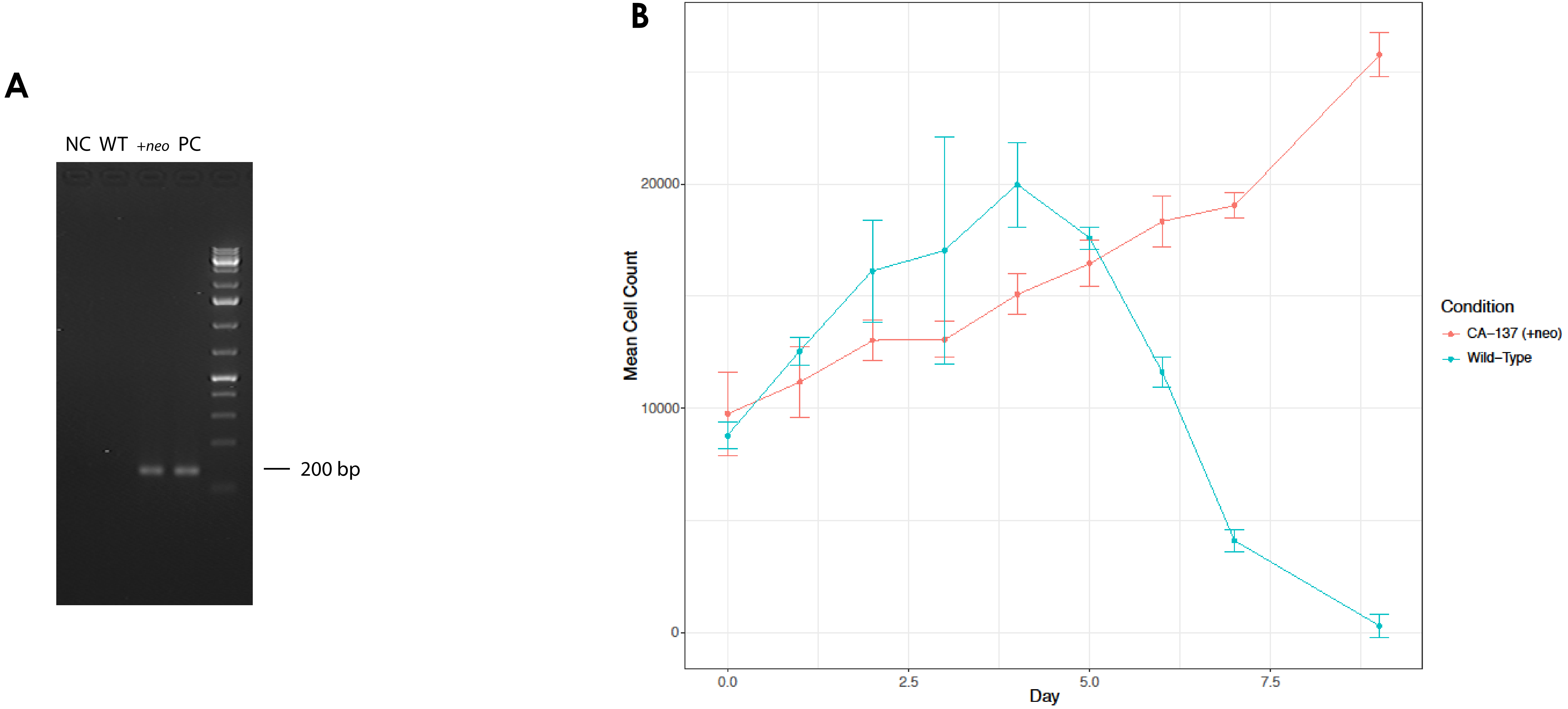


**Suppl. Fig. 8 | Transformation of *Karlodinium veneficum*.**
**(A)** The *neo* gene was detected in the genomic DNA of the transformants through a nested PCR; "NC", negative control; "WT", wild-type; "+ *neo*", transformed cells with DinoIII-*neo*; "PC", plasmid positive control for *neo* gene. Two amplification experiments were repeated independently with similar results. GeneRuler DNA ladder was used.
**(B)** Growth curves of *K. veneficum* in 150 µg/ml kanamycin and 400 µg/ml ampicillin. "CA-137 (+neo)": transformed cells with DinoIII-*neo*; "Wild-Type": wild-type cells. The centre values represent the mean and the error bars are the standard deviation. Each culture was divided into three triplicate bottles.
For each triplicate there were three technical replicates. Four replicated experiments were repeated independently following this design with similar results.

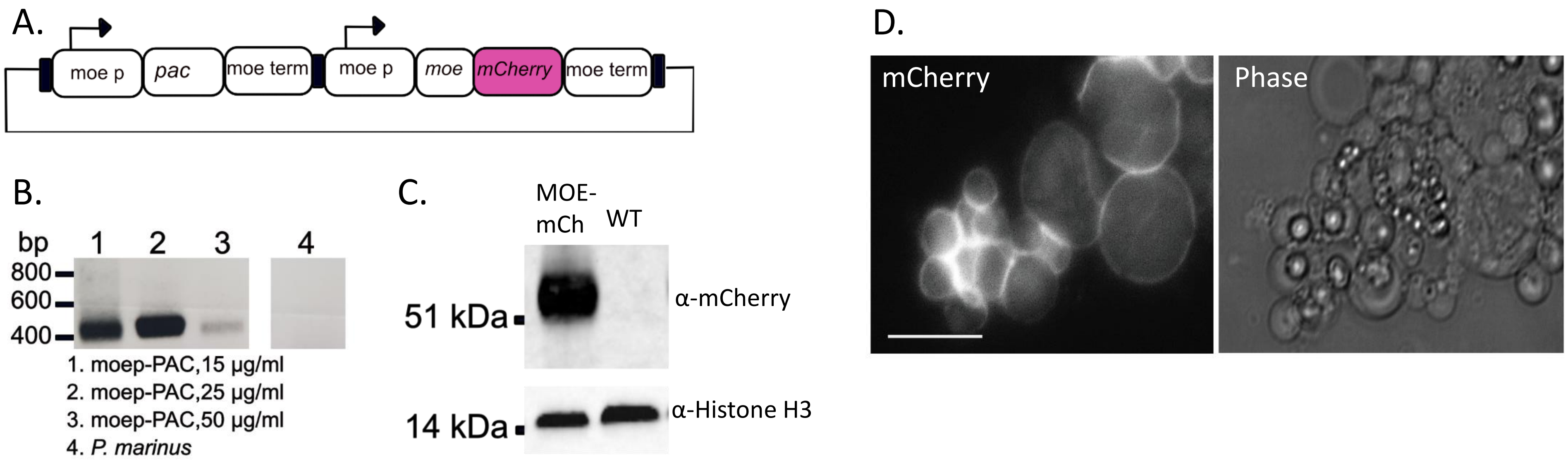


**Suppl. Fig. 9 | Transformation of *Perkinsus marinus.***

**A.** *Perkinsus marinus* cells were transformed using a plasmid that generates resistance against puromycin and expression of MOE-mCherry.

**B.** The puromycin resistance gene (490 bp) can be amplified from gDNA extracted from resistant cell lines that were subjected to different concentrations of puromycin (15, 25 and 50 µg/ml). Non-transformed *P. marinus* was used as a control. HyperLadder 1kb (BIO-33053, Bioline) was used. The experiment was replicated twice with similar results.

**C.** Western blot was used to confirm expression of MOE-mCherry (40.4 kDa) in transformed *P. marinus* cells (polyclonal rabbit α-mCherry, 1:1000, Abcam) and Histone H3 (15 kDa) was used as a loading control (polyclonal rabbit α-Histone H3, 1:1000, Invitrogen). SeeBlue Plus2 Pre-stained protein standard (ThermoFisher, LC5925) was used. The experiment was replicated five times with similar results.

**D.** Epifluorescence image of transformed cells shows positive signal for the MOE-mCherry fusion protein. Scale bar 20µm. The experiment was replicated twice with similar results.

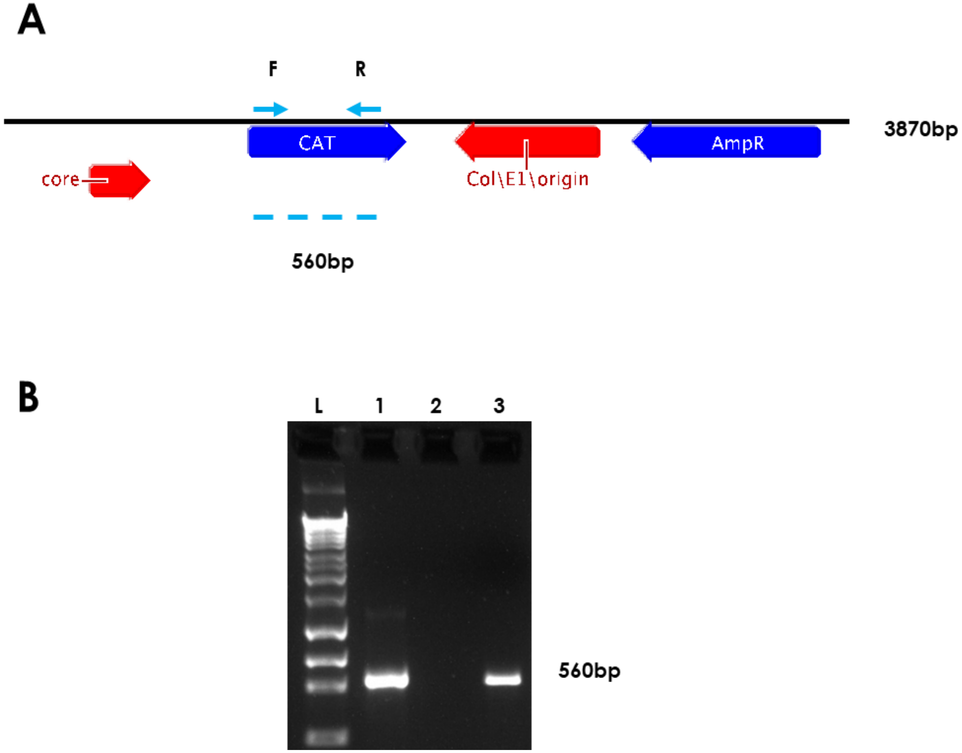


**Suppl. Fig. 10a | Transformation of *A. carterae* with the artificial minicircle pAmpAtpBChl.**
**(A)** Scheme of the pAmpAtpBChl artificial minicircle. Positions of PCR primers used for amplification of the chloramphenicol acetyl transferase gene indicated, as well as sizes of the expected products. Core refers to the minicircle core region (presumed origin of replication).
**(B)** PCR amplification of the chloramphenicol acetyl transferase gene (560 bp) was performed using F and R primers. L – HyperLadder 1kb plus (Bioline), 1 - Positive control PCR with artificial minicircle template, 2 - negative control, 3 - PCR against artificially introduced CAT gene. The experiment was repeated twice with the same results.

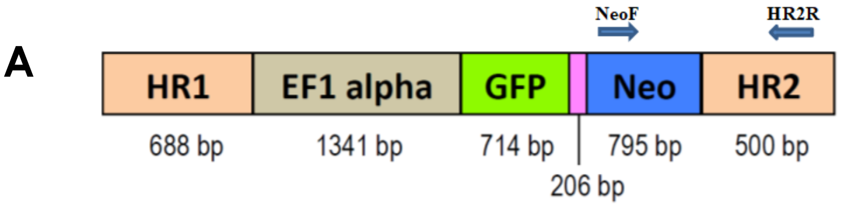


**B**

| Primer | Target sequence | Primers 5' → 3' |
|---|---|---|
| NeoF | Neo gene | CGCTTGGGTGGAGAGGCTATTCGGCTATGAC |
| NeoR | Neo gene | GATGCCTGCTTGCCGAATATCATGGTGGA |
| HR2R | EF-1 alpha 3'UTR | ATGGAAGTGGATCCGGCACTAGCGCT |

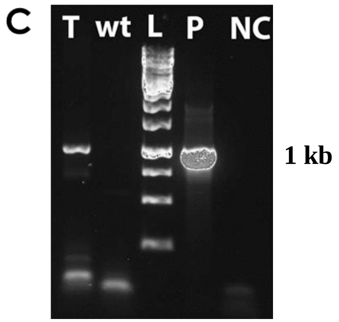


**Suppl. Fig. 10b | Transformation of *Bodo saltans*.**
DNA was extracted from pools of transfected (6 months post-transfection) and wild type cells. The presence of the *Neo* gene in the transfected cells was confirmed by PCR using primers that target the *Neo* gene (NeoF) and the EF-1 alpha 3'UTR (HR2R) included in the cassette plasmid. **(A)** Schematic representation showing the structure of the EF-1 alpha cassette contained in the plasmid. A 206 bp fragment of the intergenic region of α-β-tubulin of *B. saltans* is shown in pink. Arrows indicate primers used to verify the presence of the cassette in transfected *B. saltans* cells. HR1 and HR2 (Homologous regions 1 and 2) represent EF-1 alpha 5' and 3'UTRs, respectively. **(B)** NeoF and HR2R primers sequences. The primers set of NeoF and NeoR was used to confirm the expression of the neomycin resistance gene using RT-PCR (Figure 3k). **(C)** Agarose gel electrophoresis of amplified PCR products using NeoF and HR2R primers. The expected band size is 1 kb. Lanes from left to right: T= transfected cells (6 months post-transfection); wt = wild type cells; L = 1 kb DNA ladder; P = EF-1 alpha plasmid (positive control); NC = negative control. Sequences obtained from PCR products confirmed the presence of the Neo gene and the 3'UTR region of the cassette. Results shown are representative of 10 independent experiments.

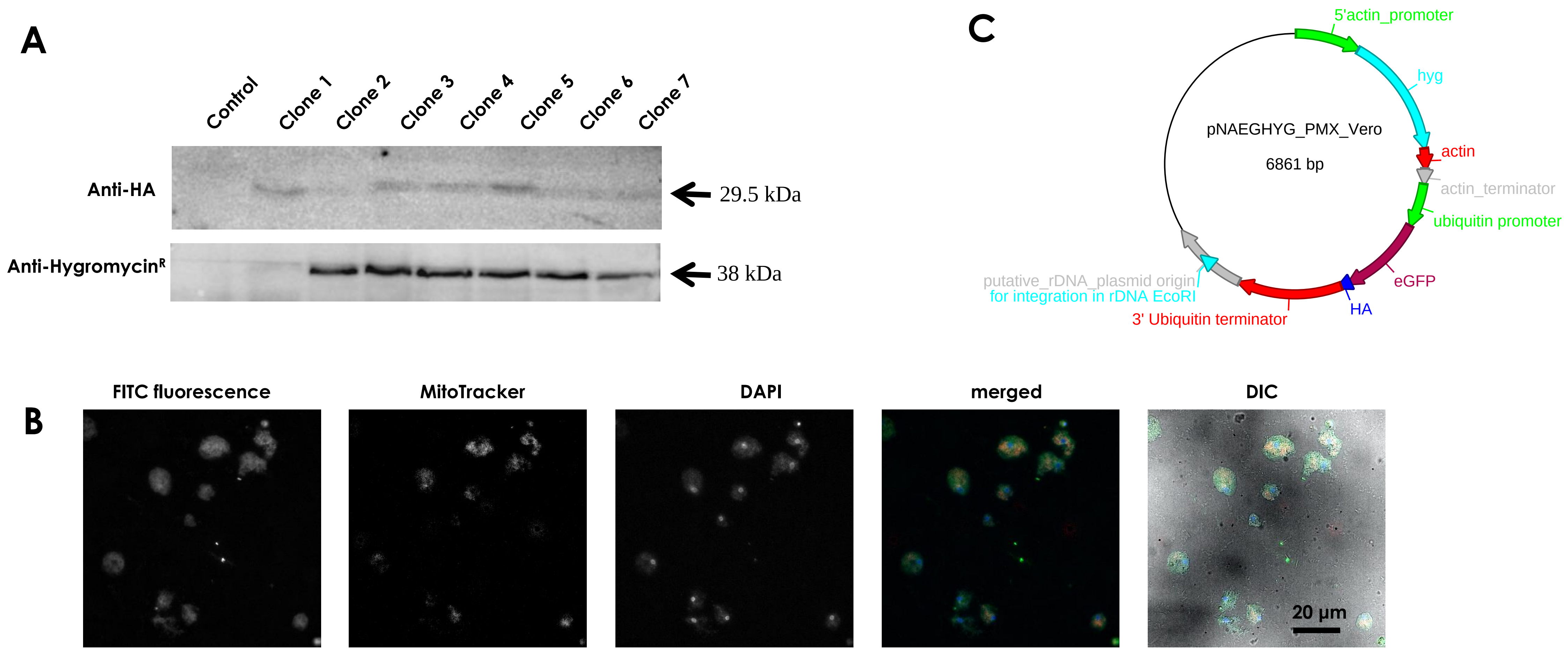


**Suppl. Fig. 11 | Transformation of *Naegleria gruberi*.**

**(A)** Western blot of transfected clonal cells and the control (non-transfected cells) demonstrating the expression of the HA-tag (anti-HA antibody, Sigma) and hygromycin B resistance gene – *hph* (anti-hygromycin$^R$ antibody, Fitzgerald Industries) in these clones but not in the control. Molecular Marker: PageRuler, Unstained Protein Ladder (Thermo Scientific, Catalog number:26614). Representative results of three independent replicates are shown.

**(B)** Cellular localization of eGFP by immunofluorescence microscopy in cells 20 days post-transfection. The different panels show the expression of eGFP (FITC fluorescence; green), the localisation of mitochondria (MitoTracker; red) and nuclear/mitochondrial localisation (DAPI; blue). The merged panel demonstrates the co-localisation of all different panels and the DIC demonstrates the localisation using Differential interference contrast. Scale bar 20 µm. Representative results of three independent replicates are shown.

**(C)** Map of the hygromycin plasmid that was used for transfection.

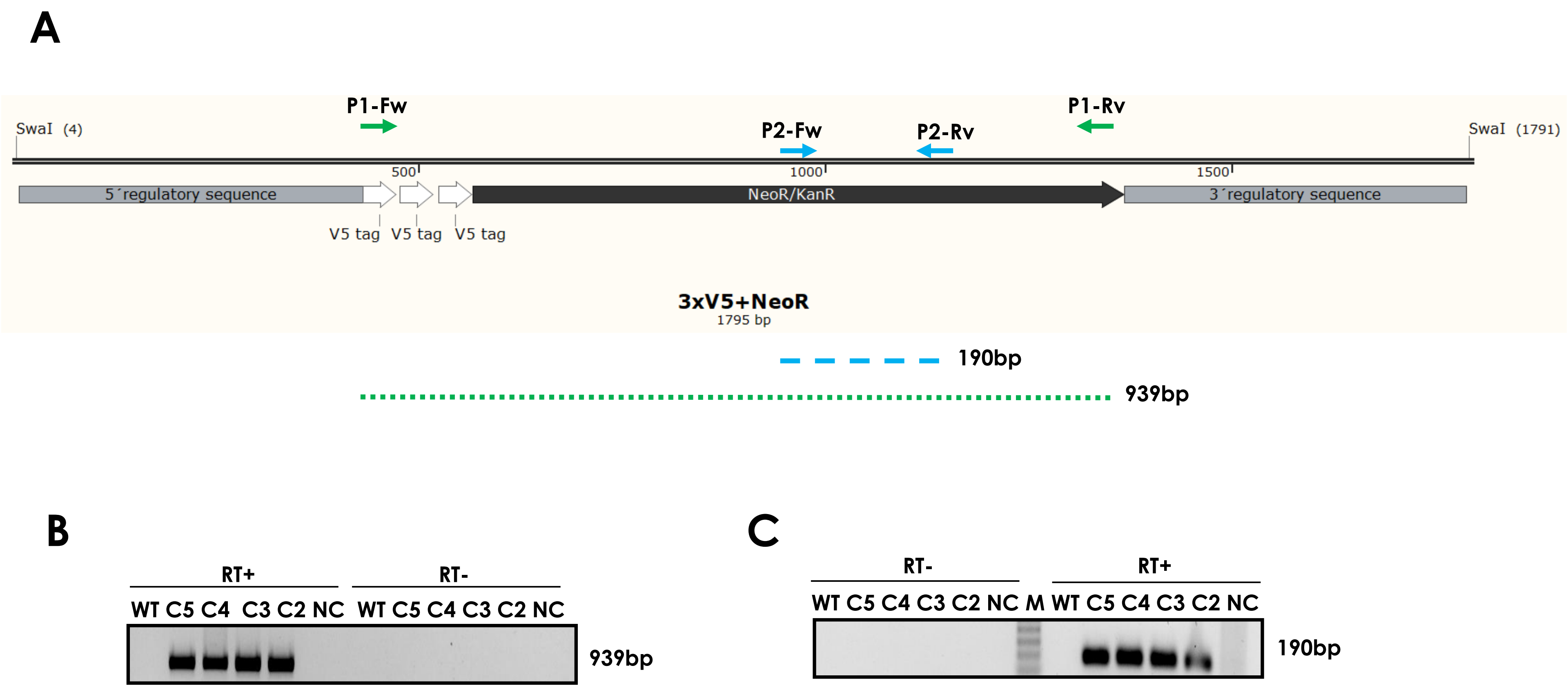


**Suppl. Fig. 12 | Validation of proper transcription of 3xV5+NeoR heterologous gene in *Diplonema papillatum* by RT-PCR**.
**(A)** Scheme of the 3xV5+Neo$^R$ construct. Positions of RT-PCR primers used for amplification of 3xV5+NeoR cDNA and its part (see below) are indicated, as well as sizes of the RT-PCR products.
**(B)** RT-PCR amplification of the 3xV5+Neo$^R$ transcript (939 bp-long region) was performed using P1-FW and P1-RV primers. **(C)** RT-PCR amplification of the 3xV5+Neo$^R$ transcript (190 bp-long region) using P2-FW and P2-RV primers. RNA from wild type cells (WT) and transformants A1, A2, A3 and A4 was used as a template for RT-PCR. A sample without cDNA (NC) served as a negative control. Reactions with and without reverse transcriptase are indicated by (RT+) and (RT-). M = 1Kb Plus DNA ladder (Invitrogen, 10381-010) was used. Representative images of two independent experiments are shown.

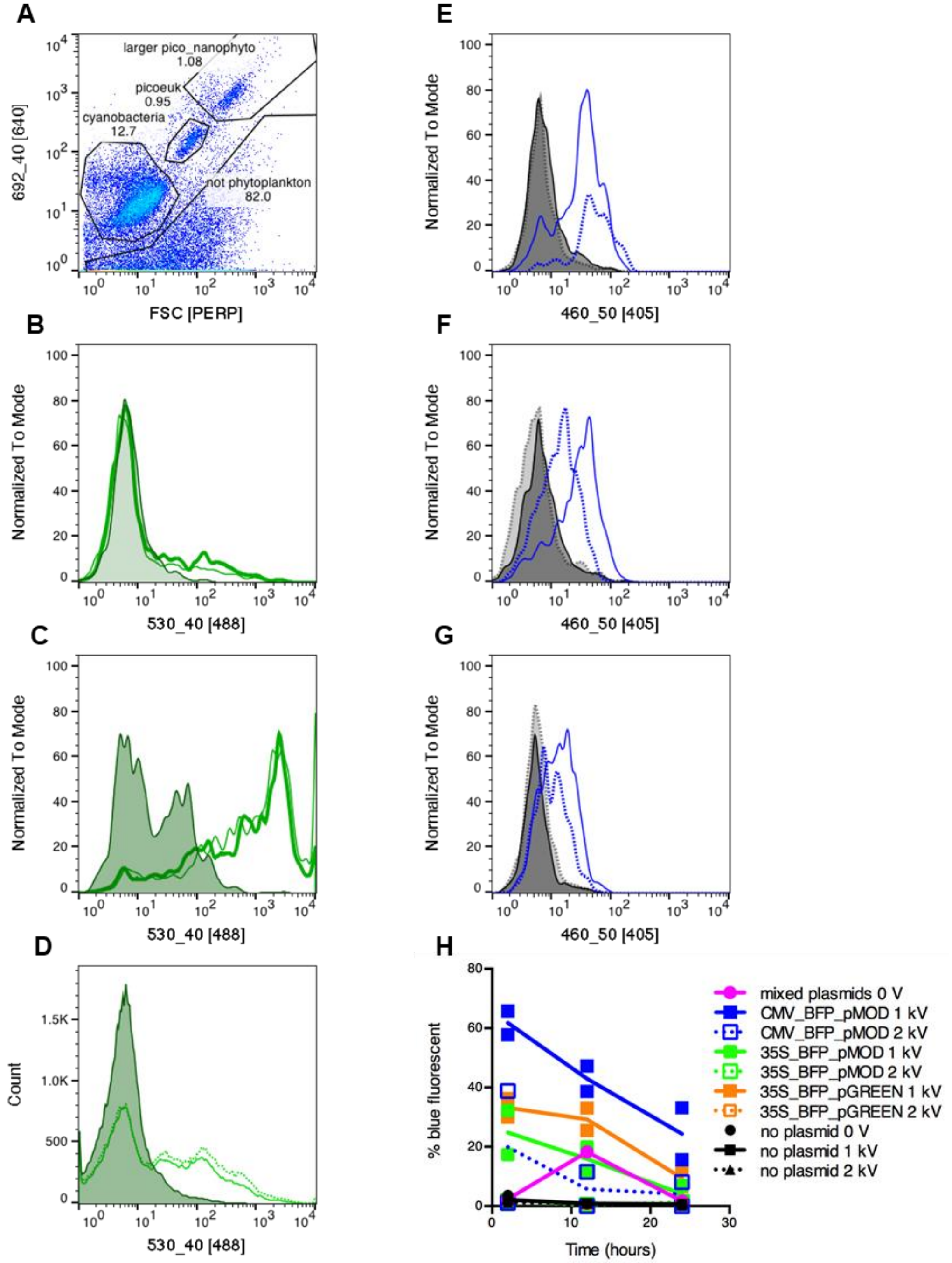


**Suppl. Fig. 13 | An exceptional case of transient expression of BFP in natural sea water communities.** Flow cytogram of red fluorescence from chlorophyll excited by the 488 nm laser (692_40 [488]) versus forward scatter of natural community obtained by concentration of 50 liters of sea water by tangential flow filtration after which cells were washed with sorbitol 800 mM (A). Histograms of green fluorescence (from fluorescein 2,000 kD dextran and autofluorescence) in picophytoeukaryote (picoeuk) (B), larger pico-nanophytoeukarotes (C), and nonphytoplankton (D) 2 h after electroporation with a pulse of 1 kV for 20 ms (solid line) or 2 kV for 8 ms (dashed line). Shaded histogram shows sub-sample incubated with dextran but without electroporation. Picophytoeukaryotes were electroporated also with the CMV_BFP_pMOD plasmid (blue) or without plasmid (grey) at 1 kV (solid lines) or 2 kV (dashed lines) at 2 hours (E), 12 hours (F), and 24 hours (G). Time course of percentage blue fluorescent cells in picoeukaryotic phytoplankton over time in all treatments (2 replicas per treatment) (H). Out of a total of six trials on independent days with different natural plankton communities, an increase in blue fluorescent cells in samples electroporated with plasmid compared to controls was seen in some picoplancton, in two other trials, but the percentages were lower than in the case shown here. In the other three trials, blue fluorescent cells were too rare for differences in samples electroporated with plasmid compared to controls to be detected by the flow cytometry protocol used. In no other case was an increase in blue fluorescence noted in samples incubated with plasmids but without electroporation.

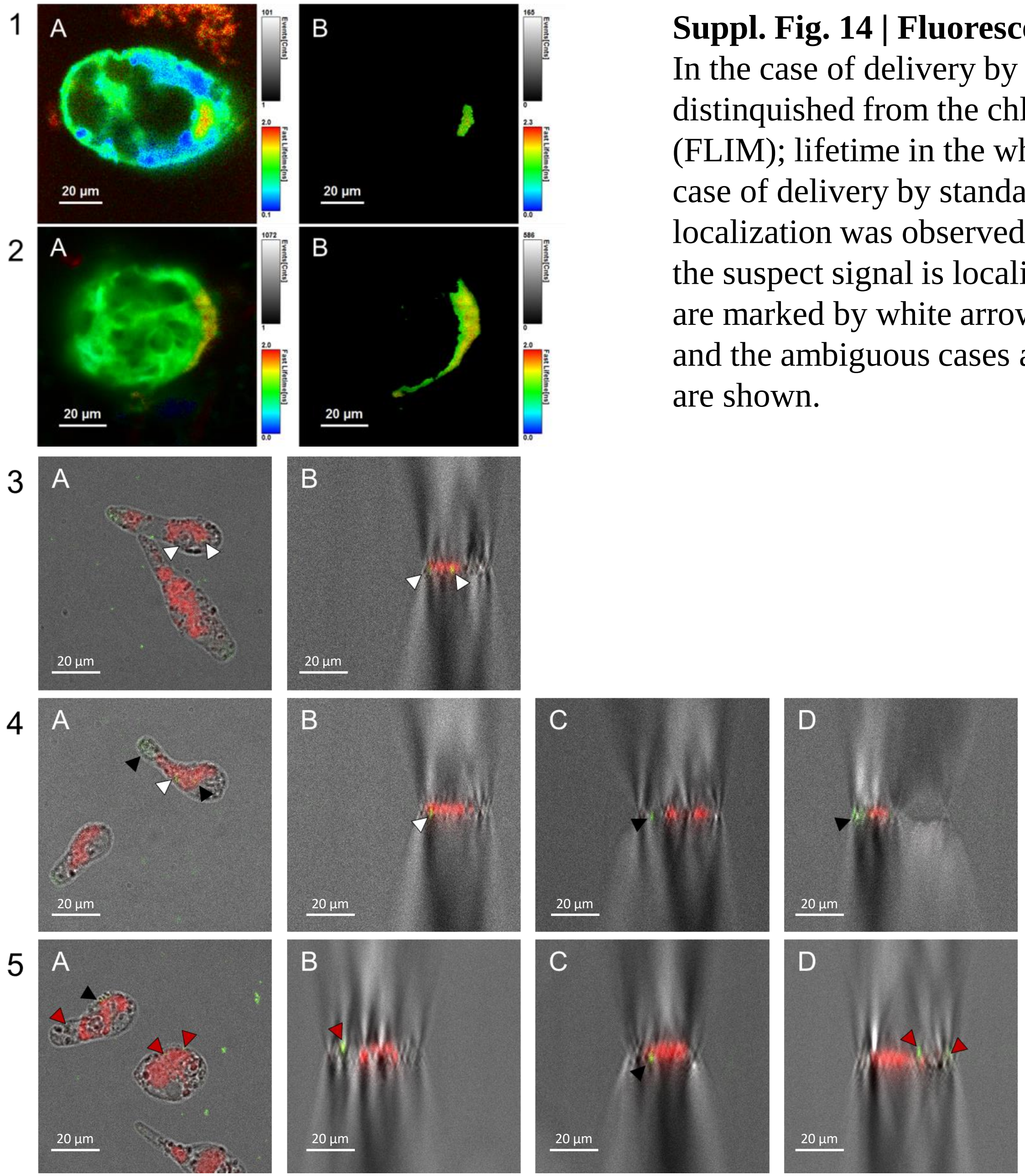


**Suppl. Fig. 14 | Fluorescent signal of the plasmid DNA in *E. gymnastica*** (1, 3-5) **and *P. parkeae*** (2). In the case of delivery by nucleofection (1-2), DNA was labelled by fluorescein and its signal was distinquished from the chlorophyll autofluorescence using fluorescence-lifetime imaging microscopy (FLIM); lifetime in the whole cells (A) and in the area with longest component (B) is shown. In the case of delivery by standard electroporation (3-5), DNA was labelled by tetramethylrhodamine and its localization was observed by confocal microscopy (A); XZY scan was performed to determine whether the suspect signal is localized inside the cell or on its surface (B-D): the seemingly successful deliveries are marked by white arrows, the probable extracellular DNA localizations are marked by red arrows, and the ambiguous cases are marked by black arrows. Representative images of a single experiments are shown.

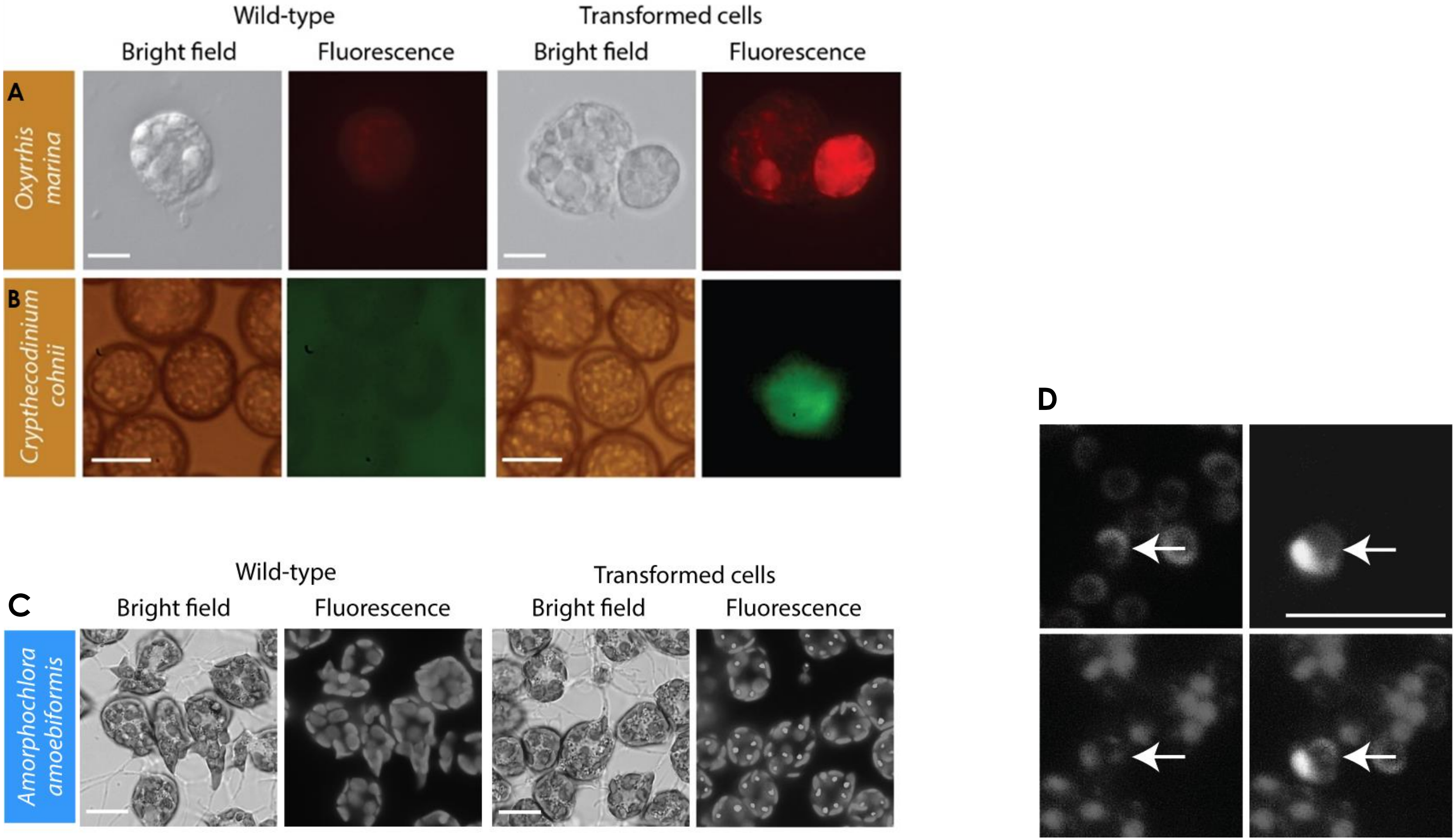


**Suppl. Fig. 15 | Transformation of *Oxyrrhis marina* and *Crypthecodinium cohnii* by labeled DNA.** Fluorescent microscopy images showing the transformants and wild type cells of *O. marina* **(A)** and *C. cohnii* **(B)**. Representative images of two (A) and three (B) independent experiments are shown. Coloured boxes behind species names reflect supergroup assignments in Fig.1 (Alveolates). Scale bars 20 μm and 11 μm for *O. marina* and *C. cohnii*, respectively.

**And trace of the different regions in the cell -** for *A. amoebiformis* shown in Fig. 2; scale bar: 10 μm **(C)** and *M. commoda* shown in Fig. 3c; the scale bar: 5 μm **(D)**.

#### SUPPLEMENTARY RESULTS

##### Archaeplastids

The prasinophyte *Micromonas pusilla* (CCMP1545) was selected for reverse genetics, as members of the genus *Micromonas* are abundant in marine ecosystems and have larger genomes than *Bathycoccus* and *Ostreococcus*, in addition to having a flagellum[1,2]. Although *M. pusilla* is typically less stable in culture in an axenic state than *M. commoda* (RCC299/NOUM17; deposited at the NCMA after being rendered axenic and genome sequenced as CCMP2709), it was initially selected for manipulation due to its interesting genomic features[3]. The procedures that worked for *Ostreococcus*[4] and *Bathycoccus* (this work) failed with *M. pusilla*. Furthermore, we made multiple attempts to use electroporation and bacterial conjugation (ATCC agrobacterium strain AGL-1) where we also monitored cells to ensure they were in exponential growth at the start of each experiment. We constructed plasmids with a codon-optimized chloramphenicol acetyltransferase (CAT) or eGFP-SV40NLS genes driven by *M. pusilla* RPS9 elements in between Ti-plasmid elements required for *Agrobacterium*-mediated transfer in the pOSCAR plasmid system[5]. We determined *M. pusilla* sensitivity to chloramphenicol by testing a range of concentrations (**Suppl. Table 3; Online Methods**). While it is difficult to say exactly why these efforts failed, we did not observe GFP fluorescence nor did chloramphenicol resistant cells emerge from culture. Because of the persistence of *Agrobacterium* in the culture after transfer, analysis by flow cytometry and recovering axenic cultures was challenging. In all assays where DNA was added directly to cultures, it was in supercoiled form as isolated from *E. coli*. Lonza nucleofection was not attempted with *M. pusilla* due to toxicity to all three Lonza buffers (SF, SG, SE), at least under the conditions tested herein, while buffers SF and SG allowed growth of *M. commoda* (SE was toxic). It should be noted that *M. commoda* and *M. pusilla* have major differences, for example, the latter retains the peptidoglycan pathway (peptidoglycan layer surrounding the plastid), while the former has lost most of this pathway like *Arabidopsis thaliana*[2]. This influences antibiotic usage as many of them work on this pathway. Furthermore, *M. pusilla* has some known RNAi pathway genes while *M. commoda* does not.

*Pyramimonas parkeae* (SCCAP K-0007), class I prasinophyte alga[9], was targeted by mRNA encoding firefly luciferase and by two circular plasmids. The cells were electroporated by Gene Pulser xCell (BioRad) with 5 µg of *in vitro*-synthesized firefly luciferase mRNA using exponential pulse at 8 different combinations of voltage and capacitance settings (**Suppl. Table 4**). Light emission was measured after 4.5, 5.5 and 22 h, yet no specific luciferase activity could be detected. Before we conducted the genetic transformations, the sensitivity to 5 antibiotics was tested. Puromycin and geneticin were identified to be most effective at a concentration of 30 µg/ml each (**Suppl. Table 3**). Two plasmids containing 5' and 3' UTRs from the *P. parkeae* histone H2 gene were synthesized, with one containing the geneticin resistance gene, while the other plasmid carried the firefly luciferase and puromycin resistance genes. The former gene was flanked by the 5´and 3 UTRs of the native histone H4 gene. Both circular and linearized plasmids were delivered by electroporation using Gene Pulser xCell or Amaxa Nucleofector II (Lonza) (**Suppl. Table 4**). However, no light emission was measured in the cultures post DNA delivery, and no cells were selected even after 4 weeks of growth on selective liquid F2 medium. Subsequently, we have tried to deliver fluorescein- or tetramethylrhodamine-labelled DNA by electroporation or nucleofection as described above, with various settings (**Suppl. Table 4**), as well as by biolistics (PDS-1000/He [BioRad]; 0.6 µm particles; 1250 psi) to test the DNA delivery efficiency. Nucleofection resulted in fluorescent signal from inside the cells suggesting the DNA delivery was succesful (**Suppl. Fig. 14; Online Methods**), although the expression was not achieved.

**Haptophytes (incertae sedis)**
*Emiliania huxleyi* (see Transformation of natural protist communities)

**Rhizarians**
*Bigelowiella natans* (CCMP 2755) is a widespread chlorarachniophyte[10], which we have attempted to transform independently with 2 circular plasmids. One plasmid contained the firefly luciferase gene, while the other carried both the luciferase and geneticin resistance cassettes. Approximately 300 nt 3' and 5' UTRs of the *B. natans*–derived histone H4 were used as endogenous promoters and terminators for driving the geneticin resistance gene. As for the firefly luciferase gene, the native regulatory sequences were derived from the up- and downstream flanking regions of an endogenous pyruvate kinase gene. Although 3 methods of delivery (electroporation of circular plasmid, lipofection, and biolistics - **Suppl. Table 4**) were tested, no transformants were obtained and no light emission was measured in the culture post-DNA delivery. Finally, we have also attempted delivery of tetramethylrhodamine-labelled DNA using electroporation and biolistics (**Suppl. Table 4; Online Methods**), yet also with an unsuccessful outcome.

**Stramenopiles**
Although genetic transformation of diatoms using biolistics has been ongoing for decades[11,12], it has only recently accelerated, partially fuelled by the availability of more genome sequences and numerous transcriptomes[13], as well as by the development of novel genome editing methods. Indeed, new model diatoms are being developed to address questions that cannot be investigated with their more established representatives *Thalassiosira pseudonana* and *Phaeodactylum tricornutum*[14]. Different diatom species have been transformed using a variety of DNA delivery methods including electroporation, biolistics, and conjugation. Targeted gene expression and genome editing have also been achieved using overexpression[15], RNAi[16], meganucleases, TALENs[17] and CRISPR/Cas9[18]. Building on theses successes and extensive resources, we here extend this work to other ecologically and economically important diatom species. Our primary goal was to increase the repertoire of tools available for their genetic manipulation, as well as to describe what was tried with the other species.

*Seminavis robusta* (D6 and VM3-4), a benthic pennate diatom with a well-defined life cycle, is accessible to genetic analysis due to the presence of 2 defined mating types[19]. It was maintained in autoclaved filtered artificial sea water at 12 h light-dark cycle at 18 °C in f/2 medium supplemented with Guillard's f/2 nutrients, and with 250 µg/ml penicillin, 250 µg/ml ampicillin, 50 µg/ml streptomycin sulfate, and 25 µg/ml gentamicin for regular maintenance. Antibiotic sensitivities were established by following for 3 days the division of small number of cells deposited in arrays of 40 µl drops on Petri dishes, which was both faster and more reliable than screening in bulk cultures. In such liquid cultures, we found that 250 µg/ml of glyphosate or 10 µg/ml of puromycin were sufficient to arrest growth, thus qualifying as selectable marker for transformation (**Suppl. Table 3**).
To create 3 transformation constructs (pBS_Act_Agro4GlyphR, pBS_Act_PurR and pBS_ActEF-AgroCP4GlyphR), the puromycin N-acetyltransferase and agrobacterium CP4

glyphosate resistance genes were codon optimized (**Online Methods - Online Methods Table 1**) and cloned into different plasmids. To drive their expression, each antibiotic resistant marker gene was 5' flanked with 1.2 kb promoter from a highly expressed actin 1, while the 3' flanking region was represented by either 1.64 kb 3' region derived again from actin 1, or 1.0 kb 3' region derived from the elongation factor 1 of *S. robusta*. However, transformation failed with either biolistic bombardment PDS1000 (BioRad) or electroporation (BTX ECM 2001 and NEPA21) (**Suppl. Table 4**). Following biolistic bombardment, we found that the recovered cultures, which were of only one mating type, contained many apparent exconjugants. This implied that the cultures had undergone self-mating[20]. This observation may be useful for future genetic studies, as it might allow inducing the formation of homozygotes from a clonal population.

***Heterosigma akashiwo*** (CCMP2393) is a harmful alga from the raphidophyte group, causing fish mortality. Although no specific toxin has been isolated, reports indicate that some raphidophytes produce a suite of compounds that contribute to their toxicity, including reactive oxygen species[21], polyunsaturated fatty acids[22] and brevetoxin-like compounds[23]. Research on *H. akashiwo* suggests that toxicity is strain-specific and linked to genetic markers[24]. *H. akashiwo* is also a candidate for biodiesel production[25] and its non-toxic strain is being developed for biofuel production coupled to bioremediation of industrial emissions[26]. While the nuclear genome has not been fully sequenced yet, transcriptomes and the plastid genome are available[27].

Antibiotic sensitivities of *H. akashiwo* are listed in **Suppl. Table 3**. Plasmids were constructed from *Chlamydomonas reinhardtii* Chlamy_3 and 4, and the diatom pPhat_T1 plasmids[28]. pChlamy_3 was modified with cLuc or cGFP reporter genes[29], while pChlamy_4 and pPhat_T1 vectors were modified with pAES-Luc[30]. All reporter genes were codon optimized for *C. reinhardtii*. Three transformation methods were compared: a glass bead beating protocol, electroporation and *Agrobacterium*-mediated transformation (**Suppl. Table 4; Online Methods**). Of these, an optimized bead-beating protocol successfully and repeatedly produced cultures transformed with a plasmid conferring hygromycin resistance. For this protocol, cells were harvested, washed and resuspended in MAX Efficiency Transformation Reagent for Algae or with 384 mM sorbitol. Cells were mixed with 1-10 µg of linearized plasmid and glass beads (425-600 µm), vortexed at high speed for 10 s and transferred to a fresh medium with cefotaxime to control bacterial growth. Transformation and expression were verified up to 9 months after the experiment by PCR and RT-PCR amplifications of antibiotic-resistance genes, respectively, but we were not successful in visualizing the fluorescent reporter by microscopy. An optimized electroporation protocol achieved lower efficiency than the bead-beating protocol. Cells resuspended as described above with at least 1 µg of linearized plasmid were electroporated using Gene Pulser xCell with a 10-15 ms pulse at 50/75 V, 25 µF and ∞Ω. Finally, when transformation with *Agrobacterium* carrying cGFP was attempted, *H. akashiwo* survived the co-cultivation step but there was little or no viability apparent after several days of hygromycin selection. A major constraint of all methods was that *H. akashiwo* could not be cultured on solid media and hence single-cell transformants were not selected for further analysis. Therefore, all work was done with cells suspended in cultivation medium. A comparison of methods revealed that PEG-mediated bead-beating protocol produced the best results.

Fluorescein isothiocyanate conjugated to dextran, a fluorescent DNA analog, has been successfully introduced into *Caecitellus* sp. using a high voltage exponential decay pulse (1000 V, 10 µF, ∞ Ω) from a GenePulser xCell (cuvette width 0.2 cm). Cell viability was confirmed by observing motility and fluorescence 24 h after electroporation. Lower voltage pulses did not result in DNA delivery.

**Alveolates**

For introduction see Results. *Euplotes crassus* (txid5936) and *Euplotes focardii* (txid36767) are marine ciliates carrying a micronuclear genome that represents the germ line, and a macronuclear genome containing single-gene nanochromosomes amplified to thousands of copies for their somatic life[31]. Both ciliates are suitable for studying DNA rearrangements and transposition mechanisms, which have been observed in macronuclei after completion of the sexual cycle[32,33]. *E. crassus* and *E. focardii* are resistant to antibiotics usually used for genetic transformations such as ampicillin (100 µg/ml). However, geneticin (200 µg/ml), paromomycin (200 µg/ml) and puromycin (40 µg/ml) are effective (**Suppl. Table 3**) when cells were grown in a modified medium (10% artificial seawater and 90% 0.3 M glucose). Here, we describe conditions for growing both *Euplotes* species in the presence of bacteria, which are being used as vectors for transmitting RNAi to knockdown genes in ciliates. Plasmids and artificial nanochromosomes ending with telomeres that contain either eGFP and/or geneticin as selectable marker, both optimized according to the *Euplotes* codon usage, have been generated. They resemble the *Euplotes* nanochromosome structure with the flanking non-coding regions belonging to the constitutive highly expressed *Euplotes* genes. Telomeres have been added via PCR using primers containing a specific sequence and ending with $C_4A_4$ repeats. The 940 nt eGFP artificial nanochromosomes are composed of *E. crassus* 5' UTR from a highly expressed gene as promoter, eGFP coding sequence, *E. crassus* 3' UTR from a highly expressed gene, and telomeres on both ends. The 3.2 kb artificial nanochromosomes containing eGFP and resistance to geneticin are composed of β-tubulin 5' UTR, β-tubulin coding sequence, a spacer, eGFP coding sequence, β-tubulin 3' UTR, α-tubulin 3' UTR, geneticin resistance gene, α-tubulin 5' UTR, and telomeres on both ends. The DNA delivery protocols (**Online Methods – Online Methods Table 1**) were successful, since we could confirm the presence of Cy3-labelled plasmids inside the cells following electroporation, lipofectamine transfection, and microinjection (**Suppl. Table 4**). However, there is no evidence for either incorporation of the plasmids into the nuclear genome or for the expression of the introduced marker genes (eGFP and/or resistance to geneticin).

*Chromera velia* (CCMP2878) is a marine photosynthetic alveolate associated with corals[34]. We were not able to identify antibiotics suitable for its selection (**Suppl. Table 3**), although a range of antibiotics at different concentrations was tested. Nevertheless, we developed 2 plasmids for the transformation of *C. velia*. The first one contained the firefly luciferase gene, while the other plasmid carried both luciferase and the puromycin resistance genes. The *C. velia*-derived ~300 nt 5' UTR of the enolase gene and equally long 3' UTR of its H4 histone gene were used as regulatory sequences of the puromycin resistance gene. In the case of the firefly luciferase gene, the native regulatory sequences were derived from the 5' and 3' UTRs of histone H1 and pyruvate kinase, respectively (**Online Methods – Online Methods Table 1**). Electroporation with 5 different settings was tested to deliver circular and linear plasmids into the cells (**Suppl. Table 4**), yet there was no evidence of successful transformation, as no luciferase activity was detected by Sirius luminometer after 5 or 24 h. The same negative result was obtained following the delivery of tetramethylrhodamine-labelled DNA using electroporation and biolistics. Finally, 1 µg of *in vitro* synthesized firefly luciferase mRNA was used for electroporation with 4 different settings (**Suppl. Table 4**), and light emission was measured after 24 h, yet again, no specific luciferase activity has been detected.

*Oxyrrhis marina* (CCMP1788) was cultured in f/2 medium with a diverse bacterial community and fed weekly with heat-killed *E. coli*. Selection trials revealed that 5 different antibiotics led to 100% mortality of *O. marina* in 6 days (**Suppl. Table 3**). Fluorescently-labelled DNA or DNA analogs such as FITC-dextran were used to test the efficiency of delivery using a variety of chemical or electrical methods. For instance, incubation with $CaCl_2$ allowed introduction of Alexa 488-labeled DNA (Molecular Probes) with 20% efficiency. Briefly, a mix of 1-8 μg of DNA and $CaCl_2$ (f.c. 0.25 M) was combined with an equal volume of HeBS (274 mM NaCl; 10 mM KCl; 1.4 mM $Na_2HPO_4$; 15 mM D-glucose; 42 mM HEPES, pH 7.1) and incubated with 1 ml of *O. marina* culture. FITC-dextran incorporation was achieved when using Gene Pulser Electroporation Buffer in combination with three 5 ms square-wave pulses (0.1 ms pause between pulses) using a field strength of 0.5 kV/cm (**Suppl. Table 4**). Transient expression of mCherry was observed after introducing a plasmid encoding mCherry gene with a flanking sequence of the *O. marina* hsp90 gene using the above-described transformation protocol (**Table 1; Suppl. Fig. 15A**).

*Hematodinium* sp. belongs to the order Syndiniales and infects the hemolymph of many marine decapod crustaceans. The genus is considered an emerging pathogen for both natural populations and aquacultured crustaceans, with an expanding host range[35,36]. The parasite has multiple stages in the hemolymph[37] and can be continuously maintained in the laboratory[38]. *Hematodinium* cells were transformed using electroporation (Amaxa Nucleofector D-023; BioRad X-100), as well as *via* the glass beads abrasion protocol, the latter having a much higher survival rate (**Suppl. Table 4**). Cells were transformed with 150 kDa FITC-dextran and HEM or Cas9/sgRNA with GFP sequence as DNA donor. We have also tested the use of a synthetic viral RNA derived from a virus known to infect the dinoflagellate *Heterocapsa circularisquama*[39]. Originally, the RNA molecule had 2 ORFs: one that codes for a putative polyprotein bearing a protease and an RNA-dependent RNA polymerase, while the other codes for a capsid protein[40]. We have exchanged the second ORF with an eGFP reporter and *in vitro* transcribed RNA was used to transform *Hematodinium* **(Online Methods – Online Methods Table 1**). However, based on microscopy and PCR verification no clear positive result was observed. Still, *Hematodinium* appears to be a promising organism to transform since in comparison to many dinoflagellates, its cell does not have any theca or cell wall.

*Fugacium* (*Symbiodinium*) *kawagutii* (CCMP 2468) and *Alexandrium catenella* (CCMP BF-5) are both dinoflagellates. *F. kawagutii* is a symbiont originally isolated from the scleractinian coral *Montipora capitate* and has a draft genome available[41]. *A. catenella* is a free-living dinoflagellate and produces potent neurotoxins resulting in paralytic shellfish poisoning[42]. Both species were tested for antibiotic sensitivities (**Suppl. Table 3**). Subsequently, an *Agrobacterium*-mediated transformation method[43] was applied on *F. kawagutii* using the published plasmids, and Basta as the selecting agent. However, this method proved to be difficult to reproduce. Thus, for both species, we developed an electroporation protocol using MicroPulser (BioRad) (**Suppl. Table 4**). Centrifugation protocols that would result in concentrated cells with minimal damage were found to be species-dependent (*F. kawagutii* 800 g for 5 min and *A. catenella* 800 g for l min). Through extensive testing, a suitable transfection solution (10% glycerol in Milli-Q water) that resulted in minimal mortality for both dinoflagellates was identified. Several pulse conditions were tested for each species: Sc2 and ShS performed best for *F. kawagutii* and *A. catenella*, respectively (**Suppl. Table 4**). Plasmids described elsewhere[43] were used in addition to our DinoIII-*gfp*/DinoIII-*pat/*DinoIII-*bsr* vectors[44]. Each plasmid contained either a eGFP reporter, a selection marker, or both. They were introduced into cells as circular or linearized plasmids, and their presence was tested by PCR. The results were inconclusive, species-dependent, with generally very low efficiency. Hence, we tested other methods such as biolistics on *F. kawagutii* and *A. catenella* and electroporation with the Nucleofector (Lonza) on *A. catenella* (**Suppl. Table 4; Online Methods**). For biolistics, a method developed previously[17] was appied using various rupture discs and microcarriers. Due to the difficulty of growing dinoflagellates on agar, 3 μm Millipore filters were used to collect cells and were placed on agarose plates during the biolistic procedure and were then transferred to liquid media for recovery and selection. While *A. catenella* and *F. kawagutii* performed best with 900 PSI and 1550 PSI rupture discs, respectively, the results were inconclusive.

The pyrimidine analog 5-fluoroorotic acid (5FOA) has been shown to successfully suppress the growth of *Breviolum (Symbiodinium)* sp. (**Suppl. Table 3**). A 5FOA-resistant uracil-requiring mutant strain was isolated, which possesses a splice variant mutation in the URA3 gene[45]. We therefore tried to develop a method to complement the phenotypes (i.e., uracil requirement and 5FOA resistance) of this URA3 mutant with the wild type homolog. The full-length URA3 gene was amplified by PCR and cloned into pBlueScript. So far multiple attempts to transform the mutant with this plasmid, either linearized or circular, using electroporator NEPA21 failed (**Suppl. Table 4**). However, we have successfully introduced fluorescein into *Breviolum* sp., using the same method as the DNA-based transformation (**Online Methods – Online Methods Table 1**), suggesting that homologous recombination alone might not be effective enough to replace the mutated version with the wild type sequence.

The cell cycle of *Crypthecodinium cohnii* consists of motile G1 cells, which encyst when they shed their flagella completing the remaining cell-cycle phases[46]. After cytokinesis, the daughter cells remain inside the mother cell, and in this stage, treatment with PEG results in the release of the non-motile spheroplasts[47]. Transfection of *C. cohnii* (CCMP316) was attempted using physical (electroporation, microfluidics, particle bombardment) and chemical (lipofection) methods in motile daughter cells and spheroplasts with both electroporation and lipofection resulting in DNA delivery. *C. cohnii* spheroplasts were electroporated with 5 µg of plasmid using Amaxa Cell Line Optimization Solution V or 3R buffer and program D-023 in Nucleofector (Lonza). The swimming cells and spheroplasts were resuspended in 1 mg of FITC-Dextran and 300 µl of glass beads, vortexed at maximum speed for 15-30 s and subsequently recovered in fresh medium. Biolistics was also tried on 2.5 x $10^7$ swimming cells and spheroplasts using 7.5 µg of HEM plasmid, FITC-dextran, 3 µg of Cas9/sgRNA and 7.5 µg of GFP-carrying plasmid. Cells were precipitated in the presence of gold beads and shooting was carried out in a vacuum at >25 Hg using a 1550 psi rupture disk. Plasmids from *P. marinus* (PmMOE:GFP-11)[48], *Hematodinium* sp. (UB-GFP and EF-GFP) under human ubiquitin promoters, PAY and PAYCO using *E. huxleyi* ubiquitin promotors, purified Cas9 protein with synthetic sgRNA and donor DNA template with the GFP gene and site-directed homology flanking sequences resulted in no fluorescent cells independently of the delivery method attempted. We also tested transformation using chemically labeled DNA (Alexa Fluor 488) and FITC-labeled 150 kDa dextran. *C. cohnii* withstands electroporation with low or no damage, and its swimming cells were difficult to disrupt with glass-bead abrasion and particle bombardment, whereas spheroplasts were much more sensitive. Electroporation of spheroplasts was successful using the Lonza program X-001 or lipofection, both with labeled DNA (**Suppl. Fig. 15B; Online Methods – Online Methods Table 1**), but expression of encoded genes was not confirmed, and the number of transfected cells was very low. We also attempted electroporation and microfluidics (5 square waves and 6 exponential decays)[49] with plasmids PmMOE:GFP-11, UB-GFP, EF-GFP, PAY and PAYCO but with no positive outcome.

**Discobids**

*Eutreptiella gymnastica* (SCCAP K-0333) is a marine green euglenophyte distantly related to the recently sequenced model euglenid, *Euglena gracilis*[50]. All attempts to transform *E. gymnastica* with either the firefly luciferase mRNA or 2 circular plasmids failed. Four methods for DNA delivery were tested, and their efficiency was checked by labelled DNA (**Suppl**. **Fig. 14; Suppl. Table 4**). Following electroporation with 5 µg of *in vitro* synthesized mRNA for firefly luciferase (10 different settings tested) (**Suppl. Table 4**), light emission was measured after 3.5, 5.5, and 22 h, yet no specific luciferase activity has been detected.

The antibiotic selection by 10 µg/ml of puromycin was used for transformations based on plasmids (**Suppl. Table 3**). One plasmid contained the firefly luciferase gene inserted between *E. gymnastica*-derived 300 nt 5' and 3' UTRs of the histone 2A gene. The other plasmid contained this region complemented with the puromycin resistance gene flanked upstream by 5' UTR region of histone-lysine N-methyltransferase and downstream by 3' UTR region of the chloroplast light-harvesting complex II gene from *E. gymnastica*. Four methods for delivery of these 2 plasmids were performed, namely electroporation, lipofection and particle bombardment-mediated delivery of circular or linearized plasmids (**Suppl. Tables 4**; **Online Methods – Online Methods Table 1**). However, we were unable to identify any transformants under selective conditions and no light emission was measured in cultures post-DNA delivery. Since none of the transformation methods was successful, we have proceeded to the delivery of fluorescein- or tetramethylrhodamine-labelled DNA using all 4 delivery methods listed above. In case of electroporation and lipofection, a seemingly intracellular fluorescent signal could be detected suggesting the DNA delivery succeeded, although expression was not achieved (**Suppl. Fig. 14**).

**Opisthokonts**

For the ichthyosporean *Pirum gemmata* we performed electroporation by Neon (Invitrogen) and calcium phosphate transfection approaches using FITC-Dextran. For another ichthyosporean *Sphaeroforma artica* homologous promoters were used for transfection, yet with no success. Although transfection of *S. arctica* did not result in successful DNA delivery (**Suppl. Table 4**), we identified 4 antibiotics (benomyl, carboxine, phleomycin and puromycin), which might be used for future work with *S. arctica* (**Suppl. Table 3; Online Methods – Online Methods Table 1**).

**Transformation of natural protist communities**

Generally, genetic transformation has been achieved in a relatively small number of marine protists, with stable transformation reliably achieved in even fewer of them[51]. Model organisms are typically selected based on well-known criteria, such as relative ease of isolation, cultivation in the laboratory and small genome size. However, these attributes may not correlate with the capacity for uptake and expression of exogenous DNA which nevertheless, is key for genetic tractability. Thus, using a culture-independent, we explored the ability of natural marine planktonic pico- and nanoeukaryotic communities approach to incorporate and express exogenous DNA.

We used a strategy based on electroporation and flow cytometry, to identify cells from natural communities, which had incorporated and expressed DNA (using FITC-dextran) tagged to the gene encoding the blue fluorescent protein (mTagBFP2). We constructed plasmids (**Online Methods – Online Methods Document and Online Methods Table 1**) expressing mTagBFP2 under the control of the cauliflower mosaic virus 35S (35S) and the

cytomegalovirus (CMV) promoters reported to function in a broad range of eukaryotes including phytoplankton[52], as well as a VCP1 promoter previously described from a pico-stramenopile[53]. Plasmids with mTagBFP2 driven by the 35S and CMV promoters were first validated in human embryonic kidney cells and yeast (not shown), allowing their functionality to be confirmed prior to testing micro-algae cultures and natural communities. The electroporation protocol was developed using both cultured *Nannochloropsis oceanica*, *Isochrysis galbana*, and *Emiliania huxleyi*, as well as natural communities collected and concentrated by tangential flow filtration (Vivaflow). Prior to electroporation, cells were washed over polycarbonate filters with different concentrations of sorbitol, in which they were resuspended. Highest recovery of both natural and cultured phytoplankton was obtained with 800 mM sorbitol (as determined by the staining of dead cells with SYBR Green I after 2 h of incubation in sorbitol concentrations ranging from 400 mM to 1.2 M). We performed preliminary tests to determine the best voltage and time constant parameters for each sample type (cultures and natural populations). Cultured organisms included *Nannochloropsis oceanica*, *Isochrysis galbana*, and *Emiliania huxleyi*. For *E. huxleyi*, we tested a range of voltages (from 9 to 11 kV/cm electric fields) based on what had been described for *N. oceanica* (Killian et al., 2011). We obtained the best results with an electric field of 9 kV/cm (capacitance of 50 μF, resistance of 500 Ω, exponential decay pulse). For *I. galbana*, 5 kV/cm with a 20 ms time constant was chosen after preliminary tests. For natural communities, as it is not possible to optimize for all potential members, we tested electric fields from 1 kV/cm to 9.5 kV/cm, with a resistance of 550 Ω (capacitance of 50 μF and exponential decay pulse), and from 10 to 12.5 kV/cm with time constant of 10 ms (capacitance of 50 μF). We defined 2 protocols from those assays to be used in parallel: an electric field of 5 kV/cm with a 20 ms time constant, and an electric field of 10 kV/cm with an 8 ms time constant. Electroporated samples were allowed to recover in moderate (<50 μmol phot/m/s) light at about 15˚C and followed for up to 72 h.
Electroporation efficiency was monitored with 10 mg/ml of 2000 kDa FITC-dextran, with its concentration lowered to 0.5 mg/ml for *I. galbana*. Electroporated cells were analyzed using an InFlux flow cytometer (BD Biosciences) equipped with a 488 nm principal laser (for determining forward and side scatter as well as green fluorescence from FITC-dextran at 530 ± 20 nm), a 405 nm laser (for blue fluorescence at 460 ± 25 nm), and a 640 nm laser (for red fluorescence from chlorophyll at 692 ± 20 nm).

The protocol was first tested with plasmids and cultures of *N. oceanica*, *I. galbana*, and *E. huxleyi*, always comparing cells electroporated with plasmids to those electroporated with the same protocol but without plasmids (negative control). In both *N. oceanica* and *I. galbana*, blue fluorescent cells appeared within several hours. However, the percentage of cells successfully transformed varied greatly, ranging from a few percent to >70%, and, similarly, the percentage of blue fluorescent cells varied from ~1% to over 20%. In *N. oceanica*, the CMV promoter was effective at driving blue fluorescence, with >20% transformants when a linearized plasmid was used (**Table 1; Suppl. Fig. 6; Online Methods**), but only <2% when the plasmid was circular (**Table 1**). The 35S promoter resulted in inconsistent results with both linearized and circular plasmids, and in 2 trials there was no change in blue fluorescence with the VCP1 promoter. In *I. galbana*, the 35S promoter resulted in blue fluorescence peaking in up to 20% of cells at 4 h and declining thereafter. In calcified *E. huxleyi* cultures, electroporation both with and without plasmid resulted in inconsistent survival, and fluorescence and light scatter signatures became much dispersed, making interpretation impossible.

In 3 out of 6 trials with natural communities, blue-fluorescent pico-eukaryotes appeared in samples electroporated with plasmids encoding mTagBFP2. In one of these natural communities, an important portion of a particular pico-phytoeukaryote (grouped based on optical properties) exhibited blue fluorescence. Indeed, 58 to 68% cells fluoresced when electroporated at 5 kV/cm after 2 h with a plasmid containing the CMV promoter. Curiously, a portion of these cells (up to 18% at 12 h after electroporation) showed blue fluorescence when exposed to plasmid DNA even in the absence of electroporation, although the time course was markedly slower (**Suppl. Fig. 13**). Unfortunately, when sorting for culturing was prepared at the 24 h time point, the signal of blue fluorescence was so diminished that we could only sort pico-phytoplankton with similar light scattering and pigment autofluorescence characteristic for live cultures. However, based on two replications, our initial results were not reproducible in the cultured strain obtained.

We further confirmed the expression of the plasmid under the control of the CMV promoter by *N. oceanica*. To do so, cells were electroporated with the plasmid and 100 cells showing blue fluorescence were sorted in a lysis buffer (NucleoSpin RNA XS, Macherey-Nagel) and immediately stored at -80˚C. Cells incubated with the plasmid but not electroporated were also sorted and used as negative controls. RNA was extracted using the NucleoSpin RNA XS (Macherey-Nagel) with the DNA digestion step extended to 80 min. A 2 step protocol was used, composed of the reverse transcription with the SuperScript™ III reverse transcriptase (ThermoFisher Scientific), followed by the DNA amplification of the BFP fragment from the plasmid with the GoTaq Flexi DNA polymerase (Promega) in the same reaction mix. An amplicon of the expected size (750 bp) was recovered from the cells that had been electroporated but not from the non-electroporated cells (**Suppl. Fig. 6**). In another electroporation trial with natural samples, blue fluorescent cells from samples electroporated with plasmids and red fluorescent chlorophyll-containing cells from samples incubated with plasmids but not electroporated, were sorted into TE buffer (100 cells in 100 μL of buffer) and stored at -80˚C until further processing. Due to the length of time it took to sort 100 of the rare blue cells seen in this trial, DNA amplification was performed instead of RNA amplification. PCR amplification of the mTagBFP2 fragment was performed directly on the cells in TE buffer in two replicates per sample (10 μL of cells sorted in TE for 20 μL reaction mix volume). In the case of the CMV_BFP_pMOD plasmid, the expected product size was successfully obtained for 2 technical replicates of the cells electroporated at 5 kV/cm and for one replicate of the cells electroporated at 10 kV/cm (**Suppl. Fig. 6; Online Methods**). In the case of the 35S_BFP_pMOD plasmid, one one technical replicate for each protocol was clearly successful, and for the 35S_BFP_pGreen plasmid no amplicon was seen. No DNA amplification occurred from the cells incubated with the plasmid without electroporation.

Taken together, transforming natural marine planktonic protist communities is challenging because of the following issues. First, the natural abundance of protist cells is usually very low (< 1000/ml), and therefore they should be concentrated by i.e. tangential flow filtration. Second, the salts from seawater need to be removed for electroporation, resuspending the cells in sorbitol. Finally, it is not possible to optimize the electroporation protocol for a natural community, as even when the same location is sampled repeatedly, the community will likely be different due to the dynamics of the marine system. For the latter reason, in most trials we used 2 different electroporation protocols in parallel on distinct sub-samples. The variable and low percentage of apparently transformed cells presents a challenge for cell sorting, for transfer into media, for determining identity-independent culturing, and for confirming the expression of the eGFP gene. A recently published method based on single cell sorting and subsequent culturing may help to address these issues[53]. Although amplification of barcoding genes (e.g. 18S rRNA) from sorted populations is becoming more routine, we were unable to obtain amplicons from populations of pico-phytoeukaryotes that exhibited a high percentage of blue fluorescent cells, suggestion an alternative protocol is required for transformed populations.

**Open Material Transfer Agreement (OpenMTA)**
For sharing biological material (e.g. cells and DNA), the available Material Transfer Agreement (MTA)[55] is not well suited for the needs of collaborative projects aiming at the development of community-sourced toolkits. The shortcomings of traditional MTAs have recently been described[55]. Although MTA barriers could be bypassed by *de novo* synthesis of DNA, this would be very impractical and costly. Instead, we highly recommend the adoption of the OpenMTA; https://www.openplant.org/openmta/) for future community-wide efforts. An OpenMTA enables more expedited processing times, decreased administrative transaction costs, broader sharing, use and redistribution of biological materials, while maintaining the rights of creators and promoting safe practices and responsible research. Perhaps most importantly, OpenMTA allows tracking the provenance and impact of projects[56] and is more compatible with public and philanthropic funding policies, as well as with open science practices.

**Supplementary Notes 1: List of plasmid maps and sequences used for particular species following the order from Fig. 1 are shown below.** (For links to protocols.io webpage where the detailed protocols can be find see Table 1 and Suppl. Table 5).

#### Archaeoplastids

##### *Ostreococcus lucimarinus*

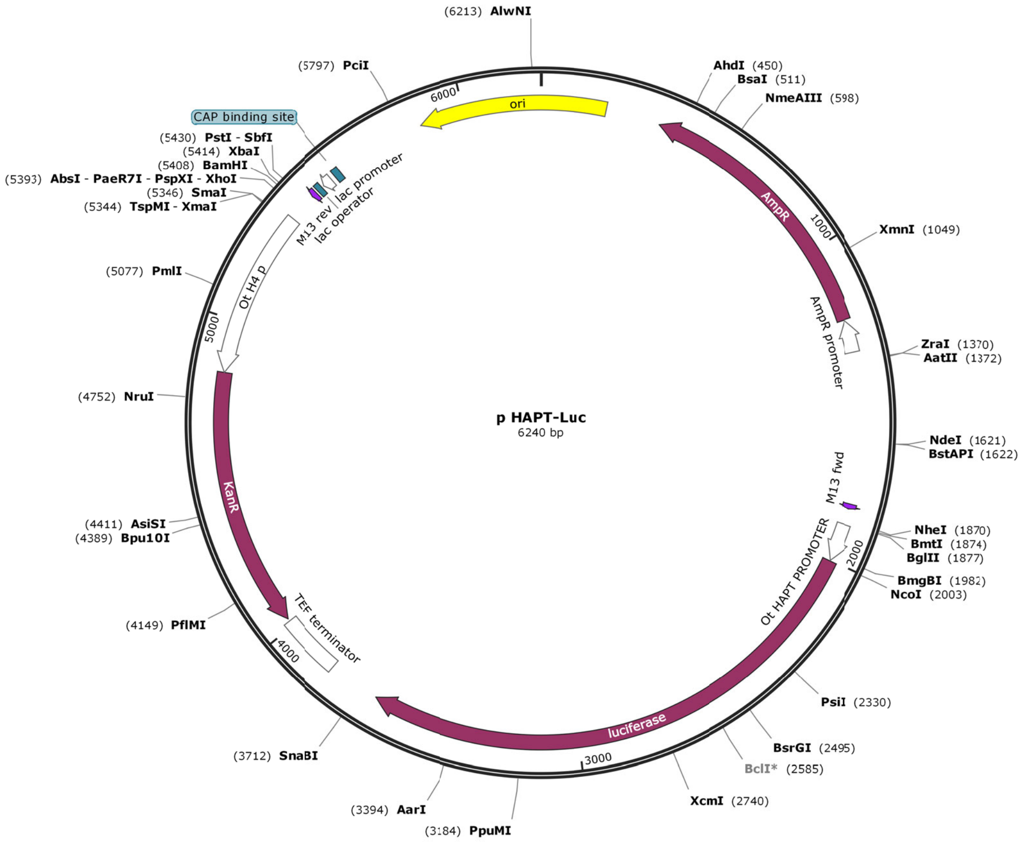


**PHAPT:Luc sequence**

```
O. tauri pHAPT:Luc                6240 bp ds-DNA     circular
DEFINITION  synthetic circular DNA

FEATURES             Location/Qualifiers
     source          1..6240
                     /organism="synthetic DNA construct"
                     /mol_type="other DNA"
     CDS             complement(377..1237)
                     /codon_start=1
                     /gene="bla"
                     /product="beta-lactamase"
                     /label=AmpR
                     /note="confers resistance to ampicillin,
               carbenicillin, and
                     related antibiotics"
                     /translation="MSIQHFRVALIPFFAAFCLPVFAHPETLVKVKDAEDQLGARVGYI
```

```
                     ELDLNSGKILESFRPEERFPMMSTFKVLLCGAVLSRIDAGQEQLGRRIHYSQNDLVEYS
                     PVTEKHLTDGMTVRELCSAAITMSDNTAANLLLTTIGGPKELTAFLHNMGDHVTRLDRW
                     EPELNEAIPNDERDTTMPVAMATTLRKLLTGELLTLASRQQLIDWMEADKVAGPLLRSA
                     LPAGWFIADKSGAGERGSRGIIAALGPDGKPSRIVVIYTTGSQATMDERNRQIAEIGAS
                     LIKHW"
     promoter        complement(1238..1342)
                     /gene="bla"
                     /label=AmpR promoter
     primer_bind     1816..1832
                     /label=M13 fwd
                     /note="common sequencing primer, one of multiple similar
                     variants"
     promoter        1883..2002
                     /label=Ostreococcus tauri High affinity
               phosphate transporte promoter
     CDS             2005..3657
                     /codon_start=1
                     /gene="luc+"
                     /product="firefly luciferase"
                     /label=luciferase
                     /note="enhanced luc+ version of the luciferase gene"
                     /translation="MEDAKNIKKGPAPFYPLEDGTAGEQLHKAMKRYALVPGTIAFTDA
                     HIEVDITYAEYFEMSVRLAEAMKRYGLNTNHRIVVCSENSLQFFMPVLGALFIGVAVAP
                     ANDIYNERELLNSMGISQPTVVFVSKKGLQKILNVQKKLPIIQKIIIMDSKTDYQGFQS
                     MYTFVTSHLPPGFNEYDFVPESFDRDKTIALIMNSSGSTGLPKGVALPHRTACVRFSHA
                     RDPIFGNQIIPDTAILSVVPFHHGFGMFTTLGYLICGFRVVLMYRFEEELFLRSLQDYK
                     IQSALLVPTLFSFFAKSTLIDKYDLSNLHEIASGGAPLSKEVGEAVAKRFHLPGIRQGY
                     GLTETTSAILITPEGDDKPGAVGKVVPFFEAKVVDLDTGKTLGVNQRGELCVRGPMIMS
                     GYVNNPEATNALIDKDGWLHSGDIAYWDEDEHFFIVDRLKSLIKYKGYQVAPAELESIL
                     LQHPNIFDAGVAGLPDDDAGELPAAVVVLEHGKTMTEKEIVDYVASQVTTAKKLRGGVV
                     FVDEVPKGLTGKLDARKIREILIKAKKGGKIAV"
     terminator      3820..4017
                     /label=TEF terminator
                     /note="Ashbya gossypii TEF terminator"
     CDS             complement(4026..4835)
                     /codon_start=1
                     /gene="aph(3')-Ia"
                     /product="aminoglycoside phosphotransferase"
                     /label=KanR
                     /note="confers resistance to kanamycin"
                     /translation="MGKEKTHVSRPRLNSNMDADLYGYKWARDNVGQSGATIYRLYGKP
                     DAPELFLKHGKGSVANDVTDEMVRLNWLTEFMPLPTIKHFIRTPDDAWLLTTAIPGKTA
                     FQVLEEYPDSGENIVDALAVFLRRLHSIPVCNCPFNSDRVFRLAQAQSRMNNGLVDASD
                     FDDERNGWPVEQVWKEMHKLLPFSPDSVVTHGDFSLDNLIFDEGKLIGCIDVGRVGIAD
                     RYQDLAILWNCLGEFSPSLQKRLFQKYGIDNPDMNKLQFHLMLDEFF"
     primer_bind     complement(5456..5472)
                     /label=M13 rev
                     /note="common sequencing primer, one of multiple similar
                     variants"
     protein_bind    5480..5496
                     /label=lac operator
                     /bound_moiety="lac repressor encoded by lacI"
                     /note="The lac repressor binds to the lac operator to
                     inhibit transcription in E. coli. This inhibition can be
                     relieved by adding lactose or
                     isopropyl-beta-D-thiogalactopyranoside (IPTG)."
     promoter        complement(5504..5534)
                     /label=lac promoter
                     /note="promoter for the E. coli lac operon"
     protein_bind    5549..5570
                     /label=CAP binding site
                     /bound_moiety="E. coli catabolite activator protein"
                     /note="CAP binding activates transcription in the presence
                     of cAMP."
     rep_origin      complement(join(5858..6240,1..206))
                     /direction=LEFT
                     /label=ori
                     /note="high-copy-number ColE1/pMB1/pBR322/pUC origin of
                     replication"
ORIGIN
        1 atgtaggcgg tgctacagag ttcttgaagt ggtggcctaa ctacggctac actagaagga
       61 cagtatttgg tatctgcgct ctgctgaagc cagttacctt cggaaaaaga gttggtagct
      121 cttgatccgg caaacaaacc accgctggta gcggtggttt ttttgtttgc aagcagcaga
      181 ttacgcgcag aaaaaaagga tctcaagaag atcctttgat cttttctacg gggtctgacg
      241 ctcagtggaa cgaaaactca cgttaaggga ttttggtcat gagattatca aaaaggatct
      301 tcacctagat ccttttaaat taaaaatgaa gttttaaatc aatctaaagt atatatgagt
      361 aaacttggtc tgacagttac caatgcttaa tcagtgaggc acctatctca gcgatctgtc
```

421 tatttcgttc atccatagtt gcctgactcc ccgtcgtgta gataactacg atacgggagg
481 gcttaccatc tggccccagt gctgcaatga taccgcgaga cccacgctca ccggctccag
541 atttatcagc aataaaccag ccagccggaa gggccgagcg cagaagtggt cctgcaactt
601 tatccgcctc catccagtct attaattgtt gccgggaagc tagagtaagt agttcgccag
661 ttaatagttt gcgcaacgtt gttgccattg ctacaggcat cgtggtgtca cgctcgtcgt
721 ttggtatggc ttcattcagc tccggttccc aacgatcaag gcgagttaca tgatccccca
781 tgttgtgcaa aaaagcggtt agctccttcg gtcctccgat cgttgtcaga agtaagttgg
841 ccgcagtgtt atcactcatg gttatggcag cactgcataa ttctcttact gtcatgccat
901 ccgtaagatg cttttctgtg actggtgagt actcaaccaa gtcattctga gaatagtgta
961 tgcggcgacc gagttgctct tgcccggcgt caatacggga taataccgcg ccacatagca
1021 gaactttaaa agtgctcatc attggaaaac gttcttcggg gcgaaaactc tcaaggatct
1081 taccgctgtt gagatccagt tcgatgtaac ccactcgtgc acccaactga tcttcagcat
1141 cttttacttt caccagcgtt tctgggtgag caaaaacagg aaggcaaaat gccgcaaaaa
1201 agggaataag ggcgacacgg aaatgttgaa tactcatact cttccttttt caatattatt
1261 gaagcattta tcagggttat tgtctcatga gcggatacat atttgaatgt atttagaaaa
1321 ataaacaaat aggggttccg cgcacatttc cccgaaaagt gccacctgac gtctaagaaa
1381 ccattattat catgacatta acctataaaa ataggcgtat cacgaggccc tttcgtctcg
1441 cgcgtttcgg tgatgacggt gaaaacctct gacacatgca gctcccggag acggtcacag
1501 cttgtctgta agcggatgcc gggagcagac aagcccgtca gggcgcgtca gcgggtgttg
1561 gcgggtgtcg gggctggctt aactatgcgg catcagagca gattgtactg agagtgcacc
1621 atatgcggtg tgaaataccg cacagatgcg taaggagaaa ataccgcatc aggcgccatt
1681 cgccattcag gctgcgcaac tgttgggaag ggcgatcggt gcgggcctct tcgctattac
1741 gccagctggc gaaaggggga tgtgctgcaa ggcgattaag ttgggtaacg ccagggtttt
1801 cccagtcacg acgttgtaaa acgacggcca gtgaattcga gctcggtacc cttcatctcc
1861 ggttctgctg ctagcaagat ctaaatactt ttccaggaaa agcaggaaaa aggaatattc
1921 tttaatattc tcggaatatt tcacgttctt gcggcctggc gacgctgacg gcgttacttc
1981 acgtcgcgct ttacccgtct cgccatggaa gacgccaaaa acataaagaa aggcccggcg
2041 ccattctatc cgctggaaga tggaaccgct ggagagcaac tgcataaggc tatgaagaga
2101 tacgccctgg ttcctggaac aattgctttt acagatgcac atatcgaggt ggacatcact
2161 tacgctgagt acttcgaaat gtccgttcgg ttggcagaag ctatgaaacg atatgggctg
2221 aatacaaatc acagaatcgt cgtatgcagt gaaaactctc ttcaattctt tatgccggtg
2281 ttgggcgcgt tatttatcgg agttgcagtt gcgcccgcga acgacattta taatgaacgt
2341 gaattgctca acagtatggg catttcgcag cctaccgtgg tgttcgtttc caaaaagggg
2401 ttgcaaaaaa ttttgaacgt gcaaaaaaag ctcccaatca tccaaaaaat tattatcatg
2461 gattctaaaa cggattacca gggatttcag tcgatgtaca cgttcgtcac atctcatcta
2521 cctcccggtt ttaatgaata cgattttgtg ccagagtcct tcgataggga caagacaatt
2581 gcactgatca tgaactcctc tggatctact ggtctgccta aaggtgtcgc tctgcctcat
2641 agaactgcct gcgtgagatt ctcgcatgcc agagatccta tttttggcaa tcaaatcatt
2701 ccggatactg cgattttaag tgttgttcca ttccatcacg gttttggaat gtttactaca
2761 ctcggatatt tgatatgtgg atttcgagtc gtcttaatgt atagatttga agaagagctg
2821 tttctgagga gccttcagga ttacaagatt caaagtgcgc tgctggtgcc aaccctattc
2881 tccttcttcg ccaaaagcac tctgattgac aaatacgatt tatctaattt acacgaaatt
2941 gcttctggtg gcgctcccct ctctaaggaa gtcggggaag cggttgccaa gaggttccat
3001 ctgccaggta tcaggcaagg atatgggctc actgagacta catcagctat tctgattaca
3061 cccgaggggg atgataaacc gggcgcggtc ggtaaagttg ttccattttt tgaagcgaag
3121 gttgtggatc tggataccgg gaaaacgctg ggcgttaatc aaagaggcga actgtgtgtg
3181 agaggtccta tgattatgtc cggttatgta aacaatccgg aagcgaccaa cgccttgatt
3241 gacaaggatg gatggctaca ttctggagac atagcttact gggacgaaga cgaacacttc
3301 ttcatcgttg accgcctgaa gtctctgatt aagtacaaag gctatcaggt ggctcccgct
3361 gaattggaat ccatcttgct ccaacacccc aacatcttcg acgcaggtgt cgcaggtctt
3421 cccgacgatg acgccggtga acttcccgcc gccgttgttg ttttggagca cggaaagacg
3481 atgacggaaa aagagatcgt ggattacgtc gccagtcaag taacaaccgc gaaaaagttg
3541 cgcggaggag ttgtgtttgt ggacgaagta ccgaaaggtc ttaccggaaa actcgacgca
3601 agaaaaatca gagagatcct cataaaggcc aagaagggcg gaaagatcgc cgtgtaattc
3661 tagctaaaaa gtggaacgat cattcactat atatatatca atttatatat acgtatgtgt
3721 aattgaagaa agatacgttt ttcctctat tgagaggatc atcgatgaat tcgagctcgt
3781 tttcgacact ggatggcggc gttagtatcg aatcgacagc agtatagcga ccagcattca
3841 catacgattg acgcatgata ttactttctg cgcacttaac ttcgcatctg ggcagatgat
3901 gtcgaggcga aaaaaaatat aaatcacgct aacatttgat taaaatagaa caactacaat
3961 ataaaaaaac tatacaaatg acaagttctt gaaaacaaga atctttttat tgtcagtact
4021 gattattaga aaaactcatc gagcatcaaa tgaaactgca atttattcat atcaggatta
4081 tcaataccat atttttgaaa aagccgtttc tgtaatgaag gagaaaactc accgaggcag
4141 ttccatagga tggcaagatc ctggtatcgg tctgcgattc cgactcgtcc aacatcaata
4201 caacctatta atttcccctc gtcaaaaata aggttatcaa gtgagaaatc accatgagtg
4261 acgactgaat ccggtgagaa tggcaaaagc ttatgcattt ctttccagac ttgttcaaca
4321 ggccagccat tacgctcgtc atcaaaatca ctcgcatcaa ccaaaccgtt attcattcgt
4381 gattgcgcct gagcgagacg aaatacgcga tcgctgttaa aaggacaatt acaaacagga
4441 atcgaatgca accggcgcag gaacactgcc agcgcatcaa caatattttc acctgaatca
4501 ggatattctt ctaatacctg gaatgctgtt ttgccgggga tcgcagtggt gagtaaccat
4561 gcatcatcag gagtacggat aaaatgcttg atggtcggaa gaggcataaa ttccgtcagc
4621 cagtttagtc tgaccatctc atctgtaaca tcattggcaa cgctaccttt gccatgtttc
4681 agaaacaact ctggcgcatc gggcttccca tacaatcgat agattgtcgc acctgattgc
4741 ccgacattat cgcgagccca tttataccca tataaatcag catccatgtt ggaatttaat
4801 cgcggcctcg aaacgtgagt cttttcctta cccatgcatg gttgttttcg aacgtgtcga
4861 ttagtgacga cctgcgcgcg cggtcggggc cacggacgcg ggcgccggtg agttcgcggt
4921 gagttgaaaa tactccgtgc gagtcacgac ccatgaaatt ataggggtaa aaagttcatt
4981 tcacacaatc tttgatgcgt tgattgtttc taaatggcac acaatgagtc atgaaccaca

5041 atgcgtggat ttaaaaacaa tcaagtgatg catgcacgtg aaaaatgggc atttcaagta
5101 aatagagagt aatgactcgt gtatgacccc ctgctttcaa actcaccgac tcaccggcgc
5161 gcgcggtgaa acgctcgatt cgatcgcacg gcgtcgtcac aactcgacag tcaatatggc
5221 ccgtacaaag caaactgccc gtaagtccac cgggggcaag gccccgcgca agcaactcgc
5281 caccaaggcg gcccgcaagt ccgccccggc gaccggcggc gtcaagaagc cgcacaggta
5341 ccgcccgggt accgttgcgc tcgtgggatc tttcacctct tagtcttctg gcctcgaggt
5401 cgacggggga tcctctagag tcgacctgca ggcatgcaag cttggcgtaa tcatggtcat
5461 agctgtttcc tgtgtgaaat tgttatccgc tcacaattcc acacaacata cgagccggaa
5521 gcataaagtg taaagcctgg ggtgcctaat gagtgagcta actcacatta attgcgttgc
5581 gctcactgcc cgctttccag tcgggaaacc tgtcgtgcca gctgcattaa tgaatcggcc
5641 aacgcgcggg gagaggcggt ttgcgtattg ggcgctcttc cgcttcctcg ctcactgact
5701 cgctgcgctc ggtcgttcgg ctgcggcgag cggtatcagc tcactcaaag gcggtaatac
5761 ggttatccac agaatcaggg gataacgcag gaaagaacat gtgagcaaaa ggccagcaaa
5821 aggccaggaa ccgtaaaaag gccgcgttgc tggcgttttt ccataggctc cgcccccctg
5881 acgagcatca caaaaatcga cgctcaagtc agaggtggcg aaacccgaca ggactataaa
5941 gataccaggc gtttccccct ggaagctccc tcgtgcgctc tcctgttccg accctgccgc
6001 ttaccggata cctgtccgcc tttctccctt cgggaagcgt ggcgctttct caatgctcac
6061 gctgtaggta tctcagttcg gtgtaggtcg ttcgctccaa gctgggctgt gtgcacgaac
6121 cccccgttca gcccgaccgc tgcgccttat ccggtaacta tcgtcttgag tccaacccgg
6181 taagacacga cttatcgcca ctggcagcag ccactggtaa caggattagc agagcgaggt

Validation of *O. lucimarinus* transgenic lines by PCR (Figure 4A)

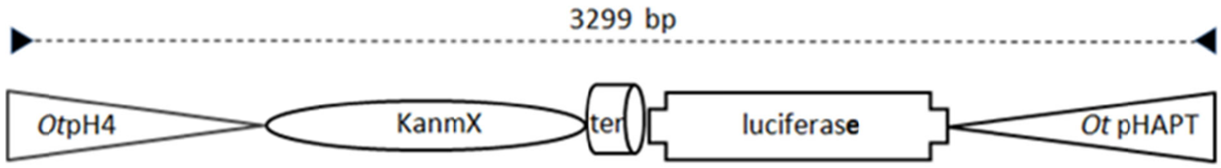


Forward primer: 1GAGCGCAACGGTACCCGGGCGGTACCTGTGCG
Reverse primer: CTGGCGACGCTGACGGCGTTACTTCACGT

### *Bathycoccus prasinos*

**pH4:KanMx pHAPT:Luc** transgene (PCR product)

Construction of  the transgene by PCR :

The transgene was generated by fusion PCR as described in Shevchuk et al. (2004) using DNA templates described below and oligonucleotides mapped on the sequence below. **Promoters of Histone H4 and high affinity phosphate (HAPT) transporters** were amplified using (Fas1/Ras1) and (Fas4/Ras4)oligonucucleotides respectively. KanMx and Luciferase sequences were amplified from pHAPT:luc plasmid template using oligonucleotides (Fas2/Ras2) and (Fas3/Ras3). Final PCR was done on amplified fragments using (Fas5/Ras5) oligonucleotides.

**Oligonucleotide sequences:**
Fas1:CCGGCTTCGTGATGCCTTGGATGTTGTCTC
Ras1: TCGAAACGTGAGTCTTTTCCTTACCcatTGTGTTTGATTTATAATGAGGTTTTCTT
Fas2: AAGAAAACCTCATTATAAATCAAACACAatgGGTAAGGAAAAGACTCACGTTTCGA
Ras2: CCAAGAAGGGCGGAAAGATCGCCGTGTAAGAAATACCGTCTATCATCGATGAATTCGA
Fas3: CGAATTCATCGATGATAGACGGTATTTCTTACACGGCGATCTTTCCGCCCTTCTTGG
Ras3: AAGCATATATACATACACACACATACAAAATGGAAGACGCCAAAAACATAAAGAAAGG
Fas4: CCTTTCTTTATGTTTTTGGCGTCTTCCATTTTGTATGTGTGTGTATGTATATATGCTT
Ras4: AAGTGTACACGACCAAACGCGCGTCGAC
Fas5: CCACCTTTACCTCTGCCGGACATTGTGA
Ras5: GCGCGGTAATATCTACGAGGTAGCACGAG

Note that transformants were checked by PCR using ( to amplify Kanx

Map of the transgene:

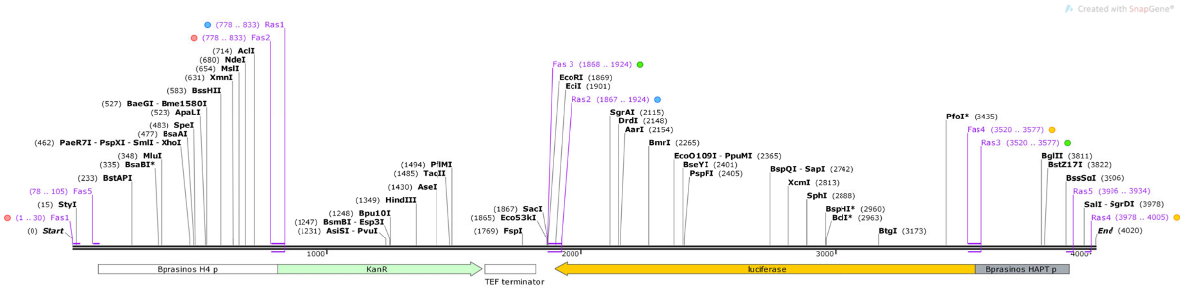


**Bprasinos transgene sequence**
4020 bp

Sequence of the transgene:

    4020 bp ds-DNA     linear

REFERENCE   1  (bases 1 to 4020)
  AUTHORS   FY Bouget and JC Lozano

FEATURES             Location/Qualifiers
     source          1..4020
                     /organism="unspecified"
                     /mol_type="genomic DNA"
     primer_bind     1..30
                     /label=Fas1
     primer_bind     78..105
                     /label=Fas5
     promoter        101..805
                     /label=Bprasinos H4 p
     primer_bind     complement(778..833)
                     /label=Ras1
     primer_bind     778..833
                     /label=Fas2
     CDS             806..1615
                     /codon_start=1
                     /gene="aph(3')-Ia"
                     /product="aminoglycoside phosphotransferase"
                     /label=KanR
                     /note="confers resistance to kanamycin"
                     /translation="MGKEKTHVSRPRLNSNMDADLYGYKWARDNVGQSGATIYRLYGKP
                     DAPELFLKHGKGSVANDVTDEMVRLNWLTEFMPLPTIKHFIRTPDDAWLLTTAIPGKTA

```
                     FQVLEEYPDSGENIVDALAVFLRRLHSIPVCNCPFNSDRVFRLAQAQSRMNNGLVDASD
                     FDDERNGWPVEQVWKEMHKLLPFSPDSVVTHGDFSLDNLIFDEGKLIGCIDVGRVGIAD
                     RYQDLAILWNCLGEFSPSLQKRLFQKYGIDNPDMNKLQFHLMLDEFF"
     terminator      1624..1821
                     /label=TEF terminator
                     /note="Ashbya gossypii TEF terminator"
     primer_bind     complement(1867..1924)
                     /label=Ras2
     primer_bind     1868..1924
                     /label=Fas 3
     CDS             complement(1896..3548)
                     /codon_start=1
                     /gene="luc+"
                     /product="firefly luciferase"
                     /label=luciferase
                     /note="enhanced luc+ version of the luciferase gene"
                     /translation="MEDAKNIKKGPAPFYPLEDGTAGEQLHKAMKRYALVPGTIAFTDA
                     HIEVDITYAEYFEMSVRLAEAMKRYGLNTNHRIVVCSENSLQFFMPVLGALFIGVAVAP
                     ANDIYNERELLNSMGISQPTVVFVSKKGLQKILNVQKKLPIIQKIIIMDSKTDYQGFQS
                     MYTFVTSHLPPGFNEYDFVPESFDRDKTIALIMNSSGSTGLPKGVALPHRTACVRFSHA
                     RDPIFGNQIIPDTAILSVVPFHHGFGMFTTLGYLICGFRVVLMYRFEEELFLRSLQDYK
                     IQSALLVPTLFSFFAKSTLIDKYDLSNLHEIASGGAPLSKEVGEAVAKRFHLPGIRQGY
                     GLTETTSAILITPEGDDKPGAVGKVVPFFEAKVVDLDTGKTLGVNQRGELCVRGPMIMS
                     GYVNNPEATNALIDKDGWLHSGDIAYWDEDEHFFIVDRLKSLIKYKGYQVAPAELESIL
                     LQHPNIFDAGVAGLPDDDAGELPAAVVVLEHGKTMTEKEIVDYVASQVTTAKKLRGGVV
                     FVDEVPKGLTGKLDARKIREILIKAKKGGKIAV"
     primer_bind     complement(3520..3577)
                     /label=Ras3
     primer_bind     3520..3577
                     /label=Fas4
     PROMOTER   3549..3920
                     /label=B. prasinos High affinity
                phosphate transporter
     primer_bind     complement(3906..3934)
                     /label=Ras5
     primer_bind     complement(3978..4005)
                     /label=Ras4
ORIGIN
        1 ccggcttcgt gatgccttgg atgttgtctc tcaacacctt gcggtgtctt ttggcgccac
       61 ctttacctaa acctttacca cctttacctc tgccggacat tgtgaatatg tgttatattt
      121 gtttttaaat gacgatttga acggacgttt tcgttttgtt tttcgtgttt ttttgtgttt
      181 tttgcctgtt ttgtaggggt tacttctcag ttttgctata aagagtgcat ttgatgcatc
      241 aaaacgagcg tttgagaagg tcgcgagtga aaaagctacg aagacgaaat tcaatcaacg
      301 acaagaaaat acaacaaaaa gaacaacctc gatccttatc acttggaacg cgtatcacgt
      361 tttaaaagaa ataattatac aatataactc ctcgtaacca tctatcatcc ctatatctat
      421 cgcatcagaa ctttttagct tttcatttgc tctccgagtt cctcgagcgt cgaatacgtg
      481 cgactagttg tgcatgtaca accctaaatg atagtgtgca aagtgcacct ttttttcgca
      541 aatgctcgaa aatgcaagac cttaaatatt cggcgacatt aagcgcgctg aaaggtacgt
      601 tactgttcac ggatttgatc gcgcgtgaac ttgttcgata gatatttaac atatctgtgt
      661 ctttgagaag ttttgattca tatgaatcag tgatacatct gatgtcgtac aaaacgttaa
      721 atcttgattt ctaaccttct tttgttttcg aatggttcaa atgaatcgtt gacacgaaag
      781 aaaacctcat tataaatcaa acacaatggg taaggaaaag actcacgttt cgaggccgcg
      841 attaaattcc aacatggatg ctgatttata tgggtataaa tgggctcgcg ataatgtcgg
      901 gcaatcaggt gcgacaatct atcgattgta tgggaagccc gatgcgccag agttgtttct
      961 gaaacatggc aaaggtagcg ttgccaatga tgttacagat gagatggtca gactaaactg
     1021 gctgacggaa tttatgcctc ttccgaccat caagcatttt atccgtactc ctgatgatgc
     1081 atggttactc accactgcga tccccggcaa aacagcattc caggtattag aagaatatcc
     1141 tgattcaggt gaaaatattg ttgatgcgct ggcagtgttc ctgcgccggt tgcattcgat
     1201 tcctgtttgt aattgtcctt ttaacagcga tcgcgtattt cgtctcgctc aggcgcaatc
     1261 acgaatgaat aacggtttgg ttgatgcgag tgattttgat gacgagcgta atggctggcc
     1321 tgttgaacaa gtctggaaag aaatgcataa gcttttgcca ttctcaccgg attcagtcgt
     1381 cactcatggt gatttctcac ttgataacct tatttttgac gaggggaaat taataggttg
     1441 tattgatgtt ggacgagtcg gaatcgcaga ccgataccag gatcttgcca tcctatggaa
     1501 ctgcctcggt gagttttctc cttcattaca gaaacggctt tttcaaaaat atggtattga
     1561 taatcctgat atgaataaat tgcagtttca tttgatgctc gatgagtttt tctaataatc
     1621 agtactgaca ataaaaagat tcttgttttc aagaacttgt catttgtata gtttttttat
     1681 attgtagttg ttctatttta atcaaatgtt agcgtgattt atattttttt tcgcctcgac
     1741 atcatctgcc cagatgcgaa gttaagtgcg cagaaagtaa tatcatgcgt caatcgtatg
     1801 tgaatgctgg tcgctatact gctgtcgatt cgatactaac gccgccatcc agtgtcgaaa
     1861 acgagctcga attcatcgat gatagacggt atttcttaca cggcgatctt tccgcccttc
     1921 ttggccttta tgaggatctc tctgattttt cttgcgtcga gttttccggt aagacctttc
     1981 ggtacttcgt ccacaaacac aactcctccg cgcaactttt tcgcggttgt tacttgactg
     2041 gcgacgtaat ccacgatctc tttttccgtc atcgtctttc cgtgctccaa aacaacaacg
     2101 gcggcgggaa gttcaccggc gtcatcgtcg ggaagacctg cgacacctgc gtcgaagatg
     2161 ttggggtgtt ggagcaagat ggattccaat tcagcgggag ccacctgata gcctttgtac
     2221 ttaatcagag acttcaggcg gtcaacgatg aagaagtgtt cgtcttcgtc ccagtaagct
     2281 atgtctccag aatgtagcca tccatccttg tcaatcaagg cgttggtcgc ttccggattg
```

```
     2341 tttacataac cggacataat cataggacct ctcacacaca gttcgcctct ttgattaacg
     2401 cccagcgttt tcccggtatc cagatccaca accttcgctt caaaaaatgg aacaacttta
     2461 ccgaccgcgc ccggtttatc atccccctcg ggtgtaatca gaatagctga tgtagtctca
     2521 gtgagcccat atccttgcct gatacctggc agatggaacc tcttggcaac cgcttccccg
     2581 acttccttag agaggggagc gccaccagaa gcaatttcgt gtaaattaga taaatcgtat
     2641 ttgtcaatca gagtgctttt ggcgaagaag gagaataggg ttggcaccag cagcgcactt
     2701 tgaatcttgt aatcctgaag gctcctcaga aacagctctt cttcaaatct atacattaag
     2761 acgactcgaa atccacatat caaatatccg agtgtagtaa acattccaaa accgtgatgg
     2821 aatggaacaa cacttaaaat cgcagtatcc ggaatgattt gattgccaaa aataggatct
     2881 ctggcatgcg agaatctcac gcaggcagtt ctatgaggca gagcgacacc tttaggcaga
     2941 ccagtagatc cagaggagtt catgatcagt gcaattgtct tgtccctatc gaaggactct
     3001 ggcacaaaat cgtattcatt aaaaccggga ggtagatgag atgtgacgaa cgtgtacatc
     3061 gactgaaatc cctggtaatc cgttttagaa tccatgataa taatttttg gatgattggg
     3121 agcttttttt gcacgttcaa aatttttgc aacccctttt tggaaacgaa caccacggta
     3181 ggctgcgaaa tgcccatact gttgagcaat tcacgttcat tataaatgtc gttcgcgggc
     3241 gcaactgcaa ctccgataaa taacgcgccc aacaccggca taaagaattg aagagagttt
     3301 tcactgcata cgacgattct gtgatttgta ttcagcccat atcgtttcat agcttctgcc
     3361 aaccgaacgg acatttcgaa gtactcagcg taagtgatgt ccacctcgat atgtgcatct
     3421 gtaaaagcaa ttgttccagg aaccagggcg tatctcttca tagccttatg cagttgctct
     3481 ccagcggttc catcttccag cggatagaat ggcgccgggc ctttctttat gtttttggcg
     3541 tcttccattt tgtatgtgtg tgtatgtata tatgctttgg gaatatatgt tcacagaatg
     3601 acgactttga aagcgcgttt gaatttttaa acgaaaatct ccgtgtggct gatatttttt
     3661 gctttttgct tttttcaac cacccggatt tttgcttttt ttcaaaacaa cccaccgacc
     3721 gtaaatgttg tgtgttcttg tttctgttgg ggctgctttc ttttagagga gggaggatgc
     3781 attcagagtt aatatattat atgtgctccg agatctgtgg tatacgagga gttggtgttg
     3841 gcttttaat acacaaaata cgcctaaacg cgaggaggcg tcgttgaaac gtaaaggtac
     3901 tattactcgt gctacctcgt agatattacc gcgcgtaatt agaagtcgtg ggagttgttg
     3961 tcgttgtcgt tgtattcgtc gacgcgcgtt tggtcgtgta cactttactc gcgccgcgac
//
```

Validation of *B. prasinos* transgenic lines by PCR (Figure 4b)

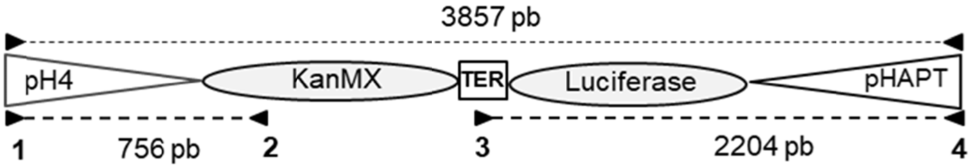


**Oligonucleotide sequences**

1 (Fas5) CCACCTTTACCTCTGCCGGACATTGTGA

2 TCGAAACGTGAGTCTTTTCCTTACC

3 TCGCCTCGACATCATCTGCCCAGATGC

4 (Ras5) GCGCGGTAATATCTACGAGGTAGCACGAG

<u>Reference:</u>

Nikolai A. Shevchuk, Anton V. Bryksin, Yevgeniya A. Nusinovich, Felipe C. Cabello, Margaret Sutherland, Stephan Ladisch /Nucleic Acids Research/, Volume 32, Issue 2, 16 January 2004, Page e19, https://doi.org/10.1093/nar/gnh014

### *Micromonas commoda*

PS9proMco-eGFP

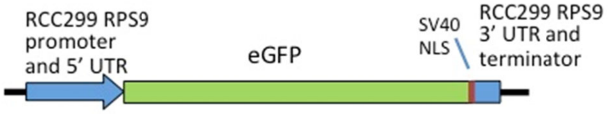


RPS9proMco-eGFP-NLS construct. The ribosomal protein S9 (RPS9) gene (found at Chr_03:1,087,263-1,088,617 of the March 2015 version3b assembly of the RCC299 genome) was used by taking the sequence and replacing the RPS9 coding region including the introns with a *Micromonas* codon optimized coding sequence for eGFP. The construct was synthesized by IDT in its entirety using an ampicillin resistant plasmid backbone.

```
LOCUS       RPS9_eGFP_SV40NL        3987 bp ds-DNA     circular

DEFINITION
ACCESSION
VERSION
SOURCE
ORGANISM
COMMENT

FEATURES             Location/Qualifiers

     misc_feature    2823..3075
                     /locus_tag="Insert-EcoHindIII"
                      /label="Insert-EcoHindIII"
     misc_feature    2823..3075
                     /locus_tag="RPS9 promoter from 299"
                     /label="RPS9 promoter from 299"
     misc_feature    3817..3976
                     /locus_tag="RPS9 Termination region 299"
                     /label="RPS9 Termination region 299"
     misc_feature    3817..3976
                     /locus_tag="Insert-EcoHindIII(1)"
                     /label="Insert-EcoHindIII(1)"
     CDS             3076..3816
                     /locus_tag="eGFP ORF"
                     /label="eGFP ORF"
ORIGIN

        1 GGAAGTTTGT CTAGATCTCA GGCGTGGATG ATCAGTTCTG GACCAGCGAG CTGTGCTGCG
       61 ACTCGTGGCG TAATCATGGT CATAGCTGTT TCCTGTGTGA AATTGTTATC CGCTCACAAT
      121 TCCACACAAC ATACGAGCCG GAAGCATAAA GTGTAAAGCC TGGGGTGCCT AATGAGTGAG
      181 CTAACTCACA TTAATTGCGT TGCGCTCACT GCCCGCTTTC CAGTCGGGAA ACCTGTCGTG
      241 CCAGCTGCAT TAATGAATCG GCCAACGCGC GGGGAGAGGC GGTTTGCGTA TTGGGCGCTC
      301 TTCCGCTTCC TCGCTCACTG ACTCGCTGCG CTCGGTCGTT CGGCTGCGGC GAGCGGTATC
      361 AGCTCACTCA AAGGCGGTAA TACGGTTATC CACAGAATCA GGGGATAACG CAGGAAAGAA
      421 CATGTGAGCA AAAGGCCAGC AAAAGGCCAG GAACCGTAAA AAGGCCGCGT TGCTGGCGTT
      481 TTTCCATAGG CTCCGCCCCC CTGACGAGCA TCACAAAAAT CGACGCTCAA GTCAGAGGTG
      541 GCGAAACCCG ACAGGACTAT AAAGATACCA GGCGTTTCCC CCTGGAAGCT CCCTCGTGCG
      601 CTCTCCTGTT CCGACCCTGT CGCTTACCGG ATACCTGTCC GCCTTTCTCC CTTCGGGAAG
      661 CGTGGCGCTT TCTCATAGCT CACGCTGTAG GTATCTCAGT TCGGTGTAGG TCGTTCGCTC
      721 CAAGCTGGGC TGTGTGCACG AACCCCCCGT TCAGCCCGAC CGCTGCGCCT TATCCGGTAA
      781 CTATCGTCTT GAGTCCAACC CGGTAAGACA CGACTTATCG CCACTGGCAG CAGCCACTGG
      841 TAACAGGATT AGCAGAGCGA GGTATGTAGG CGGTGCTACA GAGTTCTTGA AGTGGTGGCC
      901 TAACTACGGC TACACTAGAA GAACAGTATT TGGTATCTGC GCTCTGCTGA AGCCAGTTAC
      961 CTTCGGAAAA AGAGTTGGTA GCTCTTGATC CGGCAAACAA ACCACCGCTG GTAGCGGTGG
     1021 TTTTTTTGTT TGCAAGCAGC AGATTACGCG CAGAAAAAAA GGATCTCAAG AAGATCCTTT
     1081 GATCTTTTCT ACGGGGTCTG ACGCTCAGTG GAACGAAAAC TCACGTTAAG GGATTTTGGT
     1141 CATGAGATTA TCAAAAAGGA TCTTCACCTA GATCCTTTTA AATTAAAAAT GAAGTTTTAA
     1201 ATCAATCTAA AGTATATATG AGTAAACTTG GTCTGACAGT TACCAATGCT TAATCAGTGA
     1261 GGCACCTATC TCAGCGATCT GTCTATTTCG TTCATCCATA GTTGCCTGAC TCCCCGTCGT
     1321 GTAGATAACT ACGATACGGG AGGGCTTACC ATCTGGCCCC AGTGCTGCAA TGATACCGCG
```

```
1381 AGACCCACGC TCACCGGCTC CAGATTTATC AGCAATAAAC CAGCCAGCCG GAAGGGCCGA
1441 GCGCAGAAGT GGTCCTGCAA CTTTATCCGC CTCCATCCAG TCTATTAATT GTTGCCGGGA
1501 AGCTAGAGTA AGTAGTTCGC CAGTTAATAG TTTGCGCAAC GTTGTTGCCA TTGCTACAGG
1561 CATCGTGGTG TCACGCTCGT CGTTTGGTAT GGCTTCATTC AGCTCCGGTT CCCAACGATC
1621 AAGGCGAGTT ACATGATCCC CCATGTTGTG CAAAAAAGCG GTTAGCTCCT TCGGTCCTCC
1681 GATCGTTGTC AGAAGTAAGT TGGCCGCAGT GTTATCACTC ATGGTTATGG CAGCACTGCA
1741 TAATTCTCTT ACTGTCATGC CATCCGTAAG ATGCTTTTCT GTGACTGGTG AGTACTCAAC
1801 CAAGTCATTC TGAGAATAGT GTATGCGGCG ACCGAGTTGC TCTTGCCCGG CGTCAATACG
1861 GGATAATACC GCGCCACATA GCAGAACTTT AAAAGTGCTC ATCATTGGAA AACGTTCTTC
1921 GGGGCGAAAA CTCTCAAGGA TCTTACCGCT GTTGAGATCC AGTTCGATGT AACCCACTCG
1981 TGCACCCAAC TGATCTTCAG CATCTTTTAC TTTCACCAGC GTTTCTGGGT GAGCAAAAAC
2041 AGGAAGGCAA AATGCCGCAA AAAAGGGAAT AAGGGCGACA CGGAAATGTT GAATACTCAT
2101 ACTCTACCTT TTTCAATATT ATTGAAGCAT TTATCAGGGT TATTGTCTCA TGAGCGGATA
2161 CATATTTGAA TGTATTTAGA AAAATAAACA AATAGGGGTT CCGCGCACAT TTCCCCGAAA
2221 AGTGCCACCT GACGTCTAAG AAACCATTAT TATCATGACA TTAACCTATA AAAATAGGCG
2281 TATCACGAGG CCCTTTCGTC TCGCGCGTTT CGGTGATGAC GGTGAAAACC TCTGACACAT
2341 GCAGCTCCCG GAGACGGTCA CAGCTTGTCT GTAAGCGGAT GCCGGGAGCA GACAAGCCCG
2401 TCAGGGCGCG TCAGCGGGTG TTGGCGGGTG TCGGGGCTGG CTTAACTATG CGGCATCAGA
2461 GCAGATTGTA CTGAGAGTGC ACCAAATGCG GTGTGAAATA CCGCACAGAT GCGTAAGGAG
2521 AAAATACCGC ATCAGGCGCC ATTCGCCATT CAGGCTGCGC AACTGTTGGG AAGGGCGATC
2581 GGTGCGGGCC TCATCGCTAT TACGCCAGCT GGCGAAAGGG GGATGTGCTG CAAGGCGATT
2641 AAGTTGGGTA ACGCCAGGGT TTTCCCAGTC ACGACGTTGT AAAACGACGG CCAGTGCAAC
2701 GCGATGACGA TGGATAGCGA TTCATCGATG AGCTGACCCG ATCGCCGCCG CCGGAGGGTT
2761 GCGTTTGAGA CGGGCGACAG ATGAGCACAT CTCGTTCGCT ATTCAGGGAT TGTTAATTAA
2821 CAGAATTCTC GTGGATTTTT GACAAAAAGC GATCTTTACC AGTGAGATCG ACCCATTTGC
2881 AGTTGACCAT CCACTTCGTT TTCCCATTAT TTCCTATTTC GAGGTTCGCA CAGTATGTGT
2941 TCATGACAGT TAGTGAGCTT CGTTCAAAAC AGGTTCTGGT CGGATTTTCC TGGCCGGACT
3001 TTCCGCTAAA ACCCTCGTCG ACCCAGTCGT CCCGCCCTCT TACACCTAGC GCAGGAAGGC
3061 GAAGGGACAG CCAAGatggt ctcgaagggc gaggagctct tcacgggcgt cgtcccgatc
3121 ctcgtcgagc tcgacggcga cgtcaacggc cacaagttct cggtctcggg cgagggcgag
3181 ggcgacgcga cgtacggcaa gctcacgctc aagttcatct gcacgacggg caagctcccg
3241 gtcccgtggc cgacgctcgt cacgacgctc acgtacggcg tccagtgctt ctcgcgctac
3301 ccggaccaca tgaagcagca cgacttcttc aagtcggcga tgccggaggg ctacgtccag
3361 gagcgcacga tcttcttcaa ggacgacggc aactacaaga cgcgcgcgga ggtcaagttc
3421 gagggcgaca cgctcgtcaa ccgcatcgag ctcaagggca tcgacttcaa ggaggacggc
3481 aacatcctcg gccacaagct cgagtacaac tacaactcgc acaacgtcta catcatggcg
3541 gacaagcaga agaacggcat caaggtcaac ttcaagatcc gccacaacat cgaggacggc
3601 tcggtccagc tcgcggacca ctaccagcag aacacgccga tcggcgacgg cccggtcctc
3661 ctcccggaca accactacct ctcgacgcag tcggcgctct cgaaggaccc gaacgagaag
3721 cgcgaccaca tggtcctcct cgagttcgtc acggcggcgg gcatcacgct cggcatggac
3781 gagctctaca agccgaagaa gaagcgcaag gtctgaACAC CCGAGCTGAG GGCCGGGGGA
3841 TCTTCCACGC GGCGTCTAAG TGACGACGCC AGGCCTTGCG CGCCGATCTC ACGAGAGTGG
3901 CGTGCAATTT ACTAACAGTA GTGACTTGAA ATACACGGGT AACTCCTGTC CACAAACGAA
3961 CCCGCGCGAG AAGCTTGCGT TAATTAA
```

H3proMpo-LUC

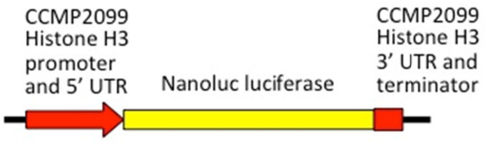


H3proMpo-Luc construct. The histone H3 gene (found at scaffold_9:539,958-541,019 of the July 2017 assembly of the CCMP2099 genome) was used by taking the sequence and replacing the histone coding region with a *Micromonas* codon optimized coding sequence for the Nanoluc Luciferase. Sequence was synthesized by IDT and cloned into the EcoRI site of pUC13.

```
LOCUS       Mp_HH3pro_NanoLu        3931 bp ds-DNA     circular
DEFINITION  .
ACCESSION
VERSION
SOURCE      .
  ORGANISM  .
COMMENT
FEATURES             Location/Qualifiers
     terminator      1656..1661
                     /locus_tag="Rps93'UTR and terminator"
                     /label="Rps93'UTR and terminator"
     misc_feature    483..488
                     /locus_tag="EcoRI"
                     /label="EcoRI"
     misc_feature    1656..1661
                     /locus_tag="EcoRI(1)"
                     /label="EcoRI(1)"
     misc_feature    489..905
                     /locus_tag="histone H3 promoter CCMP2099"
                     /label="histone H3 promoter CCMP2099"
     misc_feature    1422..1655
                     /locus_tag="HistoneH3_term"
                     /label="HistoneH3_term"
     misc_feature    906..1421
                     /locus_tag="NanoLuc optimized coding region"
                     /label="NanoLuc optimized coding region"

ORIGIN
        1 TCGCGCGTTT CGGTGATGAC GGTGAAAACC TCTGACACAT GCAGCTCCCG GAGACGGTCA
       61 CAGCTTGTCT GTAAGCGGAT GCCGGGAGCA GACAAGCCCG TCAGGGCGCG TCAGCGGGTG
      121 TTGGCGGGTG TCGGGGCTGG CTTAACTATG CGGCATCAGA GCAGATTGTA CTGAGAGTGC
      181 ACCAAATGCG GTGTGAAATA CCGCACAGAT GCGTAAGGAG AAAATACCGC ATCAGGCGCC
      241 ATTCGCCATT CAGGCTGCGC AACTGTTGGG AAGGGCGATC GGTGCGGGCC TCATCGCTAT
      301 TACGCCAGCT GGCGAAAGGG GGATGTGCTG CAAGGCGATT AAGTTGGGTA ACGCCAGGGT
      361 TTTCCCAGTC ACGACGTTGT AAAACGACGG CCAGTGCAAC GCGATGACGA TGGATAGCGA
      421 TTCATCGATG AGCTGACCCG ATCGCCGCCG CCGGAGGGTT GCGTTTGAGA CGGGCGACAG
      481 ATgaattcCT TTACCTCTGC CGGACATTGT GTGAGTTGTT TCGTGTGTGT ACGAAAGTGA
      541 TGGGTGAAGG AGTCGCGGGG AGGCTTTTAG CCAACCTGCT TTTCGAGTTG GTGGGGACGT
      601 TTTGACCCGA GCCAAAGATG CGAATCCTGT TAATGTTGTA ACGTCAATTC ATGCAGAGGG
      661 TGGCGGCGGG TACGCGATGG TTTTAGTTGT GGTATGTGAG GAATCCCTCA TATGACACAC
      721 AACGATGCTT GCGGGATTCC GCATAGGACC AGTCCAAAGA CGTATAAACC CCACGAAGCC
      781 GACGTTTTAA AAAAGAAACA GGATTCGCAA AAAAGCCCCA AAAATCGCCA AAACTGCAAG
      841 TTTCCCCACC TACTTCAAAC CCGTTCGCCA CTTCCCCTCG TCACAACCCC CCGCACTCTC
      901 TCAAGATGGT CTTCACGCTC GAGGACTTCG TCGGCGACTG GCGCCAGACG GCGGGCTACA
      961 ACCTCGACCA GGTCCTCGAG CAGGGCGGCG TCTCGTCGCT CTTCCAGAAC CTCGGCGTCT
     1021 CGGTCACGCC GATCCAGCGC ATCGTCCTCT CGGGCGAGAA CGGCCTCAAG ATCGACATCC
     1081 ACGTCATCAT CCCGTACGAG GGCCTCTCGG GCGACCAGAT GGGCCAGATC GAGAAGATCT
     1141 TCAAGGTCGT CTACCCGGTC GACGACCACC ACTTCAAGGT CATCCTCCAC TACGGCACGC
     1201 TCGTCATCGA CGGCGTCACG CCGAACATGA TCGACTACTT CGGCCGCCCG TACGAGGGCA
     1261 TCGCGGTCTT CGACGGCAAG AAGATCACGG TCACGGGCAC GCTCTGGAAC GGCAACAAGA
     1321 TCATCGACGA GCGCCTCATC AACCCGGACG GCTCGCTCCT CTTCCGCGTC ACGATCAACG
     1381 GCGTCACGGG CTGGCGCCTC TGCGAGCGCA TCCTCGCGTG AGCGCTGTGC CGACGCACGT
     1441 TCCGTCCAGT ACGAACACAT AACCAACAAC TCACAATGGT GTTAACAAAT TAAACACCAC
     1501 CACTTCTTTA AAGAACCAAA CGCAACACGT ATATACGCGA GCGTCGAACC GAGCGAGCGC
     1561 GCTTGAATCC GTAAAAAAAA ACAACGGCTA GAAAATTCAA ATCGAAAAGG CAGACTTTCT
     1621 AGATGACGAA TTTTTGTCGC AATTAAACGG TTCGTgaatt cATCAGTTCT GGACCAGCGA
     1681 GCTGTGCTGC GACTCGTGGC GTAATCATGG TCATAGCTGT TTCCTGTGTG AAATTGTTAT
     1741 CCGCTCACAA TTCCACACAA CATACGAGCC GGAAGCATAA AGTGTAAAGC CTGGGGTGCC
     1801 TAATGAGTGA GCTAACTCAC ATTAATTGCG TTGCGCTCAC TGCCCGCTTT CCAGTCGGGA
     1861 AACCTGTCGT GCCAGCTGCA TTAATGAATC GGCCAACGCG CGGGGAGAGG CGGTTTGCGT
     1921 ATTGGGCGCT CTTCCGCTTC CTCGCTCACT GACTCGCTGC GCTCGGTCGT TCGGCTGCGG
     1981 CGAGCGGTAT CAGCTCACTC AAAGGCGGTA ATACGGTTAT CCACAGAATC AGGGGATAAC
     2041 GCAGGAAAGA ACATGTGAGC AAAAGGCCAG CAAAAGGCCA GGAACCGTAA AAAGGCCGCG
     2101 TTGCTGGCGT TTTTCCATAG GCTCCGCCCC CCTGACGAGC ATCACAAAAA TCGACGCTCA
     2161 AGTCAGAGGT GGCGAAACCC GACAGGACTA TAAAGATACC AGGCGTTTCC CCCTGGAAGC
     2221 TCCCTCGTGC GCTCTCCTGT TCCGACCCTG TCGCTTACCG GATACCTGTC CGCCTTTCTC
     2281 CCTTCGGGAA GCGTGGCGCT TTCTCATAGC TCACGCTGTA GGTATCTCAG TTCGGTGTAG
     2341 GTCGTTCGCT CCAAGCTGGG CTGTGTGCAC GAACCCCCCG TTCAGCCCGA CCGCTGCGCC
     2401 TTATCCGGTA ACTATCGTCT TGAGTCCAAC CCGGTAAGAC ACGACTTATC GCCACTGGCA
     2461 GCAGCCACTG GTAACAGGAT TAGCAGAGCG AGGTATGTAG GCGGTGCTAC AGAGTTCTTG
     2521 AAGTGGTGGC CTAACTACGG CTACACTAGA AGAACAGTAT TTGGTATCTG CGCTCTGCTG
     2581 AAGCCAGTTA CCTTCGGAAA AAGAGTTGGT AGCTCTTGAT CCGGCAAACA AACCACCGCT
     2641 GGTAGCGGTG GTTTTTTTGT TTGCAAGCAG CAGATTACGC GCAGAAAAAA AGGATCTCAA
     2701 GAAGATCCTT TGATCTTTTC TACGGGGTCT GACGCTCAGT GGAACGAAAA CTCACGTTAA
     2761 GGGATTTTGG TCATGAGATT ATCAAAAAGG ATCTTCACCT AGATCCTTTT AAATTAAAAA
```

2821 TGAAGTTTTA AATCAATCTA AAGTATATAT GAGTAAACTT GGTCTGACAG TTACCAATGC
2881 TTAATCAGTG AGGCACCTAT CTCAGCGATC TGTCTATTTC GTTCATCCAT AGTTGCCTGA
2941 CTCCCCGTCG TGTAGATAAC TACGATACGG GAGGGCTTAC CATCTGGCCC CAGTGCTGCA
3001 ATGATACCGC GAGACCCACG CTCACCGGCT CCAGATTTAT CAGCAATAAA CCAGCCAGCC
3061 GGAAGGGCCG AGCGCAGAAG TGGTCCTGCA ACTTTATCCG CCTCCATCCA GTCTATTAAT
3121 TGTTGCCGGG AAGCTAGAGT AAGTAGTTCG CCAGTTAATA GTTTGCGCAA CGTTGTTGCC
3181 ATTGCTACAG GCATCGTGGT GTCACGCTCG TCGTTTGGTA TGGCTTCATT CAGCTCCGGT
3241 TCCCAACGAT CAAGGCGAGT TACATGATCC CCCATGTTGT GCAAAAAAGC GGTTAGCTCC
3301 TTCGGTCCTC CGATCGTTGT CAGAAGTAAG TTGGCCGCAG TGTTATCACT CATGGTTATG
3361 GCAGCACTGC ATAATTCTCT TACTGTCATG CCATCCGTAA GATGCTTTTC TGTGACTGGT
3421 GAGTACTCAA CCAAGTCATT CTGAGAATAG TGTATGCGGC GACCGAGTTG CTCTTGCCCG
3481 GCGTCAATAC GGGATAATAC CGCGCCACAT AGCAGAACTT TAAAAGTGCT CATCATTGGA
3541 AAACGTTCTT CGGGGCGAAA ACTCTCAAGG ATCTTACCGC TGTTGAGATC CAGTTCGATG
3601 TAACCCACTC GTGCACCCAA CTGATCTTCA GCATCTTTTA CTTTCACCAG CGTTTCTGGG
3661 TGAGCAAAAA CAGGAAGGCA AAATGCCGCA AAAAAGGGAA TAAGGGCGAC ACGGAAATGT
3721 TGAATACTCA TACTCTACCT TTTTCAATAT TATTGAAGCA TTTATCAGGG TTATTGTCTC
3781 ATGAGCGGAT ACATATTTGA ATGTATTTAG AAAAATAAAC AAATAGGGGT TCCGCGCACA
3841 TTTCCCCGAA AAGTGCCACC TGACGTCTAA GAAACCATTA TTATCATGAC ATTAACCTAT
3901 AAAAATAGGC GTATCACGAG GCCCTTTCGT C

*Micromonas pusilla*

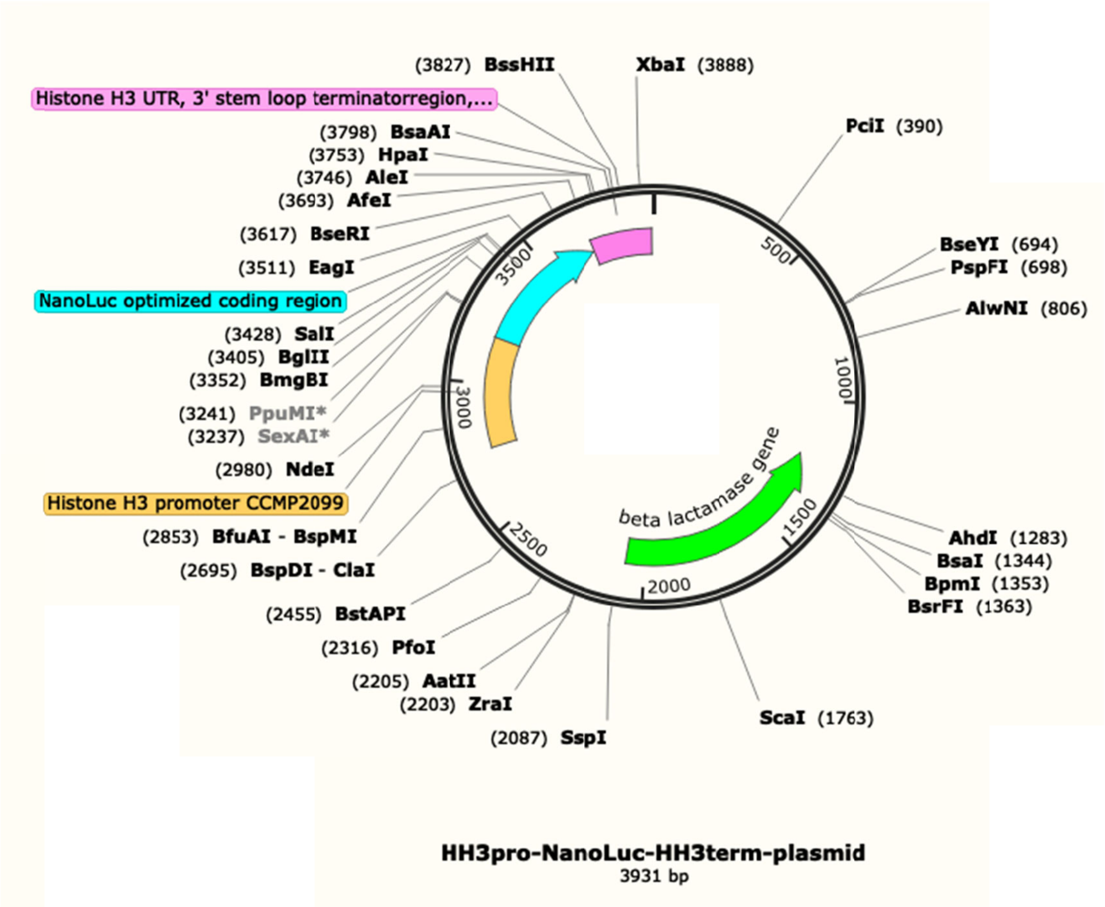


```
LOCUS       HH3pro_NanoLuc_H        3931 bp ds-DNA     circular     26-OCT-2019
DEFINITION  .
ACCESSION
VERSION
SOURCE      .
  ORGANISM  .
COMMENT
COMMENT
COMMENT     ApEinfo:methylated:1
FEATURES             Location/Qualifiers
     misc_feature    2759..3175
                     /locus_tag="Histone H3 promoter CCMP2099"
                     /label="Histone H3 promoter CCMP2099"
                     /ApEinfo_label="Histone H3 promoter CCMP2099"
                     /ApEinfo_fwdcolor="#fecf6b"
                     /ApEinfo_revcolor="green"
                     /ApEinfo_graphicformat="arrow_data {{0 1 2 0 0 -1} {} 0}
                     width 5 offset 0"
     misc_feature    3692..3925
                     /locus_tag="Histone H3 UTR, 3' stem loop terminator
                     region, CCMP2099"
                     /label="Histone H3 UTR, 3' stem loop terminator region,
                     CCMP2099"
                     /ApEinfo_label="Histone H3 UTR, 3' stem loop terminator
                     region, CCMP2099"
                     /ApEinfo_fwdcolor="#fc85e7"
                     /ApEinfo_revcolor="green"
                     /ApEinfo_graphicformat="arrow_data {{0 1 2 0 0 -1} {} 0}
                     width 5 offset 0"
     CDS             3176..3691
                     /locus_tag="NanoLuc optimized coding region"
                     /label="NanoLuc optimized coding region"
```

```
                     /ApEinfo_label="NanoLuc optimized coding region"
                     /ApEinfo_fwdcolor="cyan"
                     /ApEinfo_revcolor="green"
                     /ApEinfo_graphicformat="arrow_data {{0 1 2 0 0 -1} {} 0}
                     width 5 offset 0"
     CDS             complement(1210..2070)
                     /locus_tag="beta lactamase gene"
                     /label="beta lactamase gene"
                     /ApEinfo_label="beta lactamase gene"
                     /ApEinfo_fwdcolor="cyan"
                     /ApEinfo_revcolor="green"
                     /ApEinfo_graphicformat="arrow_data {{0 1 2 0 0 -1} {} 0}
                     width 5 offset 0"
ORIGIN
        1 ATCAGTTCTG GACCAGCGAG CTGTGCTGCG ACTCGTGGCG TAATCATGGT CATAGCTGTT
       61 TCCTGTGTGA AATTGTTATC CGCTCACAAT TCCACACAAC ATACGAGCCG GAAGCATAAA
      121 GTGTAAAGCC TGGGGTGCCT AATGAGTGAG CTAACTCACA TTAATTGCGT TGCGCTCACT
      181 GCCCGCTTTC CAGTCGGGAA ACCTGTCGTG CCAGCTGCAT TAATGAATCG GCCAACGCGC
      241 GGGGAGAGGC GGTTTGCGTA TTGGGCGCTC TTCCGCTTCC TCGCTCACTG ACTCGCTGCG
      301 CTCGGTCGTT CGGCTGCGGC GAGCGGTATC AGCTCACTCA AAGGCGGTAA TACGGTTATC
      361 CACAGAATCA GGGGATAACG CAGGAAAGAA CATGTGAGCA AAAGGCCAGC AAAAGGCCAG
      421 GAACCGTAAA AAGGCCGCGT TGCTGGCGTT TTTCCATAGG CTCCGCCCCC CTGACGAGCA
      481 TCACAAAAAT CGACGCTCAA GTCAGAGGTG GCGAAACCCG ACAGGACTAT AAAGATACCA
      541 GGCGTTTCCC CCTGGAAGCT CCCTCGTGCG CTCTCCTGTT CCGACCCTGT CGCTTACCGG
      601 ATACCTGTCC GCCTTTCTCC CTTCGGGAAG CGTGGCGCTT TCTCATAGCT CACGCTGTAG
      661 GTATCTCAGT TCGGTGTAGG TCGTTCGCTC CAAGCTGGGC TGTGTGCACG AACCCCCCGT
      721 TCAGCCCGAC CGCTGCGCCT TATCCGGTAA CTATCGTCTT GAGTCCAACC CGGTAAGACA
      781 CGACTTATCG CCACTGGCAG CAGCCACTGG TAACAGGATT AGCAGAGCGA GGTATGTAGG
      841 CGGTGCTACA GAGTTCTTGA AGTGGTGGCC TAACTACGGC TACACTAGAA GAACAGTATT
      901 TGGTATCTGC GCTCTGCTGA AGCCAGTTAC CTTCGGAAAA AGAGTTGGTA GCTCTTGATC
      961 CGGCAAACAA ACCACCGCTG GTAGCGGTGG TTTTTTTGTT TGCAAGCAGC AGATTACGCG
     1021 CAGAAAAAAA GGATCTCAAG AAGATCCTTT GATCTTTTCT ACGGGGTCTG ACGCTCAGTG
     1081 GAACGAAAAC TCACGTTAAG GGATTTTGGT CATGAGATTA TCAAAAAGGA TCTTCACCTA
     1141 GATCCTTTTA AATTAAAAAT GAAGTTTTAA ATCAATCTAA AGTATATATG AGTAAACTTG
     1201 GTCTGACAGT TACCAATGCT TAATCAGTGA GGCACCTATC TCAGCGATCT GTCTATTTCG
     1261 TTCATCCATA GTTGCCTGAC TCCCCGTCGT GTAGATAACT ACGATACGGG AGGGCTTACC
     1321 ATCTGGCCCC AGTGCTGCAA TGATACCGCG AGACCCACGC TCACCGGCTC CAGATTTATC
     1381 AGCAATAAAC CAGCCAGCCG GAAGGGCCGA GCGCAGAAGT GGTCCTGCAA CTTTATCCGC
     1441 CTCCATCCAG TCTATTAATT GTTGCCGGGA AGCTAGAGTA AGTAGTTCGC CAGTTAATAG
     1501 TTTGCGCAAC GTTGTTGCCA TTGCTACAGG CATCGTGGTG TCACGCTCGT CGTTTGGTAT
     1561 GGCTTCATTC AGCTCCGGTT CCCAACGATC AAGGCGAGTT ACATGATCCC CCATGTTGTG
     1621 CAAAAAAGCG GTTAGCTCCT TCGGTCCTCC GATCGTTGTC AGAAGTAAGT TGGCCGCAGT
     1681 GTTATCACTC ATGGTTATGG CAGCACTGCA TAATTCTCTT ACTGTCATGC CATCCGTAAG
     1741 ATGCTTTTCT GTGACTGGTG AGTACTCAAC CAAGTCATTC TGAGAATAGT GTATGCGGCG
     1801 ACCGAGTTGC TCTTGCCCGG CGTCAATACG GGATAATACC GCGCCACATA GCAGAACTTT
     1861 AAAAGTGCTC ATCATTGGAA AACGTTCTTC GGGGCGAAAA CTCTCAAGGA TCTTACCGCT
     1921 GTTGAGATCC AGTTCGATGT AACCCACTCG TGCACCCAAC TGATCTTCAG CATCTTTTAC
     1981 TTTCACCAGC GTTTCTGGGT GAGCAAAAAC AGGAAGGCAA AATGCCGCAA AAAAGGGAAT
     2041 AAGGGCGACA CGGAAATGTT GAATACTCAT ACTCTACCTT TTTCAATATT ATTGAAGCAT
     2101 TTATCAGGGT TATTGTCTCA TGAGCGGATA CATATTTGAA TGTATTTAGA AAAATAAACA
     2161 AATAGGGGTT CCGCGCACAT TTCCCCGAAA AGTGCCACCT GACGTCTAAG AAACCATTAT
     2221 TATCATGACA TTAACCTATA AAAATAGGCG TATCACGAGG CCCTTTCGTC TCGCGCGTTT
     2281 CGGTGATGAC GGTGAAAACC TCTGACACAT GCAGCTCCCG GAGACGGTCA CAGCTTGTCT
     2341 GTAAGCGGAT GCCGGGAGCA GACAAGCCCG TCAGGGCGCG TCAGCGGGTG TTGGCGGGTG
     2401 TCGGGGCTGG CTTAACTATG CGGCATCAGA GCAGATTGTA CTGAGAGTGC ACCAAATGCG
     2461 GTGTGAAATA CCGCACAGAT GCGTAAGGAG AAAATACCGC ATCAGGCGCC ATTCGCCATT
     2521 CAGGCTGCGC AACTGTTGGG AAGGGCGATC GGTGCGGGCC TCATCGCTAT TACGCCAGCT
     2581 GGCGAAAGGG GGATGTGCTG CAAGGCGATT AAGTTGGGTA ACGCCAGGGT TTTCCCAGTC
     2641 ACGACGTTGT AAAACGACGG CCAGTGCAAC GCGATGACGA TGGATAGCGA TTCATCGATG
     2701 AGCTGACCCG ATCGCCGCCG CCGGAGGGTT GCGTTTGAGA CGGGCGACAG ATgaattcCT
     2761 TTACCTCTGC CGGACATTGT GTGAGTTGTT TCGTGTGTGT ACGAAAGTGA TGGGTGAAGG
     2821 AGTCGCGGGG AGGCTTTTAG CCAACCTGCT TTTCGAGTTG GTGGGGACGT TTTGACCCGA
     2881 GCCAAAGATG CGAATCCTGT TAATGTTGTA ACGTCAATTC ATGCAGAGGG TGGCGGCGGG
     2941 TACGCGATGG TTTTAGTTGT GGTATGTGAG GAATCCCTCA TATGACACAC AACGATGCTT
     3001 GCGGGATTCC GCATAGGACC AGTCCAAAGA CGTATAAACC CCACGAAGCC GACGTTTTAA
     3061 AAAAGAAACA GGATTCGCAA AAAAGCCCCA AAAATCGCCA AAACTGCAAG TTTCCCCACC
     3121 TACTTCAAAC CCGTTCGCCA CTTCCCCTCG TCACAACCCC CCGCACTCTC TCAAGATGGT
     3181 CTTCACGCTC GAGGACTTCG TCGGCGACTG GCGCCAGACG GCGGGCTACA ACCTCGACCA
     3241 GGTCCTCGAG CAGGGCGGCG TCTCGTCGCT CTTCCAGAAC CTCGGCGTCT CGGTCACGCC
     3301 GATCCAGCGC ATCGTCCTCT CGGGCGAGAA CGGCCTCAAG ATCGACATCC ACGTCATCAT
     3361 CCCGTACGAG GGCCTCTCGG GCGACCAGAT GGGCCAGATC GAGAAGATCT TCAAGGTCGT
     3421 CTACCCGGTC GACGACCACC ACTTCAAGGT CATCCTCCAC TACGGCACGC TCGTCATCGA
     3481 CGGCGTCACG CCGAACATGA TCGACTACTT CGGCCGCCCG TACGAGGGCA TCGCGGTCTT
     3541 CGACGGCAAG AAGATCACGG TCACGGGCAC GCTCTGGAAC GGCAACAAGA TCATCGACGA
     3601 GCGCCTCATC AACCCGGACG GCTCGCTCCT CTTCCGCGTC ACGATCAACG GCGTCACGGG
     3661 CTGGCGCCTC TGCGAGCGCA TCCTCGCGTG AGCGCTGTGC CGACGCACGT TCCGTCCAGT
     3721 ACGAACACAT AACCAACAAC TCACAATGGT GTTAACAAAT TAAACACCAC CACTTCTTTA
```

```
     3781 AAGAACCAAA CGCAACACGT ATATACGCGA GCGTCGAACC GAGCGAGCGC GCTTGAATCC
     3841 GTAAAAAAAA ACAACGGCTA GAAAATTCAA ATCGAAAAGG CAGACTTTCT AGATGACGAA
     3901 TTTTTGTCGC AATTAAACGG TTCGTgaatt c
//
```

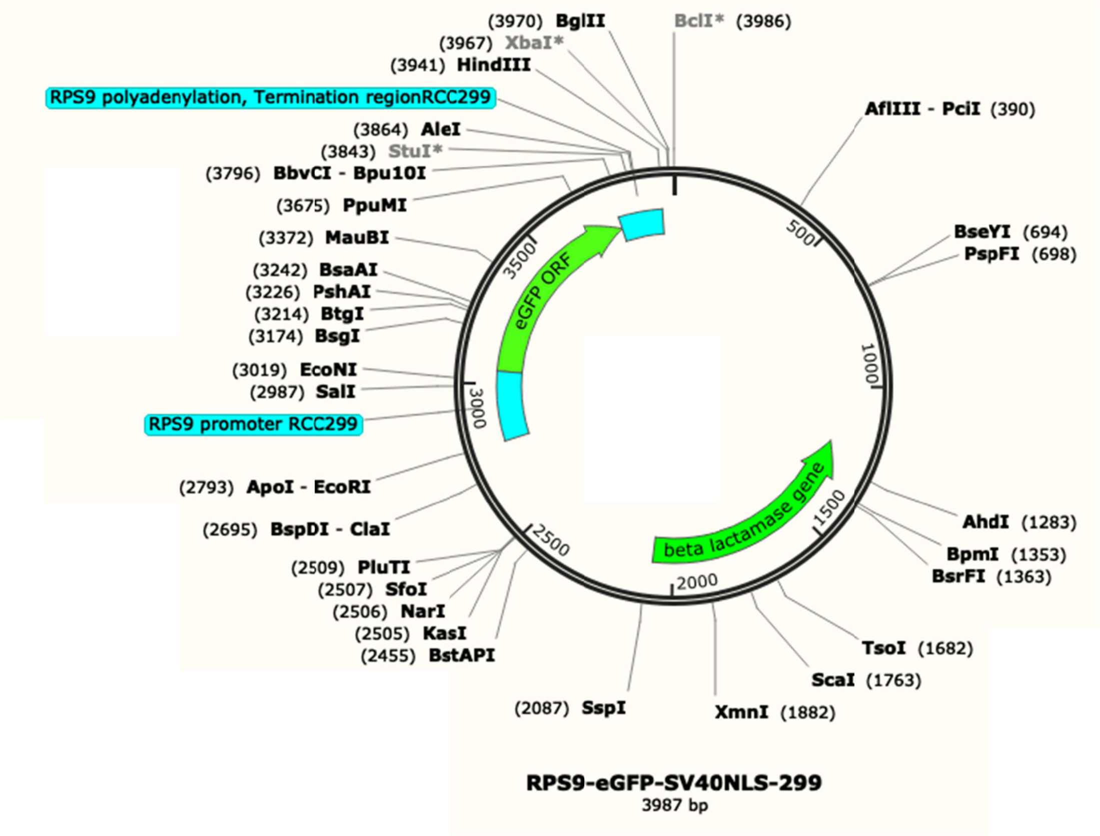


```
LOCUS       RPS9_eGFP_SV40NL        3987 bp ds-DNA     circular     26-OCT-2019
DEFINITION  .
ACCESSION
VERSION
SOURCE      .
COMMENT
COMMENT
COMMENT     ApEinfo:methylated:1
FEATURES             Location/Qualifiers
     promoter        2793..3045
                     /locus_tag="RPS9 promoter RCC299"
                     /label="RPS9 promoter RCC299"
                     /ApEinfo_label="RPS9 promoter RCC299"
                     /ApEinfo_fwdcolor="cyan"
                     /ApEinfo_revcolor="green"
                     /ApEinfo_graphicformat="arrow_data {{0 1 2 0 0 -1} {} 0}
                     width 5 offset 0"
     3'UTR           3787..3946
                     /locus_tag="RPS9 polyadenylation, Termination region
                     RCC299"
                     /label="RPS9 polyadenylation, Termination region RCC299"
                     /ApEinfo_label="RPS9 polyadenylation, Termination region
                     RCC299"
                     /ApEinfo_fwdcolor="cyan"
                     /ApEinfo_revcolor="green"
                     /ApEinfo_graphicformat="arrow_data {{0 1 2 0 0 -1} {} 0}
                     width 5 offset 0"
     CDS             3046..3786
                     /locus_tag="eGFP ORF"
                     /label="eGFP ORF"
                     /ApEinfo_label="eGFP ORF"
                     /ApEinfo_fwdcolor="#5bff37"
                     /ApEinfo_revcolor="green"
```

```
                     /ApEinfo_graphicformat="arrow_data {{0 1 2 0 0 -1} {} 0}
                     width 5 offset 0"
     CDS             complement(1210..2070)
                     /locus_tag="beta lactamase gene"
                     /label="beta lactamase gene"
                     /ApEinfo_label="beta lactamase gene"
                     /ApEinfo_fwdcolor="cyan"
                     /ApEinfo_revcolor="green"
                     /ApEinfo_graphicformat="arrow_data {{0 1 2 0 0 -1} {} 0}
                     width 5 offset 0"
ORIGIN
        1 ATCAGTTCTG GACCAGCGAG CTGTGCTGCG ACTCGTGGCG TAATCATGGT CATAGCTGTT
       61 TCCTGTGTGA AATTGTTATC CGCTCACAAT TCCACACAAC ATACGAGCCG GAAGCATAAA
      121 GTGTAAAGCC TGGGGTGCCT AATGAGTGAG CTAACTCACA TTAATTGCGT TGCGCTCACT
      181 GCCCGCTTTC CAGTCGGGAA ACCTGTCGTG CCAGCTGCAT TAATGAATCG GCCAACGCGC
      241 GGGGAGAGGC GGTTTGCGTA TTGGGCGCTC TTCCGCTTCC TCGCTCACTG ACTCGCTGCG
      301 CTCGGTCGTT CGGCTGCGGC GAGCGGTATC AGCTCACTCA AAGGCGGTAA TACGGTTATC
      361 CACAGAATCA GGGGATAACG CAGGAAAGAA CATGTGAGCA AAAGGCCAGC AAAAGGCCAG
      421 GAACCGTAAA AAGGCCGCGT TGCTGGCGTT TTTCCATAGG CTCCGCCCCC CTGACGAGCA
      481 TCACAAAAAT CGACGCTCAA GTCAGAGGTG GCGAAACCCG ACAGGACTAT AAAGATACCA
      541 GGCGTTTCCC CCTGGAAGCT CCCTCGTGCG CTCTCCTGTT CCGACCCTGT CGCTTACCGG
      601 ATACCTGTCC GCCTTTCTCC CTTCGGGAAG CGTGGCGCTT TCTCATAGCT CACGCTGTAG
      661 GTATCTCAGT TCGGTGTAGG TCGTTCGCTC CAAGCTGGGC TGTGTGCACG AACCCCCCGT
      721 TCAGCCCGAC CGCTGCGCCT TATCCGGTAA CTATCGTCTT GAGTCCAACC CGGTAAGACA
      781 CGACTTATCG CCACTGGCAG CAGCCACTGG TAACAGGATT AGCAGAGCGA GGTATGTAGG
      841 CGGTGCTACA GAGTTCTTGA AGTGGTGGCC TAACTACGGC TACACTAGAA GAACAGTATT
      901 TGGTATCTGC GCTCTGCTGA AGCCAGTTAC CTTCGGAAAA AGAGTTGGTA GCTCTTGATC
      961 CGGCAAACAA ACCACCGCTG GTAGCGGTGG TTTTTTTGTT TGCAAGCAGC AGATTACGCG
     1021 CAGAAAAAAA GGATCTCAAG AAGATCCTTT GATCTTTTCT ACGGGGTCTG ACGCTCAGTG
     1081 GAACGAAAAC TCACGTTAAG GGATTTTGGT CATGAGATTA TCAAAAAGGA TCTTCACCTA
     1141 GATCCTTTTA AATTAAAAAT GAAGTTTTAA ATCAATCTAA AGTATATATG AGTAAACTTG
     1201 GTCTGACAGT TACCAATGCT TAATCAGTGA GGCACCTATC TCAGCGATCT GTCTATTTCG
     1261 TTCATCCATA GTTGCCTGAC TCCCCGTCGT GTAGATAACT ACGATACGGG AGGGCTTACC
     1321 ATCTGGCCCC AGTGCTGCAA TGATACCGCG AGACCCACGC TCACCGGCTC CAGATTTATC
     1381 AGCAATAAAC CAGCCAGCCG GAAGGGCCGA GCGCAGAAGT GGTCCTGCAA CTTTATCCGC
     1441 CTCCATCCAG TCTATTAATT GTTGCCGGGA AGCTAGAGTA AGTAGTTCGC CAGTTAATAG
     1501 TTTGCGCAAC GTTGTTGCCA TTGCTACAGG CATCGTGGTG TCACGCTCGT CGTTTGGTAT
     1561 GGCTTCATTC AGCTCCGGTT CCCAACGATC AAGGCGAGTT ACATGATCCC CCATGTTGTG
     1621 CAAAAAAGCG GTTAGCTCCT TCGGTCCTCC GATCGTTGTC AGAAGTAAGT TGGCCGCAGT
     1681 GTTATCACTC ATGGTTATGG CAGCACTGCA TAATTCTCTT ACTGTCATGC CATCCGTAAG
     1741 ATGCTTTTCT GTGACTGGTG AGTACTCAAC CAAGTCATTC TGAGAATAGT GTATGCGGCG
     1801 ACCGAGTTGC TCTTGCCCGG CGTCAATACG GGATAATACC GCGCCACATA GCAGAACTTT
     1861 AAAAGTGCTC ATCATTGGAA AACGTTCTTC GGGGCGAAAA CTCTCAAGGA TCTTACCGCT
     1921 GTTGAGATCC AGTTCGATGT AACCCACTCG TGCACCCAAC TGATCTTCAG CATCTTTTAC
     1981 TTTCACCAGC GTTTCTGGGT GAGCAAAAAC AGGAAGGCAA AATGCCGCAA AAAAGGGAAT
     2041 AAGGGCGACA CGGAAATGTT GAATACTCAT ACTCTACCTT TTTCAATATT ATTGAAGCAT
     2101 TTATCAGGGT TATTGTCTCA TGAGCGGATA CATATTTGAA TGTATTTAGA AAAATAAACA
     2161 AATAGGGGTT CCGCGCACAT TTCCCCGAAA AGTGCCACCT GACGTCTAAG AAACCATTAT
     2221 TATCATGACA TTAACCTATA AAAATAGGCG TATCACGAGG CCCTTTCGTC TCGCGCGTTT
     2281 CGGTGATGAC GGTGAAAACC TCTGACACAT GCAGCTCCCG GAGACGGTCA CAGCTTGTCT
     2341 GTAAGCGGAT GCCGGGAGCA GACAAGCCCG TCAGGGCGCG TCAGCGGGTG TTGGCGGGTG
     2401 TCGGGGCTGG CTTAACTATG CGGCATCAGA GCAGATTGTA CTGAGAGTGC ACCAAATGCG
     2461 GTGTGAAATA CCGCACAGAT GCGTAAGGAG AAAATACCGC ATCAGGCGCC ATTCGCCATT
     2521 CAGGCTGCGC AACTGTTGGG AAGGGCGATC GGTGCGGGCC TCATCGCTAT TACGCCAGCT
     2581 GGCGAAAGGG GGATGTGCTG CAAGGCGATT AAGTTGGGTA ACGCCAGGGT TTTCCCAGTC
     2641 ACGACGTTGT AAAACGACGG CCAGTGCAAC GCGATGACGA TGGATAGCGA TTCATCGATG
     2701 AGCTGACCCG ATCGCCGCCG CCGGAGGGTT GCGTTTGAGA CGGGCGACAG ATGAGCACAT
     2761 CTCGTTCGCT ATTCAGGGAT TGTTAATTAA CAgaattcTC GTGGATTTTT GACAAAAAGC
     2821 GATCTTTACC AGTGAGATCG ACCCATTTGC AGTTGACCAT CCACTTCGTT TTCCCATTAT
     2881 TTCCTATTTC GAGGTTCGCA CAGTATGTGT TCATGACAGT TAGTGAGCTT CGTTCAAAAC
     2941 AGGTTCTGGT CGGATTTTCC TGGCCGGACT TTCCGCTAAA ACCCTCGTCG ACCCAGTCGT
     3001 CCCGCCCTCT TACACCTAGC GCAGGAAGGC GAAGGGACAG CCAAGATGGT CTCGAAGGGC
     3061 GAGGAGCTCT TCACGGGCGT CGTCCCGATC CTCGTCGAGC TCGACGGCGA CGTCAACGGC
     3121 CACAAGTTCT CGGTCTCGGG CGAGGGCGAG GGCGACGCGA CGTACGGCAA GCTCACGCTC
     3181 AAGTTCATCT GCACGACGGG CAAGCTCCCG GTCCCGTGGC CGACGCTCGT CACGACGCTC
     3241 ACGTACGGCG TCCAGTGCTT CTCGCGCTAC CCGGACCACA TGAAGCAGCA CGACTTCTTC
     3301 AAGTCGGCGA TGCCGGAGGG CTACGTCCAG GAGCGCACGA TCTTCTTCAA GGACGACGGC
     3361 AACTACAAGA CGCGCGCGGA GGTCAAGTTC GAGGGCGACA CGCTCGTCAA CCGCATCGAG
     3421 CTCAAGGGCA TCGACTTCAA GGAGGACGGC AACATCCTCG GCCACAAGCT CGAGTACAAC
     3481 TACAACTCGC ACAACGTCTA CATCATGGCG GACAAGCAGA AGAACGGCAT CAAGGTCAAC
     3541 TTCAAGATCC GCCACAACAT CGAGGACGGC TCGGTCCAGC TCGCGGACCA CTACCAGCAG
     3601 AACACGCCGA TCGGCGACGG CCCGGTCCTC CTCCCGGACA ACCACTACCT CTCGACGCAG
     3661 TCGGCGCTCT CGAAGGACCC GAACGAGAAG CGCGACCACA TGGTCCTCCT CGAGTTCGTC
     3721 ACGGCGGCGG GCATCACGCT CGGCATGGAC GAGCTCTACA AGCCGAAGAA GAAGCGCAAG
     3781 GTCTGAACAC CCGAGCTGAG GGCCGGGGGA TCTTCCACGC GGCGTCTAAG TGACGACGCC
     3841 AGGCCTTGCG CGCCGATCTC ACGAGAGTGG CGTGCAATTT ACTAACAGTA GTGACTTGAA
     3901 ATACACGGGT AACTCCTGTC CACAAACGAA CCCGCGCGAG aagcttGCGT TAATTAAGGA
```

```
     3961 AGTTTGTCTA GATCTCAGGC GTGGATG
```

#### *Tetraselmis striata*

#### GenBank: KY886895

```
LOCUS       KY886895                5787 bp    DNA     circular
DEFINITION  Synthetic plasmid pGEM-Rps12:bar:Gapdh, complete sequence.
ACCESSION   KY886895
VERSION     KY886895.1  GI:1229733783
KEYWORDS    .
SOURCE      Synthetic plasmid pGEM-Rps12:bar:Gapdh
  ORGANISM  Synthetic plasmid pGEM-Rps12:bar:Gapdh
            other sequences; artificial sequences; vectors.
REFERENCE   1  (bases 1 to 5787)
  AUTHORS   Luan,F., Wen,X., Beardslee,P. and Cerutti,H.
  TITLE     Tetraselmis transformation via microprojectile bombardment
  JOURNAL   Unpublished
REFERENCE   2  (bases 1 to 5787)
  AUTHORS   Luan,F., Wen,X., Beardslee,P. and Cerutti,H.
  TITLE     Direct Submission
  JOURNAL   Submitted (05-APR-2017) School of Biological Sciences, University
            of Nebraska-Lincoln, 1901 Vine St, Lincoln, NE 68588, USA
COMMENT     ##Assembly-Data-START##
            Sequencing Technology :: Sanger dideoxy sequencing
            ##Assembly-Data-END##
FEATURES             Location/Qualifiers
     source          1..5787
                     /organism="Synthetic plasmid pGEM-Rps12:bar:Gapdh"
                     /mol_type="other DNA"
                     /db_xref="taxon:2014915"
     misc_feature    join(1..2926,5731..5787)
                     /note="pGEM-T Easy vector backbone"
     regulatory      2927..3943
                     /regulatory_class="promoter"
                     /note="Rps12-Pro; Ribosomal protein S12 promoter including
                     5' UTR; derived from Tetraselmis striata"
     CDS             3950..4501
                     /note="phosphinothricin acetyltransferase; herbicide
                     resistance protein"
                     /codon_start=1
                     /transl_table=11
                     /product="Bar"
                     /protein_id="AST24419.1"
                     /db_xref="GI:1229733784"
                     /translation="MSPERRPADIRRATEADMPAVCTIVNHYIETSTVNFRTEPQEPQ
                     EWTDDLVRLRERYPWLVAEVDGEVAGIAYAGPWKARNAYDWTAESTVYVSPRHQRTGL
                     GSTLYTHLLKSLEAQGFKSVVAVIGLPNDPSVRMHEALGYAPRGMLRAAGFKHGNWHD
                     VGFWQLDFSLPVPPRPVLPVTEI"
     regulatory      4508..5730
                     /regulatory_class="terminator"
                     /note="Gapdh-Ter; glyceraldehyde 3-phosphate dehydrogenase
                     terminator; derived from Tetraselmis striata"
ORIGIN
        1 tatagtgagt cgtattacaa ttcactggcc gtcgttttac aacgtcgtga ctgggaaaac
       61 cctggcgtta cccaacttaa tcgccttgca gcacatcccc ctttcgccag ctggcgtaat
      121 agcgaagagg cccgcaccga tcgcccttcc caacagttgc gcagcctgaa tggcgaatgg
      181 acgcgccctg tagcggcgca ttaagcgcgg cgggtgtggt ggttacgcgc agcgtgaccg
      241 ctacacttgc cagcgcccta gcgcccgctc ctttcgcttt cttcccttcc tttctcgcca
      301 cgttcgccgg ctttccccgt caagctctaa atcgggggct ccctttaggg ttccgattta
      361 gtgctttacg gcacctcgac cccaaaaaac ttgattaggg tgatggttca cgtagtgggc
      421 catcgccctg atagacggtt tttcgccctt tgacgttgga gtccacgttc tttaatagtg
      481 gactcttgtt ccaaactgga acaacactca accctatctc ggtctattct tttgatttat
      541 aagggatttt gccgatttcg gcctattggt taaaaaatga gctgatttaa caaaaattta
      601 acgcgaattt taacaaaata ttaacgctta caatttcctg atgcggtatt ttctccttac
      661 gcatctgtgc ggtatttcac accgcatcag gtggcacttt tcggggaaat gtgcgcggaa
      721 cccctatttg tttatttttc taaatacatt caaatatgta tccgctcatg agacaataac
      781 cctgataaat gcttcaataa tattgaaaaa ggaagagtat gagtattcaa catttccgtg
      841 tcgcccttat tccctttttt gcggcatttt gccttcctgt ttttgctcac ccagaaacgc
      901 tggtgaaagt aaaagatgct gaagatcagt tgggtgcacg agtgggttac atcgaactgg
      961 atctcaacag cggtaagatc cttgagagtt ttcgccccga agaacgtttt ccaatgatga
     1021 gcacttttaa agttctgcta tgtggcgcgg tattatcccg tattgacgcc gggcaagagc
     1081 aactcggtcg ccgcatacac tattctcaga atgacttggt tgagtactca ccagtcacag
     1141 aaaagcatct tacggatggc atgacagtaa gagaattatg cagtgctgcc ataaccatga
```

1201 gtgataacac tgcggccaac ttacttctga caacgatcgg aggaccgaag gagctaaccg
1261 ctttttttgca caacatgggg gatcatgtaa ctcgccttga tcgttgggaa ccggagctga
1321 atgaagccat accaaacgac gagcgtgaca ccacgatgcc tgtagcaatg gcaacaacgt
1381 tgcgcaaact attaactggc gaactactta ctctagcttc ccggcaacaa ttaatagact
1441 ggatggaggc ggataaagtt gcaggaccac ttctgcgctc ggcccttccg gctggctggt
1501 ttattgctga taaatctgga gccggtgagc gtgggtctcg cggtatcatt gcagcactgg
1561 ggccagatgg taagccctcc cgtatcgtag ttatctacac gacggggagt caggcaacta
1621 tggatgaacg aaatagacag atcgctgaga taggtgcctc actgattaag cattggtaac
1681 tgtcagacca agtttactca tatatacttt agattgattt aaaacttcat ttttaattta
1741 aaaggatcta ggtgaagatc ctttttgata atctcatgac caaaatccct taacgtgagt
1801 tttcgttcca ctgagcgtca gaccccgtag aaaagatcaa aggatcttct tgagatcctt
1861 tttttctgcg cgtaatctgc tgcttgcaaa caaaaaaacc accgctacca gcggtggttt
1921 gtttgccgga tcaagagcta ccaactcttt ttccgaaggt aactggcttc agcagagcgc
1981 agataccaaa tactgttctt ctagtgtagc cgtagttagg ccaccacttc aagaactctg
2041 tagcaccgcc tacatacctc gctctgctaa tcctgttacc agtggctgct gccagtggcg
2101 ataagtcgtg tcttaccggg ttggactcaa gacgatagtt accggataag gcgcagcggt
2161 cgggctgaac ggggggttcg tgcacacagc ccagcttgga gcgaacgacc tacaccgaac
2221 tgagatacct acagcgtgag ctatgagaaa gcgccacgct tcccgaaggg agaaaggcgg
2281 acaggtatcc ggtaagcggc agggtcggaa caggagagcg cacgagggag cttccagggg
2341 gaaacgcctg gtatctttat agtcctgtcg ggtttcgcca cctctgactt gagcgtcgat
2401 ttttgtgatg ctcgtcaggg gggcggagcc tatggaaaaa cgccagcaac gcggcctttt
2461 tacggttcct ggccttttgc tggccttttg ctcacatgtt ctttcctgcg ttatcccctg
2521 attctgtgga taaccgtatt accgcctttg agtgagctga taccgctcgc cgcagccgaa
2581 cgaccgagcg cagcgagtca gtgagcgagg aagcggaaga gcgcccaata cgcaaaccgc
2641 ctctccccgc gcgttggccg attcattaat gcagctggca cgacaggttt cccgactgga
2701 aagcgggcag tgagcgcaac gcaattaatg tgagttagct cactcattag gcaccccagg
2761 ctttacactt tatgcttccg gctcgtatgt tgtgtggaat tgtgagcgga taacaatttc
2821 acacaggaaa cagctatgac catgattacg ccaagctatt taggtgacac tatagaatac
2881 tcaagctatg catccaacgc gttgggagct ctcccatatg gtcgactgct gctaactggt
2941 tcccggggct gaattcggaa tgtatgcgtg gttattgcgt ttcctgtgca tatacttccg
3001 ggatactttc gccaaataca tcagcagtag tttatgggca ggcggcggcc gccgcccgcc
3061 gacgacatcg tagtcgctcg tcgtcaggga agtgccgtca ggagcgtggt ggcggcggcg
3121 cggcggctcg cgccccggcg gccggtgcgg aaaggtgagc ggaggcgccc gtgacgtccc
3181 cctttcgccc accatcctct acgctcaacc cacaggtgat tcagcaagcc ctccggcccc
3241 gccctccagg caaccagtca cgctagtcca ccccagcgta tcgccggccc ttcccgcgct
3301 ggctttagcg gcaggctcgc cgcctgatcc ggcgcgccct cgccgccgcc accgagaatc
3361 ttcgcagcag ccgcaccgcc gggaagtgta ctgggccagg ggcagcaccg tgtcgcagca
3421 aatgtgcgag gcgcgcagga aacgcaggca tgccgctgga atccacgcgg acaatggagg
3481 ggaaaggggg ggattcgttt ggtggccccc agcgtagatg gggcagggtg accaggtctc
3541 ctttattcct tcaactgccc agattcgtcc gaggttgggc attccttgca ttacgggaaa
3601 taccgcgcgg ctctgtaatg ccggtcgcga tttacgcctt gaacttcaaa ttattgtttt
3661 tcggggaaac ccaccaggct gggttcgcta gggaagcgtc caagcctgac tgagtagctg
3721 cctgcctggt caccacttga ggtccaccac accgcagtca cttgccctgc cgccatggcg
3781 attctgagtg ctgccatctc tgcctctgca gctgccaggc aggctcgcca tctgcctgct
3841 gtacgccagc acatctgacc tccccccgcc cccctgtctt cctttctaat aggaacatcc
3901 tctaggacga aaagactttg agacagctct acctgtcgag gaagagctca tgagcccaga
3961 acgacgcccg gccgacatcc gccgtgccac cgaggcggac atgccggcgg tctgcaccat
4021 cgtcaaccac tacatcgaga caagcacggt caacttccgt accgagccgc aggaaccgca
4081 ggagtggacg gacgacctcg tccgtctgcg ggagcgctat ccctggctcg tcgccgaggt
4141 ggacggcgag gtcgccggca tcgcctacgc gggcccctgg aaggcacgca acgcctacga
4201 ctggacggcc gagtcgaccg ttacgtctc cccccgccac cagcggacgg gactgggctc
4261 cacgctctac acccacctgc tgaagtccct ggaggcacag ggcttcaaga gcgtggtcgc
4321 tgtcatcggg ctgcccaacg acccgagcgt gcgcatgcac gaggcgctcg gatatgcccc
4381 ccgcggcatg ctgcgggcgg ccggcttcaa gcacgggaac tggcatgacg tgggtttctg
4441 gcagctggac ttcagcctgc cggtaccgcc ccgtccggtc ctgcccgtca ccgagatctg
4501 aggatcctga ggcctatcct cgctgctcat accaccattc atccacaaat agcagtagcc
4561 agcacctgtt gccctctgtg cgcaagtgtg tcagatagga ctgcaaatat gaaaaccagc
4621 caaattcctg tactggaatt ggttactctc cactttttttt taggtagaac tgtaaccata
4681 tcttgcaaaa ataccaacaa tgcaaggaag gagacattgt gcaggaatac atgctaccta
4741 attctggggg cgacgttctg aggttttcag ttgcggacag caatgcagct tcaagacaac
4801 aatggcatgc cttgtccctt tccgcacatg gttatggcat gaatagcaag atgttcacta
4861 gcgggctgct gccattcact gattgttgat gaatgcgcac atttctctgg gtagtggagg
4921 gtaattttgg cgttttgggt attatccgaa atgcctagtt tccaacatcg tggatatgct
4981 cttcagtctc attgtagcat gctgcccgcc tgttttgaaa tgcagtgggt taaattggtg
5041 cagaagaggg ggcagattct gtcacatgca actaaacatc aagggacaat gacaaaattg
5101 tttcccacta ggacgataaa tccgcactat ataatgagac agtgagtgct tttccaccaa
5161 cctccatgtt gatagaaatg ccaacaccag atcgatgggc aacattatcg gcactcggcc
5221 atgtaggcac tcatcgcaaa tcccggcact ggtggtgcgt gttgggctgc cactgtaagc
5281 gcatgcccgc tgcagagact ccactgatga aggcgctcca gcgaagatat aagcgaaaac
5341 tgtgacacag caacccacca gttgacttca ctggcaagaa gtaccttctt tgttagcatc
5401 cagtagactg acctcaatat gcactcgcat taattcgact ataacgaatc atggaactac
5461 cgaagcatgc aatacgtaac atatccgctg tgtaagtata aatgagcggt gcgcatgtgc
5521 tatgatgttg cgcagcatag tttatgagtt gttattaagg aggttgctac tgggcttgcc
5581 aaatgattca gttctgtgag ccagatcctt gagatttggt gtcgcgacga gcacatgtaa
5641 acctttaaaa ccgttatatc acttaattaa tgacacagca tatgaaacct agcttaagtt
5701 ggtaccgtcg tggacaagta caagatcgcc gaattcccgc ggccgccatg gcggccggga
5761 gcatgcgacg tcgggcccaa ttcgccc

*Pyramimonas parkeae*

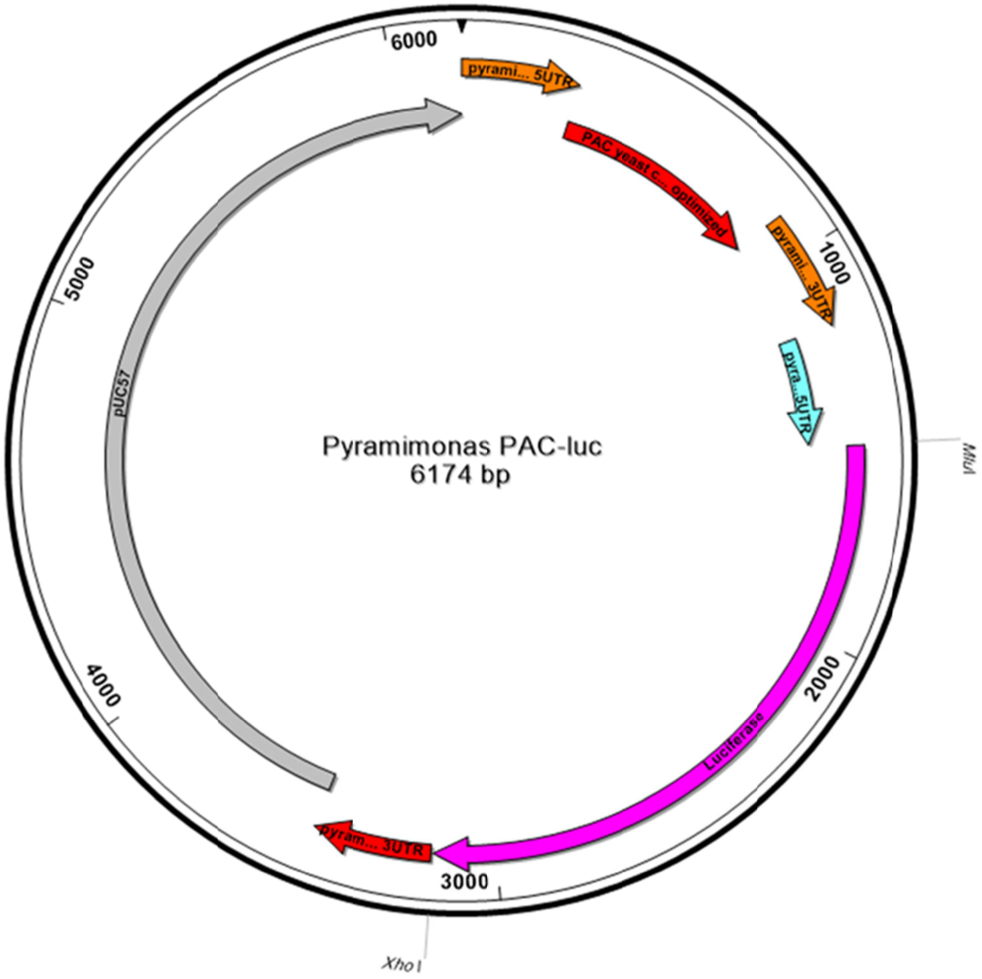


>Pyramimonas PAC-luc

CGTGCGGTGCGAGCTTGTTTGCACCTGCGAGCGTGACCCAAGCTTTATGAATCCCCCGTCGACAATCACGCATCCAATGAGACGCT
ACCATATTGTCAGAGCGGATCGTTGCTGCAACCAATGAATGGTGCTGTAGATTCGTATCGCGGATAGAGCAAGCACGATCCTCTCT
GCAGCCAATCGTGGATACTACACGTGTACCAAATTGTGAATCGTATGCTCGAAGATGGGAAGAGAAAATCACAACGCGCGCTTCAC
CCTCGCACAGCAGCAAGCTATCCACCACCGACCCAAGCGAAATGACCGAACATAAGCCATCTGTTAGATTGGCTACTAGAGATGAT
GTTCCAAGAGCTGTTAGAACTTTGGCTGCTGCTTTTGCTGATTATCCAGCTACTAGACATACTGTTGATCCAGATAGACACATCGA
AAGAGTTACCGAATTGCAAGAATTATTCTTAACCAGAGTCGGTTTGGATATCGGTAAAGTTTGGGTTGCTGATGATGGTGCTGCTG
TTGCAGTTTGGACTACTCCAGAATCTTTGGATGCTGGTGCTGTTTTCGCTGAAATTGGTCCAAGAATGGCTGAATTGTCTGGTTCT
AGATTAGCTGCTCAACAACAAATGGAAGGTTTGTTGGCTCCACATAGACCAAAAGAACCAGCTTGGTTTTTGGCTACTGTTGGTGT
TTCTCCAGATCATCAAGGTAAAGGTTTAGGTTCTGCTGTTGTTTTGCCAGGTGTTGAAGCTGCTGAAAGAGCTGGTGTTCCAGCTT
TTTTGGAAACTTCTGCTCCAAGAAATTTGCCATTCTACGAAAGATTGGGTTTCACTGTTACTGCCGATGTTGAAGTTCCTGAAGGT
CCTAGAACTTGGTGTATGACTAGAAAACCAGGTGCTTAAGCTTGTTGCTTATAATAAGAACCAACCGGTGTTTCACAACACCACCT
TCAGAGGAAGAAGTTGACTCGGGACACGTTATCGGAGGAATGTGGACATAGTTGGAATCACAGGGTGTGTACGCGCGCAGGAGTTT
TCGGTCCATGAGGAAATTGCTTCCAATATCTAAGCAGCCAAAACGTGTGCATCAAAGAAGTCATTGTACGAAACTATTGTACATCT
GATGGGGCTAAGCGCGGCTCATCACAACATGACCTTTCAATGAAGTCGATTCCCTTATATGTTACCTGCAGGATGCACTCACAGGC
TGTCAAAGTAAAGTGTTGACCAAGGTTCGTGGTTTCGAATTCGTGCCCCCGAGATCTGTTGCGTGTTGACTTTGCCAAACGAGTCC
GCATGCAGTCTGGCCAATGGCGACCCGAGGATGCGGGCGAGGTACGGTGTGGCCAATGGGCGATGCCGCTGAGGTGAGTTCCAGAT
CGATGGTCGGATTCCCATCCAACATGCGCAGCATTTCCATGTGTTCATATACTGCGGACTCGCCTTGCCTGTCCAACAAACGCACG

AGCAGCGCACCTCCATCCCACAAACAAGTAATCAAAAACGCGTATGGAAGACGCCAAAAACATAAAGAAAGGCCCGGCGCCATTCT
ATCCGCTGGAAGATGGAACCGCTGGAGAGCAACTGCATAAGGCTATGAAGAGATACGCCCTGGTTCCTGGAACAATTGCTTTTACA
GATGCACATATCGAGGTGGACATCACTTACGCTGAGTACTTCGAAATGTCCGTTCGGTTGGCAGAAGCTATGAAACGATATGGGCT
GAATACAAATCACAGAATCGTCGTATGCAGTGAAAACTCTCTTCAATTCTTTATGCCGGTGTTGGGCGCGTTATTTATCGGAGTTG
CAGTTGCGCCCGCGAACGACATTTATAATGAACGTGAATTGCTCAACAGTATGGGCATTTCGCAGCCTACCGTGGTGTTCGTTTCC
AAAAAGGGGTTGCAAAAAATTTTGAACGTGCAAAAAAAGCTCCCAATCATCCAAAAAATTATTATCATGGATTCTAAAACGGATTA
CCAGGGATTTCAGTCGATGTACACGTTCGTCACATCTCATCTACCTCCCGGTTTTAATGAATACGATTTTGTGCCAGAGTCCTTCG
ATAGGGACAAGACAATTGCACTGATCATGAACTCCTCTGGATCTACTGGTCTGCCTAAAGGTGTCGCTCTGCCTCATAGAACTGCC
TGCGTGAGATTCTCGCATGCCAGAGATCCTATTTTTGGCAATCAAATCATTCCGGATACTGCGATTTTAAGTGTTGTTCCATTCCA
TCACGGTTTTGGAATGTTTACTACACTCGGATATTTGATATGTGGATTTCGAGTCGTCTTAATGTATAGATTTGAAGAAGAGCTGT
TTCTGAGGAGCCTTCAGGATTACAAGATTCAAAGTGCGCTGCTGGTGCCAACCCTATTCTCCTTCTTCGCCAAAAGCACTCTGATT
GACAAATACGATTTATCTAATTTACACGAAATTGCTTCTGGTGGCGCTCCCCTCTCTAAGGAAGTCGGGGAAGCGGTTGCCAAGAG
GTTCCATCTGCCAGGTATCAGGCAAGGATATGGGCTCACTGAGACTACATCAGCTATTCTGATTACACCCGAGGGGGATGATAAAC
CGGGCGCGGTCGGTAAAGTTGTTCCATTTTTTGAAGCGAAGGTTGTGGATCTGGATACCGGGAAAACGCTGGGCGTTAATCAAAGA
GGCGAACTGTGTGTGAGAGGTCCTATGATTATGTCCGGTTATGTAAACAATCCGGAAGCGACCAACGCCTTGATTGACAAGGATGG
ATGGCTACATTCTGGAGACATAGCTTACTGGGACGAAGACGAACACTTCTTCATCGTTGACCGCCTGAAGTCTCTGATTAAGTACA
AAGGCTATCAGGTGGCTCCCGCTGAATTGGAATCCATCTTGCTCCAACACCCCAACATCTTCGACGCAGGTGTCGCAGGTCTTCCC
GACGATGACGCCGGTGAACTTCCCGCCGCCGTTGTTGTTTTGGAGCACGGAAAGACGATGACGGAAAAAGAGATCGTGGATTACGT
CGCCAGTCAAGTAACAACCGCGAAAAAGTTGCGCGGAGGAGTTGTGTTTGTGGACGAAGTACCGAAAGGTCTTACCGGAAAACTCG
ACGCAAGAAAAATCAGAGAGATCCTCATAAAGGCCAAGAAGGGCGGAAAGATCGCCGTGTAACTCGAGGCCACAAGGTGACCGGGA
ATCCAACGAAAAACAAAACGGTGTTTCCAAACACCACCTGTATGGACATGGATCAACTTAAGGTCGTTGAGTTTGGATGAAGTCTA
GGCTGACCAGGATATCGCCCAAGTTGGAACATAAGTCAAGGTTGCATATGTTATGTACAATTAGAATCCTTGTTTGTGTGCGAAGA
CGGATTGCAAGGGCAGCAGCATCGTAATATATCCACAATGAACATCCAAAATTCCAGACTTCTCTCACGTAAGCACGTTGCTGATT
TGCCAGTAAATGGAGTTTCGTATCATCGGATCCCGGGCCCGTCGACTGCAGAGGCCTGCATGCAAGCTTGGCGTAATCATGGTCAT
AGCTGTTTCCTGTGTGAAATTGTTATCCGCTCACAATTCCACACAACATACGAGCCGGAAGCATAAAGTGTAAAGCCTGGGGTGCC
TAATGAGTGAGCTAACTCACATTAATTGCGTTGCGCTCACTGCCCGCTTTCCAGTCGGGAAACCTGTCGTGCCAGCTGCATTAATG
AATCGGCCAACGCGCGGGGAGAGGCGGTTTGCGTATTGGGCGCTCTTCCGCTTCCTCGCTCACTGACTCGCTGCGCTCGGTCGTTC
GGCTGCGGCGAGCGGTATCAGCTCACTCAAAGGCGGTAATACGGTTATCCACAGAATCAGGGGATAACGCAGGAAAGAACATGTGA
GCAAAAGGCCAGCAAAAGGCCAGGAACCGTAAAAAGGCCGCGTTGCTGGCGTTTTTCCATAGGCTCCGCCCCCCTGACGAGCATCA
CAAAAATCGACGCTCAAGTCAGAGGTGGCGAAACCCGACAGGACTATAAAGATACCAGGCGTTTCCCCCTGGAAGCTCCCTCGTGC
GCTCTCCTGTTCCGACCCTGCCGCTTACCGGATACCTGTCCGCCTTTCTCCCTTCGGGAAGCGTGGCGCTTTCTCATAGCTCACGC
TGTAGGTATCTCAGTTCGGTGTAGGTCGTTCGCTCCAAGCTGGGCTGTGTGCACGAACCCCCCGTTCAGCCCGACCGCTGCGCCTT
ATCCGGTAACTATCGTCTTGAGTCCAACCCGGTAAGACACGACTTATCGCCACTGGCAGCAGCCACTGGTAACAGGATTAGCAGAG
CGAGGTATGTAGGCGGTGCTACAGAGTTCTTGAAGTGGTGGCCTAACTACGGCTACACTAGAAGAACAGTATTTGGTATCTGCGCT
CTGCTGAAGCCAGTTACCTTCGGAAAAAGAGTTGGTAGCTCTTGATCCGGCAAACAAACCACCGCTGGTAGCGGTGGTTTTTTTGT
TTGCAAGCAGCAGATTACGCGCAGAAAAAAAGGATCTCAAGAAGATCCTTTGATCTTTTCTACGGGGTCTGACGCTCAGTGGAACG
AAAACTCACGTTAAGGGATTTTGGTCATGAGATTATCAAAAAGGATCTTCACCTAGATCCTTTTAAATTAAAAATGAAGTTTTAAA
TCAATCTAAAGTATATATGAGTAAACTTGGTCTGACAGTTACCAATGCTTAATCAGTGAGGCACCTATCTCAGCGATCTGTCTATT
TCGTTCATCCATAGTTGCCTGACTCCCCGTCGTGTAGATAACTACGATACGGGAGGGCTTACCATCTGGCCCCAGTGCTGCAATGA
TACCGCGAGACCCACGCTCACCGGCTCCAGATTTATCAGCAATAAACCAGCCAGCCGGAAGGGCCGAGCGCAGAAGTGGTCCTGCA
ACTTTATCCGCCTCCATCCAGTCTATTAATTGTTGCCGGGAAGCTAGAGTAAGTAGTTCGCCAGTTAATAGTTTGCGCAACGTTGT
TGCCATTGCTACAGGCATCGTGGTGTCACGCTCGTCGTTTGGTATGGCTTCATTCAGCTCCGGTTCCCAACGATCAAGGCGAGTTA
CATGATCCCCCATGTTGTGCAAAAAAGCGGTTAGCTCCTTCGGTCCTCCGATCGTTGTCAGAAGTAAGTTGGCCGCAGTGTTATCA
CTCATGGTTATGGCAGCACTGCATAATTCTCTTACTGTCATGCCATCCGTAAGATGCTTTTCTGTGACTGGTGAGTACTCAACCAA
GTCATTCTGAGAATAGTGTATGCGGCGACCGAGTTGCTCTTGCCCGGCGTCAATACGGGATAATACCGCGCCACATAGCAGAACTT
TAAAAGTGCTCATCATTGGAAAACGTTCTTCGGGGCGAAAACTCTCAAGGATCTTACCGCTGTTGAGATCCAGTTCGATGTAACCC
ACTCGTGCACCCAACTGATCTTCAGCATCTTTTACTTTCACCAGCGTTTCTGGGTGAGCAAAAACAGGAAGGCAAAATGCCGCAAA
AAAGGGAATAAGGGCGACACGGAAATGTTGAATACTCATACTCTTCCTTTTTCAATATTATTGAAGCATTTATCAGGGTTATTGTC
TCATGAGCGGATACATATTTGAATGTATTTAGAAAAATAAACAAATAGGGGTTCCGCGCACATTTCCCCGAAAAGTGCCACCTGAC
GTCTAAGAAACCATTATTATCATGACATTAACCTATAAAAATAGGCGTATCACGAGGCCCTTTCGTCTCGCGCGTTTCGGTGATGA
CGGTGAAAACCTCTGACACATGCAGCTCCCGGAGACGGTCACAGCTTGTCTGTAAGCGGATGCCGGGAGCAGACAAGCCCGTCAGG
GCGCGTCAGCGGGTGTTGGCGGGTGTCGGGGCTGGCTTAACTATGCGGCATCAGAGCAGATTGTACTGAGAGTGCACCATATGCGG
TGTGAAATACCGCACAGATGCGTAAGGAGAAAATACCGCATCAGGCGCCATTCGCCATTCAGGCTGCGCAACTGTTGGGAAGGGCG
ATCGGTGCGGGCCTCTTCGCTATTACGCCAGCTGGCGAAAGGGGGATGTGCTGCAAGGCGATTAAGTTGGGTAACGCCAGGGTTTT
CCCAGTCACGACGTTGTAAAACGACGGCCAGTGAATTCGAGCTCGGTACCTCGCGAATGCATCTAGAT

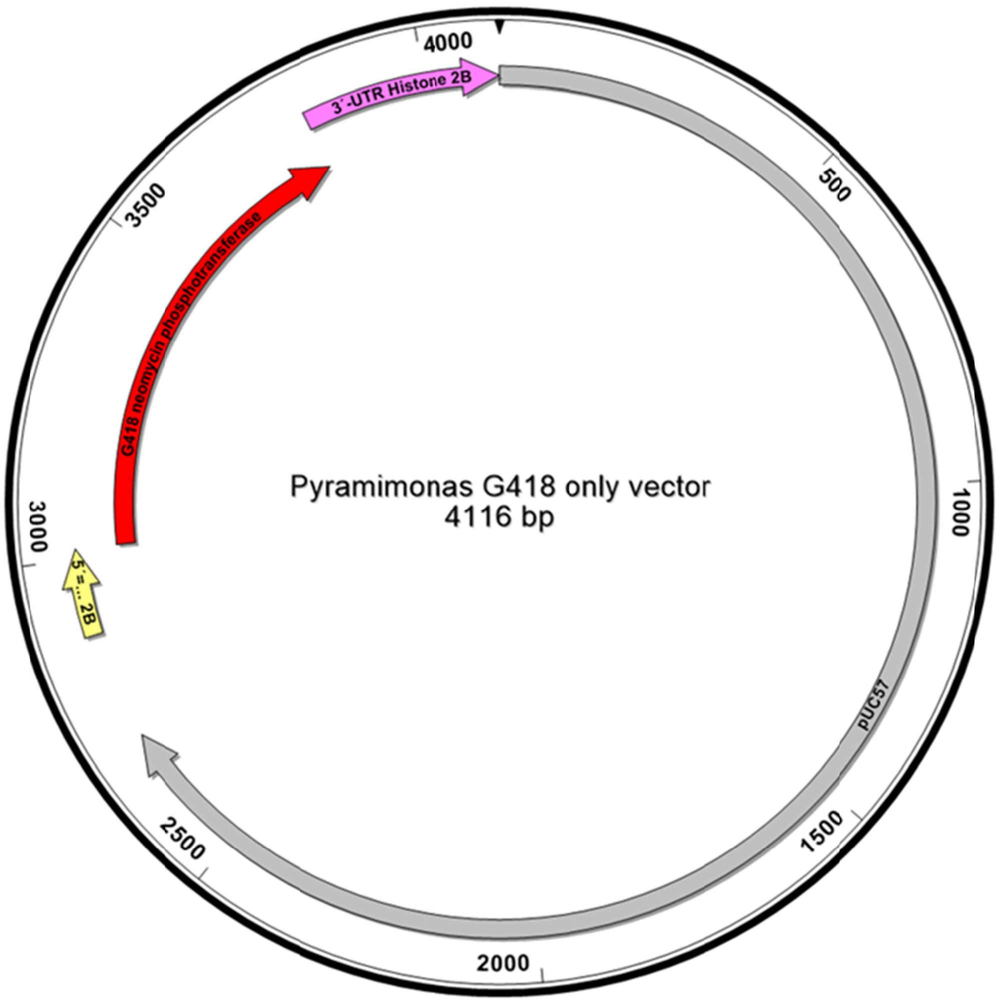


>Luc only vector

1-2710 pUC57 backbone
2711 -3015 5´-UTR Histone 2B
3016-3810 - G418 neomycin phosphotransferase
3811- 4116 3´-UTR Histone 2B

```
ATCGGATCCCGGGCCCGTCGACTGCAGAGGCCTGCATGCAAGCTTGGCGTAATCATGGTCATAGCTGTTTCCTGTGTGAAATTGTT
ATCCGCTCACAATTCCACACAACATACGAGCCGGAAGCATAAAGTGTAAAGCCTGGGGTGCCTAATGAGTGAGCTAACTCACATTA
ATTGCGTTGCGCTCACTGCCCGCTTTCCAGTCGGGAAACCTGTCGTGCCAGCTGCATTAATGAATCGGCCAACGCGCGGGGAGAGG
CGGTTTGCGTATTGGGCGCTCTTCCGCTTCCTCGCTCACTGACTCGCTGCGCTCGGTCGTTCGGCTGCGGCGAGCGGTATCAGCTC
ACTCAAAGGCGGTAATACGGTTATCCACAGAATCAGGGGATAACGCAGGAAAGAACATGTGAGCAAAAGGCCAGCAAAAGGCCAGG
AACCGTAAAAAGGCCGCGTTGCTGGCGTTTTTCCATAGGCTCCGCCCCCCTGACGAGCATCACAAAAATCGACGCTCAAGTCAGAG
GTGGCGAAACCCGACAGGACTATAAAGATACCAGGCGTTTCCCCCTGGAAGCTCCCTCGTGCGCTCTCCTGTTCCGACCCTGCCGC
TTACCGGATACCTGTCCGCCTTTCTCCCTTCGGGAAGCGTGGCGCTTTCTCATAGCTCACGCTGTAGGTATCTCAGTTCGGTGTAG
GTCGTTCGCTCCAAGCTGGGCTGTGTGCACGAACCCCCCGTTCAGCCCGACCGCTGCGCCTTATCCGGTAACTATCGTCTTGAGTC
CAACCCGGTAAGACACGACTTATCGCCACTGGCAGCAGCCACTGGTAACAGGATTAGCAGAGCGAGGTATGTAGGCGGTGCTACAG
AGTTCTTGAAGTGGTGGCCTAACTACGGCTACACTAGAAGAACAGTATTTGGTATCTGCGCTCTGCTGAAGCCAGTTACCTTCGGA
AAAAGAGTTGGTAGCTCTTGATCCGGCAAACAAACCACCGCTGGTAGCGGTGGTTTTTTTGTTTGCAAGCAGCAGATTACGCGCAG
AAAAAAAGGATCTCAAGAAGATCCTTTGATCTTTTCTACGGGGTCTGACGCTCAGTGGAACGAAAACTCACGTTAAGGGATTTTGG
TCATGAGATTATCAAAAAGGATCTTCACCTAGATCCTTTTAAATTAAAAATGAAGTTTTAAATCAATCTAAAGTATATATGAGTAA
ACTTGGTCTGACAGTTACCAATGCTTAATCAGTGAGGCACCTATCTCAGCGATCTGTCTATTTCGTTCATCCATAGTTGCCTGACT
CCCCGTCGTGTAGATAACTACGATACGGGAGGGCTTACCATCTGGCCCCAGTGCTGCAATGATACCGCGAGACCCACGCTCACCGG
CTCCAGATTTATCAGCAATAAACCAGCCAGCCGGAAGGGCCGAGCGCAGAAGTGGTCCTGCAACTTTATCCGCCTCCATCCAGTCT
ATTAATTGTTGCCGGGAAGCTAGAGTAAGTAGTTCGCCAGTTAATAGTTTGCGCAACGTTGTTGCCATTGCTACAGGCATCGTGGT
GTCACGCTCGTCGTTTGGTATGGCTTCATTCAGCTCCGGTTCCCAACGATCAAGGCGAGTTACATGATCCCCCATGTTGTGCAAAA
AAGCGGTTAGCTCCTTCGGTCCTCCGATCGTTGTCAGAAGTAAGTTGGCCGCAGTGTTATCACTCATGGTTATGGCAGCACTGCAT
AATTCTCTTACTGTCATGCCATCCGTAAGATGCTTTTCTGTGACTGGTGAGTACTCAACCAAGTCATTCTGAGAATAGTGTATGCG
GCGACCGAGTTGCTCTTGCCCGGCGTCAATACGGGATAATACCGCGCCACATAGCAGAACTTTAAAAGTGCTCATCATTGGAAAAC
GTTCTTCGGGGCGAAAACTCTCAAGGATCTTACCGCTGTTGAGATCCAGTTCGATGTAACCCACTCGTGCACCCAACTGATCTTCA
GCATCTTTTACTTTCACCAGCGTTTCTGGGTGAGCAAAAACAGGAAGGCAAAATGCCGCAAAAAAGGGAATAAGGGCGACACGGAA
```

ATGTTGAATACTCATACTCTTCCTTTTTCAATATTATTGAAGCATTTATCAGGGTTATTGTCTCATGAGCGGATACATATTTGAAT
GTATTTAGAAAAATAAACAAATAGGGGTTCCGCGCACATTTCCCCGAAAAGTGCCACCTGACGTCTAAGAAACCATTATTATCATG
ACATTAACCTATAAAAATAGGCGTATCACGAGGCCCTTTCGTCTCGCGCGTTTCGGTGATGACGGTGAAAACCTCTGACACATGCA
GCTCCCGGAGACGGTCACAGCTTGTCTGTAAGCGGATGCCGGGAGCAGACAAGCCCGTCAGGGCGCGTCAGCGGGTGTTGGCGGGT
GTCGGGGCTGGCTTAACTATGCGGCATCAGAGCAGATTGTACTGAGAGTGCACCATATGCGGTGTGAAATACCGCACAGATGCGTA
AGGAGAAAATACCGCATCAGGCGCCATTCGCCATTCAGGCTGCGCAACTGTTGGGAAGGGCGATCGGTGCGGGCCTCTTCGCTATT
ACGCCAGCTGGCGAAAGGGGGATGTGCTGCAAGGCGATTAAGTTGGGTAACGCCAGGGTTTTCCCAGTCACGACGTTGTAAAACGA
CGGCCAGTGAATTCGAGCTCGGTACCTCGCGAATGCATCTAGATCGTGCGGTGCGAGCTTGTTTGCACCTGCGAGCGTGACCCAAG
CTTTATGAATCCCCCGTCGACAATCACGCATCCAATGAGACGCTACCATATTGTCAGAGCGGATCGTTGCTGCAACCAATGAATGG
TGCTGTAGATTCGTATCGCGGATAGAGCAAGCACGATCCTCTCTGCAGCCAATCGTGGATACTACACGTGTACCAAATTGTGAATC
GTATGCTCGAAGATGGGAAGAGAAAATCACAACGCGCGCTTCACCCTCGCACAGCAGCAAGCTATCCACCACCGACCCAAGCGAAA
CGCGTATGATTGAACAAGATGGCCTACATGCAGGTTCTCCAGCTGCCTGGGTTGAGAGACTGTTTGGCTATGACTGGGCACAGCAG
ACCATTGGTTGCTCTGATGCAGCAGTTTTCAGACTTTCAGCCCAAGGCAGGCCAGTCCTTTTTGTAAAGACAGACCTCAGTGGGGC
TCTCAATGAGCTCCAGGATGAGGCTGCCAGACTCTCCTGGTTGGCAACAACTGGGGTCCCCTGTGCAGCTGTCCTTGATGTGGTCA
CAGAAGCTGGAAGGGACTGGCTCCTACTAGGTGAGGTGCCTGGGCAGGACCTCCTTTCCTCTCACCTAGCTCCAGCTGAGAAAGTG
TCAATCATGGCTGATGCCATGAGAAGACTCCACACCCTTGACCCAGCCACCTGCCCCTTTGACCACCAGGCCAAGCACAGGATAGA
GAGGGCCAGAACCAGGATGGAGGCTGGCCTGGTGGACCAAGATGACTTGGATGAAGAACACCAGGGCCTGGCCCCTGCTGAACTAT
TTGCCAGGCTCAAGGCATCCATGCCAGATGGTGAGGACCTAGTGGTGACTCATGGGGATGCCTGCCTTCCCAACATCATGGTTGAA
AATGGAAGGTTCTCTGGCTTCATAGACTGTGGCAGGCTGGGAGTGGCTGACAGGTACCAGGACATTGCCCTAGCAACCAGGGACAT
AGCAGAAGAGCTAGGGGGAGAGTGGGCAGACAGGTTCCTAGTGCTCTATGGCATTGCAGCCCCTGACTCCCAGAGAATTGCCTTCT
ACAGACTTCTTGATGAGTTCTTCTAACTCGAGGCTTGTTGCTTATAATAAGAACCAACCGGTGTTTCACAACACCACCTTCAGAGG
AAGAAGTTGACTCGGGACACGTTATCGGAGGAATGTGGACATAGTTGGAATCACAGGGTGTGTACGCGCGCAGGAGTTTTCGGTCC
ATGAGGAAATTGCTTCCAATATCTAAGCAGCCAAAACGTGTGCATCAAAGAAGTCATTGTACGAAACTATTGTACATCTGATGGGG
CTAAGCGCGGCTCATCACAACATGACCTTTCAATGAAGTCGATTCCCTTATATGTTACCTGCAGGATGCACTCA

#### **Haptophytes**

*Isochrysis galbana*

pIgNAT

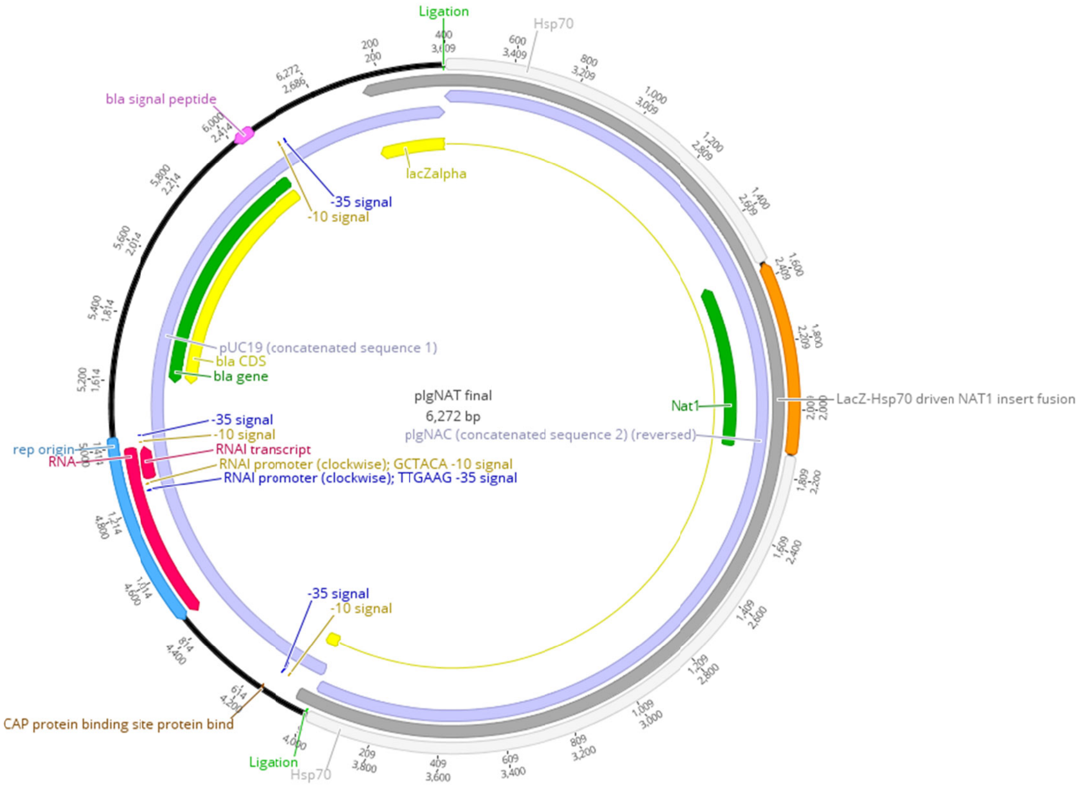


```
>pIgNAT_final
TCGCGCGTTTCGGTGATGACGGTGAAAACCTCTGACACATGCAGCTCCCGGAGACGGTCACAGCTTGTCTGTAAGCGGATGCCGGG
AGCAGACAAGCCCGTCAGGGCGCGTCAGCGGGTGTTGGCGGGTGTCGGGGCTGGCTTAACTATGCGGCATCAGAGCAGATTGTACT
GAGAGTGCACCATATGCGGTGTGAAATACCGCACAGATGCGTAAGGAGAAAATACCGCATCAGGCGCCATTCGCCATTCAGGCTGC
GCAACTGTTGGGAAGGGCGATCGGTGCGGGCCTCTTCGCTATTACGCCAGCTGGCGAAAGGGGGATGTGCTGCAAGGCGATTAAGT
TGGGTAACGCCAGGGTTTTCCCAGTCACGACGTTGTAAAACGACGGCCAGTGAATTCCTCGTCGAGTGTGGCGCGTGGCGGCGGCC
TCTGGGCGGCGCGCCTCGGAGCCCTCCCGGAGCCATGTCGGTCCCTCCCCTAGGCGCGGGGAGCGGCCTCTGGGCGGCGCTGCTCC
GCAGGCGTGGCGTCGCCGTCGCCGCCTACGACGCCGTGCCGTGGGTGTGCGAGGCTGGCTTTGGCGAGGTGCAGCGCGGCGGCCCC
GAGCGGGTGGGCCGCCATGCGGACGCGTCTCTGCTGCTCTGCTGGCCGCCGCTCGAGGCCTCCGCCTCCGGCAGCAACACGATGGG
CGTCCGAGCGCTGCGCGCCATGACGGGCAGAGGCGCGCTCCGCCCGAGCTGTGGTTGCCGGCCGCGCGGCACGCCTCACGCCCCGC
TCACGCGCCCTCCCCCGCCGCGGCGCCACACCTCAGGCGCGCTTCTGTACGTCGGCGAGTGGCGCGGCGCCGTCGGCGTCCTCTCT
TCGCTGGCGCCTTCCACCGCCGACCACGGGATGGTTGGCGGGCCGCAGCTGCAGGCCGAGGTGGAGCGGCACTGGCGGCTCGCCGA
GTCCGTCGCCCTGCCCCGCTGGCCAGGCGTGGGGAGCCCGCAGGCGGGGCACCGCCTCTACGTCTTTACTCGCCGCCAGCCACCGC
CCGAGCCCGCGGAGGAGGATGCAGCGCCGCCGCCCCCGCCGCCCCCCCGCCGCCGCCGCCGCCGCCACCGCCGCCGCCTCCGC
CGCCGCCTCCGCCGCCGCCTCCGCCGCCGCCGCCGCCTCCGCCGCCCACTCAGCGACAGCCGCCGCCCGTTTCGGATGGCTACGCC
GCCCGGCGCGAGAGGCTCTCTCGGCTGCAGGGTGAAGTGGACGCGCCTGGCCGGGAGTGGGCGTGGCTCATCGCCAGCCTGCTGTG
GGAAGAAGCTTGCCGCCCAGAGGAGATTTGATGAAATGAAAAGTAATATATCTTTACTTCCATGTTCCAACACCCGGACGCGCGTT
CCCAGGCGCCAGGGTGATGGAAGGTTCGACGCACCCCCGGACTGGGCGGAGAAATGAAACATTTCGGAAAACGGTGTGGTCCTATA
CACCCGCCCACACCGCGACACCCCTCGCACGCGGTCATCTCCGCTCGGTGCATATCCCTAGGGTGGGAGCTTCTCCTCTGAATGCA
CTCTGAGCCGAATCAGGGGCAGGGCATCGACATGTAGAGGGCCTGCTCGCCGTCCGAGGCCGTGCCGTCGTAGAGCGCCGTGTCGA
GGCCGCAGAGGGTGAAGCCCATGCGACGGTAGGCGTGGATGGCCGGGGCGTTGACGTTCGTGACCTCGAGCCAGAGGTGGCCAGCG
CCGCGCTCGCGCGCGAACTCGGTGGCGAGGCCCATGAGCGCGCGGCCGACGCCGTGGCCGCGGTGCTCGGGGGCGACCTCGATGTC
CTCGACGGTGAGGCGACGGTTCCAGCCCGAGTACGAGACGACGACGAAGCCGGCGAGGTCGCCGTCGTCGCCGTAGGCGACGAACG
TGCGCGAGTCCGGGTCGCCGTCCTCGCCGGCGTCGGACTCGTCGTCCGACTCGTCGTCGGGGAAGACCTTCGTGAGGGGCGGGTCG
ACGGGGACCTCGCGGAGCGTGAAGCCGTCGCCCGTGGCGGTGACGCGGAAGACGGTGTCCGTGGTGAACGAGCCGTCGAGCGCCTC
GATGGCCTCCGCGTCGCCGGGCACCGACGTGCGGTAGCGGTACGCCGTGTCGTCGAGCGTCGCCGCCGCCATGGTGTCCAAGAGCG
CCTACTCGCGTGACGGCGCTGCGTCACAGGCAGGGGGTCAACAGGGGCTAGGCGCCACCGCGGGCGCACGTCTCAAGTCGAGAGCC
TTGCACAACACGTGTTCGAGGCTCGCGACAAAAAATACCCCTGCTCCTCAAATCCTTGCTGGGTCGCTGTCCCTTGTCAGGAGCAG
GAGAACTAGGAGAGCGCAACCAGTAGGAGGCCCGCGCCTGCCCCCGTGCCGCCGGGTGCAACCTGTCCTATACGAGAGAGGAATCC
CGGAATCCCTGGGACCCTGTCGTGTGGGTGAACCCTACCCAGCCCCCCTCCGTGTCGCGATGCGCTCCCCGCTCCCGTTCCCCTGT
GTCCCTGATCGTCCTTCTCCAAGATCCTGCTCGTGCAGGTGGGGACCTTGTCCTCCTCCCATCCCTCGTTTGGTTTGTCTGTTCTC
TCTCTCTCTCTGCACCTTGGCCAATGTTTAGCGTTTAGGCCCAAAAGCCTCTCGAAAGGACAGACCAAGCAAGTCCGTGGCACT
```

GGCAAGAGGTGAAGAGCCACATCGGCGAACACACCTGCTTGGAGCACTTGGAGTGCAGTCGGTAACCTCACATCCAGGGCCGACCT
CTGCTGTTGCTTTCATCTCGCGCGATCTGCCTCTTGCGTCTTGGGTCTTGCCAAGAGCCAAGAGGCAAGACCTTGAAGTGGTCTTG
CGTAAGGAGATTGGCCGCCCGATAGTGCGGTCTGTGTGCTGGGGCCTTTTCTAGATTTCTACATCTCTAGCTACTCGCGTGACGGC
GCTGCGTCACAGGCAGGGGGTCAACAGGGGCTAGGCGCCACCGCGGGCGCACGTCTCAAGTCGAGAGCCTTGCACAACACGTGTTC
GAGGCTCGCGACAAAAAATACCTTCCTCCTCCTCAAATCCTTGCTGGGTCGCTGTCCCTTGTCAGGAGCAGGAGAACTAGGAGAGC
GCAACCAGTAGGAGGCCCGCGCCTGCCCCCGTGCCGCCGAGTGCAACCTGTCCTACACGAGAGAGGAATCCCGGAATCCCTGTGGG
ACCCTGTCGTGTGTGTGGACCCTGTTTCTTCTGTTTCAGCCCCCCTCCGTGTCGCGATGCGCTCCCCGCTCCCGTTCCCCTGTGTC
CCTGATCGTCCTTCTCCAAGATCCTGCTCGTGCAGGTGGGGACCTTGTCCTCCAGGCCCAAAAGCCTCTCGAAACACGAAAGGACA
GACCACAGCAAGTCTGGACTGGGGCACTGGCAAGAGCCATGCGAACACCTGCTCTTGCGAGCCTTGGAGCACTTGGAGCAGCCTTG
CTTGTTTTTTTTCACCTCTATAACTCCTGTCCACGTGGCGTGGTGAGGGACTCCGGAAACAAAGCCCATGCATGGGCTTTCAGGAG
TCATGGGCTTTCCGGAGTCCCTCACCACGCATGTGCATGTGCATGTGCATGTGCATGTGCATGTGCATGCAGCCTCCCTCCTCAAA
GGCGTGCGTGGACGCCGCCGCTTGCCCATTTTTGAACGCCGTGAAGAAGTTAACTTTGGTCATCATGCGTCATGGATGGACACTGA
TATAAGTCAGCCCCGGTCGGGAATTTCTTCCTCTGACCGAAAGGAACGTTTCTAAATTCTTCCTTTGCCCGAGAGACGCTAGAACT
CGTCCGTTCCCCGCGACGTGCGTCCCAGCGAGCGAGACATAACAAAGCAATGAGCCATCGCAGCGAGAGGAGCGACAAGAGCGATG
GCACGAGCGCGCCGCCGCCGCGTCCCGCCGCGCCACAGGCGCCGTCTCTGATGTCGAGGCTGAACCACGGCACGACGAGGTCGGGG
AGCAACGTACGGAGCAGCGTGCTGAGCGTGCGTGTCCCCTCACCTCGGATCCTCTAGAGTCGACCTGCAGGCATGCAAGCTTGGCG
TAATCATGGTCATAGCTGTTTCCTGTGTGAAATTGTTATCCGCTCACAATTCCACACAACATACGAGCCGGAAGCATAAAGTGTAA
AGCCTGGGGTGCCTAATGAGTGAGCTAACTCACATTAATTGCGTTGCGCTCACTGCCCGCTTTCCAGTCGGGAAACCTGTCGTGCC
AGCTGCATTAATGAATCGGCCAACGCGCGGGGAGAGGCGGTTTGCGTATTGGGCGCTCTTCCGCTTCCTCGCTCACTGACTCGCTG
CGCTCGGTCGTTCGGCTGCGGCGAGCGGTATCAGCTCACTCAAAGGCGGTAATACGGTTATCCACAGAATCAGGGGATAACGCAGG
AAAGAACATGTGAGCAAAAGGCCAGCAAAAGGCCAGGAACCGTAAAAAGGCCGCGTTGCTGGCGTTTTTCCATAGGCTCCGCCCCC
CTGACGAGCATCACAAAAATCGACGCTCAAGTCAGAGGTGGCGAAACCCGACAGGACTATAAAGATACCAGGCGTTTCCCCCTGGA
AGCTCCCTCGTGCGCTCTCCTGTTCCGACCCTGCCGCTTACCGGATACCTGTCCGCCTTTCTCCCTTCGGGAAGCGTGGCGCTTTC
TCATAGCTCACGCTGTAGGTATCTCAGTTCGGTGTAGGTCGTTCGCTCCAAGCTGGGCTGTGTGCACGAACCCCCCGTTCAGCCCG
ACCGCTGCGCCTTATCCGGTAACTATCGTCTTGAGTCCAACCCGGTAAGACACGACTTATCGCCACTGGCAGCAGCCACTGGTAAC
AGGATTAGCAGAGCGAGGTATGTAGGCGGTGCTACAGAGTTCTTGAAGTGGTGGCCTAACTACGGCTACACTAGAAGAACAGTATT
TGGTATCTGCGCTCTGCTGAAGCCAGTTACCTTCGGAAAAAGAGTTGGTAGCTCTTGATCCGGCAAACAAACCACCGCTGGTAGCG
GTGGTTTTTTTGTTTGCAAGCAGCAGATTACGCGCAGAAAAAAAGGATCTCAAGAAGATCCTTTGATCTTTTCTACGGGGTCTGAC
GCTCAGTGGAACGAAAACTCACGTTAAGGGATTTTGGTCATGAGATTATCAAAAAGGATCTTCACCTAGATCCTTTTAAATTAAAA
ATGAAGTTTTAAATCAATCTAAAGTATATATGAGTAAACTTGGTCTGACAGTTACCAATGCTTAATCAGTGAGGCACCTATCTCAG
CGATCTGTCTATTTCGTTCATCCATAGTTGCCTGACTCCCCGTCGTGTAGATAACTACGATACGGGAGGGCTTACCATCTGGCCCC
AGTGCTGCAATGATACCGCGAGACCCACGCTCACCGGCTCCAGATTTATCAGCAATAAACCAGCCAGCCGGAAGGGCCGAGCGCAG
AAGTGGTCCTGCAACTTTATCCGCCTCCATCCAGTCTATTAATTGTTGCCGGGAAGCTAGAGTAAGTAGTTCGCCAGTTAATAGTT
TGCGCAACGTTGTTGCCATTGCTACAGGCATCGTGGTGTCACGCTCGTCGTTTGGTATGGCTTCATTCAGCTCCGGTTCCCAACGA
TCAAGGCGAGTTACATGATCCCCCATGTTGTGCAAAAAAGCGGTTAGCTCCTTCGGTCCTCCGATCGTTGTCAGAAGTAAGTTGGC
CGCAGTGTTATCACTCATGGTTATGGCAGCACTGCATAATTCTCTTACTGTCATGCCATCCGTAAGATGCTTTTCTGTGACTGGTG
AGTACTCAACCAAGTCATTCTGAGAATAGTGTATGCGGCGACCGAGTTGCTCTTGCCCGGCGTCAATACGGGATAATACCGCGCCA
CATAGCAGAACTTTAAAAGTGCTCATCATTGGAAAACGTTCTTCGGGGCGAAAACTCTCAAGGATCTTACCGCTGTTGAGATCCAG
TTCGATGTAACCCACTCGTGCACCCAACTGATCTTCAGCATCTTTTACTTTCACCAGCGTTTCTGGGTGAGCAAAAACAGGAAGGC
AAAATGCCGCAAAAAAGGGAATAAGGGCGACACGGAAATGTTGAATACTCATACTCTTCCTTTTTCAATATTATTGAAGCATTTAT
CAGGGTTATTGTCTCATGAGCGGATACATATTTGAATGTATTTAGAAAAATAAACAAATAGGGGTTCCGCGCACATTTCCCCGAAA
AGTGCCACCTGACGTCTAAGAAACCATTATTATCATGACATTAACCTATAAAAATAGGCGTATCACGAGGCCCTTTCGTC

*Emiliania huxleyi*

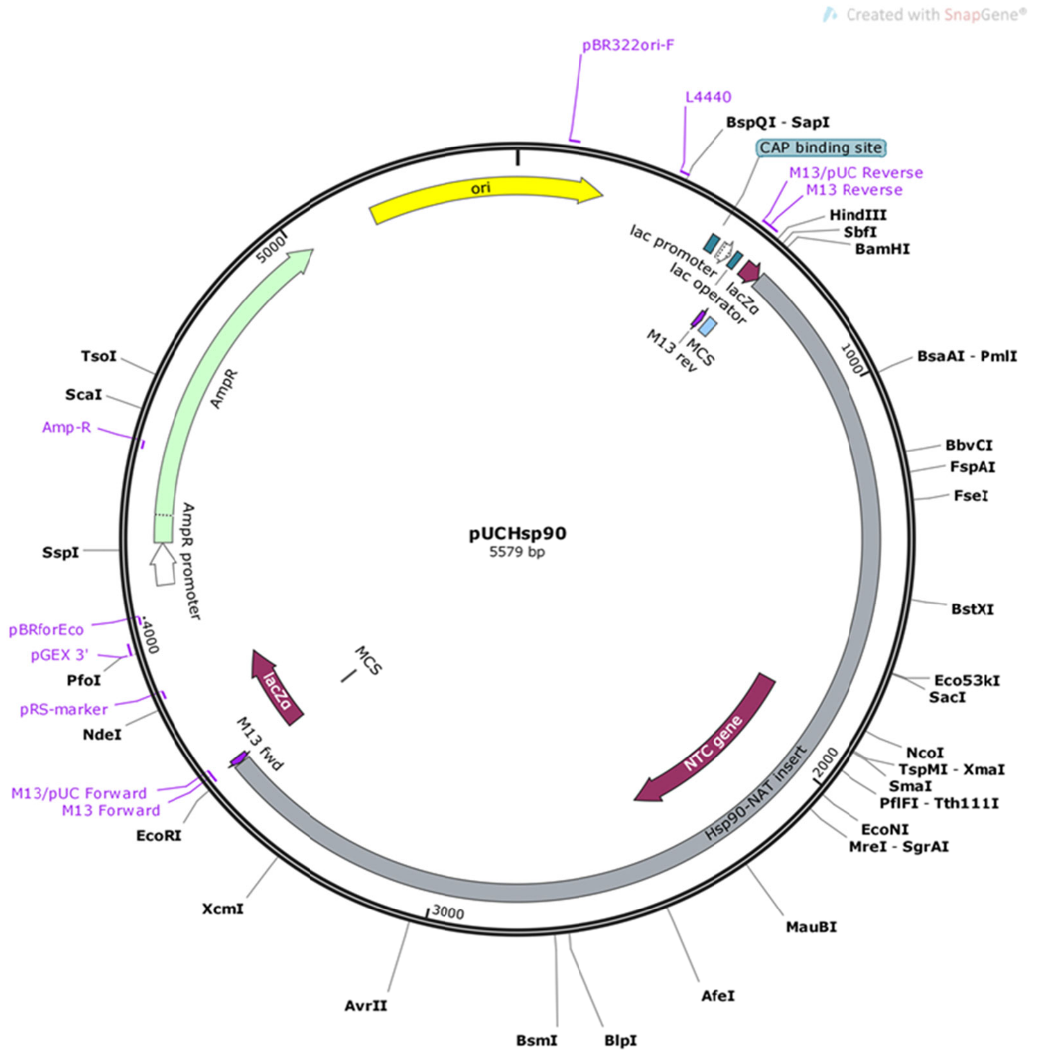


```
>pUCHsp90  (5579 bp)
gagatacctacagcgtgagctatgagaaagcgccacgcttcccgaagggagaaaggcggacaggtatccggtaagcggcagggtcg
gaacaggagagcgcacgagggagcttccagggggaaacgcctggtatctttatagtcctgtcgggtttcgccacctctgacttgag
cgtcgatttttgtgatgctcgtcaggggggcggagcctatggaaaaacgccagcaacgcggcctttttacggttcctggccttttg
ctggccttttgctcacatgttctttcctgcgttatcccctgattctgtggataaccgtattaccgcctttgagtgagctgataccg
ctcgccgcagccgaacgaccgagcgcagcgagtcagtgagcgaggaagcggaagagcgcccaatacgcaaaccgcctctccccgcg
cgttggccgattcattaatgcagctggcacgacaggtttcccgactggaaagcgggcagtgagcgcaacgcaattaatgtgagtta
gctcactcattaggcaccccaggctttacactttatgcttccggctcgtatgttgtgtggaattgtgagcggataacaatttcaca
caggaaacagctatgaccatgattacgccaagcttgcatgcctgcaggtcgactctagagGATCCAGGCCGAGGAGGTGGCGCGGC
TCGTGGTCCGGCTGGCGTACGAGCGGAAGAGGAAGCGGATGCCGCTCGCAGAGGCGCTGCCGCGGCTGGAGGCGCTAGTCGAGGCG
GCGGGGGAGGAGGTGGCGGCATCGCCGTCGCAGAGCCTCGCAGGGGTGGTGACGGCCGAGGCGGACGGGGAGGCGCGGCTGTGCGA
CCTCTGCTGGTCGGCGCCGCGGTCTGTTCGCTTCGCGTGCGGCCACGCGCTGTACTGCGAGACGTGTGCGCCGCGCGTGCTGGAGC
GCGACACCAACTGCCCGGCCTGCCGCCAGCCGGCGCTGCCGATCGCCAACTGCGGAGCGCACGTGGCGGCGCAGGCGACCTTTGTC
TACCAGCCCCCACGCGCCCCGCGCCGCCCACACAGCCCGCGGGTGGCATGGCGGTGGTCGGTGGGCGAGGCGGGCGCAGGCGGGG
TGGGCGGGGCGGGAGGGGTGGGCGGGGGAGGCCTCGGGGCGAGGGGCGTGGCACTGGCGGCTCGTAGCTTAGGCCAGCCTCAGCAG
CGAGCCGGTCTCCACGACCAGCGCAGGGCAGGGCGTGCGCATGGCAGGAGCGCATGCGGCATCGGGGCATGGCAGGAGGCGCAGAG
CTGCTGGTAGGCCGGCCGGGGCCGATTTTATTAAATTTTCAACTTTAAGCTCATGCCTCGAGACGAGTATGCAAACGTCGCCGTTT
GCGAGCTGCTCGACGCCTCGCCCCTCCCTCTCACCCGCCTTCCACTCGCCGTCGTAGACGTCGCCGGCTCGCCGCGCGTATGCTAA
GTCCTACGAAGAGTATGCAGACTACCAGACCCTATGGACCACAGAACCGCAGCGGAACGGTCCATTAGACTGGTCCCTGAGACATA
TTCTAACCGTACGCCACCCGCACTGCCACACCGCGCAGCAGCCTCGCCGTGGCCCGTGCAGCGTGCGGGTGAAGAGATTTGTCCCC
CGCCAAATAAAAAAACGGCCGCTTGCTTCTAGACGAGTCTACGCGGCACATCCACAAGTGCACAGAGCTCGCCCGCTCGCCCGCCC
CGCCCCGGCCCGGCCTCGCCCCGCCTCGCCGCCTCTCCTCCGCCACACCCGCACCGCCCTCCGCCGACTCGCCCGGCGACCCACCT
CCCCGCTCCTCCCCGCGCCGCAGAACAACCAGTCGCCATGGCCGCGGCCACGCTCGACGACACCGCGTACCGCTACCGCACGTCCG
TCCCGGGCGACGCGGAGGCCATCGAGGCCCTCGACGGCTCGTTCACCACGGACACCGTCTTCCGCGTCACCGCCACGGGCGACGGC
```

TTCACGCTCCGCGAGGTCCCCGTCGACCCGCCCCTCACGAAGGTCTTCCCCGACGACGAGTCGGACGACGAGTCCGACGCCGGCGA
GGACGGCGACCCGGACTCGCGCACGTTCGTCGCCTACGGCGACGACGGCGACCTCGCCGGCTTCGTCGTCGTCTCGTACTCGGGCT
GGAACCGTCGCCTCACCGTCGAGGACATCGAGGTCGCCCCCGAGCACCGCGGCCACGGCGTCGGCCGCGCGCTCATGGGCCTCGCC
ACCGAGTTCGCGCGCGAGCGCGGCGCTGGCCACCTCTGGCTCGAGGTCACGAACGTCAACGCCCCGGCCATCCACGCCTACCGTCG
CATGGGCTTCACCCTCTGCGGCCTCGACACGGCGCTCTACGACGGCACCGCCTCCGACGGCGAGCAGGCCCTCTACATGTCCATGC
CCTGCCCGTGAGCGCGCGAGTGTCGGGGTGAGCGCAGCGCTGCGCTCTCCCGCCTCGAGCCGCCCCGAGGGCTCTCGCGCCTCGGC
GTCCTGGCGCGCTGGGGTGTCGCTTTCGGCTCTGTTCGGTTTTGTGTTTTTATTTCTTGTCATCCGGCTCTGCCATCTGTCAAAAA
ACTGTTCAGTTTTCAAAGGCAAAACACGACGCTCTGCGGTTGGTGCACCCCTCACACTGACACATTCCCCTCATATCCCTGCGTAC
CCTTGGCTGAGCACATCTACCTGCGAGCTAGTCTCTCCGCATTCTCTCCCCTAGTCATCCGCCCGCGCCGCGCACCCGTCATGCCA
GCACCGTCCAGTCCGTCCACCGACACGCGGCACACACCCGGAAGCGCAACGACGGCCGTCCTCCGGCTCCGCCAGGTGCTGCCGCG
GCCCGCGCGTCTCGGAGCGCCTGCACACGTCACGAGTGGGTACAGAGGAGCTGTGTGCACGGACACAATGCTCACATACTCACAGG
CTCATTGAGGACCCGCTCCGTTCCGGCTTCCGCCCCCCATCGCTCACCGCCCCAGACCCGACTCCTCGATCTTCTTGGGCAGCAGC
TGCTTCGACAGGTCCTCTCCGTGCCCTAGGCAGCCGGTCTCGCCAGCGCCGCAGCTCCAGAGGGCTCCGTCGGCGGTGATGGCGAG
GCTGTGGCCCACTCCAGCCGACACAGCGACGACGCGCTGGCCGGTGAAGGCCTCAACCTTCTTCGGCTGCAGCTGGTTCTGCTCGT
CGCCGTGGCCCAGCTGGCCAACGCCTCCATAGCCCCAGCTCCAGACGGCACCGTCGGCGGTGAGGGCGAGGCTGTAGTGGCCTCCA
GCCGACACAGCGACGACGCGGTGGCCGACAAAGGCCTCGACCTTCTTGGGCAGCAGCTGGCGCTGCTCCTCGCCGTGGCCCAGCTT
GCCAAAGGCTCCCTGGCCCCAGCTCCAGACGCTGCCGTTGGCGGTGAGGGCGAGGCTGTAGTGGCCTCCAGCCGACACAGCGACGA
CGCGCCCAGCCGACACAGCGACGACGCGCTGGCCAGCAAAGGCCTCGATCTTCTTCGGCAGCAGCTGGTCCTGCTCGTCGCCGTGG
CCGAGCTGGCCTTCGCGTCCCCAACCCCAGCTCCAGACAGCTCCGTCGGGaattcactggccgtcgttttacaacgtcgtgactgg
gaaaaccctggcgttacccaacttaatcgccttgcagcacatccccctttcgccagctggcgtaatagcgaagaggcccgcaccga
tcgcccttcccaacagttgcgcagcctgaatggcgaatggcgcctgatgcggtattttctccttacgcatctgtgcggtatttcac
accgcatatggtgcactctcagtacaatctgctctgatgccgcatagttaagccagccccgacacccgccaacacccgctgacgcg
ccctgacgggcttgtctgctcccggcatccgcttacagacaagctgtgaccgtctccgggagctgcatgtgtcagaggttttcacc
gtcatcaccgaaacgcgcgagacgaaagggcctcgtgatacgcctatttttataggttaatgtcatgataataatggtttcttaga
cgtcaggtggcacttttcggggaaatgtgcgcggaacccctatttgtttatttttctaaatacattcaaatatgtatccgctcatg
agacaataaccctgataaatgcttcaataatattgaaaaaggaagagtatgagtattcaacatttccgtgtcgcccttattccctt
ttttgcggcattttgccttcctgtttttgctcacccagaaacgctggtgaaagtaaaagatgctgaagatcagttgggtgcacgag
tgggttacatcgaactggatctcaacagcggtaagatccttgagagttttcgccccgaagaacgttttccaatgatgagcactttt
aaagttctgctatgtggcgcggtattatcccgtattgacgccgggcaagagcaactcggtcgccgcatacactattctcagaatga
cttggttgagtactcaccagtcacagaaaagcatcttacggatggcatgacagtaagagaattatgcagtgctgccataaccatga
gtgataacactgcggccaacttacttctgacaacgatcggaggaccgaaggagctaaccgcttttttgcacaacatgggggatcat
gtaactcgccttgatcgttgggaaccggagctgaatgaagccataccaaacgacgagcgtgacaccacgatgcctgtagcaatggc
aacaacgttgcgcaaactattaactggcgaactacttactctagcttcccggcaacaattaatagactggatggaggcggataaag
ttgcaggaccacttctgcgctcggcccttccggctggctggtttattgctgataaatctggagccggtgagcgtgggtctcgcggt
atcattgcagcactggggccagatggtaagccctcccgtatcgtagttatctacacgacggggagtcaggcaactatggatgaacg
aaatagacagatcgctgagataggtgcctcactgattaagcattggtaactgtcagaccaagtttactcatatatactttagattg
atttaaaacttcatttttaatttaaaaggatctaggtgaagatcctttttgataatctcatgaccaaaatcccttaacgtgagttt
tcgttccactgagcgtcagaccccgtagaaaagatcaaaggatcttcttgagatcctttttttctgcgcgtaatctgctgcttgca
aacaaaaaaaccaccgctaccagcggtggtttgtttgccggatcaagagctaccaactctttttccgaaggtaactggcttcagca
gagcgcagataccaaatactgttcttctagtgtagccgtagttaggccaccacttcaagaactctgtagcaccgcctacatacctc
gctctgctaatcctgttaccagtggctgctgccagtggcgataagtcgtgtcttaccgggttggactcaagacgatagttaccgga
taaggcgcagcggtcgggctgaacggggggttcgtgcacacagcccagcttggagcgaacgacctacaccgaact

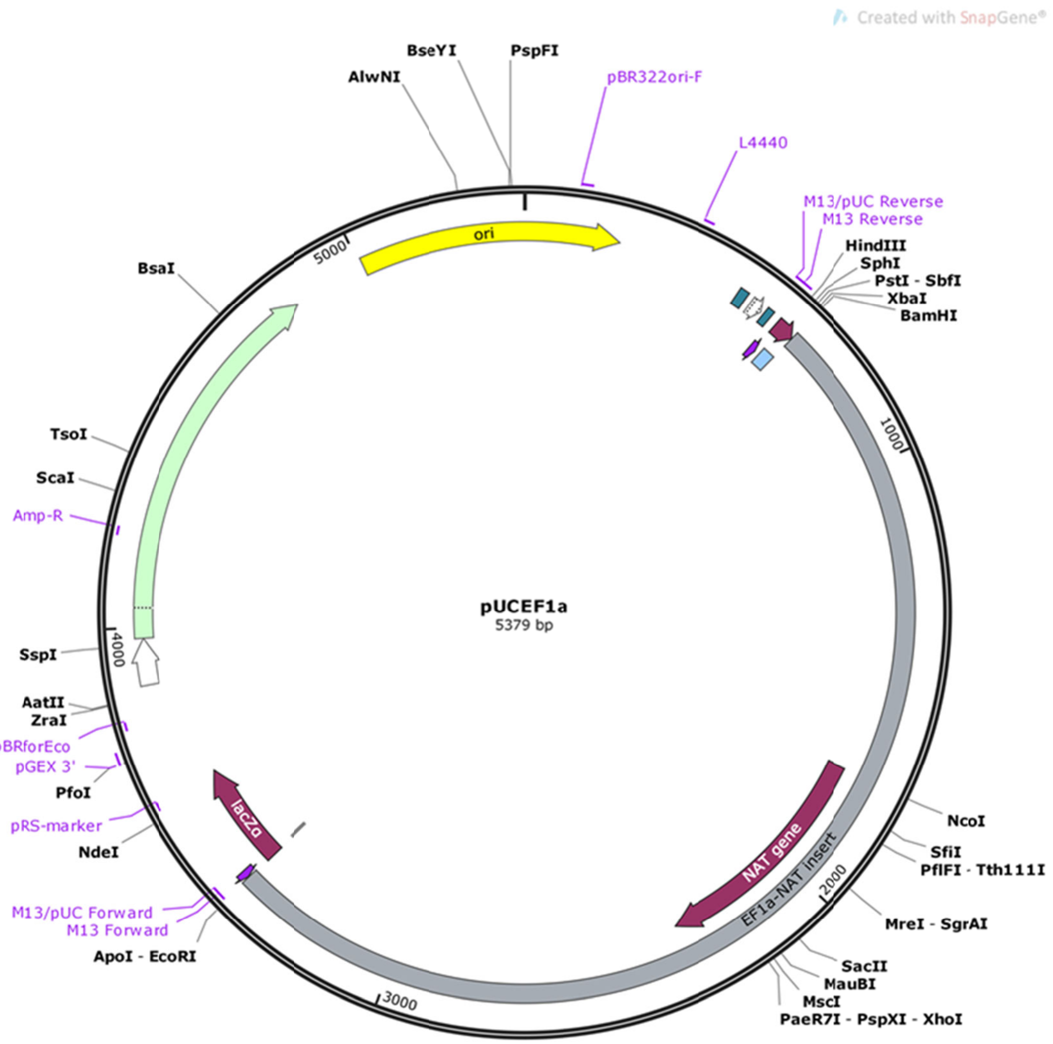


>pUCEF1a(5379bp)
gagatacctacagcgtgagctatgagaaagcgccacgcttcccgaagggagaaaggcggacaggtatccggtaagcggcagggtcg
gaacaggagagcgcacgagggagcttccagggggaaacgcctggtatctttatagtcctgtcgggtttcgccacctctgacttgag
cgtcgatttttgtgatgctcgtcaggggggcggagcctatggaaaaacgccagcaacgcggcctttttacggttcctggccttttg
ctggccttttgctcacatgttctttcctgcgttatcccctgattctgtggataaccgtattaccgcctttgagtgagctgataccg
ctcgccgcagccgaacgaccgagcgcagcgagtcagtgagcgaggaagcggaagagcgcccaatacgcaaaccgcctctccccgcg
cgttggccgattcattaatgcagctggcacgacaggtttcccgactggaaagcgggcagtgagcgcaacgcaattaatgtgagtta
gctcactcattaggcaccccaggctttacactttatgcttccggctcgtatgttgtgtggaattgtgagcggataacaatttcaca
caggaaacagctatgaccatgattacgccaagcttgcatgcctgcaggtcgactctagagGATCCCGGGAGATGACCGCACGACCA
CGTGGCGTCGGGAAATGACCGGTTTGGGGGGGATAAGGCTGATCCCCCCTGGGCGTCTTTCGGAAAACGACCGCTTCGACCACGTG
GCGTCGGGAAATGACCGGTCAAGTGACCGGTCCAATATGAAATGACCGGTCCCAACGAGGGGCCGGAATCCGAGGTAGAGCACGGC
CGCTCCAGGGTCCCACCTCCCACCTGCACGAGCAGGATTTTGGAAGGACGATCAGGGGCACAGGTGTAAACACACTGAAGGATCGA
AGAGCCGCACTTGCATCTTGCTTGCTTCCGGCCTTCTGGAAAAAGCAAGGGTTGCGTTAGGGTTAGGGGATTAGGGGGTCGTTAGG
GTTAGGGTTTGTTGTCGTCGCGCCGCTCTCGCTCGATACAGAGACACGGAAAGCAGACGTGACCCAAGCTGTGAGGCGACTATTTG
AGAGAGAGAGACTATTTGTCATCGCTCATCGGGAGAGCTCGCCGCCGACTCTACGCCGCCGCCGCCGCCCTCCCGAGGACAGATAA
TAGCCCTCAACAAGTCGACGAGTGTGTCCCGGCAAGACTCATGTGGGGTGCGAATGGAATCAGTTTTGTCCTCGTTTCCAAGGAGA
AAGAGGAAAAGAAAAACACTTGTCGAGCCACGACGTAGCAACCAAGGTGACGCAGACGCACGCCCACAAAGCCCACAGCGCCGCAG
CCGCGCGGCGCGGCGAGGCGGGCAGGTCGGCGGGGACGCGCCGCTCGCCGACCGCGCGCCCGGCGGGCGACCGCCGCCCTGCACCA
GGGGGGCGCCGGGGCACGGGAGGTCGGGAGCGCGGAGTCGCAAAACTTGCGTCCCCTGCGCTGCGCAGACCTCACCAGGGTCGGCG
CGCCGCCGGCACGCGGGCTGCGCGCCTCCCGCCGTTCCGTCGCGGCTGCCCGCCCCACGTCCCGAATCGCTCCCCCGTCCGCGCGC
TATTCCGTCTGCCTGCGCGCGTCCTGGCCGTGTCGCAGAGCCCCGGCCTCTGACACCTCGCCCGCCCCCGCGCAGAAAGCACCACC
CCTACCCGCAACCATGGCCGCCGCCACGCTCGACGACACCGCCTACCGCTACCGCACCTCCGTCCCGGGCGACGCCGAGGCCATCG
AGGCCCTCGACGGCTCGTTCACCACGGACACCGTCTTCCGCGTCACCGCCACGGGCGACGGCTTCACGCTCCGCGAGGTCCCCGTC
GACCCGCCCCTCACGAAGGTCTTCCCCGACGACGAGTCGGACGACGAGTCCGACGCCGGCGAGGACGGCGACCCGGACTCGCGCAC
GTTCGTCGCCTACGGCGACGACGGCGACCTCGCCGGCTTCGTCGTCGTCTCGTACTCGGGCTGGAACCGTCGCCTCACCGTCGAGG
ACATCGAGGTCGCCCCCGAGCACCGCGGCCACGGCGTCGGCCGCGCGCTCATGGGCCTCGCCACCGAGTTCGCGCGCGAGCGCGGC
GCTGGCCACCTCTGGCTCGAGGTCACGAACGTCAACGCCCCGGCCATCCACGCCTACCGTCGCATGGGCTTCACCCTCTGCGGCCT
CGACACGGCGCTCTACGACGGCACCGCCTCGGACGGCGAGCAGGCCCTCTACATGTCCATGCCCTGCCCGTGACGGGAGATGACCG

CACGACCACGTGGCGTCGGGAAATGACCGGTTTGGGGGGGATAAGGCTGATCCCCCCTGGGCGTCTTTCGGAAAACGACCGCTTCG
ACCACGTGGCGTCGGGAAATGACCGGTCAAGTGACCGGTCCAATATGAAATGACCGGTCCCAACGAGGGGCCGGAATCCGAGGTAG
AGCACGGCCGCTCCAGGGTCCCACCTCCCACCTGCACGAGCAGGATTTTGGAAGGACGATCAGGGGCACAGGTGTAAACACACTGA
AGGATCGAAGAGCCGCACTTGCATCTTGCTTGCTTCCGGCCTTCTGGAAAAAGCAAGGGTTGCGTTAGGGTTAGGGGATTAGGGGG
TCGTTAGGGTTAGGGTTTGTTGTCGTCGCGCCGCTCTCGCTCGATACAGAGACACGGAAAGCAGACGTGACCCAAGCTGTGAGGCG
ACTATTTGAGAGAGAGAGACTATTTGTCATCGCTCATCGGGAGAGCTCGCCGCCGACTCTACGCCGCCGCCGCCGCCCTCCCGAGG
ACAGATAATAGCCCTCAACAAGTCGACGAGTGTGTCCCGGCAAGACTCATGTGGGGTGCGAATGGAATCAGTTTTGTCCTCGTTTC
CAAGGAGAAAGAGGAAAAGAAAAACACTTGTCGAGCCACGACGTAGCAACCAAGGTGACGCAGACGCACGCCCACAAAGCCCACAG
CGCCGCAGCCGCGCGGCGCGGCGAGGCGGGCAGGTCGGCGGGGACGCGCCGCTCGCCGACCGCGCGCCCGGCGGGCGACCGCCGCC
CTGCACCAGGGGGGCGCCGGGGCACGGGAGGTCGGGAGCGCGGAGTCGCAAAACTTGCGTCCCCTGCGCTGCGCAGACCTCACCAG
GGTCGGCGCGCCGCCGGCACGCGGGCTGCGCGCCTCCCGCCGTTCCGTCGCGGCTGCCCGCCCCACGTCCCGAATCGCTCCCCCGT
CCGCGCGCTATTCCGTCTGCCTGCGCGCGTCCTGGCCGTGTCGCAGAGCCCCGGCCTCTGACACCTCGCCCGCCCCCGCGCAGAAA
GCACCACCCCTACCCGCAACCGaattcactggccgtcgttttacaacgtcgtgactgggaaaaccctggcgttacccaacttaatc
gccttgcagcacatccccctttcgccagctggcgtaatagcgaagaggcccgcaccgatcgcccttcccaacagttgcgcagcctg
aatggcgaatggcgcctgatgcggtattttctccttacgcatctgtgcggtatttcacaccgcatatggtgcactctcagtacaat
ctgctctgatgccgcatagttaagccagccccgacacccgccaacacccgctgacgcgccctgacgggcttgtctgctcccggcat
ccgcttacagacaagctgtgaccgtctccgggagctgcatgtgtcagaggttttcaccgtcatcaccgaaacgcgcgagacgaaag
ggcctcgtgatacgcctatttttataggttaatgtcatgataataatggtttcttagacgtcaggtggcacttttcggggaaatgt
gcgcggaacccctatttgtttatttttctaaatacattcaaatatgtatccgctcatgagacaataaccctgataaatgcttcaat
aatattgaaaaaggaagagtatgagtattcaacatttccgtgtcgcccttattcccttttttgcggcattttgccttcctgttttt
gctcacccagaaacgctggtgaaagtaaaagatgctgaagatcagttgggtgcacgagtgggttacatcgaactggatctcaacag
cggtaagatccttgagagttttcgccccgaagaacgttttccaatgatgagcacttttaaagttctgctatgtggcgcggtattat
cccgtattgacgccgggcaagagcaactcggtcgccgcatacactattctcagaatgacttggttgagtactcaccagtcacagaa
aagcatcttacggatggcatgacagtaagagaattatgcagtgctgccataaccatgagtgataacactgcggccaacttacttct
gacaacgatcggaggaccgaaggagctaaccgcttttttgcacaacatgggggatcatgtaactcgccttgatcgttgggaaccgg
agctgaatgaagccataccaaacgacgagcgtgacaccacgatgcctgtagcaatggcaacaacgttgcgcaaactattaactggc
gaactacttactctagcttcccggcaacaattaatagactggatggaggcggataaagttgcaggaccacttctgcgctcggccct
tccggctggctggtttattgctgataaatctggagccggtgagcgtgggtctcgcggtatcattgcagcactggggccagatggta
agccctcccgtatcgtagttatctacacgacggggagtcaggcaactatggatgaacgaaatagacagatcgctgagataggtgcc
tcactgattaagcattggtaactgtcagaccaagtttactcatatatactttagattgatttaaaacttcatttttaatttaaaag
gatctaggtgaagatcctttttgataatctcatgaccaaaatcccttaacgtgagttttcgttccactgagcgtcagaccccgtag
aaaagatcaaaggatcttcttgagatcctttttttctgcgcgtaatctgctgcttgcaaacaaaaaaaccaccgctaccagcggtg
gtttgtttgccggatcaagagctaccaactctttttccgaaggtaactggcttcagcagagcgcagataccaaatactgttcttct
agtgtagccgtagttaggccaccacttcaagaactctgtagcaccgcctacatacctcgctctgctaatcctgttaccagtggctg
ctgccagtggcgataagtcgtgtcttaccgggttggactcaagacgatagttaccggataaggcgcagcggtcgggctgaacgggg
ggttcgtgcacacagcccagcttggagcgaacgacctacaccgaact

**Rhizarians**

*Amorphochlora (Lotharella) amoebiformis*

pLaRGfp

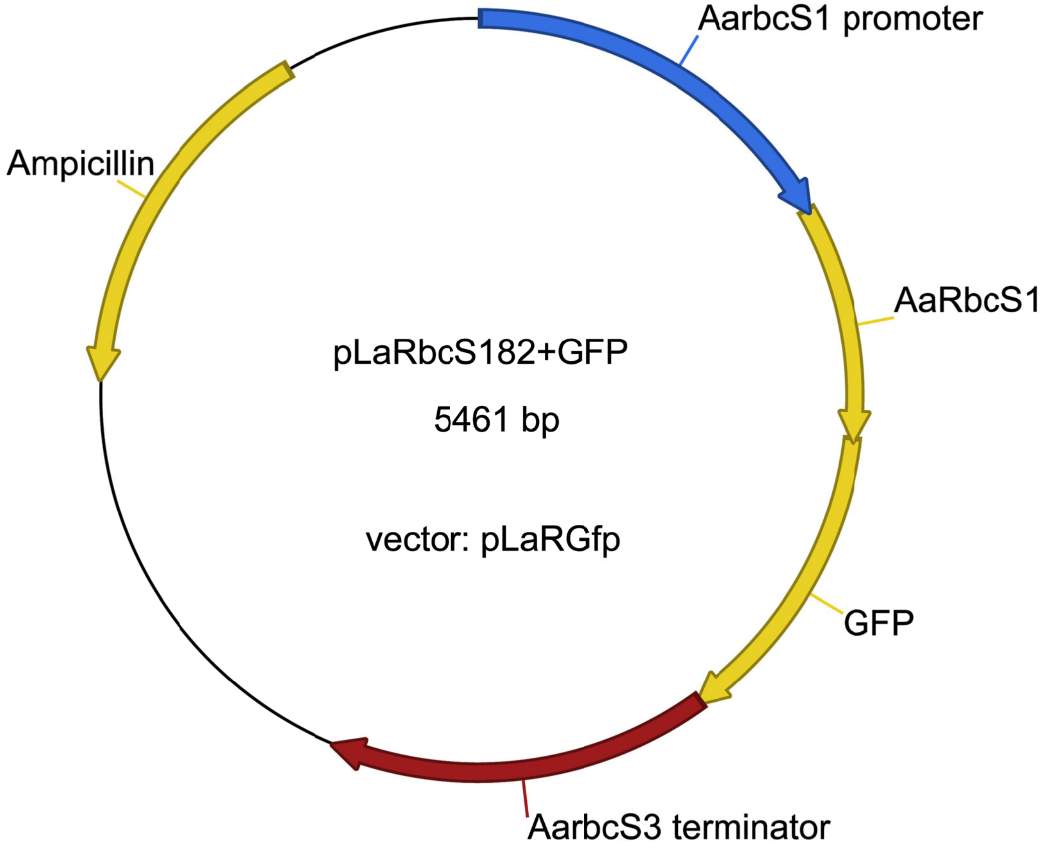


```
>pLaRbcS182+GFP
GAACTCGAGCAGCTGCACACATTAACATTCATGTGGATAATAAATATAATATATATATATATATATATATTATATATATATATATATA
CAAGCCCAAAATATTTATATATGTGTTTTAATGGAATCATGGGTCACATACATATACAATTGGTACGTGTATTTACAATTCCTTAC
ACCCCCCAAAACAACCCCCCCCCTTCTCCACAAAACGACATTTAATCGGTCAGATTCTAGTAATGATAAATTCGCAGCGCGTGGC
GGCATGGGTACCTCACTAAAAACACACACCCTCAATCTATCCCCCCCGCCCCTATCGCACAATTTCTTCGCTTCATGTCCACCTCC
TTTCGATTAGAACCCGCCCCCCCACTTTTAGGAGGGGGGGTACCTCCACGGGGTCCAGACAAGGAGGGAGAAAAGGGCCGCGTAAC
CCGTCGTTGGAAATACACGCATTCTGATAGGACTACTCGTTGGTCAAACCCATCAGATAACGGCCCGGAGGTGACGTTTTGGACAC
GTGTACACTATGTGACATAACGTTAAGTCACAGAAGAATACGGTTACATTGCGCAACGTAACCGTGATACACCATAACATTACGCT
ATGAGTTAACCGAAGAATTAAGTTCTTGGCCGGTGCGGCTTCTTGGAACATACGCACACACCGGCTGGCGGCGCCAACGCGCGATA
GCGTGGAGTATTTGAAGCAAAATAAAGTACGTCAAGAGTGAAATATATATATGGTCGACGTACGGAGGAAGTTTGGAAGCTAGAAG
GTTAGAAATGAGGGCAGGCGAACGCACCGAGCGCTCTGCCGCGTGGGTTGACAGTAGCCGCTTTATTAAATTTATATACCGTCCCA
GTTGCCGCCGGCTCGACCACCCGGCAACGCGGCAGAATCCAAGCAGACAGATACGTAAATAAGCTTATGAAGCTCGCCTTGTCCGC
CAGCATCGTCGTCAACGTTGCCCTTTGCACCGCCCTCGTGTGCCTTGTCACCTTCGGATCCTCCAACGTCGGTGCCCCCGTCGCCG
GCCGCACCATGCAGTTCTCCAGGACTGTGACCCCCGCCCCCACTTTCCGCTCCGCGGTTGCTCAGGCCTACTGTGAGGAGGTTGTT
GACCCGACTGCCAACAAGCGTTACGAGACCATGTCTTACATGCCGGACTTCTCCGAGCGTGACCAGGAGGCCCAGATCAACTACAT
GATCCAGAACGGTCTCATCCCCTGTGTTGAGTTCGACAAGAACGGTGACATGTTCCGTGAGAACTCCAAGATGCCTGGTTACTACG
ATGGCCGTTACTGGACTATGTGGAAGCTCCCCATGTTCGGTGCCACCGACGCTCGCTCTGTTATGGCTGAGGTTAAGGAGTGCCAG
AAGTACAACCCTTCCTCTTACATCCGTATCCTCGGATTCGACAACATCAAGCAGGTTCAGTGCATGGGTTTCCTCGTCGCTAAGCC
TGGTGGAGCCATGGTGAGCAAGGGCGAGGAGCTGTTCACCGGGGTGGTGCCCATCCTGGTCGAGCTGGACGGCGACGTAAACGGCC
ACAAGTTCAGCGTGTCCGGCGAGGGCGAGGGCGATGCCACCTACGGCAAGCTGACCCTGAAGTTCATCTGCACCACCGGCAAGCTG
CCCGTGCCCTGGCCCACCCTCGTGACCACCCTGACCTACGGCGTGCAGTGCTTCAGCCGCTACCCCGACCACATGAAGCAGCACGA
CTTCTTCAAGTCCGCCATGCCCGAAGGCTACGTCCAGGAGCGCACCATCTTCTTCAAGGACGACGGCAACTACAAGACCCGCGCCG
AGGTGAAGTTCGAGGGCGACACCCTGGTGAACCGCATCGAGCTGAAGGGCATCGACTTCAAGGAGGACGGCAACATCCTGGGGCAC
AAGCTGGAGTACAACTACAACAGCCACAACGTCTATATCATGGCCGACAAGCAGAAGAACGGCATCAAGGTGAACTTCAAGATCCG
CCACAACATCGAGGACGGCAGCGTGCAGCTCGCCGACCACTACCAGCAGAACACCCCCATCGGCGACGGCCCCGTGCTGCTGCCCG
ACAACCACTACCTGAGCACCCAGTCCGCCCTGAGCAAAGACCCCAACGAGAAGCGCGATCACATGGTCCTGCTGGAGTTCGTGACC
GCCGCCGGGATCACTCTCGGCATGGACGAGCTGTACAAGTAAGAATTCACTTGCTCACTTGGCACGTGTGTCTTTCCCCGCGTCAA
TGGCTGTCATTAGGCGGATCATTGAGAAACTAAGTGAGACTATGTTGTCTCGATTAGAAAGGAGACACTTACAGTAGGAGCGAATT
```

TCTTTTTAAGAATTAGGCCGACTGGTATTGTATGTTCCCAAATTTCGTCTCAGCGCAACCTGTTCGGAGCTGCTGAATGCTCGTGT
TCAATTTTTATAATAATATGTAATGTTGGCACATTTTACACTACATATCGGATCGGATCGGATGATCATCAATTAGCTGATATATT
AACTTACATATTGTAATGTTTTGACTAGTACATAATTGAGTATTAGTCGGAATTAATGACTCAGACCAGTCAAGAGTCAGATCGAG
ATGTTACTATCATTTTAGTCATCAAAAACACATGTCAGATATTAGAACAGTATTTGCAACTCGCGTCTAAACGTACAACACTAGAA
TAATAGTCTAGATAGTAGAACACACATACGTACCTGATACATATTTTTATATCATACATGTATATATATACACACCGCACATATAC
TAGATATATATATATATATATTTCTCGATGAGTTAGTTATATATGTATGATATCACTATGATATATCCTACTTACACATGTCTAGTTA
TGACTATGTTATTGATTTATGATTCAAGATTTAAACGTTCAGGCGGCCGAGTGTAAATTAAATTTTAAGTTTATCGCCTGTCGCGA
TCATTGCTGTGATCATTCCATGCACACCCCACCCCTGCATGAATCGAGTTAGACACAGACTGGCTCTTTCTAGCGGCTGATAAACG
ACCTCTACTAATTTGAATTAACGTCGGCAAGACTACGTATTATGACTAGCCAGAGGCTTGGGATCGATCCATCAGATCTGCCGGTC
TCCCTATAGTGAGTCGTATTAATTTCGATAAGCCAGGTTAACCTGCATTAATGAATCGGCCAACGCGCGGGGAGAGGCGGTTTGCG
TATTGGGCGCTCTTCCGCTTCCTCGCTCACTGACTCGCTGCGCTCGGTCGTTCGGCTGCGGCGAGCGGTATCAGCTCACTCAAAGG
CGGTAATACGGTTATCCACAGAATCAGGGGATAACGCAGGAAAGAACATGTGAGCAAAAGGCCAGCAAAAGGCCAGGAACCGTAAA
AAGGCCGCGTTGCTGGCGTTTTTCCATAGGCTCCGCCCCCCTGACGAGCATCACAAAAATCGACGCTCAAGTCAGAGGTGGCGAAA
CCCGACAGGACTATAAAGATACCAGGCGTTTCCCCCTGGAAGCTCCCTCGTGCGCTCTCCTGTTCCGACCCTGCCGCTTACCGGAT
ACCTGTCCGCCTTTCTCCCTTCGGGAAGCGTGGCGCTTTCTCAATGCTCACGCTGTAGGTATCTCAGTTCGGTGTAGGTCGTTCGC
TCCAAGCTGGGCTGTGTGCACGAACCCCCCGTTCAGCCCGACCGCTGCGCCTTATCCGGTAACTATCGTCTTGAGTCCAACCCGGT
AAGACACGACTTATCGCCACTGGCAGCAGCCACTGGTAACAGGATTAGCAGAGCGAGGTATGTAGGCGGTGCTACAGAGTTCTTGA
AGTGGTGGCCTAACTACGGCTACACTAGAAGGACAGTATTTGGTATCTGCGCTCTGCTGAAGCCAGTTACCTTCGGAAAAAGAGTT
GGTAGCTCTTGATCCGGCAAACAAACCACCGCTGGTAGCGGTGGTTTTTTTGTTTGCAAGCAGCAGATTACGCGCAGAAAAAAAGG
ATCTCAAGAAGATCCTTTGATCTTTTCTACGGGGTCTGACGCTCAGTGGAACGAAAACTCACGTTAAGGGATTTTGGTCATGAGAT
TATCAAAAAGGATCTTCACCTAGATCCTTTTAAATTAAAAATGAAGTTTTAAATCAATCTAAAGTATATATGAGTAAACTTGGTCT
GACAGTTACCAATGCTTAATCAGTGAGGCACCTATCTCAGCGATCTGTCTATTTCGTTCATCCATAGTTGCCTGACTCCCCGTCGT
GTAGATAACTACGATACGGGAGGGCTTACCATCTGGCCCCAGTGCTGCAATGATACCGCGAGACCCACGCTCACCGGCTCCAGATT
TATCAGCAATAAACCAGCCAGCCGGAAGGGCCGAGCGCAGAAGTGGTCCTGCAACTTTATCCGCCTCCATCCAGTCTATTAATTGT
TGCCGGGAAGCTAGAGTAAGTAGTTCGCCAGTTAATAGTTTGCGCAACGTTGTTGCCATTGCTACAGGCATCGTGGTGTCACGCTC
GTCGTTTGGTATGGCTTCATTCAGCTCCGGTTCCCAACGATCAAGGCGAGTTACATGATCCCCCATGTTGTGCAAAAAAGCGGTTA
GCTCCTTCGGTCCTCCGATCGTTGTCAGAAGTAAGTTGGCCGCAGTGTTATCACTCATGGTTATGGCAGCACTGCATAATTCTCTT
ACTGTCATGCCATCCGTAAGATGCTTTTCTGTGACTGGTGAGTACTCAACCAAGTCATTCTGAGAATAGTGTATGCGGCGACCGAG
TTGCTCTTGCCCGGCGTCAATACGGGATAATACCGCGCCACATAGCAGAACTTTAAAAGTGCTCATCATTGGAAAACGTTCTTCGG
GGCGAAAACTCTCAAGGATCTTACCGCTGTTGAGATCCAGTTCGATGTAACCCACTCGTGCACCCAACTGATCTTCAGCATCTTTT
ACTTTCACCAGCGTTTCTGGGTGAGCAAAAACAGGAAGGCAAAATGCCGCAAAAAAGGGAATAAGGGCGACACGGAAATGTTGAAT
ACTCATACTCTTCCTTTTTCAATATTATTGAAGCATTTATCAGGGTTATTGTCTCATGAGCGGATACATATTTGAATGTATTTAGA
AAAATAAACAAATAGGGGTTCCGCGCACATTTCCCCGAAAAGTGCCACCTGACGTCTAAGAAACCATTATTATCATGACATTAACC
TATAAAAATAGGCGTATCACGAGGCCCTTTCGTCTCGCGCGTTTCGGTGATGACGGTGAAAACCTCTGACACATGCAGCTCCCGGA
GACGGTCACAGCTTGTCTGTAAGCGGATGCCGGGAGCAGACAAGCCCGTCAGGGCGCGTCAGCGGGTGTTGGCGGGTGTCGGGGCT
GGCTTAACTATGCGGCATCAGAGCAGATTGTACTGAGAGTGCACCATATGGACATATTGTCGTTAGAACGCGGCTACAATTAATAC
ATAACCTTATGTATCATACACATACGATTTAGGTGACACTATA

#### *Bigelowiella natans*

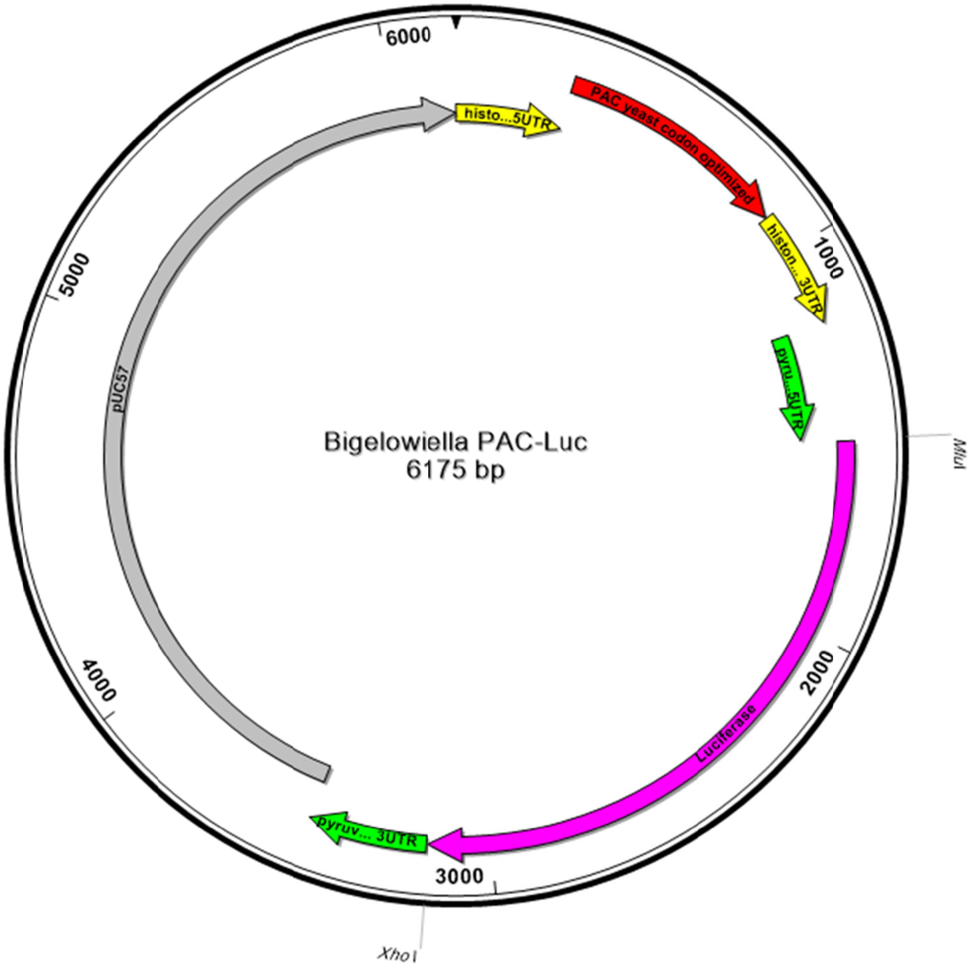


```
>Bigelowiella PAC-Luc
AGGTCATTTTAGACCAAAAATTGGTGTGAACTCCGCTAAATTATTTCGTTAATTAATTTTACGGAATAGCAATAAGGCTTCTGGAA
ATGATTTGTTAAGACAAAGCTTCCATGTTTTATAGGCTAATTTTTGTTTTTCCCCAGACAATATGATGTCGTATAGTATAGATATG
AATTTATTTTTTAATTATTTGCAGAAATAAGTATGAGTTTTTACTAACGTACGTTCAAGCTCCGATTGTGGTGAAATGAGTTGAAG
CAAAAGGAATCATAACTCAACACAACAAGTTACTAAACAAACATGACCGAACATAAGCCATCTGTTAGATTGGCTACTAGAGATGA
TGTTCCAAGAGCTGTTAGAACTTTGGCTGCTGCTTTTGCTGATTATCCAGCTACTAGACATACTGTTGATCCAGATAGACACATCG
AAAGAGTTACCGAATTGCAAGAATTATTCTTAACCAGAGTCGGTTTGGATATCGGTAAAGTTTGGGTTGCTGATGATGGTGCTGCT
GTTGCAGTTTGGACTACTCCAGAATCTTTGGATGCTGGTGCTGTTTTCGCTGAAATTGGTCCAAGAATGGCTGAATTGTCTGGTTC
TAGATTAGCTGCTCAACAACAAATGGAAGGTTTGTTGGCTCCACATAGACCAAAAGAACCAGCTTGGTTTTTGGCTACTGTTGGTG
TTTCTCCAGATCATCAAGGTAAAGGTTTAGGTTCTGCTGTTGTTTTGCCAGGTGTTGAAGCTGCTGAAAGAGCTGGTGTTCCAGCT
TTTTTGGAAACTTCTGCTCCAAGAAATTTGCCATTCTACGAAAGATTGGGTTTCACTGTTACTGCCGATGTTGAAGTTCCTGAAGG
TCCTAGAACTTGGTGTATGACTAGAAAACCAGGTGCTTAAATTGCTCACAGCATTCTCAATGCTACTACATCCGGTGTTCTTAAAC
ACCACCATTCTCTTCGGAAACTGTGCCTTAGTCAAAAATTGTAAACCGATAGAAAATAAATAGAAAATGATGTTATCTAGTGCATA
AGTAGCGAGTTGGAATACGTTCCTACGTCTCCTTGTCAACACCACGATGTAATTCAAGTGATTAGCCATGCTTGATGGCAGATGAA
CAAAGGGAATTCATATTGTACATGATTAAGCGTAGTGGAGCCAGTGAAGAACTCCCACACCCGCCCATCCTAGCTAGTGAACTTTT
TGTTTCTTCATAGCACTGGTCTATCTGATGTTCACGTTTCATTTCAACAAACATTCGCTTGTCGCAGGAGGATTGTAGACTAAATT
GTCATTGTTTTTCAGTAGACGGAAAGAGAAAGATATCTGAAACGGCACTGCAATCGGCTACGCGCACACGGTGAAATGCTACGTAT
AAGGATTTTGATGTTTTTGCCTGTCCGACAAGGAAGTCATTGTTGTCGCAGTCCATAAACGTCGGAATTCATTATGGTTTTTGATT
AGTGGGTTACCCCACCAGACCCCAGAACAACACACATAACGCGTATGGAAGACGCCAAAAACATAAAGAAAGGCCCGGCGCCATTC
TATCCGCTGGAAGATGGAACCGCTGGAGAGCAACTGCATAAGGCTATGAAGAGATACGCCCTGGTTCCTGGAACAATTGCTTTTAC
AGATGCACATATCGAGGTGGACATCACTTACGCTGAGTACTTCGAAATGTCCGTTCGGTTGGCAGAAGCTATGAAACGATATGGGC
TGAATACAAATCACAGAATCGTCGTATGCAGTGAAAACTCTCTTCAATTCTTTATGCCGGTGTTGGGCGCGTTATTTATCGGAGTT
GCAGTTGCGCCCGCGAACGACATTTATAATGAACGTGAATTGCTCAACAGTATGGGCATTTCGCAGCCTACCGTGGTGTTCGTTTC
CAAAAAGGGGTTGCAAAAAATTTTGAACGTGCAAAAAAAGCTCCCAATCATCCAAAAAATTATTATCATGGATTCTAAAACGGATT
```

ACCAGGGATTTCAGTCGATGTACACGTTCGTCACATCTCATCTACCTCCCGGTTTTAATGAATACGATTTTGTGCCAGAGTCCTTC
GATAGGGACAAGACAATTGCACTGATCATGAACTCCTCTGGATCTACTGGTCTGCCTAAAGGTGTCGCTCTGCCTCATAGAACTGC
CTGCGTGAGATTCTCGCATGCCAGAGATCCTATTTTTGGCAATCAAATCATTCCGGATACTGCGATTTTAAGTGTTGTTCCATTCC
ATCACGGTTTTGGAATGTTTACTACACTCGGATATTTGATATGTGGATTTCGAGTCGTCTTAATGTATAGATTTGAAGAAGAGCTG
TTTCTGAGGAGCCTTCAGGATTACAAGATTCAAAGTGCGCTGCTGGTGCCAACCCTATTCTCCTTCTTCGCCAAAAGCACTCTGAT
TGACAAATACGATTTATCTAATTTACACGAAATTGCTTCTGGTGGCGCTCCCCTCTCTAAGGAAGTCGGGGAAGCGGTTGCCAAGA
GGTTCCATCTGCCAGGTATCAGGCAAGGATATGGGCTCACTGAGACTACATCAGCTATTCTGATTACACCCGAGGGGGATGATAAA
CCGGGCGCGGTCGGTAAAGTTGTTCCATTTTTTGAAGCGAAGGTTGTGGATCTGGATACCGGGAAAACGCTGGGCGTTAATCAAAG
AGGCGAACTGTGTGTGAGAGGTCCTATGATTATGTCCGGTTATGTAAACAATCCGGAAGCGACCAACGCCTTGATTGACAAGGATG
GATGGCTACATTCTGGAGACATAGCTTACTGGGACGAAGACGAACACTTCTTCATCGTTGACCGCCTGAAGTCTCTGATTAAGTAC
AAAGGCTATCAGGTGGCTCCCGCTGAATTGGAATCCATCTTGCTCCAACACCCCAACATCTTCGACGCAGGTGTCGCAGGTCTTCC
CGACGATGACGCCGGTGAACTTCCCGCCGCCGTTGTTGTTTTGGAGCACGGAAAGACGATGACGGAAAAAGAGATCGTGGATTACG
TCGCCAGTCAAGTAACAACCGCGAAAAAGTTGCGCGGAGGAGTTGTGTTTGTGGACGAAGTACCGAAAGGTCTTACCGGAAAACTC
GACGCAAGAAAAATCAGAGAGATCCTCATAAAGGCCAAGAAGGGCGGAAAGATCGCCGTGTAACTCGAGGGTGAACAAACAAACGA
CACATCTAAGTAGTGCAAGAATGGCTTCTCAAGACTCAAGTGACAACCCTAGTGTAGTGTTCGCACTTTGATGATGATGATTCGAT
TAGCAGCAAGATACATTTTTGTTAATAGCTTTTGCATTTGGATGCCAATTAGTGTTAAAAAAAACCGATCTAATTCACTTGTAATA
ACGAAGAAGCCAGTGTCATTAAGCCAAGTTTAGGTGCTCAAGCCGTGTGAACAAGGGACAATTTAATGAGGCCAGGAGTAGAGCAA
TGCAACAGAGTTTGCATCATCAAAAATCGGATCCCGGGCCCGTCGACTGCAGAGGCCTGCATGCAAGCTTGGCGTAATCATGGTCA
TAGCTGTTTCCTGTGTGAAATTGTTATCCGCTCACAATTCCACACAACATACGAGCCGGAAGCATAAAGTGTAAAGCCTGGGGTGC
CTAATGAGTGAGCTAACTCACATTAATTGCGTTGCGCTCACTGCCCGCTTTCCAGTCGGGAAACCTGTCGTGCCAGCTGCATTAAT
GAATCGGCCAACGCGCGGGGAGAGGCGGTTTGCGTATTGGGCGCTCTTCCGCTTCCTCGCTCACTGACTCGCTGCGCTCGGTCGTT
CGGCTGCGGCGAGCGGTATCAGCTCACTCAAAGGCGGTAATACGGTTATCCACAGAATCAGGGGATAACGCAGGAAAGAACATGTG
AGCAAAAGGCCAGCAAAAGGCCAGGAACCGTAAAAAGGCCGCGTTGCTGGCGTTTTTCCATAGGCTCCGCCCCCCTGACGAGCATC
ACAAAAATCGACGCTCAAGTCAGAGGTGGCGAAACCCGACAGGACTATAAAGATACCAGGCGTTTCCCCCTGGAAGCTCCCTCGTG
CGCTCTCCTGTTCCGACCCTGCCGCTTACCGGATACCTGTCCGCCTTTCTCCCTTCGGGAAGCGTGGCGCTTTCTCATAGCTCACG
CTGTAGGTATCTCAGTTCGGTGTAGGTCGTTCGCTCCAAGCTGGGCTGTGTGCACGAACCCCCCGTTCAGCCCGACCGCTGCGCCT
TATCCGGTAACTATCGTCTTGAGTCCAACCCGGTAAGACACGACTTATCGCCACTGGCAGCAGCCACTGGTAACAGGATTAGCAGA
GCGAGGTATGTAGGCGGTGCTACAGAGTTCTTGAAGTGGTGGCCTAACTACGGCTACACTAGAAGAACAGTATTTGGTATCTGCGC
TCTGCTGAAGCCAGTTACCTTCGGAAAAAGAGTTGGTAGCTCTTGATCCGGCAAACAAACCACCGCTGGTAGCGGTGGTTTTTTTG
TTTGCAAGCAGCAGATTACGCGCAGAAAAAAAGGATCTCAAGAAGATCCTTTGATCTTTTCTACGGGGTCTGACGCTCAGTGGAAC
GAAAACTCACGTTAAGGGATTTTGGTCATGAGATTATCAAAAAGGATCTTCACCTAGATCCTTTTAAATTAAAAATGAAGTTTTAA
ATCAATCTAAAGTATATATGAGTAAACTTGGTCTGACAGTTACCAATGCTTAATCAGTGAGGCACCTATCTCAGCGATCTGTCTAT
TTCGTTCATCCATAGTTGCCTGACTCCCCGTCGTGTAGATAACTACGATACGGGAGGGCTTACCATCTGGCCCCAGTGCTGCAATG
ATACCGCGAGACCCACGCTCACCGGCTCCAGATTTATCAGCAATAAACCAGCCAGCCGGAAGGGCCGAGCGCAGAAGTGGTCCTGC
AACTTTATCCGCCTCCATCCAGTCTATTAATTGTTGCCGGGAAGCTAGAGTAAGTAGTTCGCCAGTTAATAGTTTGCGCAACGTTG
TTGCCATTGCTACAGGCATCGTGGTGTCACGCTCGTCGTTTGGTATGGCTTCATTCAGCTCCGGTTCCCAACGATCAAGGCGAGTT
ACATGATCCCCCATGTTGTGCAAAAAAGCGGTTAGCTCCTTCGGTCCTCCGATCGTTGTCAGAAGTAAGTTGGCCGCAGTGTTATC
ACTCATGGTTATGGCAGCACTGCATAATTCTCTTACTGTCATGCCATCCGTAAGATGCTTTTCTGTGACTGGTGAGTACTCAACCA
AGTCATTCTGAGAATAGTGTATGCGGCGACCGAGTTGCTCTTGCCCGGCGTCAATACGGGATAATACCGCGCCACATAGCAGAACT
TTAAAAGTGCTCATCATTGGAAAACGTTCTTCGGGGCGAAAACTCTCAAGGATCTTACCGCTGTTGAGATCCAGTTCGATGTAACC
CACTCGTGCACCCAACTGATCTTCAGCATCTTTTACTTTCACCAGCGTTTCTGGGTGAGCAAAAACAGGAAGGCAAAATGCCGCAA
AAAAGGGAATAAGGGCGACACGGAAATGTTGAATACTCATACTCTTCCTTTTTCAATATTATTGAAGCATTTATCAGGGTTATTGT
CTCATGAGCGGATACATATTTGAATGTATTTAGAAAAATAAACAAATAGGGGTTCCGCGCACATTTCCCCGAAAAGTGCCACCTGA
CGTCTAAGAAACCATTATTATCATGACATTAACCTATAAAAATAGGCGTATCACGAGGCCCTTTCGTCTCGCGCGTTTCGGTGATG
ACGGTGAAAACCTCTGACACATGCAGCTCCCGGAGACGGTCACAGCTTGTCTGTAAGCGGATGCCGGGAGCAGACAAGCCCGTCAG
GGCGCGTCAGCGGGTGTTGGCGGGTGTCGGGGCTGGCTTAACTATGCGGCATCAGAGCAGATTGTACTGAGAGTGCACCATATGCG
GTGTGAAATACCGCACAGATGCGTAAGGAGAAAATACCGCATCAGGCGCCATTCGCCATTCAGGCTGCGCAACTGTTGGGAAGGGC
GATCGGTGCGGGCCTCTTCGCTATTACGCCAGCTGGCGAAAGGGGGATGTGCTGCAAGGCGATTAAGTTGGGTAACGCCAGGGTTT
TCCCAGTCACGACGTTGTAAAACGACGGCCAGTGAATTCGAGCTCGGTACCTCGCGAATGCATCTAGAT

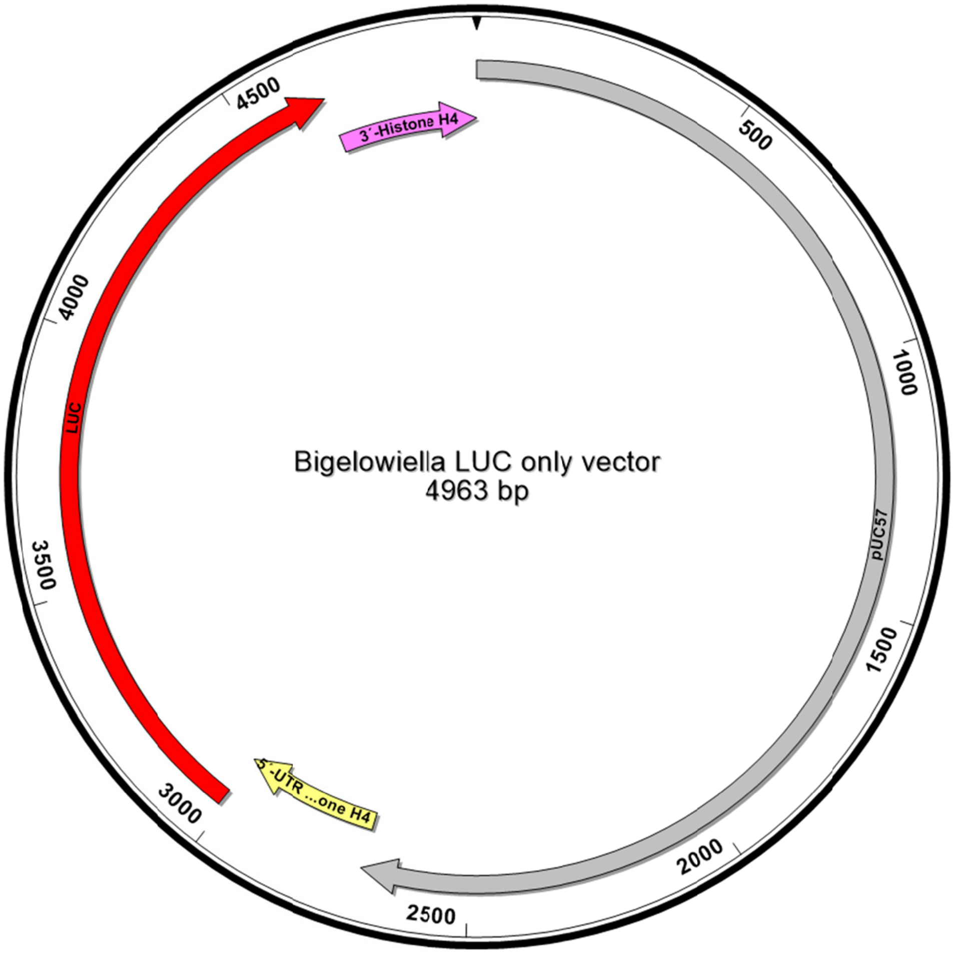


1-2270 pUC57 backbone
2771-3010 5´- UTR Histone H4
3011-4663 Luciferase
4664-4963 3´-UTR histone H4

ATCGGATCCCGGGCCCGTCGACTGCAGAGGCCTGCATGCAAGCTTGGCGTAATCATGGTCATAGCTGTTTCCTGTGTGAAATTGTT
ATCCGCTCACAATTCCACACAACATACGAGCCGGAAGCATAAAGTGTAAAGCCTGGGGTGCCTAATGAGTGAGCTAACTCACATTA
ATTGCGTTGCGCTCACTGCCCGCTTTCCAGTCGGGAAACCTGTCGTGCCAGCTGCATTAATGAATCGGCCAACGCGCGGGGAGAGG
CGGTTTGCGTATTGGGCGCTCTTCCGCTTCCTCGCTCACTGACTCGCTGCGCTCGGTCGTTCGGCTGCGGCGAGCGGTATCAGCTC
ACTCAAAGGCGGTAATACGGTTATCCACAGAATCAGGGGATAACGCAGGAAAGAACATGTGAGCAAAAGGCCAGCAAAAGGCCAGG
AACCGTAAAAAGGCCGCGTTGCTGGCGTTTTTCCATAGGCTCCGCCCCCCTGACGAGCATCACAAAAATCGACGCTCAAGTCAGAG
GTGGCGAAACCCGACAGGACTATAAAGATACCAGGCGTTTCCCCCTGGAAGCTCCCTCGTGCGCTCTCCTGTTCCGACCCTGCCGC
TTACCGGATACCTGTCCGCCTTTCTCCCTTCGGGAAGCGTGGCGCTTTCTCATAGCTCACGCTGTAGGTATCTCAGTTCGGTGTAG
GTCGTTCGCTCCAAGCTGGGCTGTGTGCACGAACCCCCCGTTCAGCCCGACCGCTGCGCCTTATCCGGTAACTATCGTCTTGAGTC
CAACCCGGTAAGACACGACTTATCGCCACTGGCAGCAGCCACTGGTAACAGGATTAGCAGAGCGAGGTATGTAGGCGGTGCTACAG
AGTTCTTGAAGTGGTGGCCTAACTACGGCTACACTAGAAGAACAGTATTTGGTATCTGCGCTCTGCTGAAGCCAGTTACCTTCGGA
AAAAGAGTTGGTAGCTCTTGATCCGGCAAACAAACCACCGCTGGTAGCGGTGGTTTTTTTGTTTGCAAGCAGCAGATTACGCGCAG
AAAAAAAGGATCTCAAGAAGATCCTTTGATCTTTTCTACGGGGTCTGACGCTCAGTGGAACGAAAACTCACGTTAAGGGATTTTGG
TCATGAGATTATCAAAAAGGATCTTCACCTAGATCCTTTTAAATTAAAAATGAAGTTTTAAATCAATCTAAAGTATATATGAGTAA
ACTTGGTCTGACAGTTACCAATGCTTAATCAGTGAGGCACCTATCTCAGCGATCTGTCTATTTCGTTCATCCATAGTTGCCTGACT
CCCCGTCGTGTAGATAACTACGATACGGGAGGGCTTACCATCTGGCCCCAGTGCTGCAATGATACCGCGAGACCCACGCTCACCGG
CTCCAGATTTATCAGCAATAAACCAGCCAGCCGGAAGGGCCGAGCGCAGAAGTGGTCCTGCAACTTTATCCGCCTCCATCCAGTCT
ATTAATTGTTGCCGGGAAGCTAGAGTAAGTAGTTCGCCAGTTAATAGTTTGCGCAACGTTGTTGCCATTGCTACAGGCATCGTGGT
GTCACGCTCGTCGTTTGGTATGGCTTCATTCAGCTCCGGTTCCCAACGATCAAGGCGAGTTACATGATCCCCCATGTTGTGCAAAA
AAGCGGTTAGCTCCTTCGGTCCTCCGATCGTTGTCAGAAGTAAGTTGGCCGCAGTGTTATCACTCATGGTTATGGCAGCACTGCAT
AATTCTCTTACTGTCATGCCATCCGTAAGATGCTTTTCTGTGACTGGTGAGTACTCAACCAAGTCATTCTGAGAATAGTGTATGCG
GCGACCGAGTTGCTCTTGCCCGGCGTCAATACGGGATAATACCGCGCCACATAGCAGAACTTTAAAAGTGCTCATCATTGGAAAAC
GTTCTTCGGGGCGAAAACTCTCAAGGATCTTACCGCTGTTGAGATCCAGTTCGATGTAACCCACTCGTGCACCCAACTGATCTTCA
GCATCTTTTACTTTCACCAGCGTTTCTGGGTGAGCAAAAACAGGAAGGCAAAATGCCGCAAAAAAGGGAATAAGGGCGACACGGAA
ATGTTGAATACTCATACTCTTCCTTTTTCAATATTATTGAAGCATTTATCAGGGTTATTGTCTCATGAGCGGATACATATTTGAAT
GTATTTAGAAAAATAAACAAATAGGGGTTCCGCGCACATTTCCCCGAAAAGTGCCACCTGACGTCTAAGAAACCATTATTATCATG

ACATTAACCTATAAAAATAGGCGTATCACGAGGCCCTTTCGTCTCGCGCGTTTCGGTGATGACGGTGAAAACCTCTGACACATGCA
GCTCCCGGAGACGGTCACAGCTTGTCTGTAAGCGGATGCCGGGAGCAGACAAGCCCGTCAGGGCGCGTCAGCGGGTGTTGGCGGGT
GTCGGGGCTGGCTTAACTATGCGGCATCAGAGCAGATTGTACTGAGAGTGCACCATATGCGGTGTGAAATACCGCACAGATGCGTA
AGGAGAAAATACCGCATCAGGCGCCATTCGCCATTCAGGCTGCGCAACTGTTGGGAAGGGCGATCGGTGCGGGCCTCTTCGCTATT
ACGCCAGCTGGCGAAAGGGGGATGTGCTGCAAGGCGATTAAGTTGGGTAACGCCAGGGTTTTCCCAGTCACGACGTTGTAAAACGA
CGGCCAGTGAATTCGAGCTCGGTACCTCGCGAATGCATCTAGATAGGTCATTTTAGACCAAAAATTGGTGTGAACTCCGCTAAATT
ATTTCGTTAATTAATTTTACGGAATAGCAATAAGGCTTCTGGAAATGATTTGTTAAGACAAAGCTTCCATGTTTTATAGGCTAATT
TTTGTTTTTCCCCAGACAATATGATGTCGTATAGTATAGATATGAATTTATTTTTTAATTATTTGCAGAAATAAGTATGAGTTTTT
ACTAACGTACGTTCAAGCTCCGATTGTGGTGAAATGAGTTGAAGCAAAAGGAATCATAACTCAACACAACAAGTTACTAAACAAAC
ATGGAAGACGCCAAAAACATAAAGAAAGGCCCGGCGCCATTCTATCCGCTGGAAGATGGAACCGCTGGAGAGCAACTGCATAAGGC
TATGAAGAGATACGCCCTGGTTCCTGGAACAATTGCTTTTACAGATGCACATATCGAGGTGGACATCACTTACGCTGAGTACTTCG
AAATGTCCGTTCGGTTGGCAGAAGCTATGAAACGATATGGGCTGAATACAAATCACAGAATCGTCGTATGCAGTGAAAACTCTCTT
CAATTCTTTATGCCGGTGTTGGGCGCGTTATTTATCGGAGTTGCAGTTGCGCCCGCGAACGACATTTATAATGAACGTGAATTGCT
CAACAGTATGGGCATTTCGCAGCCTACCGTGGTGTTCGTTTCCAAAAAGGGGTTGCAAAAAATTTTGAACGTGCAAAAAAAGCTCC
CAATCATCCAAAAAATTATTATCATGGATTCTAAAACGGATTACCAGGGATTTCAGTCGATGTACACGTTCGTCACATCTCATCTA
CCTCCCGGTTTTAATGAATACGATTTTGTGCCAGAGTCCTTCGATAGGGACAAGACAATTGCACTGATCATGAACTCCTCTGGATC
TACTGGTCTGCCTAAAGGTGTCGCTCTGCCTCATAGAACTGCCTGCGTGAGATTCTCGCATGCCAGAGATCCTATTTTTGGCAATC
AAATCATTCCGGATACTGCGATTTTAAGTGTTGTTCCATTCCATCACGGTTTTGGAATGTTTACTACACTCGGATATTTGATATGT
GGATTTCGAGTCGTCTTAATGTATAGATTTGAAGAAGAGCTGTTTCTGAGGAGCCTTCAGGATTACAAGATTCAAAGTGCGCTGCT
GGTGCCAACCCTATTCTCCTTCTTCGCCAAAAGCACTCTGATTGACAAATACGATTTATCTAATTTACACGAAATTGCTTCTGGTG
GCGCTCCCCTCTCTAAGGAAGTCGGGGAAGCGGTTGCCAAGAGGTTCCATCTGCCAGGTATCAGGCAAGGATATGGGCTCACTGAG
ACTACATCAGCTATTCTGATTACACCCGAGGGGGATGATAAACCGGGCGCGGTCGGTAAAGTTGTTCCATTTTTTGAAGCGAAGGT
TGTGGATCTGGATACCGGGAAAACGCTGGGCGTTAATCAAAGAGGCGAACTGTGTGTGAGAGGTCCTATGATTATGTCCGGTTATG
TAAACAATCCGGAAGCGACCAACGCCTTGATTGACAAGGATGGATGGCTACATTCTGGAGACATAGCTTACTGGGACGAAGACGAA
CACTTCTTCATCGTTGACCGCCTGAAGTCTCTGATTAAGTACAAAGGCTATCAGGTGGCTCCCGCTGAATTGGAATCCATCTTGCT
CCAACACCCCAACATCTTCGACGCAGGTGTCGCAGGTCTTCCCGACGATGACGCCGGTGAACTTCCCGCCGCCGTTGTTGTTTTGG
AGCACGGAAAGACGATGACGGAAAAAGAGATCGTGGATTACGTCGCCAGTCAAGTAACAACCGCGAAAAAGTTGCGCGGAGGAGTT
GTGTTTGTGGACGAAGTACCGAAAGGTCTTACCGGAAAACTCGACGCAAGAAAAATCAGAGAGATCCTCATAAAGGCCAAGAAGGG
CGGAAAGATCGCCGTGTAAATTGCTCACAGCATTCTCAATGCTACTACATCCGGTGTTCTTAAACACCACCATTCTCTTCGGAAAC
TGTGCCTTAGTCAAAAATTGTAAACCGATAGAAAATAAATAGAAAATGATGTTATCTAGTGCATAAGTAGCGAGTTGGAATACGTT
CCTACGTCTCCTTGTCAACACCACGATGTAATTCAAGTGATTAGCCATGCTTGATGGCAGATGAACAAAGGGAATTCATATTGTAC
ATGATTAAGCGTAGTGGAGCCAGTGAAGAACTCCCACACCCGCCCATCCTAGCTAGTGAAC

#### **Stramenopiles**

*Fragilariopsis cylindrus*

pUC:FCP:ShBle:FCP:EGFP Available at addgene plasmid #85987

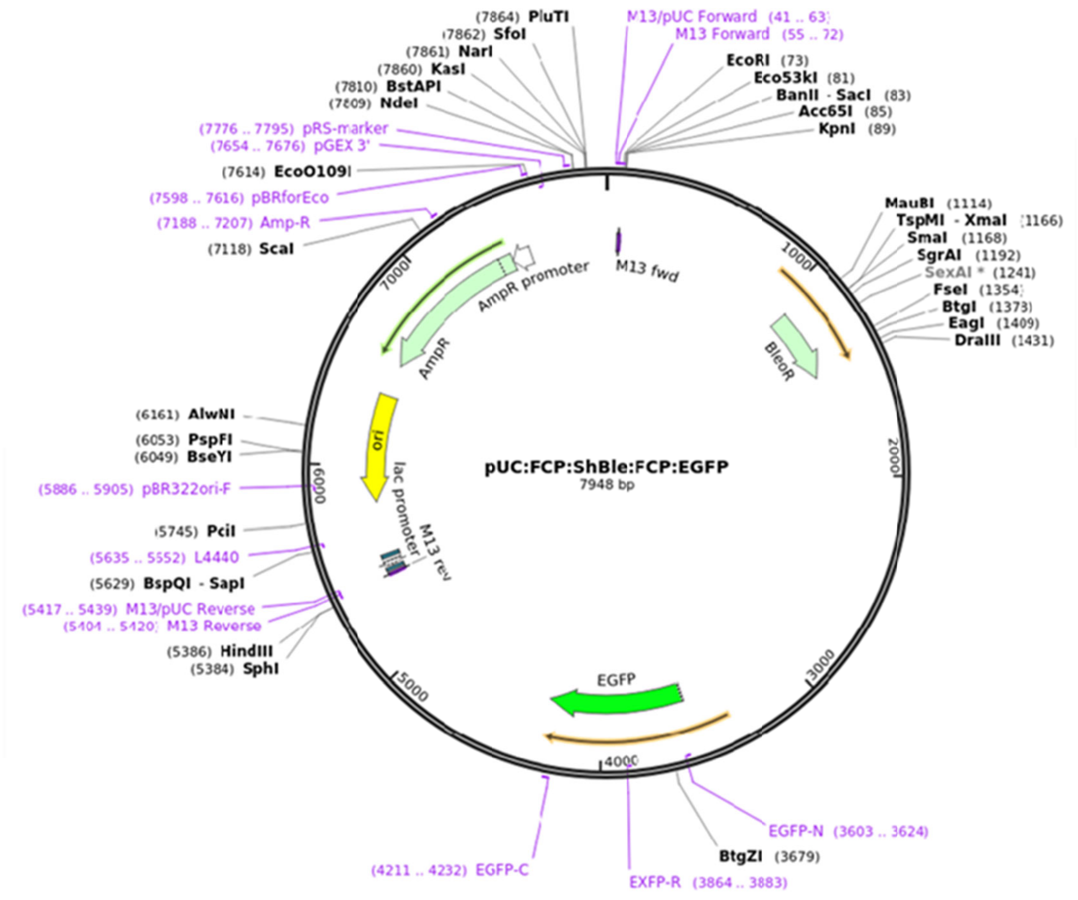


```
> pUC:FCP:ShBle:FCP:EGFP
TGTGCTGCAAGGCGATTAAGTTGGGTAACGCCAGGGTTTTCCCAGTCACGACGTTGTAAAACGACGGCCA
GTGAATTCGAGCTCGGTACCCCAAAGTAAGGCATAGAAATAATCTGCATGAAAAGATCTAACGATTTCTA
TCAGTTCCCCTACAAATGGATGCTTAGGGGTGCTTGTATGAGTGTACGAAAGAGCCAACCAAAAAATAAT
TTTTTATTCTGATTTTTAATTTTTTGTGCTAAAACATGCTGCGCTGTGTACTTCCTCCGTGCACCGGTAA
ACGCGCACGGAGGAAATACAGTGTTTTAGAAAAACAACGCGCAAAAATGCCAATCTTTATTATTTTGAAG
TGAAGCGCGCTATTTACTGTTATCGCGCTAATACTGTAAAAATTCGCGCTGATCTTGATGATACCACCTT
AACTTTGTTTCTATTTTTTGATAACAAATACAAAATCTACTAGTATTGTGCACCCTCCTACTACTAGCTA
CTACTAGGGAGTGTCGTACTAGTAGTCTAAAAATACGCGTGTTCAACATGCGCGTGCTAGAAGTATGATG
CGTGTGTAACAAATTTTTAAAAGATTAAAAAAATCTTTATTTTGAGGGATTGGATGCGTGATAGAATGGA
GAATTGTGTTCGTGGGATTCATTTTGACCTACCAATTAAAACGAATAAAAATGTCAGCGTCGATTATAGA
CACGAAGTACAGACCAGACAGCGCGTCGACGTCAGATTTTTATGGCCAATCAGATTGTTTCCAACACACA
AATGGAAAGAAAGAATGCATTCTGTGGAAGAGTGTACATCCGGGGGAATTTAGAACGAAAACCACGTCGG
TGAATTGTACACGCGACAATCCTCCAATCGTGGATAATCCAGGATTTTTTGTTGATGTTGAACTGCATAT
TTGCCGATCCTTCATGCGAACTCCAAATTTCTATATCATTTTTTCAGCAAAATCCAATATTTTTCAGCAA
AATCCAAAAAACAAGCCATCGTTTTACCGCTTTCGATCTTCTCATAACTTTATTCTTTCAACAGTATAAA
CTACCAAAAACAAACTTATATATCAAAATGGCCAAGTTGACCAGTGCCGTTCCGGTGCTCACCGCGCGCG
ACGTCGCCGGAGCGGTCGAGTTCTGGACCGACCGGCTCGGGTTCTCCCGGGACTTCGTGGAGGACGACTT
CGCCGGTGTGGTCCGGGACGACGTGACCCTGTTCATCAGCGCGGTCCAGGACCAGGTGGTGCCGGACAAC
ACCCTGGCCTGGGTGTGGGTGCGCGGCCTGGACGAGCTGTACGCCGAGTGGTCGGAGGTCGTGTCCACGA
ACTTCCGGGACGCCTCCGGGCCGGCCATGACCGAGATCGGCGAGCAGCCGTGGGGGCGGGAGTTCGCCCT
```

GCGCGACCCGGCCGGCAACTGCGTGCACTTCGTGGCCGAGGAGCAGGACTGAGCATTTTATTAATCCTTA
TTTGATCGATTCATTGATTCGAATATAGTGAGACCCTTCCGTTGACAACTACAATCGATTCCTAATTTAA
TTATTTAGTAGAGTTCCTTGAGTTTTTGTCACAAATATTATTATCTACTTCTATCATATACTACTATTCA
TAGCTAGCTACGCATTGTTTCCTCAATCCATACGCAGTTGTACTGTTACATTCACAACCCGTATCCCTCA
TGACCTCTTGACTGTATCCTACAGTACGTATTTCCTTTATTTTTCTCTTCCACGAGCTACAGTAGCCACA
CCCCTACGGTACGGTACAGTATTGATAAATCAACCGACGACAGCGCCTCTCAACTGCCAAACAGCTGGAA
TGCGTTCGTTTGAGAATGACCCCAAGACAAAAGATCCATTTTCAACCTACAGTAACGTCCGCCCACGTTT
GCAGGATCCAAAAATGCGAGGAATCAATCCAGTCGAACTAGTCGAGAAAGGGCAACAAAGATCCAATTTC
CGTACTAGACAGATGTACAGTATGGAATCGAGTGAATCAAACCAGTCGATTCAGTCGAACTAGTCGAGGA
TTGTAATCAATACTGTACATAGTTTGCGCTTTGCCTTCCACTAAGTTACCGGTACAGGAGTAGTAGATTT
TGTGAGATTTTTCATGACAAACAAATCGAAAAGTAAAAAAAAAACCTATTAGAAATCAACAATGACGACG
ATGATGATGTAACATGATGAGGAGGATAATCATAATAGTGACAATTCAACTATCGTATCACCTTATTTCT
TCACAATCAGCCAAAAGAAAATTCCCTAGAGGTCAGCAGAAAAAAATTACGAAAAGAACGGCGTACTGTAC
AGCAACGGCGAAGGATTGACAAATGAATAGTATTATTAGAATAAAAAGTGTTTGGTTTTCGTTATCGTTA
TATTTATCGGTATTGTTACTATTACTATTTAGTTTGGTTCTATTTTTTAATAGTAATAATAATAATAATA
ATGATACAACAATCATAGTAGTGGGCGTTAATGGATTTCTTCTACCAACAAGAACAACAATAACGCCGAC
ATCAGCACAGCGACTACTGTTGTCGTCTACTACAGCACAACAACAACGACTCAAAGTAAGGCATAGAAAT
AATCTGCATGAAAAGATCTAACGATTTCTATCAGTTCCCCTACAAATGGATGCTTAGGGGTGCTTGTATG
AGTGTACGAAAGAGCCAACCAAAAAAATAATTTTTTATTCTGATTTTTAATTTTTTTGTGCTAAAACATGCT
GCGCTGTGTACTTCCTCCGTGCACCGGTAAACGCGCACGGAGGAAATACAGTGTTTTAGAAAAACAACGC
GCAAAAATGCCAATCTTTATTATTTTGAAGTGAAGCGCGCTATTTACTGTTATCGCGCTAATACTGTAAA
AATTCGCGCTGATCTTGATGATACCACCTTAACTTTGTTTCTATTTTTTGATAACAAATACAAAATCTAC
TAGTATTGTGCACCCTCCTACTACTAGCTACTACTAGGGAGTGTCGTACTAGTAGTCTAAAAATACGCGT
GTTCAACATGCGCGTGCTAGAAGTATGATGCGTGTGTAACAAATTTTTAAAAGATTAAAAAAATCTTTAT
TTTGAGGGATTGGATGCGTGATAGAATGGAGAATTGTGTTCGTGGGATTCATTTTGACCTACCAATTAAA
ACGAATAAAAATGTCAGCGTCGATTATAGACACGAAGTACAGACCAGACAGCGCGTCGACGTCAGATTTT
TATGGCCAATCAGATTGTTTCCAACACACAAATGGAAAGAAAGAATGCATTCTGTGGAAGAGTGTACATC
CGGGGGAATTTAGAACGAAAACCACGTCGGTGAATTGTACACGCGACAATCCTCCAATCGTGGATAATCC
AGGATTTTTTGTTGATGTTGAACTGCATATTTGCCGATCCTTCATGCGAACTCCAAATTTCTATATCATT
TTTTCAGCAAAATCCAATATTTTTCAGCAAAATCCAAAAAACAAGCCATCGTTTTACCGCTTTCGATCTT
CTCATAACTTTATTCTTTCAACAGTATAAACTACCAAAAACAAACTTATATATCAAAATGGTGAGCAAGG
GCGAGGAGCTGTTCACCGGGGTGGTGCCCATCCTGGTCGAGCTGGACGGCGACGTAAACGGCCACAAGTT
CAGCGTGTCCGGCGAGGGCGAGGGCGATGCCACCTACGGCAAGCTGACCCTGAAGTTCATCTGCACCACC
GGCAAGCTGCCCGTGCCCTGGCCCACCCTCGTGACCACCCTGACCTACGGCGTGCAGTGCTTCAGCCGCT
ACCCCGACCACATGAAGCAGCACGACTTCTTCAAGTCCGCCATGCCCGAAGGCTACGTCCAGGAGCGCAC
CATCTTCTTCAAGGACGACGGCAACTACAAGACCCGCGCCGAGGTGAAGTTCGAGGGCGACACCCTGGTG
AACCGCATCGAGCTGAAGGGCATCGACTTCAAGGAGGACGGCAACATCCTGGGGCACAAGCTGGAGTACA
ACTACAACAGCCACAACGTCTATATCATGGCCGACAAGCAGAAGAACGGCATCAAGGTGAACTTCAAGAT
CCGCCACAACATCGAGGACGGCAGCGTGCAGCTCGCCGACCACTACCAGCAGAACACCCCCATCGGCGAC
GGCCCCGTGCTGCTGCCCGACAACCACTACCTGAGCACCCAGTCCGCCCTGAGCAAAGACCCCAACGAGA
AGCGCGATCACATGGTCCTGCTGGAGTTCGTGACCGCCGCCGGGATCACTCTCGGCATGGACGAGCTGTA
CAAGTAAGCATTTTATTAATCCTTATTTGATCGATTCATTGATTCGAATATAGTGAGACCCTTCCGTTGA
CAACTACAATCGATTCCTAATTTAATTATTTAGTAGAGTTCCTTGAGTTTTTGTCACAAATATTATTATC
TACTTCTATCATATACTACTATTCATAGCTAGCTACGCATTGTTTCCTCAATCCATACGCAGTTGTACTG
TTACATTCACAACCCGTATCCCTCATGACCTCTTGACTGTATCCTACAGTACGTATTTCCTTTATTTTTC
TCTTCCACGAGCTACAGTAGCCACACCCCTACGGTACGGTACAGTATTGATAAATCAACCGACGACAGCG
CCTCTCAACTGCCAAACAGCTGGAATGCGTTCGTTTGAGAATGACCCCAAGACAAAAGATCCATTTTCAA
CCTACAGTAACGTCCGCCCACGTTTGCAGGATCCAAAAATGCGAGGAATCAATCCAGTCGAACTAGTCGA
GAAAGGGCAACAAAGATCCAATTTCCGTACTAGACAGATGTACAGTATGGAATCGAGTGAATCAAAACAG
TCGATTCAGTCGAACTAGTCGAGGATTGTAATCAATACTGTACATAGTTTGCGCTTTGCCTTCCACTAAG
TTACCGGTACAGGAGTAGTAGATTTTGTGAGATTTTTCATGACAAACAAATCGAAAAGTAAAAAAAAAAC
CTATTAGAAATCAACAATGACGACGATGATGATGTAACATGATGAGGAGGATAATCATAATAGTGACAAT
TCAACTATCGTATCACCTTATTTCTTCACAATCAGCCAAAAGAAAATTCCCTAGAGGTCAGCAGAAAAAA
TTACGAAAAGAACGGCGTACTGTACAGCAACGGCGAAGGATTGACAAATGAATAGTATTATTAGAATAAA
AAGTGTTTGGTTTTCGTTATCGTTATATTTATCGGTATTGTTACTATTACTATTTAGTTTGGTTCTATTT
TTTAATAGTAATAATAATAATAATAATGATACAACAATCATAGTAGTGGGCGTTAATGGATTTCTTCTAC
CAACAAGAACAACAATAACGCCGACATCAGCACAGCGACTACTGTTGTCGTCGACTAAGGCATGCAAGCT
TGGCGTAATCATGGTCATAGCTGTTTCCTGTGTGAAATTGTTATCCGCTCACAATTCCACACAACATACG
AGCCGGAAGCATAAAGTGTAAAGCCTGGGGTGCCTAATGAGTGAGCTAACTCACATTAATTGCGTTGCGC
TCACTGCCCGCTTTCCAGTCGGGAAACCTGTCGTGCCAGCTGCATTAATGAATCGGCCAACGCGCGGGGA
GAGGCGGTTTGCGTATTGGGCGCTCTTCCGCTTCCTCGCTCACTGACTCGCTGCGCTCGGTCGTTCGGCT
GCGGCGAGCGGTATCAGCTCACTCAAAGGCGGTAATACGGTTATCCACAGAATCAGGGGATAACGCAGGA
AAGAACATGTGAGCAAAAGGCCAGCAAAAGGCCAGGAACCGTAAAAAGGCCGCGTTGCTGGCGTTTTTCC
ATAGGCTCCGCCCCCCTGACGAGCATCACAAAAATCGACGCTCAAGTCAGAGGTGGCGAAACCCGACAGG
ACTATAAAGATACCAGGCGTTTCCCCCTGGAAGCTCCCTCGTGCGCTCTCCTGTTCCGACCCTGCCGCTT
ACCGGATACCTGTCCGCCTTTCTCCCTTCGGGAAGCGTGGCGCTTTCTCATAGCTCACGCTGTAGGTATC
TCAGTTCGGTGTAGGTCGTTCGCTCCAAGCTGGGCTGTGTGCACGAACCCCCCGTTCAGCCCGACCGCTG
CGCCTTATCCGGTAACTATCGTCTTGAGTCCAACCCGGTAAGACACGACTTATCGCCACTGGCAGCAGCC

ACTGGTAACAGGATTAGCAGAGCGAGGTATGTAGGCGGTGCTACAGAGTTCTTGAAGTGGTGGCCTAACT
ACGGCTACACTAGAAGAACAGTATTTGGTATCTGCGCTCTGCTGAAGCCAGTTACCTTCGGAAAAAGAGT
TGGTAGCTCTTGATCCGGCAAACAAACCACCGCTGGTAGCGGTGGTTTTTTTGTTTGCAAGCAGCAGATT
ACGCGCAGAAAAAAAGGATCTCAAGAAGATCCTTTGATCTTTTCTACGGGGTCTGACGCTCAGTGGAACG
AAAACTCACGTTAAGGGATTTTGGTCATGAGATTATCAAAAAGGATCTTCACCTAGATCCTTTTAAATTA
AAAATGAAGTTTTAAATCAATCTAAAGTATATATGAGTAAACTTGGTCTGACAGTTACCAATGCTTAATC
AGTGAGGCACCTATCTCAGCGATCTGTCTATTTCGTTCATCCATAGTTGCCTGACTCCCCGTCGTGTAGA
TAACTACGATACGGGAGGGCTTACCATCTGGCCCCAGTGCTGCAATGATACCGCGAGACCCACGCTCACC
GGCTCCAGATTTATCAGCAATAAACCAGCCAGCCGGAAGGGCCGAGCGCAGAAGTGGTCCTGCAACTTTA
TCCGCCTCCATCCAGTCTATTAATTGTTGCCGGGAAGCTAGAGTAAGTAGTTCGCCAGTTAATAGTTTGC
GCAACGTTGTTGCCATTGCTACAGGCATCGTGGTGTCACGCTCGTCGTTTGGTATGGCTTCATTCAGCTC
CGGTTCCCAACGATCAAGGCGAGTTACATGATCCCCCATGTTGTGCAAAAAAGCGGTTAGCTCCTTCGGT
CCTCCGATCGTTGTCAGAAGTAAGTTGGCCGCAGTGTTATCACTCATGGTTATGGCAGCACTGCATAATT
CTCTTACTGTCATGCCATCCGTAAGATGCTTTTCTGTGACTGGTGAGTACTCAACCAAGTCATTCTGAGA
ATAGTGTATGCGGCGACCGAGTTGCTCTTGCCCGGCGTCAATACGGGATAATACCGCGCCACATAGCAGA
ACTTTAAAAGTGCTCATCATTGGAAAACGTTCTTCGGGGCGAAAACTCTCAAGGATCTTACCGCTGTTGA
GATCCAGTTCGATGTAACCCACTCGTGCACCCAACTGATCTTCAGCATCTTTTACTTTCACCAGCGTTTC
TGGGTGAGCAAAAACAGGAAGGCAAAATGCCGCAAAAAAGGGAATAAGGGCGACACGGAAATGTTGAATA
CTCATACTCTTCCTTTTTCAATATTATTGAAGCATTTATCAGGGTTATTGTCTCATGAGCGGATACATAT
TTGAATGTATTTAGAAAAATAAACAAATAGGGGTTCCGCGCACATTTCCCCGAAAAGTGCCACCTGACGT
CTAAGAAACCATTATTATCATGACATTAACCTATAAAAATAGGCGTATCACGAGGCCCTTTCGTCTCGCG
CGTTTCGGTGATGACGGTGAAAACCTCTGACACATGCAGCTCCCGGAGACGGTCACAGCTTGTCTGTAAG
CGGATGCCGGGAGCAGACAAGCCCGTCAGGGCGCGTCAGCGGGTGTTGGCGGGTGTCGGGGCTGGCTTAA
CTATGCGGCATCAGAGCAGATTGTACTGAGAGTGCACCATATGCGGTGTGAAATACCGCACAGATGCGTA
AGGAGAAAATACCGCATCAGGCGCCATTCGCCATTCAGGCTGCGCAACTGTTGGGAAGGGCGATCGGTGC
GGGCCTCTTCGCTATTACGCCAGCTGGCGAAAGGGGGA

#### *Thalassiosira pseudonana*

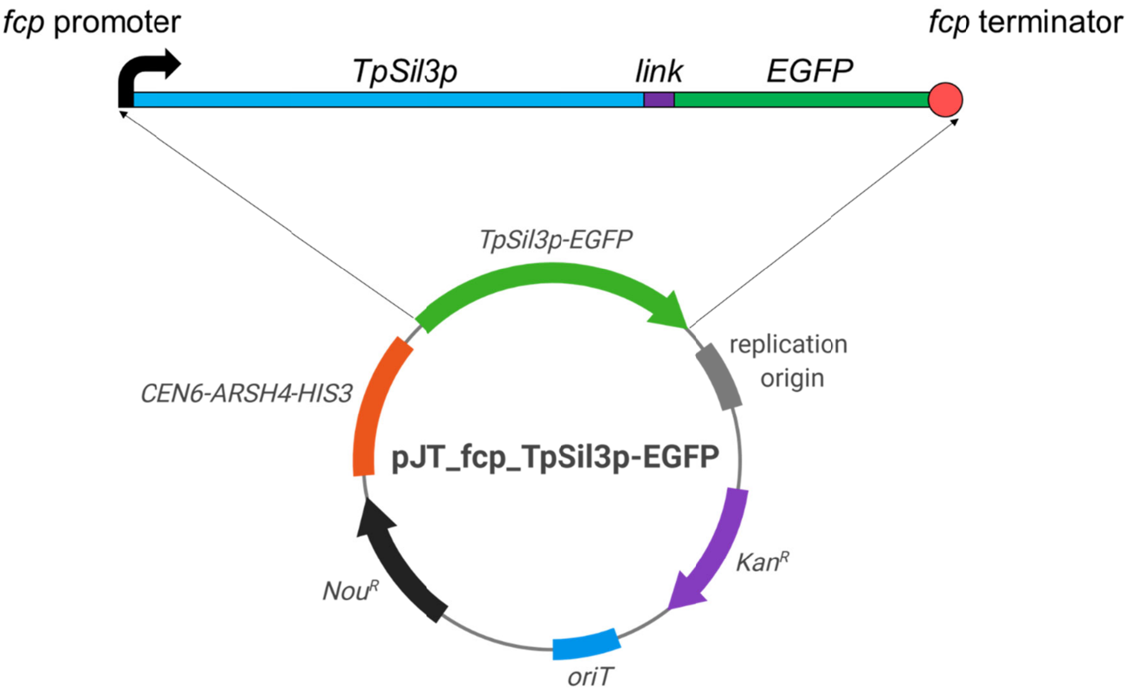


Expression cassette elements and their nucleotide sequences

```
>fcp/p-TpSil3p-link-EGFP-fcp/t
GCGCTTTTTCCGAGAACTCCCCATAAGTCAACGGCTCCAATCAAGAATGTATCCGACAACGGCGAGCATAGCAACACGTCCGTCTT
TGGAGTAGAATCATCATGTTGTGGATGAATACACAGATGAATGACATTAAAAGCATGAACATGTTAGAGAGTAGGAGGTAGAGATT
GATATGGTAGCATTGCGATGTTTGTTTTTGGTCAGCATATGATGAGTGGATACCAATATGATGAAAGTTGAATCTCGCGTTTGAGC
TCAGCGGTACGTTATTGATCGAAAGTAGCCTGATCAAAATCCTTGGAGAGTACAAGAGGATCAAAGAATCCAGTGGGGGCGATAAC
TCCAAGCTCGTTCTCAAAGAGGCAATGGAGGTAGAAACTCATCCCAGTTGAGAAGAAGTGAAGGCAGTGGCGGTGGCGAAAGCAGA
GGCAACGAGGACAGACTTCCTGTGGGTTGATGCAACGAATATTTCCAGAAGGAGAAGTTTAGAGAGTTGAACCGCTACCTACAATG
ACAAAGTATCGTATCGATTTTGATGTTGGTTGGTTATGAATTCAAACTGTAAGTTGGATTGTGAGAAGATCAGAAGTTGAACGAAC
ACATCTTTCCGATCATTCACCTCCACACTGCAACAACACGGTACTTCTTCCGCGGCAGGTCTCTGTCGCCATTCTCTTGTCCTGTT
GTTGGCTGTGAGACGAGGAAAGCAACGACAAGTTTCACAAAAGGGAGTTCCTTTAACGAGATATGTTTTTTATAAAGAGTCCCAAT
AGAAAGACAAATTGATTCCTCCGTGCAAACGCGCAAATAAACACCACGTCCATTATATCCATATCTTTCAGAGTATCCAACAAGTG
TTGAAGGACAGGTAGTTGAAGTAACGTATCTTCCCCCTCGACTGGATCCATCAACAAGGCGAACAAATCCATTCAACCTCTCATAA
ATTATCTGATTTACCAAACCATGAAGACTTCTGCCATTGCATTGCTTGCCGTTCTCGCCACCACTGCTGCCACCGAGCCCCGCCGA
TTGAGAACTCTTGAAGGACATGGGGGAGATCACTCCATCTCCATGTCCATGCACAGCTCGAAAGCTGAGAAGCAAGCCATCGAGGC
AGCTGTTGAGGAGGATGTTGCTGGCCCTGCAAAGGCAGCCAAGCTTTTCAAGCCCAAAGCAAGCAAGGCTGGTTCCATGCCTGATG
AGGCCGGTGCAAAGAGTGCCAAGATGAGCATGGACACCAAGAGTGGAAAGTCGGAGGACGCAGCTGCCGTAGATGCCAAAGCTTCC
AAGGAATCTCACATGTCTATAAGTGGTGATATGAGCATGGCCAAGTCACACAAGGCCGAGGCCGAGGACGTCACTGAGATGTCCAT
GGCAAAGGCCGGCAAGGATGAGGCTTCAACCGAGGATATGTGTATGCCCTTCGCTAAAAGTGACAAGGAAATGAGCGTCAAATCGA
AGCAAGGAAAGACCGAGATGAGTGTGGCCGATGCCAAGGCCTCAAAGGAGTCTAGCATGCCCTCTTCGAAGGCTGCCAAGATCTTC
AAGGGAAAGAGTGGGAAGTCCGGGAGTCTCTCCATGCTCAAGAGTGAAAAGGCAAGCTCCGCTCACAGCCTCAGTATGCCAAAAGC
TGAGAAGGTCCACTCCATGAGCGCTAAAGGGAGTGGTTCTACGTCTGGAAGCGGAATGGTGTCTAAGGGAGAGGAGCTTTTTACCG
GAGTAGTCCCGATTTTGGTCGAGCTTGATGGAGATGTCAACGGACACAAATTCAGTGTCAGCGGAGAGGGCGAGGGCGATGCCACG
TATGGAAAGCTTACTCTCAAGTTCATTTGTACTACGGGTAAGTTGCCAGTGCCCTGGCCCACACTCGTCACCACCCTTACCTATGG
TGTGCAATGCTTTTCTCGTTACCCCGACCACATGAAGCAGCATGACTTCTTTAAGTCCGCTATGCCAGAGGGATACGTCCAAGAAC
GTACAATCTTCTTCAAGGACGATGGAAATTATAAGACTCGTGCGGAGGTCAAGTTTGAGGGGGACACCCTCGTAAATCGCATCGAG
CTTAAGGGAATCGACTTCAAGGAAGATGGCAACATTTTGGGTCATAAGCTCGAATATAATTACAATTCCCATAACGTGTACATCAT
GGCAGACAAACAAAAGAACGGTATCAAGGTTAATTTTAAGATCCGCCATAACATCGAAGACGGAAGCGTTCAGTTGGCTGATCACT
ACCAGCAGAACACACCAATTGGAGACGGACCAGTGTTGCTTCCCGATAACCACTATCTTTCTACCCAATCTGCGTTGTCCAAAGAT
CCTAACGAGAAGCGTGATCATATGGTTCTTTTGGAGTTTGTCACAGCAGCCGGAATCACTTTGGGTATGGACGAGCTCTATAAGTA
AATACTGGATTGGTGAATCAATGAGCCGTAGCACAATGGTTACATTCGGCTAGCTAAGATCCAATGGCAAGGACCAAGTGCTGGAA
CTTGTTTTGCTTTAGCAGATCTTAGCGTGAGAGGTATTTGTCCTCTGTCAGGAGTAGATAGTAGATGTTCTTTTTAAACTAAAATG
CTAACTGTTCCGAATTCCTCATCGCAGCTAATCCGTACATCAAAAGACAAAATGCTAGGTATGTGTACTACATCTCCTGTTGCTAG
ATAAGACATATGATAGGAAACACACCATCAATAGTCATTGTAGCTTTACTTATACTACGCATTTGCACTTTCCCCTGAGTGGCAGA
```

GGCGCATTGAGAAAATCGATCTCAACATAGTTTATGTAGCATCCCCTAGATCCATTACTTTAAGTCTCCTTCGTCTTTGGTGTAGG
CATGTTGGACACAACGAGGTAAAACACAACACAAACAATGTGTCCAGCAAAGTAGTAGCTGCTCCAGTTCT

Fusion protein amino acid sequence

>TpSil3p-link-EGFP
MKTSAIALLAVLATTAATEPRRLRTLEGHGGDHSISMSMHSSKAEKQAIEAAVEEDVAGPAKAAKLFKPKASKAGSMPDEAGAKSA
KMSMDTKSGKSEDAAAVDAKASKESHMSISGDMSMAKSHKAEAEDVTEMSMAKAGKDEASTEDMCMPFAKSDKEMSVKSKQGKTEM
SVADAKASKESSMPSSKAAKIFKGKSGKSGSLSMLKSEKASSAHSLSMPKAEKVHSMSAKGSGSTSGSGMVSKGEELFTGVVPILV
ELDGDVNGHKFSVSGEGEGDATYGKLTLKFICTTGKLPVPWPTLVTTLTYGVQCFSRYPDHMKQHDFFKSAMPEGYVQERTIFFKD
DGNYKTRAEVKFEGDTLVNRIELKGIDFKEDGNILGHKLEYNYNSHNVYIMADKQKNGIKVNFKIRHNIEDGSVQLADHYQQNTPI
GDGPVLLPDNHYLSTQSALSKDPNEKRDHMVLLEFVTAAGITLGMDELYK

Full vector sequence with annotated expression cassette elements

TCGCGCGTTTCGGTGATGACGGTGAAAACCTCTGACACATGCAGCTCCCGGAGACGGTCACAGCTTGTCTGTAAGCGGATGCCGGG
AGCAGACAAGCCCGTCAGGGCGCGTCAGCGGGTGTTGGCGGGTGTCGGGGCTGGCTTAACTATGCGGCATCAGAGCAGATTGTACT
GAGAGTGCACCAGATCGTCTTGCCTTGCTCGTCGGTGATGTACTTACAGCTCGAAGTGCCTCTTCTTGATGGAGCGCATGGGGACG
TGCTTGGCAATCACGCGCACCCCCCGGCCGTTTTAGCGGCTAAAAAAGTCATGGCTCTGCCCTCGGGCGGACCACGCCCATCATGA
CCTTGCCAAGCTCGTCCTGCTTCTCTTCGATCTTCGCCAGCAGGGCGAGGATCGTGGCATCACCGAACCGCGCCGTGCGCGGGTCG
TCGGTGAGCCAGAGTTTCAGCAGGCCGCCCAGGCGGCCCAGGTCGCCATTGATGCGGGCCAGCTCGCGGACGTGCTCATAGTCCAC
GACGCCCGTGATTTTGTAGCCCTGGCCGACGGCCAGCAGGTAGGCCGACAGGCTCATGCCGGCCGCCGCCGCCTTTTCCTCAATCG
CTCTTCGTTCGTCTGGAAGGCAGTACACCTTGATAGGTGGGCTGCCCTTCCTGGTTGGCTTGGTTTCATCAGCCATCCGCTTGCCC
TCATCTGTTACGCCGGCGGTAGCCGGCCAGCCTCGCAGAGCAGGATTCCCGTTGAGCACCGCCAGGTGCGAATAAGGGACAGTGAA
GAAGGAACACCCGCTCGCGGGTGGGCCTACTTCACCTATCCTGCCCGGCTGACGCCGTTGGATACACCAAGGAAAGTCTACACGAA
CCCTTTGGCAAAATCCTGTATATCGTGCGAAAAAGGATGGATATACCGAAAAAATCGCTATAATGACCCCGAAGCAGGGTTATGCA
GCGGAAGATGGCCATTCGCCATTCAGGCTGCGCAACTGTTGGGAAGGGCGATCGGTGCGGGCCTCTTCGCTATTACGCCAGCTGGC
GAAAGGGGGATGTGCTGCAAGGCGATTAAGTTGGGTAACGCCAGGGTTTTCCCAGTCACGACGTTGTAAAACGACGGCCAGTGAGC
TTGCGCTTTTTCCGAGAACTCCCCATAAGTCAACGGCTCCAATCAAGAATGTATCCGACAACGGCGAGCATAGCAACACGTCCGTC
TTTGGAGTAGAATCATCATGTTGTGGATGAATACACAGATGAATGACATTAAAAGCATGAACATGTTAGAGAGTAGGAGGTAGAGA
TTGATATGGTAGCATTGCGATGTTTGTTTTTGGTCAGCATATGATGAGTGGATACCAATATGATGAAAGTTGAATCTCGCGTTTGA
GCTCAGCGGTACGTTATTGATCGAAAGTAGCCTGATCAAAATCCTTGGAGAGTACAAGAGGATCAAAGAATCCAGTGGGGGCGATA
ACTCCAAGCTCGTTCTCAAAGAGGCAATGGAGGTAGAAACTCATCCCAGTTGAGAAGAAGTGAAGGCAGTGGCGGTGGCGAAAGCA
GAGGCAACGAGGACAGACTTCCTGTGGGTTGATGCAACGAATATTTCCAGAAGGAGAAGTTTAGAGAGTTGAACCGCTACCTACAA
TGACAAAGTATCGTATCGATTTTGATGTTGGTTGGTTATGAATTCAAACTGTAAGTTGGATTGTGAGAAGATCAGAAGTTGAACGA
ACACATCTTTCCGATCATTCACCTCCACACTGCAACAACACGGTACTTCTTCCGCGGCAGGTCTCTGTCGCCATTCTCTTGTCCTG
TTGTTGGCTGTGAGACGAGGAAAGCAACGACAAGTTTCACAAAAGGGAGTTCCTTTAACGAGATATGTTTTTTATAAAGAGTCCCA
ATAGAAAGACAAATTGATTCCTCCGTGCAAACGCGCAAATAAACACCACGTCCATTATATCCATATCTTTCAGAGTATCCAACAAG
TGTTGAAGGACAGGTAGTTGAAGTAACGTATCTTCCCCCTCGACTGGATCCATCAACAAGGCGAACAAATCCATTCAACCTCTCAT
AAATTATCTGATTTACCAAACCGATACCAAAATGACCACTCTTGACGACACGGCTTACCGGTACCGCACCAGTGTCCCGGGGGACG
CCGAGGCCATCGAGGCACTGGATGGGTCCTTCACCACCGACACCGTCTTCCGCGTCACCGCCACCGGGGACGGCTTCACCCTGCGG
GAGGTGCCGGTGGACCCGCCCCTGACCAAGGTGTTCCCCGACGACGAATCGGACGACGAATCGGACGCCGGGGAGGACGGCGACCC
GGACTCCCGGACGTTCGTCGCGTACGGGGACGACGGCGACCTGGCGGGCTTCGTGGTCGTCTCGTACTCCGGCTGGAACCGCCGGC
TGACCGTCGAGGACATCGAGGTCGCCCCGGAGCACCGGGGGCACGGGGTCGGGCGCGCGTTGATGGGGCTCGCGACGGAGTTCGCC
CGCGAGCGGGGCGCCGGGCACCTCTGGCTGGAGGTCACCAACGTCAACGCACCGGCGATCCACGCGTACCGGCGGATGGGGTTCAC
CCTCTGCGGCCTGGACACCGCCCTGTACGACGGCACCGCCTCGGACGGCGAGCAGGCGCTCTACATGAGCATGCCCTGCCCCTGAG
CGCGGCCGCATACTGGATTGGTGAATCAATGAGCCGTAGCACAATGGTTACATTCGGCTAGCTAAGATCCAATGGCAAGGACCAAG
TGCTGGAACTTGTTTTGCTTTAGCAGATCTTAGCGTGAGAGGTATTTGTCCTCTGTCAGGAGTAGATAGTAGATGTTCTTTTTAAA
CTAAAATGCTAACTGTTCCGAATTCCTCATCGCAGCTAATCCGTACATCAAAAGACAAAATGCTAGGTATGTGTACTACATCTCCT
GTTGCTAGATAAGACATATGATAGGAAACACACCATCAATAGTCATTGTAGCTTTACTTATACTACGCATTTGCACTTTCCCCTGA
GTGGCAGAGGCGCATTGAGAAAATCGATCTCAACATAGTTTATGTAGCATCCCCTAGATCCATTACTTTAAGTCTCCTTCGTCTTT
GGTGTAGGCATGTTGGACACAACGAGGTAAAACACAACACAAACAATGTGTCCAGCAAAGTAGTAGCTGCTCCAGTTCTCCCGCGA
GCATCACGTGCTATAAAAATAATTATAATTTAAATTTTTTAATATAAATATATAAATTAAAAATAGAAAGTAAAAAAAGAAATTAA
AGAAAAAATAGTTTTTGTTTTCCGAAGATGTAAAAGACTCTAGGGGGATCGCCAACAAATACTACCTTTTATCTTGCTCTTCCTGC
TCTCAGGTATTAATGCCGAATTGTTTCATCTTGTCTGTGTAGAAGACCACACACGAAAATCCTGTGATTTTACATTTTACTTATCG
TTAATCGAATGTATATCTATTTAATCTGCTTTTCTTGTCTAATAAATATATATGTAAAGTACGCTTTTTGTTGAAATTTTTTAAAC
CTTTGTTTATTTTTTTTTCTTCATTCCGTAACTCTTCTACCTTCTTTATTTACTTTCTAAAATCCAAATACAAAACATAAAAATAA
ATAAACACAGAGTAAATTCCCAAATTATTCCATCATTAAAAGATACGAGGCGCGTGTAAGTTACAGGCAAGCGATCCTAGTACACT
CTATATTTTTTTATGCCTCGGTAATGATTTTCATTTTTTTTTTTCCACCTAGCGGATGACTCTTTTTTTTTCTTAGCGATTGGCAT
TATCACATAATGAATTATACATTATATAAAGTAATGTGATTTCTTCGAAGAATATACTAAAAAATGAGCAGGCAAGATAAACGAAG
GCAAAGATGACAGAGCAGAAAGCCCTAGTAAAGCGTATTACAAATGAAACCAAGATTCAGATTGCGATCTCTTTAAAGGGTGGTCC
CCTAGCGATAGAGCACTCGATCTTCCCAGAAAAAGAGGCAGAAGCAGTAGCAGAACAGGCCACACAATCGCAAGTGATTAACGTCC
ACACAGGTATAGGGTTTCTGGACCATATGATACATGCTCTGGCCAAGCATTCCGGCTGGTCGCTAATCGTTGAGTGCATTGGTGAC
TTACACATAGACGACCATCACACCACTGAAGACTGCGGGATTGCTCTCGGTCAAGCTTTTAAAGAGGCCCTACTGGCGCGTGGAGT
AAAAAGGTTTGGATCAGGATTTGCGCCTTTGGATGAGGCACTTTCCAGAGCGGTGGTAGATCTTTCGAACAGGCCGTACGCAGTTG
TCGAACTTGGTTTGCAAAGGGAGAAAGTAGGAGATCTCTCTTGCGAGATGATCCCGCATTTTCTTGAAAGCTTTGCAGAGGCTAGC
AGAATTACCCTCCACGTTGATTGTCTGCGAGGCAAGAATGATCATCACCGTAGTGAGAGTGCGTTCAAGGCTCTTGCGGTTGCCAT
AAGAGAAGCCACCTCGCCCAATGGTACCAACGATGTTCCCTCCACCAAAGGTGTTCTTATGTAGTTTTACACAGGAGTCTGGACTT
GACCTCTAGA**GCGCTTTTTCCGAGAACTCCCCATAAGTCAACGGCTCCAATCAAGAATGTATCCGACAACGGCGAGCATAGCAACA**
**CGTCCGTCTTTGGAGTAGAATCATCATGTTGTGGATGAATACACAGATGAATGACATTAAAAGCATGAACATGTTAGAGAGTAGGA**
**GGTAGAGATTGATATGGTAGCATTGCGATGTTTGTTTTTGGTCAGCATATGATGAGTGGATACCAATATGATGAAAGTTGAATCTC**
**GCGTTTGAGCTCAGCGGTACGTTATTGATCGAAAGTAGCCTGATCAAAATCCTTGGAGAGTACAAGAGGATCAAAGAATCCAGTGG**
**GGGCGATAACTCCAAGCTCGTTCTCAAAGAGGCAATGGAGGTAGAAACTCATCCCAGTTGAGAAGAAGTGAAGGCAGTGGCGGTGG**
**CGAAAGCAGAGGCAACGAGGACAGACTTCCTGTGGGTTGATGCAACGAATATTTCCAGAAGGAGAAGTTTAGAGAGTTGAACCGCT**

**ACCTACAATGACAAAGTATCGTATCGATTTTGATGTTGGTTGGTTATGAATTCAAACTGTAAGTTGGATTGTGAGAAGATCAGAAG**
**TTGAACGAACACATCTTTCCGATCATTCACCTCCACACTGCAACAACACGGTACTTCTTCCGCGGCAGGTCTCTGTCGCCATTCTC**
**TTGTCCTGTTGTTGGCTGTGAGACGAGGAAAGCAACGACAAGTTTCACAAAAGGGAGTTCCTTTAACGAGATATGTTTTTTATAAA**
**GAGTCCCAATAGAAAGACAAATTGATTCCTCCGTGCAAACGCGCAAATAAACACCACGTCCATTATATCCATATCTTTCAGAGTAT**
**CCAACAAGTGTTGAAGGACAGGTAGTTGAAGTAACGTATCTTCCCCCTCGACTGGATCCATCAACAAGGCGAACAAATCCATTCAA**
**CCTCTCATAAATTATCTGATTTACCAAACC**ATGAAGACTTCTGCCATTGCATTGCTTGCCGTTCTCGCCACCACTGCTGCCACCGA
GCCCCGCCGATTGAGAACTCTTGAAGGACATGGGGGAGATCACTCCATCTCCATGTCCATGCACAGCTCGAAAGCTGAGAAGCAAG
CCATCGAGGCAGCTGTTGAGGAGGATGTTGCTGGCCCTGCAAAGGCAGCCAAGCTTTTCAAGCCCAAAGCAAGCAAGGCTGGTTCC
ATGCCTGATGAGGCCGGTGCAAAGAGTGCCAAGATGAGCATGGACACCAAGAGTGGAAAGTCGGAGGACGCAGCTGCCGTAGATGC
CAAAGCTTCCAAGGAATCTCACATGTCTATAAGTGGTGATATGAGCATGGCCAAGTCACACAAGGCCGAGGCCGAGGACGTCACTG
AGATGTCCATGGCAAAGGCCGGCAAGGATGAGGCTTCAACCGAGGATATGTGTATGCCCTTCGCTAAAAGTGACAAGGAAATGAGC
GTCAAATCGAAGCAAGGAAAGACCGAGATGAGTGTGGCCGATGCCAAGGCCTCAAAGGAGTCTAGCATGCCCTCTTCGAAGGCTGC
CAAGATCTTCAAGGGAAAGAGTGGGAAGTCCGGGAGTCTCTCCATGCTCAAGAGTGAAAAGGCAAGCTCCGCTCACAGCCTCAGTA
TGCCAAAAGCTGAGAAGGTCCACTCCATGAGCGCTAAAGGGAGTGGTTCTACGTCTGGAAGCGGAATGGTGTCTAAGGGAGAGGAG
CTTTTTACCGGAGTAGTCCCGATTTTGGTCGAGCTTGATGGAGATGTCAACGGACACAAATTCAGTGTCAGCGGAGAGGGCGAGGG
CGATGCCACGTATGGAAAGCTTACTCTCAAGTTCATTTGTACTACGGGTAAGTTGCCAGTGCCCTGGCCCACACTCGTCACCACCC
TTACCTATGGTGTGCAATGCTTTTCTCGTTACCCCGACCACATGAAGCAGCATGACTTCTTTAAGTCCGCTATGCCAGAGGGATAC
GTCCAAGAACGTACAATCTTCTTCAAGGACGATGGAAATTATAAGACTCGTGCGGAGGTCAAGTTTGAGGGGGACACCCTCGTAAA
TCGCATCGAGCTTAAGGGAATCGACTTCAAGGAAGATGGCAACATTTTGGGTCATAAGCTCGAATATAATTACAATTCCCATAACG
TGTACATCATGGCAGACAAACAAAAGAACGGTATCAAGGTTAATTTTAAGATCCGCCATAACATCGAAGACGGAAGCGTTCAGTTG
GCTGATCACTACCAGCAGAACACACCAATTGGAGACGGACCAGTGTTGCTTCCCGATAACCACTATCTTTCTACCCAATCTGCGTT
GTCCAAAGATCCTAACGAGAAGCGTGATCATATGGTTCTTTTGGAGTTTGTCACAGCAGCCGGAATCACTTTGGGTATGGACGAGC
TCTATAAGTAAATACTGGATTGGTGAATCAATGAGCCGTAGCACAATGGTTACATTCGGCTAGCTAAGATCCAATGGCAAGGACCA
AGTGCTGGAACTTGTTTTGCTTTAGCAGATCTTAGCGTGAGAGGTATTTGTCCTCTGTCAGGAGTAGATAGTAGATGTTCTTTTTA
AACTAAAATGCTAACTGTTCCGAATTCCTCATCGCAGCTAATCCGTACATCAAAAGACAAAATGCTAGGTATGTGTACTACATCTC
CTGTTGCTAGATAAGACATATGATAGGAAACACACCATCAATAGTCATTGTAGCTTTACTTATACTACGCATTTGCACTTTCCCCT
GAGTGGCAGAGGCGCATTGAGAAAATCGATCTCAACATAGTTTATGTAGCATCCCCTAGATCCATTACTTTAAGTCTCCTTCGTCT
TTGGTGTAGGCATGTTGGACACAACGAGGTAAAACACAACACAAACAATGTGTCCAGCAAAGTAGTAGCTGCTCCAGTTCTCTGCA
GGCATGCAAGCTTGGCGTAATCATGGTCATAGCTGTTTCCTGTGTGAAATTGTTATCCGCTCACAATTCCACACAACATACGAGCC
GGAAGCATAAAGTGTAAAGCCTGGGGTGCCTAATGAGTGAGCTAACTCACATTAATTGCGTTGCGCTCACTGCCCGCTTTCCAGTC
GGGAAACCTGTCGTGCCAGCTGCATTAATGAATCGGCCAACGCGCGGGGAGAGGCGGTTTGCGTATTGGGCGCTCTTCCGCTTCCT
CGCTCACTGACTCGCTGCGCTCGGTCGTTCGGCTGCGGCGAGCGGTATCAGCTCACTCAAAGGCGGTAATACGGTTATCCACAGAA
TCAGGGGATAACGCAGGAAAGAACATGTGAGCAAAAGGCCAGCAAAAGGCCAGGAACCGTAAAAAGGCCGCGTTGCTGGCGTTTTT
CCATAGGCTCCGCCCCCCTGACGAGCATCACAAAAATCGACGCTCAAGTCAGAGGTGGCGAAACCCGACAGGACTATAAAGATACC
AGGCGTTTCCCCCTGGAAGCTCCCTCGTGCGCTCTCCTGTTCCGACCCTGCCGCTTACCGGATACCTGTCCGCCTTTCTCCCTTCG
GGAAGCGTGGCGCTTTCTCATAGCTCACGCTGTAGGTATCTCAGTTCGGTGTAGGTCGTTCGCTCCAAGCTGGGCTGTGTGCACGA
ACCCCCCGTTCAGCCCGACCGCTGCGCCTTATCCGGTAACTATCGTCTTGAGTCCAACCCGGTAAGACACGACTTATCGCCACTGG
CAGCAGCCACTGGTAACAGGATTAGCAGAGCGAGGTATGTAGGCGGTGCTACAGAGTTCTTGAAGTGGTGGCCTAACTACGGCTAC
ACTAGAAGAACAGTATTTGGTATCTGCGCTCTGCTGAAGCCAGTTACCTTCGGAAAAAGAGTTGGTAGCTCTTGATCCGGCAAACA
AACCACCGCTGGTAGCGGTGGTTTTTTTGTTTGCAAGCAGCAGATTACGCGCAGAAAAAAAGGATCTCAAGAAGATCCTTTGATCT
TTTCTACGGGGTCTGACGCTCAGTGGAACGAAAACTCACGTTAAGGGATTTTGGTCATGAGATTATCAAAAAGGATCTTCACCTAG
ATCCTTTTAAATTAAAAATGAAGTTTTAAATCAATCTAAAGTATATATGAGTAAACTTGGTCTGACAGTTACCAATGCTTAATCAG
TGAGGCACCTATCTCAGCGATCTGTCTATTTCGTTCATCCATAGTTGCCTGACTCCCCGTCGTGTAGATAACTACGATACGGGAGG
GCTTACCATCTGGCCCCAGTGCTGCAATGATACCGCGAGACCCACGCTCACCGGCTCCAGATTTATCAGCAATAAACCAGCCAGCC
GGAAGGGCCGAGCGCAGAAGTGGTCCTGCAACTTTATCCGCCTCCATCCAGTCTATTAATTGTTGCCGGGAAGCTAGAGTAAGTAG
TTCGCCAGTTAATAGTTTGCGCAACGTTGTTGCCATTGCTACAGGCATCGTGGTGTCACGCTCGTCGTTTGGTATGGCTTCATTCA
GCTCCGGTTCCCAACGATCAAGGCGAGTTACATGATCCCCCATGTTGTGCAAAAAAGCGGTTAGCTCCTTCGGTCCTCCGATCGTT
GTCAGAAGTAAGTTGGCCGCAGTGTTATCACTCATGGTTATGGCAGCACTGCATAATTCTCTTACTGTCATGCCATCCGTAAGATG
CTTTTCTGTGACTGGTGAGTCGATTTATTCAACAAAGCCACGTTGTGTCTCAAAATCTCTGATGTTACATTGCACAAGATAAAAAT
ATATCATCATGAACAATAAAACTGTCTGCTTACATAAACAGTAATACAAGGGGTGTTATGAGCCATATTCAACGGGAAACGTCTTG
CTCGAGGCCGCGATTAAATTCCAACATGGATGCTGATTTATATGGGTATAAATGGGCTCGCGATAATGTCGGGCAATCAGGTGCGA
CAATCTATCGATTGTATGGGAAGCCCGATGCGCCAGAGTTGTTTCTGAAACATGGCAAAGGTAGCGTTGCCAATGATGTTACAGAT
GAGATGGTCAGACTAAACTGGCTGACGGAATTTATGCCTCTTCCGACCATCAAGCATTTTATCCGTACTCCTGATGATGCATGGTT
ACTCACCACTGCGATCCCCGGGAAAACAGCATTCCAGGTATTAGAAGAATATCCTGATTCAGGTGAAAATATTGTTGATGCGCTGG
CAGTGTTCCTGCGCCGGTTGCATTCGATTCCTGTTTGTAATTGTCCTTTTAACAGCGATCGCGTATTTCGTCTCGCTCAGGCGCAA
TCACGAATGAATAACGGTTTGGTTGATGCGAGTGATTTTGATGACGAGCGTAATGGCTGGCCTGTTGAACAAGTCTGGAAAGAAAT
GCATAAGCTTTTGCCATTCTCACCGGATTCAGTCGTCACTCATGGTGATTTCTCACTTGATAACCTTATTTTTGACGAGGGGAAAT
TAATAGGTTGTATTGATGTTGGACGAGTCGGAATCGCAGACCGATACCAGGATCTTGCCATCCTATGGAACTGCCTCGGTGAGTTT
TCTCCTTCATTACAGAAACGGCTTTTTCAAAAATATGGTATTGATAATCCTGATATGAATAAATTGCAGTTTCATTTGATGCTCGA
TGAGTTTTTCTAATCAGAATTGGTTAATTGGTTGTAACACTGGCACTCAACCAAGTCATTCTGAGAATAGTGTATGCGGCGACCGA
GTTGCTCTTGCCCGGCGTCAATACGGGATAATACCGCGCCACATAGCAGAACTTTAAAAGTGCTCATCATTGGAAAACGTTCTTCG
GGGCGAAAACTCTCAAGGATCTTACCGCTGTTGAGATCCAGTTCGATGTAACCCACTCGTGCACCCAACTGATCTTCAGCATCTTT
TACTTTCACCAGCGTTTCTGGGTGAGCAAAAACAGGAAGGCAAAATGCCGCAAAAAAGGGAATAAGGGCGACACGGAAATGTTGAA
TACTCATACTCTTCCTTTTTCAATATTATTGAAGCATTTATCAGGGTTATTGTCTCATGAGCGGATACATATTTGAATGTATTTAG
AAAAATAAACAAATAGGGGTTCCGCGCACATTTCCCCGAAAAGTGCCACCTGACGTCTAAGAAACCATTATTATCATGACATTAAC
CTATAAAAATAGGCGTATCACGAGGCCCTTTCGTC

*Serninavis robusta*

pBS ActAgroCP4Gl

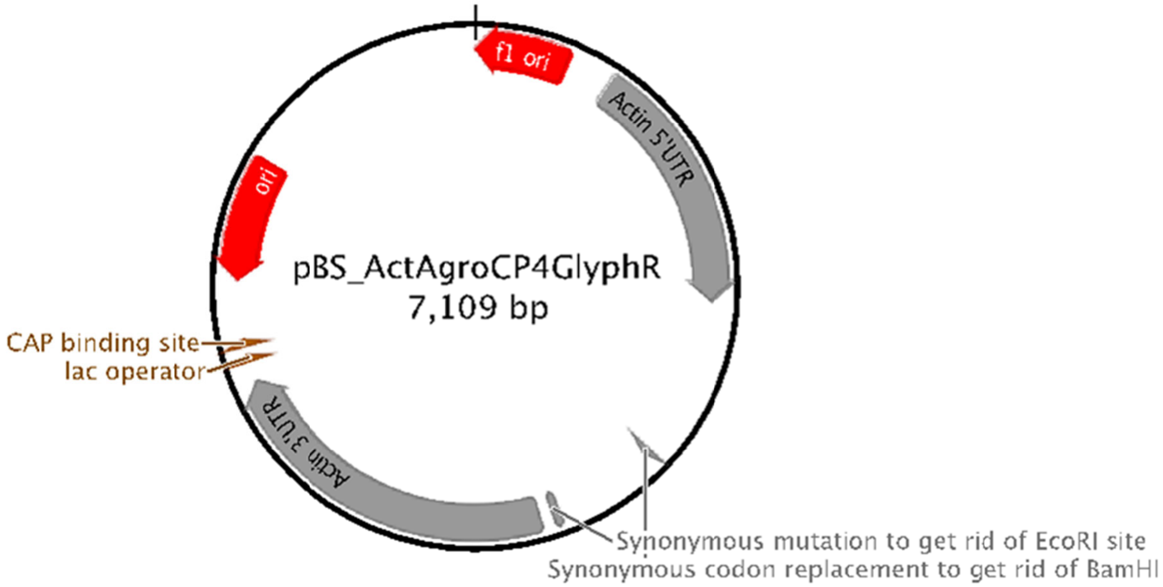


---

```
LOCUS       pBS_ActAgroCP4Gl        7109 bp ds-DNA     circular SYN 26-OCT-2019
 DEFINITION  .
 ACCESSION
 VERSION     .
 KEYWORDS    pBS_Sr_Act_AgroCP4GlyphR.
 SOURCE      null
   ORGANISM
             .
 REFERENCE   1  (bases 1 to 7109)
   AUTHORS   Turkewitz and Lab
   TITLE     Direct Submission
   JOURNAL   Exported Monday, Jun 24, 2019 from SnapGene Viewer 4.3.8
             https://www.snapgene.com
 FEATURES             Location/Qualifiers
      source          1..7109
                      /organism="recombinant plasmid"
                      /mol_type="other DNA"
      rep_origin      complement(3..458)
                      /direction="LEFT"
                      /note="f1 bacteriophage origin of replication; arrow
                      indicates direction of (+) strand synthesis"
                      /label="f1 ori"
      primer_bind     600..616
                      /note="common sequencing primer, one of multiple similar
                      variants"
                      /label="M13 fwd"
      promoter        626..644
                      /note="promoter for bacteriophage T7 RNA polymerase"
                      /label="T7 promoter"
      misc_feature    665..1864
                      /label="Actin 5'UTR"
      CDS             1871..3238
                      /codon_start=1
                      /translation="MSHGASSRPATARKSSGLSGTVRIPGDKSISHRSFMFGGLASGE
                      TRITGLLEGEDVINTGKAMQAMGARIRKEGDTWIIDGVGNGGLLAPEAPLDFGNAATG
                      CRLTMGLVGVYDFDSTFIGDASLTKRPMGRVLNPLREMGVQVKSEDGDRLPVTLRGPK
                      TPTPITYRVPMASAQVKSAVLLAGLNTPGITTVIEPIMTRDHTEKMLQGFGANLTVET
                      DADGVRTIRLEGRGKLTGQVIDVPGDPSSTAFPLVAALLVPGSDVTILNVLMNPTRTG
```

```
                     LILTLQEMGADIEVINPRLAGGEDVADLRVRSSTLKGVTVPEDRAPSMIDEYPILAVA
                     AAFAEGATVMNGLEELRVKESDRLSAVANGLKLNGVDCDEGETSLVVRGRPDGKGLGN
                     ASGAAVATHLDHRIAMSFLVMGLVSENPVTVDDATMIATSFPEFMDLMAGLGAKIELS
                     DTKAA"
                     /label="Agro CP4 AroA (GlyphR) Seminavis-Optimized"
     misc_feature    2651..2653
                     /label="Synonymous codon replacement to get rid of BamHI"
     misc_feature    3178
                     /label="Synonymous mutation to get rid of EcoRI site"
     misc_feature    3245..4884
                     /label="Actin 3'UTR"
     promoter        complement(4921..4939)
                     /note="promoter for bacteriophage T3 RNA polymerase"
                     /label="T3 promoter"
     primer_bind     complement(4960..4976)
                     /note="common sequencing primer, one of multiple similar
                     variants"
                     /label="M13 rev"
     protein_bind    4984..5000
                     /bound_moiety="lac repressor encoded by lacI"
                     /note="The lac repressor binds to the lac operator to
                     inhibit transcription in E. coli. This inhibition can be
                     relieved by adding lactose or
                     isopropyl-beta-D-thiogalactopyranoside (IPTG)."
                     /label="lac operator"
     promoter        complement(5008..5038)
                     /note="promoter for the E. coli lac operon"
                     /label="lac promoter"
     protein_bind    5053..5074
                     /bound_moiety="E. coli catabolite activator protein"
                     /note="CAP binding activates transcription in the presence
                      of cAMP."
                     /label="CAP binding site"
     rep_origin      complement(5362..5950)
                     /direction="LEFT"
                     /note="high-copy-number ColE1/pMB1/pBR322/pUC origin of
                     replication"
                     /label="ori"
     CDS             complement(6121..6981)
                     /codon_start=1
                     /gene="bla"
                     /product="beta-lactamase"
                     /note="confers resistance to ampicillin, carbenicillin,
                     and related antibiotics"
                     /translation="MSIQHFRVALIPFFAAFCLPVFAHPETLVKVKDAEDQLGARVGY
                     IELDLNSGKILESFRPEERFPMMSTFKVLLCGAVLSRIDAGQEQLGRRIHYSQNDLVE
                     YSPVTEKHLTDGMTVRELCSAAITMSDNTAANLLLTTIGGPKELTAFLHNMGDHVTRL
                     DRWEPELNEAIPNDERDTTMPVAMATTLRKLLTGELLTLASRQQLIDWMEADKVAGPL
                     LRSALPAGWFIADKSGAGERGSRGIIAALGPDGKPSRIVVIYTTGSQATMDERNRQIA
                     EIGASLIKHW"
                     /label="AmpR"
     promoter        complement(6982..7086)
                     /gene="bla"
                     /label="AmpR promoter"
ORIGIN
        1 ctaaattgta agcgttaata ttttgttaaa attcgcgtta aatttttgtt aaatcagctc
       61 attttttaac caataggccg aaatcggcaa aatcccttat aaatcaaaag aatagaccga
      121 gatagggttg agtgttgttc cagtttggaa caagagtcca ctattaaaga acgtggactc
      181 caacgtcaaa gggcgaaaaa ccgtctatca gggcgatggc ccactacgtg aaccatcacc
      241 ctaatcaagt tttttggggt cgaggtgccg taaagcacta aatcggaacc ctaaagggag
      301 cccccgattt agagcttgac ggggaaagcc ggcgaacgtg gcgagaaagg aagggaagaa
      361 agcgaaagga gcgggcgcta gggcgctggc aagtgtagcg gtcacgctgc gcgtaaccac
      421 cacacccgcc gcgcttaatg cgccgctaca gggcgcgtcc cattcgccat tcaggctgcg
      481 caactgttgg gaagggcgat cggtgcgggc ctcttcgcta ttacgccagc tggcgaaagg
      541 gggatgtgct gcaaggcgat taagttgggt aacgccaggg ttttcccagt cacgacgttg
      601 taaaacgacg gccagtgagc gcgcgtaata cgactcacta tagggcgaat tgggtaccgg
      661 gcccagtttt acatcaagac tgcatacaag atatcgagaa acgcgatgtt ggctaatggt
      721 atctttgtga ttgcaaaaag ggtgaagtcg agaagacatg atgacggtac cgtacggtgc
      781 taccggtacg tacgaaagat cgcgacttca gtcttgcttg aaggtcacac cacgtatcgt
      841 ggaaaacaaa aagcttgcac cgagggcaag tatctgtaat gttctggaag tcgctccacg
      901 gaaaaggtat caggccttcg tcgcaggata agtcacatgg ctgatgatgg gccaagcaag
      961 ggtaaccttt ctctttcagc ctttgaagac tggtaagata agctggtctg tttcttcact
     1021 catgttgttg aagtcgctcc acggaaaagg tatcaggcct tcgtcgcagg ataagtcaca
     1081 tggctgatga tgggccaagc aagggtaacc tttctcttcc tttctctttc agcctttgaa
     1141 gactggtaag ataagctggt ctgtttcttc actcatgttg ctttcgacgc aaaagcataa
     1201 accaagctgc aacgtggaga ggctatccct cgccagtgcc aagttcgagg catccaaacg
     1261 ccccattctc aacctcagtg tccccaaagc attcttgaca tgacggttct tgccaagacc
```

1321 gcttagctac tggtacagct acagcctttg tttcggcgtt gtctgacgcg acggaagcgg
1381 gtgggcttct caatatcttt tgtcgatgtt cgatgagcca gtcaaaaaag cgtgcatacc
1441 gaggataatg cttggtacca gcaaaatgtg aggtggtccc tcgttgcagc ccgtgaagcg
1501 aaagtcacaa tgactggaat gccaaagggc ggaatccaaa ccaaacgaaa tttgaaaaag
1561 ttggacctga aagatctata agatacgatt ccttctgcgt aaagatactt aacagtacaa
1621 gcaggacttc gtttcgagaa gggaattcca atgttttcct cccaagacca aacaaagatc
1681 tgccgcattc aagagagggg tcttgttggg acatctgcct tgacactgca cagcgggagg
1741 ttgcagcgac agcgcagtga acccattcaa caagattgct tggattctgc cataaaattg
1801 cttctcggta gcaacactct ctttctcgaa actctcttca aagaatcaaa caaactaatt
1861 caccctcgag atgtcccacg gagcctcctc ccgtcccgcc accgcccgta agtcctccgg
1921 attgtccgga accgtccgta ttcccggaga taagtccatt tcccaccgtt ccttcatgtt
1981 cggaggattg gcctccggag aaacccgtat taccggattg ttggaaggag aagatgtcat
2041 taacaccgga aaggccatgc aagccatggg agcccgtatt cgtaaggaag gagatacctg
2101 gattattgat ggagtcggaa acggaggatt gttggccccc gaagcccccct tggatttcgg
2161 aaacgccgcc accggatgcc gtttgaccat gggattggtc ggagtctacg atttcgattc
2221 caccttcatt ggagatgcct ccttgaccaa gcgtcccatg ggacgtgtct tgaacccctt
2281 gcgtgaaatg ggagtccaag tcaagtccga agatggagat cgtttgcccg tcaccttgcg
2341 tggacccaag acccccaccc ccattaccta ccgtgtcccc atggcctccg cccaagtcaa
2401 gtccgccgtc ttgttggccg gattgaacac ccccggaatt accaccgtca ttgaacccat
2461 tatgacccgt gatcacaccg aaaagatgtt gcaaggattc ggagccaact tgaccgtcga
2521 aaccgatgcc gatggagtcc gtaccattcg tttggaagga cgtggaaagt tgaccggaca
2581 agtcattgat gtccccggag atccctcctc caccgccttc cccttggtcg ccgccttgtt
2641 ggtccccgga agtgatgtca ccattttgaa cgtcttgatg aaccccaccc gtaccggatt
2701 gattttgacc ttgcaagaaa tgggagccga tattgaagtc attaaccccc gtttggccgg
2761 aggagaagat gtcgccgatt tgcgtgtccg ttcctccacc ttgaagggag tcaccgtccc
2821 cgaagatcgt gccccctcca tgattgatga ataccccatt ttggccgtcg ccgccgcctt
2881 cgccgaagga gccaccgtca tgaacggatt ggaagaattg cgtgtcaagg aatccgatcg
2941 tttgtccgcc gtcgccaacg gattgaagtt gaacggagtc gattgcgatg aaggagaaac
3001 ctccttggtc gtccgtggac gtcccgatgg aaagggattg ggaaacgcct ccggagccgc
3061 cgtcgccacc cacttggatc accgtattgc catgtccttc ttggtcatgg gattggtctc
3121 cgaaaacccc gtcaccgtcg atgatgccac catgattgcc acctccttcc ccgaatttat
3181 ggatttgatg gccggattgg gagccaagat tgaattgtcc gataccaagg ccgcctaact
3241 gcagggaatg ggccatgcgc aggcaaatgc cagctgctgt gtctgctctt tgtgaaatat
3301 tattgggcca gcagtctttt attttttgtt ggacctgggt tgcttgccgt tgttgaacaa
3361 catggtgatg tgaccgacgg atcaatgaga aaagcctaat tgttagaatg ttttattcat
3421 tgcggtccct tgtctctaca tagctagcta gctaactagc tagttttcga ctcctgatat
3481 caaggtattc ccgtcgttgc tggagccgat cttgccaagc cctggcaacc gcaccgtacc
3541 gattgctggg gggctgtagg tacacgtact ggaagcaacg ttgatcgatt ggcttttcgg
3601 gcaactggag ctgtgcgaca cgatctttga tagcttttcc ttagggttta accctaaaat
3661 attattttgc tggtttgtgc ccacagcgag ttcgatttag tcgaatcttt tttgggctcg
3721 atcggatcca tccatcttcg aagatctttc ttttgcacgc cattgattct ctcaaacgtt
3781 ctcgctcgtt ggtcctcaat aaccatgatg aacaacgcgt tgctaccgat tcgatccagc
3841 cttcccaagg ctatcagaag aacctcgtgt cgtcttattc gatcaggtcc gccttctcgc
3901 ttggtatcgc ccatcaggca agcatttcct gcatcacaga gtgcttttgc tttgcgtcct
3961 ttttcgagcg gcgcccagac gctttctgat gtgctggccc gagagttagc cgaagaaacc
4021 gaggaaggtc gagaccaaat ccccgaagaa cttgctgagc ttaagagttc tattgagaaa
4081 gattggaagg tcgttgacga tggtgcgatc actcggttga tccgatcgat tggagccagt
4141 aaagtggttg tctcgtttca ctgccaagac acggtggatg gcgcagagga ttatgtcgag
4201 gaagaagagg gagaagaaag aacctgcctt gccctttcga tttgagattc ttgtgtccaa
4261 ggcaggaaat acattggtgc ttaattgtat cagcaatgct ggtgaaacca cagtcgatgg
4321 ggttgccatg accactgaag acattgaatc ggttcaagcc aatggtattg gtcgcaacaa
4381 ttatcaggga cctgaatttc aagagctagc cgaggaccta caggaggcac ttcatgaata
4441 tgtcttttcg gagctaggta tcgatgaaga cgtctctgcc tttgtctcga tgtatgctga
4501 ttacaaggag caagtgcaat acatcggatt cttgacgaat gtttctaagg ttcttccgta
4561 agattcatgc tccctcttaa gtctgaacag acataattga aaatttagct agaaccaatc
4621 aacgaataaa aaatagagac tgatattgag cttgtttgga agcgggtgct ttttgggaaa
4681 cagaactggg gtactcctca cacgaagcca taaaagagtc acgcaacaac ggtatcgaca
4741 acgacaaagg catcagctgt ctcaacgtac cccgcgcagc tgatccttgt acggggtttg
4801 tatggttctc tcacgggtgg tacacaaagg gacgttcatg cagaccaagc atatcgtcag
4861 ttaccgccac tcagcagttg tactgcggcc gccaccgcgg tggagctcca gcttttgttc
4921 cctttagtga gggttaattg cgcgcttggc gtaatcatgg tcatagctgt ttcctgtgtg
4981 aaattgttat ccgctcacaa ttccacacaa catacgagcc ggaagcataa agtgtaaagc
5041 ctggggtgcc taatgagtga gctaactcac attaattgcg ttgcgctcac tgcccgcttt
5101 ccagtcggga aacctgtcgt gccagctgca ttaatgaatc ggccaacgcg cggggagagg
5161 cggtttgcgt attgggcgct cttccgcttc ctcgctcact gactcgctgc gctcggtcgt
5221 tcggctgcgg cgagcggtat cagctcactc aaaggcggta atacggttat ccacagaatc
5281 aggggataac gcaggaaaga acatgtgagc aaaaggccag caaaaggcca ggaaccgtaa
5341 aaaggccgcg ttgctggcgt ttttccatag gctccgcccc cctgacgagc atcacaaaaa
5401 tcgacgctca agtcagaggt ggcgaaaccc gacaggacta taaagatacc aggcgtttcc
5461 ccctggaagc tccctcgtgc gctctcctgt tccgaccctg ccgcttaccg gatacctgtc
5521 cgcctttctc ccttcgggaa gcgtggcgct ttctcatagc tcacgctgta ggtatctcag
5581 ttcggtgtag gtcgttcgct ccaagctggg ctgtgtgcac gaaccccccg ttcagcccga
5641 ccgctgcgcc ttatccggta actatcgtct tgagtccaac ccggtaagac acgacttatc
5701 gccactggca gcagccactg gtaacaggat tagcagagcg aggtatgtag gcggtgctac
5761 agagttcttg aagtggtggc ctaactacgg ctacactaga aggacagtat ttggtatctg
5821 cgctctgctg aagccagtta ccttcggaaa aagagttggt agctcttgat ccggcaaaca
5881 aaccaccgct ggtagcggtg gtttttttgt ttgcaagcag cagattacgc gcagaaaaaa

```
      5941 aggatctcaa gaagatcctt tgatcttttc tacggggtct gacgctcagt ggaacgaaaa
      6001 ctcacgttaa gggattttgg tcatgagatt atcaaaaagg atcttcacct agatcctttt
      6061 aaattaaaaa tgaagtttta aatcaatcta aagtatatat gagtaaactt ggtctgacag
      6121 ttaccaatgc ttaatcagtg aggcacctat ctcagcgatc tgtctatttc gttcatccat
      6181 agttgcctga ctccccgtcg tgtagataac tacgatacgg gagggcttac catctggccc
      6241 cagtgctgca atgataccgc gagacccacg ctcaccggct ccagatttat cagcaataaa
      6301 ccagccagcc ggaagggccg agcgcagaag tggtcctgca actttatccg cctccatcca
      6361 gtctattaat tgttgccggg aagctagagt aagtagttcg ccagttaata gtttgcgcaa
      6421 cgttgttgcc attgctacag gcatcgtggt gtcacgctcg tcgtttggta tggcttcatt
      6481 cagctccggt tcccaacgat caaggcgagt tacatgatcc cccatgttgt gcaaaaaagc
      6541 ggttagctcc ttcggtcctc cgatcgttgt cagaagtaag ttggccgcag tgttatcact
      6601 catggttatg gcagcactgc ataattctct tactgtcatg ccatccgtaa gatgcttttc
      6661 tgtgactggt gagtactcaa ccaagtcatt ctgagaatag tgtatgcggc gaccgagttg
      6721 ctcttgcccg gcgtcaatac gggataatac cgcgccacat agcagaactt taaaagtgct
      6781 catcattgga aaacgttctt cggggcgaaa actctcaagg atcttaccgc tgttgagatc
      6841 cagttcgatg taacccactc gtgcacccaa ctgatcttca gcatctttta ctttcaccag
      6901 cgtttctggg tgagcaaaaa caggaaggca aaatgccgca aaaaagggaa taagggcgac
      6961 acggaaatgt tgaatactca tactcttcct ttttcaatat tattgaagca tttatcaggg
      7021 ttattgtctc atgagcggat acatatttga atgtatttag aaaaataaac aaataggggt
      7081 tccgcgcaca tttccccgaa aagtgccac
 //
```

pBS ActEF AgroCP

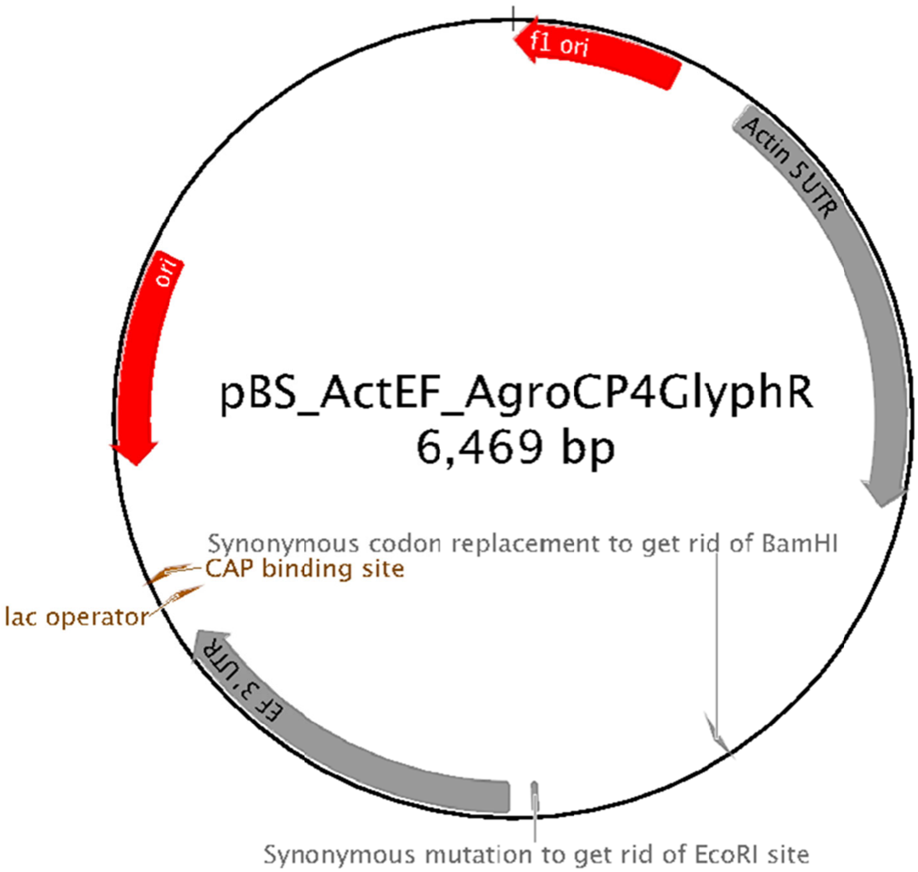


```
LOCUS       pBS_ActEF_AgroCP        6469 bp ds-DNA     circular SYN 26-OCT-2019
 DEFINITION  .
 ACCESSION
 VERSION     .
 KEYWORDS    pBS_Sr_ActEF_AgroCP4GlyphR.
 SOURCE      null
   ORGANISM
             .
 REFERENCE   1  (bases 1 to 6469)
   AUTHORS   Turkewitz and Lab
```

```
  TITLE     Direct Submission
  JOURNAL   Exported Monday, Jun 24, 2019 from SnapGene Viewer 4.3.8
            https://www.snapgene.com
FEATURES             Location/Qualifiers
     source          1..6469
                     /organism="recombinant plasmid"
                     /mol_type="other DNA"
     rep_origin      complement(3..458)
                     /direction="LEFT"
                     /note="f1 bacteriophage origin of replication; arrow
                     indicates direction of (+) strand synthesis"
                     /label="f1 ori"
     primer_bind     600..616
                     /note="common sequencing primer, one of multiple similar
                     variants"
                     /label="M13 fwd"
     promoter        626..644
                     /note="promoter for bacteriophage T7 RNA polymerase"
                     /label="T7 promoter"
     misc_feature    665..1864
                     /label="Actin 5'UTR"
     CDS             1871..3238
                     /codon_start=1
                     /translation="MSHGASSRPATARKSSGLSGTVRIPGDKSISHRSFMFGGLASGE
                     TRITGLLEGEDVINTGKAMQAMGARIRKEGDTWIIDGVGNGGLLAPEAPLDFGNAATG
                     CRLTMGLVGVYDFDSTFIGDASLTKRPMGRVLNPLREMGVQVKSEDGDRLPVTLRGPK
                     TPTPITYRVPMASAQVKSAVLLAGLNTPGITTVIEPIMTRDHTEKMLQGFGANLTVET
                     DADGVRTIRLEGRGKLTGQVIDVPGDPSSTAFPLVAALLVPGSDVTILNVLMNPTRTG
                     LILTLQEMGADIEVINPRLAGGEDVADLRVRSSTLKGVTVPEDRAPSMIDEYPILAVA
                     AAFAEGATVMNGLEELRVKESDRLSAVANGLKLNGVDCDEGETSLVVRGRPDGKGLGN
                     ASGAAVATHLDHRIAMSFLVMGLVSENPVTVDDATMIATSFPEFMDLMAGLGAKIELS
                     DTKAA"
                     /label="Agro CP4 AroA (GlyphR) Seminavis-Optimized"
     misc_feature    2651..2653
                     /label="Synonymous codon replacement to get rid of BamHI"
     misc_feature    3178
                     /label="Synonymous mutation to get rid of EcoRI site"
     misc_feature    3245..4244
                     /label="EF 3' UTR"
     promoter        complement(4281..4299)
                     /note="promoter for bacteriophage T3 RNA polymerase"
                     /label="T3 promoter"
     primer_bind     complement(4320..4336)
                     /note="common sequencing primer, one of multiple similar
                     variants"
                     /label="M13 rev"
     protein_bind    4344..4360
                     /bound_moiety="lac repressor encoded by lacI"
                     /note="The lac repressor binds to the lac operator to
                     inhibit transcription in E. coli. This inhibition can be
                     relieved by adding lactose or
                     isopropyl-beta-D-thiogalactopyranoside (IPTG)."
                     /label="lac operator"
     promoter        complement(4368..4398)
                     /note="promoter for the E. coli lac operon"
                     /label="lac promoter"
     protein_bind    4413..4434
                     /bound_moiety="E. coli catabolite activator protein"
                     /note="CAP binding activates transcription in the presence
                      of cAMP."
                     /label="CAP binding site"
     rep_origin      complement(4722..5310)
                     /direction="LEFT"
                     /note="high-copy-number ColE1/pMB1/pBR322/pUC origin of
                     replication"
                     /label="ori"
     CDS             complement(5481..6341)
                     /codon_start=1
                     /gene="bla"
                     /product="beta-lactamase"
                     /note="confers resistance to ampicillin, carbenicillin,
                     and related antibiotics"
                     /translation="MSIQHFRVALIPFFAAFCLPVFAHPETLVKVKDAEDQLGARVGY
                     IELDLNSGKILESFRPEERFPMMSTFKVLLCGAVLSRIDAGQEQLGRRIHYSQNDLVE
                     YSPVTEKHLTDGMTVRELCSAAITMSDNTAANLLLTTIGGPKELTAFLHNMGDHVTRL
                     DRWEPELNEAIPNDERDTTMPVAMATTLRKLLTGELLTLASRQQLIDWMEADKVAGPL
                     LRSALPAGWFIADKSGAGERGSRGIIAALGPDGKPSRIVVIYTTGSQATMDERNRQIA
```

```
                     EIGASLIKHW"
                     /label="AmpR"
     promoter        complement(6342..6446)
                     /gene="bla"
                     /label="AmpR promoter"
ORIGIN
        1 ctaaattgta agcgttaata ttttgttaaa attcgcgtta aatttttgtt aaatcagctc
       61 atttttaac caataggccg aaatcggcaa aatcccttat aaatcaaaag aatagaccga
      121 gatagggttg agtgttgttc cagtttggaa caagagtcca ctattaaaga acgtggactc
      181 caacgtcaaa gggcgaaaaa ccgtctatca gggcgatggc ccactacgtg aaccatcacc
      241 ctaatcaagt tttttggggt cgaggtgccg taaagcacta aatcggaacc ctaaagggag
      301 cccccgattt agagcttgac ggggaaagcc ggcgaacgtg gcgagaaagg aagggaagaa
      361 agcgaaagga gcgggcgcta gggcgctggc aagtgtagcg gtcacgctgc gcgtaaccac
      421 cacacccgcc gcgcttaatg cgccgctaca gggcgcgtcc cattcgccat tcaggctgcg
      481 caactgttgg gaagggcgat cggtgcgggc ctcttcgcta ttacgccagc tggcgaaagg
      541 gggatgtgct gcaaggcgat taagttgggt aacgccaggg ttttcccagt cacgacgttg
      601 taaaacgacg gccagtgagc gcgcgtaata cgactcacta tagggcgaat tgggtaccgg
      661 gcccagtttt acatcaagac tgcatacaag atatcgagaa acgcgatgtt ggctaatggt
      721 atctttgtga ttgcaaaaag ggtgaagtcg agaagacatg atgacggtac cgtacggtgc
      781 taccggtacg tacgaaagat cgcgacttca gtcttgcttg aaggtcacac cacgtatcgt
      841 ggaaaacaaa aagcttgcac cgagggcaag tatctgtaat gttctggaag tcgctccacg
      901 gaaaaggtat caggccttcg tcgcaggata agtcacatgg ctgatgatgg gccaagcaag
      961 ggtaaccttt ctctttcagc ctttgaagac tggtaagata agctggtctg tttcttcact
     1021 catgttgttg aagtcgctcc acggaaaagg tatcaggcct tcgtcgcagg ataagtcaca
     1081 tggctgatga tgggccaagc aagggtaacc tttctcttcc tttctctttc agcctttgaa
     1141 gactggtaag ataagctggt ctgtttcttc actcatgttg ctttcgacgc aaaagcataa
     1201 accaagctgc aacgtggaga ggctatccct cgccagtgcc aagttcgagg catccaaacg
     1261 ccccattctc aacctcagtg tccccaaagc attcttgaca tgacggttct tgccaagacc
     1321 gcttagctac tggtacagct acagcctttg tttcggcgtt gtctgacgcg acggaagcgg
     1381 gtgggcttct caatatcttt tgtcgatgtt cgatgagcca gtcaaaaaag cgtgcatacc
     1441 gaggataatg cttggtacca gcaaaatgtg aggtggtccc tcgttgcagc ccgtgaagcg
     1501 aaagtcacaa tgactggaat gccaaagggc ggaatccaaa ccaaacgaaa tttgaaaaag
     1561 ttggacctga aagatctata agatacgatt ccttctgcgt aaagatactt aacagtacaa
     1621 gcaggacttc gtttcgagaa gggaattcca atgttttcct cccaagacca aacaaagatc
     1681 tgccgcattc aagagagggg tcttgttggg acatctgcct tgacactgca cagcgggagg
     1741 ttgcagcgac agcgcagtga acccattcaa caagattgct tggattctgc cataaaattg
     1801 cttctcggta gcaacactct ctttctcgaa actctcttca aagaatcaaa caaactaatt
     1861 caccctcgag atgtcccacg gagcctcctc ccgtcccgcc accgcccgta agtcctccgg
     1921 attgtccgga accgtccgta ttcccggaga taagtccatt tcccaccgtt ccttcatgtt
     1981 cggaggattg gcctccggag aaacccgtat taccggattg ttggaaggag aagatgtcat
     2041 taacaccgga aaggccatgc aagccatggg agcccgtatt cgtaaggaag gagatacctg
     2101 gattattgat ggagtcggaa acggaggatt gttggccccc gaagcccccct tggatttcgg
     2161 aaacgccgcc accggatgcc gtttgaccat gggattggtc ggagtctacg atttcgattc
     2221 caccttcatt ggagatgcct ccttgaccaa gcgtcccatg ggacgtgtct tgaacccctt
     2281 gcgtgaaatg ggagtccaag tcaagtccga agatggagat cgtttgcccg tcaccttgcg
     2341 tggacccaag acccccaccc ccattaccta ccgtgtcccc atggcctccg cccaagtcaa
     2401 gtccgccgtc ttgttggccg gattgaacac ccccggaatt accaccgtca ttgaacccat
     2461 tatgacccgt gatcacaccg aaaagatgtt gcaaggattc ggagccaact tgaccgtcga
     2521 aaccgatgcc gatggagtcc gtaccattcg tttggaagga cgtggaaagt tgaccggaca
     2581 agtcattgat gtccccggag atccctcctc caccgccttc cccttggtcg ccgccttgtt
     2641 ggtccccgga agtgatgtca ccattttgaa cgtcttgatg aaccccaccc gtaccggatt
     2701 gattttgacc ttgcaagaaa tgggagccga tattgaagtc attaaccccc gtttggccgg
     2761 aggagaagat gtcgccgatt tgcgtgtccg ttcctccacc ttgaagggag tcaccgtccc
     2821 cgaagatcgt gccccctcca tgattgatga ataccccatt ttggccgtcg ccgccgcctt
     2881 cgccgaagga gccaccgtca tgaacggatt ggaagaattg cgtgtcaagg aatccgatcg
     2941 tttgtccgcc gtcgccaacg gattgaagtt gaacggagtc gattgcgatg aaggagaaac
     3001 ctccttggtc gtccgtggac gtcccgatgg aaagggattg ggaaacgcct ccggagccgc
     3061 cgtcgccacc cacttggatc accgtattgc catgtccttc ttggtcatgg gattggtctc
     3121 cgaaaacccc gtcaccgtcg atgatgccac catgattgcc acctccttcc ccgaatttat
     3181 ggatttgatg gccggattgg gagccaagat tgaattgtcc gataccaagg ccgcctaact
     3241 gcagtcccaa tctgttctta ccaaaaacaa aaaaacacat aatacaatgg agactgctgt
     3301 ggatggatct tctggtgcgt attgagagtt gtgtgattga tgacaaggag acaactcgac
     3361 gaaactcgga tgcattccaa agagtagtag tataatacat gtataagacc aagaactaaa
     3421 aaagggattt acaatacgat acttctagac tagctagcta gtgtaggatt taatctagac
     3481 gccacaggag ggttcgtgtg ttgtctctgc aaaggtttgg tgctgtcggt tgcttggttc
     3541 ggcgttgcat agatcctagc aacagttcca actctcagtc atcgtgcaga gtctagctag
     3601 agtagagaag ttatgacgga cgcttggctg cctagctagt cgagctagac tagtaggcgt
     3661 cgccaggaca cagccaaggt gaggaaacgg agatagctga ctagccagtc tttagaagca
     3721 gagtgagata gctagctaga ggagtcaaag gtgtcacgga atcctgtttt attgcatggc
     3781 gtctcgatcc cgctgacgcc gagccatgga tgggcgacgg aggcagtttc tctacggtca
     3841 caaggggcag catggaatgt ttggtttact cgactccctc catcgagata caccagccgc
     3901 tcaagaacca tgtatacatt atagccaagc aatgcttcgt cttggtggtg gtggtggcac
     3961 ggccgcttga tggttcggtg catccaatca tggagacgtg tggacgttgg tcgtcctatt
     4021 tgctgcttcc agagcgacct ggcaaccagc cacgtggctt gtctttgagc tagaggagtt
     4081 ttggatggcc gaatgactgc ggctttgctt gatcacccaa tgcacgccaa agaaaacagt
     4141 cgctctatct gtatgtgcag attgcttgct tctgccgttc gtctgcctcc tttgtccgtg
     4201 cacactccat catgcataga accaatgctg aagcagcagg aagagcggcc gccaccgcgg
```

```
4261 tggagctcca gcttttgttc cctttagtga gggttaattg cgcgcttggc gtaatcatgg
4321 tcatagctgt ttcctgtgtg aaattgttat ccgctcacaa ttccacacaa catacgagcc
4381 ggaagcataa agtgtaaagc ctggggtgcc taatgagtga gctaactcac attaattgcg
4441 ttgcgctcac tgcccgcttt ccagtcggga aacctgtcgt gccagctgca ttaatgaatc
4501 ggccaacgcg cggggagagg cggtttgcgt attgggcgct cttccgcttc ctcgctcact
4561 gactcgctgc gctcggtcgt tcggctgcgg cgagcggtat cagctcactc aaaggcggta
4621 atacggttat ccacagaatc aggggataac gcaggaaaga acatgtgagc aaaaggccag
4681 caaaaggcca ggaaccgtaa aaaggccgcg ttgctggcgt ttttccatag gctccgcccc
4741 cctgacgagc atcacaaaaa tcgacgctca agtcagaggt ggcgaaaccc gacaggacta
4801 taaagatacc aggcgtttcc ccctggaagc tccctcgtgc gctctcctgt tccgaccctg
4861 ccgcttaccg gatacctgtc cgcctttctc ccttcgggaa gcgtggcgct ttctcatagc
4921 tcacgctgta ggtatctcag ttcggtgtag gtcgttcgct ccaagctggg ctgtgtgcac
4981 gaaccccccg ttcagcccga ccgctgcgcc ttatccggta actatcgtct tgagtccaac
5041 ccggtaagac acgacttatc gccactggca gcagccactg gtaacaggat tagcagagcg
5101 aggtatgtag gcggtgctac agagttcttg aagtggtggc ctaactacgg ctacactaga
5161 aggacagtat ttggtatctg cgctctgctg aagccagtta ccttcggaaa aagagttggt
5221 agctcttgat ccggcaaaca aaccaccgct ggtagcggtg gtttttttgt ttgcaagcag
5281 cagattacgc gcagaaaaaa aggatctcaa gaagatcctt tgatcttttc tacggggtct
5341 gacgctcagt ggaacgaaaa ctcacgttaa gggattttgg tcatgagatt atcaaaaagg
5401 atcttcacct agatcctttt aaattaaaaa tgaagtttta aatcaatcta aagtatatat
5461 gagtaaactt ggtctgacag ttaccaatgc ttaatcagtg aggcacctat ctcagcgatc
5521 tgtctatttc gttcatccat agttgcctga ctccccgtcg tgtagataac tacgatacgg
5581 gagggcttac catctggccc cagtgctgca atgataccgc gagacccacg ctcaccggct
5641 ccagatttat cagcaataaa ccagccagcc ggaagggccg agcgcagaag tggtcctgca
5701 actttatccg cctccatcca gtctattaat tgttgccggg aagctagagt aagtagttcg
5761 ccagttaata gtttgcgcaa cgttgttgcc attgctacag gcatcgtggt gtcacgctcg
5821 tcgtttggta tggcttcatt cagctccggt tcccaacgat caaggcgagt tacatgatcc
5881 cccatgttgt gcaaaaaagc ggttagctcc ttcggtcctc cgatcgttgt cagaagtaag
5941 ttggccgcag tgttatcact catggttatg gcagcactgc ataattctct tactgtcatg
6001 ccatccgtaa gatgcttttc tgtgactggt gagtactcaa ccaagtcatt ctgagaatag
6061 tgtatgcggc gaccgagttg ctcttgcccg gcgtcaatac gggataatac cgcgccacat
6121 agcagaactt taaaagtgct catcattgga aaacgttctt cggggcgaaa actctcaagg
6181 atcttaccgc tgttgagatc cagttcgatg taacccactc gtgcacccaa ctgatcttca
6241 gcatctttta ctttcaccag cgtttctggg tgagcaaaaa caggaaggca aaatgccgca
6301 aaaaagggaa taagggcgac acggaaatgt tgaatactca tactcttcct ttttcaatat
6361 tattgaagca tttatcaggg ttattgtctc atgagcggat acatatttga atgtatttag
6421 aaaaataaac aaataggggt tccgcgcaca tttccccgaa aagtgccac
//
```

pBS_Act_PurR

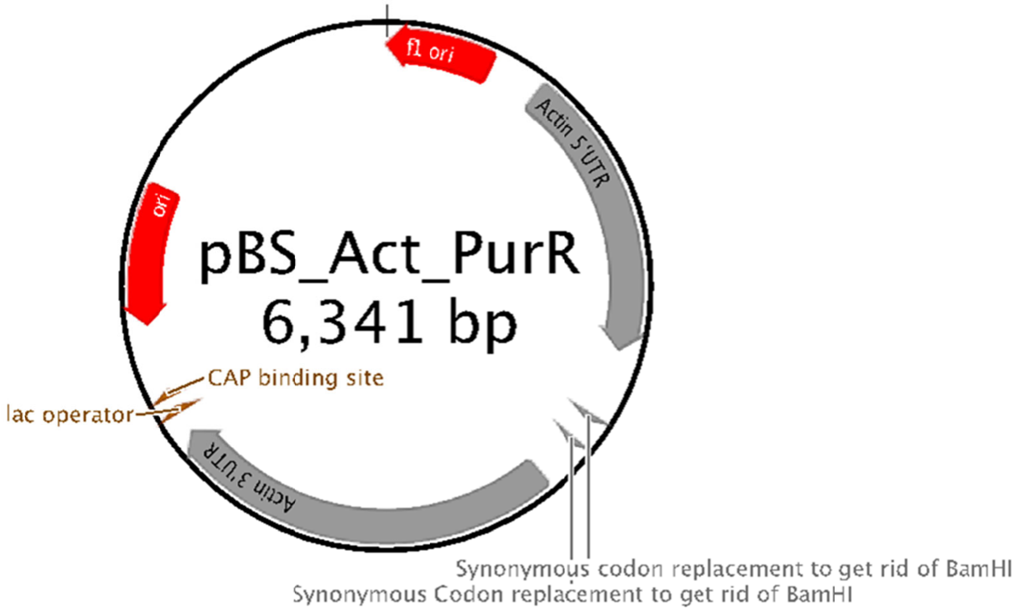

```
LOCUS       pBS_Act_PurR            6341 bp ds-DNA
 DEFINITION  .
 ACCESSION
 VERSION     .
 KEYWORDS    pBS_Sr_Act_PurR.
 SOURCE      null
   ORGANISM
             .
 REFERENCE   1  (bases 1 to 6341)
   AUTHORS   Turkewitz and Lab
   TITLE     Direct Submission
   JOURNAL   Exported Monday, Jun 24, 2019 from SnapGene Viewer 4.3.8
             https://www.snapgene.com
 FEATURES             Location/Qualifiers
      source          1..6341
                      /organism="recombinant plasmid"
                      /mol_type="other DNA"
      rep_origin      complement(3..458)
                      /direction="LEFT"
                      /note="f1 bacteriophage origin of replication; arrow
                      indicates direction of (+) strand synthesis"
                      /label="f1 ori"
      primer_bind     600..616
                      /note="common sequencing primer, one of multiple similar
                      variants"
                      /label="M13 fwd"
      promoter        626..644
                      /note="promoter for bacteriophage T7 RNA polymerase"
                      /label="T7 promoter"
      misc_feature    665..1864
                      /label="Actin 5'UTR"
      CDS             1871..2470
                      /codon_start=1
                      /translation="MTEYKPTVRLATRDDVPRAVRTLAAAFADYPATRHTVDPDRHIE
                      RVTELQELFLTRVGLDIGKVWVADDGAAVAVWTTPESVEAGAVFAEIGPRMAELSGSR
                      LAAQQQMEGLLAPHRPKEPAWFLATVGVSPDHQGKGLGSAVVLPGVEAAERAGVPAFL
                      ETSAPRNLPFYERLGFTVTADVEVPEGPRTWCMTRKPGA"
                      /label="PurR Seminavis-Optimized"
      primer_bind     2009..2043
                      /note="IDT Tm = 65.7"
                      /label="PurR RTfwd2"
      primer_bind     complement(2135..2163)
                      /note="IDT Tm = 65.9"
                      /label="PurR RTrev"
      misc_feature    2171..2173
                      /label="Synonymous codon replacement to get rid of BamHI"
      misc_feature    2291..2293
                      /label="Synonymous Codon replacement to get rid of BamHI"
      primer_bind     2385..2410
                      /note="IDT Tm = 65.2"
                      /label="PurR RTfwd1"
      misc_feature    2477..4116
                      /label="Actin 3'UTR"
      primer_bind     complement(2477..2500)
                      /note="IDT Tm = 65.1C"
                      /label="ACT RTrev"
      promoter        complement(4153..4171)
                      /note="promoter for bacteriophage T3 RNA polymerase"
                      /label="T3 promoter"
      primer_bind     complement(4192..4208)
                      /note="common sequencing primer, one of multiple similar
                      variants"
                      /label="M13 rev"
      protein_bind    4216..4232
                      /bound_moiety="lac repressor encoded by lacI"
                      /note="The lac repressor binds to the lac operator to
                      inhibit transcription in E. coli. This inhibition can be
                      relieved by adding lactose or
                      isopropyl-beta-D-thiogalactopyranoside (IPTG)."
                      /label="lac operator"
      promoter        complement(4240..4270)
                      /note="promoter for the E. coli lac operon"
                      /label="lac promoter"
      protein_bind    4285..4306
                      /bound_moiety="E. coli catabolite activator protein"
                      /note="CAP binding activates transcription in the presence
                       of cAMP."
```

```
                     /label="CAP binding site"
     rep_origin      complement(4594..5182)
                     /direction="LEFT"
                     /note="high-copy-number ColE1/pMB1/pBR322/pUC origin of
                     replication"
                     /label="ori"
     CDS             complement(5353..6213)
                     /codon_start=1
                     /gene="bla"
                     /product="beta-lactamase"
                     /note="confers resistance to ampicillin, carbenicillin,
                     and related antibiotics"
                     /translation="MSIQHFRVALIPFFAAFCLPVFAHPETLVKVKDAEDQLGARVGY
                     IELDLNSGKILESFRPEERFPMMSTFKVLLCGAVLSRIDAGQEQLGRRIHYSQNDLVE
                     YSPVTEKHLTDGMTVRELCSAAITMSDNTAANLLLTTIGGPKELTAFLHNMGDHVTRL
                     DRWEPELNEAIPNDERDTTMPVAMATTLRKLLTGELLTLASRQQLIDWMEADKVAGPL
                     LRSALPAGWFIADKSGAGERGSRGIIAALGPDGKPSRIVVIYTTGSQATMDERNRQIA
                     EIGASLIKHW"
                     /label="AmpR"
     promoter        complement(6214..6318)
                     /gene="bla"
                     /label="AmpR promoter"
ORIGIN
        1 ctaaattgta agcgttaata ttttgttaaa attcgcgtta aatttttgtt aaatcagctc
       61 atttttaac caataggccg aaatcggcaa aatcccttat aaatcaaaag aatagaccga
      121 gatagggttg agtgttgttc cagtttggaa caagagtcca ctattaaaga acgtggactc
      181 caacgtcaaa gggcgaaaaa ccgtctatca gggcgatggc ccactacgtg aaccatcacc
      241 ctaatcaagt tttttggggt cgaggtgccg taaagcacta aatcggaacc ctaaagggag
      301 cccccgattt agagcttgac ggggaaagcc ggcgaacgtg gcgagaaagg aagggaagaa
      361 agcgaaagga gcgggcgcta gggcgctggc aagtgtagcg gtcacgctgc gcgtaaccac
      421 cacacccgcc gcgcttaatg cgccgctaca gggcgcgtcc cattcgccat tcaggctgcg
      481 caactgttgg gaagggcgat cggtgcgggc ctcttcgcta ttacgccagc tggcgaaagg
      541 gggatgtgct gcaaggcgat taagttgggt aacgccaggg ttttcccagt cacgacgttg
      601 taaaacgacg gccagtgagc gcgcgtaata cgactcacta tagggcgaat tgggtaccgg
      661 gcccagtttt acatcaagac tgcatacaag atatcgagaa acgcgatgtt ggctaatggt
      721 atctttgtga ttgcaaaaag ggtgaagtcg agaagacatg atgacggtac cgtacggtgc
      781 taccggtacg tacgaaagat cgcgacttca gtcttgcttg aaggtcacac cacgtatcgt
      841 ggaaaacaaa aagcttgcac cgagggcaag tatctgtaat gttctggaag tcgctccacg
      901 gaaaaggtat caggccttcg tcgcaggata agtcacatgg ctgatgatgg gccaagcaag
      961 ggtaaccttt ctctttcagc ctttgaagac tggtaagata agctggtctg tttcttcact
     1021 catgttgttg aagtcgctcc acggaaaagg tatcaggcct tcgtcgcagg ataagtcaca
     1081 tggctgatga tgggccaagc aagggtaacc tttctcttcc tttctctttc agcctttgaa
     1141 gactggtaag ataagctggt ctgtttcttc actcatgttg ctttcgacgc aaaagcataa
     1201 accaagctgc aacgtggaga ggctatccct cgccagtgcc aagttcgagg catccaaacg
     1261 ccccattctc aacctcagtg tccccaaagc attcttgaca tgacggttct tgccaagacc
     1321 gcttagctac tggtacagct acagcctttg tttcggcgtt gtctgacgcg acggaagcgg
     1381 gtgggcttct caatatcttt tgtcgatgtt cgatgagcca gtcaaaaaag cgtgcatacc
     1441 gaggataatg cttggtacca gcaaaatgtg aggtggtccc tcgttgcagc ccgtgaagcg
     1501 aaagtcacaa tgactggaat gccaaagggc ggaatccaaa ccaaacgaaa tttgaaaaag
     1561 ttggacctga aagatctata agatacgatt ccttctgcgt aaagatactt aacagtacaa
     1621 gcaggacttc gtttcgagaa gggaattcca atgttttcct cccaagacca aacaaagatc
     1681 tgccgcattc aagagagggg tcttgttggg acatctgcct tgacactgca cagcgggagg
     1741 ttgcagcgac agcgcagtga acccattcaa caagattgct tggattctgc cataaaattg
     1801 cttctcggta gcaacactct ctttctcgaa actctcttca aagaatcaaa caaactaatt
     1861 caccctcgag atgaccgaat acaagcccac cgtccgtttg gccacccgtg atgatgtccc
     1921 ccgtgccgtc cgtaccttgg ccgccgcctt cgccgattac cccgccaccc gtcacaccgt
     1981 cgatcccgat cgtcacattg aacgtgtcac cgaattgcaa gaattgttct tgacccgtgt
     2041 cggattggat attggaaagg tctgggtcgc cgatgatgga gccgccgtcg ccgtctggac
     2101 cacccccgaa tccgtcgaag ccggagccgt cttcgccgaa attggacccc gtatggccga
     2161 attgtccgga agtcgtttgg ccgcccaaca acaaatggaa ggattgttgg ccccccaccg
     2221 tcccaaggaa cccgcctggt tcttggccac cgtcggagtc tcccccgatc accaaggaaa
     2281 gggattggga agtgccgtcg tcttgcccgg agtcgaagcc gccgaacgtg ccggagtccc
     2341 cgccttcttg gaaacctccg ccccccgtaa cttgcccttc tacgaacgtt tgggattcac
     2401 cgtcaccgcc gatgtcgaag tccccgaagg accccgtacc tggtgcatga cccgtaagcc
     2461 cggagcctaa ctgcagggaa tgggccatgc gcaggcaaat gccagctgct gtgtctgctc
     2521 tttgtgaaat attattgggc cagcagtctt ttattttttg ttggacctgg gttgcttgcc
     2581 gttgttgaac aacatggtga tgtgaccgac ggatcaatga gaaaagccta attgttagaa
     2641 tgttttattc attgcggtcc cttgtctcta catagctagc tagctaacta gctagttttc
     2701 gactcctgat atcaaggtat tcccgtcgtt gctggagccg atcttgccaa gccctggcaa
     2761 ccgcaccgta ccgattgctg gggggctgta ggtacacgta ctggaagcaa cgttgatcga
     2821 ttggcttttc gggcaactgg agctgtgcga cacgatcttt gatagctttt ccttagggtt
     2881 taaccctaaa atattatttt gctggtttgt gcccacagcg agttcgattt agtcgaatct
     2941 tttttgggct cgatcggatc catccatctt cgaagatctt tcttttgcac gccattgatt
     3001 ctctcaaacg ttctcgctcg ttggtcctca ataaccatga tgaacaacgc gttgctaccg
     3061 attcgatcca gccttcccaa ggctatcaga agaacctcgt gtcgtcttat tcgatcaggt
     3121 ccgccttctc gcttggtatc gcccatcagg caagcatttc ctgcatcaca gagtgctttt
     3181 gctttgcgtc ctttttcgag cggcgcccag acgctttctg atgtgctggc ccgagagtta
```

3241 gccgaagaaa ccgaggaagg tcgagaccaa atccccgaag aacttgctga gcttaagagt
3301 tctattgaga aagattggaa ggtcgttgac gatggtgcga tcactcggtt gatccgatcg
3361 attggagcca gtaaagtggt tgtctcgttt cactgccaag acacggtgga tggcgcagag
3421 gattatgtcg aggaagaaga gggagaagaa agaacctgcc ttgccctttc gatttgagat
3481 tcttgtgtcc aaggcaggaa atacattggt gcttaattgt atcagcaatg ctggtgaaac
3541 cacagtcgat ggggttgcca tgaccactga agacattgaa tcggttcaag ccaatggtat
3601 tggtcgcaac aattatcagg gacctgaatt tcaagagcta gccgaggacc tacaggaggc
3661 acttcatgaa tatgtctttt cggagctagg tatcgatgaa gacgtctctg cctttgtctc
3721 gatgtatgct gattacaagg agcaagtgca atacatcgga ttcttgacga atgtttctaa
3781 ggttcttccg taagattcat gctccctctt aagtctgaac agacataatt gaaaatttag
3841 ctagaaccaa tcaacgaata aaaaatagag actgatattg agcttgtttg gaagcgggtg
3901 ctttttggga aacagaactg ggtactcct cacacgaagc cataaaagag tcacgcaaca
3961 acggtatcga caacgacaaa ggcatcagct gtctcaacgt accccgcgca gctgatcctt
4021 gtacggggtt tgtatggttc tctcacgggt ggtacacaaa gggacgttca tgcagaccaa
4081 gcatatcgtc agttaccgcc actcagcagt tgtactgcgg ccgccaccgc ggtggagctc
4141 cagcttttgt tccctttagt gagggttaat tgcgcgcttg gcgtaatcat ggtcatagct
4201 gtttcctgtg tgaaattgtt atccgctcac aattccacac aacatacgag ccggaagcat
4261 aaagtgtaaa gcctggggtg cctaatgagt gagctaactc acattaattg cgttgcgctc
4321 actgcccgct ttccagtcgg gaaacctgtc gtgccagctg cattaatgaa tcggccaacg
4381 cgcggggaga ggcggtttgc gtattgggcg ctcttccgct tcctcgctca ctgactcgct
4441 gcgctcggtc gttcggctgc ggcgagcggt atcagctcac tcaaaggcgg taatacggtt
4501 atccacagaa tcaggggata acgcaggaaa gaacatgtga gcaaaaggcc agcaaaaggc
4561 caggaaccgt aaaaaggccg cgttgctggc gtttttccat aggctccgcc cccctgacga
4621 gcatcacaaa aatcgacgct caagtcagag gtggcgaaac ccgacaggac tataaagata
4681 ccaggcgttt ccccctggaa gctccctcgt gcgctctcct gttccgaccc tgccgcttac
4741 cggatacctg tccgcctttc tcccttcggg aagcgtggcg ctttctcata gctcacgctg
4801 taggtatctc agttcggtgt aggtcgttcg ctccaagctg ggctgtgtgc acgaaccccc
4861 cgttcagccc gaccgctgcg ccttatccgg taactatcgt cttgagtcca acccggtaag
4921 acacgactta tcgccactgg cagcagccac tggtaacagg attagcagag cgaggtatgt
4981 aggcggtgct acagagttct tgaagtggtg gcctaactac ggctacacta gaaggacagt
5041 atttggtatc tgcgctctgc tgaagccagt taccttcgga aaaagagttg gtagctcttg
5101 atccggcaaa caaaccaccg ctggtagcgg tggttttttt gtttgcaagc agcagattac
5161 gcgcagaaaa aaaggatctc aagaagatcc tttgatcttt tctacggggt ctgacgctca
5221 gtggaacgaa aactcacgtt aagggatttt ggtcatgaga ttatcaaaaa ggatcttcac
5281 ctagatcctt ttaaattaaa aatgaagttt taaatcaatc taaagtatat atgagtaaac
5341 ttggtctgac agttaccaat gcttaatcag tgaggcacct atctcagcga tctgtctatt
5401 tcgttcatcc atagttgcct gactccccgt cgtgtagata actacgatac gggagggctt
5461 accatctggc cccagtgctg caatgatacc gcgagaccca cgctcaccgg ctccagattt
5521 atcagcaata aaccagccag ccggaagggc cgagcgcaga agtggtcctg caactttatc
5581 cgcctccatc cagtctatta attgttgccg ggaagctaga gtaagtagtt cgccagttaa
5641 tagtttgcgc aacgttgttg ccattgctac aggcatcgtg gtgtcacgct cgtcgtttgg
5701 tatggcttca ttcagctccg gttcccaacg atcaaggcga gttacatgat cccccatgtt
5761 gtgcaaaaaa gcggttagct ccttcggtcc tccgatcgtt gtcagaagta agttggccgc
5821 agtgttatca ctcatggtta tggcagcact gcataattct cttactgtca tgccatccgt
5881 aagatgcttt tctgtgactg gtgagtactc aaccaagtca ttctgagaat agtgtatgcg
5941 gcgaccgagt tgctcttgcc cggcgtcaat acgggataat accgcgccac atagcagaac
6001 tttaaaagtg ctcatcattg gaaaacgttc ttcggggcga aaactctcaa ggatcttacc
6061 gctgttgaga tccagttcga tgtaacccac tcgtgcaccc aactgatctt cagcatcttt
6121 tactttcacc agcgtttctg ggtgagcaaa aacaggaagg caaaatgccg caaaaaaggg
6181 aataagggcg acacggaaat gttgaatact catactcttc ctttttcaat attattgaag
6241 catttatcag ggttattgtc tcatgagcgg atacatattt gaatgtattt agaaaaataa
6301 acaaataggg gttccgcgca catttccccg aaaagtgcca c
//

#### *Pseudo-nitzschia multiseries*

pPmAGFPC10 EPI

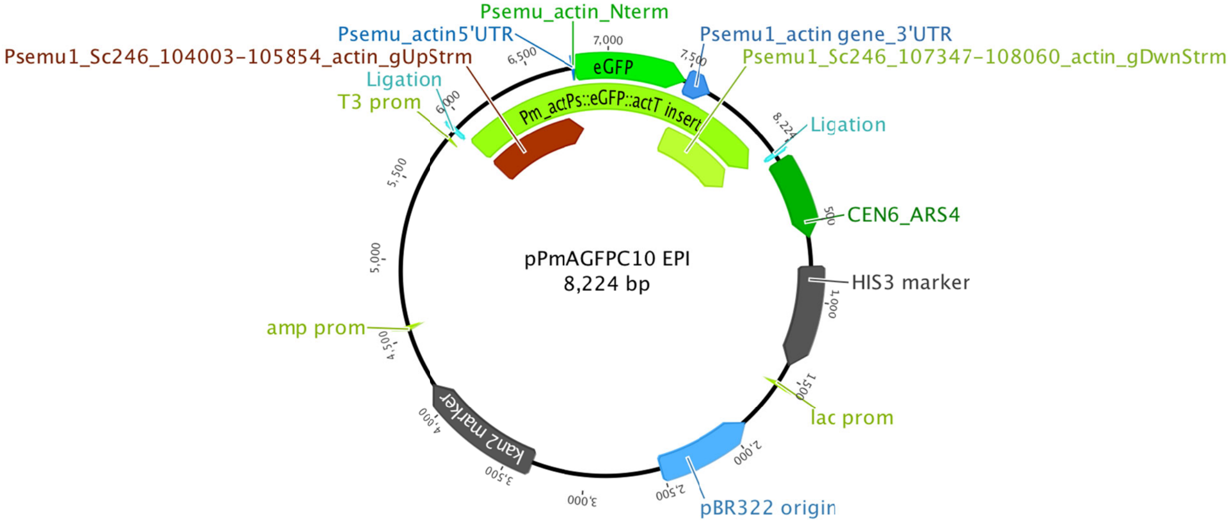


```
LOCUS       pPmAGFPC10_EPI          8224 bp    DNA     circular UNA 01-OCT-2019
DEFINITION  GOOD:Gibson Assembly of Pm_actPs::eGFP::actT insert into pPtPuc3
            PCR_GAcompatibleBackbone.
SOURCE      Pseudo-nitzschia multiseries 15091C3
  ORGANISM  Pseudo-nitzschia multiseries 15091C3
            .
FEATURES             Location/Qualifiers
     ligation        6..35
                     /X-Transferred_From="Pmu_actPsT_shble_del"
                     /X-Transferred_Similarity="100.00%"
                     /Transferred_From="Pmu_actPsT_shble, Pmu_actPsT_shble_del,
                     backup of pPm_actPs_eGFP_actT_expression cassette ,
                     pPm_actPs_eGFP_actT_consensus,
                     pPm_actPs_eGFP_actT_expression cassette ,
                     pPm_actPs_eGFP_actT_expression cassette_new"
                     /Transferred_Similarity="100.00%"
                     /label="Ligation"
     gene            72..599

                     /label="CEN6_ARS4"
     ORF             780..1442


                     /label="HIS3 marker"
     promoter        complement(1560..1589)
                     /label="lac prom"
     misc_feature    complement(1898..2517)
                     /label="pBR322 origin"
                     /note="Geneious type: origin of replication"
     marker          3349..4164
                     /label="kan2 marker"
     promoter        complement(4544..4572)
                     /label="amp prom"
     promoter        5840..5859
                     /label="T3 prom"
     ligation        5896..5925
                     /label="Ligation"
     misc_feature    5932..8224
                     /label="Psemu1_Sc246_104003-105854_actin_gUpStrm"
     misc_feature    5935
                     /Original_Bases
                     /label
     5'UTR           6778..6781
                     /created_by="User"
                     /label="Psemu_actin5'UTR"
     CDS             6782..6790
                     /created_by="User"
```

```
                     /label="Psemu_actin_Nterm"
     CDS             6791..7510
                     /label="eGFP"
     misc_feature    6983..6994
                     /created_by="User"
                     /Transferred_From="eGFP_Pmact_vector_seq"
                     /Transferred_Similarity="100.00%"
                     /label="eGFP modified chromaphore domain"
     3'utr           7511..7684
                     /label="Psemu1_Sc246_107347-108060_actin_gDwnStrm"
ORIGIN
        1 ctcgaggaag tttgtctaga tctcaggcgt ggatgcactc cgctttggtt tcacagtcag
       61 gaataacact agctcgtctt cagcgagcat cacgtgctat aaaaataatt ataatttaaa
      121 ttttttaata taaatatata aattaaaaat agaaagtaaa aaaagaaatt aaagaaaaaa
      181 tagtttttgt tttccgaaga tgtaaaagac tctaggggga tcgccaacaa atactacctt
      241 ttaccttgct cttcctgctc tcaggtatta atgccgaatt gtttcatctt gtctgtgtag
      301 aagaccacac acgaaaatcc tgtgatttta cattttactt atcgttaatc gaatgtatat
      361 ctatttaatc tgcttttctt gtctaataaa tatatatgta aagtacgctt tttgttgaaa
      421 ttttttaaac ctttgtttat ttttttttct tcattccgta actcttctac cttctttatt
      481 tactttctaa aatccaaata caaaacataa aaataaataa acacagagta aattcccaaa
      541 ttattccatc attaaaagat acgaggcgcg tgtaagttac aggcaagcga tcctagtaca
      601 ctctatattt ttttatgcct cggtaatgat tttcattttt tttttccacc tagcggatga
      661 ctcttttttt ttcttagcga ttggcattat cacataatga attatacatt atataaagta
      721 atgtgatttc ttcgaagaat atactaaaaa atgagcaggc aagataaacg aaggcaaaga
      781 tgacagagca gaaagcccta gtaaagcgta ttacaaatga aaccaagatt cagattgcga
      841 tctctttaaa gggtggtccc ctagcgatag agcactcgat cttcccagaa aaagaggcag
      901 aagcagtagc agaacaggcc acacaatcgc aagtgattaa cgtccacaca ggtatagggt
      961 ttctggacca tatgatacat gctctggcca agcattccgg ctggtcgcta atcgttgagt
     1021 gcattggtga cttacacata gacgaccatc acaccactga agactgcggg attgctctcg
     1081 gtcaagcttt taaagaggcc ctagggggccg tgcgtggagt aaaaaggttt ggatcaggat
     1141 ttgcgccttt ggatgaggca ctttccagag cggtggtaga tctttcgaac aggccgtacg
     1201 cagttgtcga acttggtttg caaagggaga aagtaggaga tctctcttgc gagatgatcc
     1261 cgcattttct tgaaagcttt gcagaggcta gcagaattac cctccacgtt gattgtctgc
     1321 gaggcaagaa tgatcatcac cgtagtgaga gtgcgttcaa ggctcttgcg gttgccataa
     1381 gagaagccac ctcgcccaat ggtaccaacg atgttccctc caccaaaggt gttcttatgt
     1441 agttttacac aggagtctgg acttgacctc tagagtcgac ctgcaggcat gcaagcttgg
     1501 cgtaatcatg gtcatagctg tttcctgtgt gaaattgtta tccgctcaca attccacaca
     1561 acatacgagc cggaagcata aagtgtaaag cctggggtgc ctaatgagtg agctaactca
     1621 cattaattgc gttgcgctca ctgcccgctt tccagtcggg aaacctgtcg tgccagctgc
     1681 attaatgaat cggccaacgc gcggggagag gcggtttgcg tattgggcgc tcttccgctt
     1741 cctcgctcac tgactcgctg cgctcggtcg ttcggctgcg gcgagcggta tcagctcact
     1801 caaaggcggt aatacggtta tccacagaat caggggataa cgcaggaaag aacatgtgag
     1861 caaaaggcca gcaaaaggcc aggaaccgta aaaaggccgc gttgctggcg tttttccata
     1921 ggctccgccc ccctgacgag catcacaaaa atcgacgctc aagtcagagg tggcgaaacc
     1981 cgacaggact ataaagatac caggcgtttc cccctggaag ctccctcgtg cgctctcctg
     2041 ttccgaccct gccgcttacc ggatacctgt ccgcctttct cccttcggga agcgtggcgc
     2101 tttctcatag ctcacgctgt aggtatctca gttcggtgta ggtcgttcgc tccaagctgg
     2161 gctgtgtgca cgaacccccc gttcagcccg accgctgcgc cttatccggt aactatcgtc
     2221 ttgagtccaa cccggtaaga cacgacttat cgccactggc agcagccact ggtaacagga
     2281 ttagcagagc gaggtatgta ggcggtgcta cagagttctt gaagtggtgg cctaactacg
     2341 gctacactag aagaacagta tttggtatct gcgctctgct gaagccagtt accttcggaa
     2401 aaagagttgg tagctcttga tccggcaaac aaaccaccgc tggtagcggt ggtttttttg
     2461 tttgcaagca gcagattacg cgcagaaaaa aaggatctca agaagatcct ttgatctttt
     2521 ctacggggtc tgacgctcag tggaacgaaa actcacgtta agggattttg gtcatgagat
     2581 tatcaaaaag gatcttcacc tagatccttt taaattaaaa atgaagtttt aaatcaatct
     2641 aaagtatata tgagtaaact tggtctgaca gttaccaatg cttaatcagt gaggcaccta
     2701 tctcagcgat ctgtctattt cgttcatcca tagttgcctg actccccgtc gtgtagataa
     2761 ctacgatacg ggagggctta ccatctggcc ccagtgctgc aatgataccg cgagacccac
     2821 gctcaccggc tccagattta tcagcaataa accagccagc cggaagggcc gagcgcagaa
     2881 gtggtcctgc aactttatcc gcctccatcc agtctattaa ttgttgccgg gaagctagag
     2941 taagtagttc gccagttaat agtttgcgca acgttgttgc cattgctaca ggcatcgtgg
     3001 tgtcacgctc gtcgtttggt atggcttcat tcagctccgg ttcccaacga tcaaggcgag
     3061 ttacatgatc ccccatgttg tgcaaaaaag cggttagctc cttcggtcct ccgatcgttg
     3121 tcagaagtaa gttggccgca gtgttatcac tcatggttat ggcagcactg cataattctc
     3181 ttactgtcat gccatccgta agatgctttt ctgtgactgg tgagtcgatt tattcaacaa
     3241 agccacgttg tgtctcaaaa tctctgatgt tacattgcac aagataaaaa tatatcatca
     3301 tgaacaataa aactgtctgc ttacataaac agtaatacaa ggggtgttat gagccatatt
     3361 caacgggaaa cgtcttgctc gaggccgcga ttaaattcca acatggatgc tgatttatat
     3421 gggtataaat gggctcgcga taatgtcggg caatcaggtg cgacaatcta tcgattgtat
     3481 gggaagcccg atgcgccaga gttgtttctg aaacatggca aaggtagcgt tgccaatgat
     3541 gttacagatg agatggtcag actaaactgg ctgacggaat ttatgcctct tccgaccatc
     3601 aagcatttta tccgtactcc tgatgatgca tggttactca ccactgcgat ccccgggaaa
     3661 acagcattcc aggtattaga agaatatcct gattcaggtg aaaatattgt tgatgcgctg
     3721 gcagtgttcc tgcgccggtt gcattcgatt cctgtttgta attgtccttt taacagcgat
     3781 cgcgtatttc gtctcgctca ggcgcaatca cgaatgaata acggtttggt tgatgcgagt
     3841 gattttgatg acgagcgtaa tggctggcct gttgaacaag tctggaaaga aatgcataag
     3901 cttttgccat tctcaccgga ttcagtcgtc actcatggtg atttctcact tgataacctt
```

3961 atttttgacg aggggaaatt aataggttgt attgatgttg gacgagtcgg aatcgcagac
4021 cgataccagg atcttgccat cctatggaac tgcctcggtg agttttctcc ttcattacag
4081 aaacggcttt ttcaaaaata tggtattgat aatcctgata tgaataaatt gcagtttcat
4141 ttgatgctcg atgagttttt ctaatcagaa ttggttaatt ggttgtaaca ctggcactca
4201 accaagtcat tctgagaata gtgtatgcgg cgaccgagtt gctcttgccc ggcgtcaata
4261 cgggataata ccgcgccaca tagcagaact ttaaaagtgc tcatcattgg aaaacgttct
4321 tcggggcgaa aactctcaag gatcttaccg ctgttgagat ccagttcgat gtaacccact
4381 cgtgcaccca actgatcttc agcatctttt actttcacca gcgtttctgg gtgagcaaaa
4441 acaggaaggc aaaatgccgc aaaaaaggga ataagggcga cacggaaatg ttgaatactc
4501 atactcttcc tttttcaata ttattgaagc atttatcagg gttattgtct catgagcgga
4561 tacatatttg aatgtattta gaaaaataaa caaatagggg ttccgcgcac atttccccga
4621 aaagtgccac ctgacgtcta agaaaccatt attatcatga cattaaccta taaaaatagg
4681 cgtatcacga ggccctttcg tctcgcgcgt ttcggtgatg acggtgaaaa cctctgacac
4741 atgcagctcc cggagacggt cacagcttgt ctgtaagcgg atgccgggag cagacaagcc
4801 cgtcagggcg cgtcagcggg tgttggcggg tgtcggggct ggcttaacta tgcggcatca
4861 gagcagattg tactgagagt gcaccagatc gtcttgcctt gctcgtcggt gatgtactta
4921 cagctcgaag tgcctcttct tgatggagcg catggggacg tgcttggcaa tcacgcgcac
4981 cccccggccg ttttagcggc taaaaaagtc atggctctgc cctcgggcgg accacgccca
5041 tcatgacctt gccaagctcg tcctgcttct cttcgatctt cgccagcagg gcgaggatcg
5101 tggcatcacc gaaccgcgcc gtgcgcgggt cgtcggtgag ccagagtttc agcaggccgc
5161 ccaggcggcc caggtcgcca ttgatgcggg ccagctcgcg gacgtgctca tagtccacga
5221 cgcccgtgat tttgtagccc tggccgacgg ccagcaggta ggccgacagg ctcatgccgg
5281 ccgccgccgc cttttcctca atcgctcttc gttcgtctgg aaggcagtac accttgatag
5341 gtgggctgcc cttcctggtt ggcttggttt catcagccat ccgcttgccc tcatctgtta
5401 cgccggcggt agccggccag cctcgcagag caggattccc gttgagcacc gccaggtgcg
5461 aataagggac agtgaagaag gaacacccgc tcgcgggtgg gcctacttca cctatcctgc
5521 ccggctgacg ccgttggata caccaaggaa agtctacacg aaccctttgg caaaatcctg
5581 tatatcgtgc gaaaaaggat ggatataccg aaaaaatcgc tataatgacc ccgaagcagg
5641 gttatgcagc ggaagatggc cattcgccat tcaggctgcg caactgttgg gaagggcgat
5701 cggtgcgggc ctcttcgcta ttacgccagc tggcgaaagg gggatgtgct gcaaggcgat
5761 taagttgggt aacgccaggg ttttcccagt cacgacgttg taaaacgacg gccagtgacc
5821 atgattacgc caagctcgaa attaaccctc actaaaggga acaaaagctg gtacctaaca
5881 ggattagtgc aattcgagca catctcgttc gctattcagg gattgctcga gtcgagtcca
5941 ggcgccctag ccttctgatg tttgtgggtc ggtctgggcg ccctagcctt ctgaaagctg
6001 attggctgaa gaataatttg agtattggag ggcctggtgg caatgtcaca tcatgttcct
6061 gctgtcacat gttcacgaac ctatttttat gtgcttcatc atctcattgg ttggaattgc
6121 tgaaagtggt tattccacca agtcagagga aattacggtt aaccgttaaa ctatgtgcaa
6181 tcgttaagac tgaggtctga cttttctctg acttagctag tattaataat ctcttacttc
6241 tattaccgga gtattgttta atcttatcat acgtagcggt actcgtagca cacctacggt
6301 actcgttgct cgtaccgtag cccctagcac ctttggtagt cgatggaaca tggagtaagg
6361 cactaatgta cggtaggagc atacgtacgt accgtatctt accaaatttt gacgagaaga
6421 atgaactgtc tgacgagacc attctatcat tattttttcg ggtgaggttt taagagaaag
6481 agaaaagttt gcctcgccgc ttgctgtgcc agcgctcaga gaaacggttt tgatttttgg
6541 ttcaaaaaac gactaaagga cataacgtag aaaacctagc cgatttcagc gagggataac
6601 tttgcttctc gttcaaaaaa aagatcgttg gcattcatcg agactcccta cccgtgagtg
6661 ttatttcaat gaaaaaataa ccgcacgtaa cttgtcacaa ataagcaatc acatcggtgc
6721 catccacact ccgtagtttt acacatcatc ttgacactag taatcaaata aatttatcaa
6781 catggctgat atggtgagca agggcgagga gctgttcacc ggggtggtgc ccatcctggt
6841 cgagctggac ggcgacgtaa acggccacaa gttcagcgtg tccggcgagg gcgagggcga
6901 tgccacctac ggcaagctga ccctgaagtt catctgcacc accggcaagc tgcccgtgcc
6961 ctggcccacc ctcgtgacca ccctgaccta cggcgtgcag tgcttcagcc gctaccccga
7021 ccacatgaag cagcacgact tcttcaagtc cgccatgccc gaaggctacg tccaggagcg
7081 caccatcttc ttcaaggacg acggcaacta caagacccgc gccgaggtga agttcgaggg
7141 cgacaccctg gtgaaccgca tcgagctgaa gggcatcgac ttcaaggagg acggcaacat
7201 cctggggcac aagctggagt acaactacaa cagccacaac gtctatatca tggccgacaa
7261 gcagaagaac ggcatcaagg tgaacttcaa gatccgccac aacatcgagg acggcagcgt
7321 gcagctcgcc gaccactacc agcagaacac ccccatcggc gacggccccg tgctgctgcc
7381 cgacaaccac tacctgagca cccagtccgc cctgagcaaa gaccccaacg agaagcgcga
7441 tcacatggtc ctgctggagt tcgtgaccgc cgccgggatc actctcggca tggacgagct
7501 gtacaagtga taaatcaaag attggggtga tagatgatca tatccaatag tgcatgcgaa
7561 tctattcgat atccgatacg aagttgcgcc tatttggttc gttgttttgc agcggtacta
7621 gtgaccataa tctttcaaag gagtagtaga actaacaata attatgatca ctaagcgtgc
7681 ctgatggatt ccccacttat cctagcgttt gatgggtttg agaacggcca gatctgaaaa
7741 tgatttgtaa tattaccatg ccgtatccgg tacgcgtaaa caatgacgac tgaattcatt
7801 agcaagccgg cccatccctt atttattggc gagagtccgg aatcaaaaaa tcattccgaa
7861 tttttcgtac cgtacggtag atatcggtat cggtaattag gagtattatc cgaaagtact
7921 ttacagctag gtagaacaag tacggtacgt acagcactca gatcgaactc gaactttatt
7981 ctcaccgcag gacccaccag tgggttgaat gtaaagacgc taataaatac ccgacagaag
8041 gcgccttttt tcttaaagtt ccgtacatgg gatgtcccaa aggtcaaagg tactcgtacg
8101 agtaccaggt accgttacgc taccggtacg gtatggggta cggcacgagc attattttct
8161 aatgtgccat agacatagtc gaactcttga gaattcgttg acgcatcaga gttgtgttat
8221 tgct

pPtPUC3

For details, see Karas et al Nat Commun. 2015 Apr 21;6:6925. doi: 10.1038/ncomms7925 available from Addgene (Plasmid #62863)

LUC –only vector

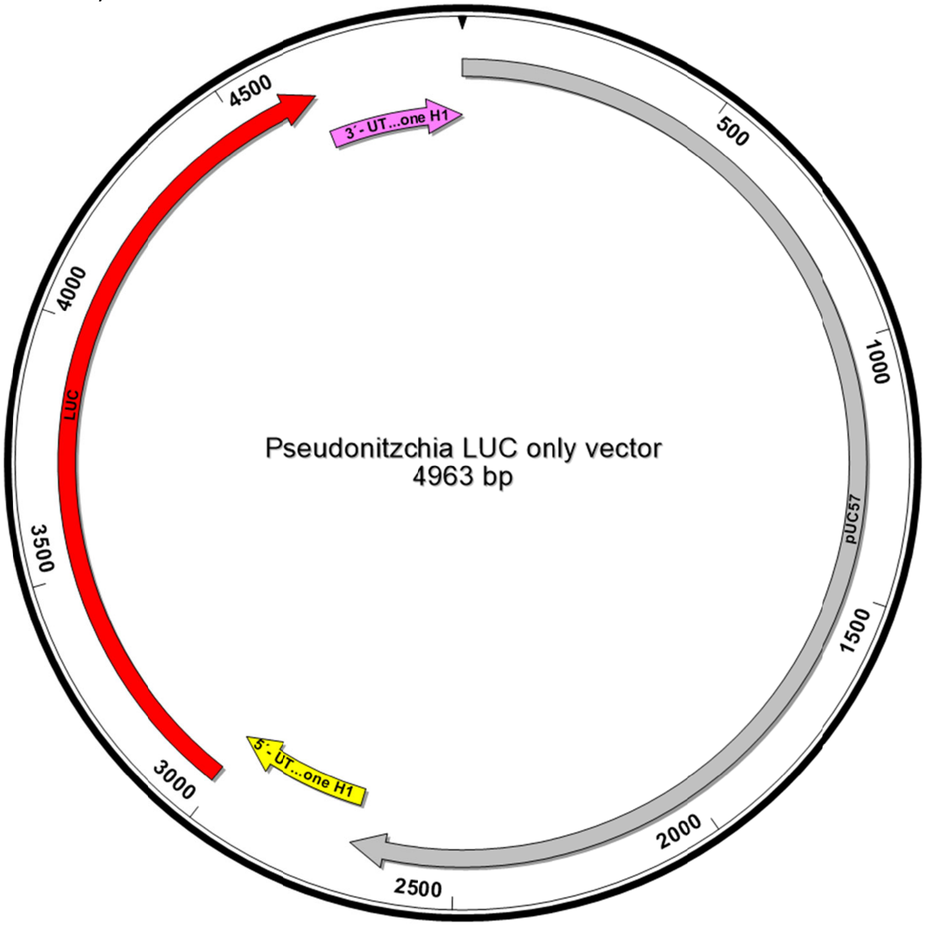


```
1-2270 pUC57 backbone
2771-3010 5´- UTR Histone H1
3011-4663 Luciferase
4664-4963 3´-UTR UTR Histone H1
```

```
ATCGGATCCCGGGCCCGTCGACTGCAGAGGCCTGCATGCAAGCTTGGCGTAATCATGGTCATAGCTGTTTCCTGTGTGAAATTGTT
ATCCGCTCACAATTCCACACAACATACGAGCCGGAAGCATAAAGTGTAAAGCCTGGGGTGCCTAATGAGTGAGCTAACTCACATTA
ATTGCGTTGCGCTCACTGCCCGCTTTCCAGTCGGGAAACCTGTCGTGCCAGCTGCATTAATGAATCGGCCAACGCGCGGGGAGAGG
CGGTTTGCGTATTGGGCGCTCTTCCGCTTCCTCGCTCACTGACTCGCTGCGCTCGGTCGTTCGGCTGCGGCGAGCGGTATCAGCTC
ACTCAAAGGCGGTAATACGGTTATCCACAGAATCAGGGGATAACGCAGGAAAGAACATGTGAGCAAAAGGCCAGCAAAAGGCCAGG
AACCGTAAAAAGGCCGCGTTGCTGGCGTTTTTCCATAGGCTCCGCCCCCCTGACGAGCATCACAAAAATCGACGCTCAAGTCAGAG
GTGGCGAAACCCGACAGGACTATAAAGATACCAGGCGTTTCCCCCTGGAAGCTCCCTCGTGCGCTCTCCTGTTCCGACCCTGCCGC
TTACCGGATACCTGTCCGCCTTTCTCCCTTCGGGAAGCGTGGCGCTTTCTCATAGCTCACGCTGTAGGTATCTCAGTTCGGTGTAG
GTCGTTCGCTCCAAGCTGGGCTGTGTGCACGAACCCCCCGTTCAGCCCGACCGCTGCGCCTTATCCGGTAACTATCGTCTTGAGTC
CAACCCGGTAAGACACGACTTATCGCCACTGGCAGCAGCCACTGGTAACAGGATTAGCAGAGCGAGGTATGTAGGCGGTGCTACAG
AGTTCTTGAAGTGGTGGCCTAACTACGGCTACACTAGAAGAACAGTATTTGGTATCTGCGCTCTGCTGAAGCCAGTTACCTTCGGA
AAAAGAGTTGGTAGCTCTTGATCCGGCAAACAAACCACCGCTGGTAGCGGTGGTTTTTTTGTTTGCAAGCAGCAGATTACGCGCAG
AAAAAAAGGATCTCAAGAAGATCCTTTGATCTTTTCTACGGGGTCTGACGCTCAGTGGAACGAAAACTCACGTTAAGGGATTTTGG
TCATGAGATTATCAAAAAGGATCTTCACCTAGATCCTTTTAAATTAAAAATGAAGTTTTAAATCAATCTAAAGTATATATGAGTAA
ACTTGGTCTGACAGTTACCAATGCTTAATCAGTGAGGCACCTATCTCAGCGATCTGTCTATTTCGTTCATCCATAGTTGCCTGACT
CCCCGTCGTGTAGATAACTACGATACGGGAGGGCTTACCATCTGGCCCCAGTGCTGCAATGATACCGCGAGACCCACGCTCACCGG
CTCCAGATTTATCAGCAATAAACCAGCCAGCCGGAAGGGCCGAGCGCAGAAGTGGTCCTGCAACTTTATCCGCCTCCATCCAGTCT
ATTAATTGTTGCCGGGAAGCTAGAGTAAGTAGTTCGCCAGTTAATAGTTTGCGCAACGTTGTTGCCATTGCTACAGGCATCGTGGT
GTCACGCTCGTCGTTTGGTATGGCTTCATTCAGCTCCGGTTCCCAACGATCAAGGCGAGTTACATGATCCCCCATGTTGTGCAAAA
AAGCGGTTAGCTCCTTCGGTCCTCCGATCGTTGTCAGAAGTAAGTTGGCCGCAGTGTTATCACTCATGGTTATGGCAGCACTGCAT
AATTCTCTTACTGTCATGCCATCCGTAAGATGCTTTTCTGTGACTGGTGAGTACTCAACCAAGTCATTCTGAGAATAGTGTATGCG
GCGACCGAGTTGCTCTTGCCCGGCGTCAATACGGGATAATACCGCGCCACATAGCAGAACTTTAAAAGTGCTCATCATTGGAAAAC
```

GTTCTTCGGGGCGAAAACTCTCAAGGATCTTACCGCTGTTGAGATCCAGTTCGATGTAACCCACTCGTGCACCCAACTGATCTTCA
GCATCTTTTACTTTCACCAGCGTTTCTGGGTGAGCAAAAACAGGAAGGCAAAATGCCGCAAAAAAGGGAATAAGGGCGACACGGAA
ATGTTGAATACTCATACTCTTCCTTTTTCAATATTATTGAAGCATTTATCAGGGTTATTGTCTCATGAGCGGATACATATTTGAAT
GTATTTAGAAAAATAAACAAATAGGGGTTCCGCGCACATTTCCCCGAAAAGTGCCACCTGACGTCTAAGAAACCATTATTATCATG
ACATTAACCTATAAAAATAGGCGTATCACGAGGCCCTTTCGTCTCGCGCGTTTCGGTGATGACGGTGAAAACCTCTGACACATGCA
GCTCCCGGAGACGGTCACAGCTTGTCTGTAAGCGGATGCCGGGAGCAGACAAGCCCGTCAGGGCGCGTCAGCGGGTGTTGGCGGGT
GTCGGGGCTGGCTTAACTATGCGGCATCAGAGCAGATTGTACTGAGAGTGCACCATATGCGGTGTGAAATACCGCACAGATGCGTA
AGGAGAAAATACCGCATCAGGCGCCATTCGCCATTCAGGCTGCGCAACTGTTGGGAAGGGCGATCGGTGCGGGCCTCTTCGCTATT
ACGCCAGCTGGCGAAAGGGGGATGTGCTGCAAGGCGATTAAGTTGGGTAACGCCAGGGTTTTCCCAGTCACGACGTTGTAAAACGA
CGGCCAGTGAATTCGAGCTCGGTACCTCGCGAATGCATCTAGATTACTCCGTGTTAGGCCGCGCCCGTTTTTACGGTAGGTACAGA
TAGCCGTGCAGGCTCGCGAGCGAACCACGTTTTTCCCCTTTTTTTCTACGGATCAATTCTACCGGATTACACAGCAAACGTTTTTG
ATTTTGGAGCCTTTTTAAGACCCAGAAAATCTCTGGCGCGAGACAGAGATTGTTCAGAGATGCAATCACTTTTTACAATTCTTCTG
CAAAAGACAGATTTTATCTCACTCGCTGCATCGCGCATAAAAAGTACCGAAAATAAAATATTTTTTCCATTCAAACAACATCAATC
ATGGAAGACGCCAAAAACATAAAGAAAGGCCCGGCGCCATTCTATCCGCTGGAAGATGGAACCGCTGGAGAGCAACTGCATAAGGC
TATGAAGAGATACGCCCTGGTTCCTGGAACAATTGCTTTTACAGATGCACATATCGAGGTGGACATCACTTACGCTGAGTACTTCG
AAATGTCCGTTCGGTTGGCAGAAGCTATGAAACGATATGGGCTGAATACAAATCACAGAATCGTCGTATGCAGTGAAAACTCTCTT
CAATTCTTTATGCCGGTGTTGGGCGCGTTATTTATCGGAGTTGCAGTTGCGCCCGCGAACGACATTTATAATGAACGTGAATTGCT
CAACAGTATGGGCATTTCGCAGCCTACCGTGGTGTTCGTTTCCAAAAAGGGGTTGCAAAAAATTTTGAACGTGCAAAAAAAGCTCC
CAATCATCCAAAAAATTATTATCATGGATTCTAAAACGGATTACCAGGGATTTCAGTCGATGTACACGTTCGTCACATCTCATCTA
CCTCCCGGTTTTAATGAATACGATTTTGTGCCAGAGTCCTTCGATAGGGACAAGACAATTGCACTGATCATGAACTCCTCTGGATC
TACTGGTCTGCCTAAAGGTGTCGCTCTGCCTCATAGAACTGCCTGCGTGAGATTCTCGCATGCCAGAGATCCTATTTTTGGCAATC
AAATCATTCCGGATACTGCGATTTTAAGTGTTGTTCCATTCCATCACGGTTTTGGAATGTTTACTACACTCGGATATTTGATATGT
GGATTTCGAGTCGTCTTAATGTATAGATTTGAAGAAGAGCTGTTTCTGAGGAGCCTTCAGGATTACAAGATTCAAAGTGCGCTGCT
GGTGCCAACCCTATTCTCCTTCTTCGCCAAAAGCACTCTGATTGACAAATACGATTTATCTAATTTACACGAAATTGCTTCTGGTG
GCGCTCCCCTCTCTAAGGAAGTCGGGGAAGCGGTTGCCAAGAGGTTCCATCTGCCAGGTATCAGGCAAGGATATGGGCTCACTGAG
ACTACATCAGCTATTCTGATTACACCCGAGGGGGATGATAAACCGGGCGCGGTCGGTAAAGTTGTTCCATTTTTTGAAGCGAAGGT
TGTGGATCTGGATACCGGGAAAACGCTGGGCGTTAATCAAAGAGGCGAACTGTGTGTGAGAGGTCCTATGATTATGTCCGGTTATG
TAAACAATCCGGAAGCGACCAACGCCTTGATTGACAAGGATGGATGGCTACATTCTGGAGACATAGCTTACTGGGACGAAGACGAA
CACTTCTTCATCGTTGACCGCCTGAAGTCTCTGATTAAGTACAAAGGCTATCAGGTGGCTCCCGCTGAATTGGAATCCATCTTGCT
CCAACACCCCAACATCTTCGACGCAGGTGTCGCAGGTCTTCCCGACGATGACGCCGGTGAACTTCCCGCCGCCGTTGTTGTTTTGG
AGCACGGAAAGACGATGACGGAAAAAGAGATCGTGGATTACGTCGCCAGTCAAGTAACAACCGCGAAAAAGTTGCGCGGAGGAGTT
GTGTTTGTGGACGAAGTACCGAAAGGTCTTACCGGAAAACTCGACGCAAGAAAAATCAGAGAGATCCTCATAAAGGCCAAGAAGGG
CGGAAAGATCGCCGTGTAAATTGCTCACAGCATTCTCAATGCTACTACATCCGGTGTTCTTAAACACCACCATTCTCTTCGGAAAC
TGTGCCTTAGTCAAAAATTGTAAACCGATAGAAAATAAATAGAAAATGATGTTATCTAGTGCATAAGTAGCGAGTTGGAATACGTT
CCTACGTCTCCTTGTCAACACCACGATGTAATTCAAGTGATTAGCCATGCTTGATGGCAGATGAACAAAGGGAATTCATATTGTAC
ATGATTAAGCGTAGTGGAGCCAGTGAAGAACTCCCACACCCGCCCATCCTAGCTAGTGAAC

*Heterosigma akashiwo*

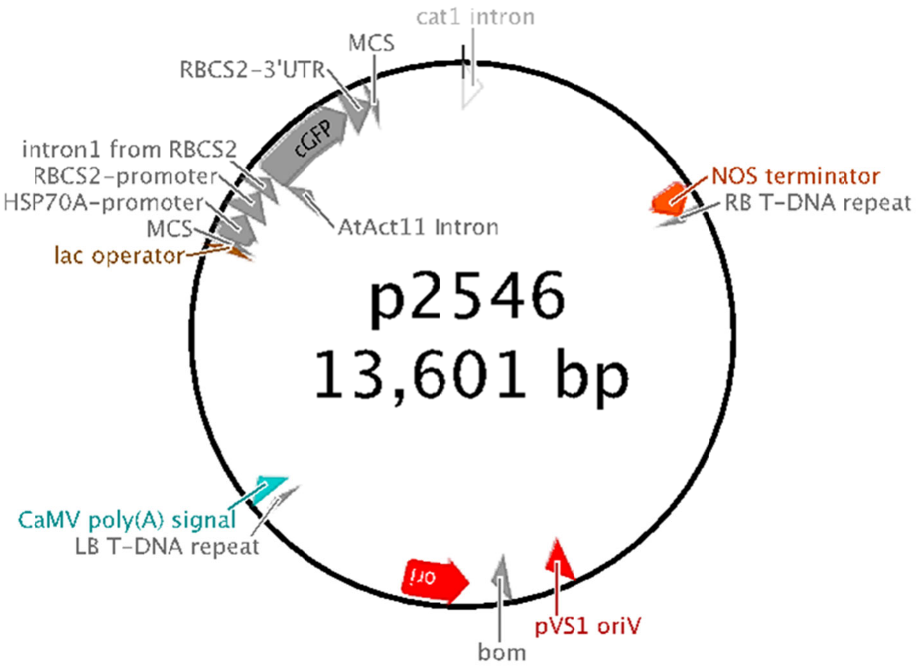


```
LOCUS       p2546                  13601 bp ds-DNA     circular     19-OCT-2016
DEFINITION  .
FEATURES             Location/Qualifiers
     misc_feature    12821..12837
                     /label="MCS"
                     /ApEinfo_revcolor=#84b0dc
                     /ApEinfo_fwdcolor=#84b0dc
     primer_bind     11004..11020
                     /label="M13 rev"
                     /ApEinfo_revcolor=#a020f0
                     /ApEinfo_fwdcolor=#a020f0
     protein_bind    10980..10996
                     /label="lac operator"
                     /ApEinfo_revcolor=#31849b
                     /ApEinfo_fwdcolor=#31849b
     promoter        10966..10972
                     /label="lac promoter"
                     /ApEinfo_revcolor=#c6c9d1
                     /ApEinfo_fwdcolor=#c6c9d1
     misc_feature    2350..2374
                     /label="RB T-DNA repeat"
                     /ApEinfo_revcolor=#f8d3a9
                     /ApEinfo_fwdcolor=#f8d3a9
     CDS             2024..2041
                     /label="6xHis"
                     /ApEinfo_revcolor=#ff9ccd
                     /ApEinfo_fwdcolor=#ff9ccd
     CDS             complement(7410..8204)
                     /label="KanR"
                     /ApEinfo_revcolor=#b7e6d7
                     /ApEinfo_fwdcolor=#b7e6d7
     misc_feature    12572..12815
                     /label="RBCS2-3'UTR"
                     /ApEinfo_revcolor=#f58a5e
                     /ApEinfo_fwdcolor=#f58a5e
     CDS             200..2017
                     /label="GUS"
                     /ApEinfo_revcolor=#993366
                     /ApEinfo_fwdcolor=#993366
```

```
misc_feature    8629..8653
                /label="LB T-DNA repeat"
                /ApEinfo_revcolor=#f8d3a9
                /ApEinfo_fwdcolor=#f8d3a9
rep_origin      complement(6735..7323)
                /label="ori"
                /ApEinfo_revcolor=#ffef86
                /ApEinfo_fwdcolor=#ffef86
CDS             10..199
                /label="GUS"
                /ApEinfo_revcolor=#a187be
                /ApEinfo_fwdcolor=#a187be
promoter        complement(10038..10715)
                /label="CaMV 35S promoter (enhanced)"
                /ApEinfo_revcolor=#c6c9d1
                /ApEinfo_fwdcolor=#c6c9d1
polyA_signal    8731..8905
                /label="CaMV poly(A) signal"
                /ApEinfo_revcolor=#b4abac
                /ApEinfo_fwdcolor=#b4abac
CDS             4732..5805
                /label="pVS1 RepA"
                /ApEinfo_revcolor=#993366
                /ApEinfo_fwdcolor=#993366
misc_feature    11699..12570
                /label="cGFP"
                /ApEinfo_revcolor=#b1ff67
                /ApEinfo_fwdcolor=#b1ff67
intron          10..199
                /label="cat1 intron"
                /ApEinfo_revcolor=#a187be
                /ApEinfo_fwdcolor=#a187be
promoter        10942..10947
                /label="lac promoter"
                /ApEinfo_revcolor=#c6c9d1
                /ApEinfo_fwdcolor=#c6c9d1
promoter        13234..13579
                /label="CaMV 35S promoter"
                /ApEinfo_revcolor=#c6c9d1
                /ApEinfo_fwdcolor=#c6c9d1
terminator      2076..2328
                /label="NOS terminator"
                /ApEinfo_revcolor=#c6c9d1
                /ApEinfo_fwdcolor=#c6c9d1
misc_feature    11040..11309
                /label="HSP70A-promoter"
                /ApEinfo_revcolor=#d59687
                /ApEinfo_fwdcolor=#d59687
CDS             12821..13000
                /label="lacZα"
                /ApEinfo_revcolor=#993366
                /ApEinfo_fwdcolor=#993366
misc_feature    11540..11693
                /label="intron1 from RBCS2"
                /ApEinfo_revcolor=#c6c9d1
                /ApEinfo_fwdcolor=#c6c9d1
misc_feature    6409..6549
                /label="bom"
                /ApEinfo_revcolor=#b4abac
                /ApEinfo_fwdcolor=#b4abac
misc_feature    11714..11868
                /label="AtAct11 Intron"
                /ApEinfo_revcolor=#9eafd2
                /ApEinfo_fwdcolor=#9eafd2
CDS             11016..11036
                /label="lacZα"
                /ApEinfo_revcolor=#993366
                /ApEinfo_fwdcolor=#993366
primer_bind     complement(12841..12857)
                /label="M13 fwd"
                /ApEinfo_revcolor=#a020f0
                /ApEinfo_fwdcolor=#a020f0
misc_feature    11310..11539
                /label="RBCS2-promoter"
                /ApEinfo_revcolor=#008080
                /ApEinfo_fwdcolor=#008080
CDS             3674..4303
```

```
                     /label="pVS1 StaA"
                     /ApEinfo_revcolor=#993366
                     /ApEinfo_fwdcolor=#993366
     rep_origin      5871..6065
                     /label="pVS1 oriV"
                     /ApEinfo_revcolor=#ffef86
                     /ApEinfo_fwdcolor=#ffef86
     CDS             13596..9
                     /label="GUS"
                     /ApEinfo_revcolor=#993366
                     /ApEinfo_fwdcolor=#993366
     CDS             complement(8945..9970)
                     /label="HygR"
                     /ApEinfo_revcolor=#b7e6d7
                     /ApEinfo_fwdcolor=#b7e6d7
     misc_feature    11030..11036
                     /label="MCS"
                     /ApEinfo_revcolor=#84b0dc
                     /ApEinfo_fwdcolor=#84b0dc
     promoter        10948..10965
                     /label="lac promoter"
                     /ApEinfo_revcolor=#c6c9d1
                     /ApEinfo_fwdcolor=#c6c9d1
ORIGIN
        1 GATCTGAGGG TAAATTTCTA GTTTTTCTCC TTCATTTTCT TGGTTAGGAC CCTTTTCTCT
       61 TTTTATTTTT TTGAGCTTTG ATCTTTCTTT AAACTGATCT ATTTTTTAAT TGATTGGTTA
      121 TGGTGTAAAT ATTACATAGC TTTAACTGAT AATCTGATTA CTTTATTTCG TGTGTCTATG
      181 ATGATGATGA TAGTTACAGA ACCGACGACT CGTCCGTCCT GTAGAAACCC CAACCCGTGA
      241 AATCAAAAAA CTCGACGGCC TGTGGGCATT CAGTCTGGAT CGCGAAAACT GTGGAATTGA
      301 TCAGCGTTGG TGGGAAAGCG CGTTACAAGA AAGCCGGGCA ATTGCTGTGC CAGGCAGTTT
      361 TAACGATCAG TTCGCCGATG CAGATATTCG TAATTATGCG GGCAACGTCT GGTATCAGCG
      421 CGAAGTCTTT ATACCGAAAG GTTGGGCAGG CCAGCGTATC GTGCTGCGTT TCGATGCGGT
      481 CACTCATTAC GGCAAAGTGT GGGTCAATAA TCAGGAAGTG ATGGAGCATC AGGGCGGCTA
      541 TACGCCATTT GAAGCCGATG TCACGCCGTA TGTTATTGCC GGGAAAAGTG TACGTATCAC
      601 CGTTTGTGTG AACAACGAAC TGAACTGGCA GACTATCCCG CCGGGAATGG TGATTACCGA
      661 CGAAAACGGC AAGAAAAAGC AGTCTTACTT CCATGATTTC TTTAACTATG CCGGAATCCA
      721 TCGCAGCGTA ATGCTCTACA CCACGCCGAA CACCTGGGTG GACGATATCA CCGTGGTGAC
      781 GCATGTCGCG CAAGACTGTA ACCACGCGTC TGTTGACTGG CAGGTGGTGG CCAATGGTGA
      841 TGTCAGCGTT GAACTGCGTG ATGCGGATCA ACAGGTGGTT GCAACTGGAC AAGGCACTAG
      901 CGGGACTTTG CAAGTGGTGA ATCCGCACCT CTGGCAACCG GGTGAAGGTT ATCTCTATGA
      961 ACTCGAAGTC ACAGCCAAAA GCCAGACAGA GTCTGATATC TACCCGCTTC GCGTCGGCAT
     1021 CCGGTCAGTG GCAGTGAAGG GCCAACAGTT CCTGATTAAC CACAAACCGT TCTACTTTAC
     1081 TGGCTTTGGT CGTCATGAAG ATGCGGACTT ACGTGGCAAA GGATTCGATA ACGTGCTGAT
     1141 GGTGCACGAC CACGCATTAA TGGACTGGAT TGGGGCCAAC TCCTACCGTA CCTCGCATTA
     1201 CCCTTACGCT GAAGAGATGC TCGACTGGGC AGATGAACAT GGCATCGTGG TGATTGATGA
     1261 AACTGCTGCT GTCGGCTTTC AGCTGTCTTT AGGCATTGGT TTCGAAGCGG GCAACAAGCC
     1321 GAAAGAACTG TACAGCGAAG AGGCAGTCAA CGGGGAAACT CAGCAAGCGC ACTTACAGGC
     1381 GATTAAAGAG CTGATAGCGC GTGACAAAAA CCACCCAAGC GTGGTGATGT GGAGTATTGC
     1441 CAACGAACCG GATACCCGTC CGCAAGGTGC ACGGGAATAT TTCGCGCCAC TGGCGGAAGC
     1501 AACGCGTAAA CTCGACCCGA CGCGTCCGAT CACCTGCGTC AATGTAATGT TCTGCGACGC
     1561 TCACACCGAT ACCATCAGCG ATCTCTTTGA TGTGCTGTGC CTGAACCGTT ATTACGGATG
     1621 GTATGTCCAA AGCGGCGATT TGGAAACGGC AGAGAAGGTA CTGGAAAAAG AACTTCTGGC
     1681 CTGGCAGGAG AAACTGCATC AGCCGATTAT CATCACCGAA TACGGCGTGG ATACGTTAGC
     1741 CGGGCTGCAC TCAATGTACA CCGACATGTG GAGTGAAGAG TATCAGTGTG CATGGCTGGA
     1801 TATGTATCAC CGCGTCTTTG ATCGCGTCAG CGCCGTCGTC GGTGAACAGG TATGGAATTT
     1861 CGCCGATTTT GCGACCTCGC AAGGCATATT GCGCGTTGGC GGTAACAAGA AAGGGATCTT
     1921 CACTCGCGAC CGCAAACCGA AGTCGGCGGC TTTTCTGCTG CAAAAACGCT GGACTGGCAT
     1981 GAACTTCGGT GAAAAACCGC AGCAGGGAGG CAAACAAGCT AGCCACCACC ACCACCACCA
     2041 CGTGTGAATT ACAGGTGACC AGCTCGAATT TCCCCGATCG TTCAAACATT TGGCAATAAA
     2101 GTTTCTTAAG ATTGAATCCT GTTGCCGGTC TTGCGATGAT TATCATATAA TTTCTGTTGA
     2161 ATTACGTTAA GCATGTAATA ATTAACATGT AATGCATGAC GTTATTTATG AGATGGGTTT
     2221 TTATGATTAG AGTCCCGCAA TTATACATTT AATACGCGAT AGAAAACAAA ATATAGCGCG
     2281 CAAACTAGGA TAAATTATCG CGCGCGGTGT CATCTATGTT ACTAGATCGG GAATTAAACT
     2341 ATCAGTGTTT GACAGGATAT ATTGGCGGGT AAACCTAAGA GAAAAGAGCG TTTATTAGAA
     2401 TAACGGATAT TTAAAAGGGC GTGAAAAGGT TTATCCGTTC GTCCATTTGT ATGTGCATGC
     2461 CAACCACAGG GTTCCCCTCG GGATCAAAGT ACTTTGATCC AACCCCTCCG CTGCTATAGT
     2521 GCAGTCGGCT TCTGACGTTC AGTGCAGCCG TCTTCTGAAA ACGACATGTC GCACAAGTCC
     2581 TAAGTTACGC GACAGGCTGC CGCCCTGCCC TTTTCCTGGC GTTTTCTTGT CGCGTGTTTT
     2641 AGTCGCATAA AGTAGAATAC TTGCGACTAG AACCGGAGAC ATTACGCCAT GAACAAGAGC
     2701 GCCGCCGCTG GCCTGCTGGG CTATGCCCGC GTCAGCACCG ACGACCAGGA CTTGACCAAC
     2761 CAACGGGCCG AACTGCACGC GGCCGGCTGC ACCAAGCTGT TTTCCGAGAA GATCACCGGC
     2821 ACCAGGCGCG ACCGCCCGGA GCTGGCCAGG ATGCTTGACC ACCTACGCCC TGGCGACGTT
     2881 GTGACAGTGA CCAGGCTAGA CCGCCTGGCC CGCAGCACCC GCGACCTACT GGACATTGCC
     2941 GAGCGCATCC AGGAGGCCGG CGCGGGCCTG CGTAGCCTGG CAGAGCCGTG GGCCGACACC
     3001 ACCACGCCGG CCGGCCGCAT GGTGTTGACC GTGTTCGCCG GCATTGCCGA GTTCGAGCGT
     3061 TCCCTAATCA TCGACCGCAC CCGGAGCGGG CGCGAGGCCG CCAAGGCCCG AGGCGTGAAG
     3121 TTTGGCCCCC GCCCTACCCT CACCCCGGCA CAGATCGCGC ACGCCCGCGA GCTGATCGAC
```

3181 CAGGAAGGCC GCACCGTGAA AGAGGCGGCT GCACTGCTTG GCGTGCATCG CTCGACCCTG
3241 TACCGCGCAC TTGAGCGCAG CGAGGAAGTG ACGCCCACCG AGGCCAGGCG GCGCGGTGCC
3301 TTCCGTGAGG ACGCATTGAC CGAGGCCGAC GCCCTGGCGG CCGCCGAGAA TGAACGCCAA
3361 GAGGAACAAG CATGAAACCG CACCAGGACG GCCAGGACGA ACCGTTTTTC ATTACCGAAG
3421 AGATCGAGGC GGAGATGATC GCGGCCGGGT ACGTGTTCGA GCCGCCCGCG CACGTCTCAA
3481 CCGTGCGGCT GCATGAAATC CTGGCCGGTT TGTCTGATGC CAAGCTGGCG GCCTGGCCGG
3541 CCAGCTTGGC CGCTGAAGAA ACCGAGCGCC GCCGTCTAAA AAGGTGATGT GTATTTGAGT
3601 AAAACAGCTT GCGTCATGCG GTCGCTGCGT ATATGATGCG ATGAGTAAAT AAACAAATAC
3661 GCAAGGGGAA CGCATGAAGG TTATCGCTGT ACTTAACCAG AAAGGCGGGT CAGGCAAGAC
3721 GACCATCGCA ACCCATCTAG CCCGCGCCCT GCAACTCGCC GGGGCCGATG TTCTGTTAGT
3781 CGATTCCGAT CCCCAGGGCA GTGCCCGCGA TTGGGCGGCC GTGCGGGAAG ATCAACCGCT
3841 AACCGTTGTC GGCATCGACC GCCCGACGAT TGACCGCGAC GTGAAGGCCA TCGGCCGGCG
3901 CGACTTCGTA GTGATCGACG GAGCGCCCCA GGCGGCGGAC TTGGCTGTGT CCGCGATCAA
3961 GGCAGCCGAC TTCGTGCTGA TTCCGGTGCA GCCAAGCCCT TACGACATAT GGGCCACCGC
4021 CGACCTGGTG GAGCTGGTTA AGCAGCGCAT TGAGGTCACG GATGGAAGGC TACAAGCGGC
4081 CTTTGTCGTG TCGCGGGCGA TCAAAGGCAC GCGCATCGGC GGTGAGGTTG CCGAGGCGCT
4141 GGCCGGGTAC GAGCTGCCCA TTCTTGAGTC CCGTATCACG CAGCGCGTGA GCTACCCAGG
4201 CACTGCCGCC GCCGGCACAA CCGTTCTTGA ATCAGAACCC GAGGGCGACG CTGCCCGCGA
4261 GGTCCAGGCG CTGGCCGCTG AAATTAAATC AAAACTCATT TGAGTTAATG AGGTAAAGAG
4321 AAAATGAGCA AAAGCACAAA CACGCTAAGT GCCGGCCGTC CGAGCGCACG CAGCAGCAAG
4381 GCTGCAACGT TGGCCAGCCT GGCAGACACG CCAGCCATGA AGCGGGTCAA CTTTCAGTTG
4441 CCGGCGGAGG ATCACACCAA GCTGAAGATG TACGCGGTAC GCCAAGGCAA GACCATTACC
4501 GAGCTGCTAT CTGAATACAT CGCGCAGCTA CCAGAGTAAA TGAGCAAATG AATAAATGAG
4561 TAGATGAATT TTAGCGGCTA AAGGAGGCGG CATGGAAAAT CAAGAACAAC CAGGCACCGA
4621 CGCCGTGGAA TGCCCCATGT GTGGAGGAAC GGGCGGTTGG CCAGGCGTAA GCGGCTGGGT
4681 TGTCTGCCGG CCCTGCAATG GCACTGGAAC CCCCAAGCCC GAGGAATCGG CGTGAGCGGT
4741 CGCAAACCAT CCGGCCCGGT ACAAATCGGC GCGGCGCTGG GTGATGACCT GGTGGAGAAG
4801 TTGAAGGCCG CGCAGGCCGC CCAGCGGCAA CGCATCGAGG CAGAAGCACG CCCCGGTGAA
4861 TCGTGGCAAG CGGCCGCTGA TCGAATCCGC AAAGAATCCC GGCAACCGCC GGCAGCCGGT
4921 GCGCCGTCGA TTAGGAAGCC GCCCAAGGGC GACGAGCAAC CAGATTTTTT CGTTCCGATG
4981 CTCTATGACG TGGGCACCCG CGATAGTCGC AGCATCATGG ACGTGGCCGT TTTCCGTCTG
5041 TCGAAGCGTG ACCGACGAGC TGGCGAGGTG ATCCGCTACG AGCTTCCAGA CGGGCACGTA
5101 GAGGTTTCCG CAGGGCCGGC CGGCATGGCC AGTGTGTGGG ATTACGACCT GGTACTGATG
5161 GCGGTTTCCC ATCTAACCGA ATCCATGAAC CGATACCGGG AAGGGAAGGG AGACAAGCCC
5221 GGCCGCGTGT TCCGTCCACA CGTTGCGGAC GTACTCAAGT TCTGCCGGCG AGCCGATGGC
5281 GGAAAGCAGA AAGACGACCT GGTAGAAACC TGCATTCGGT TAAACACCAC GCACGTTGCC
5341 ATGCAGCGTA CGAAGAAGGC CAAGAACGGC CGCCTGGTGA CGGTATCCGA GGGTGAAGCC
5401 TTGATTAGCC GCTACAAGAT CGTAAAGAGC GAAACCGGGC GGCCGGAGTA CATCGAGATC
5461 GAGCTAGCTG ATTGGATGTA CCGCGAGATC ACAGAAGGCA AGAACCCGGA CGTGCTGACG
5521 GTTCACCCCG ATTACTTTTT GATCGATCCC GGCATCGGCC GTTTTCTCTA CCGCCTGGCA
5581 CGCCGCGCCG CAGGCAAGGC AGAAGCCAGA TGGTTGTTCA AGACGATCTA CGAACGCAGT
5641 GGCAGCGCCG GAGAGTTCAA GAAGTTCTGT TTCACCGTGC GCAAGCTGAT CGGGTCAAAT
5701 GACCTGCCGG AGTACGATTT GAAGGAGGAG GCGGGGCAGG CTGGCCCGAT CCTAGTCATG
5761 CGCTACCGCA ACCTGATCGA GGGCGAAGCA TCCGCCGGTT CCTAATGTAC GGAGCAGATG
5821 CTAGGGCAAA TTGCCCTAGC AGGGGAAAAA GGTCGAAAAG GTCTCTTTCC TGTGGATAGC
5881 ACGTACATTG GGAACCCAAA GCCGTACATT GGGAACCGGA ACCCGTACAT TGGGAACCCA
5941 AAGCCGTACA TTGGGAACCG GTCACACATG TAAGTGACTG ATATAAAAGA GAAAAAAGGC
6001 GATTTTTCCG CCTAAAACTC TTTAAAACTT ATTAAAACTC TTAAAACCCG CCTGGCCTGT
6061 GCATAACTGT CTGGCCAGCG CACAGCCGAA GAGCTGCAAA AAGCGCCTAC CCTTCGGTCG
6121 CTGCGCTCCC TACGCCCCGC CGCTTCGCGT CGGCCTATCG CGGCCGCTGG CCGCTCAAAA
6181 ATGGCTGGCC TACGGCCAGG CAATCTACCA GGGCGCGGAC AAGCCGCGCC GTCGCCACTC
6241 GACCGCCGGC GCCCACATCA AGGCACCCTG CCTCGCGCGT TTCGGTGATG ACGGTGAAAA
6301 CCTCTGACAC ATGCAGCTCC CGGAGACGGT CACAGCTTGT CTGTAAGCGG ATGCCGGGAG
6361 CAGACAAGCC CGTCAGGGCG CGTCAGCGGG TGTTGGCGGG TGTCGGGGCG CAGCCATGAC
6421 CCAGTCACGT AGCGATAGCG GAGTGTATAC TGGCTTAACT ATGCGGCATC AGAGCAGATT
6481 GTACTGAGAG TGCACCATAT GCGGTGTGAA ATACCGCACA GATGCGTAAG GAGAAAATAC
6541 CGCATCAGGC GCTCTTCCGC TTCCTCGCTC ACTGACTCGC TGCGCTCGGT CGTTCGGCTG
6601 CGGCGAGCGG TATCAGCTCA CTCAAAGGCG GTAATACGGT TATCCACAGA ATCAGGGGAT
6661 AACGCAGGAA AGAACATGTG AGCAAAAGGC CAGCAAAAGG CCAGGAACCG TAAAAAGGCC
6721 GCGTTGCTGG CGTTTTTCCA TAGGCTCCGC CCCCCTGACG AGCATCACAA AAATCGACGC
6781 TCAAGTCAGA GGTGGCGAAA CCCGACAGGA CTATAAAGAT ACCAGGCGTT TCCCCCTGGA
6841 AGCTCCCTCG TGCGCTCTCC TGTTCCGACC CTGCCGCTTA CCGGATACCT GTCCGCCTTT
6901 CTCCCTTCGG GAAGCGTGGC GCTTTCTCAT AGCTCACGCT GTAGGTATCT CAGTTCGGTG
6961 TAGGTCGTTC GCTCCAAGCT GGGCTGTGTG CACGAACCCC CCGTTCAGCC CGACCGCTGC
7021 GCCTTATCCG GTAACTATCG TCTTGAGTCC AACCCGGTAA GACACGACTT ATCGCCACTG
7081 GCAGCAGCCA CTGGTAACAG GATTAGCAGA GCGAGGTATG TAGGCGGTGC TACAGAGTTC
7141 TTGAAGTGGT GGCCTAACTA CGGCTACACT AGAAGGACAG TATTTGGTAT CTGCGCTCTG
7201 CTGAAGCCAG TTACCTTCGG AAAAAGAGTT GGTAGCTCTT GATCCGGCAA ACAAACCACC
7261 GCTGGTAGCG GTGGTTTTTT TGTTTGCAAG CAGCAGATTA CGCGCAGAAA AAAAGGATCT
7321 CAAGAAGATC CTTTGATCTT TTCTACGGGG TCTGACGCTC AGTGGAACGA AAACTCACGT
7381 TAAGGGATTT TGGTCATGCA TTCTAGGTAC TAAAACAATT CATCCAGTAA AATATAATAT
7441 TTTATTTTCT CCCAATCAGG CTTGATCCCC AGTAAGTCAA AAAATAGCTC GACATACTGT
7501 TCTTCCCCGA TATCCTCCCT GATCGACCGG ACGCAGAAGG CAATGTCATA CCACTTGTCC
7561 GCCCTGCCGC TTCTCCCAAG ATCAATAAAG CCACTTACTT TGCCATCTTT CACAAAGATG
7621 TTGCTGTCTC CCAGGTCGCC GTGGGAAAAG ACAAGTTCCT CTTCGGGCTT TTCCGTCTTT
7681 AAAAAATCAT ACAGCTCGCG CGGATCTTTA AATGGAGTGT CTTCTTCCCA GTTTTCGCAA
7741 TCCACATCGG CCAGATCGTT ATTCAGTAAG TAATCCAATT CGGCTAAGCG GCTGTCTAAG

7801 CTATTCGTAT AGGGACAATC CGATATGTCG ATGGAGTGAA AGAGCCTGAT GCACTCCGCA
7861 TACAGCTCGA TAATCTTTTC AGGGCTTTGT TCATCTTCAT ACTCTTCCGA GCAAAGGACG
7921 CCATCGGCCT CACTCATGAG CAGATTGCTC CAGCCATCAT GCCGTTCAAA GTGCAGGACC
7981 TTTGGAACAG GCAGCTTTCC TTCCAGCCAT AGCATCATGT CCTTTTCCCG TTCCACATCA
8041 TAGGTGGTCC CTTTATACCG GCTGTCCGTC ATTTTTAAAT ATAGGTTTTC ATTTTCTCCC
8101 ACCAGCTTAT ATACCTTAGC AGGAGACATT CCTTCCGTAT CTTTTACGCA GCGGTATTTT
8161 TCGATCAGTT TTTTCAATTC CGGTGATATT CTCATTTTAG CCATTTATTA TTTCCTTCCT
8221 CTTTTCTACA GTATTTAAAG ATACCCCAAG AAGCTAATTA TAACAAGACG AACTCCAATT
8281 CACTGTTCCT TGCATTCTAA AACCTTAAAT ACCAGAAAAC AGCTTTTTCA AAGTTGTTTT
8341 CAAAGTTGGC GTATAACATA GTATCGACGG AGCCGATTTT GAAACCGCGG TGATCACAGG
8401 CAGCAACGCT CTGTCATCGT TACAATCAAC ATGCTACCCT CCGCGAGATC ATCCGTGTTT
8461 CAAACCCGGC AGCTTAGTTG CCGTTCTTCC GAATAGCATC GGTAACATGA GCAAAGTCTG
8521 CCGCCTTACA ACGGCTCTCC CGCTGACGCC GTCCCGGACT GATGGGCTGC CTGTATCGAG
8581 TGGTGATTTT GTGCCGAGCT GCCGGTCGGG GAGCTGTTGG CTGGCTGGTG GCAGGATATA
8641 TTGTGGTGTA AACAAATTGA CGCTTAGACA ACTTAATAAC ACATTGCGGA CGTTTTTAAT
8701 GTACTGAATT AACGCCGAAT TAATTCGGGG GATCTGGATT TTAGTACTGG ATTTTGGTTT
8761 TAGGAATTAG AAATTTTATT GATAGAAGTA TTTTACAAAT ACAAATACAT ACTAAGGGTT
8821 TCTTATATGC TCAACACATG AGCGAAACCC TATAGGAACC CTAATTCCCT TATCTGGGAA
8881 CTACTCACAC ATTATTATGG AGAAACTCGA GCTTGTCGAT CGACAGATCC GGTCGGCATC
8941 TACTCTATTT CTTTGCCCTC GGACGAGTGC TGGGGCGTCG GTTTCCACTA TCGGCGAGTA
9001 CTTCTACACA GCCATCGGTC CAGACGGCCG CGCTTCTGCG GGCGATTTGT GTACGCCCGA
9061 CAGTCCCGGC TCCGGATCGG ACGATTGCGT CGCATCGACC CTGCGCCCAA GCTGCATCAT
9121 CGAAATTGCC GTCAACCAAG CTCTGATAGA GTTGGTCAAG ACCAATGCGG AGCATATACG
9181 CCCGGAGTCG TGGCGATCCT GCAAGCTCCG GATGCCTCCG CTCGAAGTAG CGCGTCTGCT
9241 GCTCCATACA AGCCAACCAC GGCCTCCAGA AGAAGATGTT GGCGACCTCG TATTGGGAAT
9301 CCCCGAACAT CGCCTCGCTC CAGTCAATGA CCGCTGTTAT GCGGCCATTG TCCGTCAGGA
9361 CATTGTTGGA GCCGAAATCC GCGTGCACGA GGTGCCGGAC TTCGGGGCAG TCCTCGGCCC
9421 AAAGCATCAG CTCATCGAGA GCCTGCGCGA CGGACGCACT GACGGTGTCG TCCATCACAG
9481 TTTGCCAGTG ATACACATGG GGATCAGCAA TCGCGCATAT GAAATCACGC CATGTAGTGT
9541 ATTGACCGAT TCCTTGCGGT CCGAATGGGC CGAACCCGCT CGTCTGGCTA AGATCGGCCG
9601 CAGCGATCGC ATCCATAGCC TCCGCGACCG GTTGTAGAAC AGCGGGCAGT TCGGTTTCAG
9661 GCAGGTCTTG CAACGTGACA CCCTGTGCAC GGCGGGAGAT GCAATAGGTC AGGCTCTCGC
9721 TAAACTCCCC AATGTCAAGC ACTTCCGGAA TCGGGAGCGC GGCCGATGCA AAGTGCCGAT
9781 AAACATAACG ATCTTTGTAG AAACCATCGG CGCAGCTATT TACCCGCAGG ACATATCCAC
9841 GCCCTCCTAC ATCGAAGCTG AAAGCACGAG ATTCTTCGCC CTCCGAGAGC TGCATCAGGT
9901 CGGAGACGCT GTCGAACTTT TCGATCAGAA ACTTCTCGAC AGACGTCGCG GTGAGTTCAG
9961 GCTTTTTCAT ATCTCATTGC CCCCCGGGAT CTGCGAAAGC TCGAGAGAGA TAGATTTGTA
10021 GAGAGAGACT GGTGATTTCA GCGTGTCCTC TCCAAATGAA ATGAACTTCC TTATATAGAG
10081 GAAGGTCTTG CGAAGGATAG TGGGATTGTG CGTCATCCCT TACGTCAGTG GAGATATCAC
10141 ATCAATCCAC TTGCTTTGAA GACGTGGTTG GAACGTCTTC TTTTTCCACG ATGCTCCTCG
10201 TGGGTGGGGG TCCATCTTTG GGACCACTGT CGGCAGAGGC ATCTTGAACG ATAGCCTTTC
10261 CTTTATCGCA ATGATGGCAT TTGTAGGTGC CACCTTCCTT TTCTACTGTC CTTTTGATGA
10321 AGTGACAGAT AGCTGGGCAA TGGAATCCGA GGAGGTTTCC CGATATTACC CTTTGTTGAA
10381 AAGTCTCAAT AGCCCTTTGG TCTTCTGAGA CTGTATCTTT GATATTCTTG GAGTAGACGA
10441 GAGTGTCGTG CTCCACCATG TTATCACATC AATCCACTTG CTTTGAAGAC GTGGTTGGAA
10501 CGTCTTCTTT TTCCACGATG CTCCTCGTGG GTGGGGGTCC ATCTTTGGGA CCACTGTCGG
10561 CAGAGGCATC TTGAACGATA GCCTTTCCTT TATCGCAATG ATGGCATTTG TAGGTGCCAC
10621 CTTCCTTTTC TACTGTCCTT TTGATGAAGT GACAGATAGC TGGGCAATGG AATCCGAGGA
10681 GGTTTCCCGA TATTACCCTT TGTTGAAAAG TCTCAATAGC CCTTTGGTCT TCTGAGACTG
10741 TATCTTTGAT ATTCTTGGAG TAGACGAGAG TGTCGTGCTC CACCATGTTG GCAAGCTGCT
10801 CTAGCCAATA CGCAAACCGC CTCTCCCCGC GCGTTGGCCG ATTCATTAAT GCAGCTGGCA
10861 CGACAGGTTT CCCGACTGGA AAGCGGGCAG TGAGCGCAAC GCAATTAATG TGAGTTAGCT
10921 CACTCATTAG GCACCCCAGG CTTTACACTT TATGCTTCCG GCTCGTATGT TGTGTGGAAT
10981 TGTGAGCGGA TAACAATTTC ACACAGGAAA CAGCTATGAC CATGATTACG AATTCGAGCT
11041 CGCTGAGGCT TGACATGATT GGTGCGTATG TTTGTATGAA GCTACAGGAC TGATTTGGCG
11101 GGCTATGAGG GCGGGGGAAG CTCTGGAAGG GCCGCGATGG GGCGCGCGGC GTCCAGAAGG
11161 CGCCATACGG CCCGCTGGCG GCACCCATCC GGTATAAAAG CCCGCGACCC CGAACGGTGA
11221 CCTCCACTTT CAGCGACAAA CGAGCACTTA TACATACGCG ACTATTCTGC CGCTATACAT
11281 AACCACTCAG CTAGCTTAAG ATCCCATCAA GCTTGCATGC CGGGCGCGCC AGAAGGAGCG
11341 CAGCCAAACC AGGATGATGT TTGATGGGGT ATTTGAGCAC TTGCAACCCT TATCCGGAAG
11401 CCCCCTGGCC CACAAAGGCT AGGCGCCAAT GCAAGCAGTT CGCATGCAGC CCCTGGAGCG
11461 GTGCCCTCCT GATAAACCGG CCAGGGGGCC TATGTTCTTT ACTTTTTTAC AAGAGAAGTC
11521 ACTCAACATC TTAAAATGGC CAGGTGAGTC GACGAGCAAG CCCGGCGGAT CAGGCAGCGT
11581 GCTTGCAGAT TTGACTTGCA ACGCCCGCAT TGTGTCGACG AAGGCTTTTG GCTCCTCTGT
11641 CGCTGTCTCA AGCAGCATCT AACCCTGCGT CGCCGTTTCC ATTTGCAGGA TGCtcgagAT
11701 GGCCAAGGGC GAGgtaataa gatcttcaac acctacacca tttttttaat cactactacc
11761 cattgcattg aacaaacttc caagttcttc ttagcttcag attaagaaag taccctttct
11821 tggctttgtt gatgtggtac cattgtccat tgtcttgtgt gtttccagGA GCTGTTCACC
11881 GGTGTGGTCC CCATCCTGGT GGAGCTGGAC GGCGACGTGA ACGGCCACAA GTTCTCCGTC
11941 TCCGGCGAGG GTGAGGGTGA CGCCACCTAC GGCAAGCTGA CCCTGAAGTT CATCTGCACC
12001 ACCGGCAAGC TGCCCGTGCC CTGGCCCACC CTGGTCACCA CcctgaccTA CGGTGTGCAG
12061 TGCTTCTCCC GCTACCCCGA CCACATGAAG CAGCACGACT TCTTCAAGTC CGCCATGCCC
12121 GAGGGCTACG TGCAGGAGCG CACCATCTTC TTCAAGGACG ACGGCAACTA CAAGACCCGC
12181 GCCGAGGTCA AGTTCGAGGG CGACACCCTG GTGAACCGCA TCGAGCTGAA GGGCATCGAC
12241 TTCAAGGAGG ACGGCAACAT CCTGGGCCAC AAGCTGGAGT ACAACTACAA CTCCCACAAC
12301 GTGTACATCA TGGCCGACAA GCAGAAGAAC GGCATCAAGG TGAACTTCAA GATCCGCCAC
12361 AACATCGAGG ACGGCTCCGT GCAGCTGGCC GACCACTACC AGCAGAACAC CCCCATCGGC

```
    12421 GATGGCCCCG TGCTGCTGCC CGACAACCAC TACCTGTCCA TCCAGTCCGC CCTGTCCAAG
    12481 GACCCCAACG AGAAGCGCGA CCACATGGTC CTGCTGGAGT TCGTCACCGC TGCCGGCATC
    12541 ACCCACGGCA TGGACGAGCT GTACAAGTAA gGATCCCCGC TCCGTGTAAA TGGAGGCGCT
    12601 CGTTGATCTG AGCCTTGCCC CCTGACGAAC GGCGGTGGAT GGAAGATACT GCTCTCAAGT
    12661 GCTGAAGCGG TAGCTTAGCT CCCCGTTTCG TGCTGATCAG TCTTTTTCAA CACGTAAAAA
    12721 GCGGAGGAGT TTTGCAATTT TGTTGGTTGT AACGATCCTC CGTTGATTTT GGCCTCTTTC
    12781 TCCATGGGCG GGCTGGGCGT ATTTGAAGCG GGTACCCCcc TGCAGGCATG CAAGCTTGGC
    12841 ACTGGCCGTC GTTTTACAAC GTCGTGACTG GGAAAACCCT GGCGTTACCC AACTTAATCG
    12901 CCTTGCAGCA CATCCCCCTT TCGCCAGCTG GCGTAATAGC GAAGAGGCCC GCACCGATCG
    12961 CCCTTCCCAA CAGTTGCGCA GCCTGAATGG CGAATGCTAG AGCAGCTTGA GCTTGGATCA
    13021 GATTGTCGTT TCCCGCCTTC AGTTTAGCTT CATGGAGTCA AAGATTCAAA TAGAGGACCT
    13081 AACAGAACTC GCCGTAAAGA CTGGCGAACA GTTCATACAG AGTCTCTTAC GACTCAATGA
    13141 CAAGAAGAAA ATCTTCGTCA ACATGGTGGA GCACGACACA CTTGTCTACT CCAAAAATAT
    13201 CAAAGATACA GTCTCAGAAG ACCAAAGGGC AATTGAGACT TTTCAACAAA GGGTAATATC
    13261 CGGAAACCTC CTCGGATTCC ATTGCCCAGC TATCTGTCAC TTTATTGTGA AGATAGTGGA
    13321 AAAGGAAGGT GGCTCCTACA AATGCCATCA TTGCGATAAA GGAAAGGCCA TCGTTGAAGA
    13381 TGCCTCTGCC GACAGTGGTC CCAAAGATGG ACCCCCACCC ACGAGGAGCA TCGTGGAAAA
    13441 AGAAGACGTT CCAACCACGT CTTCAAAGCA AGTGGATTGA TGTGATATCT CCACTGACGT
    13501 AAGGGATGAC GCACAATCCC ACTATCCTTC GCAAGACCCT TCCTCTATAT AAGGAAGTTC
    13561 ATTTCATTTG GAGAGAACAC GGGGGACTCT TGACCATGGT A
//
```

## p2552

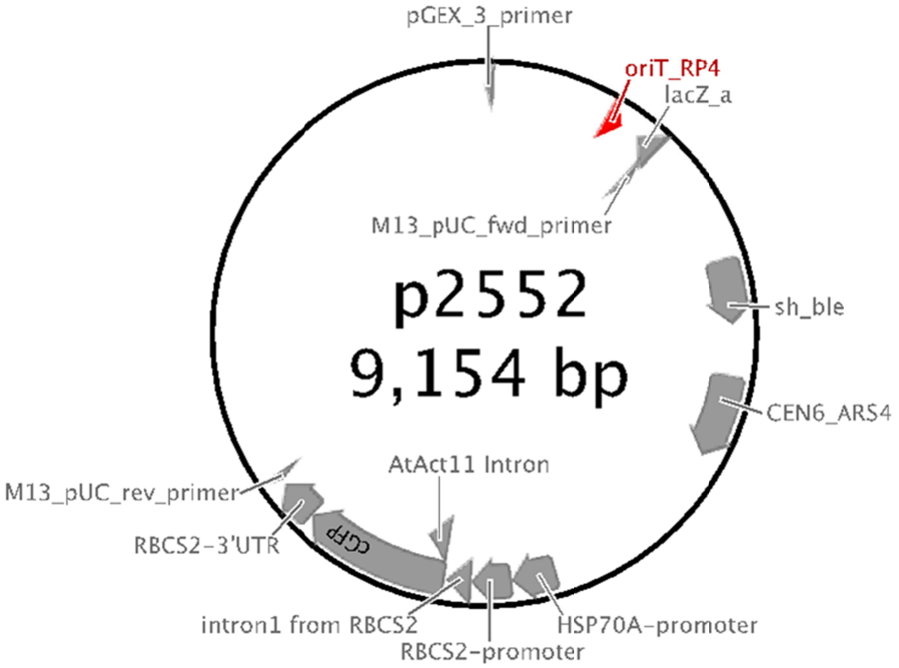


```
LOCUS       p2552                   9154 bp ds-DNA     circular     26-OCT-2019
 DEFINITION  .
 ACCESSION   urn.local...s-b4f8v7w
 KEYWORDS    .
 SOURCE      null
   ORGANISM
             .
 FEATURES             Location/Qualifiers
      source          1..3920
                      /ApEinfo_revcolor="#ff9ccd"
                      /ApEinfo_fwdcolor="#ff9ccd"
                      /label="source:pPtPuc3"
      misc_feature    complement(29..51)
```

```
                     /ApEinfo_revcolor="#b1ff67"
                     /ApEinfo_fwdcolor="#b1ff67"
                     /label="pGEX_3_primer"
     rep_origin      758..856
                     /ApEinfo_revcolor="#b4abac"
                     /ApEinfo_fwdcolor="#b4abac"
                     /label="oriT_RP4"
     misc_feature    complement(969..1112)
                     /ApEinfo_revcolor="#b1ff67"
                     /ApEinfo_fwdcolor="#b1ff67"
                     /label="lacZ_a"
     misc_feature    1083..1105
                     /ApEinfo_revcolor="#b1ff67"
                     /ApEinfo_fwdcolor="#b1ff67"
                     /label="M13_pUC_fwd_primer"
     promoter        1098..1114
                     /ApEinfo_revcolor="#346ee0"
                     /ApEinfo_fwdcolor="#346ee0"
                     /label="M13_forward20_primer"
     promoter        1138..1157
                     /ApEinfo_revcolor="#346ee0"
                     /ApEinfo_fwdcolor="#346ee0"
                     /label="T3_promoter"
     CDS             complement(1815..2303)
                     /ApEinfo_revcolor="#ffef86"
                     /ApEinfo_fwdcolor="#ffef86"
                     /label="ORF frame 3"
     CDS             complement(1844..2476)
                     /ApEinfo_revcolor="#ffef86"
                     /ApEinfo_fwdcolor="#ffef86"
                     /label="ORF frame 1"
     gene            1860..2231
                     /ApEinfo_revcolor="#d59687"
                     /ApEinfo_fwdcolor="#d59687"
                     /label="bleo"
     misc_feature    1860..2234
                     /ApEinfo_revcolor="#b1ff67"
                     /ApEinfo_fwdcolor="#b1ff67"
                     /label="sh_ble"
     misc_feature    2542..3033
                     /ApEinfo_revcolor="#b1ff67"
                     /ApEinfo_fwdcolor="#b1ff67"
                     /label="CEN6_ARS4"
     gene            3231..3890
                     /ApEinfo_revcolor="#d59687"
                     /ApEinfo_fwdcolor="#d59687"
                     /label="HIS3"
     CDS             3231..3893
                     /ApEinfo_revcolor="#ffef86"
                     /ApEinfo_fwdcolor="#ffef86"
                     /label="ORF frame 3"
     misc_feature    4153..4422
                     /ApEinfo_revcolor="#d59687"
                     /ApEinfo_fwdcolor="#d59687"
                     /label="HSP70A-promoter"
     misc_feature    4423..4652
                     /ApEinfo_revcolor="#008080"
                     /ApEinfo_fwdcolor="#008080"
                     /label="RBCS2-promoter"
     misc_feature    4653..4806
                     /ApEinfo_revcolor="#c6c9d1"
                     /ApEinfo_fwdcolor="#c6c9d1"
                     /label="intron1 from RBCS2"
     misc_feature    4812..5683
                     /ApEinfo_revcolor="#b1ff67"
                     /ApEinfo_fwdcolor="#b1ff67"
                     /label="cGFP"
     misc_feature    4827..4981
                     /ApEinfo_revcolor="#9eafd2"
                     /ApEinfo_fwdcolor="#9eafd2"
                     /label="AtAct11 Intron"
     misc_feature    5685..5928
                     /ApEinfo_revcolor="#f58a5e"
                     /ApEinfo_fwdcolor="#f58a5e"
                     /label="RBCS2-3'UTR"
     source          5934..9154
                     /ApEinfo_revcolor="#ff9ccd"
```

```
                     /ApEinfo_fwdcolor="#ff9ccd"
                     /label="source:pPtPuc3"
     promoter        complement(5959..5977)
                     /ApEinfo_revcolor="#346ee0"
                     /ApEinfo_fwdcolor="#346ee0"
                     /label="M13_reverse_primer"
     misc_feature    complement(5976..5998)
                     /ApEinfo_revcolor="#b1ff67"
                     /ApEinfo_fwdcolor="#b1ff67"
                     /label="M13_pUC_rev_primer"
     promoter        complement(6012..6041)
                     /ApEinfo_revcolor="#346ee0"
                     /ApEinfo_fwdcolor="#346ee0"
                     /label="lac_promoter"
     CDS             complement(7124..7642)
                     /ApEinfo_revcolor="#ffef86"
                     /ApEinfo_fwdcolor="#ffef86"
                     /label="ORF frame 1"
     gene            7801..8616
                     /ApEinfo_revcolor="#d59687"
                     /ApEinfo_fwdcolor="#d59687"
                     /label="KanR2"
     CDS             7801..8616
                     /ApEinfo_revcolor="#ffef86"
                     /ApEinfo_fwdcolor="#ffef86"
                     /label="ORF frame 1"
     promoter        complement(8996..9024)
                     /ApEinfo_revcolor="#346ee0"
                     /ApEinfo_fwdcolor="#346ee0"
                     /label="AmpR_promoter"
ORIGIN
        1 tcgcgcgttt cggtgatgac ggtgaaaacc tctgacacat gcagctcccg gagacggtca
       61 cagcttgtct gtaagcggat gccgggagca gacaagcccg tcagggcgcg tcagcgggtg
      121 ttggcgggtg tcggggctgg cttaactatg cggcatcaga gcagattgta ctgagagtgc
      181 accagatcgt cttgccttgc tcgtcggtga tgtacttaca gctcgaagtg cctcttcttg
      241 atggagcgca tggggacgtg cttggcaatc acgcgcaccc cccggccgtt ttagcggcta
      301 aaaaagtcat ggctctgccc tcgggcggac cacgcccatc atgaccttgc caagctcgtc
      361 ctgcttctct tcgatcttcg ccagcagggc gaggatcgtg gcatcaccga accgcgccgt
      421 gcgcgggtcg tcggtgagcc agagtttcag caggccgccc aggcggccca ggtcgccatt
      481 gatgcgggcc agctcgcgga cgtgctcata gtccacgacg cccgtgattt tgtagccctg
      541 gccgacggcc agcaggtagg ccgacaggct catgccggcc gccgccgcct tttcctcaat
      601 cgctcttcgt tcgtctggaa ggcagtacac cttgataggt gggctgccct tcctggttgg
      661 cttggtttca tcagccatcc gcttgccctc atctgttacg ccggcggtag ccggccagcc
      721 tcgcagagca ggattcccgt tgagcaccgc caggtgcgaa taagggacag tgaagaagga
      781 acacccgctc gcgggtgggc ctacttcacc tatcctgccc ggctgacgcc gttggataca
      841 ccaaggaaag tctacacgaa ccctttggca aaatcctgta tatcgtgcga aaaaggatgg
      901 atataccgaa aaaatcgcta taatgacccc gaagcagggt tatgcagcgg aagatggcca
      961 ttcgccattc aggctgcgca actgttggga agggcgatcg gtgcgggcct cttcgctatt
     1021 acgccagctg gcgaaagggg gatgtgctgc aaggcgatta agttgggtaa cgccagggtt
     1081 ttcccagtca cgacgttgta aaacgacggc cagtgaccat gattacgcca agctcgaaat
     1141 taaccctcac taaagggaac aaaagctggt acctaacagg attagtgcaa ttcgagttga
     1201 atcactggga aaaacattgt cttctttttt atattatcat ttgcattagt gctgcagtcg
     1261 tagatacttg ttggttgaaa gacatcagct gggagggact ggactagcgt ttggtaagga
     1321 gacatacctg ttaacgttgg ttgcaaaatt ccatttcgcg atttatgtta tctgtaaatc
     1381 ctgatttgtc tggaattctt gatacttccg tttttttaga ggccaatgat tagcatcggc
     1441 gattctcaaa atagcatttt cgacatgcgg tgctgatttc ataaacatag acaacgcttt
     1501 tacatgtaaa agtaacttgc ggacttggaa cagtgctctg tttttggtgt gaacgtaact
     1561 cagcaatatt tctgtgctag caaggttttt tatgatcgac cgaagatctc aaaactccgg
     1621 gtctttcaac tgtctgacta gaccatgttc gtaacgtcgg acagcagctt tcgttgtact
     1681 ggtagaattt ctacgtgcga agcacgtgta ggcaggttga acgacgatcc ctgccgatgg
     1741 atggattggc acgcggcgga acgctttcgt gatctacacc acctggatct tcacatatct
     1801 tcgaaatcga aaaattaacc aagtcgacgg tatcgataat attctagctg agggtaccca
     1861 tggccaagtt gaccagtgcc gttccggtgc tcaccgcgcg cgacgtcgcc ggagcggtcg
     1921 agttctggac cgaccggctc gggttctccc gggacttcgt ggaggacgac ttcgccggtg
     1981 tggtccggga cgacgtgacc ctgttcatca gcgcggtcca ggaccaggtg gtgccggaca
     2041 acaccctggc ctgggtgtgg gtgcgcggcc tggacgagct gtacgccgag tggtcggagg
     2101 tcgtgtccac gaacttccgg gacgcctccg ggccggccat gaccgagatc ggcgagcagc
     2161 cgtgggggcg ggagttcgcc ctgcgcgacc cggccggcaa ctgcgtgcac ttcgtggccg
     2221 aggagcagga ctgaccgacg ccgaccaaca ccgccggtcc gacgcggccc gacgggtccg
     2281 aggcctcgga gatctgggcc catgcggccg caacaactac ctcgactttg gctgggacac
     2341 tttcagtgag gacaagaagc ttcagaagcg tgctatcgaa ctcaaccagg gacgtgcggc
     2401 acaaatgggc atccttgctc tcatggtgca cgaacagttg ggagtctcta tccttcctta
     2461 aaaatttaat tttcattagt tgcagtcact ccgctttggt ttcacagtca ggaataacac
     2521 tagctcgtct tcagcgagca tcacgtgcta taaaaataat tataatttaa atttttttaat
     2581 ataaatatat aaattaaaaa tagaaagtaa aaaaagaaat taaagaaaaa atagtttttg
     2641 ttttccgaag atgtaaaaga ctctaggggg atcgccaaca aatactacct tttaccttgc
     2701 tcttcctgct ctcaggtatt aatgccgaat tgtttcatct tgtctgtgta gaagaccaca
```

2761 cacgaaaatc ctgtgatttt acattttact tatcgttaat cgaatgtata tctatttaat
2821 ctgcttttct tgtctaataa atatatatgt aaagtacgct tttgttgaa attttttaaa
2881 cctttgttta tttttttttc ttcattccgt aactcttcta ccttctttat ttactttcta
2941 aaatccaaat acaaaacata aaaataaata aacacagagt aaattcccaa attattccat
3001 cattaaaaga tacgaggcgc gtgtaagtta caggcaagcg atcctagtac actctatatt
3061 tttttatgcc tcggtaatga ttttcatttt ttttttccac ctagcggatg actctttttt
3121 tttcttagcg attggcatta tcacataatg aattatacat tatataaagt aatgtgattt
3181 cttcgaagaa tatactaaaa aatgagcagg caagataaac gaaggcaaag atgacagagc
3241 agaaagccct agtaaagcgt attacaaatg aaaccaagat tcagattgcg atctctttaa
3301 agggtggtcc cctagcgata gagcactcga tcttcccaga aaaagaggca gaagcagtag
3361 cagaacaggc cacacaatcg caagtgatta acgtccacac aggtataggg tttctggacc
3421 atatgataca tgctctggcc aagcattccg gctggtcgct aatcgttgag tgcattggtg
3481 acttacacat agacgaccat cacaccactg aagactgcgg gattgctctc ggtcaagctt
3541 ttaaagaggc cctaggggcc gtgcgtggag taaaaaggtt tggatcagga tttgcgcctt
3601 tggatgaggc actttccaga gcggtggtag atctttcgaa caggccgtac gcagttgtcg
3661 aacttggttt gcaaagggag aaagtaggag atctctcttg cgagatgatc ccgcattttc
3721 ttgaaagctt tgcagaggct agcagaatta ccctccacgt tgattgtctg cgaggcaaga
3781 atgatcatca ccgtagtgag agtgcgttca aggctcttgc ggttgccata agagaagcca
3841 cctcgcccaa tggtaccaac gatgttccct ccaccaaagg tgttcttatg tagttttaca
3901 caggagtctg gacttgacct ctagaatccc cagcatgcct gctattgtct tcccaatcct
3961 ccccccttgct gtcctgcccc accccacccc ccagaataga atgacaccta ctcagacaat
4021 gcgatgcaat ttcctcattt tattaggaaa ggacagtggg agtggcacct tccagggtca
4081 aggaaggcac gggggagggg caaacaacag atggctggca actagaaggc acagtcgagg
4141 ctgatagcga gctcgctgag gcttgacatg attggtgcgt atgtttgtat gaagctacag
4201 gactgatttg gcgggctatg agggcggggg aagctctgga agggccgcga tggggcgcgc
4261 ggcgtccaga aggcgccata cggcccgctg gcggcaccca tccggtataa aagcccgcga
4321 ccccgaacgg tgacctccac tttcagcgac aaacgagcac ttatacatac gcgactattc
4381 tgccgctata cataaccact cagctagctt aagatcccat caagcttgca tgccgggcgc
4441 gccagaagga gcgcagccaa accaggatga tgtttgatgg ggtatttgag cacttgcaac
4501 ccttatccgg aagcccccctg gcccacaaag gctaggcgcc aatgcaagca gttcgcatgc
4561 agcccctgga gcggtgccct cctgataaac cggccagggg gcctatgttc tttacttttt
4621 tacaagagaa gtcactcaac atcttaaaat ggccaggtga gtcgacgagc aagcccggcg
4681 gatcaggcag cgtgcttgca gatttgactt gcaacgcccg cattgtgtcg acgaaggctt
4741 ttggctcctc tgtcgctgtc tcaagcagca tctaaccctg cgtcgccgtt tccatttgca
4801 ggatgctcga gatggccaag ggcgaggtaa taagatcttc aacacctaca ccattttttt
4861 aatcactact acccattgca ttgaacaaac ttccaagttc ttcttagctt cagattaaga
4921 aagtaccctt tcttggcttt gttgatgtgg taccattgtc cattgtcttg tgtgtttcca
4981 ggagctgttc accggtgtgg tccccatcct ggtggagctg gacggcgacg tgaacggcca
5041 caagttctcc gtctccggcg agggtgaggg tgacgccacc tacggcaagc tgaccctgaa
5101 gttcatctgc accaccggca agctgcccgt gccctggccc accctggtca ccaccctgac
5161 ctacggtgtg cagtgcttct cccgctaccc cgaccacatg aagcagcacg acttcttcaa
5221 gtccgccatg cccgagggct acgtgcagga gcgcaccatc ttcttcaagg acgacggcaa
5281 ctacaagacc cgcgccgagg tcaagttcga gggcgacacc ctggtgaacc gcatcgagct
5341 gaagggcatc gacttcaagg aggacggcaa catcctgggc cacaagctgg agtacaacta
5401 caactcccac aacgtgtaca tcatggccga caagcagaag aacggcatca aggtgaactt
5461 caagatccgc cacaacatcg aggacggctc cgtgcagctg gccgaccact accagcagaa
5521 cacccccatc ggcgatggcc ccgtgctgct gcccgacaac cactacctgt ccatccagtc
5581 cgccctgtcc aaggacccca acgagaagcg cgaccacatg gtcctgctgg agttcgtcac
5641 cgctgccggc atcacccacg gcatggacga gctgtacaag taaggatccc cgctccgtgt
5701 aaatggaggc gctcgttgat ctgagccttg cccccctgacg aacggcggtg gatggaagat
5761 actgctctca agtgctgaag cggtagctta gctccccgtt tcgtgctgat cagtcttttt
5821 caacacgtaa aaagcggagg agttttgcaa ttttgttggt tgtaacgatc ctccgttgat
5881 tttggcctct ttctccatgg gcgggctggg cgtatttgaa gcgggtaccc ccctgcaggc
5941 atgcaagctt ggcgtaatca tggtcatagc tgtttcctgt gtgaaattgt tatccgctca
6001 caattccaca caacatacga gccggaagca taaagtgtaa agcctggggt gcctaatgag
6061 tgagctaact cacattaatt gcgttgcgct cactgcccgc tttccagtcg ggaaacctgt
6121 cgtgccagct gcattaatga atcggccaac gcgcggggag aggcggtttg cgtattgggc
6181 gctcttccgc ttcctcgctc actgactcgc tgcgctcggt cgttcggctg cggcgagcgg
6241 tatcagctca ctcaaaggcg gtaatacggt tatccacaga atcaggggat aacgcaggaa
6301 agaacatgtg agcaaaaggc cagcaaaagg ccaggaaccg taaaaaggcc gcgttgctgg
6361 cgtttttcca taggctccgc cccccctgacg agcatcacaa aaatcgacgc tcaagtcaga
6421 ggtggcgaaa cccgacagga ctataaagat accaggcgtt tcccccctgga agctccctcg
6481 tgcgctctcc tgttccgacc ctgccgctta ccggatacct gtccgccttt ctcccttcgg
6541 gaagcgtggc gctttctcat agctcacgct gtaggtatct cagttcggtg taggtcgttc
6601 gctccaagct gggctgtgtg cacgaacccc ccgttcagcc cgaccgctgc gccttatccg
6661 gtaactatcg tcttgagtcc aacccggtaa gacacgactt atcgccactg gcagcagcca
6721 ctggtaacag gattagcaga gcgaggtatg taggcggtgc tacagagttc ttgaagtggt
6781 ggcctaacta cggctacact agaagaacag tatttggtat ctgcgctctg ctgaagccag
6841 ttaccttcgg aaaaagagtt ggtagctctt gatccggcaa acaaaccacc gctggtagcg
6901 gtggtttttt tgtttgcaag cagcagatta cgcgcagaaa aaaaggatct caagaagatc
6961 ctttgatctt ttctacgggg tctgacgctc agtggaacga aaactcacgt taagggattt
7021 tggtcatgag attatcaaaa aggatcttca cctagatcct tttaaattaa aaatgaagtt
7081 ttaaatcaat ctaaagtata tatgagtaaa cttggtctga cagttaccaa tgcttaatca
7141 gtgaggcacc tatctcagcg atctgtctat ttcgttcatc catagttgcc tgactccccg
7201 tcgtgtagat aactacgata cgggagggct taccatctgg ccccagtgct gcaatgatac
7261 cgcgagaccc acgctcaccg gctccagatt tatcagcaat aaaccagcca gccggaaggg
7321 ccgagcgcag aagtggtcct gcaactttat ccgcctccat ccagtctatt aattgttgcc

```
     7381 gggaagctag agtaagtagt tcgccagtta atagtttgcg caacgttgtt gccattgcta
     7441 caggcatcgt ggtgtcacgc tcgtcgtttg gtatggcttc attcagctcc ggttcccaac
     7501 gatcaaggcg agttacatga tcccccatgt tgtgcaaaaa agcggttagc tccttcggtc
     7561 ctccgatcgt tgtcagaagt aagttggccg cagtgttatc actcatggtt atggcagcac
     7621 tgcataattc tcttactgtc atgccatccg taagatgctt ttctgtgact ggtgagtcga
     7681 tttattcaac aaagccacgt tgtgtctcaa aatctctgat gttacattgc acaagataaa
     7741 aatatatcat catgaacaat aaaactgtct gcttacataa acagtaatac aaggggtgtt
     7801 atgagccata ttcaacggga aacgtcttgc tcgaggccgc gattaaattc caacatggat
     7861 gctgatttat atgggtataa atgggctcgc gataatgtcg ggcaatcagg tgcgacaatc
     7921 tatcgattgt atgggaagcc cgatgcgcca gagttgtttc tgaaacatgg caaaggtagc
     7981 gttgccaatg atgttacaga tgagatggtc agactaaact ggctgacgga atttatgcct
     8041 cttccgacca tcaagcattt tatccgtact cctgatgatg catggttact caccactgcg
     8101 atccccggga aaacagcatt ccaggtatta gaagaatatc ctgattcagg tgaaaatatt
     8161 gttgatgcgc tggcagtgtt cctgcgccgg ttgcattcga ttcctgtttg taattgtcct
     8221 tttaacagcg atcgcgtatt tcgtctcgct caggcgcaat cacgaatgaa taacggtttg
     8281 gttgatgcga gtgattttga tgacgagcgt aatggctggc ctgttgaaca agtctggaaa
     8341 gaaatgcata agcttttgcc attctcaccg gattcagtcg tcactcatgg tgatttctca
     8401 cttgataacc ttatttttga cgaggggaaa ttaataggtt gtattgatgt tggacgagtc
     8461 ggaatcgcag accgatacca ggatcttgcc atcctatgga actgcctcgg tgagttttct
     8521 ccttcattac agaaacggct ttttcaaaaa tatggtattg ataatcctga tatgaataaa
     8581 ttgcagtttc atttgatgct cgatgagttt ttctaatcag aattggttaa ttggttgtaa
     8641 cactggcact caaccaagtc attctgagaa tagtgtatgc ggcgaccgag ttgctcttgc
     8701 ccggcgtcaa tacgggataa taccgcgcca catagcagaa ctttaaaagt gctcatcatt
     8761 ggaaaacgtt cttcggggcg aaaactctca aggatcttac cgctgttgag atccagttcg
     8821 atgtaaccca ctcgtgcacc caactgatct tcagcatctt ttactttcac cagcgtttct
     8881 gggtgagcaa aaacaggaag gcaaaatgcc gcaaaaaagg gaataagggc gacacggaaa
     8941 tgttgaatac tcatactctt cctttttcaa tattattgaa gcatttatca gggttattgt
     9001 ctcatgagcg gatacatatt tgaatgtatt tagaaaaata aacaaatagg ggttccgcgc
     9061 acatttcccc gaaaagtgcc acctgacgtc taagaaacca ttattatcat gacattaacc
     9121 tataaaaata ggcgtatcac gaggcccttt cgtc
//
```

<u>*Aurantiochytrium limacinum*</u>

*pUC19_18GZG*
*(Addgene Plasmid #117228)*

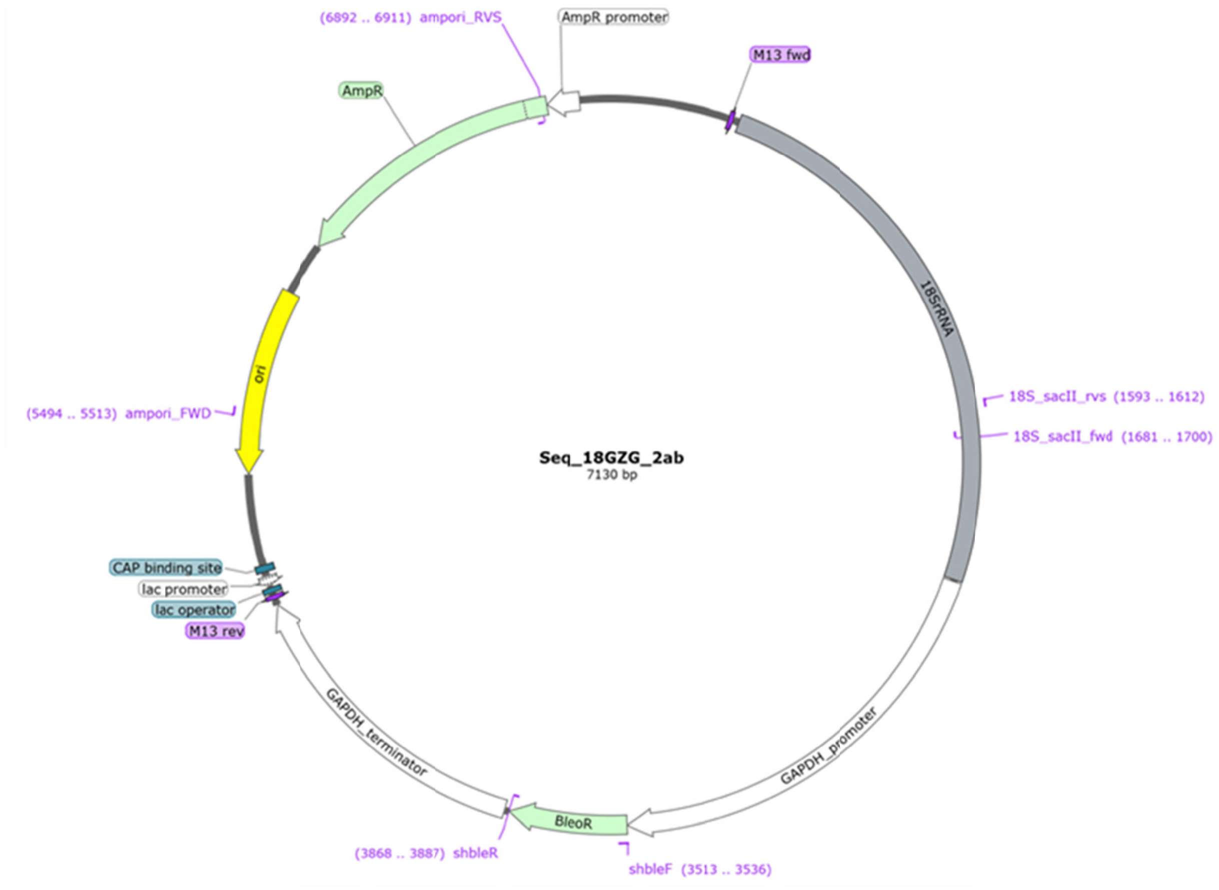


```
> pUC19_18GZG
TCGCGCGTTTCGGTGATGACGGTGAAAACCTCTGACACATGCAGCTCCCGGAGACGGTCACAGCTTGTCT
GTAAGCGGATGCCGGGAGCAGACAAGCCCGTCAGGGCGCGTCAGCGGGTGTTGGCGGGTGTCGGGGCTGG
CTTAACTATGCGGCATCAGAGCAGATTGTACTGAGAGTGCACCATATGCGGTGTGAAATACCGCACAGAT
GCGTAAGGAGAAAATACCGCATCAGGCGCCATTCGCCATTCAGGCTGCGCAACTGTTGGGAAGGGCGATC
GGTGCGGGCCTCTTCGCTATTACGCCAGCTGGCGAAAGGGGGATGTGCTGCAAGGCGATTAAGTTGGGTA
ACGCCAGGGTTTTCCCAGTCACGACGTTGTAAAACGACGGCCAGTGAATTCGAGCTCGGTACCCCTTGTT
ACGACTTCACCTTCCTCTAAACAATAAGATTCACCCGAGTTCTGCCTCTGTCCAAAAATTAATCCAAACA
GAAACATCCCATGGTTTCATCGGACCGTTCAATCGGTAGGTGCGACGGGCGGTGTGTACAAAGGGCAGGG
ACGTATTCAATGCAAGCTGATGACTTGCGTTTACTAGGAATTCCTCGTTGGAGATTAATAATTGCAAAAA
TCTAGCCCCAGCACGATGAGCGTTCCAAGGATTAGCCAGGCCTTCCGACCAAGCACTCAATTCCAAAAAT
TGAAATTAAAACCCGATGAACCCATCAGTGTAGCGCGCGTGCGGCCCAGAACATCTAAGGGCATCACAGA
CCTGTTATTGCCTCGAACTTCCTGCCCGTAAACCGGACATGTCCCTCTAAGAAGTAAAAAACGTACTATG
TTGCCATACCACGCACTATTTAGTAGGCCGAGGTCTCGTTCGTTAACGGAATTAACCAGACAAATCACTC
CACCAACTAAGAACGGCCATGCACCACCACCCATAGAATCATGAAAGAGCTCTCAATCTGTCAATCCTAC
CTATGTCTGGACCTGGTAAGTTTTCCCGTGTTGAGTCAAATTAAGCCGCAGGCTCCACTCCTGGTGGTGC
CCTTCCGTCAATTCCTTTAAGTTTCAGCCTTGCGACCATACTCCCCCCGGAACCCAAAGACTTTGATTTC
TCATGTGCTGCTGCTGAGGCCCATTAAAAAAAGCACTCAACAATCGCAAGTCGGCATCGTTTACGGTCTA
GACTACGATGGTATCTAATCATCTTCGATCCCCAGACTTTCGTTCTTGATTAATGAAAACATGCTTGGTA
AATGCCTTCGCTCTAGTTCGTCTTTCGGAAATCCAAGAATTTCACCTCTAGCTCCTAAATACGAATACCC
CCAACTGTTCCTATTAACCATTACTCAGGCGTGCAAACCAACAAAATAGCACCCAAGTCCTATCTTATCA
TCCCATAATAAACATACCGGTCATACGACCTGCTTGGAACACTCTGCTTTGATTACAGTGAAAGATTTCT
CCCCTATAAAGAAAAGAAAAAGATGGCCAAGGCAACACAGACAATCAATCCCCATTCAGGGAAAGCACCG
GTCGCCCATGCCAGAAATTCAACTACGAGCTTTTTAACTGCAACAACTTTAGCATATGCTTCTGGAGCTG
GAATTACCGCGGCTGCTGGCACCAGACTTGCCCTCCAGTTGATCCTCGATGAGGGTTTTACATTGCTCTC
ATTCCGATAGCAAAACGCATACACGCTTCGCATCGATATTTCTCGTCACTACCTCGTGGAGTCCACAGTG
```

GGTAATTTACGCGCCTGCTGCTATCCTTGGATATGGTAGCCGTCTCTCAGGCTCCCTCTCCGGAGTCGAG
CCCTAACTCTCCGTCACCCGTTATAGTCACCGTAGTCCAATACACTACCGTCGACAACTGATGGGGCAGA
AACTCAAACGATTCATCGACCAAAATAGTCAATCTGCTCAATTATCATGATTCACCAATAAAATCGGCTT
CAATCTAATAAGTGCAGCCCCATACAGGGCTCTTACAGCATGTATTATTTCCAGAATTACTGCAGGTATC
CATATAAAAGAAACTACCGAAGAAATTATTACTGATATAATGAGCCGTTCGCAGTCTCACAGTACAATCG
CTTATACTTACACATGCATGGCTTAATCTTTGAGACGAGCATATGACTACTGGGGTACCCTTGATCTTGT
GAGGGCTCCACTGGAATTTTCCGACACAAGAAAGTGCCAGTGGACAAAGGGGGAAGTGGGCCTCGAAATA
GCGGTATGGTTATGTGAGGGATACGGGGCGGAGTTCCGGCGTTCCCTCAGGCCTTCCTCTCTGCGCCTCA
GGCAAGTTTCTAAGAAAAGTGCTTTTGGATAATAATCTTGTATGGAAATGTGAAGCGCAAGTGCGCAGAA
CCTAGAAAATCTAAAACAAAAGAAAAAAGAGATCGTCACCCGAGCAGCAGAATCAATTCACTCCAAGGTA
AGTAAGGCGTGCTGTGCGGGAGCGCCCCCGGCCAGTCAGTGTTGGAGCTTGGGATGAGTTGCATCTGCGC
GAAGGGTTGGCCCATCCATCTCAAGTCCTTTCCTGGCGGTTTCGCCCGCGTCACCAGCCGCCTGGTGCTC
CTGATGGATGGGGACTCTGGATGGTCGAAAGAGGATGTGTTGTATCTATCTGTCTGATAAGGTAAAAGGG
GACGGCTTCTGTACTTTCCTTCTTCGCTCGCTCCCGATCGTGTCTTCTAGGATGCGCGCTTTTGATGTGC
TGAAGATTCCGAGCACCGTGTGCTATGCCGTCTCCGTCTTGCCTGTGTTGGCGGCTAGCTAGCGCTGAGG
GTTGTGGACGGCCTTGGGAATTAGGAAAAGAAAGATACTCGTGACGTTGAAACGTCCTGTTCTTTTTCTT
TTCTAGTATTTTCTATTCCAGTCTTTACTTTCTGTTCCTTTCATTTCACTTCAAGTACATCGTCACCTCT
ATCGATCTAGTGTGAGCCGAATTGAGCCTCCTGTCATGTAGCAGGAGGAACTACAGAGGTTCCGACTGGT
ACGAACAAACGAATCGGCGAACGGAGAGTAGGCATGAAGTTGTTGTTGTCCAAGATCAAAAAGATAGAAA
CGAATATCTTTCTTGTTCTGCTACCTACATTGAAACGGACATGAAAGACGACAATTCTAGATGAAGACAC
TGCCAAAAGGGAAGAAGCGCTCGGCCACCGCAGCAAGAAAGGCAAGAGAGAGAAGATAAAGTAAATTTTC
GAGAGAACAAAAGAAATGAATGCAGGAAGGAAGGAAGGAAGGAGGGGAGGTAGTGAAAGAACGCGTGACA
AGATTTGTAATGAAGAACATGGCATGAAAGAACGAACAGGGGGGACTGACGATTTGAGGGACTGATGTGC
GCAATTGAATCTTTTTCATTTGCATTGCGGCTGCGGCGGCAAACAAAACAAAAATAATTATTCGGCATTC
ACTTTGCTTGCGTTGTTGGACAACATAACCATAACAGAAACAAAAGCAAGGAAACCTAGCGCACACATAA
ACACCAAGCAAAATGGCCAAGTTGACCAGTGCCGTTCCGGTGCTCACCGCGCGCGACGTCGCCGGAGCGG
TCGAGTTCTGGACCGACCGGCTCGGGTTCTCCCGGGACTTCGTGGAGGACGACTTCGCCGGTGTGGTCCG
GGACGACGTGACCCTGTTCATCAGCGCGGTCCAGGACCAGGTGGTGCCGGACAACACCCTGGCCTGGGTG
TGGGTGCGCGGCCTGGACGAGCTGTACGCCGAGTGGTCGGAGGTCGTGTCCACGAACTTCCGGGACGCCT
CCGGGCCGGCCATGACCGAGATCGGCGAGCAGCCGTGGGGGCGGGAGTTCGCCCTGCGCGACCCGGCCGG
CAACTGCGTGCACTTCGTGGCCGAGGAGCAGGACTGAGGGGATCCTCTAGAGTCGACATGTACCCAATAC
CACACCGGTAGCTTCTCGCGGCGGCTGACAAGAAAGATTGTTTTTACACATTTCGAGGCATTGATGACCC
TTATCGACCTATCGTCTCAGATCATAAAATGCACGAGCATGAAGTACGCGTGTTGTGACTTGCTTGCGTC
CCACCTTCTAGATGGCTTTCTTCTTTTCAAGTGATTAAAACACCACAGTAGATGAGGACTCTTAGTAAGC
ACTGAAGAAGCGAGTAAATAGCCCTCATCCCGTTTCCCTCTTTTCTAACACACTTTGTTTGGAATTCTAA
AATATCTTTTATCATCTCTTTTCATTCACAAAACTAGTATTTCTGCATTAGAAATCATTATCCTATCTGG
CACTTTCACTCTGACAAGAACTTGCGTACATGGCGGATCCTGGCAATCATTTTACTTGTACAGACCCAAG
ACTTTGTGAGTCAGTAAATAGTAAAGAAATGCAGAATAATCATTAGATAATTGCAAAACCCTGATCTTCA
AAATGTTATATCACAAGTACCTACCAAGACATTTGTATCTTCTTTTTGTCTTATGCATTTTTATTTCCCT
TAGCGCAATTAAAAATATTAAATAACTGACGACTCTTTTCTTAAGGATTCGCGGTTTGCTAGTTGCACTA
TTGAAAAGAGCACGGTATTAATTTCATAGTTTTACTTTGTTGGCAATCTCAGCAAACATGGGTTTTAATT
TTAAAATTAAATAATATTTGTGTTAGACTCAAAAGATACCTATACAACCCTTAGGGCTCATCGTCAACAT
TTGCAAATTAAAACGATTCAGCTCCTTTGGTATGGTCATGTCTGTTTCTGATGCACTGAAAAACCTTGGA
CGAACGACACATTTTATTTTCATTAGGACATCTAAAAAACGTTGTAAAGAGTATACAATAAGCATAAAAA
AGGAGAAGAAAAAAAAGAAAGACATACATGGTTCTGCTCTTCTAGTGCTTCAAGGCATGCAAGCTTGGCG
TAATCATGGTCATAGCTGTTTCCTGTGTGAAATTGTTATCCGCTCACAATTCCACACAACATACGAGCCG
GAAGCATAAAGTGTAAAGCCTGGGGTGCCTAATGAGTGAGCTAACTCACATTAATTGCGTTGCGCTCACT
GCCCGCTTTCCAGTCGGGAAACCTGTCGTGCCAGCTGCATTAATGAATCGGCCAACGCGCGGGGAGAGGC
GGTTTGCGTATTGGGCGCTCTTCCGCTTCCTCGCTCACTGACTCGCTGCGCTCGGTCGTTCGGCTGCGGC
GAGCGGTATCAGCTCACTCAAAGGCGGTAATACGGTTATCCACAGAATCAGGGGATAACGCAGGAAAGAA
CATGTGAGCAAAAGGCCAGCAAAAGGCCAGGAACCGTAAAAAGGCCGCGTTGCTGGCGTTTTTCCATAGG
CTCCGCCCCCCTGACGAGCATCACAAAAATCGACGCTCAAGTCAGAGGTGGCGAAACCCGACAGGACTAT
AAAGATACCAGGCGTTTCCCCCTGGAAGCTCCCTCGTGCGCTCTCCTGTTCCGACCCTGCCGCTTACCGG
ATACCTGTCCGCCTTTCTCCCTTCGGGAAGCGTGGCGCTTTCTCATAGCTCACGCTGTAGGTATCTCAGT
TCGGTGTAGGTCGTTCGCTCCAAGCTGGGCTGTGTGCACGAACCCCCCGTTCAGCCCGACCGCTGCGCCT
TATCCGGTAACTATCGTCTTGAGTCCAACCCGGTAAGACACGACTTATCGCCACTGGCAGCAGCCACTGG
TAACAGGATTAGCAGAGCGAGGTATGTAGGCGGTGCTACAGAGTTCTTGAAGTGGTGGCCTAACTACGGC
TACACTAGAAGAACAGTATTTGGTATCTGCGCTCTGCTGAAGCCAGTTACCTTCGGAAAAAGAGTTGGTA
GCTCTTGATCCGGCAAACAAACCACCGCTGGTAGCGGTGGTTTTTTTGTTTGCAAGCAGCAGATTACGCG
CAGAAAAAAAGGATCTCAAGAAGATCCTTTGATCTTTTCTACGGGGTCTGACGCTCAGTGGAACGAAAAC
TCACGTTAAGGGATTTTGGTCATGAGATTATCAAAAAGGATCTTCACCTAGATCCTTTTAAATTAAAAAT
GAAGTTTTAAATCAATCTAAAGTATATATGAGTAAACTTGGTCTGACAGTTACCAATGCTTAATCAGTGA
GGCACCTATCTCAGCGATCTGTCTATTTCGTTCATCCATAGTTGCCTGACTCCCCGTCGTGTAGATAACT
ACGATACGGGAGGGCTTACCATCTGGCCCCAGTGCTGCAATGATACCGCGAGACCCACGCTCACCGGCTC
CAGATTTATCAGCAATAAACCAGCCAGCCGGAAGGGCCGAGCGCAGAAGTGGTCCTGCAACTTTATCCGC
CTCCATCCAGTCTATTAATTGTTGCCGGGAAGCTAGAGTAAGTAGTTCGCCAGTTAATAGTTTGCGCAAC
GTTGTTGCCATTGCTACAGGCATCGTGGTGTCACGCTCGTCGTTTGGTATGGCTTCATTCAGCTCCGGTT
CCCAACGATCAAGGCGAGTTACATGATCCCCCATGTTGTGCAAAAAAGCGGTTAGCTCCTTCGGTCCTCC
GATCGTTGTCAGAAGTAAGTTGGCCGCAGTGTTATCACTCATGGTTATGGCAGCACTGCATAATTCTCTT
ACTGTCATGCCATCCGTAAGATGCTTTTCTGTGACTGGTGAGTACTCAACCAAGTCATTCTGAGAATAGT
GTATGCGGCGACCGAGTTGCTCTTGCCCGGCGTCAATACGGGATAATACCGCGCCACATAGCAGAACTTT
AAAAGTGCTCATCATTGGAAAACGTTCTTCGGGGCGAAAACTCTCAAGGATCTTACCGCTGTTGAGATCC
AGTTCGATGTAACCCACTCGTGCACCCAACTGATCTTCAGCATCTTTTACTTTCACCAGCGTTTCTGGGT
GAGCAAAAACAGGAAGGCAAAATGCCGCAAAAAAGGGAATAAGGGCGACACGGAAATGTTGAATACTCAT
ACTCTTCCTTTTTCAATATTATTGAAGCATTTATCAGGGTTATTGTCTCATGAGCGGATACATATTTGAA
TGTATTTAGAAAAATAAACAAATAGGGGTTCCGCGCACATTTCCCCGAAAAGTGCCACCTGACGTCTAAG
AAACCATTATTATCATGACATTAACCTATAAAAATAGGCGTATCACGAGGCCCTTTCGTC

#### *Caecitellus sp.*

No vector: experiment inserted fluorescein isothiocyanate conjugated to dextran.

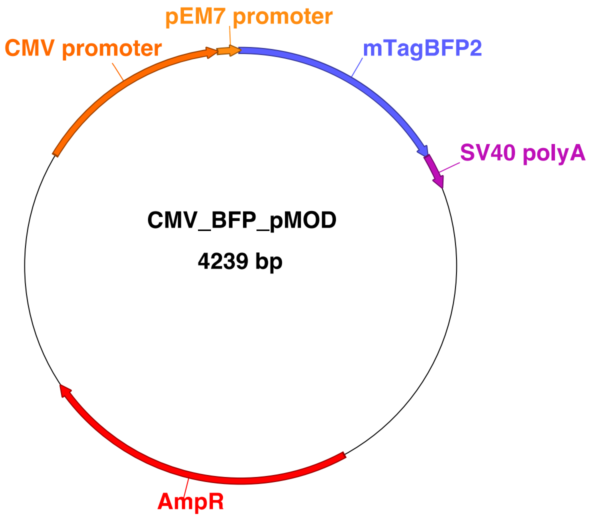


**Figure 1:** CMV_BFP_pMOD plasmid used for electroporation of model organism *Nannochloropsis oceanica* and marine natural samples. BFP: Blue fluorescent protein (mTagBFP2).

```
FEATURES             Location/Qualifiers
     misc_feature    1..705
                     /locus_tag="mTagBFP2"
                     /ApEinfo_label="mTagBFP2"
                     /ApEinfo_fwdcolor="#5a5eff"
                     /ApEinfo_revcolor="green"
                     /ApEinfo_graphicformat="arrow_data {{0 1 2 0 0 -1} {} 0}
                     width 5 offset 0"
     misc_feature    1788..2783
                     /locus_tag="AmpR"
                     /ApEinfo_label="AmpR"
                     /ApEinfo_fwdcolor="#ff0000"
                     /ApEinfo_revcolor="green"
                     /ApEinfo_graphicformat="arrow_data {{0 1 2 0 0 -1} {} 0}
                     width 5 offset 0"
     misc_feature    4172..4239
                     /locus_tag="pEM7 promoter"
                     /ApEinfo_label="pEM7 promoter"
                     /ApEinfo_fwdcolor="#ff8c10"
                     /ApEinfo_revcolor="green"
                     /ApEinfo_graphicformat="arrow_data {{0 1 2 0 0 -1} {} 0}
                     width 5 offset 0"
     misc_feature    3549..4167
                     /locus_tag="CMV promoter"
                     /ApEinfo_label="CMV promoter"
```

```
                     /ApEinfo_fwdcolor="#ff6a00"
                     /ApEinfo_revcolor="green"
                     /ApEinfo_graphicformat="arrow_data {{0 1 2 0 0 -1} {} 0}
                     width 5 offset 0"
     misc_feature    706..810
                     /locus_tag="SV40 polyA signal sequence"
                     /ApEinfo_label="SV40 polyA signal sequence"
                     /ApEinfo_fwdcolor="#be0fb7"
                     /ApEinfo_revcolor="green"
                     /ApEinfo_graphicformat="arrow_data {{0 1 2 0 0 -1} {} 0}
                     width 5 offset 0"
ORIGIN
        1 atgagcgagc tgattaagga gaacatgcac atgaagctgt acatggaggg caccgtggac
       61 aaccatcact tcaagtgcac atccgagggc gaaggcaagc cctacgaggg cacccagacc
      121 atgagaatca aggtggtcga gggcggccct ctcccttcg ccttcgacat cctggctact
      181 agcttcctct acggcagcaa gaccttcatc aaccacaccc agggcatccc cgacttcttc
      241 aagcagtcct tccctgaggg cttcacatgg gagagagtca ccacatacga agacgggggc
      301 gtgctgaccg ctacccagga caccagcctc caggacggct gcctcatcta caacgtcaag
      361 atcagagggg tgaacttcac atccaacggc cctgtgatgc agaagaaaac actcggctgg
      421 gaggccttca ccgagacgct gtaccccgct gacggcggcc tggaaggcag aaacgacatg
      481 gccctgaagc tcgtgggcgg gagccatctg atcgcaaacg ccaagaccac atatagatcc
      541 aagaaacccg ctaagaacct caagatgcct ggcgtctact atgtggacta cagactggaa
      601 agaatcaagg aggccaacaa cgagacctac gtcgagcagc acgaggtggc agtggccaga
      661 tactgcgacc tccctagcaa actggggcac aagcttaatt aaTGATGGTT ACAAATAAAG
      721 CAATAGCATC ACAAATTTCA CAAATAAAGC ATTTTTTTCA CTGCATTCTA GTTGTGGTTT
      781 GTCCAAACTC ATCAATGTAT CTTATCATGT CTGATTCGAG CTCGGTACCC GGGGATCCTC
      841 TAGAGTCGAC CTGCAGGCAT GCAAGCTTGC CAACGACTAC GCACTAGCCA ACAAGAGCTT
      901 CAGGGTTGAG ATGTGTATAA GAGACAGCTG TCTTAATGAA TCGGCCAACG CGCGGGGAGA
      961 GGCGGTTTGC GTATTGGGCG CTCTTCCGCT TCCTCGCTCA CTGACTCGCT GCGCTCGGTC
     1021 GTTCGGCTGC GGCGAGCGGT ATCAGCTCAC TCAAAGGCGG TAATACGGTT ATCCACAGAA
     1081 TCAGGGGATA ACGCAGGAAA GAACATGTGA GCAAAAGGCC AGCAAAAGGC CAGGAACCGT
     1141 AAAAAGGCCG CGTTGCTGGC GTTTTTCCAT AGGCTCCGCC CCCCTGACGA GCATCACAAA
     1201 AATCGACGCT CAAGTCAGAG GTGGCGAAAC CCGACAGGAC TATAAAGATA CCAGGCGTTT
     1261 CCCCCTGGAA GCTCCCTCGT GCGCTCTCCT GTTCCGACCC TGCCGCTTAC CGGATACCTG
     1321 TCCGCCTTTC TCCCTTCGGG AAGCGTGGCG CTTTCTCATA GCTCACGCTG TAGGTATCTC
     1381 AGTTCGGTGT AGGTCGTTCG CTCCAAGCTG GGCTGTGTGC ACGAACCCCC CGTTCAGCCC
     1441 GACCGCTGCG CCTTATCCGG TAACTATCGT CTTGAGTCCA ACCCGGTAAG ACACGACTTA
     1501 TCGCCACTGG CAGCAGCCAC TGGTAACAGG ATTAGCAGAG CGAGGTATGT AGGCGGTGCT
     1561 ACAGAGTTCT TGAAGTGGTG GCCTAACTAC GGCTACACTA GAAGGACAGT ATTTGGTATC
     1621 TGCGCTCTGC TGAAGCCAGT TACCTTCGGA AAAAGAGTTG GTAGCTCTTG ATCCGGCAAA
     1681 CAAACCACCG CTGGTAGCGG TGGTTTTTTT GTTTGCAAGC AGCAGATTAC GCGCAGAAAA
     1741 AAAGGATCTC AAGAAGATCC TTTGATCTTT TCTACGGGGT CTGACGCTCA GTGGAACGAA
     1801 AACTCACGTT AAGGGATTTT GGTCATGAGA TTATCAAAAA GGATCTTCAC CTAGATCCTT
     1861 TTAAATTAAA AATGAAGTTT TAAATCAATC TAAAGTATAT ATGAGTAAAC TTGGTCTGAC
     1921 AGTTACCAAT GCTTAATCAG TGAGGCACCT ATCTCAGCGA TCTGTCTATT TCGTTCATCC
     1981 ATAGTTGCCT GACTCCCCGT CGTGTAGATA ACTACGATAC GGGAGGGCTT ACCATCTGGC
     2041 CCCAGTGCTG CAATGATACC GCGAGACCCA CGCTCACCGG CTCCAGATTT ATCAGCAATA
     2101 AACCAGCCAG CCGGAAGGGC CGAGCGCAGA AGTGGTCCTG CAACTTTATC CGCCTCCATC
     2161 CAGTCTATTA ATTGTTGCCG GGAAGCTAGA GTAAGTAGTT CGCCAGTTAA TAGTTTGCGC
     2221 AACGTTGTTG CCATTGCTAC AGGCATCGTG GTGTCACGCT CGTCGTTTGG TATGGCTTCA
     2281 TTCAGCTCCG GTTCCCAACG ATCAAGGCGA GTTACATGAT CCCCCATGTT GTGCAAAAAA
     2341 GCGGTTAGCT CCTTCGGTCC TCCGATCGTT GTCAGAAGTA AGTTGGCCGC AGTGTTATCA
     2401 CTCATGGTTA TGGCAGCACT GCATAATTCT CTTACTGTCA TGCCATCCGT AAGATGCTTT
     2461 TCTGTGACTG GTGAGTACTC AACCAAGTCA TTCTGAGAAT AGTGTATGCG GCGACCGAGT
     2521 TGCTCTTGCC CGGCGTCAAT ACGGGATAAT ACCGCGCCAC ATAGCAGAAC TTTAAAAGTG
     2581 CTCATCATTG GAAAACGTTC TTCGGGGCGA AAACTCTCAA GGATCTTACC GCTGTTGAGA
     2641 TCCAGTTCGA TGTAACCCAC TCGTGCACCC AACTGATCTT CAGCATCTTT TACTTTCACC
     2701 AGCGTTTCTG GGTGAGCAAA AACAGGAAGG CAAAATGCCG CAAAAAAGGG AATAAGGGCG
     2761 ACACGGAAAT GTTGAATACT CAtACTCTTC CTTTTTCAAT ATTATTGAAG CATTTATCAG
     2821 GGTTATTGTC TCATGAGCGG ATACATATTT GAATGTATTT AGAAAAATAA ACAAATAGGG
     2881 GTTCCGCGCA CATTTCCCCG AAAAGTGCCA CCTGACGTCT AAGAAACCAT TATTATCATG
     2941 ACATTAACCT ATAAAAATAG GCGTATCACG AGGCCCTTTC GTCTCGCGCG TTTCGGTGAT
     3001 GACGGTGAAA ACCTCTGACA CATGCAGCTC CCGGAGACGG TCACAGCTTG TCTGTAAGCG
     3061 GATGCCGGGA GCAGACAAGC CCGTCAGGGC GCGTCAGCGG GTGTTGGCGG GTGTCGGGGC
     3121 TGGCTTAACT ATGCGGCATC AGAGCAGATT GTACTGAGAG TGCACCATAT GCGGTGTGAA
     3181 ATACCGCACA GATGCGTAAG GAGAAAATAC CGCATCAGGC GCCATTCGCC ATTCAGGCTG
     3241 CGCAACTGTT GGGAAGGGCG ATCGGTGCGG GCCTCTTCGC TATTACGACA GCTGTCTCTT
     3301 ATACACATCT CAACCATCAT CGATGAATTT TCTCGGGTGT TCTCGCATAT TGGCTCGAaa
     3361 ttcTTCGCCA CCTCTGACTT GAGCGTCGAT TTTTGTGATG CTCGTCAGGG GGGCGGAGCC
     3421 TATGGAAAAA CGCCAGCAAC GCGGCCTTTT TACGGTTCCT GGCCTTTTGC TGGCCTTTTG
     3481 CTCACATGTG TGCTGGGCCC AGCCGGCCAG ATCTGAGCTC GCGGCCGCGA TATCGCTAGc
     3541 tcgaggtccg ttACATAACT TACGGTAAAT GGCCCGCCTG GCTGACCGCC CAACGACCCC
     3601 CGCCCATTGA CGTCAATAAT GACGTATGTT CCCATAGTAA CGCCAATAGG GACTTTCCAT
     3661 TGACGTCAAT GGGTGGAGTA TTTACGGTAA ACTGCCCACT TGGCAGTACA TCAAGTGTAT
     3721 CATATGCCAA GTACGCCCCC TATTGACGTC AATGACGGTA AATGGCCCGC CTGGCATTAT
     3781 GCCCAGTACA TGACCTTATG GGACTTTCCT ACTTGGCAGT ACATCTACGT ATTAGTCATC
     3841 GCTATTACCA TGGTGATGCG GTTTTGGCAG TACATCAATG GGCGTGGATA GCGGTTTGAC
```

```
     3901 TCACGGGGAT TTCCAAGTCT CCACCCCATT GACGTCAATG GGAGTTTGTT TTGGCACCAA
     3961 AATCAACGGG ACTTTCCAAA ATGTCGTAAC AACTCCGCCC CATTGACGCA AATGGGCGGT
     4021 AGGCGTGTAC GGTGGGAGGT CTATATAAGC AGAGCTCGTT TAGTGAACCG TCAGATCGCC
     4081 TGGAGACGCC ATCCACGCTG TTTTGACCTC CATAGAAGAC ACCGGGACCG ATCCAGCCTC
     4141 CGCGGCCGGG AAcggtgcat tggaacggac cGTGTTGACA ATTAATCATC GGCATAGTAT
     4201 ATCGGCATAG TATAATACGA CAAGGTGAGG AACTAAACC
//
```

### *Phaeodactylum tricornutum*

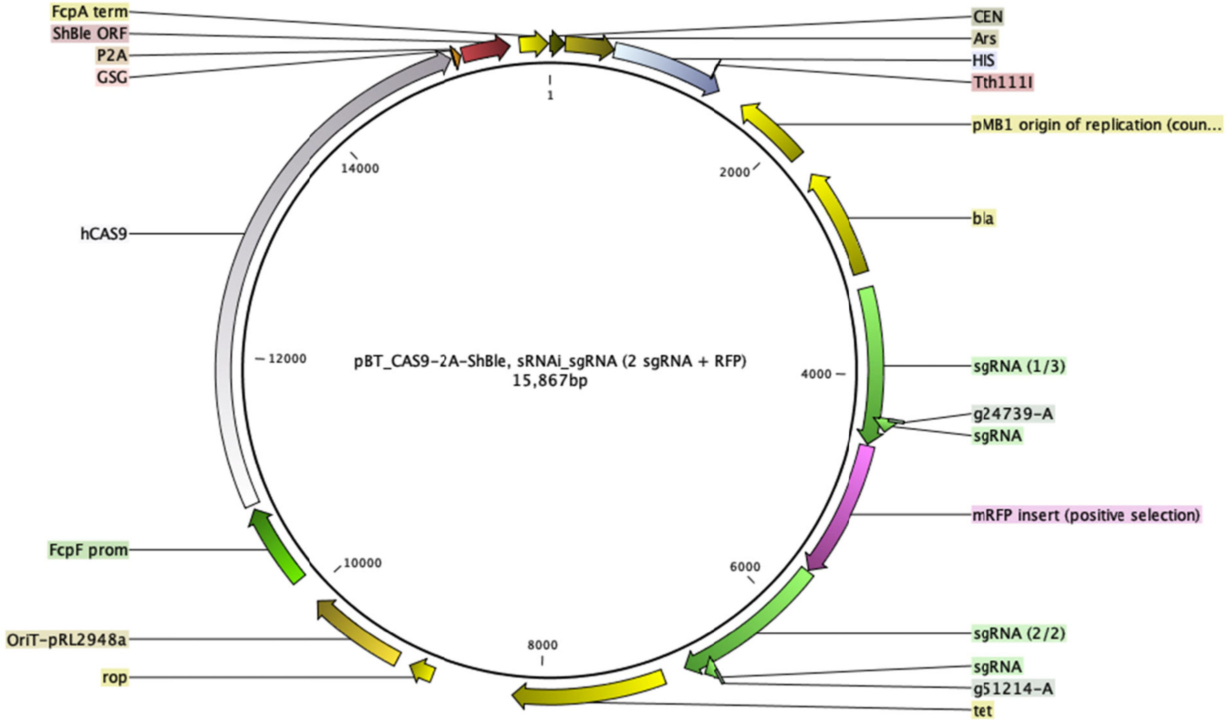


```
>pBT_CAS9-2A-ShBle
ATCACGTGCTATAAAAATAATTATAATTTAAATTTTTTAATATAAATATATAAATTAAAAATAGAAAGTAAAAAAA
GAAATTAAAGAAAAAATAGTTTTTGTTTTCCGAAGATGTAAAAGACTCTAGGGGGATCGCCAACAAATACTACCTT
TTATCTTGCTCTTCCTGCTCTCAGGTATTAATGCCGAATTGTTTCATCTTGTCTGTGTAGAAGACCACACACGAAA
ATCCTGTGATTTTACATTTTACTTATCGTTAATCGAATGTATATCTATTTAATCTGCTTTTCTTGTCTAATAAATA
TATATGTAAAGTACGCTTTTTGTTGAAATTTTTTAAACCTTTGTTTATTTTTTTTTCTTCATTCCGTAACTCTTCT
ACCTTCTTTATTTACTTTCTAAAATCCAAATACAAAACATAAAAATAAATAAACACAGAGTAAATTCCCAAATTAT
TCCATCATTAAAAGATACGAGGCGCGTGTAAGTTACAGGCAAGCGATCCTAGTACACTCTATATTTTTTTATGCCT
CGGTAATGATTTTCATTTTTTTTTTTTCCACCTAGCGGATGACTCTTTTTTTTTTCTTAGCGATTGGCATTATCACAT
AATGAATTATACATTATATAAAGTAATGTGATTTCTTCGAAGAATATACTAAAAAATGAGCAGGCAAGATAAACGA
AGGCAAAGATGACAGAGCAGAAAGCCCTAGTAAAGCGTATTACAAATGAAACCAAGATTCAGATTGCGATCTCTTT
AAAGGGTGGTCCCCTAGCGATAGAGCACTCGATCTTCCCAGAAAAAGAGGCAGAAGCAGTAGCAGAACAGGCCACA
CAATCGCAAGTGATTAACGTCCACACAGGTATAGGGTTTCTGGACCATATGATACATGCTCTGGCCAAGCATTCCG
GCTGGTCGCTAATCGTTGAGTGCATTGGTGACTTACACATAGACGACCATCACACCACTGAAGACTGCGGGATTGC
TCTCGGTCAAGCTTTTAAAGAGGCCCTACTGGCGCGTGGAGTAAAAAGGTTTGGATCAGGATTTGCGCCTTTGGAT
GAGGCACTTTCCAGAGCGGTGGTAGATCTTTCGAACAGGCCGTACGCAGTTGTCGAACTTGGTTTGCAAAGGGAGA
AAGTAGGAGATCTCTCTTGCGAGATGATCCCGCATTTTCTTGAAAGCTTTGCAGAGGCTAGCAGAATTACCCTCCA
CGTTGATTGTCTGCGAGGCAAGAATGATCATCACCGTAGTGAGAGTGCGTTCAAGGCTCTTGCGGTTGCCATAAGA
GAAGCCACCTCGCCCAATGGTACCAACGATGTTCCCTCCACCAAAGGTGTTCTTATGTAGTTTTACACAGGAGTCT
GGACTTGACCTCTAGAGTCGACCTGCAGGCATCTCTTCCGCTTCCTCGCTCACTGACTCGCTGCGCTCGGTCGTTC
GGCTGCGGCGAGCGGTATCAGCTCACTCAAAGGCGGTAATACGGTTATCCACAGAATCAGGGGATAACGCAGGAAA
GAACATGTGAGCAAAAGGCCAGCAAAAGGCCAGGAACCGTAAAAAGGCCGCGTTGCTGGCGTTTTTCCATAGGCTC
CGCCCCCCTGACGAGCATCACAAAAATCGACGCTCAAGTCAGAGGTGGCGAAACCCGACAGGACTATAAAGATACC
AGGCGTTTCCCCCTGGAAGCTCCCTCGTGCGCTCTCCTGTTCCGACCCTGCCGCTTACCGGATACCTGTCCGCCTT
TCTCCCTTCGGGAAGCGTGGCGCTTTCTCATAGCTCACGCTGTAGGTATCTCAGTTCGGTGTAGGTCGTTCGCTCC
AAGCTGGGCTGTGTGCACGAACCCCCCGTTCAGCCCGACCGCTGCGCCTTATCCGGTAACTATCGTCTTGAGTCCA
ACCCGGTAAGACACGACTTATCGCCACTGGCAGCAGCCACTGGTAACAGGATTAGCAGAGCGAGGTATGTAGGCGG
```

TGCTACAGAGTTCTTGAAGTGGTGGCCTAACTACGGCTACACTAGAAGGACAGTATTTGGTATCTGCGCTCTGCTG
AAGCCAGTTACCTTCGGAAAAAGAGTTGGTAGCTCTTGATCCGGCAAACAAACCACCGCTGGTAGCGGTGGTTTTT
TTGTTTGCAAGCAGCAGATTACGCGCAGAAAAAAAGGATCTCAAGAAGATCCTTTGATCTTTTCTACGGGGTCTGA
CGCTCAGTGGAACGAAAACTCACGTTAAGGGATTTTGGTCATGAGATTATCAAAAAGGATCTTCACCTAGATCCTT
TTAAATTAAAAATGAAGTTTTAAATCAATCTAAAGTATATATGAGTAAACTTGGTCTGACAGTTACCAATGCTTAA
TCAGTGAGGCACCTATCTCAGCGATCTGTCTATTTCGTTCATCCATAGTTGCCTGACTCCCCGTCGTGTAGATAAC
TACGATACGGGAGGGCTTACCATCTGGCCCCAGTGCTGCAATGATACCGCGgGAtCCACGCTCACCGGCTCCAGAT
TTATCAGCAATAAACCAGCCAGCCGGAAGGGCCGAGCGCAGAAGTGGTCCTGCAACTTTATCCGCCTCCATCCAGT
CTATTAATTGTTGCCGGGAAGCTAGAGTAAGTAGTTCGCCAGTTAATAGTTTGCGCAACGTTGTTGCCATTGCTGC
AGGCATCGTGGTGTCACGCTCGTCGTTTGGTATGGCTTCATTCAGCTCCGGTTCCCAACGATCAAGGCGAGTTACA
TGATCCCCCATGTTGTGCAAAAAAGCGGTTAGCTCCTTCGGTCCTCCGATCGTTGTCAGAAGTAAGTTGGCCGCAG
TGTTATCACTCATGGTTATGGCAGCACTGCATAATTCTCTTACTGTCATGCCATCCGTAAGATGCTTTTCTGTGAC
TGGTGAGTACTCAACCAAGTCATTCTGAGAATAGTGTATGCGGCGACCGAGTTGCTCTTGCCCGGCGTCAACACGG
GATAATACCGCGCCACATAGCAGAACTTTAAAAGTGCTCATCATTGGAAAACGTTCTTCGGGGCGAAAACTCTCAA
GGATCTTACCGCTGTTGAGATCCAGTTCGATGTAACCCACTCGTGCACCCAACTGATCTTCAGCATCTTTTACTTT
CACCAGCGTTTCTGGGTGAGCAAAAACAGGAAGGCAAAATGCCGCAAAAAAGGGAATAAGGGCGACACGGAAATGT
TGAATACTCATACTCTTCCTTTTTCAATATTATTGAAGCATTTATCAGGGTTATTGTCTCATGAGCGGATACATAT
TTGAATGTATTTAGAAAAATAAACAAATAGGGGTTCCGCGCACATTTCCCCGAATCGACTTCTGCAGCTCTTCCAA
ATCGTACACGCCACTGGGCATTGAAGTGGACTCGCCTGCAGCATGAAAATTATCATAATAGCTGCAGGTTTCGATC
GAGCCCGGACGCTTCTGTGCTTCTTCCAAAACCCAGCTCGCCGTTAGACTTCGGCAAGCCGCATCGACGGCTTCCC
GATCGCTTTCGGCCATGACTCGCTCGTGAACACACATATTCCGTCGACTGCTTAAACACAATGCTAGAGCCCCGCT
TCCACCAGCTCCGCGACCGTCCGAGATCGGTCCCATGGGAGGTTTTTGTAAATTGGCGGTATAGACCTTTCGAGGC
TTCTTGCGATTGGGCGAGCCATTGCTGGTAATGTGGGTGATGGCAGTGACGTTTACATCCCCGCTACTGCCCGCAA
CCGGAACGTTTTCAATATCCCCGTCAGTGTCCATTGGCAAGATGTTTTCTTCCTGGTGCCGTTGTAGTTCTTGCGA
CCGGTACGCTAAAACTGTCGCGAGTTCTTCCATAACGTGATTCATTTCCGGAACGGTTCGAGTGCAATACACGAGC
TTACCCGCCGACGGATTCGCAAACTGGTAGGAAGTAATCAGCGATAGCAAACATACGGTTTTTCCCGTTCCCGTAG
GCATCTCCAGGAGACAGTGCCCACCTGCGTCCAGCGACTGTTTCAGTGCTCGCATGTACTGATGCTGCTCAAGGTA
GATGCGATCGTACGGAAAGAACACATCCAGGCCGTCCAAATCGAATCTCATTTTGACGCCTTTTTCACACCTTCCG
CTTATGCAAAGCTTTCAGGAATTGCCTATGAAGAGAACGGTCAATTCCTCATTGATGACTGACAAGACTTGTTTCT
GTTTACGGTAATGAAGTCTCTCGCAATCCTTTTGCAATAGATCGATCGATCGACAGAATGGAAGCAACTTTTGATT
ACTTACTGTTACAGTTATTGTGTATAGGTATACTCTTTCTATTTATATATTTGCAGAAAATCATAGTTTTGTCAAA
CTTGTTTCCTCCATGTTTTAGAGCTAGAAATAGCAAGTTAAAATAAGGCTAGTCCGTTATCAACTTGAAAAAGTGG
CACCGAGTCGGTGCTTTTTTTTTTTGCCTTTTGACCGAAAGCTATCTTACTTACTACCTACCCAGCCACCGACAGC
CTGTTTTCTATAGTACTGGAGCAAGACCAGAGTATAGAATTTGCTGCTGTTATAGCCGTTgaattcctgtCTGTAA
TACGCAAACCGCCTCTCCCCGCGCGTTGGCCGATTCATTAATGCAGCTGGCACGACAGGTTTCCCGACTGGAAAGC
GGGCAGTGAGCGCAACGCAATTAATGTGAGTTAGCTCACTCATTAGGCACCCCAGGCTTTACACTTTATGCTTCCG
GCTCGTATGTTGTGTGGAATTGTGAGCGGATAACAATTTCACACATACTAGAGAAAGAGGAGAAATACTAGATGGC
TTCCTCCGAAGACGTTATCAAAGAGTTCATGCGTTTCAAAGTTCGTATGGAAGGTTCCGTTAACGGTCACGAGTTC
GAAATCGAAGGTGAAGGTGAAGGTCGTCCGTACGAAGGTACCCAGACCGCTAAACTGAAAGTTACCAAAGGTGGTC
CGCTGCCGTTCGCTTGGGACATCCTGTCCCCGCAGTTCCAGTACGGTTCCAAAGCTTACGTTAAACACCCGGCTGA
CATCCCGGACTACCTGAAACTGTCCTTCCCGGAAGGTTTCAAATGGGAACGTGTTATGAACTTCGAAGACGGTGGT
GTTGTTACCGTTACCCAGGACTCCTCCCTGCAAGACGGTGAGTTCATCTACAAAGTTAAACTGCGTGGTACCAACT
TCCCGTCCGACGGTCCGGTTATGCAGAAAAAAACCATGGGTTGGGAAGCTTCCACCGAACGTATGTACCCGGAAGA
CGGTGCTCTGAAAGGTGAAATCAAAATGCGTCTGAAACTGAAAGACGGTGGTCACTACGACGCTGAAGTTAAAACC
ACCTACATGGCTAAAAAACCGGTTCAGCTGCCGGGTGCTTACAAAACCGACATCAAACTGGACATCACCTCCCACA
ACGAAGACTACACCATCGTTGAACAGTACGAACGTGCTGAAGGTCGTCACTCCACCGGTGCTTAATAACGCTGATA
GTGCTAGTGTAGATCGCTACTAGAGCCAGGCATCAAATAAAACGAAAGGCTCAGTCGAAAGACTGGGCCTTTCGTT
TTATCTGTTGTTTGTCGGTGAACGCTCTCTACTAGAGTCACACTGGCTCACCTTCGGGTGGGCCTTTCTGCGTTTA
TACGTTATCGATTGAGTCGACTTCTGCAGCTCTTCCAAATCGTACACGCCACTGGGCATTGAAGTGGACTCGCCTG
CAGCATGAAAATTATCATAATAGCTGCAGGTTTCGATCGAGCCCGGACGCTTCTGTGCTTCTTCCAAAACCCAGCT
CGCCGTTAGACTTCGGCAAGCCGCATCGACGGCTTCCCGATCGCTTTCGGCCATGACTCGCTCGTGAACACACATA
TTCCGTCGACTGCTTAAACACAATGCTAGAGCCCCGCTTCCACCAGCTCCGCGACCGTCCGAGATCGGTCCCATGG
GAGGTTTTTGTAAATTGGCGGTATAGACCTTTCGAGGCTTCTTGCGATTGGGCGAGCCATTGCTGGTAATGTGGGT
GATGGCAGTGACGTTTACATCCCCGCTACTGCCCGCAACCGGAACGTTTTCAATATCCCCGTCAGTGTCCATTGGC
AAGATGTTTTCTTCCTGGTGCCGTTGTAGTTCTTGCGACCGGTACGCTAAAACTGTCGCGAGTTCTTCCATAACGT
GATTCATTTCCGGAACGGTTCGAGTGCAATACACGAGCTTACCCGCCGACGGATTCGCAAACTGGTAGGAAGTAAT
CAGCGATAGCAAACATACGGTTTTTCCCGTTCCCGTAGGCATCTCCAGGAGACAGTGCCCACCTGCGTCCAGCGAC
TGTTTCAGTGCTCGCATGTACTGATGCTGCTCAAGGTAGATGCGATCGTACGGAAAGAACACATCCAGGCCGTCCA
AATCGAATCTCATTTTGACGCCTTTTTCACACCTTCCGCTTATGCAAAGCTTTCAGGAATTGCCTATGAAGAGAAC
GGTCAATTCCTCATTGATGACTGACAAGACTTGTTTCTGTTTACGGTAATGAAGTCTCTCGCAATCCTTTTGCAAT
AGATCGATCGATCGACAGAATGGAAGCAACTTTTGATTACTTACTGTTACAGTTATTGTGTATAGGTATACTCTTT
CTATTTATATATTTGCAGAAAATCATAGTTTTGTTGGCTCCACTTCTTCAATGTTTTAGAGCTAGAAATAGCAAGT
TAAAATAAGGCTAGTCCGTTATCAACTTGAAAAAGTGGCACCGAGTCGGTGCTTTTTTTTTTTGCCTTTTGACCGA
AAGCTATCTTACTTACTACCTACCCAGCCACCGACAGCCTGTTTTCTATAGTACTGGAGCAAGACCAGAGTATAGA
ATTTGCTGCTGTTATAGCCGTTAAGTGCCACCTGACGTCTAAGAAACCATTATTATCATGACATTAACCTATAAAA
ATAGGCGTATCACGAGGCCCTTTCGTCTTCAAGAATTCTCATGTTTGACAGCTTATCATCGATAAGCTTTAATGCG
GTAGTTTATCACAGTTAAATTGCTAACGCAGTCAGGCACCGTGTATGAAATCTAACAATGCGCTCATCGTCATCCT
CGGCACCGTCACCCTGGATGCTGTAGGCATAGGCTTGGTTATGCCGGTACTGCCGGGCCTCTTGCGGATATCGTC

CATTCCGACAGCATCGCCAGTCACTATGGCGTGCTGCTAGCGCTATATGCGTTGATGCAATTTCTATGCGCACCCG
TTCTCGGAGCACTGTCCGACCGCTTTGGCCGCCGCCCAGTCCTGCTCGCTTCGCTACTTGGAGCCACTATCGACTA
CGCGATCATGGCGACCACACCCGTCCTGTGGATCCTCTACGCCGGACGCATCGTGGCCGGCATCACCGGCGCCACA
GGTGCGGTTGCTGGCGCCTATATCGCCGACATCACCGATGGGGAAGATCGGGCTCGCCACTTCGGGCTCATGAGCG
CTTGTTTCGGCGTGGGTATGGTGGCAGGCCCCGTGGCCGGGGGACTGTTGGGCGCCATCTCCTTGCATGCACCATT
CCTTGCGGCGGCGGTGCTCAACGGCCTCAACCTACTACTGGGCTGCTTCCTAATGCAGGAGTCGCATAAGGGAGAG
CGTCGACCGATGCCCTTGAGAGCCTTCAACCCAGTCAGCTCCTTCCGGTGGGCGCGGGGCATGACTATCGTCGCCG
CACTTATGACTGTCTTCTTTATCATGCAACTCGTAGGACAGGTGCCGGCAGCGCTCTGGGTCATTTTCGGCGAGGA
CCGCTTTCGCTGGAGCGCGACGATGATCGGCCTGTCGCTTGCGGTATTCGGAATCTTGCACGCCCTCGCTCAAGCC
TTCGTCACTGGTCCCGCCACCAAACGTTTCGGCGAGAAGCAGGCCATTATCGCCGGCATGGCGGCCGACGCGCTGG
GCTACGTCTTGCTGGCGTTCGCGACGCGAGGCTGGATGGCCTTCCCCATTATGATTCTTCTCGCTTCCGGCGGCAT
CGGGATGCCCGCGTTGCAGGCCATGCTGTCCAGGCAGGTAGATGACGACCATCAGGGACAGCTTCAAGGATCGCTC
GCGGCTCTTACCAGCCTAACTTCGATCATTGGACCGCTGATCGTCACGGCGATTTATGCCGCCTCGGCGAGCACAT
GGAACGGGTTGGCATGGATTGTAGGCGCCGCCCTATACCTTGTCTGCCTCCCCGCGTTGCGTCGCGGTGCATGGAG
CCGGGCCACCTCGACCTGAATGGAAGCCGGCGGCACCTCGCTAACGGATTCACCACTCCAAGAATTGGAGCCAATC
AATTCTTGCGGAGAACTGTGAATGCGCAAACCAACCCTTGGCAGAACATATCCATCGCGTCCGCCATCTCCAGCAG
CCGCACGCGGCGCATCTCGGGCAGCGTTGGGTCCTGGCCACGGGTGCGCATGATCGTGCTCCTGTCGTTGAGGACC
CGGCTAGGCTGGCGGGGTTGCCTTACTGGTTAGCAGAATGAATCACCGATACGCGAGCGAACGTGAAGCGACTGCT
GCTGCAAAACGTCTGCGACCTGAGCAACAACATGAATGGTCTTCGGTTTCCGTGTTTCGTAAAGTCTGGAAACGCG
GAAGTCAGCGCCCTGCACCATTATGTTCCGGATCTGCATCGCAGGATGCTGCTGGCTACCCTGTGGAACACCTACA
TCTGTATTAACGAAGCGCTGGCATTGACCCTGAGTGATTTTTCTCTGGTCCCGCCGCATCCATACCGCCAGTTGTT
TACCCTCACAACGTTCCAGTAACCGGGCATGTTCATCATCAGTAACCCGTATCGTGAGCATCCTCTCTCGTTTCAT
CGGTATCATTACCCCCATGAACAGAAATCCCCCTTACACGGAGGCATCAGTGACCAAACAGGAAAAAACCGCCCTT
AACATGGCCCGCTTTATCAGAAGCCAGACATTAACGCTTCTGGAGAAACTCAACGAGCTGGACGCGGATGAACAGG
CAGACATCTGTGAATCGCTTCACGACCACGCTGATGAGCTTTACCGCAGCTGCCTCGCGCGTTTCGGTGATGACGG
TGAAAACCTCTGACACATGCAGCTCCCGGAGACGGTCACAGCTTGTCTGTAAGCGGATGCCGGGAGCAGACAAGCC
CGTCAGGGCGCGTCAGCGGGTGTTGGCGGGTGTCGGGGCGGATCGTCTTGCCTTGCTCGTCGGTGATGTACTTACA
GCTCGAAGTGCCTCTTCTTGATGGAGGCCATGGAACGTGCTTCCGAATCACGCGCACCCCGATCCGTTACGTGAAA
AAAGTCATGGCTCTGCCCTCGGGCGGACCACGCCCATCATGACCTTGCCAAGCTCGTCCTGCTTCTCTTCGATCTT
CGCCAGCAGGGCGAGGATCGTGGCATCACCGAACCGCGCCGTGCGCGGGTCGTCGGTGAGCCAGAGTTTCAGCAGG
CCGCCCAGGCGGCCCAGGTCGCCATTGATGCGGGCCAGCTCGCGGACGTGCTCATAGTCCACGACGCCCGTGATTT
TGTAGCCCTGGCCGACGGCCAGCAGGTAGGCCGACAGGCTCATGCCGGCCGCCGCCGCCTTTTCCTCAATCGCTCT
TCGTTCGTCTGGAAGGCAGTACACCTTGATAGGTGGGCTGCCCTTCCTGGTTGGCTTGGTTTCATCAGCCATCCGC
TTGCCCTCATCTGTTACGCCGGCGGTAGCCGGCCAGCCTCGCAGAGCAGGATTCCCGTTGAGCACCGCCAGGTGCG
AATAAGGGACAGTGAAGAAGGAACACCCGCTCGCGGGTGGGCCTACTTCACCTATCCTGCCCGGCTGACGCCGTTG
GATACACCAAGGAAAGTCTACGCGAACCCTTTGGCAAAATCCTGTATATCGTGCGAAAAAGGATGGATATACCGAA
AAAATCGCTATAATGACCCCGAAGCAGGGTTATGCAGCGGAAGATGCCATTCGCCATTCAGGCTGCGCAACTGTTG
GGAAGGGCGATCGGTGCGGGCCTCTTCGCTATTACGCCAGCTGGCGAAAGGGGGATGTGCTGCAAGGCGATTAAGT
TGGGTAACGCCAGGGTTTTCCCAGTCACGACGTTGTAAAACGACGGCCAGTGACCATGATTACGCCAAGCTCGAAA
TTAACCCTCACTAAAGGGAACAAAAGCTGGTACCTAACAGGATTAGTGCAATTCGAGTTGAATCACTGGGAAAAAC
ATTGTCTTCTTTTTTATATTATCATTTGCATTAGTGCTGCAGTCGTAGATACTTGTTGGTTGAAAGACATCAGCTG
GGAGGGACTGGACTAGCGTTTGGTAAGGAGACATACCTGTTAACGTTGGTTGCAAAATTCCATTTCGCGATTTATG
TTATCTGTAAATCCTGATTTGTCTGGAATTCTTGATACTTCCGTTTTTTTAGAGGCCAATGATTAGCATCGGCGAT
TCTCAAAATAGCATTTTCGACATGCGGTGCTGATTTCATAAACATAGACAACGCTTTTACATGTAAAAGTAACTTG
CGGACTTGGAACAGTGCTCTGTTTTTGGTGTGAACGTAACTCAGCAATATTTCTGTGCTAGCAAGGTTTTTTATGA
TCGACCGAAGATCTCAAAACTCCGGGTCTTTCAACTGTCTGACTAGACCATGTTCGTAACGTCGGACAGCAGCTTT
CGTTGTACTGGTAGAATTTCTACGTGCGAAGCACGTGTAGGCAGGTTGAACGACGATCCCTGCCGATGGATGGATT
GGCACGCGGCGGAACGCTTTCGTGATCTACACCACCTGGATCTTCACATATCTTCGAAATCGAAAAATTAACCAAG
TCGACGGTATCGATAATATTCTAGCTGAGGGTACCCATGGACTATAAGGACCACGACGGAGACTACAAGGATCATG
ATATTGATTACAAAGACGATGACGATAAGATGGCCCCAAAGAAGAAGCGGAAGGTCGGTATCCACGGAGTCCCAGC
AGCCGACAAGAAGTACAGCATCGGCCTGGACATCGGCACCAACTCTGTGGGCTGGGCCGTGATCACCGACGAGTAC
AAGGTGCCCAGCAAGAAATTCAAGGTGCTGGGCAACACCGACCGGCACAGCATCAAGAAGAACCTGATCGGAGCCC
TGCTGTTCGACAGCGGCGAAACAGCCGAGGCCACCCGGCTGAAGAGAACCGCCAGAAGAAGATACACCAGACGGAA
GAACCGGATCTGCTATCTGCAAGAGATCTTCAGCAACGAGATGGCCAAGGTGGACGACAGCTTCTTCCACAGACTG
GAAGAGTCCTTCCTGGTGGAAGAGGATAAGAAGCACGAGCGGCACCCCATCTTCGGCAACATCGTGGACGAGGTGG
CCTACCACGAGAAGTACCCCACCATCTACCACCTGAGAAAGAAACTGGTGGACAGCACCGACAAGGCCGACCTGCG
GCTGATCTATCTGGCCCTGGCCCACATGATCAAGTTCCGGGGCCACTTCCTGATCGAGGGCGACCTGAACCCCGAC
AACAGCGACGTGGACAAGCTGTTCATCCAGCTGGTGCAGACCTACAACCAGCTGTTCGAGGAAAACCCCATCAACG
CCAGCGGCGTGGACGCCAAGGCCATCCTGTCTGCCAGACTGAGCAAGAGCAGACGGCTGGAAAATCTGATCGCCCA
GCTGCCCGGCGAGAAGAAGAATGGCCTGTTCGGAAACCTGATTGCCCTGAGCCTGGGCCTGACCCCCAACTTCAAG
AGCAACTTCGACCTGGCCGAGGATGCCAAACTGCAGCTGAGCAAGGACACCTACGACGACGACCTGGACAACCTGC
TGGCCCAGATCGGCGACCAGTACGCCGACCTGTTTCTGGCCGCCAAGAACCTGTCCGACGCCATCCTGCTGAGCGA
CATCCTGAGAGTGAACACCGAGATCACCAAGGCCCCCCTGAGCGCCTCTATGATCAAGAGATACGACGAGCACCAC
CAGGACCTGACCCTGCTGAAAGCTCTCGTGCGGCAGCAGCTGCCTGAGAAGTACAAAGAGATTTTCTTCGACCAGA
GCAAGAACGGCTACGCCGGCTACATTGACGGCGGAGCCAGCCAGGAAGAGTTCTACAAGTTCATCAAGCCCATCCT
GGAAAAGATGGACGGCACCGAGGAACTGCTCGTGAAGCTGAACAGAGAGGACCTGCTGCGGAAGCAGCGGACCTTC
GACAACGGCAGCATCCCCCACCAGATCCACCTGGGAGAGCTGCACGCCATTCTGCGGCGGCAGGAAGATTTTTACC
CATTCCTGAAGGACAACCGGGAAAAGATCGAGAAGATCCTGACCTTCCGCATCCCCTACTACGTGGGCCCTCTGGC

CAGGGGAAACAGCAGATTCGCCTGGATGACCAGAAAGAGCGAGGAAACCATCACCCCCTGGAACTTCGAGGAAGTG
GTGGACAAGGGCGCTTCCGCCCAGAGCTTCATCGAGCGGATGACCAACTTCGATAAGAACCTGCCCAACGAGAAGG
TGCTGCCCAAGCACAGCCTGCTGTACGAGTACTTCACCGTGTATAACGAGCTGACCAAAGTGAAATACGTGACCGA
GGGAATGAGAAAGCCCGCCTTCCTGAGCGGCGAGCAGAAAAAGGCCATCGTGGACCTGCTGTTCAAGACCAACCGG
AAAGTGACCGTGAAGCAGCTGAAAGAGGACTACTTCAAGAAAATCGAGTGCTTCGACTCCGTGGAAATCTCCGGCG
TGGAAGATCGGTTCAACGCCTCCCTGGGCACATACCACGATCTGCTGAAAATTATCAAGGACAAGGACTTCCTGGA
CAATGAGGAAAACGAGGACATTCTGGAAGATATCGTGCTGACCCTGACACTGTTTGAGGACAGAGAGATGATCGAG
GAACGGCTGAAAACCTATGCCCACCTGTTCGACGACAAAGTGATGAAGCAGCTGAAGCGGCGGAGATACACCGGCT
GGGGCAGGCTGAGCCGGAAGCTGATCAACGGCATCCGGGACAAGCAGTCCGGCAAGACAATCCTGGATTTCCTGAA
GTCCGACGGCTTCGCCAACAGAAACTTCATGCAGCTGATCCACGACGACAGCCTGACCTTTAAAGAGGACATCCAG
AAAGCCCAGGTGTCCGGCCAGGGCGATAGCCTGCACGAGCACATTGCCAATCTGGCCGGCAGCCCCGCCATTAAGA
AGGGCATCCTGCAGACAGTGAAGGTGGTGGACGAGCTCGTGAAAGTGATGGGCCGGCACAAGCCCGAGAACATCGT
GATCGAAATGGCCAGAGAGAACCAGACCACCCAGAAGGGACAGAAGAACAGCCGCGAGAGAATGAAGCGGATCGAA
GAGGGCATCAAAGAGCTGGGCAGCCAGATCCTGAAAGAACACCCCGTGGAAAACACCCAGCTGCAGAACGAGAAGC
TGTACCTGTACTACCTGCAGAATGGGCGGGATATGTACGTGGACCAGGAACTGGACATCAACCGGCTGTCCGACTA
CGATGTGGACCATATCGTGCCTCAGAGCTTTCTGAAGGACGACTCCATCGACAACAAGGTGCTGACCAGAAGCGAC
AAGAACCGGGGCAAGAGCGACAACGTGCCCTCCGAAGAGGTCGTGAAGAAGATGAAGAACTACTGGCGGCAGCTGC
TGAACGCCAAGCTGATTACCCAGAGAAAGTTCGACAATCTGACCAAGGCCGAGAGAGGCGGCCTGAGCGAACTGGA
TAAGGCCGGCTTCATCAAGAGACAGCTGGTGGAAACCCGGCAGATCACAAAGCACGTGGCACAGATCCTGGACTCC
CGGATGAACACTAAGTACGACGAGAATGACAAGCTGATCCGGGAAGTGAAAGTGATCACCCTGAAGTCCAAGCTGG
TGTCCGATTTCCGGAAGGATTTCCAGTTTTACAAAGTGCGCGAGATCAACAACTACCACCACGCCCACGACGCCTA
CCTGAACGCCGTCGTGGGAACCGCCCTGATCAAAAAGTACCCTAAGCTGGAAAGCGAGTTCGTGTACGGCGACTAC
AAGGTGTACGACGTGCGGAAGATGATCGCCAAGAGCGAGCAGGAAATCGGCAAGGCTACCGCCAAGTACTTCTTCT
ACAGCAACATCATGAACTTTTTCAAGACCGAGATTACCCTGGCCAACGGCGAGATCCGGAAGCGGCCTCTGATCGA
GACAAACGGCGAAACCGGGGAGATCGTGTGGGATAAGGGCCGGGATTTTGCCACCGTGCGGAAAGTGCTGAGCATG
CCCCAAGTGAATATCGTGAAAAAGACCGAGGTGCAGACAGGCGGCTTCAGCAAAGAGTCTATCCTGCCCAAGAGGA
ACAGCGATAAGCTGATCGCCAGAAAGAAGGACTGGGACCCTAAGAAGTACGGCGGCTTCGACAGCCCCACCGTGGC
CTATTCTGTGCTGGTGGTGGCCAAAGTGGAAAAGGGCAAGTCCAAGAAACTGAAGAGTGTGAAAGAGCTGCTGGGG
ATCACCATCATGGAAAGAAGCAGCTTCGAGAAGAATCCCATCGACTTTCTGGAAGCCAAGGGCTACAAAGAAGTGA
AAAAGGACCTGATCATCAAGCTGCCTAAGTACTCCCTGTTCGAGCTGGAAAACGGCCGGAAGAGAATGCTGGCCTC
TGCCGGCGAACTGCAGAAGGGAAACGAACTGGCCCTGCCCTCCAAATATGTGAACTTCCTGTACCTGGCCAGCCAC
TATGAGAAGCTGAAGGGCTCCCCCGAGGATAATGAGCAGAAACAGCTGTTTGTGGAACAGCACAAGCACTACCTGG
ACGAGATCATCGAGCAGATCAGCGAGTTCTCCAAGAGAGTGATCCTGGCCGACGCTAATCTGGACAAAGTGCTGTC
CGCCTACAACAAGCACCGGGATAAGCCCATCAGAGAGCAGGCCGAGAATATCATCCACCTGTTTACCCTGACCAAT
CTGGGAGCCCCTGCCGCCTTCAAGTACTTTGACACCACCATCGACCGGAAGAGGTACACCAGCACCAAAGAGGTGC
TGGACGCCACCCTGATCCACCAGAGCATCACCGGCCTGTACGAGACACGGATCGACCTGTCTCAGCTGGGAGGCGA
CAAAAGGCCGGCGGCCACGAAAAAGGCCGGCCAGGCAAAAAAGAAAAAGgcgcgcgagacgggaagcggagctact
aacttcagcctgctgaagcaggctggcgacgtggaggagaaccctggacctggactccggcgcATGGCCAAGTTGA
CCAGTGCCGTTCCGGTGCTCACCGCGCGCGACGTCGCCGGAGCGGTCGAGTTCTGGACCGACCGGCTCGGGTTCTC
CCGGGACTTCGTGGAGGACGACTTCGCCGGTGTGGTCCGGGACGACGTGACCCTGTTCATCAGCGCGGTCCAGGAC
CAGGTGGTGCCGGACAACACCCTGGCCTGGGTGTGGGTGCGCGGCCTGGACGAGCTGTACGCCGAGTGGTCGGAGG
TCGTGTCCACGAACTTCCGGGACGCCTCCGGGCCGGCCATGACCGAGATCGGCGAGCAGCCGTGGGGGCGGGAGTT
CGCCCTGCGCGACCCGGCCGGCAACTGCGTGCACTTCGTGGCCGAGGAGCAGGACTGACCGACGCCGACCAACACC
GCCGGTCCGACGCGGCCCGACGGGTCCGAGGCCTCGGAGATCTGGGCCCATGCGGCCGCAACAACTACCTCGACTT
TGGCTGGGACACTTTCAGTGAGGACAAGAAGCTTCAGAAGCGTGCTATCGAACTCAACCAGGGACGTGCGGCACAA
ATGGGCATCCTTGCTCTCATGGTGCACGAACAGTTGGGAGTCTCTATCCTTCCTTAAAAATTTAATTTTCATTAGT
TGCAGTCACTCCGCTTTGGTTTCACAGTCAGGAATAACACTAGCTCGTCTTCAGCGAGC

#### ***Alveolates***

##### *Euplotes crassus and focardii*

*Euplotes crassus* GFP artificial nanochromosome sequence

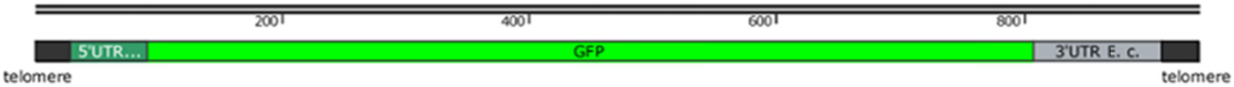


```
CCCCAAAACCCCAAAACCCCAAAACCCCTGAAGGAAGGCCGTCAAGGCCGCATATTAACTAGGTGAAAT
GTTGGTTATAAATATTTAAAAATGTCTGGAGGAGAAGAACTTTTCGCTGGAATTGTTCCAGTTCTTATT
GAACTTGATGGAGATGTTCATGGACATAAGTTCTCTGTTAGAGGAGAAGGAGAAGGAGATGCTGATTAT
GGAAAGCTTGAAATTAAGTTCATTTGTACTACTGGAAAGCTTCCAGTTCCATGGCCAACTCTTGTTACT
ACTCTTTGTTATGGAATTCAATGTTTCGCTAGATATCCAGAACATATGAAGATGAATGATTTCTTCAAG
TCTGCTATGCCAGAAGGATATATTCAAGAAAGAACTATTCAATTCCAAGATGATGGAAAGTATAAGACT
AGAGGAGAAGTTAAGTTCGAAGGAGATACTCTTGTTAATAGAATTGAACTTAAGGGAAAGGATTTCAAG
GAAGATGGAAATATTCTTGGACATAAGCTTGAATATTCTTTCAATTCTCATAATGTTTATATTAGACCA
GATAAGGCTAATAATGGACTTGAAGCTAATTTCAAGACTAGACATAATATTGAAGGAGGAGGAGTTCAA
CTTGCTGATCATTATCAAACTAATGTTCCACTTGGAGATGGACCAGTTCTTATTCCAATTAATCATTAT
CTTTCTACTCAAACTAAGATTTCTAAGGATAGAAATGAAGCTAGAGATCATATGGTTCTTCTTGAATCT
TTCTCTGCTTGTTGTCATACTCATGGAATGGATGAACTTTATAGATAAAGGAACTCAAATTTTTGCTAA
AGATTTGTGGGAGAATCTTAGTCAAATTGGCAGAATTATTCAACTGTGTATTCTAATAAACTGGGCCTC
ATGGGCCTTCCTTTCGGGGTTTTGGGGTTTTGGGGTTTTGGGG
```

*Chromera velia*

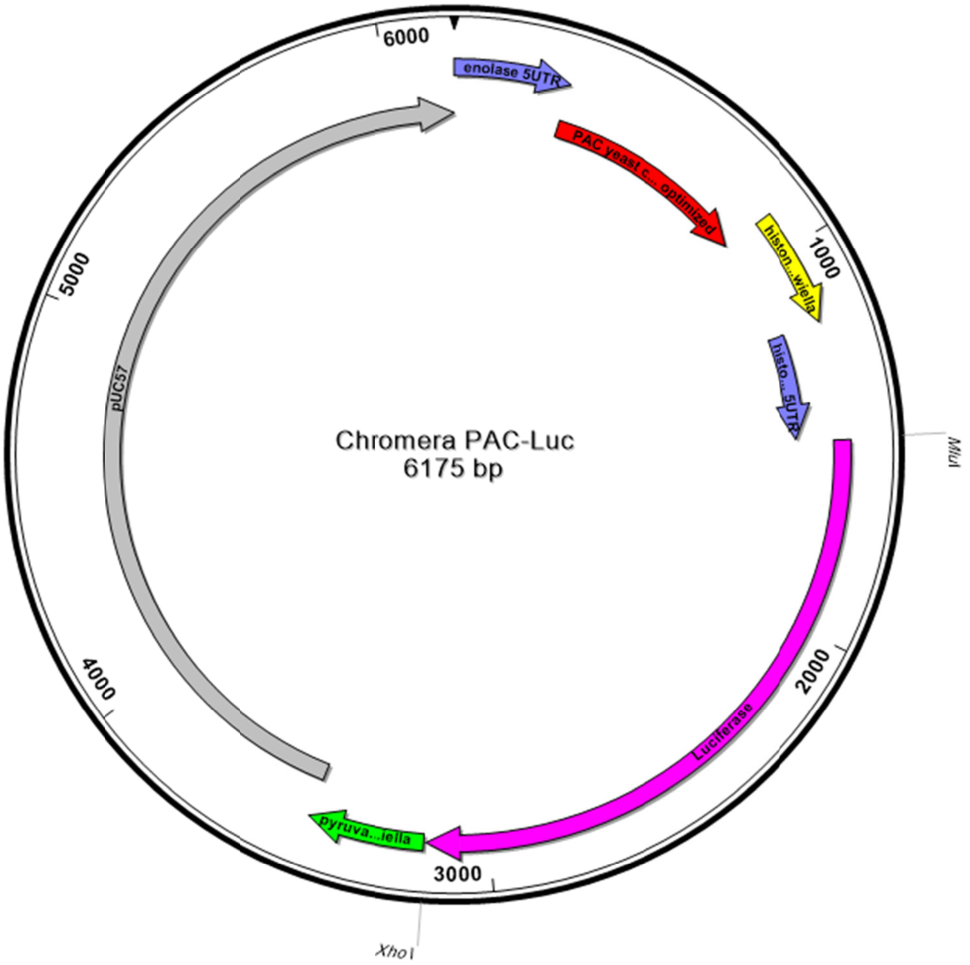


```
>Chromera PAC-Luc
CGGTCGACCGAAAAAGCGATACACAAGCGAAATTTAGACGGGAAGCACTTCTTCCGGCCTGTTTCAAAACTGCGCCGGTCTCTGAC
AAGTTTCGCAAGGCCTTCTGTAGATCCAAACCCCGTCAGCCAAAACAAGTGTGCTGTGCTGTGGAACGAACACAAAATTCTCCAGC
TTTCTTTGCGTAAGTGTCGCAGCGTGGCGGCGAGAGTTTCCACTTCCTTTCTATTCGTCTCAAACGGTGCTTTTAGCTCCACTTCT
TCCCTCTCCCGTCGCCGTTTTCTTTTGAATCCTCTCTCCGTCATGACCGAACATAAGCCATCTGTTAGATTGGCTACTAGAGATGA
TGTTCCAAGAGCTGTTAGAACTTTGGCTGCTGCTTTTGCTGATTATCCAGCTACTAGACATACTGTTGATCCAGATAGACACATCG
AAAGAGTTACCGAATTGCAAGAATTATTCTTAACCAGAGTCGGTTTGGATATCGGTAAAGTTTGGGTTGCTGATGATGGTGCTGCT
GTTGCAGTTTGGACTACTCCAGAATCTTTGGATGCTGGTGCTGTTTTCGCTGAAATTGGTCCAAGAATGGCTGAATTGTCTGGTTC
TAGATTAGCTGCTCAACAACAAATGGAAGGTTTGTTGGCTCCACATAGACCAAAAGAACCAGCTTGGTTTTTGGCTACTGTTGGTG
TTTCTCCAGATCATCAAGGTAAAGGTTTAGGTTCTGCTGTTGTTTTGCCAGGTGTTGAAGCTGCTGAAAGAGCTGGTGTTCCAGCT
TTTTTGGAAACTTCTGCTCCAAGAAATTTGCCATTCTACGAAAGATTGGGTTTCACTGTTACTGCCGATGTTGAAGTTCCTGAAGG
TCCTAGAACTTGGTGTATGACTAGAAAACCAGGTGCTTAAATTGCTCACAGCATTCTCAATGCTACTACATCCGGTGTTCTTAAAC
ACCACCATTCTCTTCGGAAACTGTGCCTTAGTCAAAAATTGTAAACCGATAGAAAATAAATAGAAAATGATGTTATCTAGTGCATA
AGTAGCGAGTTGGAATACGTTCCTACGTCTCCTTGTCAACACCACGATGTAATTCAAGTGATTAGCCATGCTTGATGGCAGATGAA
CAAAGGGAATTCATATTGTACATGATTAAGCGTAGTGGAGCCAGTGAAGAACTCCCACACCCGCCCATCCTAGCTAGTGAACcttg
gacaacgtggaattctacgggcgtctcttacaaggttacgtacaggggaccggaaggggagagcgaaggggaaaaaaatcaggggga
gaaaactgtgctgagggaccctcaactttgcacggtcagaccaaacgaaaaaatgattcgacgttggaacgtctcgtcatttcctt
ccgagatcccaacgtcgaatcatttttcgttttgatttttcagatacagaaaaaagaaaaatgaaatatttcatttttttaaaaaaa
gaaattaattttccgatacagagaaaattatatataggACGCGTATGGAAGACGCCAAAAACATAAAGAAAGGCCCGGCGCCATTC
TATCCGCTGGAAGATGGAACCGCTGGAGAGCAACTGCATAAGGCTATGAAGAGATACGCCCTGGTTCCTGGAACAATTGCTTTTAC
AGATGCACATATCGAGGTGGACATCACTTACGCTGAGTACTTCGAAATGTCCGTTCGGTTGGCAGAAGCTATGAAACGATATGGGC
TGAATACAAATCACAGAATCGTCGTATGCAGTGAAAACTCTCTTCAATTCTTTATGCCGGTGTTGGGCGCGTTATTTATCGGAGTT
GCAGTTGCGCCCGCGAACGACATTTATAATGAACGTGAATTGCTCAACAGTATGGGCATTTCGCAGCCTACCGTGGTGTTCGTTTC
CAAAAAGGGGTTGCAAAAAATTTTGAACGTGCAAAAAAAGCTCCCAATCATCCAAAAAATTATTATCATGGATTCTAAAACGGATT
ACCAGGGATTTCAGTCGATGTACACGTTCGTCACATCTCATCTACCTCCCGGTTTTAATGAATACGATTTTGTGCCAGAGTCCTTC
```

GATAGGGACAAGACAATTGCACTGATCATGAACTCCTCTGGATCTACTGGTCTGCCTAAAGGTGTCGCTCTGCCTCATAGAACTGC
CTGCGTGAGATTCTCGCATGCCAGAGATCCTATTTTTGGCAATCAAATCATTCCGGATACTGCGATTTTAAGTGTTGTTCCATTCC
ATCACGGTTTTGGAATGTTTACTACACTCGGATATTTGATATGTGGATTTCGAGTCGTCTTAATGTATAGATTTGAAGAAGAGCTG
TTTCTGAGGAGCCTTCAGGATTACAAGATTCAAAGTGCGCTGCTGGTGCCAACCCTATTCTCCTTCTTCGCCAAAAGCACTCTGAT
TGACAAATACGATTTATCTAATTTACACGAAATTGCTTCTGGTGGCGCTCCCCTCTCTAAGGAAGTCGGGGAAGCGGTTGCCAAGA
GGTTCCATCTGCCAGGTATCAGGCAAGGATATGGGCTCACTGAGACTACATCAGCTATTCTGATTACACCCGAGGGGGATGATAAA
CCGGGCGCGGTCGGTAAAGTTGTTCCATTTTTTGAAGCGAAGGTTGTGGATCTGGATACCGGGAAAACGCTGGGCGTTAATCAAAG
AGGCGAACTGTGTGTGAGAGGTCCTATGATTATGTCCGGTTATGTAAACAATCCGGAAGCGACCAACGCCTTGATTGACAAGGATG
GATGGCTACATTCTGGAGACATAGCTTACTGGGACGAAGACGAACACTTCTTCATCGTTGACCGCCTGAAGTCTCTGATTAAGTAC
AAAGGCTATCAGGTGGCTCCCGCTGAATTGGAATCCATCTTGCTCCAACACCCCAACATCTTCGACGCAGGTGTCGCAGGTCTTCC
CGACGATGACGCCGGTGAACTTCCCGCCGCCGTTGTTGTTTTGGAGCACGGAAAGACGATGACGGAAAAAGAGATCGTGGATTACG
TCGCCAGTCAAGTAACAACCGCGAAAAAGTTGCGCGGAGGAGTTGTGTTTGTGGACGAAGTACCGAAAGGTCTTACCGGAAAACTC
GACGCAAGAAAAATCAGAGAGATCCTCATAAAGGCCAAGAAGGGCGGAAAGATCGCCGTGTAACTCGAGGGTGAACAAACAAACGA
CACATCTAAGTAGTGCAAGAATGGCTTCTCAAGACTCAAGTGACAACCCTAGTGTAGTGTTCGCACTTTGATGATGATGATTCGAT
TAGCAGCAAGATACATTTTTGTTAATAGCTTTTGCATTTGGATGCCAATTAGTGTTAAAAAAAACCGATCTAATTCACTTGTAATA
ACGAAGAAGCCAGTGTCATTAAGCCAAGTTTAGGTGCTCAAGCCGTGTGAACAAGGGACAATTTAATGAGGCCAGGAGTAGAGCAA
TGCAACAGAGTTTGCATCATCAAAAATCGGATCCCGGGCCCGTCGACTGCAGAGGCCTGCATGCAAGCTTGGCGTAATCATGGTCA
TAGCTGTTTCCTGTGTGAAATTGTTATCCGCTCACAATTCCACACAACATACGAGCCGGAAGCATAAAGTGTAAAGCCTGGGGTGC
CTAATGAGTGAGCTAACTCACATTAATTGCGTTGCGCTCACTGCCCGCTTTCCAGTCGGGAAACCTGTCGTGCCAGCTGCATTAAT
GAATCGGCCAACGCGCGGGGAGAGGCGGTTTGCGTATTGGGCGCTCTTCCGCTTCCTCGCTCACTGACTCGCTGCGCTCGGTCGTT
CGGCTGCGGCGAGCGGTATCAGCTCACTCAAAGGCGGTAATACGGTTATCCACAGAATCAGGGGATAACGCAGGAAAGAACATGTG
AGCAAAAGGCCAGCAAAAGGCCAGGAACCGTAAAAAGGCCGCGTTGCTGGCGTTTTTCCATAGGCTCCGCCCCCCTGACGAGCATC
ACAAAAATCGACGCTCAAGTCAGAGGTGGCGAAACCCGACAGGACTATAAAGATACCAGGCGTTTCCCCCTGGAAGCTCCCTCGTG
CGCTCTCCTGTTCCGACCCTGCCGCTTACCGGATACCTGTCCGCCTTTCTCCCTTCGGGAAGCGTGGCGCTTTCTCATAGCTCACG
CTGTAGGTATCTCAGTTCGGTGTAGGTCGTTCGCTCCAAGCTGGGCTGTGTGCACGAACCCCCCGTTCAGCCCGACCGCTGCGCCT
TATCCGGTAACTATCGTCTTGAGTCCAACCCGGTAAGACACGACTTATCGCCACTGGCAGCAGCCACTGGTAACAGGATTAGCAGA
GCGAGGTATGTAGGCGGTGCTACAGAGTTCTTGAAGTGGTGGCCTAACTACGGCTACACTAGAAGAACAGTATTTGGTATCTGCGC
TCTGCTGAAGCCAGTTACCTTCGGAAAAAGAGTTGGTAGCTCTTGATCCGGCAAACAAACCACCGCTGGTAGCGGTGGTTTTTTTG
TTTGCAAGCAGCAGATTACGCGCAGAAAAAAAGGATCTCAAGAAGATCCTTTGATCTTTTCTACGGGGTCTGACGCTCAGTGGAAC
GAAAACTCACGTTAAGGGATTTTGGTCATGAGATTATCAAAAAGGATCTTCACCTAGATCCTTTTAAATTAAAAATGAAGTTTTAA
ATCAATCTAAAGTATATATGAGTAAACTTGGTCTGACAGTTACCAATGCTTAATCAGTGAGGCACCTATCTCAGCGATCTGTCTAT
TTCGTTCATCCATAGTTGCCTGACTCCCCGTCGTGTAGATAACTACGATACGGGAGGGCTTACCATCTGGCCCCAGTGCTGCAATG
ATACCGCGAGACCCACGCTCACCGGCTCCAGATTTATCAGCAATAAACCAGCCAGCCGGAAGGGCCGAGCGCAGAAGTGGTCCTGC
AACTTTATCCGCCTCCATCCAGTCTATTAATTGTTGCCGGGAAGCTAGAGTAAGTAGTTCGCCAGTTAATAGTTTGCGCAACGTTG
TTGCCATTGCTACAGGCATCGTGGTGTCACGCTCGTCGTTTGGTATGGCTTCATTCAGCTCCGGTTCCCAACGATCAAGGCGAGTT
ACATGATCCCCCATGTTGTGCAAAAAAGCGGTTAGCTCCTTCGGTCCTCCGATCGTTGTCAGAAGTAAGTTGGCCGCAGTGTTATC
ACTCATGGTTATGGCAGCACTGCATAATTCTCTTACTGTCATGCCATCCGTAAGATGCTTTTCTGTGACTGGTGAGTACTCAACCA
AGTCATTCTGAGAATAGTGTATGCGGCGACCGAGTTGCTCTTGCCCGGCGTCAATACGGGATAATACCGCGCCACATAGCAGAACT
TTAAAAGTGCTCATCATTGGAAAACGTTCTTCGGGGCGAAAACTCTCAAGGATCTTACCGCTGTTGAGATCCAGTTCGATGTAACC
CACTCGTGCACCCAACTGATCTTCAGCATCTTTTACTTTCACCAGCGTTTCTGGGTGAGCAAAAACAGGAAGGCAAAATGCCGCAA
AAAAGGGAATAAGGGCGACACGGAAATGTTGAATACTCATACTCTTCCTTTTTCAATATTATTGAAGCATTTATCAGGGTTATTGT
CTCATGAGCGGATACATATTTGAATGTATTTAGAAAAATAAACAAATAGGGGTTCCGCGCACATTTCCCCGAAAAGTGCCACCTGA
CGTCTAAGAAACCATTATTATCATGACATTAACCTATAAAAATAGGCGTATCACGAGGCCCTTTCGTCTCGCGCGTTTCGGTGATG
ACGGTGAAAACCTCTGACACATGCAGCTCCCGGAGACGGTCACAGCTTGTCTGTAAGCGGATGCCGGGAGCAGACAAGCCCGTCAG
GGCGCGTCAGCGGGTGTTGGCGGGTGTCGGGGCTGGCTTAACTATGCGGCATCAGAGCAGATTGTACTGAGAGTGCACCATATGCG
GTGTGAAATACCGCACAGATGCGTAAGGAGAAAATACCGCATCAGGCGCCATTCGCCATTCAGGCTGCGCAACTGTTGGGAAGGGC
GATCGGTGCGGGCCTCTTCGCTATTACGCCAGCTGGCGAAAGGGGGATGTGCTGCAAGGCGATTAAGTTGGGTAACGCCAGGGTTT
TCCCAGTCACGACGTTGTAAAACGACGGCCAGTGAATTCGAGCTCGGTACCTCGCGAATGCATCTAGAT

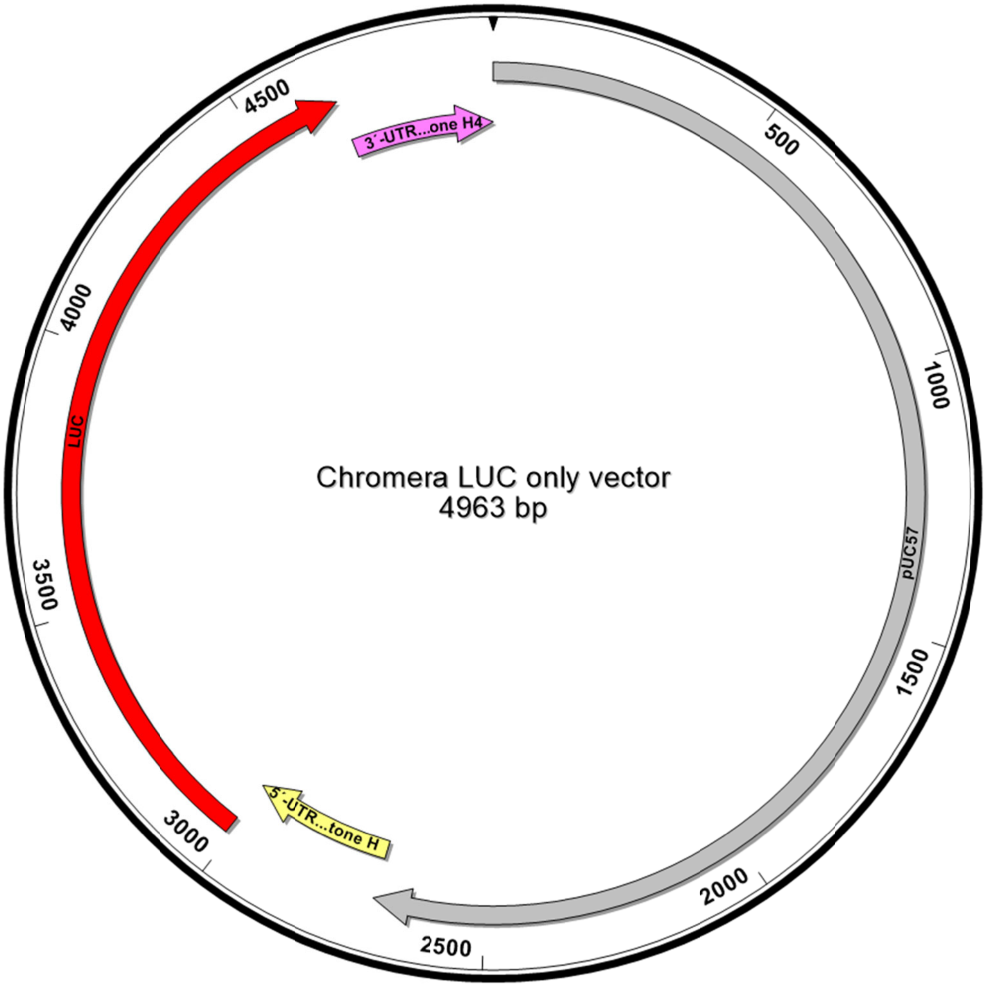


>Luc only vector

1-2270 pUC57 backbone
2771-3010 5´- UTR Histone H1
3011-4663 Luciferase
4664-4963 3´-UTR *Bigelowiella natans* histone H4

ATCGGATCCCGGGCCCGTCGACTGCAGAGGCCTGCATGCAAGCTTGGCGTAATCATGGTCATAGCTGTTTCCTGTGTGAAATTGTT
ATCCGCTCACAATTCCACACAACATACGAGCCGGAAGCATAAAGTGTAAAGCCTGGGGTGCCTAATGAGTGAGCTAACTCACATTA
ATTGCGTTGCGCTCACTGCCCGCTTTCCAGTCGGGAAACCTGTCGTGCCAGCTGCATTAATGAATCGGCCAACGCGCGGGGAGAGG
CGGTTTGCGTATTGGGCGCTCTTCCGCTTCCTCGCTCACTGACTCGCTGCGCTCGGTCGTTCGGCTGCGGCGAGCGGTATCAGCTC
ACTCAAAGGCGGTAATACGGTTATCCACAGAATCAGGGGATAACGCAGGAAAGAACATGTGAGCAAAAGGCCAGCAAAAGGCCAGG
AACCGTAAAAAGGCCGCGTTGCTGGCGTTTTTCCATAGGCTCCGCCCCCCTGACGAGCATCACAAAAATCGACGCTCAAGTCAGAG
GTGGCGAAACCCGACAGGACTATAAAGATACCAGGCGTTTCCCCCTGGAAGCTCCCTCGTGCGCTCTCCTGTTCCGACCCTGCCGC
TTACCGGATACCTGTCCGCCTTTCTCCCTTCGGGAAGCGTGGCGCTTTCTCATAGCTCACGCTGTAGGTATCTCAGTTCGGTGTAG
GTCGTTCGCTCCAAGCTGGGCTGTGTGCACGAACCCCCCGTTCAGCCCGACCGCTGCGCCTTATCCGGTAACTATCGTCTTGAGTC
CAACCCGGTAAGACACGACTTATCGCCACTGGCAGCAGCCACTGGTAACAGGATTAGCAGAGCGAGGTATGTAGGCGGTGCTACAG
AGTTCTTGAAGTGGTGGCCTAACTACGGCTACACTAGAAGAACAGTATTTGGTATCTGCGCTCTGCTGAAGCCAGTTACCTTCGGA
AAAAGAGTTGGTAGCTCTTGATCCGGCAAACAAACCACCGCTGGTAGCGGTGGTTTTTTTGTTTGCAAGCAGCAGATTACGCGCAG
AAAAAAAGGATCTCAAGAAGATCCTTTGATCTTTTCTACGGGGTCTGACGCTCAGTGGAACGAAAACTCACGTTAAGGGATTTTGG
TCATGAGATTATCAAAAAGGATCTTCACCTAGATCCTTTTAAATTAAAAATGAAGTTTTAAATCAATCTAAAGTATATATGAGTAA
ACTTGGTCTGACAGTTACCAATGCTTAATCAGTGAGGCACCTATCTCAGCGATCTGTCTATTTCGTTCATCCATAGTTGCCTGACT
CCCCGTCGTGTAGATAACTACGATACGGGAGGGCTTACCATCTGGCCCCAGTGCTGCAATGATACCGCGAGACCCACGCTCACCGG
CTCCAGATTTATCAGCAATAAACCAGCCAGCCGGAAGGGCCGAGCGCAGAAGTGGTCCTGCAACTTTATCCGCCTCCATCCAGTCT
ATTAATTGTTGCCGGGAAGCTAGAGTAAGTAGTTCGCCAGTTAATAGTTTGCGCAACGTTGTTGCCATTGCTACAGGCATCGTGGT
GTCACGCTCGTCGTTTGGTATGGCTTCATTCAGCTCCGGTTCCCAACGATCAAGGCGAGTTACATGATCCCCCATGTTGTGCAAAA
AAGCGGTTAGCTCCTTCGGTCCTCCGATCGTTGTCAGAAGTAAGTTGGCCGCAGTGTTATCACTCATGGTTATGGCAGCACTGCAT
AATTCTCTTACTGTCATGCCATCCGTAAGATGCTTTTCTGTGACTGGTGAGTACTCAACCAAGTCATTCTGAGAATAGTGTATGCG
GCGACCGAGTTGCTCTTGCCCGGCGTCAATACGGGATAATACCGCGCCACATAGCAGAACTTTAAAAGTGCTCATCATTGGAAAAC
GTTCTTCGGGGCGAAAACTCTCAAGGATCTTACCGCTGTTGAGATCCAGTTCGATGTAACCCACTCGTGCACCCAACTGATCTTCA
GCATCTTTTACTTTCACCAGCGTTTCTGGGTGAGCAAAAACAGGAAGGCAAAATGCCGCAAAAAAGGGAATAAGGGCGACACGGAA

ATGTTGAATACTCATACTCTTCCTTTTTCAATATTATTGAAGCATTTATCAGGGTTATTGTCTCATGAGCGGATACATATTTGAAT
GTATTTAGAAAAATAAACAAATAGGGGTTCCGCGCACATTTCCCCGAAAAGTGCCACCTGACGTCTAAGAAACCATTATTATCATG
ACATTAACCTATAAAAATAGGCGTATCACGAGGCCCTTTCGTCTCGCGCGTTTCGGTGATGACGGTGAAAACCTCTGACACATGCA
GCTCCCGGAGACGGTCACAGCTTGTCTGTAAGCGGATGCCGGGAGCAGACAAGCCCGTCAGGGCGCGTCAGCGGGTGTTGGCGGGT
GTCGGGGCTGGCTTAACTATGCGGCATCAGAGCAGATTGTACTGAGAGTGCACCATATGCGGTGTGAAATACCGCACAGATGCGTA
AGGAGAAAATACCGCATCAGGCGCCATTCGCCATTCAGGCTGCGCAACTGTTGGGAAGGGCGATCGGTGCGGGCCTCTTCGCTATT
ACGCCAGCTGGCGAAAGGGGGATGTGCTGCAAGGCGATTAAGTTGGGTAACGCCAGGGTTTTCCCAGTCACGACGTTGTAAAACGA
CGGCCAGTGAATTCGAGCTCGGTACCTCGCGAATGCATCTAGATCCTATATATAATTTTCTCTGTATCGGAAAATTAATTTCTTTT
TTTAAAAAATGAAATATTTCATTTTTCTTTTTTCTGTATCTGAAAAATCAAAACGAAAAATGATTCGACGTTGGGATCTCGGAAGG
AAATGACGAGACGTTCCAACGTCGAATCATTTTTTCGTTTGGTCTGACCGTGCAAAGTTGAGGGTCCCTCAGCACAGTTTTCTCCC
CTGATTTTTTTCCCCTTCGCTCTCCCCTTCCGGTCCCCTGTACGTAACCTTGTAAGAGACGCCCGTAGAATTCCACGTTGTCCAAG
ATGGAAGACGCCAAAAACATAAAGAAAGGCCCGGCGCCATTCTATCCGCTGGAAGATGGAACCGCTGGAGAGCAACTGCATAAGGC
TATGAAGAGATACGCCCTGGTTCCTGGAACAATTGCTTTTACAGATGCACATATCGAGGTGGACATCACTTACGCTGAGTACTTCG
AAATGTCCGTTCGGTTGGCAGAAGCTATGAAACGATATGGGCTGAATACAAATCACAGAATCGTCGTATGCAGTGAAAACTCTCTT
CAATTCTTTATGCCGGTGTTGGGCGCGTTATTTATCGGAGTTGCAGTTGCGCCCGCGAACGACATTTATAATGAACGTGAATTGCT
CAACAGTATGGGCATTTCGCAGCCTACCGTGGTGTTCGTTTCCAAAAAGGGGTTGCAAAAAATTTTGAACGTGCAAAAAAAGCTCC
CAATCATCCAAAAAATTATTATCATGGATTCTAAAACGGATTACCAGGGATTTCAGTCGATGTACACGTTCGTCACATCTCATCTA
CCTCCCGGTTTTAATGAATACGATTTTGTGCCAGAGTCCTTCGATAGGGACAAGACAATTGCACTGATCATGAACTCCTCTGGATC
TACTGGTCTGCCTAAAGGTGTCGCTCTGCCTCATAGAACTGCCTGCGTGAGATTCTCGCATGCCAGAGATCCTATTTTTGGCAATC
AAATCATTCCGGATACTGCGATTTTAAGTGTTGTTCCATTCCATCACGGTTTTGGAATGTTTACTACACTCGGATATTTGATATGT
GGATTTCGAGTCGTCTTAATGTATAGATTTGAAGAAGAGCTGTTTCTGAGGAGCCTTCAGGATTACAAGATTCAAAGTGCGCTGCT
GGTGCCAACCCTATTCTCCTTCTTCGCCAAAAGCACTCTGATTGACAAATACGATTTATCTAATTTACACGAAATTGCTTCTGGTG
GCGCTCCCCTCTCTAAGGAAGTCGGGGAAGCGGTTGCCAAGAGGTTCCATCTGCCAGGTATCAGGCAAGGATATGGGCTCACTGAG
ACTACATCAGCTATTCTGATTACACCCGAGGGGGATGATAAACCGGGCGCGGTCGGTAAAGTTGTTCCATTTTTTGAAGCGAAGGT
TGTGGATCTGGATACCGGGAAAACGCTGGGCGTTAATCAAAGAGGCGAACTGTGTGTGAGAGGTCCTATGATTATGTCCGGTTATG
TAAACAATCCGGAAGCGACCAACGCCTTGATTGACAAGGATGGATGGCTACATTCTGGAGACATAGCTTACTGGGACGAAGACGAA
CACTTCTTCATCGTTGACCGCCTGAAGTCTCTGATTAAGTACAAAGGCTATCAGGTGGCTCCCGCTGAATTGGAATCCATCTTGCT
CCAACACCCCAACATCTTCGACGCAGGTGTCGCAGGTCTTCCCGACGATGACGCCGGTGAACTTCCCGCCGCCGTTGTTGTTTTGG
AGCACGGAAAGACGATGACGGAAAAAGAGATCGTGGATTACGTCGCCAGTCAAGTAACAACCGCGAAAAAGTTGCGCGGAGGAGTT
GTGTTTGTGGACGAAGTACCGAAAGGTCTTACCGGAAAACTCGACGCAAGAAAAATCAGAGAGATCCTCATAAAGGCCAAGAAGGG
CGGAAAGATCGCCGTGTAAATTGCTCACAGCATTCTCAATGCTACTACATCCGGTGTTCTTAAACACCACCATTCTCTTCGGAAAC
TGTGCCTTAGTCAAAAATTGTAAACCGATAGAAAATAAATAGAAAATGATGTTATCTAGTGCATAAGTAGCGAGTTGGAATACGTT
CCTACGTCTCCTTGTCAACACCACGATGTAATTCAAGTGATTAGCCATGCTTGATGGCAGATGAACAAAGGGAATTCATATTGTAC
ATGATTAAGCGTAGTGGAGCCAGTGAAGAACTCCCACACCCGCCCATCCTAGCTAGTGAAC

*Perkinsus marinus*

Plasmids for transfection of *Perkinsus marinus* using Golden Gate modules

LevelM:BLEO:MOE:mCHERRY 6647 bp

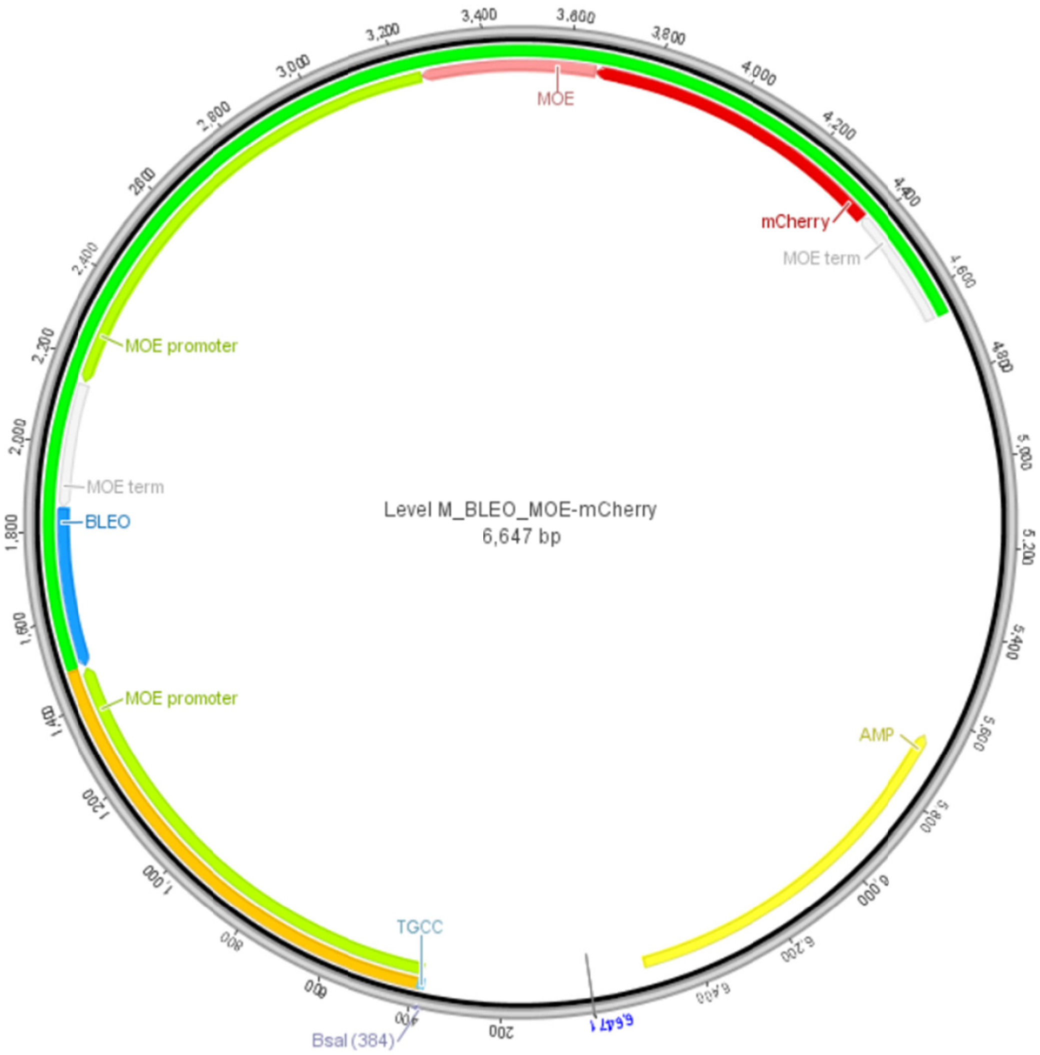


| Name | Part | Start | End |
|---|---|---|---|
| MOE promoter | Promoter | 388 | 1,481 |
| BLEO | CDS | 1,486 | 1,857 |
| MOE term | 3'UTR | 1,858 | 2,148 |
| MOE promoter | Promoter | 2,153 | 3,253 |
| MOE | CDS | 3,254 | 3,659 |
| mCherry | CDS | 3,66 | 4,373 |

| | | | |
|---|---|---|---|
| MOE term | 3'UTR | 4,374 | 4,663 |
| AMP | CDS | 5,659 | 6,519 |

**> LevelM:BLEO:MOE:mCHERRY**

```
CTAAATTGTAAGCGTTAATATTTTGTTAAAATTCGCGTTAAATTTTTGTTAAATCAGCTCATTTTTTAACCAATAG
GCCGAAATCGGCAAAATCCCTTATAAATCAAAAGAATAGACCGAGATAGGGTTGAGTGGCCGCTACAGGGCGCTCC
CATTCGCCATTCAGGCTGCGCAACTGTTGGGAAGGGCGTTTCGGTGCGGGCCTCTTCGCTATTACGCCAGCTGGCG
AAAGGGGGATGTGCTGCAAGGCGATTAAGTTGGGTAACGCCAGGGTTTTCCCAGTCACGACGTTGTAAAACGACGG
CCAGTGAGCGCGACGTAATACGACTCACTATAGGGCGAATTGGCGGAAGGCCGTCAAGGCCTAGGCGCGCCAGGTC
TCATGCCCTCGTAATGAGCCCAACCATTATCGGATCGGTCTTGCGGGCGAACGAACATCTGCGGCAGGTGAGAGGT
GGGAGTTTCCGGATGCGGTCCGAGTCACCGGCGATGGAACTCCCCACAGGCCAGTGTGACGGTGAGCACTGATGTT
CGATGGAAACCAAAGTGGCAGGGTTCGTACTTCCCAGCCATGGTTGCGCCGTTTGTAATTGATGTGATGGCAGAGC
TGACCGGTAGCAGTAGAAGGTGCGTTGATTCAGCTCAACAGCTCTCTTGTGTTAGCAGAAAGTAGAAGCAGATCTA
GATGATTCAGGAATCCCTCATCTTAGCGGTAGTATGTTACTGTGCACCCATCATGCCATGGATGATCCGGATCCCT
TGTTGGGTGAGGTAGGTTCCTTGGTGAGTTGGATGATTGTGGATCTGCCGATGGCAGAGATCATGGCTCCATTCCC
CATCGGGACCTCACGGTATCACTGATCATTTCATGGCTCAAGGGCGGGCTTGTCGACTTCGACAAGTGGGTGGGAC
TGCGGTGAGCAAGGTGCTTTGCACTTGCCGAACCCATCGCTTTGCTCCGGCTTGTAGTGAACTAACCGGAGCAAGA
ATAGGGCCCGGGCTAGTGGGACGCAACACATGAGGAACGTGTCTCAAATCCATAGATTTTATGTTGGATTCGTCTC
GTCGGAGCCAAGTGCTCCACGATGTGCATCCCTCGAGTCGAATCTTCCACTAGAACCCTTGAGCGTCAGCGAGCTA
GAGAACTTAGGAGCGGAGGGACGGGTTCTGCTAGTAGACATCAACAGATATCCCTTCACGCTCCCTTCATTGTATG
CGTGAGTATGTATCGCGGAGCAACACAGTCTTGTCCATCAAAGGGCATTTCTCCACGAATCTCGCACTAGCTTGGA
AATCATAATCTCGAGTCACTTGATTTCGTAACGCTGAATCCTTCTAAATGGACTCTATGTTGTAAGGCGAGACGCT
ATAACGATGTCTATCGACCGGACGCATTGAACGCATACACATAGCTAAGTGAGTTGTGTGGAACTCTTGATGAACT
CGCGTCACTCTTGAATCTCTATCCCGCTGTTCAACGCAATGGCCAAGTTGACCAGTGCCGTTCCGGTGCTCACCGC
GCGCGACGTCGCCGGAGCGGTCGAGTTCTGGACCGACCGGCTCGGGTTCTCCCGGGACTTCGTGGAGGACGACTTC
GCCGGTGTGGTCCGGGACGACGTGACCCTGTTCATCAGCGCGGTCCAGGACCAGGTGGTGCCGGACAACACCCTGG
CCTGGGTGTGGGTGCGCGGCCTGGACGAGCTGTACGCCGAGTGGTCGGAGGTCGTGTCCACGAACTTCCGGGACGC
CTCCGGGCCGGCCATGACCGAGATCGGCGAGCAGCCGTGGGGGCGGGAGTTCGCCCTGCGCGACCCGGCCGGCAAC
TGCGTGCACTTCGTGGCCGAGGAGCAGGACTGAGCTTGACACACATCTAGTTGAGATCTACATCAATGTTCGAAAT
CAACCATGCTTATAGCCATTGATATCTCTCTACTCTTTTCCGACCCTTATTACAAGGTCTATGACCAAACTATTGA
GCCTACGGTTTGTGACTCGCCAGGTTAAGCGATGTCACATCGCAAGTCGTTGAGGATGTACTGGTTATGAATGGCT
CTTCTTTCTTTCTGTGACTCGTTGGCATTGTTAGCGGATCTACAAGTAACCGTCGACGATCACGAGAGCTTCCGGC
TCCATGATTGGGTCATCGCTGCAAGAGGCTCGTAATGAGCCCAACCATTATCGGATCGGTCTTGCGGGCGAACGAA
CATCTGCGGCAGGTGAGAGGTGGGAGTTTCCGGATGCGGTCCGAGTCACCGGCGATGGAACTCCCCACAGGCCAGT
GTGACGGTGAGCACTGATGTTCGATGGAAACCAAAGTGGCAGGGTTCGTACTTCCCAGCCATGGTTGCGCCGTTTG
TAATTGATGTGATGGCAGAGCTGACCGGTAGCAGTAGAAGGTGCGTTGATTCAGCTCAACAGCTCTCTTGTGTTAG
CAGAAAGTAGAAGCAGATCTAGATGATTCAGGAATCCCTCATCTTAGCGGTAGTATGTTACTGTGCACCCATCATG
CCATGGATGATCCGGATCCCTTGTTGGGTGAGGTAGGTTCCTTGGTGAGTTGGATGATTGTGGATCTGCCGATGGC
AGAGATCATGGCTCCATTCCCCATCGGGACCTCACGGTATCACTGATCATTTCATGGCTCAAGGGCGGGCTTGTCG
ACTTCGACAAGTGGGTGGGACTGCGGTGAGCAAGGTGCTTTGCACTTGCCGAACCCATCGCTTTGCTCCGGCTTGT
AGTGAACTAACCGGAGCAAGAATAGGGCCCGGGCTAGTGGGACGCAACACATGAGGAACGTGTCTCAAATCCATAG
ATTTTATGTTGGATTCGTCTCGTCGGAGCCAAGTGCTCCACGATGTGCATCCCTCGAGTCGAATCTTCCACTAGAA
CCCTTGAGCGTCAGCGAGCTAGAGAACTTAGGAGCGGAGGGACGGGTTCTGCTAGTAGACATCAACAGATATCCCT
TCACGCTCCCTTCATTGTATGCGTGAGTATGTATCGCGGAGCAACACAGTCTTGTCCATCAAAGGGCATTTCTCCA
CGAATCTCGCACTAGCTTGGAAATCATAATCTCGAGTCACTTGATTTCGTAACGCTGAATCCTTCTAAATGGACTC
TATGTTGTAAGGCGAGACGCTATAACGATGTCTATCGACCGGACGCATTGAACGCATACACATAGCTAAGTGAGTT
GTGTGGAACTCTTGATGAACTCGCGTCACTCTTGAATCTCTATCCCGCTGTTCAACGCAATAATGCGCTTCATTGT
TGGTCTGTACTCCTGCCTGGCCGTTTTGGTTCTTGGGCAGTCCTCTTGTCCCACAGGGGATGCTCAGTGTCACAGT
GAGAGTCCCGGTTCGTACTGCAAGTTCTATCAAGGTCACAATGTCTGCCAGGGAACTGACACCCCCTGCTCTTGCA
CCACTACTCCAGCACCGCCGCCACCCACTCCCACTGGTGGATCATGTCCTGATGGTGATGCTTTTTGTCAGAAGCA
GTCTGGTAATTCCGGCTCCTATTGCAAGGCTGCTTACCAGACTGGTCCCGGTGGAGGTGTTTGCCAGGGTACTGAT
ATCCCCTGTAAATGTGGTGGTGGCGGATCAACTACGATGAAGCCGACTACGACGAAGCCGACTACGACGAAGCCGA
CATGTCCTACTTCGATGGTGAGCAAGGGCGAGGAGGATAACATGGCCATCATCAAGGAGTTCATGCGCTTCAAGGT
GCACATGGAGGGCTCCGTGAACGGCCACGAGTTCGAGATCGAGGGCGAGGGCGAGGGCCGCCCCTACGAGGGCACC
CAGACCGCCAAGCTGAAGGTGACCAAGGGTGGCCCCCTGCCCTTCGCCTGGGACATCCTGTCCCCTCAGTTCATGT
ACGGCTCCAAGGCCTACGTGAAGCACCCCGCCGACATCCCCGACTACTTGAAGCTGTCCTTCCCCGAGGGCTTCAA
GTGGGAGCGCGTGATGAACTTCGAGGACGGCGGCGTGGTGACCGTGACCCAGGACTCCTCCCTGCAGGACGGCGAG
TTCATCTACAAGGTGAAGCTGCGCGGCACCAACTTCCCCTCCGACGGCCCCGTAATGCAGAAGAAAACCATGGGCT
GGGAGGCCTCCTCCGAGCGGATGTACCCCGAGGACGGCGCCCTGAAGGGCGAGATCAAGCAGAGGCTGAAGCTGAA
GGACGGCGGCCACTACGACGCTGAGGTCAAGACCACCTACAAGGCCAAGAAGCCCGTGCAGCTGCCCGGCGCCTAC
AACGTCAACATCAAGTTGGACATCACCTCCCACAACGAGGACTACACCATCGTGGAACAGTACGAACGCGCCGAGG
GCCGCCACTCCACCGGCGGCATGGACGAGCTGTACAAGTGAGCTTGACACACATCTAGTTGAGATCTACATCAATG
TTCGAAATCAACCATGCTTATAGCCATTGATATCTCTCTACTCTTTTCCGACCCTTATTACAAGGTCTATGACCAA
ACTATTGAGCCTACGGTTTGTGACTCGCCAGGTTAAGCGATGTCACATCGCAAGTCGTTGAGGATGTACTGGTTAT
GAATGGCTCTTCTTTCTTTCTGTGACTCGTTGGCATTGTTAGCGGATCTACAAGTAACCGTCGACGATCACGAGAG
```

CTTCCGGCTCCATGATTGGGTCAGCAAGGGAATTAATTAACTGGCCTCATGGGCCTTCCGCTCACTGCCCGCTTTC
CAGTCGGGAAACCTGTCGTGCCAGCTGCATTAACATGGTCATAGCTGTTTCCTTGCGTATTGGGCGCTCTCCGCTT
CCTCGCTCACTGACTCGCTGCGCTCGGTCGTTCGGGTAAAGCCTGGGGTGCCTAATGAGCAAAAGGCCAGCAAAAG
GCCAGGAACCGTAAAAAGGCCGCGTTGCTGGCGTTTTTCCATAGGCTCCGCCCCCCTGACGAGCATCACAAAAATC
GACGCTCAAGTCAGAGGTGGCGAAACCCGACAGGACTATAAAGATACCAGGCGTTTCCCCCTGGAAGCTCCCTCGT
GCGCTCTCCTGTTCCGACCCTGCCGCTTACCGGATACCTGTCCGCCTTTCTCCCTTCGGGAAGCGTGGCGCTTTCT
CATAGCTCACGCTGTAGGTATCTCAGTTCGGTGTAGGTCGTTCGCTCCAAGCTGGGCTGTGTGCACGAACCCCCCG
TTCAGCCCGACCGCTGCGCCTTATCCGGTAACTATCGTCTTGAGTCCAACCCGGTAAGACACGACTTATCGCCACT
GGCAGCAGCCACTGGTAACAGGATTAGCAGAGCGAGGTATGTAGGCGGTGCTACAGAGTTCTTGAAGTGGTGGCCT
AACTACGGCTACACTAGAAGAACAGTATTTGGTATCTGCGCTCTGCTGAAGCCAGTTACCTTCGGAAAAAGAGTTG
GTAGCTCTTGATCCGGCAAACAAACCACCGCTGGTAGCGGTGGTTTTTTTGTTTGCAAGCAGCAGATTACGCGCAG
AAAAAAAGGATCTCAAGAAGATCCTTTGATCTTTTCTACGGGGTCTGACGCTCAGTGGAACGAAAACTCACGTTAA
GGGATTTTGGTCATGAGATTATCAAAAAGGATCTTCACCTAGATCCTTTTAAATTAAAAATGAAGTTTTAAATCAA
TCTAAAGTATATATGAGTAAACTTGGTCTGACAGTTACCAATGCTTAATCAGTGAGGCACCTATCTCAGCGATCTG
TCTATTTCGTTCATCCATAGTTGCCTGACTCCCCGTCGTGTAGATAACTACGATACGGGAGGGCTTACCATCTGGC
CCCAGTGCTGCAATGATACCGCGAGAACCACGCTCACCGGCTCCAGATTTATCAGCAATAAACCAGCCAGCCGGAA
GGGCCGAGCGCAGAAGTGGTCCTGCAACTTTATCCGCCTCCATCCAGTCTATTAATTGTTGCCGGGAAGCTAGAGT
AAGTAGTTCGCCAGTTAATAGTTTGCGCAACGTTGTTGCCATTGCTACAGGCATCGTGGTGTCACGCTCGTCGTTT
GGTATGGCTTCATTCAGCTCCGGTTCCCAACGATCAAGGCGAGTTACATGATCCCCCATGTTGTGCAAAAAAGCGG
TTAGCTCCTTCGGTCCTCCGATCGTTGTCAGAAGTAAGTTGGCCGCAGTGTTATCACTCATGGTTATGGCAGCACT
GCATAATTCTCTTACTGTCATGCCATCCGTAAGATGCTTTTCTGTGACTGGTGAGTACTCAACCAAGTCATTCTGA
GAATAGTGTATGCGGCGACCGAGTTGCTCTTGCCCGGCGTCAATACGGGATAATACCGCGCCACATAGCAGAACTT
TAAAAGTGCTCATCATTGGAAAACGTTCTTCGGGGCGAAAACTCTCAAGGATCTTACCGCTGTTGAGATCCAGTTC
GATGTAACCCACTCGTGCACCCAACTGATCTTCAGCATCTTTTACTTTCACCAGCGTTTCTGGGTGAGCAAAAACA
GGAAGGCAAAATGCCGCAAAAAAGGGAATAAGGGCGACACGGAAATGTTGAATACTCATACTCTTCCTTTTTCAAT
ATTATTGAAGCATTTATCAGGGTTATTGTCTCATGAGCGGATACATATTTGAATGTATTTAGAAAAATAAACAAAT
AGGGGTTCCGCGCACATTTCCCCGAAAAGTGCCAC

LevelM:BSR:MOE:mCHERRY 6665 bp

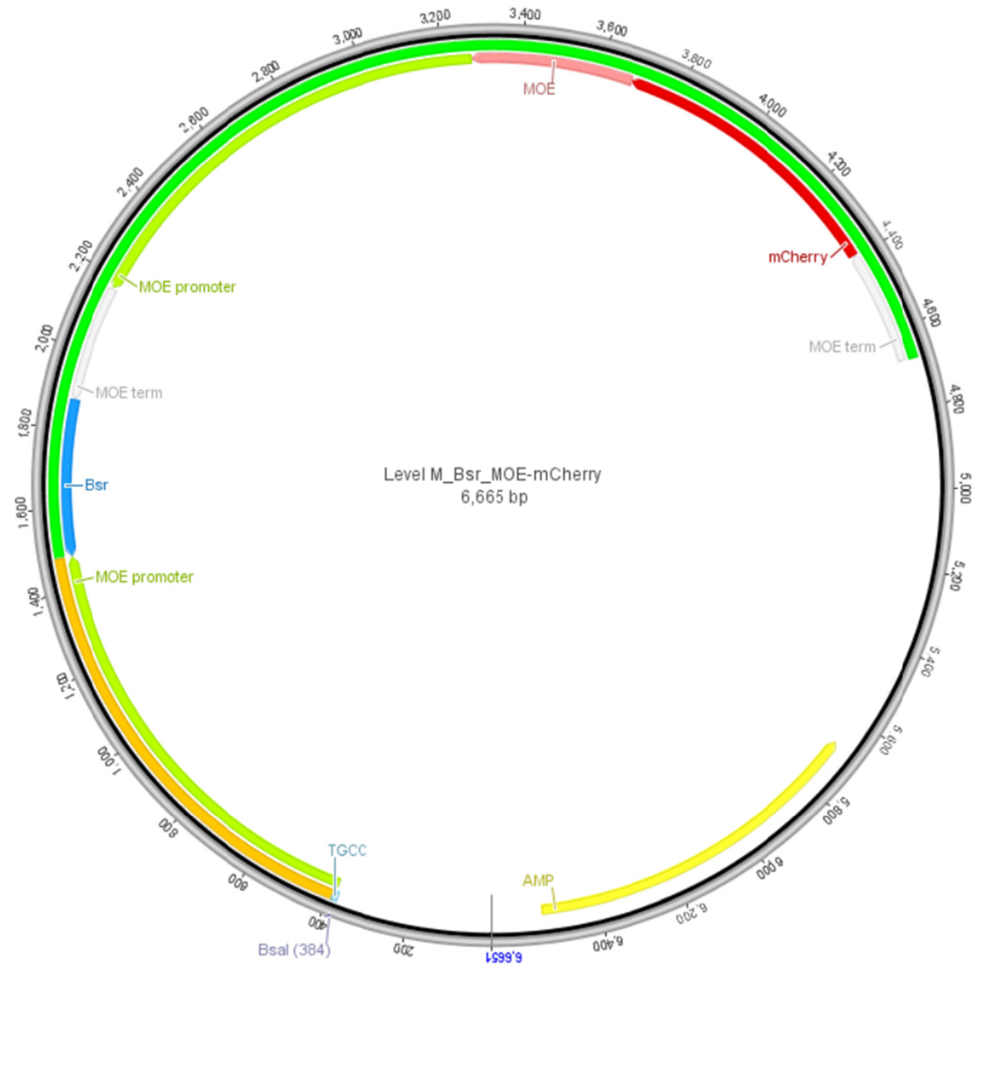

| **Name** | **Part** | **Start** | **End** |
|---|---|---|---|
| MOE promoter | Promoter | 388 | 1,481 |
| Bsr | CDS | 1,486 | 1,875 |
| MOE term | 3'UTR | 1,876 | 2,166 |
| MOE promoter | Promoter | 2,171 | 3,271 |
| MOE | CDS | 3,272 | 3,677 |
| mCherry | CDS | 3,678 | 4,391 |
| MOE term | 3'UTR | 4,392 | 4,681 |
| AMP | CDS | 5,677 | 6,537 |

```
> LevelM:BSR:MOE:mCHERRY
CTAAATTGTAAGCGTTAATATTTTGTTAAAATTCGCGTTAAATTTTTGTTAAATCAGCTCATTTTTTAACCAATAG
GCCGAAATCGGCAAAATCCCTTATAAATCAAAAGAATAGACCGAGATAGGGTTGAGTGGCCGCTACAGGGCGCTCC
CATTCGCCATTCAGGCTGCGCAACTGTTGGGAAGGGCGTTTCGGTGCGGGCCTCTTCGCTATTACGCCAGCTGGCG
AAAGGGGGATGTGCTGCAAGGCGATTAAGTTGGGTAACGCCAGGGTTTTCCCAGTCACGACGTTGTAAAACGACGG
CCAGTGAGCGCGACGTAATACGACTCACTATAGGGCGAATTGGCGGAAGGCCGTCAAGGCCTAGGCGCGCCAGGTC
TCATGCCCTCGTAATGAGCCCAACCATTATCGGATCGGTCTTGCGGGCGAACGAACATCTGCGGCAGGTGAGAGGT
GGGAGTTTCCGGATGCGGTCCGAGTCACCGGCGATGGAACTCCCCACAGGCCAGTGTGACGGTGAGCACTGATGTT
CGATGGAAACCAAAGTGGCAGGGTTCGTACTTCCCAGCCATGGTTGCGCCGTTTGTAATTGATGTGATGGCAGAGC
TGACCGGTAGCAGTAGAAGGTGCGTTGATTCAGCTCAACAGCTCTCTTGTGTTAGCAGAAAGTAGAAGCAGATCTA
GATGATTCAGGAATCCCTCATCTTAGCGGTAGTATGTTACTGTGCACCCATCATGCCATGGATGATCCGGATCCCT
TGTTGGGTGAGGTAGGTTCCTTGGTGAGTTGGATGATTGTGGATCTGCCGATGGCAGAGATCATGGCTCCATTCCC
CATCGGGACCTCACGGTATCACTGATCATTTCATGGCTCAAGGGCGGGCTTGTCGACTTCGACAAGTGGGTGGGAC
TGCGGTGAGCAAGGTGCTTTGCACTTGCCGAACCCATCGCTTTGCTCCGGCTTGTAGTGAACTAACCGGAGCAAGA
ATAGGGCCCGGGCTAGTGGGACGCAACACATGAGGAACGTGTCTCAAATCCATAGATTTTATGTTGGATTCGTCTC
GTCGGAGCCAAGTGCTCCACGATGTGCATCCCTCGAGTCGAATCTTCCACTAGAACCCTTGAGCGTCAGCGAGCTA
GAGAACTTAGGAGCGGAGGGACGGGTTCTGCTAGTAGACATCAACAGATATCCCTTCACGCTCCCTTCATTGTATG
CGTGAGTATGTATCGCGGAGCAACACAGTCTTGTCCATCAAAGGGCATTTCTCCACGAATCTCGCACTAGCTTGGA
AATCATAATCTCGAGTCACTTGATTTCGTAACGCTGAATCCTTCTAAATGGACTCTATGTTGTAAGGCGAGACGCT
ATAACGATGTCTATCGACCGGACGCATTGAACGCATACACATAGCTAAGTGAGTTGTGTGGAACTCTTGATGAACT
CGCGTCACTCTTGAATCTCTATCCCGCTGTTCAACGCAATGCCTTTGTCTCAAGAAGAATCCACCCTCATTGAAAG
AGCAACGGCTACAATCAACAGCATCCCCATCTCTGAGGACTACAGCGTCGCCAGCGCAGCTCTCTCTAGCGACGGC
CGCATCTTCACTGGTGTCAATGTATATCATTTTACTGGGGGACCTTGTGCAGAACTCGTGGTGCTGGGCACTGCTG
CTGCTGCGGCAGCTGGCAACCTGACTTGTATCGTCGCGATCGGAAATGAGAACAGGGGCATCTTGAGCCCCTGCGG
ACGGTGCCGACAGGTGCTTCTCGATCTGCATCCTGGGATCAAAGCCATAGTGAAGGACAGTGATGGACAGCCGACG
GCAGTTGGGATTCGTGAATTGCTGCCCTCTGGTTATGTGTGGGAGGGCTAAGCTTGACACACATCTAGTTGAGATC
TACATCAATGTTCGAAATCAACCATGCTTATAGCCATTGATATCTCTCTACTCTTTTCCGACCCTTATTACAAGGT
CTATGACCAAACTATTGAGCCTACGGTTTGTGACTCGCCAGGTTAAGCGATGTCACATCGCAAGTCGTTGAGGATG
TACTGGTTATGAATGGCTCTTCTTTCTTTCTGTGACTCGTTGGCATTGTTAGCGGATCTACAAGTAACCGTCGACG
ATCACGAGAGCTTCCGGCTCCATGATTGGGTCATCGCTGCAAGAGGCTCGTAATGAGCCCAACCATTATCGGATCG
GTCTTGCGGGCGAACGAACATCTGCGGCAGGTGAGAGGTGGGAGTTTCCGGATGCGGTCCGAGTCACCGGCGATGG
AACTCCCCACAGGCCAGTGTGACGGTGAGCACTGATGTTCGATGGAAACCAAAGTGGCAGGGTTCGTACTTCCCAG
CCATGGTTGCGCCGTTTGTAATTGATGTGATGGCAGAGCTGACCGGTAGCAGTAGAAGGTGCGTTGATTCAGCTCA
ACAGCTCTCTTGTGTTAGCAGAAAGTAGAAGCAGATCTAGATGATTCAGGAATCCCTCATCTTAGCGGTAGTATGT
TACTGTGCACCCATCATGCCATGGATGATCCGGATCCCTTGTTGGGTGAGGTAGGTTCCTTGGTGAGTTGGATGAT
TGTGGATCTGCCGATGGCAGAGATCATGGCTCCATTCCCCATCGGGACCTCACGGTATCACTGATCATTTCATGGC
TCAAGGGCGGGCTTGTCGACTTCGACAAGTGGGTGGGACTGCGGTGAGCAAGGTGCTTTGCACTTGCCGAACCCAT
CGCTTTGCTCCGGCTTGTAGTGAACTAACCGGAGCAAGAATAGGGCCCGGGCTAGTGGGACGCAACACATGAGGAA
CGTGTCTCAAATCCATAGATTTTATGTTGGATTCGTCTCGTCGGAGCCAAGTGCTCCACGATGTGCATCCCTCGAG
TCGAATCTTCCACTAGAACCCTTGAGCGTCAGCGAGCTAGAGAACTTAGGAGCGGAGGGACGGGTTCTGCTAGTAG
ACATCAACAGATATCCCTTCACGCTCCCTTCATTGTATGCGTGAGTATGTATCGCGGAGCAACACAGTCTTGTCCA
TCAAAGGGCATTTCTCCACGAATCTCGCACTAGCTTGGAAATCATAATCTCGAGTCACTTGATTTCGTAACGCTGA
ATCCTTCTAAATGGACTCTATGTTGTAAGGCGAGACGCTATAACGATGTCTATCGACCGGACGCATTGAACGCATA
CACATAGCTAAGTGAGTTGTGTGGAACTCTTGATGAACTCGCGTCACTCTTGAATCTCTATCCCGCTGTTCAACGC
AATAATGCGCTTCATTGTTGGTCTGTACTCCTGCCTGGCCGTTTTGGTTCTTGGGCAGTCCTCTTGTCCCACAGGG
GATGCTCAGTGTCACAGTGAGAGTCCCGGTTCGTACTGCAAGTTCTATCAAGGTCACAATGTCTGCCAGGGAACTG
```

ACACCCCCTGCTCTTGCACCACTACTCCAGCACCGCCGCCACCCACTCCCACTGGTGGATCATGTCCTGATGGTGA
TGCTTTTTGTCAGAAGCAGTCTGGTAATTCCGGCTCCTATTGCAAGGCTGCTTACCAGACTGGTCCCGGTGGAGGT
GTTTGCCAGGGTACTGATATCCCCTGTAAATGTGGTGGTGGCGGATCAACTACGATGAAGCCGACTACGACGAAGC
CGACTACGACGAAGCCGACATGTCCTACTTCGATGGTGAGCAAGGGCGAGGAGGATAACATGGCCATCATCAAGGA
GTTCATGCGCTTCAAGGTGCACATGGAGGGCTCCGTGAACGGCCACGAGTTCGAGATCGAGGGCGAGGGCGAGGGC
CGCCCCTACGAGGGCACCCAGACCGCCAAGCTGAAGGTGACCAAGGGTGGCCCCCTGCCCTTCGCCTGGGACATCC
TGTCCCCTCAGTTCATGTACGGCTCCAAGGCCTACGTGAAGCACCCCGCCGACATCCCCGACTACTTGAAGCTGTC
CTTCCCCGAGGGCTTCAAGTGGGAGCGCGTGATGAACTTCGAGGACGGCGGCGTGGTGACCGTGACCCAGGACTCC
TCCCTGCAGGACGGCGAGTTCATCTACAAGGTGAAGCTGCGCGGCACCAACTTCCCCTCCGACGGCCCCGTAATGC
AGAAGAAAACCATGGGCTGGGAGGCCTCCTCCGAGCGGATGTACCCCGAGGACGGCGCCCTGAAGGGCGAGATCAA
GCAGAGGCTGAAGCTGAAGGACGGCGGCCACTACGACGCTGAGGTCAAGACCACCTACAAGGCCAAGAAGCCCGTG
CAGCTGCCCGGCGCCTACAACGTCAACATCAAGTTGGACATCACCTCCCACAACGAGGACTACACCATCGTGGAAC
AGTACGAACGCGCCGAGGGCCGCCACTCCACCGGCGGCATGGACGAGCTGTACAAGTGAGCTTGACACACATCTAG
TTGAGATCTACATCAATGTTCGAAATCAACCATGCTTATAGCCATTGATATCTCTCTACTCTTTTCCGACCCTTAT
TACAAGGTCTATGACCAAACTATTGAGCCTACGGTTTGTGACTCGCCAGGTTAAGCGATGTCACATCGCAAGTCGT
TGAGGATGTACTGGTTATGAATGGCTCTTCTTTCTTTCTGTGACTCGTTGGCATTGTTAGCGGATCTACAAGTAAC
CGTCGACGATCACGAGAGCTTCCGGCTCCATGATTGGGTCAGCAAGGGAATTAATTAACTGGCCTCATGGGCCTTC
CGCTCACTGCCCGCTTTCCAGTCGGGAAACCTGTCGTGCCAGCTGCATTAACATGGTCATAGCTGTTTCCTTGCGT
ATTGGGCGCTCTCCGCTTCCTCGCTCACTGACTCGCTGCGCTCGGTCGTTCGGGTAAAGCCTGGGGTGCCTAATGA
GCAAAAGGCCAGCAAAAGGCCAGGAACCGTAAAAAGGCCGCGTTGCTGGCGTTTTTCCATAGGCTCCGCCCCCCTG
ACGAGCATCACAAAAATCGACGCTCAAGTCAGAGGTGGCGAAACCCGACAGGACTATAAAGATACCAGGCGTTTCC
CCCTGGAAGCTCCCTCGTGCGCTCTCCTGTTCCGACCCTGCCGCTTACCGGATACCTGTCCGCCTTTCTCCCTTCG
GGAAGCGTGGCGCTTTCTCATAGCTCACGCTGTAGGTATCTCAGTTCGGTGTAGGTCGTTCGCTCCAAGCTGGGCT
GTGTGCACGAACCCCCCGTTCAGCCCGACCGCTGCGCCTTATCCGGTAACTATCGTCTTGAGTCCAACCCGGTAAG
ACACGACTTATCGCCACTGGCAGCAGCCACTGGTAACAGGATTAGCAGAGCGAGGTATGTAGGCGGTGCTACAGAG
TTCTTGAAGTGGTGGCCTAACTACGGCTACACTAGAAGAACAGTATTTGGTATCTGCGCTCTGCTGAAGCCAGTTA
CCTTCGGAAAAAGAGTTGGTAGCTCTTGATCCGGCAAACAAACCACCGCTGGTAGCGGTGGTTTTTTTGTTTGCAA
GCAGCAGATTACGCGCAGAAAAAAAGGATCTCAAGAAGATCCTTTGATCTTTTCTACGGGGTCTGACGCTCAGTGG
AACGAAAACTCACGTTAAGGGATTTTGGTCATGAGATTATCAAAAAGGATCTTCACCTAGATCCTTTTAAATTAAA
AATGAAGTTTTAAATCAATCTAAAGTATATATGAGTAAACTTGGTCTGACAGTTACCAATGCTTAATCAGTGAGGC
ACCTATCTCAGCGATCTGTCTATTTCGTTCATCCATAGTTGCCTGACTCCCCGTCGTGTAGATAACTACGATACGG
GAGGGCTTACCATCTGGCCCCAGTGCTGCAATGATACCGCGAGAACCACGCTCACCGGCTCCAGATTTATCAGCAA
TAAACCAGCCAGCCGGAAGGGCCGAGCGCAGAAGTGGTCCTGCAACTTTATCCGCCTCCATCCAGTCTATTAATTG
TTGCCGGGAAGCTAGAGTAAGTAGTTCGCCAGTTAATAGTTTGCGCAACGTTGTTGCCATTGCTACAGGCATCGTG
GTGTCACGCTCGTCGTTTGGTATGGCTTCATTCAGCTCCGGTTCCCAACGATCAAGGCGAGTTACATGATCCCCCA
TGTTGTGCAAAAAAGCGGTTAGCTCCTTCGGTCCTCCGATCGTTGTCAGAAGTAAGTTGGCCGCAGTGTTATCACT
CATGGTTATGGCAGCACTGCATAATTCTCTTACTGTCATGCCATCCGTAAGATGCTTTTCTGTGACTGGTGAGTAC
TCAACCAAGTCATTCTGAGAATAGTGTATGCGGCGACCGAGTTGCTCTTGCCCGGCGTCAATACGGGATAATACCG
CGCCACATAGCAGAACTTTAAAAGTGCTCATCATTGGAAAACGTTCTTCGGGGCGAAAACTCTCAAGGATCTTACC
GCTGTTGAGATCCAGTTCGATGTAACCCACTCGTGCACCCAACTGATCTTCAGCATCTTTTACTTTCACCAGCGTT
TCTGGGTGAGCAAAAACAGGAAGGCAAAATGCCGCAAAAAAGGGAATAAGGGCGACACGGAAATGTTGAATACTCA
TACTCTTCCTTTTTCAATATTATTGAAGCATTTATCAGGGTTATTGTCTCATGAGCGGATACATATTTGAATGTAT
TTAGAAAAATAAACAAATAGGGGTTCCGCGCACATTTCCCCGAAAAGTGCCAC

LevelM:PAC:MOE:mCHERRY 6872 bp

| Name | Part | Start | End |
|---|---|---|---|
| MOE promoter | Promoter | 388 | 1,481 |
| PAC | CDS | 1,486 | 2,082 |
| MOE term | 3'UTR | 2,083 | 2,373 |
| MOE promoter | Promoter | 2,378 | 3,478 |
| MOE | CDS | 3,479 | 3,884 |
| mCherry | CDS | 3,885 | 4,598 |
| MOE term | 3'UTR | 4,599 | 4,888 |

AMP CDS 5,884 6,744

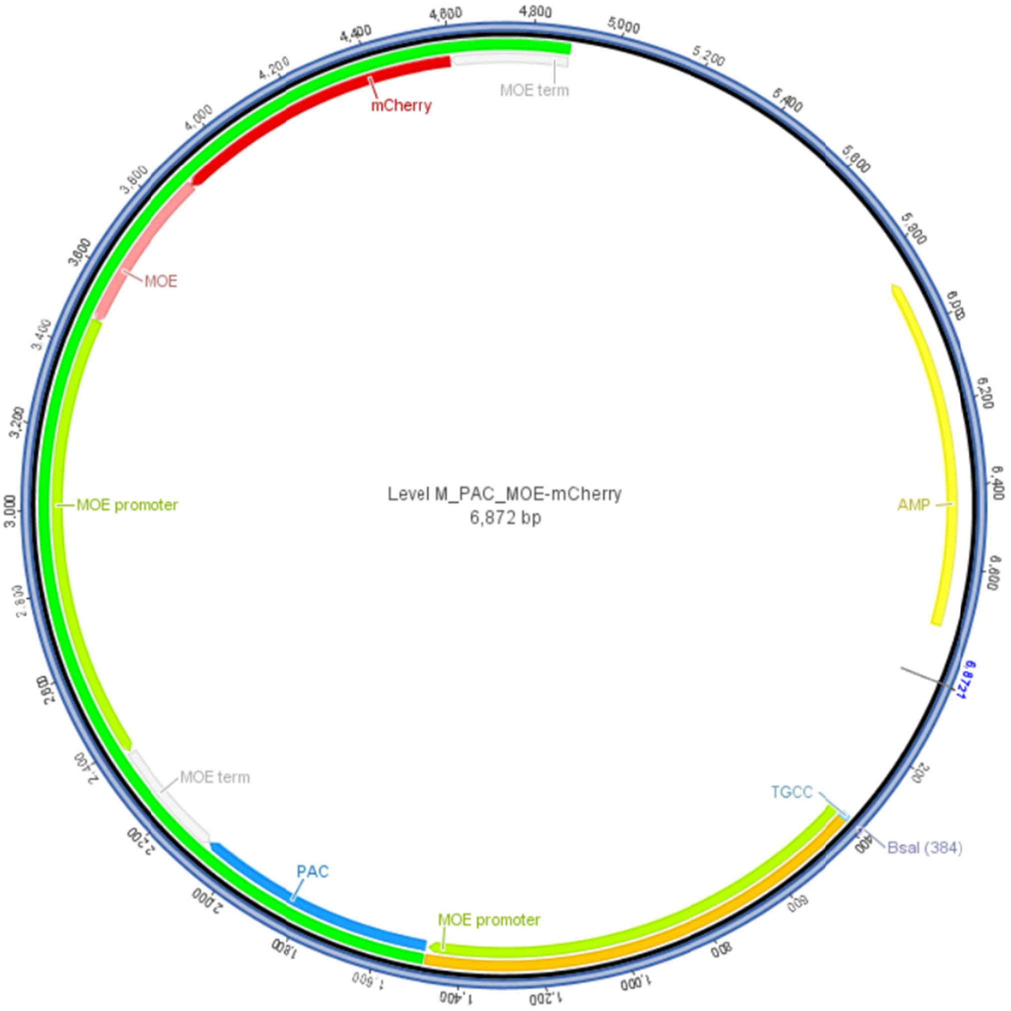


> LevelM:PAC:MOE:mCHERRY

CTAAATTGTAAGCGTTAATATTTTGTTAAAATTCGCGTTAAATTTTTGTTAAATCAGCTCATTTTTTAACCAATAG
GCCGAAATCGGCAAAATCCCTTATAAATCAAAAGAATAGACCGAGATAGGGTTGAGTGGCCGCTACAGGGCGCTCC
CATTCGCCATTCAGGCTGCGCAACTGTTGGGAAGGGCGTTTCGGTGCGGGCCTCTTCGCTATTACGCCAGCTGGCG
AAAGGGGGATGTGCTGCAAGGCGATTAAGTTGGGTAACGCCAGGGTTTTCCCAGTCACGACGTTGTAAAACGACGG
CCAGTGAGCGCGACGTAATACGACTCACTATAGGGCGAATTGGCGGAAGGCCGTCAAGGCCTAGGCGCGCCAGGTC
TCATGCCCTCGTAATGAGCCCAACCATTATCGGATCGGTCTTGCGGGCGAACGAACATCTGCGGCAGGTGAGAGGT
GGGAGTTTCCGGATGCGGTCCGAGTCACCGGCGATGGAACTCCCCACAGGCCAGTGTGACGGTGAGCACTGATGTT
CGATGGAAACCAAAGTGGCAGGGTTCGTACTTCCCAGCCATGGTTGCGCCGTTTGTAATTGATGTGATGGCAGAGC
TGACCGGTAGCAGTAGAAGGTGCGTTGATTCAGCTCAACAGCTCTCTTGTGTTAGCAGAAAGTAGAAGCAGATCTA
GATGATTCAGGAATCCCTCATCTTAGCGGTAGTATGTTACTGTGCACCCATCATGCCATGGATGATCCGGATCCCT
TGTTGGGTGAGGTAGGTTCCTTGGTGAGTTGGATGATTGTGGATCTGCCGATGGCAGAGATCATGGCTCCATTCCC
CATCGGGACCTCACGGTATCACTGATCATTTCATGGCTCAAGGGCGGGCTTGTCGACTTCGACAAGTGGGTGGGAC
TGCGGTGAGCAAGGTGCTTTGCACTTGCCGAACCCATCGCTTTGCTCCGGCTTGTAGTGAACTAACCGGAGCAAGA
ATAGGGCCCGGGCTAGTGGGACGCAACACATGAGGAACGTGTCTCAAATCCATAGATTTTATGTTGGATTCGTCTC
GTCGGAGCCAAGTGCTCCACGATGTGCATCCCTCGAGTCGAATCTTCCACTAGAACCCTTGAGCGTCAGCGAGCTA
GAGAACTTAGGAGCGGAGGGACGGGTTCTGCTAGTAGACATCAACAGATATCCCTTCACGCTCCCTTCATTGTATG
CGTGAGTATGTATCGCGGAGCAACACAGTCTTGTCCATCAAAGGGCATTTCTCCACGAATCTCGCACTAGCTTGGA

AATCATAATCTCGAGTCACTTGATTTCGTAACGCTGAATCCTTCTAAATGGACTCTATGTTGTAAGGCGAGACGCT
ATAACGATGTCTATCGACCGGACGCATTGAACGCATACACATAGCTAAGTGAGTTGTGTGGAACTCTTGATGAACT
CGCGTCACTCTTGAATCTCTATCCCGCTGTTCAACGCAATGACCGAGTACAAGCCCACGGTGCGCCTCGCCACCCG
CGACGACGTCCCCCGGGCCGTACGCACCCTCGCCGCCGCGTTCGCCGACTACCCCGCCACGCGCCACACCGTCGAC
CCGGACCGCCACATCGAGCGGGTCACCGAGCTGCAAGAACTCTTCCTCACGCGCGTCGGGCTCGACATCGGCAAGG
TGTGGGTCGCGGACGACGGCGCCGCGGTGGCGGTCTGGACCACGCCGGAGAGCGTCGAAGCGGGGGCGGTGTTCGC
CGAGATCGGCCCGCGCATGGCCGAGTTGAGCGGTTCCCGGCTGGCCGCGCAGCAACAGATGGAAGGCCTCCTGGCG
CCGCACCGGCCCAAGGAGCCCGCGTGGTTCCTGGCCACCGTCGGCGTCTCGCCCGACCACCAGGGCAAGGGTCTGG
GCAGCGCCGTCGTGCTCCCCGGAGTGGAGGCGGCCGAGCGCGCCGGGGTGCCCGCCTTCCTGGAGACTTCCGCGCC
CCGCAACCTCCCCTTCTACGAGCGGCTCGGCTTCACCGTCACCGCCGACGTCGAGGTGCCCGAAGGACCGCGCACC
TGGTGCATGACCCGCAAGCCCGGTGCCTGAGCTTGACACACATCTAGTTGAGATCTACATCAATGTTCGAAATCAA
CCATGCTTATAGCCATTGATATCTCTCTACTCTTTTCCGACCCTTATTACAAGGTCTATGACCAAACTATTGAGCC
TACGGTTTGTGACTCGCCAGGTTAAGCGATGTCACATCGCAAGTCGTTGAGGATGTACTGGTTATGAATGGCTCTT
CTTTCTTTCTGTGACTCGTTGGCATTGTTAGCGGATCTACAAGTAACCGTCGACGATCACGAGAGCTTCCGGCTCC
ATGATTGGGTCATCGCTGCAAGAGGCTCGTAATGAGCCCAACCATTATCGGATCGGTCTTGCGGGCGAACGAACAT
CTGCGGCAGGTGAGAGGTGGGAGTTTCCGGATGCGGTCCGAGTCACCGGCGATGGAACTCCCCACAGGCCAGTGTG
ACGGTGAGCACTGATGTTCGATGGAAACCAAAGTGGCAGGGTTCGTACTTCCCAGCCATGGTTGCGCCGTTTGTAA
TTGATGTGATGGCAGAGCTGACCGGTAGCAGTAGAAGGTGCGTTGATTCAGCTCAACAGCTCTCTTGTGTTAGCAG
AAAGTAGAAGCAGATCTAGATGATTCAGGAATCCCTCATCTTAGCGGTAGTATGTTACTGTGCACCCATCATGCCA
TGGATGATCCGGATCCCTTGTTGGGTGAGGTAGGTTCCTTGGTGAGTTGGATGATTGTGGATCTGCCGATGGCAGA
GATCATGGCTCCATTCCCCATCGGGACCTCACGGTATCACTGATCATTTCATGGCTCAAGGGCGGGCTTGTCGACT
TCGACAAGTGGGTGGGACTGCGGTGAGCAAGGTGCTTTGCACTTGCCGAACCCATCGCTTTGCTCCGGCTTGTAGT
GAACTAACCGGAGCAAGAATAGGGCCCGGGCTAGTGGGACGCAACACATGAGGAACGTGTCTCAAATCCATAGATT
TTATGTTGGATTCGTCTCGTCGGAGCCAAGTGCTCCACGATGTGCATCCCTCGAGTCGAATCTTCCACTAGAACCC
TTGAGCGTCAGCGAGCTAGAGAACTTAGGAGCGGAGGGACGGGTTCTGCTAGTAGACATCAACAGATATCCCTTCA
CGCTCCCTTCATTGTATGCGTGAGTATGTATCGCGGAGCAACACAGTCTTGTCCATCAAAGGGCATTTCTCCACGA
ATCTCGCACTAGCTTGGAAATCATAATCTCGAGTCACTTGATTTCGTAACGCTGAATCCTTCTAAATGGACTCTAT
GTTGTAAGGCGAGACGCTATAACGATGTCTATCGACCGGACGCATTGAACGCATACACATAGCTAAGTGAGTTGTG
TGGAACTCTTGATGAACTCGCGTCACTCTTGAATCTCTATCCCGCTGTTCAACGCAATAATGCGCTTCATTGTTGG
TCTGTACTCCTGCCTGGCCGTTTTGGTTCTTGGGCAGTCCTCTTGTCCCACAGGGGATGCTCAGTGTCACAGTGAG
AGTCCCGGTTCGTACTGCAAGTTCTATCAAGGTCACAATGTCTGCCAGGGAACTGACACCCCCTGCTCTTGCACCA
CTACTCCAGCACCGCCGCCACCCACTCCCACTGGTGGATCATGTCCTGATGGTGATGCTTTTTGTCAGAAGCAGTC
TGGTAATTCCGGCTCCTATTGCAAGGCTGCTTACCAGACTGGTCCCGGTGGAGGTGTTTGCCAGGGTACTGATATC
CCCTGTAAATGTGGTGGTGGCGGATCAACTACGATGAAGCCGACTACGACGAAGCCGACTACGACGAAGCCGACAT
GTCCTACTTCGATGGTGAGCAAGGGCGAGGAGGATAACATGGCCATCATCAAGGAGTTCATGCGCTTCAAGGTGCA
CATGGAGGGCTCCGTGAACGGCCACGAGTTCGAGATCGAGGGCGAGGGCGAGGGCCGCCCCTACGAGGGCACCCAG
ACCGCCAAGCTGAAGGTGACCAAGGGTGGCCCCCTGCCCTTCGCCTGGGACATCCTGTCCCCTCAGTTCATGTACG
GCTCCAAGGCCTACGTGAAGCACCCCGCCGACATCCCCGACTACTTGAAGCTGTCCTTCCCCGAGGGCTTCAAGTG
GGAGCGCGTGATGAACTTCGAGGACGGCGGCGTGGTGACCGTGACCCAGGACTCCTCCCTGCAGGACGGCGAGTTC
ATCTACAAGGTGAAGCTGCGCGGCACCAACTTCCCCTCCGACGGCCCCGTAATGCAGAAGAAAACCATGGGCTGGG
AGGCCTCCTCCGAGCGGATGTACCCCGAGGACGGCGCCCTGAAGGGCGAGATCAAGCAGAGGCTGAAGCTGAAGGA
CGGCGGCCACTACGACGCTGAGGTCAAGACCACCTACAAGGCCAAGAAGCCCGTGCAGCTGCCCGGCGCCTACAAC
GTCAACATCAAGTTGGACATCACCTCCCACAACGAGGACTACACCATCGTGGAACAGTACGAACGCGCCGAGGGCC
GCCACTCCACCGGCGGCATGGACGAGCTGTACAAGTGAGCTTGACACACATCTAGTTGAGATCTACATCAATGTTC
GAAATCAACCATGCTTATAGCCATTGATATCTCTCTACTCTTTTCCGACCCTTATTACAAGGTCTATGACCAAACT
ATTGAGCCTACGGTTTGTGACTCGCCAGGTTAAGCGATGTCACATCGCAAGTCGTTGAGGATGTACTGGTTATGAA
TGGCTCTTCTTTCTTTCTGTGACTCGTTGGCATTGTTAGCGGATCTACAAGTAACCGTCGACGATCACGAGAGCTT
CCGGCTCCATGATTGGGTCAGCAAGGGAATTAATTAACTGGCCTCATGGGCCTTCCGCTCACTGCCCGCTTTCCAG
TCGGGAAACCTGTCGTGCCAGCTGCATTAACATGGTCATAGCTGTTTCCTTGCGTATTGGGCGCTCTCCGCTTCCT
CGCTCACTGACTCGCTGCGCTCGGTCGTTCGGGTAAAGCCTGGGGTGCCTAATGAGCAAAAGGCCAGCAAAAGGCC
AGGAACCGTAAAAAGGCCGCGTTGCTGGCGTTTTTCCATAGGCTCCGCCCCCCTGACGAGCATCACAAAAATCGAC
GCTCAAGTCAGAGGTGGCGAAACCCGACAGGACTATAAAGATACCAGGCGTTTCCCCCTGGAAGCTCCCTCGTGCG
CTCTCCTGTTCCGACCCTGCCGCTTACCGGATACCTGTCCGCCTTTCTCCCTTCGGGAAGCGTGGCGCTTTCTCAT
AGCTCACGCTGTAGGTATCTCAGTTCGGTGTAGGTCGTTCGCTCCAAGCTGGGCTGTGTGCACGAACCCCCCGTTC
AGCCCGACCGCTGCGCCTTATCCGGTAACTATCGTCTTGAGTCCAACCCGGTAAGACACGACTTATCGCCACTGGC
AGCAGCCACTGGTAACAGGATTAGCAGAGCGAGGTATGTAGGCGGTGCTACAGAGTTCTTGAAGTGGTGGCCTAAC
TACGGCTACACTAGAAGAACAGTATTTGGTATCTGCGCTCTGCTGAAGCCAGTTACCTTCGGAAAAAGAGTTGGTA
GCTCTTGATCCGGCAAACAAACCACCGCTGGTAGCGGTGGTTTTTTTGTTTGCAAGCAGCAGATTACGCGCAGAAA
AAAAGGATCTCAAGAAGATCCTTTGATCTTTTCTACGGGGTCTGACGCTCAGTGGAACGAAAACTCACGTTAAGGG
ATTTTGGTCATGAGATTATCAAAAAGGATCTTCACCTAGATCCTTTTAAATTAAAAATGAAGTTTTAAATCAATCT
AAAGTATATATGAGTAAACTTGGTCTGACAGTTACCAATGCTTAATCAGTGAGGCACCTATCTCAGCGATCTGTCT
ATTTCGTTCATCCATAGTTGCCTGACTCCCCGTCGTGTAGATAACTACGATACGGGAGGGCTTACCATCTGGCCCC
AGTGCTGCAATGATACCGCGAGAACCACGCTCACCGGCTCCAGATTTATCAGCAATAAACCAGCCAGCCGGAAGGG
CCGAGCGCAGAAGTGGTCCTGCAACTTTATCCGCCTCCATCCAGTCTATTAATTGTTGCCGGGAAGCTAGAGTAAG
TAGTTCGCCAGTTAATAGTTTGCGCAACGTTGTTGCCATTGCTACAGGCATCGTGGTGTCACGCTCGTCGTTTGGT
ATGGCTTCATTCAGCTCCGGTTCCCAACGATCAAGGCGAGTTACATGATCCCCCATGTTGTGCAAAAAAGCGGTTA
GCTCCTTCGGTCCTCCGATCGTTGTCAGAAGTAAGTTGGCCGCAGTGTTATCACTCATGGTTATGGCAGCACTGCA
TAATTCTCTTACTGTCATGCCATCCGTAAGATGCTTTTCTGTGACTGGTGAGTACTCAACCAAGTCATTCTGAGAA

TAGTGTATGCGGCGACCGAGTTGCTCTTGCCCGGCGTCAATACGGGATAATACCGCGCCACATAGCAGAACTTTAA
AAGTGCTCATCATTGGAAAACGTTCTTCGGGGCGAAAACTCTCAAGGATCTTACCGCTGTTGAGATCCAGTTCGAT
GTAACCCACTCGTGCACCCAACTGATCTTCAGCATCTTTTACTTTCACCAGCGTTTCTGGGTGAGCAAAAACAGGA
AGGCAAAATGCCGCAAAAAAGGGAATAAGGGCGACACGGAAATGTTGAATACTCATACTCTTCCTTTTTCAATATT
ATTGAAGCATTTATCAGGGTTATTGTCTCATGAGCGGATACATATTTGAATGTATTTAGAAAAATAAACAAATAGG
GGTTCCGCGCACATTTCCCCGAAAAGTGCCAC

#### *Oxyrrhis marina*

```
>Om-mCherry-hsp90 OmHSP90-HpaI-pUC57 with mCherry
TCGCGCGTTTCGGTGATGACGGTGAAAACCTCTGACACATGCAGCTCCCGGAGACGGTCACAGCTTGTCTGTAAGCGGATGCCGGG
AGCAGACAAGCCCGTCAGGGCGCGTCAGCGGGTGTTGGCGGGTGTCGGGGCTGGCTTAACTATGCGGCATCAGAGCAGATTGTACT
GAGAGTGCACCATATGCGGTGTGAAATACCGCACAGATGCGTAAGGAGAAAATACCGCATCAGGCGCCATTCGCCATTCAGGCTGC
GCAACTGTTGGGAAGGGCGATCGGTGCGGGCCTCTTCGCTATTACGCCAGCTGGCGAAAGGGGGATGTGCTGCAAGGCGATTAAGT
TGGGTAACGCCAGGGTTTTCCCAGTCACGACGTTGTAAAACGACGGCCAGTGAATTCGAGCTCGGTACCTCGCGAATGCATCTAGA
TGGAGGAGACTTGCGTTGTCCTGGTCCATTGCGCGCAGCGCATCGCATCTTCGCAGTGGTGCATTGACATGGCAGCAGCTTCACCA
TCGCTTTTCATTTCTGACGAGGGCAGTTTTTCGTTGGATGGGTTCGGTTTCATGACGGTTGATGGGCTTGGTTTTGAGTTTTGTCT
TTGGAGTGTACGTTCCACGGCTTGAGTTTGCGAAGCGTTTAATACGACTCACTATAGGGCCGTAGCCATTTTGGCTCAAGCTTGGT
TTTGAGTTTTGTCTTTGGAGTGTACGTTCCACGGCTTGAGTTTGCGAAGCATGGTGTCGAAAGGTGAGGAAGATAATATGGCTATC
ATTAAGGAGTTCATGCGCTTTAAGGTACATATGGAGGGTAGCGTCAATGGACACGAGTTCGAAATTGAAGGCGAGGGGGAAGGCAG
GCCATACGAGGGTACACAAACGGCCAAGCTTAAGGTTACTAAGGGGGGTCCTCTCCCGTTTGCTTGGGATATCTTGTCTCCCCAGT
TCATGTATGGGTCGAAGGCTTACGTCAAGCACCCCGCCGATATCCCCGACTATCTTAAGCTGAGCTTCCCGGAGGGTTTCAACTGG
GAGCGGGTGATGAACTTCGAGGACGGGGGCGTCGTTACTGTGACCCAGGATAGCAGTTTGCAGGACGGGGAGTTCATCTACAAGGT
GAAGCTGCGTGGCACAAACTTTCCGTCTGATGGCCCTGTGATGCAGTGCCGTACCATGGGTTGGGAGGCATCTACCGAGCGAATGT
ACCCGGAGGACGGGGCATTGAAGGGAGAGATCAAGCAGCGCCTGAAGTTGAAGGATGGTGGACACTACGATGCTGAGGTGAAGACG
ACGTATAAGGCGAAGAAGCCCGTGCAGCTGCCGGGGGCGTACAACGTGGACATTAAGTTGGACATCCTGTCCCACAACGAGGACTA
CACGATCGTGGAGCAGTACGAGCGAGCGGAGGGGCGCCACTCCACCGGCGGAATGGACGAGTTGTACAAGTAGGATTGTGCAGAGC
TTGAGTGTCCAAAGACATTTTGCCCTGCGCTTCTATAGTGTCACCTAAATAACGATTGTGCAGAGCTTGAGTGTCCAAAGACATTT
TGCCCTGTGAATTGGAGATGCATGTAGGCTGTCGTCCCAGTAGGCCAGCGCAAGGAGCACAGCGCACCATCACGTGAGCTCCTGTG
GCGTGTGCTGCGGCGAACTCTGGGGAGAGTGGTGGCGTGCATGGTTTTTTTTTTTGGATGTCGCGCGTGAGCAAGCCAGGAGGCACT
GGCACCGGAAGATAGCCACGGAGATCGGATCCCGGGCCCGTCGACTGCAGAGGCCTGCATGCAAGCTTGGCGTAATCATGGTCATA
GCTGTTTCCTGTGTGAAATTGTTATCCGCTCACAATTCCACACAACATACGAGCCGGAAGCATAAAGTGTAAAGCCTGGGGTGCCT
AATGAGTGAGCTAACTCACATTAATTGCGTTGCGCTCACTGCCCGCTTTCCAGTCGGGAAACCTGTCGTGCCAGCTGCATTAATGA
ATCGGCCAACGCGCGGGGAGAGGCGGTTTGCGTATTGGGCGCTCTTCCGCTTCCTCGCTCACTGACTCGCTGCGCTCGGTCGTTCG
GCTGCGGCGAGCGGTATCAGCTCACTCAAAGGCGGTAATACGGTTATCCACAGAATCAGGGGATAACGCAGGAAAGAACATGTGAG
CAAAAGGCCAGCAAAAGGCCAGGAACCGTAAAAAGGCCGCGTTGCTGGCGTTTTTCCATAGGCTCCGCCCCCCTGACGAGCATCAC
AAAAATCGACGCTCAAGTCAGAGGTGGCGAAACCCGACAGGACTATAAAGATACCAGGCGTTTCCCCCTGGAAGCTCCCTCGTGCG
CTCTCCTGTTCCGACCCTGCCGCTTACCGGATACCTGTCCGCCTTTCTCCCTTCGGGAAGCGTGGCGCTTTCTCATAGCTCACGCT
GTAGGTATCTCAGTTCGGTGTAGGTCGTTCGCTCCAAGCTGGGCTGTGTGCACGAACCCCCCGTTCAGCCCGACCGCTGCGCCTTA
TCCGGTAACTATCGTCTTGAGTCCAACCCGGTAAGACACGACTTATCGCCACTGGCAGCAGCCACTGGTAACAGGATTAGCAGAGC
GAGGTATGTAGGCGGTGCTACAGAGTTCTTGAAGTGGTGGCCTAACTACGGCTACACTAGAAGAACAGTATTTGGTATCTGCGCTC
TGCTGAAGCCAGTTACCTTCGGAAAAAGAGTTGGTAGCTCTTGATCCGGCAAACAAACCACCGCTGGTAGCGGTGGTTTTTTTGTT
TGCAAGCAGCAGATTACGCGCAGAAAAAAAGGATCTCAAGAAGATCCTTTGATCTTTTCTACGGGGTCTGACGCTCAGTGGAACGA
AAACTCACGTTAAGGGATTTTGGTCATGAGATTATCAAAAAGGATCTTCACCTAGATCCTTTTAAATTAAAAATGAAGTTTTAAAT
CAATCTAAAGTATATATGAGTAAACTTGGTCTGACAGTTACCAATGCTTAATCAGTGAGGCACCTATCTCAGCGATCTGTCTATTT
CGTTCATCCATAGTTGCCTGACTCCCCGTCGTGTAGATAACTACGATACGGGAGGGCTTACCATCTGGCCCCAGTGCTGCAATGAT
ACCGCGAGACCCACGCTCACCGGCTCCAGATTTATCAGCAATAAACCAGCCAGCCGGAAGGGCCGAGCGCAGAAGTGGTCCTGCAA
CTTTATCCGCCTCCATCCAGTCTATTAATTGTTGCCGGGAAGCTAGAGTAAGTAGTTCGCCAGTTAATAGTTTGCGCAACGTTGTT
GCCATTGCTACAGGCATCGTGGTGTCACGCTCGTCGTTTGGTATGGCTTCATTCAGCTCCGGTTCCCAACGATCAAGGCGAGTTAC
ATGATCCCCCATGTTGTGCAAAAAAGCGGTTAGCTCCTTCGGTCCTCCGATCGTTGTCAGAAGTAAGTTGGCCGCAGTGTTATCAC
TCATGGTTATGGCAGCACTGCATAATTCTCTTACTGTCATGCCATCCGTAAGATGCTTTTCTGTGACTGGTGAGTACTCAACCAAG
TCATTCTGAGAATAGTGTATGCGGCGACCGAGTTGCTCTTGCCCGGCGTCAATACGGGATAATACCGCGCCACATAGCAGAACTTT
AAAAGTGCTCATCATTGGAAAACGTTCTTCGGGGCGAAAACTCTCAAGGATCTTACCGCTGTTGAGATCCAGTTCGATGTAACCCA
CTCGTGCACCCAACTGATCTTCAGCATCTTTTACTTTCACCAGCGTTTCTGGGTGAGCAAAAACAGGAAGGCAAAATGCCGCAAAA
AAGGGAATAAGGGCGACACGGAAATGTTGAATACTCATACTCTTCCTTTTTCAATATTATTGAAGCATTTATCAGGGTTATTGTCT
CATGAGCGGATACATATTTGAATGTATTTAGAAAAATAAACAAATAGGGGTTCCGCGCACATTTCCCCGAAAAGTGCCACCTGACG
TCTAAGAAACCATTATTATCATGACATTAACCTATAAAAATAGGCGTATCACGAGGCCCTTTCGTC
```

*Hematodinium sp.*

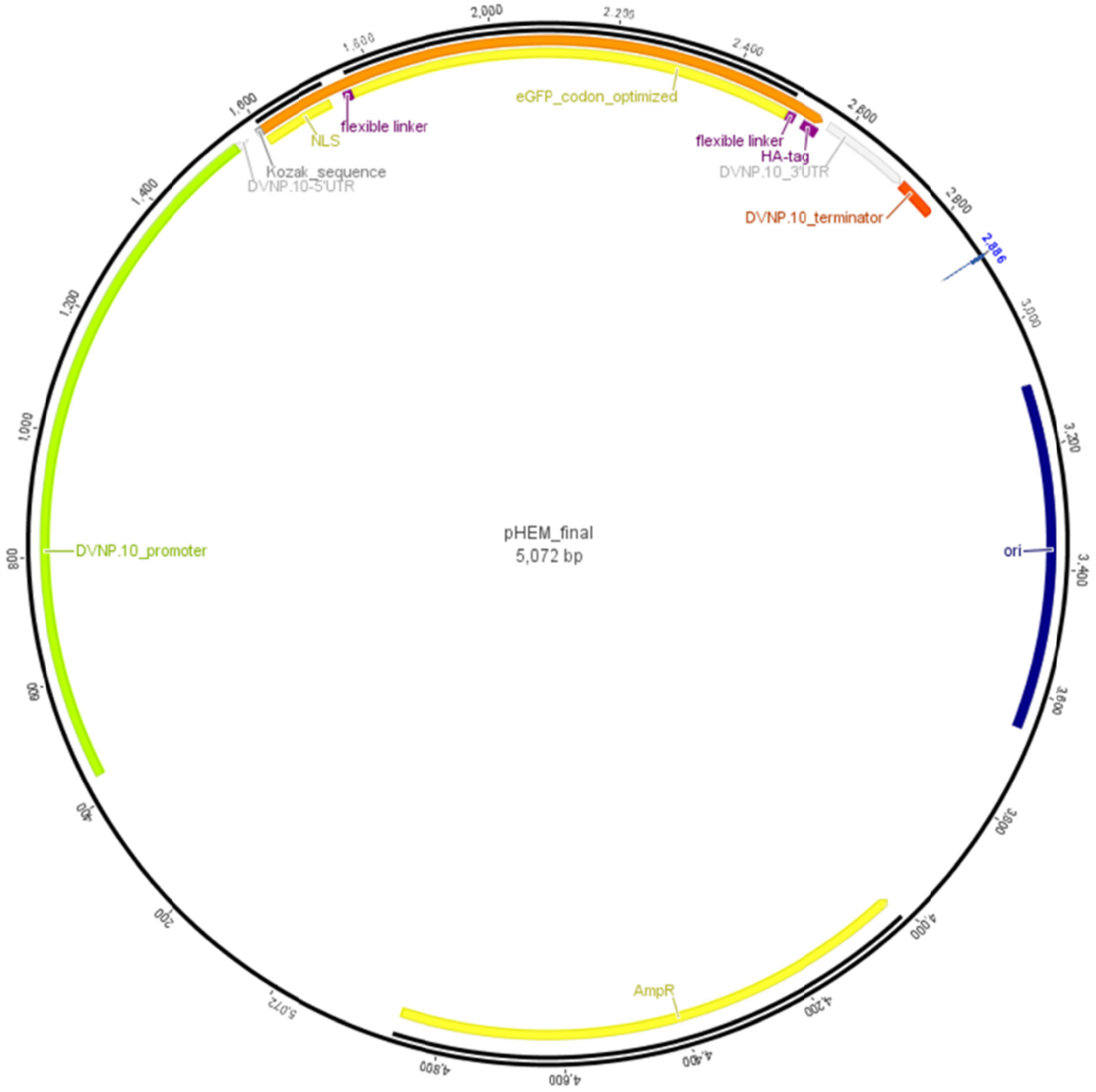


| Name | Type | Start | Stop |
|---|---|---|---|
| ori | site | 3102 | 3659 |
| NLS | CDS | 1602 | 1721 |
| Kozak_sequence | misc_feature | 1598 | 1601 |
| HA-tag | motif | 2529 | 2558 |
| flexible linker | motif | 1746 | 1760 |
| flexible linker | motif | 2499 | 2513 |
| eGFP_codon_optimized | CDS | 1761 | 2498 |
| DVNP.10_terminator | terminator | 2719 | 2784 |
| DVNP.10_promoter | promoter | 440 | 1553 |
| DVNP.10_3'UTR | 3'UTR | 257 | 2718 |

| | | | |
|---|---|---|---|
| DVNP.10-5'UTR | 5'UTR | 1554 | 1578 |
| complete ORF | ORF | 1602 | 2558 |
| AmpR | CDS | 4012 | 4872 |

```
>pHem_final
TCGCGCGTTTCGGTGATGACGGTGAAAACCTCTGACACATGCAGCTCCCGGAGACGGTCACAGCTTGTCTGTAAGCGGATGCCGGG
AGCAGACAAGCCCGTCAGGGCGCGTCAGCGGGTGTTGGCGGGTGTCGGGGCTGGCTTAACTATGCGGCATCAGAGCAGATTGTACT
GAGAGTGCACCATATGCGGTGTGAAATACCGCACAGATGCGTAAGGAGAAAATACCGCATCAGGCGCCATTCGCCATTCAGGCTGC
GCAACTGTTGGGAAGGGCGATCGGTGCGGGCCTCTTCGCTATTACGCCAGCTGGCGAAAGGGGGATGTGCTGCAAGGCGATTAAGT
TGGGTAACGCCAGGGTTTTCCCAGTCACGACGTTGTAAAACGACGGCCAGTGAATTCGAGCTCGGTACCTCGCGAATGCATCTAGA
TGACCCAATGCGGCCGCGCAGTGAACGCTCAGATTTCCTCCTTCAATATTGCATTTGGCTGTTAATCTCTTTCTTTCATTGAGATC
ACGCTCGTTTTGTTGAGCGGATTGGGGATCAGCAAGTGTCTGACTAGTTAAACGCTTGCTCCCAAGTGTGTGCTAATGCGACGCTC
ACCGATCCCTGCCAGAGGCACATTCTATTGGATGTGCCTCCCCCCCCCGCGCTTCCGCGCTATTCATTACGCAAGCTATCTACCGA
AATTCATCAAATTATTGTATGCAGGTAACTTTTTGGTTCGACAATCAAACGGGATAGTGGGGGGGGAGATGATACGAAATGGAAGA
AGGAAAGAAAAACCAACACATCGTCGATCCCCCCCTGTATGTTGTTTAGGGCGCCACACGCTGAACTTGTCGCGATTCTACGATAA
TAATCCCATTGAGTAGCTATGGCTGCCCCCCCAAAACCAAAAAATATCATATTCCCGCAAAATTTCCCCGCAAGCACTTCGAGTAC
GGGTTTAAACGGGCCCTTCGGGGGGCCGTTTGGCGCGCGGCAAGCAAAACATGGGGGGCCCGTTTAAATCGTACACCGCGCCACGC
CCGATTTTGTTTTTTTTGCTCCATATCCACGATGTTATTGCACTGAGCGGCCCCCCTCGATGGGGGGGCCCCATATGGCGTGCGGC
AGCAGAAAGATAGAGCGGGGCCCGTTTAAACCCGTACTCGCAAGTCGCTTCGATTTTTTCCCTCGATGTCAGAAACGGGTATAAAT
GGTATACTTCCCTCAACTATACGAAGTTGTTCACTGTGCGGGCGGCGGAGCTTCAATATATACTGTGACTTCCTTGTCGTGTGTTC
CGCACCCCAGGCGTATCAATGCCCGCGCTTTCGCGATCCTCCCGCAAATTTCACGGTAAAATAGAAAACGCAACGAAATTCAACCG
TTGTTGAACAACGTTGTGCCAAGGTTCACCAATAATTACTTCATATTTACCCCGGATCTCGTATAGTGATGATACGAAATTCCTCT
TGCGTCGTTTCCAAGCTTCCTCAAAACGTCCCACTCACTCGCCCTCCGTAAATACATGAAATCCAACAGCATTTTGTGTGTTCTCT
TTCTCAGCGATTTTTGTTTTTCTGTTACGCAGATCTGTCCGCGGTGGCCAAACATGGCGATGAAGGGAATGAAAGCGATGAAAGTG
AAAAAGTCGGCAAAGAAGTCGGGGAAAGGCATGAAGAAAAAGGCGAAACGCGTCTCGAAAGTCGCGCGAGGAAAGCGCGCGAAATC
GCCTAGGAGTAGTTGGCCAGCGGCCGGCGGCTCGGGCTCGATGAGCAAGGGCGAGGAGCTGTTCACCGGCGTGGTGCCCATCCTGG
TCGAGCTGGACGGCGACGTAAACGGCCACAAGTTCAGCGTGTCCGGCGAGGGCGAGGGCGATGCCACCTACGGCAAGCTGACCCTG
AAGTTCATCTGCACCACCGGCAAGCTGCCCGTGCCCTGGCCCACCCTCGTGACCACCCTGACCTACGGCGTGCAGTGCTTCAGCCG
CTACCCCGACCACATGAAGCAGCACGACTTCTTCAAGTCCGCCATGCCCGAAGGCTACGTCCAGGAGCGCACCATCTTCTTCAAGG
ACGACGGCAACTACAAGACCCGCGCCGAGGTGAAGTTCGAGGGCGACACCCTGGTGAACCGCATCGAGCTGAAGGGCATCGACTTC
AAGGAGGACGGCAACATCCTGGGCCACAAGCTGGAGTACAACTACAACAGCCACAACGTCTATATCATGGCCGACAAGCAGAAGAA
CGGCATCAAGGTGAACTTCAAGATCCGCCACAACATCGAGGACGGCAGCGTGCAGCTCGCCGACCACTACCAGCAGAACACCCCCA
TCGGCGACGGCCCCGTGCTGCTGCCCGACAACCACTACCTGAGCACCCAGTCCGCCCTGAGCAAAGACCCCAACGAGAAGCGCGAT
CACATGGTCCTGCTGGAGTTCGTGACCGCCGCCGGAATCACCCTCGGCATGGACGAGCTGTACAAGCCCGGCGACGTCGATATCGA
GCTCGGCGGCTCGGGCAGCGCCGGCATCGATATTTACCCATACGATGTTCCAGATTACGCTTAACCATGGCATAATCAATTTCCTT
TCAGGCATTTGGGAAAATAAGTCGGCGAAATTGATTTTGTCCTGTTCCCCCCCCAGCCGCTGCGTGGGGTCTTTCCACGGCGGACC
TATTGGTTTGGATATAATTAGTGGAGGAAGTTGTATAATCAGAGTTAAATCCAGTATGCCTTCTTGATTAATGAACTCATAAATAA
TACCATGATTCTGTAAATCGTGCGACGAGTAGTATTGGGTCATCGGATCCCGGGCCCGTCGACTGCAGAGGCCTGCATGCAAGCTT
GGCGTAATCATGGTCATAGCTGTTTCCTGTGTGAAATTGTTATCCGCTCACAATTCCACACAACATACGAGCCGGAAGCATAAAGT
GTAAAGCCTGGGGTGCCTAATGAGTGAGCTAACTCACATTAATTGCGTTGCGCTCACTGCCCGCTTTCCAGTCGGGAAACCTGTCG
TGCCAGCTGCATTAATGAATCGGCCAACGCGCGGGGAGAGGCGGTTTGCGTATTGGGCGCTCTTCCGCTTCCTCGCTCACTGACTC
GCTGCGCTCGGTCGTTCGGCTGCGGCGAGCGGTATCAGCTCACTCAAAGGCGGTAATACGGTTATCCACAGAATCAGGGGATAACG
CAGGAAAGAACATGTGAGCAAAAGGCCAGCAAAAGGCCAGGAACCGTAAAAAGGCCGCGTTGCTGGCGTTTTTCCATAGGCTCCGC
CCCCCTGACGAGCATCACAAAAATCGACGCTCAAGTCAGAGGTGGCGAAACCCGACAGGACTATAAAGATACCAGGCGTTTCCCCC
TGGAAGCTCCCTCGTGCGCTCTCCTGTTCCGACCCTGCCGCTTACCGGATACCTGTCCGCCTTTCTCCCTTCGGGAAGCGTGGCGC
TTTCTCATAGCTCACGCTGTAGGTATCTCAGTTCGGTGTAGGTCGTTCGCTCCAAGCTGGGCTGTGTGCACGAACCCCCCGTTCAG
CCCGACCGCTGCGCCTTATCCGGTAACTATCGTCTTGAGTCCAACCCGGTAAGACACGACTTATCGCCACTGGCAGCAGCCACTGG
TAACAGGATTAGCAGAGCGAGGTATGTAGGCGGTGCTACAGAGTTCTTGAAGTGGTGGCCTAACTACGGCTACACTAGAAGAACAG
TATTTGGTATCTGCGCTCTGCTGAAGCCAGTTACCTTCGGAAAAAGAGTTGGTAGCTCTTGATCCGGCAAACAAACCACCGCTGGT
AGCGGTGGTTTTTTTGTTTGCAAGCAGCAGATTACGCGCAGAAAAAAAGGATCTCAAGAAGATCCTTTGATCTTTTCTACGGGGTC
TGACGCTCAGTGGAACGAAAACTCACGTTAAGGGATTTTGGTCATGAGATTATCAAAAAGGATCTTCACCTAGATCCTTTTAAATT
AAAAATGAAGTTTTAAATCAATCTAAAGTATATATGAGTAAACTTGGTCTGACAGTTACCAATGCTTAATCAGTGAGGCACCTATC
TCAGCGATCTGTCTATTTCGTTCATCCATAGTTGCCTGACTCCCCGTCGTGTAGATAACTACGATACGGGAGGGCTTACCATCTGG
CCCCAGTGCTGCAATGATACCGCGAGACCCACGCTCACCGGCTCCAGATTTATCAGCAATAAACCAGCCAGCCGGAAGGGCCGAGC
GCAGAAGTGGTCCTGCAACTTTATCCGCCTCCATCCAGTCTATTAATTGTTGCCGGGAAGCTAGAGTAAGTAGTTCGCCAGTTAAT
AGTTTGCGCAACGTTGTTGCCATTGCTACAGGCATCGTGGTGTCACGCTCGTCGTTTGGTATGGCTTCATTCAGCTCCGGTTCCCA
ACGATCAAGGCGAGTTACATGATCCCCCATGTTGTGCAAAAAAGCGGTTAGCTCCTTCGGTCCTCCGATCGTTGTCAGAAGTAAGT
TGGCCGCAGTGTTATCACTCATGGTTATGGCAGCACTGCATAATTCTCTTACTGTCATGCCATCCGTAAGATGCTTTTCTGTGACT
GGTGAGTACTCAACCAAGTCATTCTGAGAATAGTGTATGCGGCGACCGAGTTGCTCTTGCCCGGCGTCAATACGGGATAATACCGC
GCCACATAGCAGAACTTTAAAAGTGCTCATCATTGGAAAACGTTCTTCGGGGCGAAAACTCTCAAGGATCTTACCGCTGTTGAGAT
CCAGTTCGATGTAACCCACTCGTGCACCCAACTGATCTTCAGCATCTTTTACTTTCACCAGCGTTTCTGGGTGAGCAAAAACAGGA
AGGCAAAATGCCGCAAAAAAGGGAATAAGGGCGACACGGAAATGTTGAATACTCATACTCTTCCTTTTTCAATATTATTGAAGCAT
TTATCAGGGTTATTGTCTCATGAGCGGATACATATTTGAATGTATTTAGAAAAATAAACAAATAGGGGTTCCGCGCACATTTCCCC
GAAAAGTGCCACCTGACGTCTAAGAAACCATTATTATCATGACATTAACCTATAAAAATAGGCGTATCACGAGGCCCTTTCGTC
```

### *Fugacium (Symbiodinium) kawagutii and Alexandrium catenella*

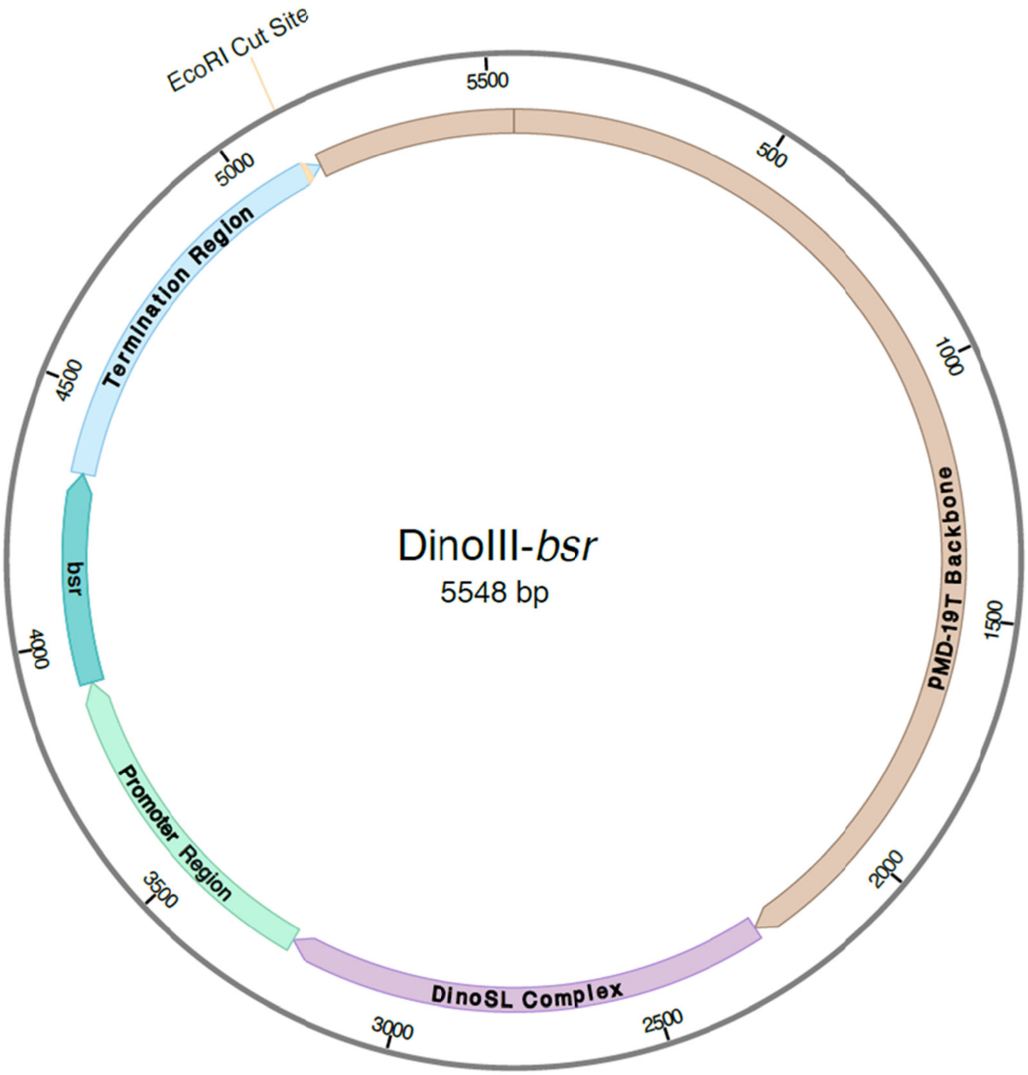


---

DinoIII-*bsr* sequence, containing the pMD™19 T-Vector (shown in lowercase; Takara, Japan), the RNA Elements (highlighted in yellow), the Promoter Region (highlighted in blue), the *bsr* gene (highlighted in teal), and the Termination Region (highlighted in pink).

gacgaaagggcctcgtgatacgcctatttttataggttaatgtcatgataataatggtttcttagacgtcaggtggcacttttcgg
ggaaatgtgcgcggaacccctatttgtttatttttctaaatacattcaaatatgtatccgctcatgagacaataaccctgataaat
gcttcaataatattgaaaaaggaagagtatgagtattcaacatttccgtgtcgcccttattcccttttttgcggcattttgccttc
ctgtttttgctcacccagaaacgctggtgaaagtaaaagatgctgaagatcagttgggtgcacgagtgggttacatcgaactggat
ctcaacagcggtaagatccttgagagttttcgccccgaagaacgttttccaatgatgagcacttttaaagttctgctatgtggcgc
ggtattatcccgtattgacgccgggcaagagcaactcggtcgccgcatacactattctcagaatgacttggttgagtactcaccag
tcacagaaaagcatcttacggatggcatgacagtaagagaattatgcagtgctgccataaccatgagtgataacactgcggccaac
ttacttctgacaacgatcggaggaccgaaggagctaaccgcttttttgcacaacatgggggatcatgtaactcgccttgatcgttg
ggaaccggagctgaatgaagccataccaaacgacgagcgtgacaccacgatgcctgtagcaatggcaacaacgttgcgcaaactat
taactggcgaactacttactctagcttcccggcaacaattaatagactggatggaggcggataaagttgcaggaccacttctgcgc
tcggcccttccggctggctggtttattgctgataaatctggagccggtgagcgtgggtctcgcggtatcattgcagcactggggcc
agatggtaagccctcccgtatcgtagttatctacacgacggggagtcaggcaactatggatgaacgaaatagacagatcgctgaga
taggtgcctcactgattaagcattggtaactgtcagaccaagtttactcatatatactttagattgatttaaaacttcatttttaa
tttaaaaggatctaggtgaagatcctttttgataatctcatgaccaaaatcccttaacgtgagttttcgttccactgagcgtcaga
ccccgtagaaaagatcaaaggatcttcttgagatcctttttttctgcgcgtaatctgctgcttgcaaacaaaaaaaccaccgctac
cagcggtggtttgtttgccggatcaagagctaccaactctttttccgaaggtaactggcttcagcagagcgcagataccaaatact

gttcttctagtgtagccgtagttaggccaccacttcaagaactctgtagcaccgcctacatacctcgctctgctaatcctgttacc
agtggctgctgccagtggcgataagtcgtgtcttaccgggttggactcaagacgatagttaccggataaggcgcagcggtcgggct
gaacggggggttcgtgcacacagcccagcttggagcgaacgacctacaccgaactgagatacctacagcgtgagctatgagaaagc
gccacgcttcccgaagggagaaaggcggacaggtatccggtaagcggcagggtcggaacaggagagcgcacgagggagcttccagg
gggaaacgcctggtatctttatagtcctgtcgggtttcgccacctctgacttgagcgtcgatttttgtgatgctcgtcaggggggc
ggagcctatggaaaaacgccagcaacgcggcctttttacggttcctggccttttgctggccttttgctcacatgttctttcctgcg
ttatcccctgattctgtggataaccgtattaccgcctttgagtgagctgataccgctcgccgcagccgaacgaccgagcgcagcga
gtcagtgagcgaggaagcggaagagcgcccaatacgcaaaccgcctctccccgcgcgttggccgattcattaatgcagctggcacg
acaggtttcccgactggaaagcgggcagtgagcgcaacgcaattaatgtgagttagctcactcattaggcaccccaggctttacac
tttatgcttccggctcgtatgttgtgtggaattgtgagcggataacaatttcacacaggaaacagctatgaccatgattacgccaa
gcttgcatgcctgcaggtcgacgattCCCGTAGCCATTTTGGCTCAAGGTACAAGTCGGGCTGATGCGGTCACGCAGGCCTCTTTT
GTATCAGATAAGACAGCAGACGTACATGATATAAATATTTATTATCGCGATGCTCCATCTGCAAAATTCCACTTGGCAGAGAAGAT
TTTCAATGCATGAAAACCTCGCAGAACCAAGCACCAATGTCAGATCATGTATGAGTCCTAGGCGCCAGTAAGTCTTTCAATTGGGT
GATGGCCACATTACCTGTTCTCTTCCTGAGGGATGCAGAATGTGGTGTGAGTGGTTGCCGAATGCTGTGCTGAACATGTAGATGGG
CTGCCCCAGCCAAGTGGATAACCTACAAGGACTGCAGGATGTAGATCCAGCAGTCCTGGTAGATAATGATTCAGACACCGGCTCAG
GCTGGCAACAGAGCAGGCAAACCTGAACACTATCGCGCAGTGGTGGGAATACAGCCCAGAATGGGAAGCAACCTGGCCTCTTCAGC
CAGCCATGACCACAAGCAACTCTGCAGTTTTGCAACACGAAATCTTTCAGGGTCTGCTCCCTGTGCTTACACATATTCAAAGATAT
TCTTCTGCGCATTTCAATGCAAATTCTTGGCGATGATTCGCTTCCGCAGGTATAGTGTTACTAGTATATAAGTCTAATCATATACT
TCAATTTATTAGTAATATTGTGAATTTCTAGGCAAGAAGGATGCCTGGACTCTGAATTATTATGGCATTGAGTAGAATCTGGATCT
TGATGATTATGCATTAATATCTTGAAATGCATTTGGATTCCCTTCGGGGATCATCCGTTAAAATTGGAACGATACAGAGAAGATTA
GCATGGCCCCTGCGCAAGGATGACACGCACAAATCGAGAAGTGTAAACAATTTTTTTGAAATTAATTGCCACTTTATTTTGAATAC
CTGAATATGCAGGTGAAGTAGTATAAGGTATTCATTATCATTGCGATTTTCTAGTAAGTCCCGTGGCTGTCATATCTAGTAACCCT
CACCTGGCAGGTGGGGAAAAGGCGAAGACAAATAGAATCAAATAGAATGTATGTGCTGACGTAGGGCTACATTACCTGAGGCTTGA
GGAACCTGTGGGTATGAATGCTACTGTTGGCAGCATCCAGTAGGTTCAGAAACACACGTCCCCACGTAATTTTTTGCATGTCTAAA
TGCACTGTGATGATTGATGATGATTGATTCCTTGAAGGCCATGGTGGGTTACTCTGTGGCCTCTTGCCACTGACTTCCAGGAAAAC
GCGGATTTTCTGACCATCACAAGGGACCACTACGAGAAGTATGCAGCGATCTCAGCAGAGTTCATTTACTGACCATTCTCAGGAAG
ATCGCTCGGGATCACTTGGGTAATATTGCTTTCTTCGCAGATGTTTTTTCCCTATTAAGCTTTTTGAGTCCCCTCATCGAGTGCCG
CATAAGTTTTTGCGTTGACATTGGCGGCAAAAAGTAAGACTAAACGATAGTTGCTTCAAGCACGCTTCTCAATCAACATTTTTTCA
AATCAAATGGGCACAGGCAGGTACCATCCAGTCCCGCAGAAGTGCTTATATATATATATATTATACACTCACGTGAGTGTGTTCAA
TGCAGGTCTGTTCCTCAAGTGCTTAGACCAAATTTCGGAGCCtctagtATGAAGACCTTCAACATCTCCCAGCAGGATCTAGAATT
AGTAGAAGTAGCGACAGAGAAGATTACAATGCTTTATGAGGATAATAAACATCATGTGGGAGCGGCAATTCGTACGAAAACAGGAG
AAATCATTTCGGCAGTACATATTGAAGCGTATATAGGACGAGTAACTGTTTGTGCAGAAGCCATTGCGATTGGTAGTGCAGTTTCG
AATGGACAAAAGGATTTTGACACGATTGTAGCTGTTAGACACCCTTATTCTGACGAAGTAGATAGAAGTATTCGAGTGGTAAGTCC
TTGTGGTATGTGTAGGGAGTTGATTTCAGACTATGCACCAGATTGTTTTGTGTTAATAGAAATGAATGGCAAGTTAGTCAAAACTA
CGATTGAAGAACTCATTCCACTCAAATATACCCGAAATTgAtgatctCGGCCAGGAGTCACAGAAAACAAGATCACTTGGAGATGT
TTCAATCCCGACTTGTGTCGTGCCAGAGTGCTACTTGAAAACTTGAAAATTGCGGACTGTCATGGATTCGCCCTTGTCTTGTGATC
CTTTTTTTGGGGGAGCCAGGTGAGAACAATGTTGTCGATGTGCTTATTTGGCTTCGCAGTCAAAACATGGGATACTTGAGACATGA
AAGAAAAATGCCGCAACGATAGCTCCATCCAATTCCATTCAGCTCCGACTACAGATGATAGCGCTTGACACCAATGACATGCTTGT
ACAGCTGCCATTTGGAAGGCAGGGAAGCTCCATAAGCTCGGGTCCCCAGGACTTTGGTCGGTCTCACATCAGATTCGGCTAGCCAG
CCCATAGCAGCCGCGGGAGATTTCGGTTGTTTGCTACAATGATTGGGGCGCCTTTCTGCGAACTTTGTGACATGTTTCCTCAAAAT
GTCAAGCAATTTTGATCTTAAAAGTTTTGATAATGCTTGCTTCCACAAGCGACCTACAGTAGGAAATGTCTCCACAATCTCCACAG
ATTCAGGACTCATCACTATGTGTGCCGTGCAGGGGTAGGGCGCAGACATGACAACATACAACACACATGAACTAAAGAATCCAAGT
CGCGGACAAAAAAATCTGATCTTACACTTACACAGAATGCAGGTTATTAGCGACGCTTCCATTGCCACCGGAGTGGCAATCGTTGA
GGCGCTTCATCGAACAGAGGGTGAACTTCTTGAGGCTGGGAGGACCGCGCAGATGCGGCTGATAAAAGTTTCACAAGCACACGGAg
aattcactggccgtcgttttacaacgtcgtgactgggaaaaccctggcgttacccaacttaatcgccttgcagcacatccccctttt
cgccagctggcgtaatagcgaagaggcccgcaccgatcgcccttcccaacagttgcgcagcctgaatggcgaatggcgcctgatgc
ggtattttctccttacgcatctgtgcggtatttcacaccgcatatggtgcactctcagtacaatctgctctgatgccgcatagtta
agccagccccgacacccgccaacacccgctgacgcgccctgacgggcttgtctgctcccggcatccgcttacagacaagctgtgac
cgtctccgggagctgcatgtgtcagaggttttcaccgtcatcaccgaaacgcgcga

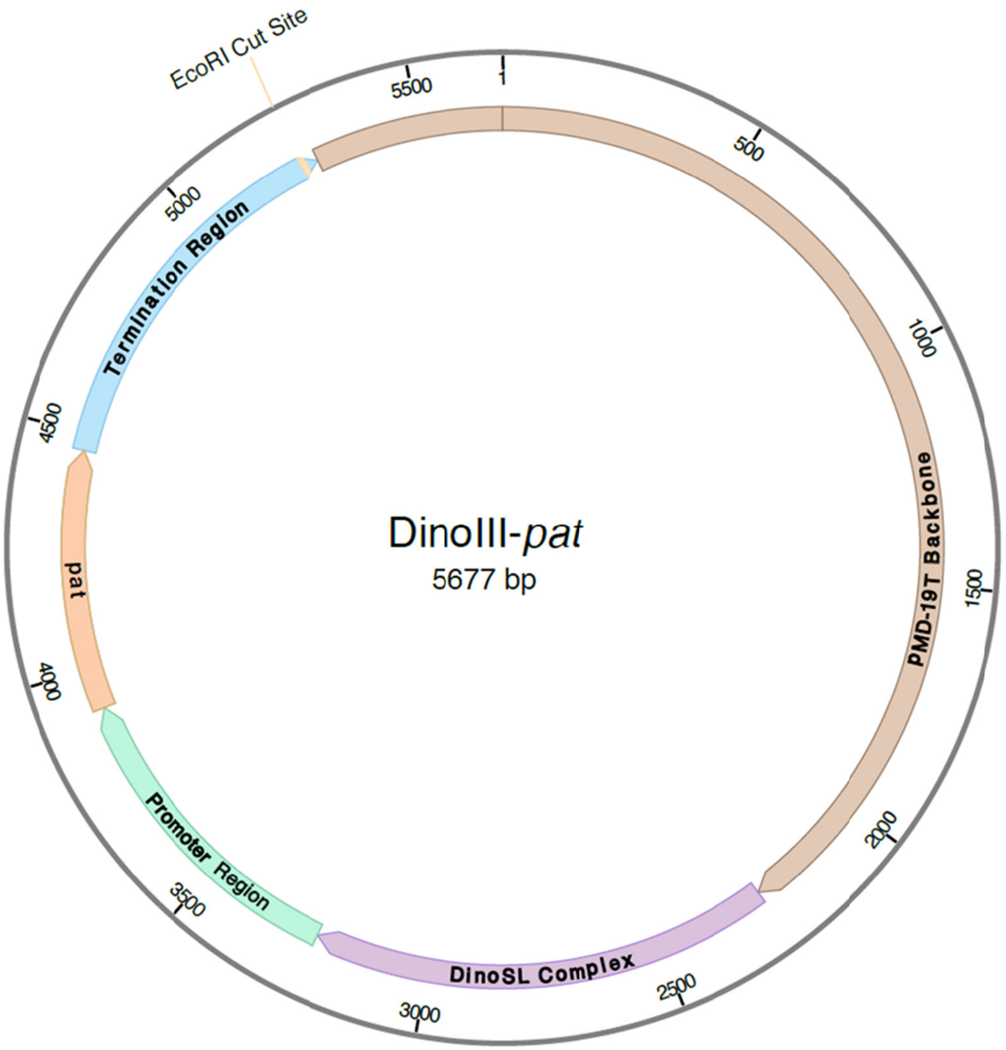


DinoIII-*pat* sequence, containing the pMD™19 T-Vector (shown in lowercase; Takara, Japan), the RNA Elements (highlighted in yellow), the Promoter Region (highlighted in blue), *pat* gene (highlighted in orange), and the Termination Region (highlighted in pink).

gacgaaagggcctcgtgatacgcctatttttataggttaatgtcatgataataatggtttcttagacgtcaggtggcacttttcgg
ggaaatgtgcgcggaacccctatttgtttatttttctaaatacattcaaatatgtatccgctcatgagacaataaccctgataaat
gcttcaataatattgaaaaaggaagagtatgagtattcaacatttccgtgtcgcccttattcccttttttgcggcattttgccttc
ctgtttttgctcacccagaaacgctggtgaaagtaaaagatgctgaagatcagttgggtgcacgagtgggttacatcgaactggat
ctcaacagcggtaagatccttgagagttttcgccccgaagaacgttttccaatgatgagcacttttaaagttctgctatgtggcgc
ggtattatcccgtattgacgccgggcaagagcaactcggtcgccgcatacactattctcagaatgacttggttgagtactcaccag
tcacagaaaagcatcttacggatggcatgacagtaagagaattatgcagtgctgccataaccatgagtgataacactgcggccaac
ttacttctgacaacgatcggaggaccgaaggagctaaccgcttttttgcacaacatgggggatcatgtaactcgccttgatcgttg
ggaaccggagctgaatgaagccataccaaacgacgagcgtgacaccacgatgcctgtagcaatggcaacaacgttgcgcaaactat
taactggcgaactacttactctagcttcccggcaacaattaatagactggatggaggcggataaagttgcaggaccacttctgcgc
tcggcccttccggctggctggtttattgctgataaatctggagccggtgagcgtgggtctcgcggtatcattgcagcactggggcc
agatggtaagccctcccgtatcgtagttatctacacgacggggagtcaggcaactatggatgaacgaaatagacagatcgctgaga
taggtgcctcactgattaagcattggtaactgtcagaccaagtttactcatatatactttagattgatttaaaacttcatttttaa
tttaaaaggatctaggtgaagatcctttttgataatctcatgaccaaaatcccttaacgtgagttttcgttccactgagcgtcaga
ccccgtagaaaagatcaaaggatcttcttgagatcctttttttctgcgcgtaatctgctgcttgcaaacaaaaaaaccaccgctac
cagcggtggtttgtttgccggatcaagagctaccaactctttttccgaaggtaactggcttcagcagagcgcagataccaaatact
gttcttctagtgtagccgtagttaggccaccacttcaagaactctgtagcaccgcctacatacctcgctctgctaatcctgttacc
agtggctgctgccagtggcgataagtcgtgtcttaccgggttggactcaagacgatagttaccggataaggcgcagcggtcgggct
gaacggggggttcgtgcacacagcccagcttggagcgaacgacctacaccgaactgagatacctacagcgtgagctatgagaaagc
gccacgcttcccgaagggagaaaggcggacaggtatccggtaagcggcagggtcggaacaggagagcgcacgagggagcttccagg
gggaaacgcctggtatctttatagtcctgtcgggtttcgccacctctgacttgagcgtcgatttttgtgatgctcgtcaggggggc
ggagcctatggaaaaacgccagcaacgcggcctttttacggttcctggccttttgctggccttttgctcacatgttctttcctgcg
ttatcccctgattctgtggataaccgtattaccgcctttgagtgagctgataccgctcgccgcagccgaacgaccgagcgcagcga

gtcagtgagcgaggaagcggaagagcgcccaatacgcaaaccgcctctccccgcgcgttggccgattcattaatgcagctggcacg
acaggtttcccgactggaaagcgggcagtgagcgcaacgcaattaatgtgagttagctcactcattaggcaccccaggctttacac
tttatgcttccggctcgtatgttgtgtggaattgtgagcggataacaatttcacacaggaaacagctatgaccatgattacgccaa
gcttgcatgcctgcaggtcgacgattCCCGTAGCCATTTTGGCTCAAGGTACAAGTCGGGCTGATGCGGTCACGCAGGCCTCTTTT
GTATCAGATAAGACAGCAGACGTACATGATATAAATATTTATTATCGCGATGCTCCATCTGCAAAATTCCACTTGGCAGAGAAGAT
TTTCAATGCATGAAAACCTCGCAGAACCAAGCACCAATGTCAGATCATGTATGAGTCCTAGGCGCCAGTAAGTCTTTCAATTGGGT
GATGGCCACATTACCTGTTCTCTTCCTGAGGGATGCAGAATGTGGTGTGAGTGGTTGCCGAATGCTGTGCTGAACATGTAGATGGG
CTGCCCCAGCCAAGTGGATAACCTACAAGGACTGCAGGATGTAGATCCAGCAGTCCTGGTAGATAATGATTCAGACACCGGCTCAG
GCTGGCAACAGAGCAGGCAAACCTGAACACTATCGCGCAGTGGTGGGAATACAGCCCAGAATGGGAAGCAACCTGGCCTCTTCAGC
CAGCCATGACCACAAGCAACTCTGCAGTTTTGCAACACGAAATCTTTCAGGGTCTGCTCCCTGTGCTTACACATATTCAAAGATAT
TCTTCTGCGCATTTCAATGCAAATTCTTGGCGATGATTCGCTTCCGCAGGTATAGTGTTACTAGTATATAAGTCTAATCATATACT
TCAATTTATTAGTAATATTGTGAATTTCTAGGCAAGAAGGATGCCTGGACTCTGAATTATTATGGCATTGAGTAGAATCTGGATCT
TGATGATTATGCATTAATATCTTGAAATGCATTTGGATTCCCTTCGGGGATCATCCGTTAAAATTGGAACGATACAGAGAAGATTA
GCATGGCCCCTGCGCAAGGATGACACGCACAAATCGAGAAGTGTAAACAATTTTTTTGAAATTAATTGCCACTTTATTTTGAATAC
CTGAATATGCAGGTGAAGTAGTATAAGGTATTCATTATCATTGCGATTTTCTAGTAAGTCCCGTGGCTGTCATATCTAGTAACCCT
CACCTGGCAGGTGGGGAAAAGGCGAAGACAAATAGAATCAAATAGAATGTATGTGCTGACGTAGGGCTACATTACCTGAGGCTTGA
GGAACCTGTGGGTATGAATGCTACTGTTGGCAGCATCCAGTAGGTTCAGAAACACACGTCCCCACGTAATTTTTTGCATGTCTAAA
TGCACTGTGATGATTGATGATGATTGATTCCTTGAAGGCCATGGTGGGTTACTCTGTGGCCTCTTGCCACTGACTTCCAGGAAAAC
GCGGATTTTCTGACCATCACAAGGGACCACTACGAGAAGTATGCAGCGATCTCAGCAGAGTTCATTTACTGACCATTCTCAGGAAG
ATCGCTCGGGATCACTTGGGTAATATTGCTTTCTTCGCAGATGTTTTTTCCCTATTAAGCTTTTTGAGTCCCCTCATCGAGTGCCG
CATAAGTTTTTGCGTTGACATTGGCGGCAAAAAGTAAGACTAAACGATAGTTGCTTCAAGCACGCTTCTCAATCAACATTTTTTCA
AATCAAATGGGCACAGGCAGGTACCATCCAGTCCCGCAGAAGTGCTTATATATATATATATTATACACTCACGTGAGTGTGTTCAA
TGCAGGTCTGTTCCTCAAGTGCTTAGACCAAATTTCGGAGCCtctagtATGAGCCCAGAACGACGCCCGGCCGACATCCGCCGTGC
CACCGAGGCGGACATGCCGGCGGTCTGCACCATCGTCAACCACTACATCGAGACAAGCACGGTCAACTTCCGTACCGAGCCGCAGG
AACCGCAGGAGTGGACGGACGACCTCGTCCGTCTGCGGGAGCGCTATCCCTGGCTCGTCGCCGAGGTGGACGGCGAGGTCGCCGGC
ATCGCCTACGCGGGCCCCTGGAAGGCACGCAACGCCTACGACTGGACGGCCGAGTCGACCGTGTACGTCTCCCCCCGCCACCAGCG
GACGGGACTGGGCTCCACGCTCTACACCCACCTGCTGAAGTCCCTGGAGGCACAGGGCTTCAAGAGCGTGGTCGCTGTCATCGGGC
TGCCCAACGACCCGAGCGTGCGCATGCACGAGGCGCTCGGATATGCCCCCCGCGGCATGCTGCGGGCGGCCGGCTTCAAGCACGGG
AACTGGCATGACGTGGGTTTCTGGCAGCTGGACTTCAGCCTGCCGGTGCCGCCCCGTCCGGTCCTGCCCGTCACCGAAATCTGAtg
atctCGGCCAGGAGTCACAGAAAACAAGATCACTTGGAGATGTTTCAATCCCGACTTGTGTCGTGCCAGAGTGCTACTTGAAAACT
TGAAAATTGCGGACTGTCATGGATTCGCCCTTGTCTTGTGATCCTTTTTTTGGGGGAGCCAGGTGAGAACAATGTTGTCGATGTGC
TTATTTGGCTTCGCAGTCAAAACATGGGATACTTGAGACATGAAAGAAAAATGCCGCAACGATAGCTCCATCCAATTCCATTCAGC
TCCGACTACAGATGATAGCGCTTGACACCAATGACATGCTTGTACAGCTGCCATTTGGAAGGCAGGGAAGCTCCATAAGCTCGGGT
CCCCAGGACTTTGGTCGGTCTCACATCAGATTCGGCTAGCCAGCCCATAGCAGCCGCGGGAGATTTCGGTTGTTTGCTACAATGAT
TGGGGCGCCTTTCTGCGAACTTTGTGACATGTTTCCTCAAAATGTCAAGCAATTTTGATCTTAAAAGTTTTGATAATGCTTGCTTC
CACAAGCGACCTACAGTAGGAAATGTCTCCACAATCTCCACAGATTCAGGACTCATCACTATGTGTGCCGTGCAGGGGTAGGGCGC
AGACATGACAACATACAACACACATGAACTAAAGAATCCAAGTCGCGGACAAAAAAATCTGATCTTACACTTACACAGAATGCAGG
TTATTAGCGACGCTTCCATTGCCACCGGAGTGGCAATCGTTGAGGCGCTTCATCGAACAGAGGGTGAACTTCTTGAGGCTGGGAGG
ACCGCGCAGATGCGGCTGATAAAAGTTTCACAAGCACACGGAgaattcactggccgtcgttttacaacgtcgtgactgggaaaacc
ctggcgttacccaacttaatcgccttgcagcacatccccctttcgccagctggcgtaatagcgaagaggcccgcaccgatcgccct
tcccaacagttgcgcagcctgaatggcgaatggcgcctgatgcggtattttctccttacgcatctgtgcggtatttcacaccgcat
atggtgcactctcagtacaatctgctctgatgccgcatagttaagccagccccgacacccgccaacacccgctgacgcgccctgac
gggcttgtctgctcccggcatccgcttacagacaagctgtgaccgtctccgggagctgcatgtgtcagaggttttcaccgtcatca
ccgaaacgcgcga

*Karlodinium veneficum*

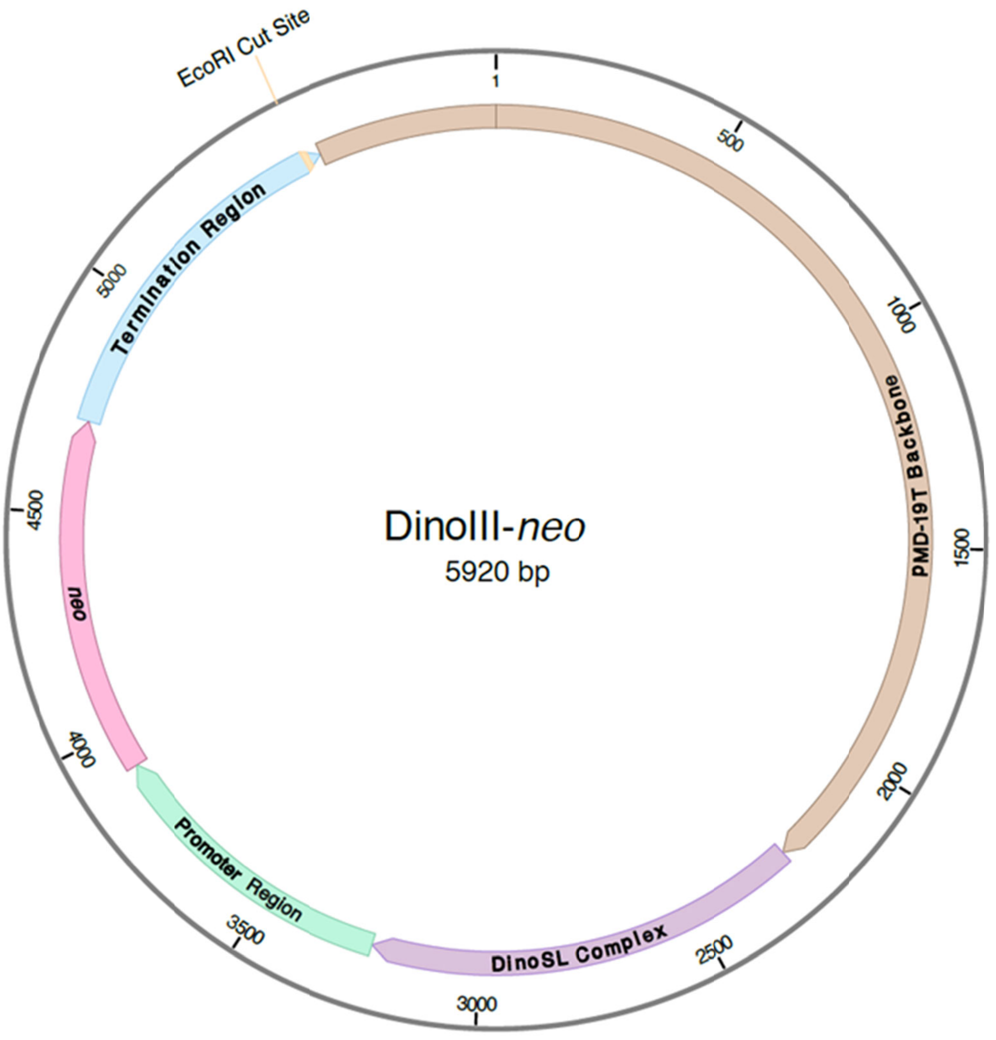


```
> DinoIII-neo
gacgaaagggcctcgtgatacgcctatttttataggttaatgtcatgataataatggtttcttagacgtcaggtggcacttttcgg
ggaaatgtgcgcggaacccctatttgtttatttttctaaatacattcaaatatgtatccgctcatgagacaataaccctgataaat
gcttcaataatattgaaaaaggaagagtatgagtattcaacatttccgtgtcgcccttattcccttttttgcggcattttgccttc
ctgtttttgctcacccagaaacgctggtgaaagtaaaagatgctgaagatcagttgggtgcacgagtgggttacatcgaactggat
ctcaacagcggtaagatccttgagagttttcgccccgaagaacgttttccaatgatgagcacttttaaagttctgctatgtggcgc
ggtattatcccgtattgacgccgggcaagagcaactcggtcgccgcatacactattctcagaatgacttggttgagtactcaccag
tcacagaaaagcatcttacggatggcatgacagtaagagaattatgcagtgctgccataaccatgagtgataacactgcggccaac
ttacttctgacaacgatcggaggaccgaaggagctaaccgcttttttgcacaacatgggggatcatgtaactcgccttgatcgttg
ggaaccggagctgaatgaagccataccaaacgacgagcgtgacaccacgatgcctgtagcaatggcaacaacgttgcgcaaactat
taactggcgaactacttactctagcttcccggcaacaattaatagactggatggaggcggataaagttgcaggaccacttctgcgc
tcggcccttccggctggctggtttattgctgataaatctggagccggtgagcgtgggtctcgcggtatcattgcagcactggggcc
agatggtaagccctcccgtatcgtagttatctacacgacggggagtcaggcaactatggatgaacgaaatagacagatcgctgaga
taggtgcctcactgattaagcattggtaactgtcagaccaagtttactcatatatactttagattgatttaaaacttcatttttaa
tttaaaaggatctaggtgaagatcctttttgataatctcatgaccaaaatcccttaacgtgagttttcgttccactgagcgtcaga
ccccgtagaaaagatcaaaggatcttcttgagatcctttttttctgcgcgtaatctgctgcttgcaaacaaaaaaaccaccgctac
cagcggtggtttgtttgccggatcaagagctaccaactctttttccgaaggtaactggcttcagcagagcgcagataccaaatact
gttcttctagtgtagccgtagttaggccaccacttcaagaactctgtagcaccgcctacatacctcgctctgctaatcctgttacc
agtggctgctgccagtggcgataagtcgtgtcttaccgggttggactcaagacgatagttaccggataaggcgcagcggtcgggct
gaacggggggttcgtgcacacagcccagcttggagcgaacgacctacaccgaactgagatacctacagcgtgagctatgagaaagc
gccacgcttcccgaagggagaaaggcggacaggtatccggtaagcggcagggtcggaacaggagagcgcacgagggagcttccagg
gggaaacgcctggtatctttatagtcctgtcgggtttcgccacctctgacttgagcgtcgatttttgtgatgctcgtcaggggggc
ggagcctatggaaaaacgccagcaacgcggcctttttacggttcctggccttttgctggccttttgctcacatgttctttcctgcg
ttatcccctgattctgtggataaccgtattaccgcctttgagtgagctgataccgctcgccgcagccgaacgaccgagcgcagcga
```

gtcagtgagcgaggaagcggaagagcgcccaatacgcaaaccgcctctccccgcgcgttggccgattcattaatgcagctggcacg
acaggtttcccgactggaaagcgggcagtgagcgcaacgcaattaatgtgagttagctcactcattaggcaccccaggctttacac
tttatgcttccggctcgtatgttgtgtggaattgtgagcggataacaatttcacacaggaaacagctatgaccatgattacgccaa
gcttgcatgcctgcaggtcgacgattCCCGTAGCCATTTTGGCTCAAGGTACAAGTCGGGCTGATGCGGTCACGCAGGCCTCTTTT
GTATCAGATAAGACAGCAGACGTACATGATATAAATATTTATTATCGCGATGCTCCATCTGCAAAATTCCACTTGGCAGAGAAGAT
TTTCAATGCATGAAAACCTCGCAGAACCAAGCACCAATGTCAGATCATGTATGAGTCCTAGGCGCCAGTAAGTCTTTCAATTGGGT
GATGGCCACATTACCTGTTCTCTTCCTGAGGGATGCAGAATGTGGTGTGAGTGGTTGCCGAATGCTGTGCTGAACATGTAGATGGG
CTGCCCCAGCCAAGTGGATAACCTACAAGGACTGCAGGATGTAGATCCAGCAGTCCTGGTAGATAATGATTCAGACACCGGCTCAG
GCTGGCAACAGAGCAGGCAAACCTGAACACTATCGCGCAGTGGTGGGAATACAGCCCAGAATGGGAAGCAACCTGGCCTCTTCAGC
CAGCCATGACCACAAGCAACTCTGCAGTTTTGCAACACGAAATCTTTCAGGGTCTGCTCCCTGTGCTTACACATATTCAAAGATAT
TCTTCTGCGCATTTCAATGCAAATTCTTGGCGATGATTCGCTTCCGCAGGTATAGTGTTACTAGTATATAAGTCTAATCATATACT
TCAATTTATTAGTAATATTGTGAATTTCTAGGCAAGAAGGATGCCTGGACTCTGAATTATTATGGCATTGAGTAGAATCTGGATCT
TGATGATTATGCATTAATATCTTGAAATGCATTTGGATTCCCTTCGGGGATCATCCGTTAAAATTGGAACGATACAGAGAAGATTA
GCATGGCCCCTGCGCAAGGATGACACGCACAAATCGAGAAGTGTAAACAATTTTTTTGAAATTAATTGCCACTTTATTTTGAATAC
CTGAATATGCAGGTGAAGTAGTATAAGGTATTCATTATCATTGCGATTTTCTAGTAAGTCCCGTGGCTGTCATATCTAGTAACCCT
CACCTGGCAGGTGGGGAAAAGGCGAAGACAAATAGAATCAAATAGAATGTATGTGCTGACGTAGGGCTACATTACCTGAGGCTTGA
GGAACCTGTGGGTATGAATGCTACTGTTGGCAGCATCCAGTAGGTTCAGAAACACACGTCCCCACGTAATTTTTTGCATGTCTAAA
TGCACTGTGATGATTGATGATGATTGATTCCTTGAAGGCCATGGTGGGTTACTCTGTGGCCTCTTGCCACTGACTTCCAGGAAAAC
GCGGATTTTCTGACCATCACAAGGGACCACTACGAGAAGTATGCAGCGATCTCAGCAGAGTTCATTTACTGACCATTCTCAGGAAG
ATCGCTCGGGATCACTTGGGTAATATTGCTTTCTTCGCAGATGTTTTTTCCCTATTAAGCTTTTTGAGTCCCCTCATCGAGTGCCG
CATAAGTTTTTGCGTTGACATTGGCGGCAAAAAGTAAGACTAAACGATAGTTGCTTCAAGCACGCTTCTCAATCAACATTTTTTCA
AATCAAATGGGCACAGGCAGGTACCATCCAGTCCCGCAGAAGTGCTTATATATATATATATTATACACTCACGTGAGTGTGTTCAA
TGCAGGTCTGTTCCTCAAGTGCTTAGACCAAATTTCGGAGCCtctagtATGATTGAACAAGATGGATTGCACGCAGGTTCTCCGGC
CGCTTGGGTGGAGAGGCTATTCGGCTATGACTGGGCACAACAGACAATCGGCTGCTCTGATGCCGCCGTGTTCCGGCTGTCAGCGC
AGGGGCGCCCGGTTCTTTTTGTCAAGACCGACCTGTCCGGTGCCCTGAATGAACTGCAAGACGAGGCAGCGCGGCTATCGTGGCTG
GCCACGACGGGCGTTCCTTGCGCAGCTGTGCTCGACGTTGTCACTGAAGCGGGAAGGGACTGGCTGCTATTGGGCGAAGTGCCGGG
GCAGGATCTCCTGTCATCTCACCTTGCTCCTGCCGAGAAAGTATCCATCATGGCTGATGCAATGCGGCGGCTGCATACGCTTGATC
CGGCTACCTGCCCATTCGACCACCAAGCGAAACATCGCATCGAGCGAGCACGTACTCGGATGGAAGCCGGTCTTGTCGATCAGGAT
GATCTGGACGAAGAGCATCAGGGGCTCGCGCCAGCCGAACTGTTCGCCAGGCTCAAGGCGAGCATGCCCGACGGCGAGGATCTCGT
CGTGACCCATGGCGATGCCTGCTTGCCGAATATCATGGTGGAAAATGGCCGCTTTTCTGGATTCATCGACTGTGGCCGGCTGGGTG
TGGCGGACCGCTATCAGGACATAGCGTTGGCTACCCGTGATATTGCTGAAGAGCTTGGCGGCGAATGGGCTGACCGCTTCCTCGTG
CTTTACGGTATCGCCGCTCCCGATTCGCAGCGCATCGCCTTCTATCGCCTTCTTGACGAGTTCTTCTGAtgatctCGGCCAGGAGT
CACAGAAAACAAGATCACTTGGAGATGTTTCAATCCCGACTTGTGTCGTGCCAGAGTGCTACTTGAAAACTTGAAAATTGCGGACT
GTCATGGATTCGCCCTTGTCTTGTGATCCTTTTTTTGGGGGAGCCAGGTGAGAACAATGTTGTCGATGTGCTTATTTGGCTTCGCA
GTCAAAACATGGGATACTTGAGACATGAAAGAAAAATGCCGCAACGATAGCTCCATCCAATTCCATTCAGCTCCGACTACAGATGA
TAGCGCTTGACACCAATGACATGCTTGTACAGCTGCCATTTGGAAGGCAGGGAAGCTCCATAAGCTCGGGTCCCCAGGACTTTGGT
CGGTCTCACATCAGATTCGGCTAGCCAGCCCATAGCAGCCGCGGGAGATTTCGGTTGTTTGCTACAATGATTGGGGCGCCTTTCTG
CGAACTTTGTGACATGTTTCCTCAAAATGTCAAGCAATTTTGATCTTAAAAGTTTTGATAATGCTTGCTTCCACAAGCGACCTACA
GTAGGAAATGTCTCCACAATCTCCACAGATTCAGGACTCATCACTATGTGTGCCGTGCAGGGGTAGGGCGCAGACATGACAACATA
CAACACACATGAACTAAAGAATCCAAGTCGCGGACAAAAAAATCTGATCTTACACTTACACAGAATGCAGGTTATTAGCGACGCTT
CCATTGCCACCGGAGTGGCAATCGTTGAGGCGCTTCATCGAACAGAGGGTGAACTTCTTGAGGCTGGGAGGACCGCGCAGATGCGG
CTGATAAAAGTTTCACAAGCACACGGAgaattcactggccgtcgttttacaacgtcgtgactgggaaaaccctggcgttacccaac
ttaatcgccttgcagcacatccccctttcgccagctggcgtaatagcgaagaggcccgcaccgatcgcccttcccaacagttgcgc
agcctgaatggcgaatggcgcctgatgcggtattttctccttacgcatctgtgcggtatttcacaccgcatatggtgcactctcag
tacaatctgctctgatgccgcatagttaagccagccccgacacccgccaacacccgctgacgcgccctgacgggcttgtctgctcc
cggcatccgcttacagacaagctgtgaccgtctccgggagctgcatgtgtcagaggttttcaccgtcatcaccgaaacgcgcga

#### *Breviolum (Symbiodinium) sp.*

no vector; transformation was with fluoroscein only.

#### *Crypthecodinium cohnii*

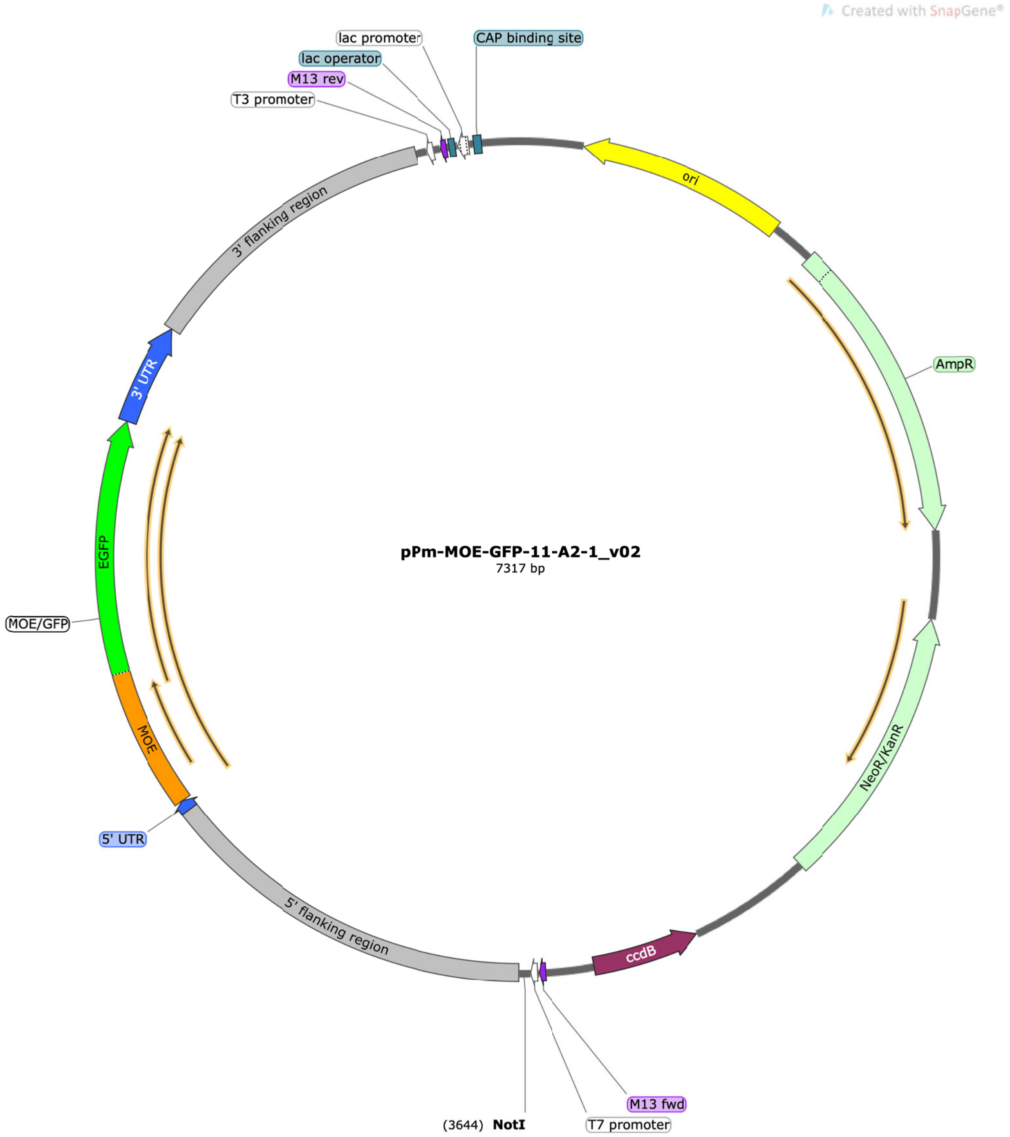


>pPmMOE[MOE]:GFP-11

Published: Fernández Robledo, J.A., Lin, Z., Vasta, G.R., 2008. Transfection of the protozoan parasite *Perkinsus marinus*. Mol Biochem Parasitol 157, 44-53.

From which a labelled probe is obtained:

Sequence of the labeled probe
Forward primer – CCGCACATGTATGGTGAGCAAGGGCGAGGAGC
Reverse primer – CGTAGGACATGTCTTGTACA
**CCGCACATGTATGGTGAGCAAGGGCGAGGAGC**TGTTCACCGGGGTGGTGCCCATCCTGGTCGAGCTGGACGGCGACGTAAACGGCC
ACAAGTTCAGCGTGTCCGGCGAGGGCGAGGGCGATGCCACCTACGGCAAGCTGACCCTGAAGTTCATCTGCACCACCGGCAAGCTG
CCCGTGCCCTGGCCCACCCTCGTGACCACCCTGACCTACGGCGTGCAGTGCTTCAGCCGCTACCCCGACCACATGAAGCAGCACGA

CTTCTTCAAGTCCGCCATGCCCGAAGGCTACGTCCAGGAGCGCACCATCTTCTTCAAGGACGACGGCAACTACAAGACCCGCGCCG
AGGTGAAGTTCGAGGGCGACACCCTGGTGAACCGCATCGAGCTGAAGGGCATCGACTTCAAGGAGGACGGCAACATCCTGGGGCAC
AAGCTGGAGTACAACTACAACAGCCACAACGTCTATATCATGGCCGACAAGCAGAAGAACGGCATCAAGGTGAACTTCAAGATCCG
CCACAACATCGAGGACGGCAGCGTGCAGCTCGCCGACCACTACCAGCAGAACACCCCCATCGGCGACGGCCCCGTGCTGCTGCCCG
ACAACCACTACCTGAGCACCCAGTCCGCCCTGAGCAAAGACCCCAACGAGAAGCGCGATCACATGGTCCTGCTGGAGTTCGTGACC
GCCGCCGGGATCACTCTCGGCATGGACGAGC**TGTACAAGACATGTCCTACG**

#### *Amphidinium carterae*

Note that the region labelled ‘atpB_CAT_MC’ is a linearized *A. carterae* atpB minicircle (Genbank AJ250266) with the atpB gene directly replaced with a *A. carterae* codon-optomized chloramphenicol acetyl transferase gene.

```
LOCUS       pAmpatpBChl        3870 bp   DNA      circular

SOURCE

  ORGANISM

FEATURES             Location/Qualifiers

     CDS             complement(2882..3742)

                     /label="AmpR"

                     /vntifkey="4"

     rep_origin      complement(2067..2734)

                     /label="Col\E1\origin"

                     /vntifkey="33"

     CDS             387..1882

                     /label="atpB_CAT_MC"

ORIGIN

        1 CTAAATTGTA AGCGTTAATA TTTTGTTAAA ATTCGCGTTA AATTTTTGTT AAATCAGCTC

       61 ATTTTTTAAC CAATAGGCCG AAATCGGCAA AATCCCTTAT AAATCAAAAG AATAGACCGA

      121 GATAGGGTTG AGTGGCCGCT ACAGGGCGCT CCCATTCGCC ATTCAGGCTG CGCAACTGTT

      181 GGGAAGGGCG TTTCGGTGCG GGCCTCTTCG CTATTACGCC AGCTGGCGAA AGGGGGATGT

      241 GCTGCAAGGC GATTAAGTTG GGTAACGCCA GGGTTTTCCC AGTCACGACG TTGTAAAACG

      301 ACGGCCAGTG AGCGCGACGT AATACGACTC ACTATAGGGC GAATTGGCGG AAGGCCGTCA

      361 AGGCCACGTG TCTTGTCCGC GGTACCAGAA ACGTACGAAG GATAACGAAG GCTGTTGAAG

      421 TCACAATCGA CAGGAGTGAT TAACTGAGGA CAAATGTTGT CAGACAACCC CTCGTCCTGG

      481 TCATGTGATA GGCTTCTCGA TGAAACTGTT CCATCTCTAC CCCAGTAGAG AAAAATCCAG

      541 GTCATATCAT AGGAGTGAGA ATGTAACAAT CGAGAAGAAT GAATGTAGAA GGTGGTAAAT

      601 CATAGGGAGC TGGAGATCTC GCAATACTGC AGAAGGCTCA GACAGGGACG ATGATAATTG

      661 ATGAAACGAC AATTGATAAA GATAAATAAG ATGAAACACT CCCTCCTGAT ATGAAGGAAG

      721 GGATTAATTA ACAAATGCCT CCTGGTTAGG CCAGAAGTAT ACCAGGAAAT GAGTTGATAA

      781 AAGCGGTCTC GGGGGTGTGC CTAAAAAGGT AAAGAAGCAT CAAACAGACT TAGATCCCTA

      841 CCTATCCGGG GATGTTGTCC CGACTGGGAT TGACAACGGG GATGGGGTAA CCGGTGTGAC

      901 GAAGTATGTT GCACTGCTGG GTTACAATGG CTGATGATAA GGCGCACCCG TAATCATAAT

      961 TCCGTCCTCA GGGGCAGGTG GGGTGGTCTT TAGGCTGATT CCTGCATACT CAATTGGTTT

     1021 AGCGTTTGAA CCGTGAAAAG CTGTTCTCTC ATCTCCTGTA CTGGTCCTGT AATTGATATC
```

1081 GCAACTGTAA ACGTGGACCC TGCGTCCCGT GAGTTGTCTA AGTACTTAGT AGGTAGGAAC

1141 CTCTATCAAC CTTCGGTCTA TGACTCCGTC TGTATCTACC GTCCGCTTGT TTCTATGGAA

1201 AAGAAGATCA CCGGTTACAC AACCGTAGAT ATTTCTCAGT GGCATCGTAA GGAACATTTC

1261 GAGGCATTCC AGTCTGTAGC ACAGTGTACA TATAACCAGA CAGTACAGCT TGATATCACC

1321 GCATTCCTTA AGACAGTAAA GAAGAACAAG CACAAGTTCT ACCCAGCATT CATCCATATC

1381 CTTGCACGTC TTATGAACGC ACATCCAGAG TTCCGTATGG CAATGAAGGA TGGTGAGCTT

1441 GTCATCTGGG ATTCTGTACA TCCATGTTAT ACAGTATTCC ATGAGCAGAC AGAGACATTC

1501 TCTTCCCTTT GGTCTGAGTA TCATGATGAT TTCCGTCAGT TCCTTCATAT CTACTCTCAG

1561 GATGTAGCAT GTTATGGTGA GAACCTTGCA TATTTCCCAA AGGGTTTCAT CGAAAATATG

1621 TTCTTCGTAT CTGCAAACCC TTGGGTATCT TTCACATCTT TCGATCTTAA CGTCGCGAAC

1681 ATGGACAATT TCTTCGCACC AGTATTCACC ATGGGTAAGT ATTATACACA GGGTGATAAG

1741 GTCCTTATGC CACTTGCAAT CCAGGTACAT CATGCAGTAT GTGATGGTTT CCATGTAGGT

1801 CGTATGCTTA ACGAGCTTCA GCAGTATTGT GACGAGTGGC AAGGTGGTGC ATAGTTGAAT

1861 TGTGTAGGTA ATAATATTAT TAGAGCTCGG AGCACAAGAC TGGCCTCATG GGCCTTCCGC

1921 TCACTGCCCG CTTTCCAGTC GGGAAACCTG TCGTGCCAGC TGCATTAACA TGGTCATAGC

1981 TGTTTCCTTG CGTATTGGGC GCTCTCCGCT TCCTCGCTCA CTGACTCGCT GCGCTCGGTC

2041 GTTCGGGTAA AGCCTGGGGT GCCTAATGAG CAAAAGGCCA GCAAAAGGCC AGGAACCGTA

2101 AAAAGGCCGC GTTGCTGGCG TTTTTCCATA GGCTCCGCCC CCCTGACGAG CATCACAAAA

2161 ATCGACGCTC AAGTCAGAGG TGGCGAAACC CGACAGGACT ATAAAGATAC CAGGCGTTTC

2221 CCCCTGGAAG CTCCCTCGTG CGCTCTCCTG TTCCGACCCT GCCGCTTACC GGATACCTGT

2281 CCGCCTTTCT CCCTTCGGGA AGCGTGGCGC TTTCTCATAG CTCACGCTGT AGGTATCTCA

2341 GTTCGGTGTA GGTCGTTCGC TCCAAGCTGG GCTGTGTGCA CGAACCCCCC GTTCAGCCCG

2401 ACCGCTGCGC CTTATCCGGT AACTATCGTC TTGAGTCCAA CCCGGTAAGA CACGACTTAT

2461 CGCCACTGGC AGCAGCCACT GGTAACAGGA TTAGCAGAGC GAGGTATGTA GGCGGTGCTA

2521 CAGAGTTCTT GAAGTGGTGG CCTAACTACG GCTACACTAG AAGAACAGTA TTTGGTATCT

2581 GCGCTCTGCT GAAGCCAGTT ACCTTCGGAA AAAGAGTTGG TAGCTCTTGA TCCGGCAAAC

2641 AAACCACCGC TGGTAGCGGT GGTTTTTTTG TTTGCAAGCA GCAGATTACG CGCAGAAAAA

2701 AAGGATCTCA AGAAGATCCT TTGATCTTTT CTACGGGGTC TGACGCTCAG TGGAACGAAA

2761 ACTCACGTTA AGGGATTTTG GTCATGAGAT TATCAAAAAG GATCTTCACC TAGATCCTTT

2821 TAAATTAAAA ATGAAGTTTT AAATCAATCT AAAGTATATA TGAGTAAACT TGGTCTGACA

2881 GTTACCAATG CTTAATCAGT GAGGCACCTA TCTCAGCGAT CTGTCTATTT CGTTCATCCA

2941 TAGTTGCCTG ACTCCCCGTC GTGTAGATAA CTACGATACG GGAGGGCTTA CCATCTGGCC

3001 CCAGTGCTGC AATGATACCG CGAGAACCAC GCTCACCGGC TCCAGATTTA TCAGCAATAA

3061 ACCAGCCAGC CGGAAGGGCC GAGCGCAGAA GTGGTCCTGC AACTTTATCC GCCTCCATCC

3121 AGTCTATTAA TTGTTGCCGG GAAGCTAGAG TAAGTAGTTC GCCAGTTAAT AGTTTGCGCA

3181 ACGTTGTTGC CATTGCTACA GGCATCGTGG TGTCACGCTC GTCGTTTGGT ATGGCTTCAT

3241 TCAGCTCCGG TTCCCAACGA TCAAGGCGAG TTACATGATC CCCCATGTTG TGCAAAAAAG

3301 CGGTTAGCTC CTTCGGTCCT CCGATCGTTG TCAGAAGTAA GTTGGCCGCA GTGTTATCAC

3361 TCATGGTTAT GGCAGCACTG CATAATTCTC TTACTGTCAT GCCATCCGTA AGATGCTTTT

3421 CTGTGACTGG TGAGTACTCA ACCAAGTCAT TCTGAGAATA GTGTATGCGG CGACCGAGTT

3481 GCTCTTGCCC GGCGTCAATA CGGGATAATA CCGCGCCACA TAGCAGAACT TTAAAAGTGC

3541 TCATCATTGG AAAACGTTCT TCGGGGCGAA AACTCTCAAG GATCTTACCG CTGTTGAGAT

3601 CCAGTTCGAT GTAACCCACT CGTGCACCCA ACTGATCTTC AGCATCTTTT ACTTTCACCA

3661 GCGTTTCTGG GTGAGCAAAA ACAGGAAGGC AAAATGCCGC AAAAAAGGGA ATAAGGGCGA

3721 CACGGAAATG TTGAATACTC ATACTCTTCC TTTTTCAATA TTATTGAAGC ATTTATCAGG

3781 GTTATTGTCT CATGAGCGGA TACATATTTG AATGTATTTA GAAAAATAAA CAAATAGGGG

3841 TTCCGCGCAC ATTTCCCCGA AAAGTGCCAC

//

***Discobans***

*Bodo saltans*

### Cassettes for BsEF1α gene C-terminal tagging

***Bodo saltans*** **genome (EF1 alpha gene: 10,296--11,639 nt):**
**scaffold5.1_size271480.3.119183-132508.final**

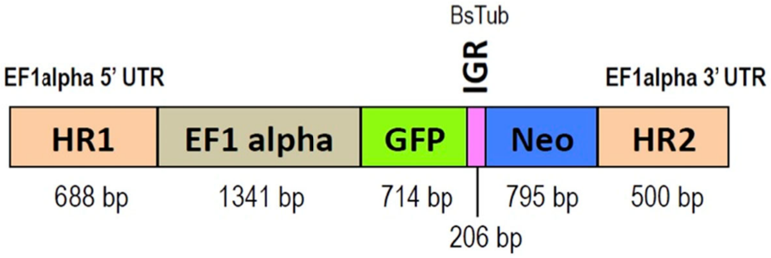


```
>Cassette for tagging EF1alpha gene_IG BsTub_Final_NoPAM
1        TGGGTGCACA CACAGACAGC GGCATCGTGA CAGTCATGTT TGTCCGGCAT GGTATCTCAG
61       ATATCAAAGC CTTTGAGTTT TTATGCAGCA ATCCCGTCAC GCAAACACGC AGCTGGGTAT
121      CATCGCTGGA CATCATGAAA CTGGCGTCCT CTACACTCAG CGGCGACGAC GATAACGACG
181      ACACCGAGTG CATCGTGTGC TTCTTTGGCG AAATGGTAGT CCCGTTCGCT GAATTTCTTC
241      TCAGGAGGCA ACTTAGAGAA CACCATAATC AACTCACCAT CCTGGTGCAG TCGATACGCT
301      CCGAGCTGTC GACGGCTGTT GGAGTGCACC GCGTCAACTC AGCTCTTCTC ACTTTCGGCG
361      TACCATCACC ACAGAAACCG CTTCTGCGGT GCCGCCACAG CTACCCATTC CAGCTCCGCT
421      CTGATCCTGA CCTTCTGCTT GAGAGCAGAA GCAGGCAACT GCCGCTCGGT GCATTCACTC
481      AACGCTTTCT GATGTTGTAA CCTTAGCGTG GAAGGTGTCA GACAATCACG CGTGGAATGT
541      TTTTTTTCTT CATCTTTTGT TGCTCCTCCC CGCTGTTTGC GCGCAATCTC TTCCTTTGTT
601      ACTTGCTACA GTCTTGTAAA CGTACGATTT CCGTTCAACT TCTGTTCATC ACACTCCTCT
661      CTCATTGGCA AGCACAAAAA GCAGCAAGAT GGGTAAGGAT AAGGTTCACA TGTCCCTGGT
721      CGTCATCGGC CACGTCGATG CCGGCAAGTC CACTGCCACC GGTCACTTGA TCTACAAGTG
781      CGGTGGTATT GACAAGCGCA CGATGGAGAA GTTCGAGAAG GAACCCGCCG AAATCGGCAA
841      GGCCTCCTTC AAGTACGCGT GGGTTCTTGA CAAGCTGAAG GCTGAGCGCG AGCGTGGTAT
901      CACCATCGAT ATCGCCCTCT GGAAGTTCGA GTCCCCCAAG TCCGTCTTCA CCATCATCGA
961      TGCCCCCGGA CACCGTGACT TCATCAAGAA CATGATTACC GGAACCTCCC AGGCTGATGC
1021     CGCCGTCCTC GTGATTGCCT CCGGCACTGG CGAGTTCGAG GCTGGCTTCT CCAAGGACGG
1081     CCAGACCCGC GAGCACGCCC TGCTCGCCTT CACCCTCGGC GTCAAGCAGA TGGTGGTCGC
1141     TTGCAACAAG ATGGACTCGA TCAACTTCTC CCAGGCCCGC TACGACGAGA TCGTCTCGAA
1201     CGTTGGCCAG TACCTCAAGA AGGTCGGTTA CAGCATCGAC AAGGTGCGCT TCGTCCCGAT
1261     CTCCGGTTGG GAAGGCGACA ACATGATCGA GAAGTCCGCC CGCATGGAGT GGTACAAGGG
1321     CCCGACCCTG CTGGAGGCCC TCGACCTGCT GGAGGCGCCG AGCCGCCCCT CGGACAAGCC
1381     CCTGCGCCTG CCCCTCCAGG ACGTGTACAA GATCGGCGGT ATCGGCACCG TGCCCGTCGG
1441     CCGTGTTGAG ACCGGCATCC TGCGCCCCGG TGATGTCGTC ACCTTCGCCC CCGCGAACAT
1501     CACCACTGAG GTGAAGAGGC AGCTCTACGT GGTGCAGTCC CTCCCCGAGG CCATCCCCGG
1561     AGACAACGTT GGCTTCAACG TGAAGAACTT GTCGATCAAG GATATCCGCC GTGGCTTCGT
1621     CTGCGGATCC GCCAAGGTGG ACCCCCCGAA GGAGTGCGAG AGCTTCACCG CTCAGGTGAT
1681     CATCCTGAAC CACCCCGGTC AGGTCGGTAA CGGCTACTCC CCCGTGCTCG ATTGCCACAC
1741     CAGCCACATC GCCTGCAAGT TCGCGATGAT CGAGTCGAAG ATCGACCGTC GCTCCGGCAA
1801     GGAGGTCGAG AAGGAGCCCA AGGCGATCAA GTCTGGTGAT GCCGCGATCG TGAAGATGGT
1861     CCCCCAGAAG CCGATGTGCG TCGAAAGTTT TGTGGAGTAC CCACCCCTCG GCCGCTTCGC
1921     CGTGCGTGAC ATGCGCCAGA CCGTTGCCGT CGGTGTCATC AAGTCCGTCG CCAAGAAGGA
1981     GGCGTCCAGC GGCAAGGTCA CCAAGGCTGC CGTCAAGGCC GGCAAGAAGG GTAGTGGGAG
2041     CAACGGCAGC AGCGGATCTG TGAGCAAGGG CGAGGAGCTG TTCACCGGGG TGGTGCCCAT
2101     CCTGGTCGAG CTGGACGGCG ACGTAAACGG CCACAAGTTC AGCGTGCGCG GCGAGGGCGA
```

```
2161     GGGCGATGCC ACCAACGGCA AGCTGACCCT GAAGTTCATC TGCACCACCG GCAAGCTGCC
2221     CGTGCCCTGG CCCACCCTCG TGACCACCCT GACCTACGGC GTGCAGTGCT TCAGCCGCTA
2281     CCCCGACCAC ATGAAGCAGC ACGACTTCTT CAAGTCCGCC ATGCCCGAAG GCTACGTCCA
2341     GGAGCGCACC ATCTCCTTCA AGGACGACGG CACCTACAAG ACCCGCGCCG AGGTGAAGTT
2401     CGAGGGCGAC ACCCTGGTGA ACCGCATCGA GCTGAAGGGC ATCGACTTCA AGGAGGACGG
2461     CAACATCCTG GGGCACAAGC TGGAGTACAA CTTCAACAGC CACAACGTCT ATATCACGGC
2521     CGACAAGCAG AAGAACGGCA TCAAGGCGAA CTTCAAGATC CGCCACAACG TCGAGGACGG
2581     CAGCGTGCAG CTCGCCGACC ACTACCAGCA GAACACCCCC ATCGGCGACG GCCCCGTGCT
2641     GCTGCCCGAC AACCACTACC TGAGCACCCA GTCCAAGCTG AGCAAAGACC CCAACGAGAA
2701     GCGCGATCAC ATGGTCCTGC TGGAGTTCGT GACCGCCGCC GGGATCACTC TCGGCATGGA
2761     CGAGCTGTAC AAGTAGGTGC GATGTACACA CCGTCTCGGA AGAGAACACT TGTGCGTTAG
2821     AGCGCTTGGC TGATGTGTTC GCATTCTTGG GTCGGCGCGC TGTGTTTCTG TGCCGATGCA
2881     CTTGCCGCGT GCTGCTATGC CATTCCTCTC CTCGCTTTTC TTCTTTTCTC ACTGCATACT
2941     AACAGAAAAG AAAAGCTACC ACTCTTCAAA CCACAAAAGA ACATGCTTGA ACAAGATGGA
3001     TTGCACGCAG GTTCTCCGGC CGCTTGGGTG GAGAGGCTAT TCGGCTATGA CTGGGCACAA
3061     CAGACAATCG GCTGCTCTGA TGCCGCCGTG TTCCGGCTGT CAGCGCAGGG GCGCCCGGTT
3121     CTTTTTGTCA AGACCGACCT GTCCGGTGCC CTGAATGAAC TGCAGGACGA GGCAGCGCGG
3181     CTATCGTGGC TGGCCACGAC GGGCGTTCCT TGCGCAGCTG TGCTCGACGT TGTCACTGAA
3241     GCGGGAAGGG ACTGGCTGCT ATTGGGCGAA GTGCCGGGGC AGGATCTCCT GTCATCTCAC
3301     CTTGCTCCTG CCGAGAAAGT ATCCATCATG GCTGATGCAA TGCGGCGGCT GCATACGCTT
3361     GATCCGGCTA CCTGCCCATT CGACCACCAA GCGAAACATC GCATCGAGCG AGCACGTACT
3421     CGGATGGAAG CCGGTCTTGT CGATCAGGAT GATCTGGACG AAGAGCATCA GGGGCTCGCG
3481     CCAGCCGAAC TGTTCGCCAG GCTCAAGGCG CGCATGCCCG ACGGCGAGGA TCTCGTCGTG
3541     ACCCATGGCG ATGCCTGCTT GCCGAATATC ATGGTGGAAA ATGGCCGCTT TTCTGGATTC
3601     ATCGACTGTG GCCGGCTGGG TGTGGCGGAC CGCTATCAGG ACATAGCGTT GGCTACCCGT
3661     GATATTGCTG AAGAGCTTGG CGGCGAATGG GCTGACCGCT TCCTCGTGCT TTACGGTATC
3721     GCCGCTCCCG ATTCGCAGCG CATCGCCTTC TATCGCCTTC TTGACGAGTT CTTCTAGTCG
3781     CGCTGCGCGG TCTACGTTTG CACCAAGAGT GAGTAATTGC AGAGATGTAA GCTCCCCGCA
3841     CGAGTCAATG ACTCGCTCCG ACTAAGAAAA GAAACAAACA AAAGCGGATG ACGCGAACGA
3901     TGGAATTGTG AACGATCTTG ACAGTCTTCG CATTTGTTTG TTGCTTTCCT TATTGCCAAT
3961     CTCGAAGTGA GTCACACAGC AATACACAAC CACACATGTC CACATCTCCG TGTCAGGCAG
4021     TAGCGCTAGT GCCGGATCCA CTTCCATTCT TTGTAGTATT TTCTTAACTC TTGATCAACT
4081     CGCAAATATT CCACGTGACG CACCTTGTCT CTGGTTCTTC TTGAGTGATG GATAAACTCT
4141     TTGATCTGCT TTCTGTCGGC TGCGTACGTA TGGTTTCGCC TCTCGAATGT GCTCAGGAAA
4201     ATGTTTGTGG ATTTCTTTGA GGATAATTCC ATCCTCGACT AAAAACCCTT GAAGTTGTTT
4261     GTTAGAAAAA TTGTGGA
```

#### *Diplonema papillatum*

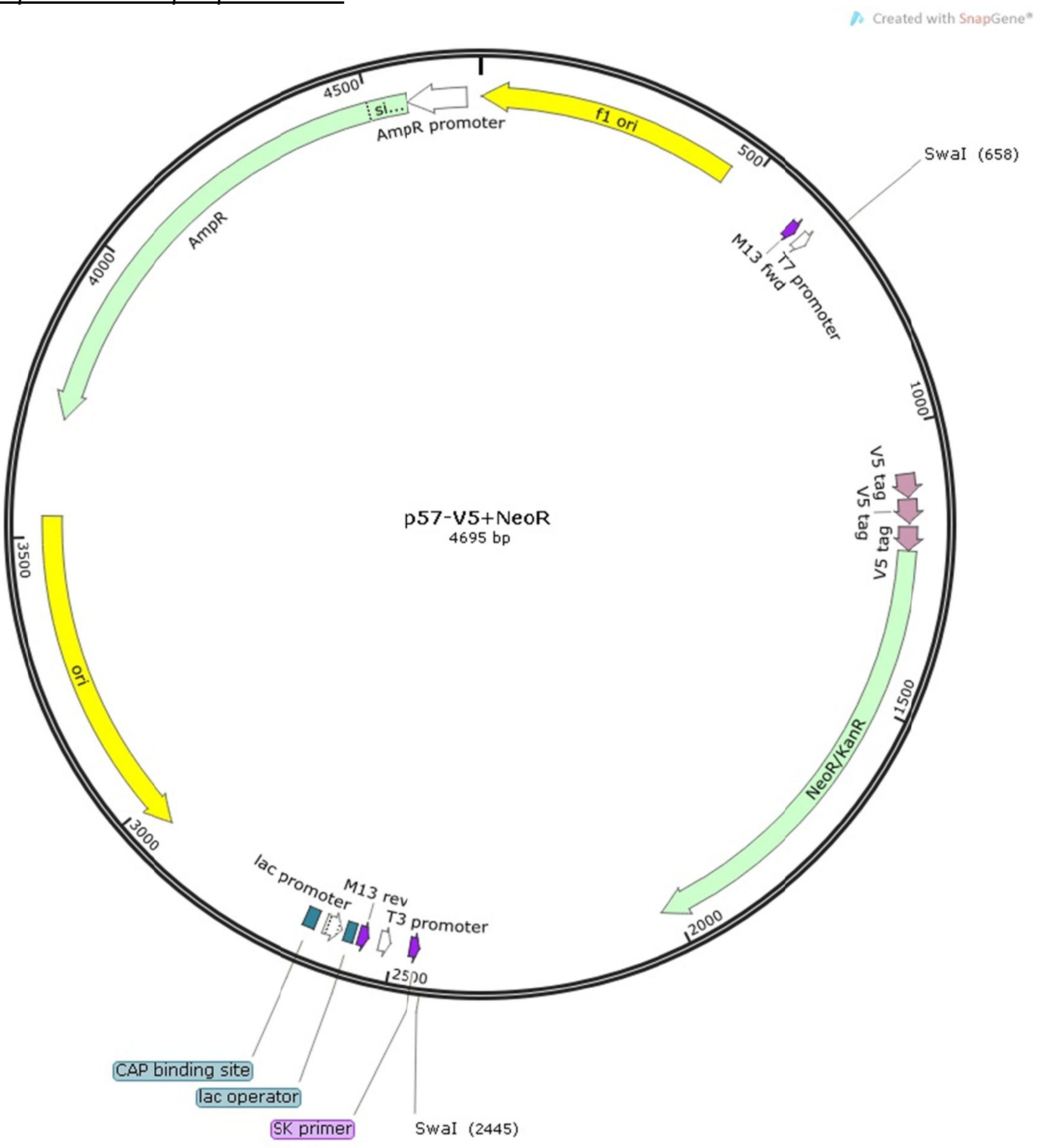


p57-V5+Neo$^{R}$
Plasmid GenBank accession number = MN047315

```
1 ctaaattgta agcgttaata ttttgttaaa attcgcgtta aatttttgtt aaatcagctc
61 atttttaac caataggccg aaatcggcaa aatcccttat aaatcaaaag aatagaccga
121 gatagggttg agtgttgttc cagtttggaa caagagtcca ctattaaaga acgtggactc
181 caacgtcaaa gggcgaaaaa ccgtctatca gggcgatggc ccactacgtg aaccatcacc
241 ctaatcaagt tttttggggt cgaggtgccg taaagcacta aatcggaacc ctaaagggag
301 cccccgattt agagcttgac ggggaaagcc ggcgaacgtg gcgagaaagg aagggaagaa
361 agcgaaagga gcgggcgcta gggcgctggc aagtgtagcg gtcacgctgc gcgtaaccac
421 cacacccgcc gcgcttaatg cgccgctaca gggcgcgtcc cattcgccat tcaggctgcg
481 caactgttgg gaagggcgat cggtgcgggc ctcttcgcta ttacgccagc tggcgaaagg
541 gggatgtgct gcaaggcgat taagttgggt aacgccaggg ttttcccagt cacgacgttg
601 taaaacgacg gccagtgagc gcgcgtaata cgactcacta tagggcgaat tgctatttaa
661 atattgccca tgtccggtgc tcatgcgtta ttgtcgaacg atcgatgaaa ggcgaaaaag
721 ttatttatac tgcagcgatc tacaataaat ttgaataaat acagaacaac taaagttccg
781 gatgtactaa tacattagac acttatacaa ataatataat taaataatct ggaaaaaaat
```

841 aattatcaaa gccataaaaa atgatgaact tcccatatac aaatattcaa atttataaca
901 ataatttagc tcatatatca catgtaaaaa taactttatt aatgtttagt ctatgttatt
961 ttaatttatt attctcatat aattatattt ttctcaattt aacaagcaat ctaagcaact
1021 atcattcaga taattcaata caattttatc tattataaaa caaaataaat tttggagtaa
1081 taatgggtaa gcctatccct aaccctctcc tcggtctcga ttctacgcga ggtaagccaa
1141 tcccaaaccc acttcttggt ctcgactcaa cacagctggg taagcccatc cccaaccccc
1201 tcctcggtct tgatagcacg ggatcggcca ttgaacaaga tggattgcac gcaggttctc
1261 cggccgcttg ggtggagagg ctattcggct atgactgggc acaacagaca atcggctgct
1321 ctgatgccgc cgtgttccgg ctgtcagcgc aggggcgccc ggttcttttt gtcaagaccg
1381 acctgtccgg tgccctgaat gaactgcagg acgaggcagc gcggctatcg tggctggcca
1441 cgacgggcgt tccttgcgca gctgtgctcg acgttgtcac tgaagcggga agggactggc
1501 tgctattggg cgaagtgccg gggcaggatc tcctgtcatc tcaccttgct cctgccgaga
1561 aagtatccat catggctgat gcaatgcggc ggctgcatac gcttgatccg gctacctgcc
1621 cattcgacca ccaagcgaaa catcgcatcg agcgagcacg tactcggatg gaagccggtc
1681 ttgtcgatca ggatgatctg gacgaagagc atcaggggct cgcgccagcc gaactgttcg
1741 ccaggctcaa ggcgcgcatg cccgacggcg aggatctcgt cgtgacgcat ggcgatgcct
1801 gcttgccgaa tatcatggtg gaaaatggcc gcttttctgg attcatcgac tgtggccggc
1861 tgggtgtggc ggaccgctat caggacatag cgttggctac ccgtgatatt gctgaagagc
1921 ttggcggcga atgggctgac cgcttcctcg tgctttacgg tatcgccgct cccgattcgc
1981 agcgcatcgc cttctatcgc cttcttgacg agttcttcta aagttgcagt taaggtcata
2041 ttctttttga ccagttaaag ttaaatgttt gcattaaaca ccaaaagtaa ataactaatt
2101 acttttactc cctctttctc ttgataacat ttctttgaac ataaaacaca tagcttatca
2161 tattttcttt gaaaagttta ataaacaaat ttaaactttc ttttccctat gtattgctta
2221 ataattcaat aacactttct tctatacctg taaatctgca taaagaagtt attttagagt
2281 aatgtacttt tgatcaattc gacacgtaca aaatttatta agcttaacta aaaataatct
2341 ccgttttagc aataatagaa gtatatatcc ttccacctta tatttttgca cattattaaa
2401 acggtgcaat tatttatttt ataaatgatc tggaagtatt tatttaaatc taggatccac
2461 tagttctaga gcggccgcca ccgcggtgga gctccagctt ttgttccctt tagtgagggt
2521 taattgcgcg cttggcgtaa tcatggtcat agctgtttcc tgtgtgaaat tgttatccgc
2581 tcacaattcc acacaacata cgagccggaa gcataaagtg taaagcctgg ggtgcctaat
2641 gagtgagcta actcacatta attgcgttgc gctcactgcc cgctttccag tcgggaaacc
2701 tgtcgtgcca gctgcattaa tgaatcggcc aacgcgcggg gagaggcggt ttgcgtattg
2761 ggcgctcttc cgcttcctcg ctcactgact cgctgcgctc ggtcgttcgg ctgcggcgag
2821 cggtatcagc tcactcaaag gcggtaatac ggttatccac agaatcaggg gataacgcag
2881 gaaagaacat gtgagcaaaa ggccagcaaa aggccaggaa ccgtaaaaag gccgcgttgc
2941 tggcgttttt ccataggctc cgcccccctg acgagcatca caaaaatcga cgctcaagtc
3001 agaggtggcg aaacccgaca ggactataaa gataccaggc gtttccccct ggaagctccc
3061 tcgtgcgctc tcctgttccg accctgccgc ttaccggata cctgtccgcc tttctccctt
3121 cgggaagcgt ggcgctttct catagctcac gctgtaggta tctcagttcg gtgtaggtcg
3181 ttcgctccaa gctgggctgt gtgcacgaac cccccgttca gcccgaccgc tgcgccttat
3241 ccggtaacta tcgtcttgag tccaacccgg taagacacga cttatcgcca ctggcagcag
3301 ccactggtaa caggattagc agagcgaggt atgtaggcgg tgctacagag ttcttgaagt
3361 ggtggcctaa ctacggctac actagaagga cagtatttgg tatctgcgct ctgctgaagc
3421 cagttacctt cggaaaaaga gttggtagct cttgatccgg caaacaaacc accgctggta
3481 gcggtggttt ttttgtttgc aagcagcaga ttacgcgcag aaaaaaagga tctcaagaag
3541 atcctttgat cttttctacg gggtctgacg ctcagtggaa cgaaaactca cgttaaggga
3601 ttttggtcat gagattatca aaaaggatct tcacctagat ccttttaaat taaaaatgaa
3661 gttttaaatc aatctaaagt atatatgagt aaacttggtc tgacagttac caatgcttaa
3721 tcagtgaggc acctatctca gcgatctgtc tatttcgttc atccatagtt gcctgactcc
3781 ccgtcgtgta gataactacg atacgggagg gcttaccatc tggccccagt gctgcaatga
3841 taccgcgaga cccacgctca ccggctccag atttatcagc aataaaccag ccagccggaa
3901 gggccgagcg cagaagtggt cctgcaactt tatccgcctc catccagtct attaattgtt
3961 gccgggaagc tagagtaagt agttcgccag ttaatagttt gcgcaacgtt gttgccattg
4021 ctacaggcat cgtggtgtca cgctcgtcgt ttggtatggc ttcattcagc tccggttccc
4081 aacgatcaag gcgagttaca tgatccccca tgttgtgcaa aaaagcggtt agctccttcg
4141 gtcctccgat cgttgtcaga agtaagttgg ccgcagtgtt atcactcatg gttatggcag
4201 cactgcataa ttctcttact gtcatgccat ccgtaagatg cttttctgtg actggtgagt
4261 actcaaccaa gtcattctga gaatagtgta tgcggcgacc gagttgctct tgcccggcgt
4321 caatacggga taataccgcg ccacatagca gaactttaaa agtgctcatc attggaaaac
4381 gttcttcggg gcgaaaactc tcaaggatct taccgctgtt gagatccagt tcgatgtaac
4441 ccactcgtgc acccaactga tcttcagcat cttttacttt caccagcgtt tctgggtgag
4501 caaaaacagg aaggcaaaat gccgcaaaaa agggaataag ggcgacacgg aaatgttgaa
4561 tactcatact cttccttttt caatattatt gaagcattta tcagggttat tgtctcatga
4621 gcggatacat atttgaatgt atttagaaaa ataaacaaat aggggttccg cgcacatttc
4681 cccgaaaagt gccac

*Eutreptiella gymnastica*

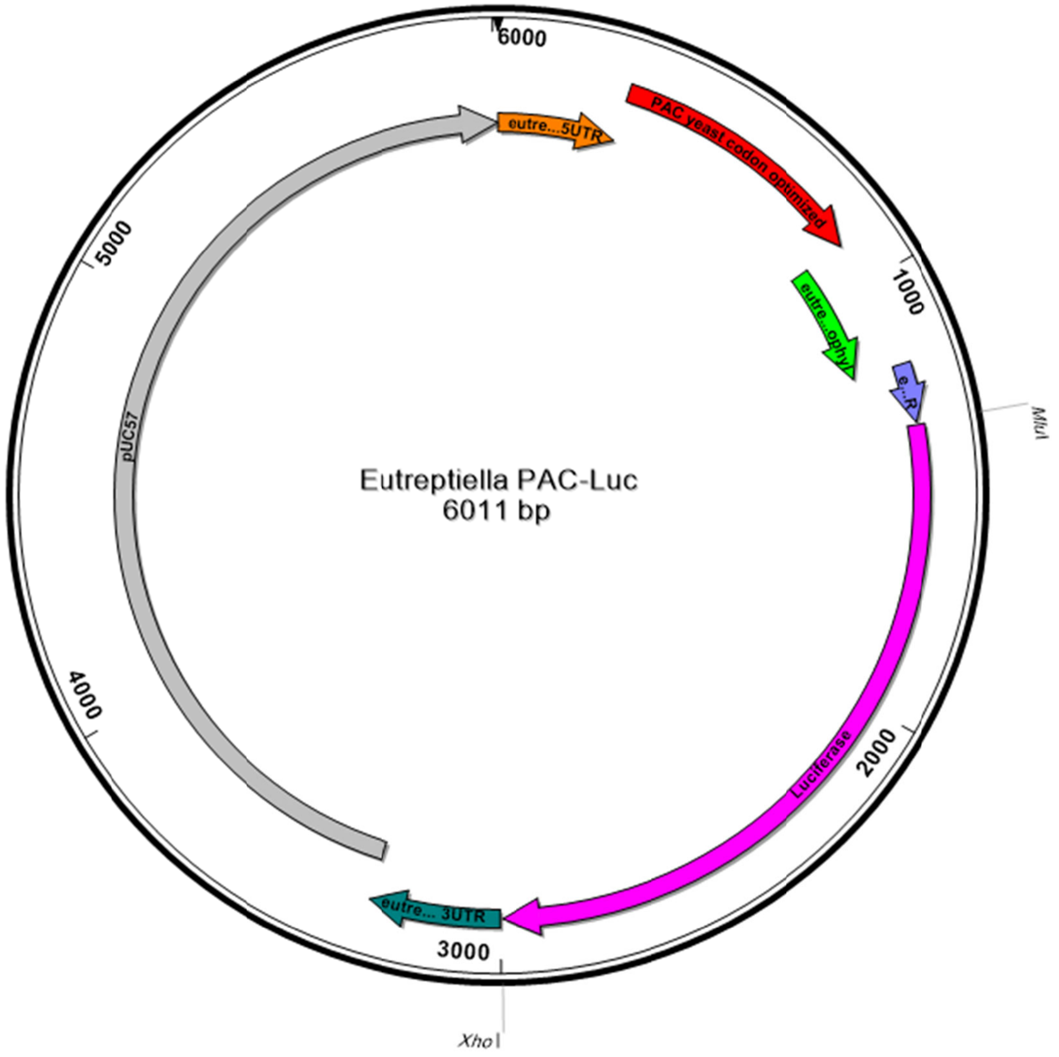


```
>Eutreptiella PAC-Luc
GCTCGTCGTCAAAAATGATCAAAAAGGCCATTCCTATGATGATATGGTGGTAAGCACAATGAAGAAAATGCAAGAC
ATCAGCAGGTTGGCAGAAGTTGCTGAAGAACTGTTTTGCATCATCATACAGCGGGAAGGGCTCTATTACCCCTTCA
ACAACCCCTGGCCTGCACACAAACGGCACTACAACGTACGAAAAAATAGCTCACTTGCATACGCTCCGAATGGAGG
ACCCAGCCAGTCCACTCATGCTCAACAAAAACAAACCACAAATCACTTACAGCTGGGTTTCAACTTTTCACTATGA
CCGAACATAAGCCATCTGTTAGATTGGCTACTAGAGATGATGTTCCAAGAGCTGTTAGAACTTTGGCTGCTGCTTT
TGCTGATTATCCAGCTACTAGACATACTGTTGATCCAGATAGACACATCGAAAGAGTTACCGAATTGCAAGAATTA
TTCTTAACCAGAGTCGGTTTGGATATCGGTAAAGTTTGGGTTGCTGATGATGGTGCTGCTGTTGCAGTTTGGACTA
CTCCAGAATCTTTGGATGCTGGTGCTGTTTTCGCTGAAATTGGTCCAAGAATGGCTGAATTGTCTGGTTCTAGATT
AGCTGCTCAACAACAAATGGAAGGTTTGTTGGCTCCACATAGACCAAAAGAACCAGCTTGGTTTTTGGCTACTGTT
GGTGTTTCTCCAGATCATCAAGGTAAAGGTTTAGGTTCTGCTGTTGTTTTGCCAGGTGTTGAAGCTGCTGAAAGAG
CTGGTGTTCCAGCTTTTTTGGAAACTTCTGCTCCAAGAAATTTGCCATTCTACGAAAGATTGGGTTTCACTGTTAC
TGCCGATGTTGAAGTTCCTGAAGGTCCTAGAACTTGGTGTATGACTAGAAAACCAGGTGCTTAACCCCCGCGTTCT
TGGCCAAGTACGCTGGTGTGGAGTTCGGTGAGCCCATGTGGTTCAAGGCTGGTGCCCAGTTCGCCCCTGGGTCCGT
TGCCATGTTCTCTGCCACCTCCGCCTGGTACGGTGCTGACCGTGTCAAGTGGCTGGGCCCCTACCCTGCTGGGTCC
ACCCCCGACTACTTGGCTGGTGAGTTCCCTGGTGACTACGGTTGGGACCCCGCTGGTTTGGCTGCTGACCCTGTTT
GACCCTGTTTGAACTACCTTGGCAGCGAGGGTTTGGTCCACGCCCAGTCCATCCTGGCTGGAATGAGTGACCGTTT
```

TCGATGAGTTGTCTTTCCTCTGTCATCTAATTTAGGCCATGGGGTATTGCATTGCATCTGCAGATTTCAGTGACCT
GTGTTCTCATTTTGTCCTCATCTCAGCTCTCTTTTCCAGCTAACACGCGTATGGAAGACGCCAAAAACATAAAGAA
AGGCCCGGCGCCATTCTATCCGCTGGAAGATGGAACCGCTGGAGAGCAACTGCATAAGGCTATGAAGAGATACGCC
CTGGTTCCTGGAACAATTGCTTTTACAGATGCACATATCGAGGTGGACATCACTTACGCTGAGTACTTCGAAATGT
CCGTTCGGTTGGCAGAAGCTATGAAACGATATGGGCTGAATACAAATCACAGAATCGTCGTATGCAGTGAAAACTC
TCTTCAATTCTTTATGCCGGTGTTGGGCGCGTTATTTATCGGAGTTGCAGTTGCGCCCGCGAACGACATTTATAAT
GAACGTGAATTGCTCAACAGTATGGGCATTTCGCAGCCTACCGTGGTGTTCGTTTCCAAAAAGGGGTTGCAAAAAA
TTTTGAACGTGCAAAAAAAGCTCCCAATCATCCAAAAAATTATTATCATGGATTCTAAAACGGATTACCAGGGATT
TCAGTCGATGTACACGTTCGTCACATCTCATCTACCTCCCGGTTTTAATGAATACGATTTTGTGCCAGAGTCCTTC
GATAGGGACAAGACAATTGCACTGATCATGAACTCCTCTGGATCTACTGGTCTGCCTAAAGGTGTCGCTCTGCCTC
ATAGAACTGCCTGCGTGAGATTCTCGCATGCCAGAGATCCTATTTTTGGCAATCAAATCATTCCGGATACTGCGAT
TTTAAGTGTTGTTCCATTCCATCACGGTTTTGGAATGTTTACTACACTCGGATATTTGATATGTGGATTTCGAGTC
GTCTTAATGTATAGATTTGAAGAAGAGCTGTTTCTGAGGAGCCTTCAGGATTACAAGATTCAAAGTGCGCTGCTGG
TGCCAACCCTATTCTCCTTCTTCGCCAAAAGCACTCTGATTGACAAATACGATTTATCTAATTTACACGAAATTGC
TTCTGGTGGCGCTCCCCTCTCTAAGGAAGTCGGGGAAGCGGTTGCCAAGAGGTTCCATCTGCCAGGTATCAGGCAA
GGATATGGGCTCACTGAGACTACATCAGCTATTCTGATTACACCCGAGGGGGATGATAAACCGGGCGCGGTCGGTA
AAGTTGTTCCATTTTTTGAAGCGAAGGTTGTGGATCTGGATACCGGGAAAACGCTGGGCGTTAATCAAAGAGGCGA
ACTGTGTGTGAGAGGTCCTATGATTATGTCCGGTTATGTAAACAATCCGGAAGCGACCAACGCCTTGATTGACAAG
GATGGATGGCTACATTCTGGAGACATAGCTTACTGGGACGAAGACGAACACTTCTTCATCGTTGACCGCCTGAAGT
CTCTGATTAAGTACAAAGGCTATCAGGTGGCTCCCGCTGAATTGGAATCCATCTTGCTCCAACACCCCAACATCTT
CGACGCAGGTGTCGCAGGTCTTCCCGACGATGACGCCGGTGAACTTCCCGCCGCCGTTGTTGTTTTGGAGCACGGA
AAGACGATGACGGAAAAAGAGATCGTGGATTACGTCGCCAGTCAAGTAACAACCGCGAAAAAGTTGCGCGGAGGAG
TTGTGTTTGTGGACGAAGTACCGAAAGGTCTTACCGGAAAACTCGACGCAAGAAAAATCAGAGAGATCCTCATAAA
GGCCAAGAAGGGCGGAAAGATCGCCGTGTAACTCGAGACGGTTTGTCCATCTCCCACATGGCCCAGTTTGCCCCTG
AGCCCGTTGCCATGTTCTCTGCCACCTCCGCCTGGTACGGTGCTGACCGTGTCAAGTGGCTGGGACTCTACTCTGC
TAGGTCCACCCCTGACTACTTGACTGGTGAGTTCCCTGGCGACTACGGTGGGGACACCGCTGGATTGGCTGCTGAC
CCTGTCACCTTCAAGGCTACCGTGAGGCTGAGATTGTCCATGCTCGTTTCGCCATGTTGGGAACAGTGGGCTGTTT
GACTCCTGAGCTCTTGTCCAAGTACACTGGTGTATCGGATCCCGGGCCCGTCGACTGCAGAGGCCTGCATGCAAGC
TTGGCGTAATCATGGTCATAGCTGTTTCCTGTGTGAAATTGTTATCCGCTCACAATTCCACACAACATACGAGCCG
GAAGCATAAAGTGTAAAGCCTGGGGTGCCTAATGAGTGAGCTAACTCACATTAATTGCGTTGCGCTCACTGCCCGC
TTTCCAGTCGGGAAACCTGTCGTGCCAGCTGCATTAATGAATCGGCCAACGCGCGGGGAGAGGCGGTTTGCGTATT
GGGCGCTCTTCCGCTTCCTCGCTCACTGACTCGCTGCGCTCGGTCGTTCGGCTGCGGCGAGCGGTATCAGCTCACT
CAAAGGCGGTAATACGGTTATCCACAGAATCAGGGGATAACGCAGGAAAGAACATGTGAGCAAAAGGCCAGCAAAA
GGCCAGGAACCGTAAAAAGGCCGCGTTGCTGGCGTTTTTCCATAGGCTCCGCCCCCCTGACGAGCATCACAAAAAT
CGACGCTCAAGTCAGAGGTGGCGAAACCCGACAGGACTATAAAGATACCAGGCGTTTCCCCCTGGAAGCTCCCTCG
TGCGCTCTCCTGTTCCGACCCTGCCGCTTACCGGATACCTGTCCGCCTTTCTCCCTTCGGGAAGCGTGGCGCTTTC
TCATAGCTCACGCTGTAGGTATCTCAGTTCGGTGTAGGTCGTTCGCTCCAAGCTGGGCTGTGTGCACGAACCCCCC
GTTCAGCCCGACCGCTGCGCCTTATCCGGTAACTATCGTCTTGAGTCCAACCCGGTAAGACACGACTTATCGCCAC
TGGCAGCAGCCACTGGTAACAGGATTAGCAGAGCGAGGTATGTAGGCGGTGCTACAGAGTTCTTGAAGTGGTGGCC
TAACTACGGCTACACTAGAAGAACAGTATTTGGTATCTGCGCTCTGCTGAAGCCAGTTACCTTCGGAAAAAGAGTT
GGTAGCTCTTGATCCGGCAAACAAACCACCGCTGGTAGCGGTGGTTTTTTTGTTTGCAAGCAGCAGATTACGCGCA
GAAAAAAAGGATCTCAAGAAGATCCTTTGATCTTTTCTACGGGGTCTGACGCTCAGTGGAACGAAAACTCACGTTA
AGGGATTTTGGTCATGAGATTATCAAAAAGGATCTTCACCTAGATCCTTTTAAATTAAAAATGAAGTTTTAAATCA
ATCTAAAGTATATATGAGTAAACTTGGTCTGACAGTTACCAATGCTTAATCAGTGAGGCACCTATCTCAGCGATCT
GTCTATTTCGTTCATCCATAGTTGCCTGACTCCCCGTCGTGTAGATAACTACGATACGGGAGGGCTTACCATCTGG
CCCCAGTGCTGCAATGATACCGCGAGACCCACGCTCACCGGCTCCAGATTTATCAGCAATAAACCAGCCAGCCGGA
AGGGCCGAGCGCAGAAGTGGTCCTGCAACTTTATCCGCCTCCATCCAGTCTATTAATTGTTGCCGGGAAGCTAGAG
TAAGTAGTTCGCCAGTTAATAGTTTGCGCAACGTTGTTGCCATTGCTACAGGCATCGTGGTGTCACGCTCGTCGTT
TGGTATGGCTTCATTCAGCTCCGGTTCCCAACGATCAAGGCGAGTTACATGATCCCCCATGTTGTGCAAAAAAGCG
GTTAGCTCCTTCGGTCCTCCGATCGTTGTCAGAAGTAAGTTGGCCGCAGTGTTATCACTCATGGTTATGGCAGCAC
TGCATAATTCTCTTACTGTCATGCCATCCGTAAGATGCTTTTCTGTGACTGGTGAGTACTCAACCAAGTCATTCTG
AGAATAGTGTATGCGGCGACCGAGTTGCTCTTGCCCGGCGTCAATACGGGATAATACCGCGCCACATAGCAGAACT
TTAAAAGTGCTCATCATTGGAAAACGTTCTTCGGGGCGAAAACTCTCAAGGATCTTACCGCTGTTGAGATCCAGTT
CGATGTAACCCACTCGTGCACCCAACTGATCTTCAGCATCTTTTACTTTCACCAGCGTTTCTGGGTGAGCAAAAAC
AGGAAGGCAAAATGCCGCAAAAAAGGGAATAAGGGCGACACGGAAATGTTGAATACTCATACTCTTCCTTTTTCAA
TATTATTGAAGCATTTATCAGGGTTATTGTCTCATGAGCGGATACATATTTGAATGTATTTAGAAAAATAAACAAA
TAGGGGTTCCGCGCACATTTCCCCGAAAAGTGCCACCTGACGTCTAAGAAACCATTATTATCATGACATTAACCTA
TAAAAATAGGCGTATCACGAGGCCCTTTCGTCTCGCGCGTTTCGGTGATGACGGTGAAAACCTCTGACACATGCAG
CTCCCGGAGACGGTCACAGCTTGTCTGTAAGCGGATGCCGGGAGCAGACAAGCCCGTCAGGGCGCGTCAGCGGGTG
TTGGCGGGTGTCGGGGCTGGCTTAACTATGCGGCATCAGAGCAGATTGTACTGAGAGTGCACCATATGCGGTGTGA
AATACCGCACAGATGCGTAAGGAGAAAATACCGCATCAGGCGCCATTCGCCATTCAGGCTGCGCAACTGTTGGGAA
GGGCGATCGGTGCGGGCCTCTTCGCTATTACGCCAGCTGGCGAAAGGGGGATGTGCTGCAAGGCGATTAAGTTGGG
TAACGCCAGGGTTTTCCCAGTCACGACGTTGTAAAACGACGGCCAGTGAATTCGAGCTCGGTACCTCGCGAATGCA
TCTAGAT

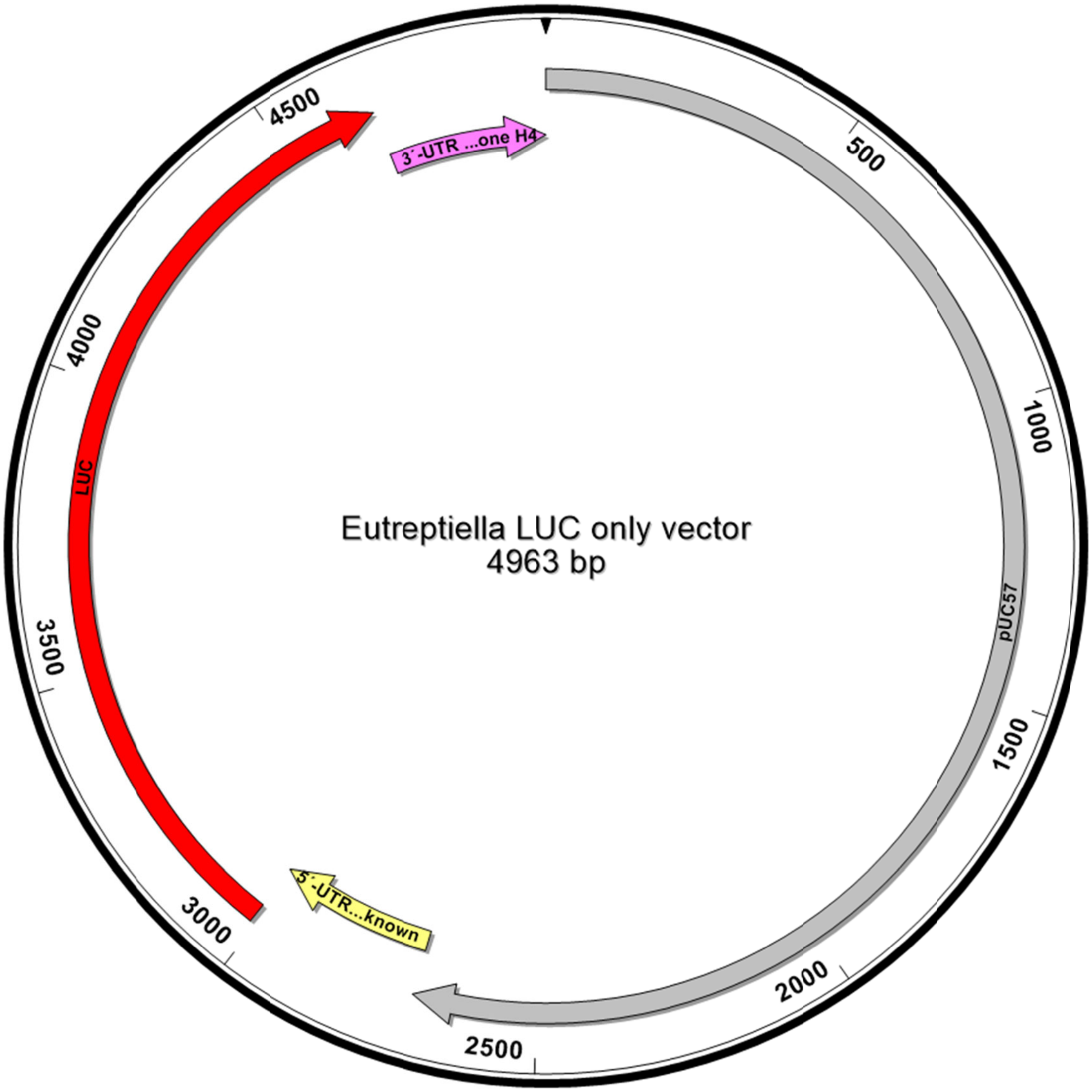


>Luc only vector

1-2270 pUC57 backbone
2771-3010 5´- UTR *Eutreptiella* unknown ORF
3011-4663 Luciferase
4664-4963 3´-UTR *Bigelowiella natans* histone H4

ATCGGATCCCGGGCCCGTCGACTGCAGAGGCCTGCATGCAAGCTTGGCGTAATCATGGTCATAGCTGTTTCCTGTG
TGAAATTGTTATCCGCTCACAATTCCACACAACATACGAGCCGGAAGCATAAAGTGTAAAGCCTGGGGTGCCTAAT
GAGTGAGCTAACTCACATTAATTGCGTTGCGCTCACTGCCCGCTTTCCAGTCGGGAAACCTGTCGTGCCAGCTGCA
TTAATGAATCGGCCAACGCGCGGGGAGAGGCGGTTTGCGTATTGGGCGCTCTTCCGCTTCCTCGCTCACTGACTCG
CTGCGCTCGGTCGTTCGGCTGCGGCGAGCGGTATCAGCTCACTCAAAGGCGGTAATACGGTTATCCACAGAATCAG
GGGATAACGCAGGAAAGAACATGTGAGCAAAAGGCCAGCAAAAGGCCAGGAACCGTAAAAAGGCCGCGTTGCTGGC
GTTTTTCCATAGGCTCCGCCCCCCTGACGAGCATCACAAAAATCGACGCTCAAGTCAGAGGTGGCGAAACCCGACA
GGACTATAAAGATACCAGGCGTTTCCCCCTGGAAGCTCCCTCGTGCGCTCTCCTGTTCCGACCCTGCCGCTTACCG
GATACCTGTCCGCCTTTCTCCCTTCGGGAAGCGTGGCGCTTTCTCATAGCTCACGCTGTAGGTATCTCAGTTCGGT
GTAGGTCGTTCGCTCCAAGCTGGGCTGTGTGCACGAACCCCCCGTTCAGCCCGACCGCTGCGCCTTATCCGGTAAC
TATCGTCTTGAGTCCAACCCGGTAAGACACGACTTATCGCCACTGGCAGCAGCCACTGGTAACAGGATTAGCAGAG
CGAGGTATGTAGGCGGTGCTACAGAGTTCTTGAAGTGGTGGCCTAACTACGGCTACACTAGAAGAACAGTATTTGG
TATCTGCGCTCTGCTGAAGCCAGTTACCTTCGGAAAAAGAGTTGGTAGCTCTTGATCCGGCAAACAAACCACCGCT
GGTAGCGGTGGTTTTTTTGTTTGCAAGCAGCAGATTACGCGCAGAAAAAAAGGATCTCAAGAAGATCCTTTGATCT
TTTCTACGGGGTCTGACGCTCAGTGGAACGAAAACTCACGTTAAGGGATTTTGGTCATGAGATTATCAAAAAGGAT
CTTCACCTAGATCCTTTTAAATTAAAAATGAAGTTTTAAATCAATCTAAAGTATATATGAGTAAACTTGGTCTGAC

AGTTACCAATGCTTAATCAGTGAGGCACCTATCTCAGCGATCTGTCTATTTCGTTCATCCATAGTTGCCTGACTCC
CCGTCGTGTAGATAACTACGATACGGGAGGGCTTACCATCTGGCCCCAGTGCTGCAATGATACCGCGAGACCCACG
CTCACCGGCTCCAGATTTATCAGCAATAAACCAGCCAGCCGGAAGGGCCGAGCGCAGAAGTGGTCCTGCAACTTTA
TCCGCCTCCATCCAGTCTATTAATTGTTGCCGGGAAGCTAGAGTAAGTAGTTCGCCAGTTAATAGTTTGCGCAACG
TTGTTGCCATTGCTACAGGCATCGTGGTGTCACGCTCGTCGTTTGGTATGGCTTCATTCAGCTCCGGTTCCCAACG
ATCAAGGCGAGTTACATGATCCCCCATGTTGTGCAAAAAAGCGGTTAGCTCCTTCGGTCCTCCGATCGTTGTCAGA
AGTAAGTTGGCCGCAGTGTTATCACTCATGGTTATGGCAGCACTGCATAATTCTCTTACTGTCATGCCATCCGTAA
GATGCTTTTCTGTGACTGGTGAGTACTCAACCAAGTCATTCTGAGAATAGTGTATGCGGCGACCGAGTTGCTCTTG
CCCGGCGTCAATACGGGATAATACCGCGCCACATAGCAGAACTTTAAAAGTGCTCATCATTGGAAAACGTTCTTCG
GGGCGAAAACTCTCAAGGATCTTACCGCTGTTGAGATCCAGTTCGATGTAACCCACTCGTGCACCCAACTGATCTT
CAGCATCTTTTACTTTCACCAGCGTTTCTGGGTGAGCAAAAACAGGAAGGCAAAATGCCGCAAAAAAGGGAATAAG
GGCGACACGGAAATGTTGAATACTCATACTCTTCCTTTTTCAATATTATTGAAGCATTTATCAGGGTTATTGTCTC
ATGAGCGGATACATATTTGAATGTATTTAGAAAAATAAACAAATAGGGGTTCCGCGCACATTTCCCCGAAAAGTGC
CACCTGACGTCTAAGAAACCATTATTATCATGACATTAACCTATAAAAATAGGCGTATCACGAGGCCCTTTCGTCT
CGCGCGTTTCGGTGATGACGGTGAAAACCTCTGACACATGCAGCTCCCGGAGACGGTCACAGCTTGTCTGTAAGCG
GATGCCGGGAGCAGACAAGCCCGTCAGGGCGCGTCAGCGGGTGTTGGCGGGTGTCGGGGCTGGCTTAACTATGCGG
CATCAGAGCAGATTGTACTGAGAGTGCACCATATGCGGTGTGAAATACCGCACAGATGCGTAAGGAGAAAATACCG
CATCAGGCGCCATTCGCCATTCAGGCTGCGCAACTGTTGGGAAGGGCGATCGGTGCGGGCCTCTTCGCTATTACGC
CAGCTGGCGAAAGGGGGATGTGCTGCAAGGCGATTAAGTTGGGTAACGCCAGGGTTTTCCCAGTCACGACGTTGTA
AAACGACGGCCAGTGAATTCGAGCTCGGTACCTCGCGAATGCATCTAGATGCTCGTCGTCAAAAATGATCAAAAAG
GCCATTCCTATGATGAtATGGTGGTAAGCACAATGAaGAAAATGCAAGacATCaGCAGgTTGGCAGAAGTTGCTGA
AGAACTGTTTtGCATCATCATACAGCGGGAaGGgCTCTATTACCCCTTcAACAACCCcTGGCCTGCACACAAACGG
cACTACAacgTACGAAAAAaTAGCTCACTTGCATACGCTCCGAATGGAGGACCCAGCCaGTCCacTCAtgCTCAAC
AAAaaCAAaCCACAAaTcACTTACAGCTGGGTTTCAACTTTTCACTATGGAAGACGCCAAAAACATAAAGAAAGGC
CCGGCGCCATTCTATCCGCTGGAAGATGGAACCGCTGGAGAGCAACTGCATAAGGCTATGAAGAGATACGCCCTGG
TTCCTGGAACAATTGCTTTTACAGATGCACATATCGAGGTGGACATCACTTACGCTGAGTACTTCGAAATGTCCGT
TCGGTTGGCAGAAGCTATGAAACGATATGGGCTGAATACAAATCACAGAATCGTCGTATGCAGTGAAAACTCTCTT
CAATTCTTTATGCCGGTGTTGGGCGCGTTATTTATCGGAGTTGCAGTTGCGCCCGCGAACGACATTTATAATGAAC
GTGAATTGCTCAACAGTATGGGCATTTCGCAGCCTACCGTGGTGTTCGTTTCCAAAAAGGGGTTGCAAAAAATTTT
GAACGTGCAAAAAAAGCTCCCAATCATCCAAAAAATTATTATCATGGATTCTAAAACGGATTACCAGGGATTTCAG
TCGATGTACACGTTCGTCACATCTCATCTACCTCCCGGTTTTAATGAATACGATTTTGTGCCAGAGTCCTTCGATA
GGGACAAGACAATTGCACTGATCATGAACTCCTCTGGATCTACTGGTCTGCCTAAAGGTGTCGCTCTGCCTCATAG
AACTGCCTGCGTGAGATTCTCGCATGCCAGAGATCCTATTTTTGGCAATCAAATCATTCCGGATACTGCGATTTTA
AGTGTTGTTCCATTCCATCACGGTTTTGGAATGTTTACTACACTCGGATATTTGATATGTGGATTTCGAGTCGTCT
TAATGTATAGATTTGAAGAAGAGCTGTTTCTGAGGAGCCTTCAGGATTACAAGATTCAAAGTGCGCTGCTGGTGCC
AACCCTATTCTCCTTCTTCGCCAAAAGCACTCTGATTGACAAATACGATTTATCTAATTTACACGAAATTGCTTCT
GGTGGCGCTCCCCTCTCTAAGGAAGTCGGGGAAGCGGTTGCCAAGAGGTTCCATCTGCCAGGTATCAGGCAAGGAT
ATGGGCTCACTGAGACTACATCAGCTATTCTGATTACACCCGAGGGGGATGATAAACCGGGCGCGGTCGGTAAAGT
TGTTCCATTTTTTGAAGCGAAGGTTGTGGATCTGGATACCGGGAAAACGCTGGGCGTTAATCAAAGAGGCGAACTG
TGTGTGAGAGGTCCTATGATTATGTCCGGTTATGTAAACAATCCGGAAGCGACCAACGCCTTGATTGACAAGGATG
GATGGCTACATTCTGGAGACATAGCTTACTGGGACGAAGACGAACACTTCTTCATCGTTGACCGCCTGAAGTCTCT
GATTAAGTACAAAGGCTATCAGGTGGCTCCCGCTGAATTGGAATCCATCTTGCTCCAACACCCCAACATCTTCGAC
GCAGGTGTCGCAGGTCTTCCCGACGATGACGCCGGTGAACTTCCCGCCGCCGTTGTTGTTTTGGAGCACGGAAAGA
CGATGACGGAAAAAGAGATCGTGGATTACGTCGCCAGTCAAGTAACAACCGCGAAAAAGTTGCGCGGAGGAGTTGT
GTTTGTGGACGAAGTACCGAAAGGTCTTACCGGAAAACTCGACGCAAGAAAAATCAGAGAGATCCTCATAAAGGCC
AAGAAGGGCGGAAAGATCGCCGTGTAAATTGCTCACAGCATTCTCAATGCTACTACATCCGGTGTTCTTAAACACC
ACCATTCTCTTCGGAAACTGTGCCTTAGTCAAAAATTGTAAACCGATAGAAAATAAATAGAAAATGATGTTATCTA
GTGCATAAGTAGCGAGTTGGAATACGTTCCTACGTCTCCTTGTCAACACCACGATGTAATTCAAGTGATTAGCCAT
GCTTGATGGCAGATGAACAAAGGGAATTCATATTGTACATGATTAAGCGTAGTGGAGCCAGTGAAGAACTCCCACA
CCCGCCCATCCTAGCTAGTGAAC

*Naegleria gruberi*

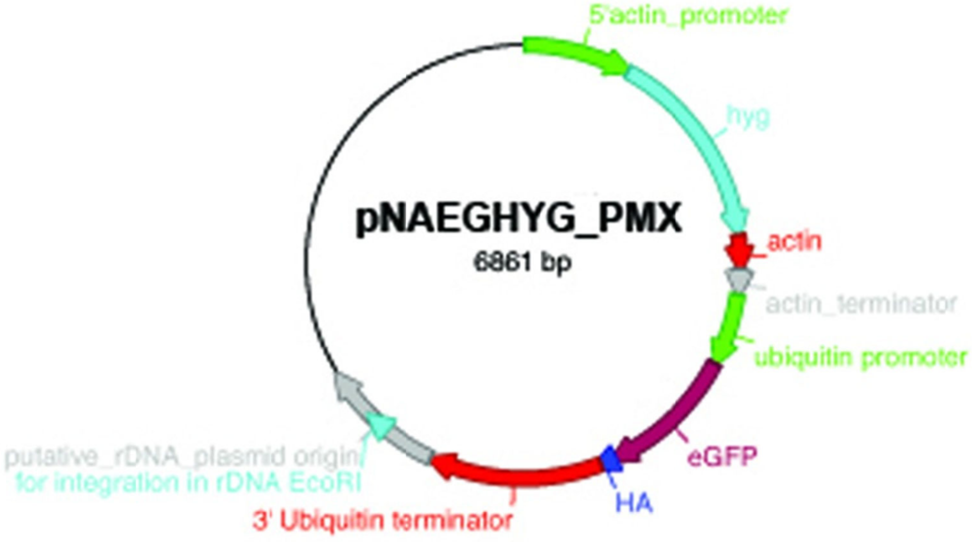


---

>pNaeHYG plasmid seqeunce
CTAAATTGTAAGCGTTAATATTTTGTTAAAATTCGCGTTAAATTTTTGTTAAATCAGCTCATTTTTTAACCAATAG
GCCGAAATCGGCAAAATCCCTTATAAATCAAAAGAATAGACCGAGATAGGGTTGAGTGTTGTTCCAGTTTGGAACA
AGAGTCCACTATTAAAGAACGTGGACTCCAACGTCAAAGGGCGAAAAACCGTCTATCAGGGCGATGGCCCACTACG
TGAACCATCACCCTAATCAAGTTTTTTGGGGTCGAGGTGCCGTAAAGCACTAAATCGGAACCCTAAAGGGAGCCCC
CGATTTAGAGCTTGACGGGGAAAGCCGGCGAACGTGGCGAGAAAGGAAGGGAAGAAAGCGAAAGGAGCGGGCGCTA
GGGCGCTGGCAAGTGTAGCGGTCACGCTGCGCGTAACCACCACACCCGCCGCGCTTAATGCGCCGCTACAGGGCGC
GTCCCATTCGCCATTCAGGCTGCGCAACTGTTGGGAAGGGCGATCGGTGCGGGCCTCTTCGCTATTACGCCAGCTG
GCGAAAGGGGGATGTGCTGCAAGGCGATTAAGTTGGGTAACGCCAGGGTTTTCCCAGTCACGACGTTGTAAAACGA
CGGCCAGTGAGCGCGCGTAATACGACTCACTATAGGGCGAATTGGAGCTCCACCGCGGTGGCGGCCGCTCTAGATC
TGACGAAAACAACATGTGGATACTCAATACTGTTGAAACCTTTCAAATAGGGACTGTGGCAAGATAAATGATCTGG
TGGTTCCAAAACACTGCTGTGATAGCAGCATTCCCAATATTTCTTATTGTGCCATCCAAAAGAGATAATTTACTAG
TAAATGATTGCATATCCTTTCACTTCAGCCTCACAAACAACTCTACAGTATCCAAAGGAAGCCTTCTCGTTTTTCC
TTTCTATTCTTGATATCCACCAGTGGCTGTATCCTTGTTCATTTCGGTTTTTCCTTCCATGAATTGTTTAATTAAG
TCCACTGTTTGATTTTTCCTTCCATGTATTGTTTAATGAAGTCCACTGTTATTGATTCTTGTTGATGGTTATTCTG
TTATTATTGTTGATGATGATGCAGTTTGTTGCCAAGAGTCAGAATTTTATGAATCATCAGACTAGGATGATACCTC
CTTTCTCTTTCTTGCCATTTTTTTTAAACATTAAATTAAACTTTGTTACAAAGGCGGGAACAAGTTCATAGTCTTA
CAAAAAAAAATGAAAAAACCAGAATTGACTGCTACTAGTGTTGAAAAATTTTTGATTGAAAAATTTGATTCTGTTT
CTGATTTGATGCAATTGTCTGAAGGTGAAGAATCAAGAGCTTTTTCTTTTGATGTTGGTGGTAGAGGTTATGTTTT
GAGAGTTAATTCTTGTGCTGATGGTTTTTATAAAGATAGATATGTTTATAGACATTTTGCTTCTGCTGCTTTGCCA
ATTCCAGAAGTTTTGGATATTGGTGAATTTTCTGAATCTTTGACTTATTGTATTTCAAGAAGAGCTCAAGGTGTTA
CTTTGCAAGATTTGCCAGAAACTGAATTGCCAGCTGTTTTGCAACCAGTTGCTGAAGCTATGGATGCTATTGCTGC
TGCTGATTTGTCTCAAACTTCTGGTTTTGGTCCATTTGGTCCACAAGGTATTGGTCAATATACTACTTGGAGAGAT
TTTATTTGTGCTATTGCTGATCCACATGTTTATCATTGGCAAACTGTTATGGATGATACTGTTTCTGCTTCTGTTG
CTCAAGCATTGGATGAATTGATGTTGTGGGCTGAAGATTGTCCAGAAGTTAGACATTTGGTTCATGCTGATTTTGG
TTCTAATAATGTTTTGACTGATAATGGTAGAATTACTGCTGTTATTGATTGGTCTGAAGCTATGTTTGGTGATTCT
CAATATGAAGTTGCTAATATTTTTTTTTGGAGACCATGGTTGGCTTGTATGGAACAACAAACTAGATATTTTGAAA
GAAGACATCCAGAATTGGCTGGTTCTCCAAGATTGAGAGCTTATATGTTGAGAATTGGTTTGGATCAATTGTATCA
ATCTTTGGTTGATGGTAATTTTGATGATGCTGCTTGGGCTCAAGGTAGATGTGATGCTATTGTTAGATCAGGTGCT
GGTACTGTTGGTAGAACTCAAATTGCTAGAAGATCAGCTGCTGTTTGGACTGATGGTTGTGTTGAAGTTTTGGCTG
ATTCTGGTAATAGAAGACCATCTACTAGACCAAGAGCTAAAGAATAAGGGCCCTGATTTAAATTATTCCAGTAATT

TTGGACGATTCTTCTTTCCTTATTGAAGCAATAAGGTAAAAAAAAAGCACCCAACAACATTGGAAAGCCACAAATGG
CTGTCTGAACACAAAAGAATGTGTAAACAGATGTTCTTTCATAACAGGAATAATTATTGAAGATGAAAACCTACAT
GATACAAAATACACACTGCACTGGATTCAAATTAAAGGTTGACAAGGTGATACAAACCACTTGTGGCCATCCTCGT
GGTCGTAAGATTGAATCAACACTTTTGTCCTGCTTTGTGAATGTCGACACTTGTTCAGTTAGTAACTAATTGGTAT
CATATCTAATACAACAGCATTGGAAGGAAAATGGATTATTACCTTACCATATCTCTCTCATCCAATCGATTGAAGT
AATGAGGTTGATGATGTTAAAAGTTGATTGGTTGTGTAAAATGGCAATAGTGTATTAATGGATTGAAAGCGAATAT
GGATGTGGAGTTGAATCTCTTCAATCTAAACGTGCTGAAGCGAGGATGAACTCACGTTGTAACAGCAAACCGCCAT
TAATTCTATTTTTAACGTCCAATAACCTATTAATACAAGTACCCATCATTGATCGTCACTACAACAAATCAAAAAA
AATCAAATAACTGCTTATTGTATACTAAAGCATATGGTGAGTAAGGGCGAGGAGCTGTTCACCGGTGTGGTGCCTA
TCCTGGTCGAGCTGGATGGCGATGTAAACGGCCACAAGTTCAGTGTGTCGGGCGAGGGCGAGGGCGATGCCACCTA
CGGCAAGCTGACCCTGAAGTTCATCTGTACCACCGGCAAGCTGCCTGTGCCTTGGCCAACCCTCGTGACCACCCTG
ACCTACGGCGTGCAATGTTTCAGTCGTTACCCCGATCACATGAAGCAACACGATTTCTTCAAGTCGGCCATGCCAG
AAGGCTACGTCCAAGAGCGTACCATCTTCTTCAAGGATGACGGCAACTACAAGACCAGAGCCGAGGTGAAGTTCGA
GGGCGATACCCTGGTGAACCGTATCGAGCTGAAGGGCATCGACTTCAAGGAGGATGGCAACATCCTGGGCCACAAG
CTGGAGTACAACTACAACAGTCACAACGTCTATATCATGGCCGATAAGCAAAAGAACGGCATCAAGGTGAACTTCA
AGATCAGACACAACATCGAGGATGGCAGTGTGCAACTCGCCGATCACTACCAACAAAACACCCCTATCGGCGACGG
CCCTGTGCTGCTGCCAGATAACCACTACCTGAGTACCCAATCAGCCCTGAGTAAAGATCCTAACGAGAAGCGTGAT
CACATGGTCCTGCTGGAGTTCGTGACCGCCGCCGGCATCACTCTCGGCATGGATGAGCTGTACAAGGGATCCTACC
CATACGATGTTCCAGATTACGCTTACCCATACGATGTTCCAGATTACGCTTAACTCGAGATAAAAACAGTTAACTT
TTGATTTCACAGTCAATTTTAATTCTTCCAAACATGATGGTCGAGCAGCTGCATTGTTTGGTCCAATATTCACTGC
CATTAATTCATTGGGTGAAATTTATATTGCTGATAGATTCGTAACGTTTTCAATAATGGTACAATTATTACAATTG
CTGGAACTGGCATAAGTGGTTACAATGGTGATTATCAAGATGCAACAAGTGCCAACTTGAATATTCCATCAGGTAT
TTTGATTGCCCCTAATGGAGAAATGATTATTTCAGATTCAGGAAGACACAGTATTCGTAGAGTATTAACCAATGGA
ACTATCACACCACTAGCTGGTGATGGAAGTGCATCTTATTGTGCAGGTATTACGGCAACTAACACTCCATTGACAA
ATCCTAGAGGAACTCTTTTTAATTCGAATGGAGATATTTATGTTGCCGAAACACAAGGTAATAGAATTAGTAGAAT
TTCAAAGAATGGAAAAGCTTTGATTGACTCTATTGCAGGTACAGGAAATTATGGTAATAATGGAGGTGGTCGGCTG
GCAACAGATACTCAATTAGCATTACCTAATTCGATTGCAACTGATCTCGATGGTCAATTACTTGTGGCAGATTCGT
ATAACCATGTCATTCGTAAAATTCTTAATAATGGAACCATGGTTATCATTGCAGGAACTGGTGTCTCAGGTTATAA
TGGAGATGGTATCGATGCAAGAAGTGCAAAATTAAACAATCCATTGTTGTTGATTCTAAAAGTGGTGAGCTCTATA
TTGCTGATAGTAATAATTATAGGATTAGAAAGATTTTAACAAATGGAACGATTGTTACAGCTGTAGGAACTGGTTC
TTCAGGTTATAATGGAGATGGCTGCAGCAATAGGACTCTCTAACAAAAAAAATTGCTTTATTTTTGTTCGCTTAAT
AAAACCCTATCAACTCGTCTTAATTGACAAGTGGTAGGGTTTTTTTTTTCGAAAATTTTTCAATCCTGACTTGGAT
ATTTTTTCAAGTTCGGGAAAGTATTTTCTTTTTTTTTAATATAATAAAACCCTATCAACTCGTCTTAATTGACAAGT
GGTAGGGTTTTTTTTTTCGAAAATTTTTCAATCCTGACTTGGATATTTTTTCAAGTTCGGGAAAGTATTTTCTTTT
TTTTAATATAATAAAACCCTATCAACGAATTCTCGTCTTAATTGACAAGTGGTAGGGTTTTTTTTTTCGAAAATTT
TTCAATCCTGACTTGGATATTTTTTCAAGTTCGGGAAAGTATTTTCTTTTTTTTTAATATAATAAAACCCTATCAAC
TCGTCTTAATTGACAAGTGGTAGGTTTTTTTTTTCGAAAATTTTTCAATCCTGACTTGGATATTTTTTCAAGTTCG
GGAAAGTATTTTCTTTTTTTTTAATATAATAAAACCCTATCAACTCGTCTTAATTGACAAGTGGTAGGGTTTTTTTTT
TCGAAAATTTTTCAATCCTGACTTGGATATTTTTTCATAATTTGGGGGATATTTTGGTACCCAGCTTTTGTTCCCT
TTAGTGAGGGTTAATTGCGCGCTTGGCGTAATCATGGTCATAGCTGTTTCCTGTGTGAAATTGTTATCCGCTCACA
ATTCCACACAACATACGAGCCGGAAGCATAAAGTGTAAAGCCTGGGGTGCCTAATGAGTGAGCTAACTCACATTAA
TTGCGTTGCGCTCACTGCCCGCTTTCCAGTCGGGAAACCTGTCGTGCCAGCTGCATTAATGAATCGGCCAACGCGC
GGGGAGAGGCGGTTTGCGTATTGGGCGCTCTTCCGCTTCCTCGCTCACTGACTCGCTGCGCTCGGTCGTTCGGCTG
CGGCGAGCGGTATCAGCTCACTCAAAGGCGGTAATACGGTTATCCACAGAATCAGGGGATAACGCAGGAAAGAACA
TGTGAGCAAAAGGCCAGCAAAAGGCCAGGAACCGTAAAAAGGCCGCGTTGCTGGCGTTTTTCCATAGGCTCCGCCC
CCCTGACGAGCATCACAAAAATCGACGCTCAAGTCAGAGGTGGCGAAACCCGACAGGACTATAAAGATACCAGGCG
TTTCCCCCTGGAAGCTCCCTCGTGCGCTCTCCTGTTCCGACCCTGCCGCTTACCGGATACCTGTCCGCCTTTCTCC
CTTCGGGAAGCGTGGCGCTTTCTCATAGCTCACGCTGTAGGTATCTCAGTTCGGTGTAGGTCGTTCGCTCCAAGCT
GGGCTGTGTGCACGAACCCCCCGTTCAGCCCGACCGCTGCGCCTTATCCGGTAACTATCGTCTTGAGTCCAACCCG
GTAAGACACGACTTATCGCCACTGGCAGCAGCCACTGGTAACAGGATTAGCAGAGCGAGGTATGTAGGCGGTGCTA
CAGAGTTCTTGAAGTGGTGGCCTAACTACGGCTACACTAGAAGGACAGTATTTGGTATCTGCGCTCTGCTGAAGCC
AGTTACCTTCGGAAAAAGAGTTGGTAGCTCTTGATCCGGCAAACAAACCACCGCTGGTAGCGGTGGTTTTTTTGTT
TGCAAGCAGCAGATTACGCGCAGAAAAAAAGGATCTCAAGAAGATCCTTTGATCTTTTCTACGGGGTCTGACGCTC
AGTGGAACGAAAACTCACGTTAAGGGATTTTGGTCATGAGATTATCAAAAAGGATCTTCACCTAGATCCTTTTAAA
TTAAAAATGAAGTTTTAAATCAATCTAAAGTATATATGAGTAAACTTGGTCTGACAGTTACCAATGCTTAATCAGT
GAGGCACCTATCTCAGCGATCTGTCTATTTCGTTCATCCATAGTTGCCTGACTCCCCGTCGTGTAGATAACTACGA
TACGGGAGGGCTTACCATCTGGCCCCAGTGCTGCAATGATACCGCGAGACCCACGCTCACCGGCTCCAGATTTATC
AGCAATAAACCAGCCAGCCGGAAGGGCCGAGCGCAGAAGTGGTCCTGCAACTTTATCCGCCTCCATCCAGTCTATT
AATTGTTGCCGGGAAGCTAGAGTAAGTAGTTCGCCAGTTAATAGTTTGCGCAACGTTGTTGCCATTGCTACAGGCA
TCGTGGTGTCACGCTCGTCGTTTGGTATGGCTTCATTCAGCTCCGGTTCCCAACGATCAAGGCGAGTTACATGATC
CCCCATGTTGTGCAAAAAAGCGGTTAGCTCCTTCGGTCCTCCGATCGTTGTCAGAAGTAAGTTGGCCGCAGTGTTA
TCACTCATGGTTATGGCAGCACTGCATAATTCTCTTACTGTCATGCCATCCGTAAGATGCTTTTCTGTGACTGGTG
AGTACTCAACCAAGTCATTCTGAGAATAGTGTATGCGGCGACCGAGTTGCTCTTGCCCGGCGTCAATACGGGATAA
TACCGCGCCACATAGCAGAACTTTAAAAGTGCTCATCATTGGAAAACGTTCTTCGGGGCGAAAACTCTCAAGGATC
TTACCGCTGTTGAGATCCAGTTCGATGTAACCCACTCGTGCACCCAACTGATCTTCAGCATCTTTTACTTTCACCA
GCGTTTCTGGGTGAGCAAAAACAGGAAGGCAAAATGCCGCAAAAAAGGGAATAAGGGCGACACGGAAATGTTGAAT
ACTCATACTCTTCCTTTTTCAATATTATTGAAGCATTTATCAGGGTTATTGTCTCATGAGCGGATACATATTTGAA
TGTATTTAGAAAAATAAACAAATAGGGGTTCCGCGCACATTTCCCCGAAAAGTGCCAC

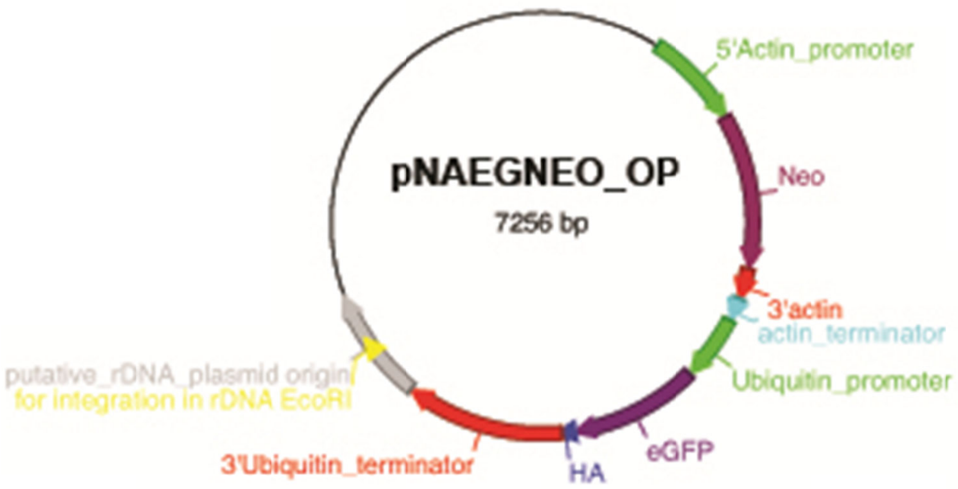


>pNaeNEO plasmid sequence
CTAAATTGTAAGCGTTAATATTTTGTTAAAATTCGCGTTAAATTTTTGTTAAATCAGCTCATTTTTTAACCAATAG
GCCGAAATCGGCAAAATCCCTTATAAATCAAAAGAATAGACCGAGATAGGGTTGAGTGTTGTTCCAGTTTGGAACA
AGAGTCCACTATTAAAGAACGTGGACTCCAACGTCAAAGGGCGAAAAACCGTCTATCAGGGCGATGGCCCACTACG
TGAACCATCACCCTAATCAAGTTTTTTGGGGTCGAGGTGCCGTAAAGCACTAAATCGGAACCCTAAAGGGAGCCCC
CGATTTAGAGCTTGACGGGGAAAGCCGGCGAACGTGGCGAGAAAGGAAGGGAAGAAAGCGAAAGGAGCGGGCGCTA
GGGCGCTGGCAAGTGTAGCGGTCACGCTGCGCGTAACCACCACACCCGCCGCGCTTAATGCGCCGCTACAGGGCGC
GTCCCATTCGCCATTCAGGCTGCGCAACTGTTGGGAAGGGCGATCGGTGCGGGCCTCTTCGCTATTACGCCAGCTG
GCGAAAGGGGGATGTGCTGCAAGGCGATTAAGTTGGGTAACGCCAGGGTTTTCCCAGTCACGACGTTGTAAAACGA
CGGCCAGTGAGCGCGCGTAATACGACTCACTATAGGGCGAATTGGAGCTCCACCGCGGTGGCGGCCGCTCTAGAAT
CTGACGAAAACAACATGTGGATACTCAATACTGTTGAAACCTTTCAAATAGGGACTGTGGCAAGATAAATGATCTG
GTGGTTCCAAAACACTGCTGTGATAGCAGCATTCCCAATATTTCTTATTGTGCCATCCAAAAGAGATAATTTACTA
GTAAATGATTGCATATCCTTTCACTTCAGCCTCACAAACAACTCTACAGTATCCAAAGGAAGCCTTCTCGTTTTTC
CTTTCTATTCTTGATATCCACCAGTGGCTGTATCCTTGTTCATTTCGGTTTTTCCTTCCATGAATTGTTTAATTAA
GTCCACTGTTTGATTTTTCCTTCCATGTATTGTTTAATGAAGTCCACTGTTATTGATTCTTGTTGATGGTTATTCT
GTTATTATTGTTGATGATGATGCAGTTTGTTGCCAAGAGTCAGAATTTTATGAATCATCAGACTAGGATGATACCT
CCTTTCTCTTTCTTGCCATTTTTTTTAAACATTAAATTAAACTTTGTTACAAAGGCGGGAACAAGTTCATAGTCTT
ACAAAAAAAAATGTCTGATCCAATCAGTTTTAGATCTGATCAAGAGACAGTAGGAGGATCGTTTAGAATGATTGAA
CAAGATGGATTGCACGCAGGATCTCCTGCCGCTTGGGTAGAGAGACTATTCGGCTATGATTGGGCACAACAAACAA
TCGGCTGTTCTGATGCCGCCGTGTTCAGGCTGTCAGCTCAAGGAAGACCAGTTCTTTTTGTAAAGACCGATCTGTC
TGGTGCCCTGAATGAACTGCAAGATGAGGCAGCTCGACTATCGTGGCTGGCCACTACAGGAGTTCCTTGTGCAGCT
GTGCTCGATGTTGTCACTGAAGCCGGAAGGGATTGGCTGCTATTGGGCGAAGTGCCAGGACAGGATCTCCTGTCAT
CTCACCTTGCTCCTGCCGAGAAAGTATCTATCATGGCTGATGCAATGAGAAGGCTGCATACTCTTGATCCTGCTAC
CTGTCCATTCGATCACCAAGCAAAACATAGAATCGAGCGAGCAAGAACTAGAATGGAAGCCGGTCTTGTCGATCAG
GATGATCTGGATGAAGAGCATCAAGGTCTCGCTCCAGCCGAACTGTTCGCCAGGCTCAAGGCCCGTATGCCAGACG
GCGAGGATCTCGTCGTGACCCATGGCGATGCCTGTTTGCCTAATATCATGGTGGAAAATGGCCGTTTTTCTGGATT
CATCGACTGTGGCCGTCTGGGTGTGGCAGACAGATATCAAGACATTGCTTTGGCTACCCGTGATATTGCTGAAGAG
CTTGGCGGCGAATGGGCTGACCGTTTCCTCGTGCTTTACGGTATCGCCGCTCCAGATTCTCAACGTATCGCCTTCT
ATCGTCTTCTTGACGAGTTCTTCTAAGGGCCCTGATTTAAATTATTCCAGTAATTTTGGACGATTCTTCTTTCCTT
ATTGAAGCAATAAGGTAAAAAAAAGCACCCAACAACATTGGAAAGCCACAAATGGCTGTCTGAACACAAAAGAATG
TGTAAACAGATGTTCTTTCATAACAGGAATAATTATTGAAGATGAAAACCTACATGATACAAAATACACACTGCAC
TGGATTCAAATTAAAGGTTGACAAGGTGATACAAACCACTTGTGGCCATCCTCGTGGTCGTAAGATTGAATCAACA
CTTTTGTCCTGCTTTGTGAATGTCGACACTTGTTCAGTTAGTAACTAATTGGTATCATATCTAATACAACAGCATT
GGAAGGAAAATGGATTATTACCTTACCATATCTCTCTCATCCAATCGATTGAAGTAATGAGGTTGATGATGTTAAA
AGTTGATTGGTTGTGTAAAATGGCAATAGTGTATTAATGGATTGAAAGCGAATATGGATGTGGAGTTGAATCTCTT
CAATCTAAACGTGCTGAAGCGAGGATGAACTCACGTTGTAACAGCAAACCGCCATTAATTCTATTTTTAACGTCCA
ATAACCTATTAATACAAGTACCCATCATTGATCGTCACTACAACAAATCAAAAAAAATCAAATAACTGCTTATTGT

ATACTAAAGCATATGGTGAGTAAGGGCGAGGAGCTGTTCACCGGTGTGGTGCCTATCCTGGTCGAGCTGGATGGCG
ATGTAAACGGCCACAAGTTCAGTGTGTCGGGCGAGGGCGAGGGCGATGCCACCTACGGCAAGCTGACCCTGAAGTT
CATCTGTACCACCGGCAAGCTGCCTGTGCCTTGGCCAACCCTCGTGACCACCCTGACCTACGGCGTGCAATGTTTC
AGTCGTTACCCCGATCACATGAAGCAACACGATTTCTTCAAGTCGGCCATGCCAGAAGGCTACGTCCAAGAGCGTA
CCATCTTCTTCAAGGATGACGGCAACTACAAGACCAGAGCCGAGGTGAAGTTCGAGGGCGATACCCTGGTGAACCG
TATCGAGCTGAAGGGCATCGACTTCAAGGAGGATGGCAACATCCTGGGCCACAAGCTGGAGTACAACTACAACAGT
CACAACGTCTATATCATGGCCGATAAGCAAAAGAACGGCATCAAGGTGAACTTCAAGATCAGACACAACATCGAGG
ATGGCAGTGTGCAACTCGCCGATCACTACCAACAAAACACCCCTATCGGCGACGGCCCTGTGCTGCTGCCAGATAA
CCACTACCTGAGTACCCAATCAGCCCTGAGTAAAGATCCTAACGAGAAGCGTGATCACATGGTCCTGCTGGAGTTC
GTGACCGCCGCCGGCATCACTCTCGGCATGGATGAGCTGTACAAGGGATCCTACCCATACGATGTTCCAGATTACG
CTTACCCATACGATGTTCCAGATTACGCTTAACTCGAGATAAAAACAGTTAACTTTTGATTTCACAGTCAATTTTA
ATTCTTCCAAACATGATGGTCGAGCAGCTGCATTGTTTGGTCCAATATTCACTGCCATTAATTCATTGGGTGAAAT
TTATATTGCTGATAGATTCGTAACGTTTTCAATAATGGTACAATTATTACAATTGCTGGAACTGGCATAAGTGGTT
ACAATGGTGATTATCAAGATGCAACAAGTGCCAACTTGAATATTCCATCAGGTATTTTGATTGCCCCTAATGGAGA
AATGATTATTTCAGATTCAGGAAGACACAGTATTCGTAGAGTATTAACCAATGGAACTATCACACCACTAGCTGGT
GATGGAAGTGCATCTTATTGTGCAGGTATTACGGCAACTAACACTCCATTGACAAATCCTAGAGGAACTCTTTTTA
ATTCGAATGGAGATATTTATGTTGCCGAAACACAAGGTAATAGAATTAGTAGAATTTCAAAGAATGGAAAAGCTTT
GATTGACTCTATTGCAGGTACAGGAAATTATGGTAATAATGGAGGTGGTCGGCTGGCAACAGATACTCAATTAGCA
TTACCTAATTCGATTGCAACTGATCTCGATGGTCAATTACTTGTGGCAGATTCGTATAACCATGTCATTCGTAAAA
TTCTTAATAATGGAACCATGGTTATCATTGCAGGAACTGGTGTCTCAGGTTATAATGGAGATGGTATCGATGCAAG
AAGTGCAAAATTAAACAATCCATTGTTGTTGATTCTAAAAGTGGTGAGCTCTATATTGCTGATAGTAATAATTATA
GGATTAGAAAGATTTTAACAAATGGAACGATTGTTACAGCTGTAGGAACTGGTTCTTCAGGTTATAATGGAGATGG
CTGCAGCAATAGGACTCTCTAACAAAAAAAATTGCTTTATTTTTGTTCGCTTAATAAAACCCTATCAACTCGTCTT
AATTGACAAGTGGTAGGGTTTTTTTTTTCGAAAATTTTTCAATCCTGACTTGGATATTTTTTCAAGTTCGGGAAAG
TATTTTCTTTTTTTTAATATAATAAAACCCTATCAACTCGTCTTAATTGACAAGTGGTAGGGTTTTTTTTTTCGAA
AATTTTTCAATCCTGACTTGGATATTTTTTCAAGTTCGGGAAAGTATTTTCTTTTTTTTAATATAATAAAACCCTA
TCAACGAATTCTCGTCTTAATTGACAAGTGGTAGGGTTTTTTTTTTCGAAAATTTTTCAATCCTGACTTGGATATT
TTTTCAAGTTCGGGAAAGTATTTTCTTTTTTTTAATATAATAAAACCCTATCAACTCGTCTTAATTGACAAGTGGT
AGGTTTTTTTTTTCGAAAATTTTTCAATCCTGACTTGGATATTTTTTCAAGTTCGGGAAAGTATTTTCTTTTTTTT
AATATAATAAAACCCTATCAACTCGTCTTAATTGACAAGTGGTAGGGTTTTTTTTTCGAAAATTTTTCAATCCTGA
CTTGGATATTTTTTCATAATTTGGGGGATATTTTGGTACCCAGCTTTTGTTCCCTTTAGTGAGGGTTAATTGCGCG
CTTGGCGTAATCATGGTCATAGCTGTTTCCTGTGTGAAATTGTTATCCGCTCACAATTCCACACAACATACGAGCC
GGAAGCATAAAGTGTAAAGCCTGGGGTGCCTAATGAGTGAGCTAACTCACATTAATTGCGTTGCGCTCACTGCCCG
CTTTCCAGTCGGGAAACCTGTCGTGCCAGCTGCATTAATGAATCGGCCAACGCGCGGGGAGAGGCGGTTTGCGTAT
TGGGCGCTCTTCCGCTTCCTCGCTCACTGACTCGCTGCGCTCGGTCGTTCGGCTGCGGCGAGCGGTATCAGCTCAC
TCAAAGGCGGTAATACGGTTATCCACAGAATCAGGGGATAACGCAGGAAAGAACATGTGAGCAAAAGGCCAGCAAA
AGGCCAGGAACCGTAAAAAGGCCGCGTTGCTGGCGTTTTTCCATAGGCTCCGCCCCCCTGACGAGCATCACAAAAA
TCGACGCTCAAGTCAGAGGTGGCGAAACCCGACAGGACTATAAAGATACCAGGCGTTTCCCCCTGGAAGCTCCCTC
GTGCGCTCTCCTGTTCCGACCCTGCCGCTTACCGGATACCTGTCCGCCTTTCTCCCTTCGGGAAGCGTGGCGCTTT
CTCATAGCTCACGCTGTAGGTATCTCAGTTCGGTGTAGGTCGTTCGCTCCAAGCTGGGCTGTGTGCACGAACCCCC
CGTTCAGCCCGACCGCTGCGCCTTATCCGGTAACTATCGTCTTGAGTCCAACCCGGTAAGACACGACTTATCGCCA
CTGGCAGCAGCCACTGGTAACAGGATTAGCAGAGCGAGGTATGTAGGCGGTGCTACAGAGTTCTTGAAGTGGTGGC
CTAACTACGGCTACACTAGAAGGACAGTATTTGGTATCTGCGCTCTGCTGAAGCCAGTTACCTTCGGAAAAAGAGT
TGGTAGCTCTTGATCCGGCAAACAAACCACCGCTGGTAGCGGTGGTTTTTTTGTTTGCAAGCAGCAGATTACGCGC
AGAAAAAAAGGATCTCAAGAAGATCCTTTGATCTTTTCTACGGGGTCTGACGCTCAGTGGAACGAAAACTCACGTT
AAGGGATTTTGGTCATGAGATTATCAAAAAGGATCTTCACCTAGATCCTTTTAAATTAAAAATGAAGTTTTAAATC
AATCTAAAGTATATATGAGTAAACTTGGTCTGACAGTTACCAATGCTTAATCAGTGAGGCACCTATCTCAGCGATC
TGTCTATTTCGTTCATCCATAGTTGCCTGACTCCCCGTCGTGTAGATAACTACGATACGGGAGGGCTTACCATCTG
GCCCCAGTGCTGCAATGATACCGCGAGACCCACGCTCACCGGCTCCAGATTTATCAGCAATAAACCAGCCAGCCGG
AAGGGCCGAGCGCAGAAGTGGTCCTGCAACTTTATCCGCCTCCATCCAGTCTATTAATTGTTGCCGGGAAGCTAGA
GTAAGTAGTTCGCCAGTTAATAGTTTGCGCAACGTTGTTGCCATTGCTACAGGCATCGTGGTGTCACGCTCGTCGT
TTGGTATGGCTTCATTCAGCTCCGGTTCCCAACGATCAAGGCGAGTTACATGATCCCCCATGTTGTGCAAAAAAGC
GGTTAGCTCCTTCGGTCCTCCGATCGTTGTCAGAAGTAAGTTGGCCGCAGTGTTATCACTCATGGTTATGGCAGCA
CTGCATAATTCTCTTACTGTCATGCCATCCGTAAGATGCTTTTCTGTGACTGGTGAGTACTCAACCAAGTCATTCT
GAGAATAGTGTATGCGGCGACCGAGTTGCTCTTGCCCGGCGTCAATACGGGATAATACCGCGCCACATAGCAGAAC
TTTAAAAGTGCTCATCATTGGAAAACGTTCTTCGGGGCGAAAACTCTCAAGGATCTTACCGCTGTTGAGATCCAGT
TCGATGTAACCCACTCGTGCACCCAACTGATCTTCAGCATCTTTTACTTTCACCAGCGTTTCTGGGTGAGCAAAAA
CAGGAAGGCAAAATGCCGCAAAAAAGGGAATAAGGGCGACACGGAAATGTTGAATACTCATACTCTTCCTTTTTCA
ATATTATTGAAGCATTTATCAGGGTTATTGTCTCATGAGCGGATACATATTTGAATGTATTTAGAAAAATAAACAA
ATAGGGGTTCCGCGCACATTTCCCCGAAAAGTGCCAC

**Opisthokonts**

*Sphaeroforma arctica*

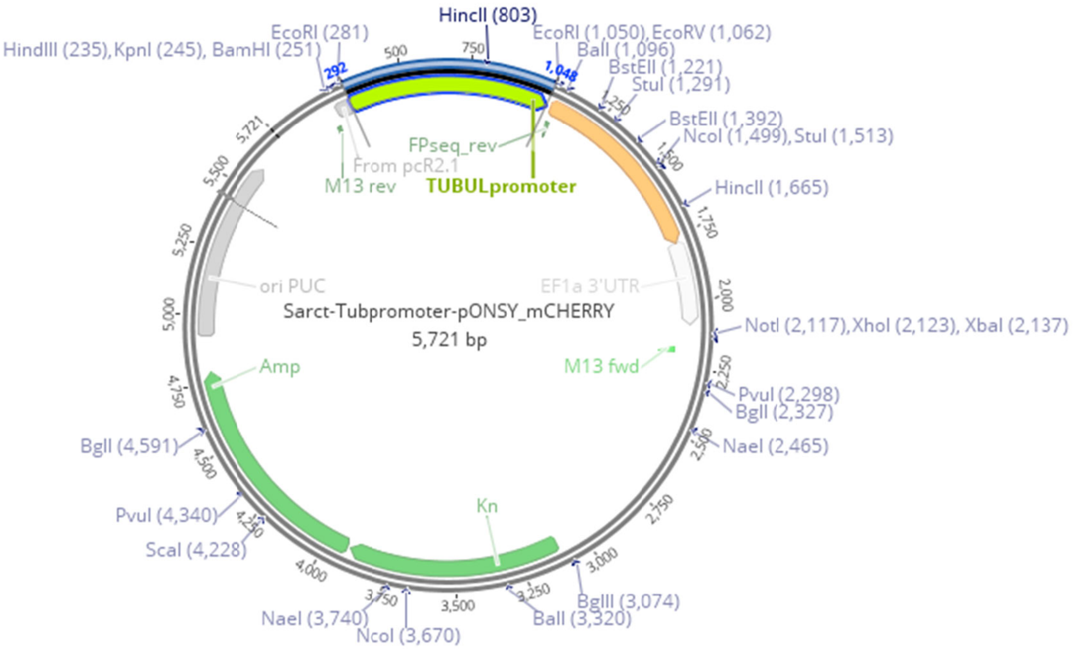


---

```
AGCGCCCAATACGCAAACCGCCTCTCCCCGCGCGTTGGCCGATTCATTAA
TGCAGCTGGCACGACAGGTTTCCCGACTGGAAAGCGGGCAGTGAGCGCAA
CGCAATTAATGTGAGTTAGCTCACTCATTAGGCACCCCAGGCTTTACACT
TTATGCTTCCGGCTCGTATGTTGTGTGGAATTGTGAGCGGATAACAATTT
CACACAGGAAACAGCTATGACCATGATTACGCCAAGCTTGGTACCGAGCG
GATCCACTAGTAACGGCCGCCAGTGTGCTGAATTCGGCTTCGTCGCTAAT
GTACACTTCTCAACCAACCTGGAGAAAATACCAGGTCAGTCAAGGGCTGT
ATAGAATTGATTACCTACGTAAACGCATGTTATATATTCACTTTAGCACA
CTATATATGCTGTCCCTTAGGCTTGAATGCGACTTGTTCCTACCCATACT
TGCTAAATACTATAGGGTACATACCGTGGTCAGGTGTGTACTCTATATGA
TACTATATATGCATACACATGCATACTACTTTGTATGCAAATAGACAGAG
AATACATACTGCTGCTTTATTTTACTTTATTAAACATCAGGATTTGCCCG
CGCAAATTTCAGGGTTTTGGGTTTTCAGATTGTTCGATATACCTACTATA
TTCTCTAATGCAAATCGCCCGAACCCATTTCCCGCGTAAACCCATAACCG
AATGCTACTGGCGGAAATTGGGACCCCAAAAATAAATGCAAATCTTGGTC
GCTCCAAAAACTTCACAGTCTAAAGAATACACTCTATTTACAGTGAACAG
TCAACAAAGCGCATGCAAAAAAAAATCATTGTTGTGCGATGAATTGCAGC
GCGTCATGGTATTATGCCTTATGGTCGTATTTGCTTCGATACCGTCTGAA
CTTGAATTTCCTGATTCAAATTAGAGAGGCGATATAAATAGTCCGAAGAT
TTGCGGAAGCCTATCAGTTGTTTCTTAGACAAACGACCATTTAATTACCT
TACCACTCTGCTTTCGTATTCGAAGCATCCACGCCTTTTATCATAGCCGA
ATTCTGCAGATATCATGGTGAGCAAGGGCGAGGAGGACAACATGGCCATC
ATCAAGGAGTTCATGCGCTTCAAGGTGCACATGGAGGGCTCCGTGAACGG
CCACGAGTTCGAGATCGAGGGCGAGGGCGAGGGCCGCCCCTACGAGGGCA
CCCAGACCGCCAAGCTGAAGGTGACCAAGGGCGGCCCCCTGCCCTTCGCC
TGGGACATCCTGTCCCCTCAGTTCATGTACGGCTCCAAGGCCTACGTGAA
GCACCCCGCCGACATCCCCGACTACTTGAAGCTGTCCTTCCCCGAGGGCT
TCAAGTGGGAGCGCGTGATGAACTTCGAGGACGGCGGCGTGGTGACCGTG
ACCCAGGACTCCTCCCTGCAGGACGGCGAGTTCATCTACAAGGTGAAGCT
GCGCGGCACCAACTTCCCCTCCGACGGCCCCGTAATGCAGAAGAAGACCA
TGGGCTGGGAGGCCTCCTCCGAGCGGATGTACCCCGAGGACGGCGCCCTG
AAGGGCGAGATCAAGCAGAGGCTGAAGCTGAAGGACGGCGGCCACTACGA
CGCCGAGGTCAAGACCACCTACAAGGCCAAGAAGCCCGTGCAGCTGCCCG
GCGCCTACAACGTCAACATCAAGCTGGACATCACCTCCCACAACGAGGAC
```

TACACCATCGTGGAACAGTACGAGCGCGCCGAGGGCCGCCACTCCACCGG
CGGCATGGACGAGCTGTACAAGTAAATTTTGTGTTTGCCAAGACACATTT
GTTTATTTCAATCATTTCTAAGACCAAAAAGGGATTTCATGATTTTTGTT
GCTTGCATTGAAGTTTTTTTTGGTTGTCATTTTTTTTGCGTTGGTTATGA
GCGCAATATATTCCATTATGGCAAGAAACACGACGAGTTGTTTTTACAGT
GGAGGGATGAGCAAGGAATGGGGAAATGGAGAACAAGTGGAATGAAATGG
TTTGACTTTGTGCGGCGAAAGTGTTGCCATTTTCAATCATGCTGCGCCTG
GCACCTAGATGCGTGTGATTGCTCATTGCTAGTGCTGTTCTCACCGCGGT
CATCCATCACACTGGCGGCCGCTCGAGCATGCATCTCTAGAGGGCCCAAT
TCGCCCTATAGTGAGTCGTATTACAATTCACTGGCCGTCGTTTTACAACG
TCGTGACTGGGAAAACCCTGGCGTTACCCAACTTAATCGCCTTGCAGCAC
ATCCCCCTTTCGCCAGCTGGCGTAATAGCGAAGAGGCCCGCACCGATCGC
CCTTCCCAACAGTTGCGCAGCCTGAATGGCGAATGGACGCGCCCTGTAGC
GGCGCATTAAGCGCGGCGGGTGTGGTGGTTACGCGCAGCGTGACCGCTAC
ACTTGCCAGCGCCCTAGCGCCCGCTCCTTTCGCTTTCTTCCCTTCCTTTC
TCGCCACGTTCGCCGGCTTTCCCCGTCAAGCTCTAAATCGGGGGCTCCCT
TTAGGGTTCCGATTTAGTGCTTTACGGCACCTCGACCCCAAAAAACTTGA
TTAGGGTGATGGTTCACGTAGTGGGCCATCGCCCTGATAGACGGTTTTTC
GCCCTTTGACGTTGGAGTCCACGTTCTTTAATAGTGGACTCTTGTTCCAA
ACTGGAACAACACTCAACCCTATCTCGGTCTATTCTTTTGATTTATAAGG
GATTTTGCCGATTTCGGCCTATTGGTTAAAAAATGAGCTGATTTAACAAA
AATTTAACGCGAATTTTAACAAAATTCAGGGCGCAAGGGCTGCTAAAGGA
AGCGGAACACGTAGAAAGCCAGTCCGCAGAAACGGTGCTGACCCCGGATG
AATGTCAGCTACTGGGCTATCTGGACAAGGGAAAACGCAAGCGCAAAGAG
AAAGCAGGTAGCTTGCAGTGGGCTTACATGGCGATAGCTAGACTGGGCGG
TTTTATGGACAGCAAGCGAACCGGAATTGCCAGCTGGGGCGCCCTCTGGT
AAGGTTGGGAAGCCCTGCAAAGTAAACTGGATGGCTTTCTTGCCGCCAAG
GATCTGATGGCGCAGGGGATCAAGATCTGATCAAGAGACAGGATGAGGAT
CGTTTCGCATGATTGAACAAGATGGATTGCACGCAGGTTCTCCGGCCGCT
TGGGTGGAGAGGCTATTCGGCTATGACTGGGCACAACAGACAATCGGCTG
CTCTGATGCCGCCGTGTTCCGGCTGTCAGCGCAGGGGCGCCCGGTTCTTT
TTGTCAAGACCGACCTGTCCGGTGCCCTGAATGAACTGCAGGACGAGGCA
GCGCGGCTATCGTGGCTGGCCACGACGGGCGTTCCTTGCGCAGCTGTGCT
CGACGTTGTCACTGAAGCGGGAAGGGACTGGCTGCTATTGGGCGAAGTGC
CGGGGCAGGATCTCCTGTCATCCCACCTTGCTCCTGCCGAGAAAGTATCC
ATCATGGCTGATGCAATGCGGCGGCTGCATACGCTTGATCCGGCTACCTG
CCCATTCGACCACCAAGCGAAACATCGCATCGAGCGAGCACGTACTCGGA
TGGAAGCCGGTCTTGTCGATCAGGATGATCTGGACGAAGAGCATCAGGGG
CTCGCGCCAGCCGAACTGTTCGCCAGGCTCAAGGCGCGCATGCCCGACGG
CGAGGATCTCGTCGTGACCCATGGCGATGCCTGCTTGCCGAATATCATGG
TGGAAAATGGCCGCTTTTCTGGATTCATCGACTGTGGCCGGCTGGGTGTG
GCGGACCGCTATCAGGACATAGCGTTGGCTACCCGTGATATTGCTGAAGA
GCTTGGCGGCGAATGGGCTGACCGCTTCCTCGTGCTTTACGGTATCGCCG
CTCCCGATTCGCAGCGCATCGCCTTCTATCGCCTTCTTGACGAGTTCTTC
TGAATTGAAAAAGGAAGAGTATGAGTATTCAACATTTCCGTGTCGCCCTT
ATTCCCTTTTTTGCGGCATTTTGCCTTCCTGTTTTTGCTCACCCAGAAAC
GCTGGTGAAAGTAAAAGATGCTGAAGATCAGTTGGGTGCACGAGTGGGTT
ACATCGAACTGGATCTCAACAGCGGTAAGATCCTTGAGAGTTTTCGCCCC
GAAGAACGTTTTCCAATGATGAGCACTTTTAAAGTTCTGCTATGTGGCGC
GGTATTATCCCGTATTGACGCCGGGCAAGAGCAACTCGGTCGCCGCATAC
ACTATTCTCAGAATGACTTGGTTGAGTACTCACCAGTCACAGAAAAGCAT
CTTACGGATGGCATGACAGTAAGAGAATTATGCAGTGCTGCCATAACCAT
GAGTGATAACACTGCGGCCAACTTACTTCTGACAACGATCGGAGGACCGA
AGGAGCTAACCGCTTTTTTGCACAACATGGGGGATCATGTAACTCGCCTT
GATCGTTGGGAACCGGAGCTGAATGAAGCCATACCAAACGACGAGCGTGA
CACCACGATGCCTGTAGCAATGGCAACAACGTTGCGCAAACTATTAACTG
GCGAACTACTTACTCTAGCTTCCCGGCAACAATTAATAGACTGGATGGAG
GCGGATAAAGTTGCAGGACCACTTCTGCGCTCGGCCCTTCCGGCTGGCTG
GTTTATTGCTGATAAATCTGGAGCCGGTGAGCGTGGGTCTCGCGGTATCA
TTGCAGCACTGGGGCCAGATGGTAAGCCCTCCCGTATCGTAGTTATCTAC
ACGACGGGGAGTCAGGCAACTATGGATGAACGAAATAGACAGATCGCTGA
GATAGGTGCCTCACTGATTAAGCATTGGTAACTGTCAGACCAAGTTTACT
CATATATACTTTAGATTGATTTAAAACTTCATTTTTAATTTAAAAGGATC
TAGGTGAAGATCCTTTTTGATAATCTCATGACCAAAATCCCTTAACGTGA
GTTTTCGTTCCACTGAGCGTCAGACCCCGTAGAAAAGATCAAAGGATCTT
CTTGAGATCCTTTTTTTCTGCGCGTAATCTGCTGCTTGCAAACAAAAAAA
CCACCGCTACCAGCGGTGGTTTGTTTGCCGGATCAAGAGCTACCAACTCT
TTTTCCGAAGGTAACTGGCTTCAGCAGAGCGCAGATACCAAATACTGTTC
TTCTAGTGTAGCCGTAGTTAGGCCACCACTTCAAGAACTCTGTAGCACCG
CCTACATACCTCGCTCTGCTAATCCTGTTACCAGTGGCTGCTGCCAGTGG
CGATAAGTCGTGTCTTACCGGGTTGGACTCAAGACGATAGTTACCGGATA
AGGCGCAGCGGTCGGGCTGAACGGGGGGTTCGTGCACACAGCCCAGCTTG
GAGCGAACGACCTACACCGAACTGAGATACCTACAGCGTGAGCTATGAGA
AAGCGCCACGCTTCCCGAAGGGAGAAAGGCGGACAGGTATCCGGTAAGCG
GCAGGGTCGGAACAGGAGAGCGCACGAGGGAGCTTCCAGGGGGAAACGCC
TGGTATCTTTATAGTCCTGTCGGGTTTCGCCACCTCTGACTTGAGCGTCG
ATTTTTGTGATGCTCGTCAGGGGGGCGGAGCCTATGGAAAAACGCCAGCA

ACGCGGCCTTTTTACGGTTCCTGGCCTTTTGCTGGCCTTTTGCTCACATG
TTCTTTCCTGCGTTATCCCCTGATTCTGTGGATAACCGTATTACCGCCTT
TGAGTGAGCTGATACCGCTCGCCGCAGCCGAACGACCGAGCGCAGCGAGT
CAGTGAGCGAGGAAGCGGAAG

#### *Abeoforma whisleri*

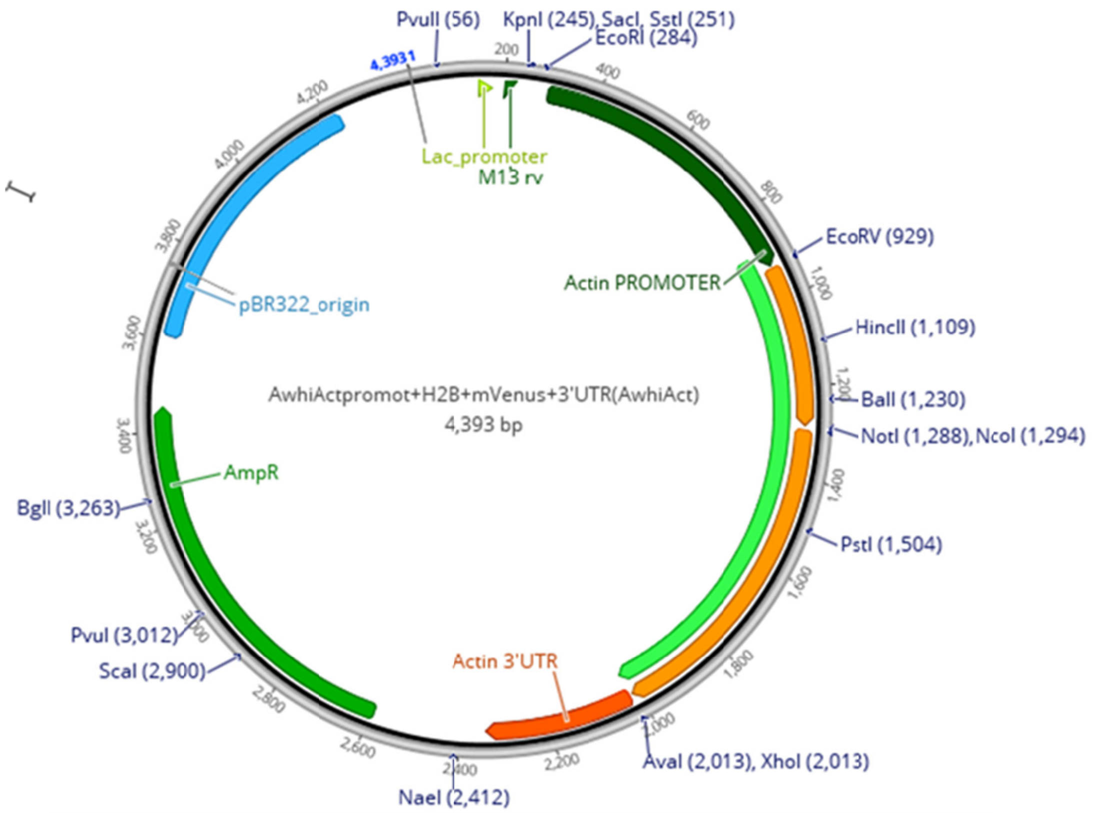


Plasmid sequence for AwH2BVenus
AwhiActpromot+H2B+mVenus+3'UTR(AwhiAct) :
AGCGCCCAATACGCAAACCGCCTCTCCCCGCGCGTTGGCCGATTCATTAATGCAGCTGGCACGACAGGTTTCCCGACTGGAAAGCG
GGCAGTGAGCGCAACGCAATTAATGTGAGTTAGCTCACTCATTAGGCACCCCAGGCTTTACACTTTATGCTTCCGGCTCGTATGTT
GTGTGGAATTGTGAGCGGATAACAATTTCACACAGGAAACAGCTATGACCATGATTACGCCAAGCTTGGTACCGAGCTCGGATCCA
CTAGTAACGGCCGCCAGTGTGCTGGAATTCGGCTTGCGCTAGCTAAATGAATAGTTAAATAGCTATAATACCGTTACGTTTAGAAA
TTTTTTATCATTCTTATATAATAAATAATTGAAAGTACAGACTGATTTTCATGATCAGTTAAATTGTTAAGGCACCTATTTATTCC
ACTCTATTTTATATGCATAGCAATCGTATGTTCGAAGTCATGTTTAAAATTGCAATTTTCAAAATTGAAAATTCAATAATTATACT
TTAGCAATTACAAATACTTAACCTTACATGTACGTTTTTACCTTTTATCGTAACCTGAAAAATCCATACAGTCTAAAATGTCTATA
TTGTGTGCCATAACAACAGTTTCCTTCTAGAAAAGAAAGAACAAAACAGAATCTTAAAAAAATCCATTGTTCAAATTCGGAGAAAT
TGAAAAAAAAAAGAAGACAAAGAAAAAGAAAGTTAATTGTCTAGCCAATCAAAAGTGAAAAAACAAAAAATGTTACAAAAAGTTAA
TTATTCATTAAAAAAAAAAAGAAAAGAGAAGAAAAAAAAATGTATAAAGGGGATCCCTATTTTTTGAATTTACAAACTATTCTAAT
TCATTGAATTATCTAAATAACATCTAATAAACATGTCTGACGAAGAAGTTGAAGCCGAATTTGCAGATATCATGCCACCTGCACCA
TCTAAAGGGGACGCTGCCAAAGGCAAACGTGGAGCTAAACCTGGAGCCAGAAAGAAAAAAAGAACAGAATCTTACTCCATCTACAT
TTACAAAGTCTTGAAACAAGTACATCCTAAAATCGGTATCTCTTCCCGTGCCATGTCTATTATGAACTCTTTTGTCAACGACATCT
TTGGCCGTGTTGCTCTTGAAGCTTCCAAGCTAGCTCAGTACAACAAGAAATCCACCATTAATGGACGCGAAATTCAAACTGCTGTT
AGATTACTTCTTCCTGGGGAATTGGCCAAACATGCTGTATCTGAAGGAACAAAAGCTGTTACTAAATACACTTCTAACAAAGCGGC
CGCCATGGTGAGCAAGGGCGAGGAGCTGTTCACCGGGGTGGTGCCCATCCTGGTCGAGCTGGACGGCGACGTAAACGGCCACAAGT
TCAGCGTGTCCGGCGAGGGCGAGGGCGATGCCACCTACGGCAAGCTGACCCTGAAGCTGATCTGCACCACCGGCAAGCTGCCCGTG
CCCTGGCCCACCCTCGTGACCACCCTGGGCTACGGCCTGCAGTGCTTCGCCCGCTACCCCGACCACATGAAGCAGCACGACTTCTT
CAAGTCCGCCATGCCCGAAGGCTACGTCCAGGAGCGCACCATCTTCTTCAAGGACGACGGCAACTACAAGACCCGCGCCGAGGTGA
AGTTCGAGGGCGACACCCTGGTGAACCGCATCGAGCTGAAGGGCATCGACTTCAAGGAGGACGGCAACATCCTGGGGCACAAGCTG
GAGTACAACTACAACAGCCACAACGTCTATATCACCGCCGACAAGCAGAAGAACGGCATCAAGGCCAACTTCAAGATCCGCCACAA
CATCGAGGACGGCGGCGTGCAGCTCGCCGACCACTACCAGCAGAACACCCCCATCGGCGACGGCCCCGTGCTGCTGCCCGACAACC
ACTACCTGAGCTACCAGTCCGCCCTGAGCAAAGACCCCAACGAGAAGCGCGATCACATGGTCCTGCTGGAGTTCGTGACCGCCGCC
GGGATCACTCTCGGCATGGACGAGCTGTACAAGCTCGAGTAAACAAATGTTTTGTTGGAAGATTTGATTTGTCTTTTTTGGAGTGC
TTAGTTAGTGCACTCCTAGATGAGCAAACATTACCACAGTGTATCGCAGCATAAAATAAAATTATAAAATGTCTTTGTAGATCAAA
AGCATTACAAACAAATATTTAAACTTTGATCACTTTTTTCAAATTCGTCAATATGACTAATTAACATCTGGGTTATAGATGACAAG
CATGAATGTTTCATTTATTTTAAAAAAAAAATAAATAAGTATTTTTTCATTTGCATCACCATCAAAAATATTTCCATTGAAAAGTC
ATCGACTGGCTATTTAAATTTCTAGAGGGCCTGCTTGCCGAATATCATGGTGGAAAATGGCCGCTTTTCTGGATTCATCGACTGTG
GCCGGCTGGGTGTGGCGGACCGCTATCAGGACATAGCGTTGGCTACCCGTGATATTGCTGAAGAGCTTGGCGGCGAATGGGCTGAC
CGCTTCCTCGTGCTTTACGGTATCGCCGCTCCCGATTCGCAGCGCATCGCCTTCTATCGCCTTCTTGACGAGTTCTTCTGAATTGA

AAAAGGAAGAGTATGAGTATTCAACATTTCCGTGTCGCCCTTATTCCCTTTTTTGCGGCATTTTGCCTTCCTGTTTTTGCTCACCC
AGAAACGCTGGTGAAAGTAAAAGATGCTGAAGATCAGTTGGGTGCACGAGTGGGTTACATCGAACTGGATCTCAACAGCGGTAAGA
TCCTTGAGAGTTTTCGCCCCGAAGAACGTTTTCCAATGATGAGCACTTTTAAAGTTCTGCTATGTGGCGCGGTATTATCCCGTATT
GACGCCGGGCAAGAGCAACTCGGTCGCCGCATACACTATTCTCAGAATGACTTGGTTGAGTACTCACCAGTCACAGAAAAGCATCT
TACGGATGGCATGACAGTAAGAGAATTATGCAGTGCTGCCATAACCATGAGTGATAACACTGCGGCCAACTTACTTCTGACAACGA
TCGGAGGACCGAAGGAGCTAACCGCTTTTTTGCACAACATGGGGGATCATGTAACTCGCCTTGATCGTTGGGAACCGGAGCTGAAT
GAAGCCATACCAAACGACGAGCGTGACACCACGATGCCTGTAGCAATGGCAACAACGTTGCGCAAACTATTAACTGGCGAACTACT
TACTCTAGCTTCCCGGCAACAATTAATAGACTGGATGGAGGCGGATAAAGTTGCAGGACCACTTCTGCGCTCGGCCCTTCCGGCTG
GCTGGTTTATTGCTGATAAATCTGGAGCCGGTGAGCGTGGGTCTCGCGGTATCATTGCAGCACTGGGGCCAGATGGTAAGCCCTCC
CGTATCGTAGTTATCTACACGACGGGGAGTCAGGCAACTATGGATGAACGAAATAGACAGATCGCTGAGATAGGTGCCTCACTGAT
TAAGCATTGGTAACTGTCAGACCAAGTTTACTCATATATACTTTAGATTGATTTAAAACTTCATTTTTAATTTAAAAGGATCTAGG
TGAAGATCCTTTTTGATAATCTCATGACCAAAATCCCTTAACGTGAGTTTTCGTTCCACTGAGCGTCAGACCCCGTAGAAAAGATC
AAAGGATCTTCTTGAGATCCTTTTTTTCTGCGCGTAATCTGCTGCTTGCAAACAAAAAAACCACCGCTACCAGCGGTGGTTTGTTT
GCCGGATCAAGAGCTACCAACTCTTTTTCCGAAGGTAACTGGCTTCAGCAGAGCGCAGATACCAAATACTGTTCTTCTAGTGTAGC
CGTAGTTAGGCCACCACTTCAAGAACTCTGTAGCACCGCCTACATACCTCGCTCTGCTAATCCTGTTACCAGTGGCTGCTGCCAGT
GGCGATAAGTCGTGTCTTACCGGGTTGGACTCAAGACGATAGTTACCGGATAAGGCGCAGCGGTCGGGCTGAACGGGGGGTTCGTG
CACACAGCCCAGCTTGGAGCGAACGACCTACACCGAACTGAGATACCTACAGCGTGAGCTATGAGAAAGCGCCACGCTTCCCGAAG
GGAGAAAGGCGGACAGGTATCCGGTAAGCGGCAGGGTCGGAACAGGAGAGCGCACGAGGGAGCTTCCAGGGGGAAACGCCTGGTAT
CTTTATAGTCCTGTCGGGTTTCGCCACCTCTGACTTGAGCGTCGATTTTTGTGATGCTCGTCAGGGGGGCGGAGCCTATGGAAAAA
CGCCAGCAACGCGGCCTTTTTACGGTTCCTGGCCTTTTGCTGGCCTTTTGCTCACATGTTCTTTCCTGCGTTATCCCCTGATTCTG
TGGATAACCGTATTACCGCCTTTGAGTGAGCTGATACCGCTCGCCGCAGCCGAACGACCGAGCGCAGCGAGTCAGTGAGCGAGGAA
GCGGAAG

*Salpingoeca rosetta*

Plasmid available at addgene, plasmid number 109096

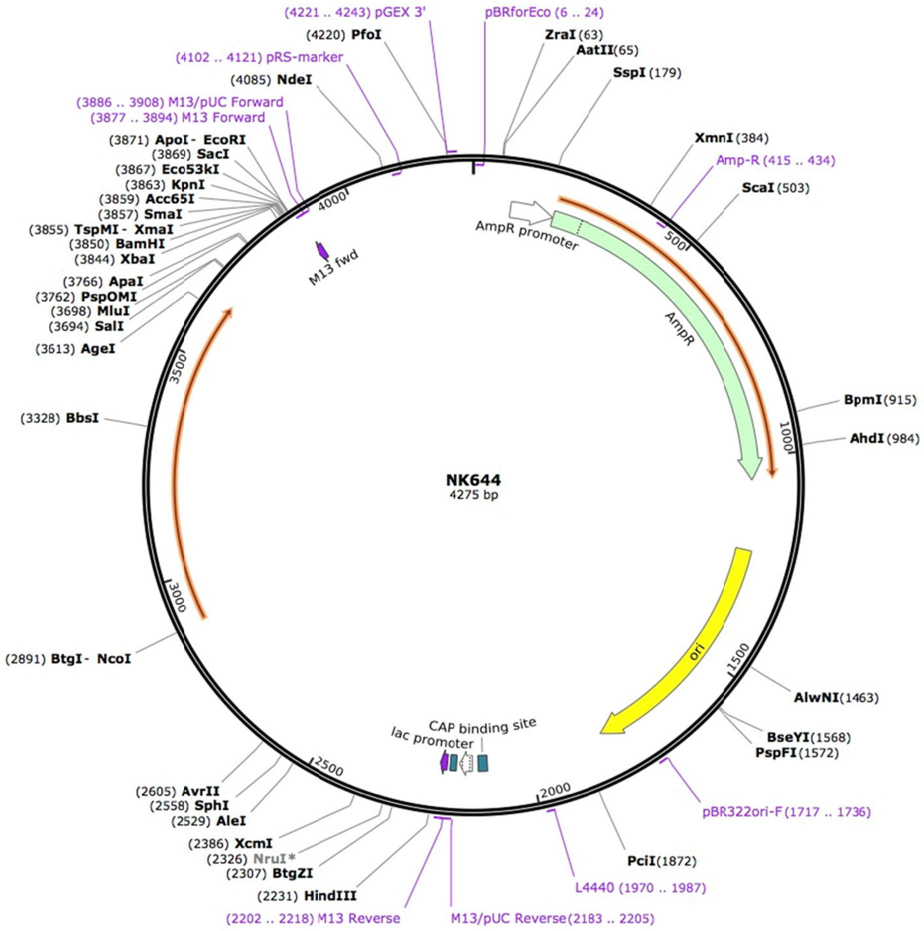


```
LOCUS       Exported                4275 bp ds-DNA     circular SYN 06-AUG-2018
DEFINITION  Reporter gene expression in S. rosetta.

ACCESSION   .

VERSION     .

KEYWORDS    NK644

SOURCE      synthetic DNA construct

  ORGANISM  synthetic DNA construct

REFERENCE   1  (bases 1 to 4275)

  AUTHORS   David Booth, Heather Middleton, Nicole King

  TITLE     A robust method for transfection in choanoflagellates illuminates

            their cell biology and the ancestry of animal septins

  JOURNAL   bioRxiv
```

```
REFERENCE   2  (bases 1 to 4275)
  AUTHORS   .
  TITLE     Direct Submission
  JOURNAL   Exported Aug 6, 2018 from SnapGene Server 1.1.58
            http://www.snapgene.com
FEATURES             Location/Qualifiers
     source          1..4275
                     /organism="synthetic DNA construct"
                     /mol_type="other DNA"
     primer_bind     complement(6..24)
                     /label=pBRforEco
                     /note="pBR322 vectors, upsteam of EcoRI site, forward
                     primer"
     promoter        92..196
                     /gene="bla"
                     /label=AmpR promoter
     CDS             197..1057
                     /codon_start=1
                     /gene="bla"
                     /product="beta-lactamase"
                     /label=AmpR
                     /note="confers resistance to ampicillin, carbenicillin, and
                     related antibiotics"
                     /translation="MSIQHFRVALIPFFAAFCLPVFAHPETLVKVKDAEDQLGARVGYI
                     ELDLNSGKILESFRPEERFPMMSTFKVLLCGAVLSRIDAGQEQLGRRIHYSQNDLVEYS
                     PVTEKHLTDGMTVRELCSAAITMSDNTAANLLLTTIGGPKELTAFLHNMGDHVTRLDRW
                     EPELNEAIPNDERDTTMPVAMATTLRKLLTGELLTLASRQQLIDWMEADKVAGPLLRSA
                     LPAGWFIADKSGAGERGSRGIIAALGPDGKPSRIVVIYTTGSQATMDERNRQIAEIGAS
                     LIKHW"
     primer_bind     complement(415..434)
                     /label=Amp-R
                     /note="Ampicillin resistance gene, reverse primer"
     rep_origin      1228..1816
                     /direction=RIGHT
                     /label=ori
                     /note="high-copy-number ColE1/pMB1/pBR322/pUC origin of
                     replication"
     primer_bind     1717..1736
                     /label=pBR322ori-F
```

/note="pBR322 origin, forward primer"

primer_bind 1970..1987

/label=L4440

/note="L4440 vector, forward primer"

protein_bind 2104..2125

/label=CAP binding site

/bound_moiety="E. coli catabolite activator protein"

/note="CAP binding activates transcription in the presence of cAMP."

promoter 2140..2170

/label=lac promoter

/note="promoter for the E. coli lac operon"

protein_bind 2178..2194

/label=lac operator

/bound_moiety="lac repressor encoded by lacI"

/note="The lac repressor binds to the lac operator to inhibit transcription in E. coli. This inhibition can be relieved by adding lactose or isopropyl-beta-D-thiogalactopyranoside (IPTG)."

primer_bind 2183..2205

/label=M13/pUC Reverse

/note="In lacZ gene"

primer_bind 2202..2218

/label=M13 rev

/note="common sequencing primer, one of multiple similar variants"

primer_bind 2202..2218

/label=M13 Reverse

/note="In lacZ gene. Also called M13-rev"

primer_bind complement(3877..3894)

/label=M13 Forward

/note="In lacZ gene. Also called M13-F20 or M13 (-21) Forward"

primer_bind complement(3877..3893)

/label=M13 fwd

/note="common sequencing primer, one of multiple similar variants"

primer_bind complement(3886..3908)

```
                     /label=M13/pUC Forward
                     /note="In lacZ gene"
     primer_bind     complement(4102..4121)
                     /label=pRS-marker
                     /note="pRS vectors, use to sequence yeast selectable
                     marker"
     primer_bind     4221..4243
                     /label=pGEX 3'
                     /note="pGEX vectors, reverse primer"
ORIGIN
        1 aaagggcctc gtgatacgcc tatttttata ggttaatgtc atgataataa tggtttctta
       61 gacgtcaggt ggcacttttc ggggaaatgt gcgcggaacc cctatttgtt tatttttcta
      121 aatacattca aatatgtatc cgctcatgag acaataaccc tgataaatgc ttcaataata
      181 ttgaaaaagg aagagtatga gtattcaaca tttccgtgtc gcccttattc ccttttttgc
      241 ggcattttgc cttcctgttt ttgctcaccc agaaacgctg gtgaaagtaa aagatgctga
      301 agatcagttg ggtgcacgag tgggttacat cgaactggat ctcaacagcg gtaagatcct
      361 tgagagtttt cgccccgaag aacgttttcc aatgatgagc acttttaaag ttctgctatg
      421 tggcgcggta ttatcccgta ttgacgccgg gcaagagcaa ctcggtcgcc gcatacacta
      481 ttctcagaat gacttggttg agtactcacc agtcacagaa aagcatctta cggatggcat
      541 gacagtaaga gaattatgca gtgctgccat aaccatgagt gataacactg cggccaactt
      601 acttctgaca acgatcggag gaccgaagga gctaaccgct tttttgcaca acatggggga
      661 tcatgtaact cgccttgatc gttgggaacc ggagctgaat gaagccatac caaacgacga
      721 gcgtgacacc acgatgcctg tagcaatggc aacaacgttg cgcaaactat taactggcga
      781 actacttact ctagcttccc ggcaacaatt aatagactgg atggaggcgg ataaagttgc
      841 aggaccactt ctgcgctcgg cccttccggc tggctggttt attgctgata aatctggagc
      901 cggtgagcgt gggtctcgcg gtatcattgc agcactgggg ccagatggta agccctcccg
      961 tatcgtagtt atctacacga cggggagtca ggcaactatg gatgaacgaa atagacagat
     1021 cgctgagata ggtgcctcac tgattaagca ttggtaactg tcagaccaag tttactcata
     1081 tatactttag attgatttaa aacttcattt ttaatttaaa aggatctagg tgaagatcct
     1141 ttttgataat ctcatgacca aaatccctta acgtgagttt tcgttccact gagcgtcaga
     1201 ccccgtagaa aagatcaaag gatcttcttg agatcctttt tttctgcgcg taatctgctg
     1261 cttgcaaaca aaaaaaccac cgctaccagc ggtggtttgt ttgccggatc aagagctacc
     1321 aactcttttt ccgaaggtaa ctggcttcag cagagcgcag ataccaaata ctgttcttct
     1381 agtgtagccg tagttaggcc accacttcaa gaactctgta gcaccgccta catacctcgc
     1441 tctgctaatc ctgttaccag tggctgctgc cagtggcgat aagtcgtgtc ttaccgggtt
     1501 ggactcaaga cgatagttac cggataaggc gcagcggtcg ggctgaacgg ggggttcgtg
     1561 cacacagccc agcttggagc gaacgaccta caccgaactg agatacctac agcgtgagct
     1621 atgagaaagc gccacgcttc ccgaagggag aaaggcggac aggtatccgg taagcggcag
     1681 ggtcggaaca ggagagcgca cgagggagct tccaggggga aacgcctggt atctttatag
```

1741 tcctgtcggg tttcgccacc tctgacttga gcgtcgattt ttgtgatgct cgtcaggggg

1801 gcggagccta tggaaaaacg ccagcaacgc ggccttttta cggttcctgg ccttttgctg

1861 gccttttgct cacatgttct ttcctgcgtt atcccctgat tctgtggata accgtattac

1921 cgcctttgag tgagctgata ccgctcgccg cagccgaacg accgagcgca gcgagtcagt

1981 gagcgaggaa gcggaagagc gcccaatacg caaaccgcct ctccccgcgc gttggccgat

2041 tcattaatgc agctggcacg acaggtttcc cgactggaaa gcgggcagtg agcgcaacgc

2101 aattaatgtg agttagctca ctcattaggc accccaggct ttacacttta tgcttccggc

2161 tcgtatgttg tgtggaattg tgagcggata acaatttcac acaggaaaca gctatgacca

2221 tgattacgcc aagcttgtaa caaactgccc tgcacctcgt ctgccacctc ggttgccacc

2281 ttgtcctcgt tccccaccac cttgtcctcg tccgctgcca gcatcgcgat cagcgtgctt

2341 gcgcttggcg ctgctgctgc cggacttccc gttatcggcc atactttttg tggtgtttcc

2401 aggcgagatg cgctgttggc aaagcaaaat ggcaaaacgg cctttgccac acgccatcct

2461 cgcctcacct cgacgtgcgg ctttcctaaa caggaaacct gacgaaacga attgagcgca

2521 ttggcacacg tgtgctgtgc gctttgtgtc gctgcatgct atccaccacg caccacccaa

2581 tcagtccaac tgtgacgtgt cacgcctagg cgaaaaagcg gcgagagcgc caaagcaaac

2641 gccaaacaaa aaccaaagaa gagcaaacgc agcgaactga tgtgagacaa gggggcggtg

2701 caatgcgtca ctgaacgtgc aaaagccaaa aacaaccaag aagcagctgc acaacacaac

2761 aacaacaaca acaacaacaa caccttgttg tcgcagctct acgtttggca ttttcctgtt

2821 gttctctctt gcgatcagtc cgtgtctctt ttctccccct tttcccagac cacaaccaca

2881 aaacaaccag ccatggtctc caagggcgag gaggacaaca tggctatcat caaggagttc

2941 atgcgcttca aggtccacat ggagggctcc gtcaacggcc acgagttcga gatcgagggc

3001 gagggcgagg gccgccccta cgagggcacc cagaccgcca agctcaaggt gacgaagggc

3061 ggcccccctcc ccttcgcctg ggacatcctc tccccccagt tcatgtacgg ctccaaggct

3121 tacgtcaagc accccgccga catccccgac tacctcaagc tctccttccc cgagggcttc

3181 aagtgggagc gcgtcatgaa cttcgaggac ggcggcgtcg tcaccgtcac ccaggactcc

3241 tctcttcagg acggcgagtt catctacaag gtcaagctgc gcggcaccaa cttcccctcc

3301 gacggccccg tcatgcagaa gaagacgatg ggttgggagg cttcctccga gcgcatgtac

3361 cccgaggacg gcgccctcaa gggcgagatc aagcagcgcc tcaagctcaa ggacggcggc

3421 cactacgacg ccgaggtcaa gaccacctac aaggccaaga agcccgtcca gctccccggc

3481 gcctacaacg tcaacatcaa gctcgacatc acctcccaca acgaggacta caccatcgtc

3541 gagcagtacg agcgtgccga gggccgccac agcacgggcg gtatggacga gctttacaag

3601 tctaccgcaa agaccggtgg tgggtgctgc acgctgctct aagcgcctcg ttggtcattg

3661 ccaccttgtt gcctcttttg ctgttgtcct cgtgtcgacg cgtgccacgt gcgcttgttg

3721 tgagttgatt ctgcttcttg tgcacgtcat gccgagccct tgggccccgc aataaaccca

3781 ctatgccagt gtaactatcc ttccccccaa tgtataactg tgacgatcaa tgagcggtgc

3841 ccgtctagag gatccccggg taccgagctc gaattcactg gccgtcgttt tacaacgtcg

3901 tgactgggaa aaccctggcg ttacccaact taatcgcctt gcagcacatc ccccttttcgc

3961 cagctggcgt aatagcgaag aggcccgcac cgatcgccct tcccaacagt tgcgcagcct

4021 gaatggcgaa tggcgcctga tgcggtattt tctccttacg catctgtgcg gtatttcaca

4081 ccgcatatgg tgcactctca gtacaatctg ctctgatgcc gcatagttaa gccagccccg

4141 acacccgcca acacccgctg acgcgccctg acgggcttgt ctgctcccgg catccgctta

4201 cagacaagct gtgaccgtct ccgggagctg catgtgtcag aggttttcac cgtcatcacc

4261 gaaacgcgcg agacg

//

#### Supplementary Notes 2

**Detailed description of Figs. 3 and 4:**

**Fig. 3. | Various methods were used to demonstrate successful transformation in different species** – Luminescence and Fluorescence.

Luminescence and Fluorescence (by FACS and epifluorescence) were used to verify expression of introduced constructs in three archaeplastids - *O. lucimarinus, B. prasinos,* and *M. commoda* (**a, b, c**).

**Fig. 4. | Various methods were used to demonstrate successful transformation in different species -** RT-PCR, Western blot and sequencing.

Western blot, RT-PCR or sequencing (in case of Cas9-induced excision by CRISPR) were used to verify expression of introduced constructs in one haptophyte – *I. galbana* (**a**), one rhizarian – *A. amoebiformis* (**b**), two stramenopiles – *F. cylindrus* and *P. tricornutum* (**c, d**), three alveolates – *K. veneficum, P. marinus* and *A. carterae* (**e, f, g**), two discobans – *B. saltans* and *D. papillatum* (**h, i**) and one opisthokont – *A. whisleri* (**j**). Note that *nptII/neo* is used synonymously with amino 3'-glycosyl phosphotransferase gene (*aph(3')*) conferring resistance to kanamycin and neomycin. Representative data of at least two independent experiments are shown. For detailed figure description see **Suppl. Notes 2**.

Luminescence, Fluorescence (by FACS and epifluorescence), Western blot, RT-PCR or sequencing (in case of Cas9-induced excision by CRISPR) were used to verify expression of introduced constructs in three archaeplastids (**3a, 3b, 3c**), one haptophyte (**4a**), one rhizarian (**4b**), two stramenopiles (**4c, 4d**), three alveolates (**4e, 4f, 4g**), two discobans (**4h, 4i**) and one opisthokont (**4j**).; for more details see below, the Results, the Online Methods and Source data. Note that *nptII/neo* is used synonymously with amino 3'-glycosyl phosphotransferase gene (*aph(3')*) conferring resistance to kanamycin and neomycin. Representative data of at least two independent experiments are shown.

(**3a**) *In vivo* luminescence of 8 *O. lucimarinus* G418-resistant transformants depicted as relative luminescence unit (RLU) per 5 s and a corresponding gel showing PCR amplification from DNA of transformants of the whole *pH4:KanMX pHAPT:luc* transgene. In total 480 transformants resistant to G418 were obtained in two independent electroporation experiments. The mean luminescence of biological triplicates is plotted for the 8 analyzed transformants exhibiting luminescence values ranging between 95 and 389 fold above the control. Note that the sd values are too low to be seen on the bar plot. A band of the expected size was amplified in all transformants and absent in WT (control cells electroporated without constructs) and the PCR negative control ($H_2O$). (**3b**) *In vivo* luminescence of 14 *B. prasinos* transformants resistant to G418 depicted as RLU per 5 s. In total 48 lines were obtained in three independent electroporation experiments. In another experiment using a different transgene (*BCCT*), 384 transformants were obtained in a single electroporation experiment (see Suppl. Fig. 1). The mean luminescence of biological triplicates is plotted for the 14 analyzed transformants. Five transformants showed luminescence values ranging between 5 and 33 fold above the control. Note that the sd values are too low to be seen on the bar plot. Corresponding gel showing PCR amplification from DNA of transformants of *pH4:KanMx* (primers 1 and 2) and *pHAPT:luc* sequences (primers 3 and 4) is shown in the lower panel. 1Kb ladder (Promega) was used in (a) and (b). (**c**) shows PCR amplification from DNA of transformants of *pH4:KanMx* (primers 1 and 2) and *pHAPT:luc* sequences (primers 3 and 4. 1Kb ladder (Promega) was used in (a) and (b). (**3c**) FACS analysis of *M. commoda* cells transformed with *eGFP* contructs or where constructs were added but no electroporation pulse applied (Wild Type). Lower panels only include the population of healthy cells selected using the depicted gates in upper panels. Bars and error bars reflect the mean and standard deviation of eGFP fluorescence and transformation efficiency from biological triplicates (n=3). Statistical analyses were performed using one-way ANOVA with an F-value of 695.241 and 11 degrees of freedom. Images show natural chlorophyll fluorescence (red, top left), eGFP (green, top right) from a transformed cell (indicated by white arrow in all panels), with DAPI DNA staining (blue, bottom left), and the overlay of all three (bottom right) demonstrating the eGFP fluorescence is localized to the nucleus as designed (see **Suppl. Fig. 15D**). A representative image from at least five experiments, each performed in biological duplicates or triplicates is shown. The scale bar represents 5 μm.

(**4a**) Expression of nourseothricin N-acetyl transferase (*NAT*) in pIgNAT transformed *I. galbana*, from cells exhibiting resistance to nourseothricin. Expression of the *NAT* transgene in WT (lane 3) and transformants (lanes 1 and 2) of *I. galbana* verified by RT-PCR. RT-negative and no template controls are shown in lanes 4 and 5, respectively, and 50 bp ladder (Bioline, lane 6), positive control PCR of genomic DNA from transformed cells (lane 7). The RT-PCR was performed once on two independently transformed *I. galbana* lines. (**4b**) Western blot of expressed eGFP in *A. amoebiformis,* with total proteins extracted from transformants (TF) and WT cells using α-GFP monoclonal antibody (Takara; 1:1,000), and an α-mouse horseradish peroxidase (HRP)-coupled antibody (GE Healthcare; 1:10,000). Precision plus protein standard (Biorad, 161-0374) was used. A representative image from two independent experiments is shown. (**4c**) Expression of *eGFP*, *ShBle* and *Rhodopsin* (endogenous gene used as control) in WT (lane 2) and transformants (1350 FC2: lane 4, 1100 FC2: lane 6) of *F. cylindrus* verified by RT-PCR. RT- negative controls are shown in lanes 3, 5 and 7 and the 100 bp ladder (NEB) is shown in lane 1. RT-PCR was carried out twice on each strain. (**4d**) Cas9-induced excision of a 38-bp region in the *cGOGAT* gene

(Phatr3_J24739). The sequence alignments from 10 *P. tricornutum* mutants (24739 KO-1 to KO-10) and two WT cell lines are shown. The two sgRNA target loci (g24739-A and g24739-B) are shown (turquoise rectangle) and include both the 20-nt sequence loci and the 3-nt PAM (protospacer-adjacent motif) sequence (yellow rectangle) and the Cas9 double-stranded break sites are also represented (dark pink rectangle). The predicted excision genotype is shown and represents the excision by dashes; the 38-bp excision is followed by an in-frame TGA stop codon. The experiment was performed twice independently. (**4e**) RT-PCR of the *nptII/neo* gene in kanamycin resistant *K. veneficum* transformants. No template negative control (lane 1); RT- negative controls of WT and transformed cells (lanes 2 and 4, respectively); cDNA from WT cells grown without antibiotics (lane 3), cDNA from transformants grown under kanamycin selection (lane 5), PCR positive control using the original vector DNA (DinoIII-*neo*, lane 6), GeneRuler DNA ladder (lane 7). (**4f**) Western blot was used to confirm the expression of MOE-GFP (40.3 kDa) in transfected *P. marinus* cells. Mock transfected cells (i.e. electroporated without plasmid) and WT were included as a control. Polyclonal rabbit α-GFP antibody (Invitrogen, 1:1000) and secondary goat α-rabbit coupled to HRP (1:10000) were used for visualization. Histone H3 (polyclonal rabbit, 1:1000, Invitrogen) was used as a loading control. SeeBlue Plus2 Pre-stained protein standard (LC5925, ThermoFisher) was used. (**4g**) RT-PCR verification of *A. carterae* transformants showing transcription from an artificial minicircle based on *atpB*. HyperLadder 1 kb plus (Bioline, lane 1), positive control PCR with artificial minicircle template (lane 2), negative control with no template (lane 3), RT-PCR against chloramphenicol acetyltransferase (*CAT*) transcript from artificial minicircle (RT+, lane 4), RT-PCR against *CAT* transcript from artificial minicircle (RT- negative control, lane 5). (**4h**) RT-PCR confirming expression of *nptII/neo* gene expression in *B. saltans* transformants. GeneRuler 1 kb DNA Ladder (Thermo Fisher Scientific); lane 1), *nptII/neo* expressed in *B. saltans* cells transformed with EF-1 alpha plasmid (lane 2); RT-negative control from transformed *B. saltans* cells (lane 3), PCR for the EF -1 alpha plasmid on RNA after DNase treatment (to verify absence of DNA, lane 4), PCR positive control using the EF- 1 alpha plasmid DNA (lane 5), negative control with no template (lane 6). NeoF and NeoR primers sequences are listed in Suppl. Fig. 10b. Results were repeated twice independently. (**4i**) Western blot of *D. papillatum* WT and C5 and C4 transformants that express the V5-tagged *nptII/neo* gene. Monoclonal mouse α-V5 antibody (Invitrogen; 1:2,000) and secondary α-mouse HRP-coupled antibody (Sigma; 1:1,000) were used. V5-tagged-mNeonGreen *Trypanosoma brucei* cells served as a positive control and mouse α-alpha-tubulin antibody (Sigma; 1:5,000) was used as a loading control. Precision plus protein standard (Biorad, 161-0374) was used. Results were repeated twice independently. (**4j**) For top and bottom gels, RT-PCR of *A. whisleri* cells transfected with either 1 μg pAwhi_H2Bvenus vector plus 10 mg carrier DNA (pUC19) (lanes 1-3), carrier DNA only (lane 4) or without DNA (lane 5), and pAwhi_H2Bvenus used as positive control (lane 7). A 1 kb DNA ladder was used (lane 6). Transfection has been repeated more than 30 times and results were repeated twice independently. All gel images are at least repeated twice with similar results.
