## Supplementary material for "Genetic tool development in marine protists: Emerging model organisms for experimental cell biology": Suppl. Table 1

**Suppl. Table 1. Overview of protists for which transformation systems have been developed prior to the EMS initiative.**

| Phylum | Species | Transformation method | Construct (promoter) | Selectable marker; selecting agent | Reporter | Reference |
|---|---|---|---|---|---|---|
| **Alveolates** | *Symbiodinium microadriaticum* | Si carbide whiskers (SiCaW) | pMT Npt/GUS (nos, CaMV35S); pMT Hpt/GUS *(Agrobacterium* [*Agro]* p1'2') | *nptII, hpt*; kanamycin, hygromycin*, G418 | Beta-glucuronidase (GUS) | ten Lohuis & Miller (1998) |
| | *Symbiodinium* spp. | Agitation with Glass beads | *pCB302-gfp-AtRACK1C*, *pCAMBIA-FABD2-gfp* (CaMV35S) | *bar, hpt; Basta*, hygromycin | Green fluorescent protein (GFP) | Ortiz-Matamoros et al. (2015a) |
| | | Glass beads and co-incubation with *Agro* | pCB302-*gfp-AtRACK1C*, pCB302-*gfp-MBD*, pCB302-*gfp-FABD2* (CaMV35S) | *bar*; Basta | Green fluorescent protein (GFP) | Ortiz-Matamoros et al. (2015b) |
| | *Amphidinium carterae* | SiCaW | pMT Npt/GUS (nos, CaMV35S); pMT Hpt/GUS (*Agro* p1'2') | *nptII, hpt*; hygromycin*, G418, kanamycin | GUS | ten Lohuis & Miller (1998) |
| **Stramenopiles** | *Phaeodactylum tricornutum* | Biolistics | (*fcp* [fucoxanthin chlorophyll-a or -c binding protein]) | *ShBle* (phleomycin/zeocin) | Luciferase (LUC) | Falciatore et al. (1999) |
| | | Biolistics | pfcpA (fcp [fucoxanthin chlorophyll-a or -c binding protein]) | *ShBle* (phleomycin/zeocin) | Chloramphenicol acetyltransferase (CAT) | Apt et al. (1996) |
| | | Electroporation | pHY11-*cat* (native NR promoter) | *cat*; chloramphenicol | CAT | Niu et al. (2012) |
| | | Conjugation | p0521S/CEN6-ARSH4-HIS3/pfcpA | *shble* | Cyan fluorescence protein (CFP), GFP, Yellow fluorescence protein (YFP) | Karas et al. (2015) |
| | *Cyclotella criptica* | Biolistics | plasmids with nptII (diatom acetyl-CoA carboxylase promoter) | *nptII;* G418 | Expression of NPTII | Dunahay et al. (1995) |

| | | | | | | |
|---|---|---|---|---|---|---|
| | *Navicular saprophila* | Biolistics | plasmids with nptII (diatom acetyl-CoA carboxylase promoter) | nptII; G419 | Expression of NPTII | Dunahay et al. (1995) |
| | *Cylindrotheca fusiformis* | Biolistics | pHUPtag-*fcp* (fucoxanthin chlorophyll-a or -c binding protein) | *ble*/*frue*, *ble*/HUPtag; zeocin, kanamycin, hygromycin B | | Fischer et al. (1999) |
| | *Conticribra (Thalassiosira) weissflogii* | Biolistics | (*fcp* [fucoxanthin chlorophyll-a or -c binding protein]) | | Luciferase (LUC) | Falciatore et al. (1999) |
| | *Pseudo-nitzschia multistriata, P. arenysensis* | Bioballistic | Histone H4 | *Shble* , zeocin | Growth on selective medium | Sabatino et al. 2015 |
| | *Thalassiosira pseudonana* | Biolistics | pTpNR/GFP (nitrate inducible NR promoter) | *Nat* (nourseothricin) | EGFP | Poulsen & Chesley (2006) |
| | *Thalassosira pseudonana* | Conjugation | pTpExpPEPCK-YFP (LHCF9 promoter_ | *Nat* (nourseothricin) | Yellow fluorescent protein (YFP) | Karas et al. (2015) |
| | *Nannochloropsis* sp. | Electroporation | *pVCP1* | *shble* | | Kilian et al. (2011) |
| | *Nannochloropsis gaditana* | Electroporation | pTUB/pHSP/pUEP/ | *shble* | | Radakovits et al. (2013) |
| | *Nannochloropsis oceanica* | Electroporation | pCMV-EM7 | *sheble* | | Osorio et al. (2019) |
| **Archaeplastids** | *Chlamydomonas reinhardtii* | Glass beads | pMN24 (containing native fragments) | *nr;* nitrate | Growth on selective medium (nitrate) | Kindle (1990) |
| | | Electroporation | pJD67 carrying ARG7 | *arg7* | Growth on corn starch | Shimogawara et al. (1998) |
| | | Si carbide whiskers | pMN24 based on pUC19 (containing native NR) | *nr*, nitrate | Growth on selective medium (nitrate) | Dunahay (1993) |
| | | Biolistics | pU12.6 (containing ASL) | argininosuccinate lyase (ASL); AS | Growth on selective medium | Debuchy et al. (1989) |

| | | | | | |
|---|---|---|---|---|---|
| | Biolistics | *p*UC19 | OEE1, oxygen-evolving enhancer protein 1 | Growth on selective medium (-acetate) | Mayfield & Kindle (1990) |
| | *Agrobacterium tumefaciens*** | T-DNA | *hpt*; hygromycin | GUS, GFP | Kumar et al. (2004) |
| *Chlorella ellipsoidea* | Biolistics | *p*DO432 | | LUC | Jarvis & Brown (1991) |
| | Electroporation | (Ubil-Ω) | | GUS | Chen et al. (2001) |
| *Chlorella saccharophila* | Electroporation | PBI221 (CaMVS35) | | GUS | Maruyama et al. (1994) |
| *Chlorella vulgaris* | Protoplast transformation | | *kanr*; G418 | Expression of human growth hormone (hGH) | Hawkins & Nakamura (1999) |
| | Electroporation | pMD18-Ap*cat* (NR promoter) | *cat*; chloramphenicol | | Niu et al. (2011) |
| *Haematococcus pluvialis* | Biolistics | *p*SV40-*LacZ* (SV40) | | β-galactosidase (LacZ) | Teng et al. (2002) |
| *Ostreococcus tauri* | Electroporation | *Potluc*<br>*Potox* | KanMX, G418<br>Nat1, cloNat | LUC | Corellou et al. [2009] |
| | Electroporation | PCR product including 1kbp of ferritin homologous sequence | KanMx, G418 | LUC (Knock in) | Lozano et al. [2014] |
| *Dunaliella salina* | Electroporation | PBI221, pUGUS, pUΩGUS, P35UΩGUS (CaMVS35, Ubil-Ω) | | GUS | Geng et al. (2003) |
| | Electroporation | | *ble*; Zeocin | | Sun et al. (2005)*** |
| | Biolistics | (CaMV35S promoter) | *bar*; Basta | GUS | Tan et al. (2005) |
| | Biolistics | *p*DM307 (native CA [carbonic anhydrase] promoter) | *bar*; Basta | Nitric oxide synthase (NOS) | Lü et al. (2005) |

*found to be most effective; ** shown to work better than glass beads; *** some introduced DNA stayed as episomal plasmid DNA

### Suppl. Table 2. Physiological and ecological properties of protists selected for this study.

| Species | Lifestyle (auto/hetero/mixo) | Axenic culture | Habitat | Cell wall | Cell structure | Life cycle | Planktonic |
|---|---|---|---|---|---|---|---|
| ***Archaeplastids*** | | | | | | | |
| *Ostreococcus lucimarinus* | photoautotroph | no | marine | no visible cell wall, just thin membrane | small (~1 µm); atypical prasinophyte (no flagella, no scales) | asexual by binary fission, possible sexual cycle | yes |
| *Bathycoccus prasinos* | photoautotroph | no | marine | organic scales | small (~2-3 µm); cells covered by scales of unknown organic material. no flagellum. | asexual by binary fission, possible sexual cycle | yes |
| *Micromonas commoda* | photoautotroph | yes (Rendered axenic using antibiotics for Worden et al. Science 2009. Tested here at start and end of each experiment using DAPI and test medium.) | marine | no visible cell wall, just thin membrane | small (~1-2 µm); single flagellum, no scales; phototactic (positive) | asexual by binary fission, possible sexual cycle | yes |
| *Micromonas pusilla* | photoautotroph | yes (Rendered axenic using antibiotics for Worden et al. Science 2009. Tested here at start and end of each experiment using DAPI and test medium.) | marine | no visible cell wall, just thin membrane | small (~1-2 µm); single flagellum; phototactic (positive); Encodes most of peptidoglycan pathway from cyanobacterial endosymbiont that formed plastid | asexual by binary fission, possible sexual cycle | yes |
| *Tetraselmis striata* | photoautotroph | no | marine coastal; most free-living but some animal symbionts | complex polysaccharide scales fuse to form a wall; cell anchored to wall by microtublues in four places | 3-25 µm, two pairs of flagella with complex scales etc covering, one large chloroplast; accumulates HUFAs | flagellated stage, vegetative no-motile stage (usually dominant), cyst | yes |
| *Pyramimonas parkeae* | photoautotroph | not tested | marine | wall-less | 20 x 15 µm | four flagella | yes |
| ***Haptophytans*** | | | | | | | |
| *Isochrysis galbana* | photoautotrophic | no | marine | thin layer of organic scales, non-calcified | 5-6µm, two flagella, 2 chloroplasts, stigma, oil droplets | Asexual by binary fission, cyst | yes |

| | | | | | | | |
|---|---|---|---|---|---|---|---|
| *Emiliana huxleyi* | photoautotrophic | no | marine, broadly distributed | calcium carbonate plates (coccoliths) | ~5 µm, no flagella | diploid calcified stage, motile non-calcified haploid stage | yes |
| ***Rhizarians*** | | | | | | | |
| *Amorphochlora (Lotharella) amoebiformis* | photomixotrophic (eat bacteria) | no | marine | none | Amoeboid cell (8-15 µm) with many filopodia. | asexual by binary fission | no |
| *Bigelowiella natans* | photomixotrophic (eat bacteria) | not tested | marine, broadly distributed | vegetative cells usually naked; this cell wall only in 'cysts' in old cultures | Amoeboid, thin filopodia (thread-like pseudopodia; sometimes reticulopodia); or coccoid cells with multilayered cell wall; or uniflagellated zoospores. store b-1,3 glucan (not starch). Mitochondria with tubular cristae. Nucleomorph genome ~400 kb (3 linear chromosomes), coding 17 plastid genes and ~350 housekeeping genes with lots of small (~20 bp) introns. Secondary endosymbiosis between cercozoan host and green algal symbiont | some species have all three cell types, many are missing one or another; which form is the main vegetative stage differs among species. Chlorarachnion may have amoeboid gamete! | yes |
| ***Stramenopiles*** | | | | | | | |
| *Fragilariopsis cylindrus* | photoautotrophic | not tested | polar oceans and sea ice; ice edge blooms | silica | ~4 µm, pennate | | yes |
| *Thalassiosira pseudonana* | photoautotrophic | yes (Bacterial growth checked in LB plates) | oceanic, coastal temperate, possible bacterial symbiont | silica and organics (long-chain polyamines, chitin, proteins) | ~5-10 µm | size reduction–restitution cycle (SRRC): asexual by binary fission accompanied by cell size reduction, cell size restored in a sexual cycle | yes |
| *Seminavis robusta* | photoautotrophic | not tested, but maintained with a cocktail of Penicillin, Ampicillin, Gentamycin, and Streptomycin | benthic | silica | ~10x50 µm; pennate | Size reduction-restitution cycle; sexual cycle with two known mating types | no |
| *Pseudo-nitzscha multiseries* | photoautotrophic | no | HAB-forming; common during coastal upwelling; significant | silica | ~5x100 µm; pennate, chain-forming | asexual by binary fission accompanied by cell size reduction, cell | yes |

| | | | | | | | |
|---|---|---|---|---|---|---|---|
| | | | player in local food web; can be toxic; neretic to open ocean | | | size restored in a sexual cycle | |
| *Heterosigma akashiwo* | photoautotrophic | no | diverse; some form HAB | naked | ~50-100 µm, heterokont flagella | | yes |
| *Aurantiochytrium limacinum* | osmoheterotrophic; stores PUFAS | yes (checked by 16S PCR, by microscopy, also, neither genomic nor transcriptomic sequencing have suggested the presence of any other organism) | marine (plant detritus) | sulfated polysaccharide scales | 4 - 20 µm; grow attached by ectoplasmic net elements or in suspension | probably asexual reproduction by zooospores | mero[1] |
| *Caecitellus sp.* | phagoheterotrophic | not tested | | none? | <5 µm; gliding motility, raptorial feeding | | yes |
| *Nannochloropsis oceanica* | photoautotroph | not tested (bought as axenic, but not retested) | marine waters worldwide | Smooth cell wall, composed of principally cellulose and algaenan, with papilla | <5 µm ovoid cells, non-motile | asexual reproduction; may be ameiotic | yes |
| *Phaeodactylum tricornutum* | photoautotrophic | yes (Provasoli antibiotic treatment, DAPI staining) | brackish and marine waters worldwide | oval morphotype silicified, other morphotypes very lightly silicified, sulfated a-mannan decorated with glucuronic residues | polymorphic (exists as oval, fusiform, triradiate, cruciform morphotype), ~5x30 µm (fusiform), ~10 µm ovoiate gliding motility (oval), can form chains | asexual by binary fission, possible sexual cycle | yes (oval type benthic) |
| ***Alveolates*** | | | | | | | |
| *Euplotes crassus* | phagoheterotrophic | no | marine | rigid pellicle | ~50-100 µm; hypotrichous ciliate | asexual by binary fission and sexual by conjugation or autogamy | yes |
| *Euplotes focardi* | phagoheterotrophic | no | marine f=antarctic | rigid pellicle | ~50-100 µm; hypotrichous ciliate | asexual by binary fission and sexual by conjugation | yes |
| *Chromera velia* | photoautotroph? | not tested | coral reef coral-associated, possibly non-facultative symbiont | thick cell wall | 7.0 × 7.6 µm, coccoid stage with no flagella | sex not observed, forms flagellate stages | mero[1] |
| *Perkinsus marinus* | osmoheterotroph, obligate parasite | yes (The ATCC repository sells it as axenic) | parasite of marine molluscs | Yes | 2-10 µm; suggested apicoplast | direct cycle, zoospores and | Both as trophozoite and |

| | | | | | | | |
|---|---|---|---|---|---|---|---|
| | | | | | | trophozoites, sex not observed | flagellated stage |
| *Oxyrrhis marina* | heterotrophic (phagotrophic; omnivorous) | no | plankton; global coastally except polar; can form blooms | naked (no theca) except has scales | 20-30 µm | | yes |
| *Hematodinium sp.* | parasite | yes | crustacean hemolymph | naked at trophont stage | Cultured as asexual multinucleated trophonts (up to 50 µm) | Complex asexual stages, non-flagellate; mononucleated sexual zoospores | Only at zoospore stage |
| *Fugacium (Symbiodinium) kawagutii* | photoautotrophic symbionts (maybe some phagotrophy?) | no (Grown under 100 µg/ml Ampicillin, 50 µg /ml Kanamycin and 50 µg/ml Streptomycin) | intra- or intercellular symbionts of marine invertebrates | naked swimming cell but cellulose wall in nonmotile phase | ~10 µm | motile (free-swimming) and non-motile (in host) phases; only the latter grows and divides | |
| *Alexandrium catenella* | photoautotroph | no (Grown under 100 µg/ml Ampicillin, 50 µg /ml Kanamycin and 50 µg/ml Streptomycin) | HAB-forming phytoplankton; produce saxitoxin (PSP) | cellulose plates | ~25 µm, usually chains of 2, 4, 8 | asexual reproduction by binary fission; sexual reproduction to form a cyst | yes |
| *Breviolum (Symbiodinium) sp.* | photoautotrophic symbionts (maybe some phagotrophy?) | not tested | intra- or intercellular symbionts of marine invertebrates | naked swimming cell but cellulose wall in nonmotile phase | ~10 µm | motile (free-swimming) and non-motile (in host) phases; only the latter grows and divides | |
| *Crypthecodinium cohnii* | heterotrophic (glucose, acetate, propionic acid); may predate via peduncle | yes (The NCMA repository sells it as axenic; the samples are routinely tested for bacterial growth in marine liquid bacterial medium) | brackish, littoral, neritic; often around macrophytes like Fucus. temperate and tropical | very thin cellulose | ~15-20 µm; produce carotenoids in the light | stores starch during log phase, DHA mainly in cysts?; swimming cells and cysts. | |
| *Amphidinium carterae* | photoautotroph | no (wild-type is not axenic, but can be grown axenically) | HAB-forming phytoplankton | naked? | ~25 µm, solitary | Asexual by binary fission | yes |
| *Karlodinium veneficum* | mixotroph (predatory) | no | HAB-forming, toxic, marine, planktonic | naked | ~10 µm | | yes |

| | | | | | | | |
|---|---|---|---|---|---|---|---|
| | | (Grown under 400 µg/ml Ampicillin) | | | | | |
| ***Discobans*** | | | | | | | |
| *Bodo saltans* | phagoheterotrophic | no (the cultures which carried bacterial populations from the original isolation were inoculated with bacteria (Klebsiella pseudomonas) as a food source | broadly distributed | none? | *B. saltans* is 4-5 µm with two flagellae | | no? *B. saltans* attaches to surfaces by tip of long (posterior) flagellum |
| *Diplonema papillatum* | phagoheterotrophic | yes (checked by microscopy, 100µg/ml of chloramphenicol added in the media) | free-living planktonic? | no pellica | ~16 µm, two flagella of equal lenth, two subapical openings | | yes |
| *Eutreptiella gymnastica* | photoautotrophic | not tested | neritic, cosmopolitan | flexible pellicle | 15-20 µm, two flagella, reddish eyespot, paramylon granules in cytoplasm, vigorous metaboly often observed | can form a cyst with layered cell wall | yes |
| *Naegleria gruberi* | phagoheterotrophic | yes (Axenic culture was tested microscopically along with DAPI straining). | wet soil and freshwater | naked | amoeboflagellate; the amoeba lacks microtubule cytoskeleton, flagellate has elaborate one including the flagella; de novo synthesis of basal body during transformation from former to latter | apparently the genome revealed two distinct haplotypes | no |
| ***Opisthokonts*** | | | | | | | |
| *Pirum gemmata* | osmoheterotroph; stores glycogen | yes (Sold axenic by ATCC) | peanut worm (*Phascolosoma agassizii*) gut contents, BC, 2004 | composition unknown but fibrous and woven; contain membrane-bound tubular extensions of the cytoplasm with tubules. | vegetative cells ~50-150 µm; large central vacuole with cytoplasm mostly pressed against periphery; multinucleate with nuclei 2 to 4 µm; sporulation happens in half an hour; endospores ~5 µm and 'weakly amoeboid' | walled cells divide internally to produce lots of endospores, which are released through parental cell wall in spurts | no |
| *Sphaeroforma arctica* | phagoheterotrophic | yes (axenic growth checked in agar plates) | invertebrate symbiont; isolated from Gammarus; arctic | carbohydrate, mostly N-acetyl-glucosamine (chitin?) | simple round cells, lots of DHA and EPA in cell membranes but not accumulated to high levels in lipid bodies | very simple growth from 5-7 µm cells for 48 hr to 35-40 µm, then | no |

| | | | | | | | |
|---|---|---|---|---|---|---|---|
| | | | | | | releasing ~120 new cells | |
| *Abeoforma whisleri* | osmoheterotroph; stores glycogen | yes (axenic growth checked in agar plates) | mussel (*Mytilus* sp.) gut contents, BC, 2007 | have both cell wall and extracellular matrix; wall sometimes very thick; composition unknown but fibrous and woven; contain membrane-bound tubular extensions of the cytoplasm with tubules. | vegetative cells mostly spherical ~50 µm but some plasmodial; endospores, plasmodia, and hyphae-like structures observed; say dispersal amoebae 'function in post reproductive dispersal' and are uninucleate | walled spherical cells, plasmodia, amoebae; asexual reproduction by dispersal amoebae, endospores, binary fission and budding. | no |
| *Salpingoeca rosetta* | phagoheterotrophic | no | free-living planktonic and benthic thecate | a proteinaceous and polysaccharide matrix | cell body 3-10 µm; single apical flagellum surrounded by a collar of 30-40 actin-filled microvilli | asexual by longitudinal fission; sexual reproduction triggered by nutrient deprivation and a secreted chondroitin lyase from *Vibrio* bacteria; dynamic life history includes unicellular and multicellular forms | yes |

[1] Meroplanktonic organisms spend only a portion of their lives as plankton.

**Suppl. Table 4. Transformation methods applied in this study.**
A. Electroporation, B. Biolistics, C. Microinjection, D. Chemical transformation, E. Conjugation and F. Glass bead abrasion conditions.

### A. Electroporation

| Species | Electroporation devices (type of transformation) | Used program/setting | Survival rate |
|---|---|---|---|
| **Archaeplastids** | | | |
| *Ostreococcus lucimarinus* | Gene Pulser Xcell | 1200V, 25μF | 30-50% |
| *Bathycoccus prasinos* | Gene Pulser Xcell | 1500V, 25μF | 50% |
| *Micromonas commoda* | LONZA® | SF Buffer with pulse EH-100, EO-100, EN-138 and EW-113 | 10-11% |
| *Micromonas pusilla* | BioRad Gene Pulser | 1000V, 10μF, 400Ω<br>800V, 25μF,400Ω<br>600V, 50μF,400Ω | n/a |
| *Pyramimonas parkeae* | Gene Pulser Xcell | 1500V, 25μF | n/a |
| | Gene Pulser Xcell | 300V, 500μF | n/a |
| | Gene Pulser Xcell | 2500V, 25μF | n/a |
| | Gene Pulser Xcell | 310V, 960μF | n/a |
| | Gene Pulser Xcell | 420V, 960μF | n/a |
| | Gene Pulser Xcell | 300V, 200μF | n/a |
| | Gene Pulser Xcell | 100V, 100μF | n/a |
| | Gene Pulser Xcell | 100V, 300μF | n/a |
| | Amaxa Nucleofector II | preset program X-001 | n/a |
| | Amaxa Nucleofector II | preset program T-020 | n/a |
| | Amaxa Nucleofector II | preset program T-023 | n/a |
| | Amaxa Nucleofector II | preset program U-035 | n/a |
| **Rhizarians (Chlorarachniophytes)** | | | |
| *Amorphochlora (Lotharella) amoebiformis* | Gene Pulser Xcell | 120 V, 25 ms square wave, 0.2 cm cuvette | 20-30% |
| *Bigelowiella natans* | Amaxa Nucleofector II | preset program X-001 | n/a |
| | Amaxa Nucleofector II | preset program T-020 | n/a |
| | Amaxa Nucleofector II | preset program T-023 | n/a |
| | Amaxa Nucleofector II | preset program U-035 | n/a |
| | Gene Pulser Xcell | 7× poring pulse: 300V, 5ms; 5× transfer pulse: 10V, 50ms | n/a |
| | Gene Pulser Xcell | 7× poring pulse: 250V, 5ms; 5× transfer pulse: 10V, 50ms | n/a |
| | Gene Pulser Xcell | 7× poring pulse: 350V, 5ms; 5× transfer pulse: 10V, 50ms | n/a |
| **Stramenopiles (Diatoms, Bacillariophytes, Raphidophytes)** | | | |

| | | | |
|---|---|---|---|
| *Seminavis robusta* | BTX ECM 2001 | Single pulse conditions: 5 V AC for 5 sec, 300 V pulse for 5, 2.5, 0.5, or 0.1 msec.<br>5 pulse conditions: 5 V AC for 5 sec, 300 V pulse for 5, 2, 1, or 0.1 msec<br>10 pulse conditions: 5 V AC for 5 sec, 300 or 500 V pulse for 0.1 or 0.05 msec (only for 500 V) | n/a |
| | NEPA21 | Poring pulse: 150, 200, 225, 250, 275, or 300 V, 10% decay rate; 5 ms length, 50 ms interval<br>Transfer pulse: 8 V, 40% decay rate, 50 ms length, 50 ms interval, alternating polarity | n/a |
| *Heterosigma akashiwo* | Gene Pulser Xcell | 50/75 V, 25 μF and ∞Ω | 80% |
| *Aurantiochytrium limacinum* | Gene Pulser (BIO-RAD) | 2 pulses: 450V, 25 μF, 1000 Ω (~5 ms) | 2.64% |
| | NEPA | 2 pulses: 250 V, 4 ms | 2.48% |
| | NEPA | 2 pulses: 275 V, 8 ms | 4.13% |
| | NEPA | 2 pulses: 300 V, 12 ms | 7.27% |
| *Nannochloropsis oceanica* | Gene Pulser II | 1800V, 50 μF, 20 ms | n/a |
| | Gene Pulser II | 1000V, 50 μF, 20 ms | n/a |
| **Alveolates** | | | |
| *Euplotes crassus* | BioRad Gene Pulser | 200V, 25μF, 100Ω | 50-60% |
| *Chromera velia* | Amaxa Nucleofector II | preset programs X-001 | n/a |
| | Amaxa Nucleofector II | preset program T-020 | n/a |
| | Amaxa Nucleofector II | preset program T-023 | n/a |
| | Gene Pulser Xcell | 1500V, 25μF | n/a |
| | Gene Pulser Xcell | 300V, 500μF | n/a |
| | Gene Pulser Xcell | 2500V, 25μF | n/a |
| | Gene Pulser Xcell | 310V, 960μF | n/a |
| *Perkinsus marinus* | Amaxa Nucleofector II | preset program D-023 | n/a |
| *Oxyrrhis marina* | BioRad's MicroPulser | DIC | n/a |
| | BioRad's MicroPulser | DIC -2 shocks | n/a |
| | BioRad's MicroPulser | SHS | n/a |
| | BioRad's MicroPulser | SC2 | n/a |
| *Hematodinium sp.* | Amaxa Nucleofector II | D-023 and X-001 | <1% |
| *Fugacium kawagutii* | BioRad's MicroPulser | DIC | n/a |
| | BioRad's MicroPulser | DIC -2 shocks | n/a |
| | BioRad's MicroPulser | SHS | n/a |

| | | | |
|---|---|---|---|
| | BioRad's MicroPulser | SC2 | n/a |
| *Alexandrium catenella* | BioRad's MicroPulser | DIC | n/a |
| | BioRad's MicroPulser | DIC -2 shocks | n/a |
| | BioRad's MicroPulser | SHS | n/a |
| | BioRad's MicroPulser | SC2 | n/a |
| *Breviolum (Symbiodinium) sp.* | NePA21 Electro-kinetic transfection system | Poring pulse: 300V, 10ms<br>Transfer pulse: 8V, 50ms | n/a |
| *Crypthecodinium cohnii* | Amaxa Nucleofector II | preset program D-023 | 79% |
| | Amaxa Nucleofector II | preset program A-020 | 85% |
| | Amaxa Nucleofector II | preset program T-020 | 67% |
| | Amaxa Nucleofector II | preset program T-030 | 53% |
| | Amaxa Nucleofector II | preset program X-001 | 62% |
| | Amaxa Nucleofector II | preset program X-001 | 71% |
| | Amaxa Nucleofector II | preset program L-029 | 77% |
| | Amaxa Nucleofector II | preset program X-003 | n/a |
| | Amaxa Nucleofector II | preset program X-005 | n/a |
| | Amaxa Nucleofector II | preset program X-033 | n/a |
| | Amaxa Nucleofector II | preset program Y-003 | n/a |
| | Amaxa Nucleofector II | preset program Y-005 | n/a |
| | Amaxa Nucleofector II | preset program Y-033 | n/a |
| | Amaxa Nucleofector II | preset program Z-007 | n/a |
| | Amaxa Nucleofector II | preset program Z-023 | n/a |
| | Amaxa Nucleofector II | preset program Z032 | n/a |
| | Amaxa Nucleofector II | preset program BAC1 | n/a |
| | Amaxa Nucleofector II | preset program BAC2 | n/a |
| | Amaxa Nucleofector II | preset program BAC3 | n/a |
| | Amaxa Nucleofector II | preset program BAC4 | n/a |
| | Amaxa Nucleofector II | preset program BAC5 | n/a |
| | Amaxa Nucleofector II | preset program BAC6 | n/a |
| | Amaxa Nucleofector II | preset program BAC7 | n/a |
| | Amaxa Nucleofector II | preset program L-029 | n/a |
| | Amaxa Nucleofector II | preset program L-029 | n/a |
| | Microfluidics | Straight 313 V | n/a |
| | Microfluidics | Straight 625 V | n/a |
| | Microfluidics | Straight 938 V | n/a |
| | Microfluidics | Straight 1250V | n/a |
| | Microfluidics | Straight 1563 V | n/a |
| | Microfluidics | Divergent 333 V | n/a |
| | Microfluidics | Divergent 667 V | n/a |
| | Microfluidics | Divergent 1000 V | n/a |
| | Microfluidics | Divergent 1333 V | n/a |
| | Microfluidics | Divergent 1667 V | n/a |

| | | | |
|---|---|---|---|
| | Microfluidics | Divergent 5000 V | n/a |
| | Lipofectamine | | 90-100% |
| *Amphidinium carterae* | Amaxa Nucleofector 4D | X-100, D-023, L-029 and EH 100. | n/a |
| | NEPA electroporator | Poring pulse: 150V, length: 5 ms, interval 50 ms, number 7<br>Transfer pulse: 8V, length 50 ms, interval 50 ms, number 5 | n/a |
| | NEPA electroporator | Poring pulse: 200V, length: 5 ms, interval 50 ms, number 7<br>Transfer pulse: 8V, length 50 ms, interval 50 ms, number 5 | n/a |
| | NEPA electroporator | Poring pulse: 225V, length: 5 ms, interval 50 ms, number 7<br>Transfer pulse: 8V, length 50 ms, interval 50 ms, number 5 | n/a |
| | NEPA electroporator | Poring pulse: 250V, length: 5 ms, interval 50 ms, number 7<br>Transfer pulse: 8V, length 50 ms, interval 50 ms, number 5 | n/a |
| | NEPA electroporator | Poring pulse: 275V, length: 5 ms, interval 50 ms, number 7<br>Transfer pulse: 8V, length 50 ms, interval 50 ms, number 5 | n/a |
| | NEPA electroporator | Poring pulse: 300V, length: 5 ms, interval 50 ms, number 1<br>Transfer pulse: 8V, length 50 ms, interval 50 ms, number 5 | n/a |
| | NEPA electroporator | Poring pulse: 300V, length: 5 ms, interval 50 ms, number 4<br>Transfer pulse: 8V, length 50 ms, interval 50 ms, number 5 | n/a |
| | NEPA electroporator | Poring pulse: 300V, length: 5 ms, interval 50 ms, number 7<br>Transfer pulse: 8V, length 50 ms, interval 50 ms, number 5 | n/a |
| | NEPA electroporator | Poring pulse: 300V, length: 5 ms, interval 50 ms, number 9<br>Transfer pulse: 8V, length 50 ms, interval 50 ms, number 5 | n/a |
| *Karlodinium veneficum* | BioRad's MicroPulser | DIC | n/a |
| | BioRad's MicroPulser | DIC -2 shocks | n/a |
| | BioRad's MicroPulser | SHS | n/a |
| | BioRad's MicroPulser | SC2 | n/a |
| **Discobans (Euglenozoans and Heteroloboseans)** | | | |
| *Bodo saltans* | NePA21 Electro-kinetic transfection system | Poring pulse: 250V, 25ms<br>Transfer pulse: 60V, 99ms | 30-50% |
| *Diplonema papillatum* | BTX | 1600V, 25W, 50mF | 10 % |
| | Amaxa Nucleofector II | preset program X-001 | 80-90 % |
| | Amaxa Nucleofector II | preset program X-014 | 40-50 % |

| | | | |
|---|---|---|---|
| *Eutreptiella gymnastica* | Gene Pulser Xcell | 1500V, 25µF | n/a |
| | Gene Pulser Xcell | 300V, 500µF | n/a |
| | Gene Pulser Xcell | 2500V, 25µF | n/a |
| | Gene Pulser Xcell | 310V, 960µF | n/a |
| | Gene Pulser Xcell | 420V, 960µF | n/a |
| | Gene Pulser Xcell | 300V, 200µF | n/a |
| | Gene Pulser Xcell | 100V, 100µF | n/a |
| | Gene Pulser Xcell | 100V, 300µF | n/a |
| | Gene Pulser Xcell | 200V, 100µF | n/a |
| | Gene Pulser Xcell | 200V, 300µF | n/a |
| | Gene Pulser Xcell | 350V, 1000µF | n/a |
| | Gene Pulser Xcell | 7× poring pulse: 300V, 5ms; 5× transfer pulse: 10V, 50ms | n/a |
| | Gene Pulser Xcell | 7× poring pulse: 250V, 5ms; 5× transfer pulse: 10V, 50ms | n/a |
| | Amaxa Nucleofector II | preset program X-001 | n/a |
| | Amaxa Nucleofector II | preset program T-020 | n/a |
| | Amaxa Nucleofector II | preset program T-023 | n/a |
| *Naegleria gruberi* | BioRad Gene Pulser xCell | 175V, 500µF, 400Ω | 10-20 % |
| | Amaxa Nucleofector II | preset program X-29 | 40-50% |
| **Opisthokonts** | | | |
| *Sphaeroforma arctica* | Neon | 1000-2500V, 10-40 ms, 1-3 pulses | n/a (not successful) |
| | Lipofectamina | | n/a (not successful) |
| | LONZA® | 16 preset codes P3/P4/P5 buffer | n/a (not successful) |
| *Abeoforma whisleri* | Neon (invitrogen) | 1300V, 25ms, pulse | 60% |
| | LONZA® | preset program EN-138 P3 buffer | 70% |
| | CaCl+Glycerol | | n/a (not successful) |
| | Lipofectamine | | 100% (not successful) |
| *Salpingoeca rosetta* | LONZA® | SF Buffer with pulse CM156 | 50% |

### B. Biolistics

| Species | Biolistics device | Settings | Survival rate |
|---|---|---|---|
| **Archaeplastids** | | | |
| *Tetraselmis striata* | Bio-Rad Biolistic PDS-1000/He Particle Delivery System | 0.6µm AuNPs, rupture disc 1550 or 2000 psi, 6cm gap | n/a |
| *Pyramimonas parkeae* | PDS-1000/He | 0.6 or 1µm AuNPs, rupture disc 1350 psi, 6cm gap | n/a |

| | | | |
|---|---|---|---|
| **Haptophytes** | | | |
| *Isochrysis galbana* | PDS-1000/He | 0.7µm Tungsten beads rupture disc 1350 psi, 6cm gap | n/a |
| **Rhizarians (Chlorarachniophytes)** | | | |
| *Amorphochlora (Lotharella) amoebiformis* | PDS-1000/He | 1 µm AuNPs, rupture disc 450 psi, 4cm gap | n/a |
| *Bigelowiella natans* | PDS-1000/He | 0.6µm AuNPs, rupture disc 1350 psi, 6cm gap | n/a |
| **Stramenopiles (Diatoms, Bacillariophytes, Raphidophytes)** | | | |
| *Fragilariopsis cylindrus* | PDS-1000/He | 0.7µm Tungsten beads, rupture disc 1550 psi, 6cm gap | n/a |
| *Seminavis robusta* | PDS-1000/He | 0.55 µm AuNPs or 1.1 µm WNPs, 1550 psi, 3 µg/mL DNA non-linearized and linearized (but not CIP-treated), at 3, 6, 9, and 12 cm gap distances | n/a |
| **Alveolates** | | | |
| *Euplotes crassus* | Bio-Rad Biolistic PDS-1000/He Particle Delivery System | 0.6 µm or 1.6 µm AuNPs, rupture disk 1550 psi, helium pressure 1750 psi, vacuum 26 inches Hg, gap distance ⅜ inches, in 10 mM HEPES pH 7.4 | 80-90% |
| *Chromera velia* | PDS-1000/He | 0.6µm AuNPs, rupture disc 1350 psi, 6cm gap | n/a |
| *Hematodinium* sp. | Bio-Rad Biolistic PDS-1000/He Particle Delivery System | rupture disk 1550 psi<br>550 nm diameter gold particles | 10% |
| *Fugacium kawagutii** | Bio-Rad Biolistic PDS-1000/He Particle Delivery System | 0.7 or 1.1 µm Tungsten; rupture disc 450, 650, 900, 1100, 1350, 1550 psi; vacuum 28 inches Hg; 7.5cm gap | n/a |
| *Alexandrium* catenella* | Bio-Rad Biolistic PDS-1000/He Particle Delivery System | 0.7 or 1.1 µm Tungsten; rupture disc 450, 650, 900, 1100, 1350, 1550 psi; vacuum 28 inches Hg; 7.5cm gap | n/a |
| *Crypthecodinium cohnii* | Bio-Rad Biolistic PDS-1000/He Particle Delivery System | rupture disk 1550 psi<br>550 nm diameter gold particles | 70-80% |
| *Amphidium carterae* | Bio-Rad Biolistics PDS-1000/He | rupture disk 1550 psi<br>550 nm diameter gold particles | n.d. |
| **Discobans (Euglenozoans and Heteroloboseans)** | | | |
| *Eutreptiella gymnastica* | PDS-1000/He | 0.6 or 1µm AuNPs, rupture disc 1350 psi, 6cm gap | n/a |

*This part of work was assisted by Kaidian Zhang from Xiamen University, China.

### C. Microinjection

| Species | Microinjection device | Used setting | Survival rate |
|---|---|---|---|
| **Alveolates** | | | |
| *Euplotes crassus* | Eppendorf InjectMan NI 2 | With Eppendorf Femtotips Microinjection Capillary Tip | 2-10% |

### D. Chemical transformation

| Species | Transfection reagent | Used setting | Survival rate |
|---|---|---|---|
| **Archaeplastids** | | | |
| *Pyramimonas parkeae* | Lipofectamine® 3000 Transfection Reagent (Invitrogen) | DNA–Lipofectamine complex prepared according to the supplier. | n/a |
| **Rhizarians (Chlorarachniophytes)** | | | |
| *Bigelowiella natans* | Lipofectamine® 3000 Transfection Reagent (Invitrogen) | DNA–Lipofectamine complex prepared according to the supplier. | n/a |
| **Alveolates** | | | |
| *Euplotes crassus* | Lipofectamine® 2000 or Lipofectamine® 3000 Transfection Reagent (Invitrogen) | DNA–Lipofectamine complex prepared according to the supplier. | 100% |
| | Effectene Transfection Reagent (QIAGEN) | DNA–Effectene complex prepared according to the supplier with a double amount of DNA | 10-20% |
| | FuGENE HD Transfection Reagent (Promega) | DNA–FuGENE complex prepared according to the supplier. | 50-60% |
| **Discobans (Euglenozoans and Heteroloboseans)** | | | |
| *Eutreptiella gymnastica* | Lipofectamine® 3000 Transfection Reagent (Invitrogen) | DNA–Lipofectamine complex prepared according to the supplier. | n/a |

### E. Conjugation

| Species | Coincugation (species co-incubated) | Survival rate | Efficiency |
|---|---|---|---|
| **Stramenopiles (Diatoms, Bacillariophytes, Raphidophytes)** | | | |
| *Thalassiosira pseudonana* | *E. coli* EPI300 | n/a | ~10% |
| *Heterosigma akashiwo* | Agrobacterium | 10-15% | n/a |
| **Alveolates** | | | |
| *Oxyrrhis marina* | *E. coli* | 100% | 1-5% |
| *Karlodinium veneficum* | *E. coli* | 100% | n/a |

| | | | |
|---|---|---|---|
| *Alexandrium catenella* | *E. coli* | 100% | n/a |

### F. Glass bead abrasion

| Species | Co-incubation (species co-incubated) | Survival rate | Efficiency |
|---|---|---|---|
| **Stramenopiles (Diatoms, Bacillariophytes, Raphidophytes)** | | | |
| *Heterosigma akashiwo* | n/a | 80% | n/a |
| **Alveolates** | | | |
| *Perkinsus marinus* | n/a | 80-90% | 0.01%-1% |
| *Hematodinium sp.* | n/a | 40-50% | 0% |
| *Amphidinium carterae* | None | No data | 0% |
| | Polyethylene glycol | No data | 0% |

**Suppl. Table 5: List of protists selected for this study including links to their transformation protocols (protocols.io) and vector sequences.** For contacting particular laboratories, see Suppl. Table 6. For the vector sequences and maps, see Suppl. Notes 1.

| Species | Source of organism/ Strain/Culture collection number | Principal Investigator (PI) | Other Investigators | protocols.io links (including construct maps and their sequences) | Sequences submitted (Accession No.) / published |
|---|---|---|---|---|---|
| **Archaeplastids** | | | | | |
| *Ostreococcus lucimarinus* | RCC802 | François-Yves Bouget | Jean-Claude Lozano<br>Valérie Vergé | https://www.protocols.io/view/selection-of-stable-transformants-in-ostreococcus-zj2f4qe<br>https://www.protocols.io/view/transient-luciferase-expression-in-ostreococcus-ot-hcib2ue<br>https://www.protocols.io/view/transient-transformation-of-ostreococcus-species-o-g86bzze<br>http://dx.doi.org/10.17504/protocols.io.g86bzze | *p*HAPT:luc vector (Djouani-Tahri *et al.*, 2011) was used as a template for preparation of linear construct |
| *Bathycoccus prasinos* | RCC4222 | François-Yves Bouget | Jean-Claude Lozano<br>Valérie Vergé | https://www.protocols.io/view/selection-of-stable-transformants-in-ostreococcus-zj2f4qe<br>http://dx.doi.org/10.17504/protocols.io.hcib2ue<br>http://dx.doi.org/10.17504/protocols.io.g86bzze | *p*HAPT:luc vector (Djouani-Tahri *et al.*, 2011)was used as a template for preparation of linear construct |
| *Micromonas commoda*[1] | CCMP 2709 (genome sequenced, axenic version of RCC299) | Alexandra Z. Worden | Manuel Ares<br>Jian Guo<br>Lisa Sudek | http://dx.doi.org/10.17504/protocols.io.8p9hvr6<br>http://dx.doi.org/10.17504/protocols.io.8p8hvrw | |
| *Micromonas pusilla* | CCMP 1545 | François-Yves Bouget<br>Alexandra Z. Worden | Manuel Ares<br>Jian Guo<br>Jean-Claude Lozano<br>Lisa Sudek<br>Valérie Vergé | https://www.protocols.io/view/plasmid-dnas-designed-for-expression-in-micromonas-i9wch7e | |
| *Tetraselmis striata* | KAS-836 | Heriberto Cerutti<br>Thomas Clemente | Patrick Beardslee<br>Fulei Luan<br>Xiaoxue Wen | http://dx.doi.org/10.17504/protocols.io.hjtb4nn | GenBank (KY886895) |
| *Pyramimonas parkeae* | SCCAP K-0007 | Vladimír Hampl | Natalia Ewa Janowicz<br>Anna M.G. Novák Vanclová | https://www.protocols.io/view/protocols-for-mrna-electroporation-hh4b38w<br>https://www.protocols.io/view/nucleofection-of-pyramimonas-parkeae-chromera-veli-ibucanw<br>https://www.protocols.io/view/biolistic-transformation-experiment-on-eutreptiell-ibvcan6 | |

| | | | | | |
|---|---|---|---|---|---|
| **Haptophytes** | | | | | |
| *Isochrysis galbana* | CCMP 1323 | Colin Brownlee | Cecilia Balestreri<br>Andrea Highfield<br>Rowena Stern<br>Glen Wheeler | https://www.protocols.io/view/biolistic-transformation-of-isochrysis-galbana-2pugdnw<br>https://www.protocols.io/view/method-for-electroporation-of-isochrysis-galbana-c-hmab42e | GenBank<br>(MK903009 - pIgNAT construct)<br>(MK903010 -PCR product of transgene) |
| *Emiliania huxleyi* | CCMP 1516 | Colin Brownlee | Cecilia Balestreri<br>Andrea Highfield<br>Rowena Stern<br>Glen Wheeler | http://dx.doi.org/10.17504/protocols.io.8tzhwp6 | |
| **Rhizarians** | | | | | |
| *Amorphochlora (Lotharella) amoebiformis* | CCMP 2058 | Yoshihisa Hirakawa | Kodai Fukuda | http://dx.doi.org/10.17504/protocols.io.35hgq36 | |
| *Bigelowiella natans* | CCMP 2755 | Vladimír Hampl | Natalia Ewa Janowicz<br>Anna M.G. Novák Vanclová | https://www.protocols.io/view/protocols-for-mrna-electroporation-hh4b38w<br>https://www.protocols.io/view/nucleofection-of-pyramimonas-parkeae-chromera-veli-ibucanw<br>https://www.protocols.io/view/biolistic-transformation-experiment-on-eutreptiell-ibvcan6 | |
| **Stramenopiles** | | | | | |
| *Fragilariopsis cylindrus* | CCMP 1102 | Thomas Mock | Amanda Hopes | http://dx.doi.org/10.17504/protocols.io.z39f8r6<br>https://www.protocols.io/view/biolistic-transformation-of-polar-diatom-fragilari-z39f8r6 | |
| *Thalassiosira pseudonana* | CCMP 1335 | Christopher L. Dupont<br>Pamela Silver | Jernej Turnsek | http://dx.doi.org/10.17504/protocols.io.jfncjme<br>http://dx.doi.org/10.17504/protocols.io.nbzdap6<br>http://dx.doi.org/10.17504/protocols.io.7ghhjt6 | |
| *Seminavis robusta* | DCG 0498 (D6)<br>DCG 0514 (VM3-4) | Aaron Turkewitz | Luke Noble<br>Matthew Rockman<br>Lev Tsypin | http://dx.doi.org/10.17504/protocols.io.4p8gvrw | |
| *Pseudo-nitzschia multiseries* | MLML-EBL culture collection, strain 15091C3<br>Unavailable due to culture collapse (after 3 years of cultivation), but DNA and RNA stocks are available | G. Jason Smith | Deborah Robertson<br>April Woods | http://dx.doi.org/10.17504/protocols.io.7vhhn36 | http://dx.doi.org/10.17504/protocols.io.7vhhn36 |

| | | | | | |
|---|---|---|---|---|---|
| *Heterosigma akashiwo* | CCMP 2393 | Kathryn Coyne | Pamela Green | http://dx.doi.org/10.17504/protocols.io.4qggvtw<br>http://dx.doi.org/10.17504/protocols.io.4qhgvt6<br>https://www.protocols.io/view/modified-genomic-dna-extraction-method-for-heteros-himb4c6<br>https://www.protocols.io/view/modified-total-rna-extraction-for-heterosigma-akas-hipb4dn<br>https://www.protocols.io/view/efforts-to-transform-heterosigma-akashiwo-using-an-4ytgxwn | |
| *Aurantiochytrium limacinum* | ATCC MYA-1381 | Jackie Collier | Joshua Rest<br>Mariana Rius | http://dx.doi.org/10.17504/protocols.io.8xyhxpw<br>http://dx.doi.org/10.17504/protocols.io.hg6b3ze<br>http://dx.doi.org/10.17504/protocols.io.pgtdjwn | https://www.addgene.org/Jackie_Collier/<br>(for pUC19_GZG, pUC19_18GZG) |
| *Caecitellus sp.* | Unavailable due to culture collapse | Patrick Keeling | Elisabeth Hehenberger<br>Nicholas A. T. Irwin | https://www.protocols.io/view/electroporation-of-caecitellus-sp-with-fitc-dextra-35kgq4w | |
| *Nannochloropsis oceanica* | CCMP 1779 | Peter von Dassow | Fernan Federichi<br>Isaac Nuñez<br>Tamara Matute<br>Albane Ruaud<br>Jorge Ibañez | http://dx.doi.org/10.17504/protocols.io.7r8hm9w<br>http://dx.doi.org/10.17504/protocols.io.h3nb8me | https://doi.org/10.5281/zenodo.3463694 |
| *Phaeodactylum tricornutum* | CCAP1055/1 | Andrew E. Allen | Mark Moosburner<br>Chris Bowler | http://dx.doi.org/10.17504/protocols.io.4abgsan<br>http://dx.doi.org/10.17504/protocols.io.4acgsaw<br>http://dx.doi.org/10.17504/protocols.io.4bmgsk6<br>http://dx.doi.org/10.17504/protocols.io.7gihjue | http://dx.doi.org/10.17504/protocols.io.7gnhjve |
| **Alveolates** | | | | | |
| *Euplotes crassus* | CCAP 1624/31 | Cristina Miceli | Rachele Cesaroni<br>Lawrence A. Klobutcher<br>Mariusz Nowacki<br>Angela Piersanti<br>Sandra Pucciarelli<br>Estienne Swart | https://www.protocols.io/view/euplotes-miceli-lab-2a8gahw/protocols | |
| *Euplotes focardii* | CCAP 1624/34 | Cristina Miceli | Angela Piersanti<br>Sandra Pucciarelli | https://www.protocols.io/view/euplotes-miceli-lab-2a8gahw/protocols | |
| *Chromera velia* | CCMP 2878 | Vladimír Hampl | Natalia Ewa Janowicz<br>Anna M.G. Novák Vanclová | https://www.protocols.io/view/protocols-for-mrna-electroporation-hh4b38w | |

| | | | | | |
|---|---|---|---|---|---|
| | | | | https://www.protocols.io/view/nucleofection-of-pyramimonas-parkeae-chromera-veli-ibucanw<br>https://www.protocols.io/view/biolistic-transformation-experiment-on-eutreptiell-ibvcan6 | |
| *Perkinsus marinus*[2] | ATCC PRA240 | José A. Fernández Robledo<br>Senjie Lin<br>Ross Waller | Duncan B. Coles<br>Elin Einarsson<br>Nastasia J. Freyria<br>Sebastian Gornik<br>Imen Lassadi<br>Arnab Pain | https://www.protocols.io/view/oyster-parasite-perkinsus-marinus-transformation-u-gv9bw96<br>https://www.protocols.io/view/glass-beads-based-transformation-protocol-for-perk-g36byre<br>https://www.protocols.io/view/fluorescence-activated-cell-sorting-facs-of-perkin-hh2b38e<br>https://www.protocols.io/view/golden-gate-plasmids-used-for-transfection-of-perk-37egrje | Genebank (EF632302, EF632303, KX423758–60) |
| *Oxyrrhis marina* | CCMP 1788<br>CCMP 1795 | Patrick Keeling<br>Claudio Slamovits<br>Senjie Lin | Elizabeth C. Cooney<br>Nicholas A. T. Irwin<br>Elisabeth Hehenberger<br>Yoshihisa Hirakawa<br>Brittany Sprecher<br>Lu Wang<br>Huan Zhang | https://www.protocols.io/view/transfection-of-alexa488-labelled-dna-into-oxyrrhi-ha8b2hw<br>https://www.protocols.io/view/calcium-phosphate-transfection-of-oxyrrhis-marina-ha4b2gw<br>https://www.protocols.io/view/electroporation-transformation-of-fitc-dextran-int-3cmgiu6<br>https://www.protocols.io/view/Dinoflagellate-transformation-e6bbhan<br>https://www.protocols.io/view/co-incubation-protocol-for-transforming-heterotrop-7pphmmn<br>https://www.protocols.io/view/electroporation-of-oxyrrhis-marina-vcne2ve<br>https://www.protocols.io/view/co-incubation-protocol-for-transforming-heterotrop-hmzb476 | |
| *Hematodinium sp.* | Submitted to ATCC collection (in the meantime please contact Waller´s lab if interested) | Ross Waller | Sebastian Gornik<br>I lan Hu<br>Imen Lassadi<br>Arnab Pain | https://www.protocols.io/view/plasmid-used-for-transfection-trails-of-hematodini-4nigvce | |

| | | | | | |
|---|---|---|---|---|---|
| *Fugacium (Symbiodinium) kawagutii* | CCMP 2468 | Senjie Lin | Brittany Sprecher<br>Lu Wang<br>Huan Zhang | https://www.protocols.io/view/dinoflagellate-transformation-7prhmm6 | |
| *Alexandrium catenella* | CCMP BF-5 | Senjie Lin | Brittany Sprecher<br>Lu Wang<br>Huan Zhang | https://www.protocols.io/view/dinoflagellate-transformation-7prhmm6<br>https://www.protocols.io/view/nucleofector-protocol-for-dinoflagellates-using-lo-7n8hmhw | |
| *Karlodinium veneficum* | CCMP 1975 | Senjie Lin | Brittany Sprecher<br>Huan Zhang | https://www.protocols.io/view/dinoflagellate-transformation-7prhmm6<br>https://www.protocols.io/view/nucleofector-protocol-for-dinoflagellates-using-lo-7n8hmhw | |
| *Breviolum* (*Symbiodinium*) sp. | NIES-4271 | Jun Minagawa | Yuu Ishii<br>Konomi Kamada<br>Shinichiro Maruyama | https://www.protocols.io/view/electroporation-of-fluorescein-into-the-coral-symb-hdcb22w | https://www.protocols.io/view/dna-construct-for-genetic-transformation-of-the-co-7udhns6/document |
| *Crypthecodinium cohnii* | CCMP 316 | José Fernández Robledo<br>Ross Waller | Duncan B. Coles<br>Nastasia J. Freyria<br>Paulo A. García<br>Imen Lassadi | https://www.protocols.io/view/transfection-of-crypthecodinium-cohnii-using-label-z26f8he | Probe amplified from EF632302 |
| *Amphidinium carterae* | CCMP 1314 | Christopher Howe | Adrain Barbrook<br>Isabel Nimmo<br>Ellen Nisbet | http://dx.doi.org/10.17504/protocols.io.4r2gv8e<br>https://www.protocols.io/view/biolistic-transformation-of-amphidnium-hnmb5c6 | |
| **Discobans (Euglenozoans and Heteroloboseans)** | | | | | |
| *Bodo saltans* | submitted to ATCC collection | Virginia Edgcomb | Miguel A. Chiurillo<br>Roberto Decampo<br>Fatma Gomaa<br>Noelia Lander<br>Zhuhong Li | http://dx.doi.org/10.17504/protocols.io.s5peg5n<br>http://dx.doi.org/10.17504/protocols.io.s5meg46<br>http://dx.doi.org/10.17504/protocols.io.s5jeg4n<br>http://dx.doi.org/10.17504/protocols.io.sh4eb8w<br>http://dx.doi.org/10.17504/protocols.io.sh6eb9e | http://dx.doi.org/10.17504/protocols.io.7fchjiw<br><br>GenBank (MN608152) |
| *Diplonema papillatum* | ATCC 50162 | Julius Lukeš | Drahomíra Faktorová<br>Ambar Kachale<br>Binnypreet Kaur<br>Getraud Burger<br>Matus Valach | https://www.protocols.io/groups/julius-lukes<br>https://dx.doi.org/10.17504/protocols.io.4digs4e | GenBank (MN047315) |
| *Eutreptiella gymnastica* | SCCAP K-0333 | Vladimír Hampl | Natalia Ewa Janowicz<br>Anna M.G. Novák Vanclová | https://www.protocols.io/view/protocols-for-mrna-electroporation-hh4b38w | |

| | | | | | |
|---|---|---|---|---|---|
| | | | | https://www.protocols.io/view/nucleofection-of-pyramimonas-parkeae-chromera-veli-ibucanw https://www.protocols.io/view/biolistic-transformation-experiment-on-eutreptiell-ibvcan6 | |
| *Naegleria gruberi* | ATCC 30224 | Anastasios Tsaousis | Veronica Freire-Beneitez Eleanna Kazana Jan Pyrih Tobias von der Haar | http://dx.doi.org/10.17504/protocols.io.hnhb5b6 http://dx.doi.org/10.17504/protocols.io.hpub5nw http://dx.doi.org/10.17504/protocols.io.hpvb5n6 | http://dx.doi.org/10.17504/protocols.io.7w4hpgw |
| **Opisthokonts** | | | | | |
| *Pirum gemmata* | ATCC PR-280 | Elena Casacuberta Iñaki Ruiz-Trillo | Cristina Aresté | http://dx.doi.org/10.17504/protocols.io.z5nf85e | |
| *Sphaeroforma arctica* | ATCC PRA-297 | Elena Casacuberta Iñaki Ruiz-Trillo | Cristina Aresté Omaya Dudin | http://dx.doi.org/10.17504/protocols.io.z5nf85e http://dx.doi.org/10.17504/protocols.io.z6ef9be | |
| *Abeoforma whisleri* | ATCC PRA-279 | Elena Casacuberta Iñaki Ruiz-Trillo | Elena Casacuberta Cristina Aresté Sebastián Najle | https://www.protocols.io/view/abeoforma-whisleri-transient-transfection-protocol-zexf3fn http://dx.doi.org/10.17504/protocols.io.zexf3fn | |
| *Salpingoeca rosetta* | ATCC PRA-390 | Nicole King | David Booth Monika Sigg | http://dx.doi.org/10.17504/protocols.io.h68b9hw | |

-American Type Culture Collection (ATCC) (https://www.atcc.org/)
-Culture Collection of Marine Phytoplankton (CCMP) now The Provasoli-Guillard National Center for Marine Algae and Microbiota (NCMA) (https://ncma.bigelow.org/cms/index/index/)
-Kuehnle AgroSystems Inc (KAS) (http://www.kuehnleagro.com/)
-Scandinavian Culture Collection of Algae and Protozoa (SCCAP) (http://www.sccap.dk/dk/soeg/detaljer.asp?Cunr=K-0007)
-Microbial Culture Collection at the National Institute for Environmental Studies (NIES Collection, Tsukuba, JAPAN) (https://mcc.nies.go.jp/)
-Culture collection of the BCCM-DCC (http://bccm.belspo.be/about-us/bccm-dcg)
-Culture collection MLML-EBL (https://www.mlml.calstate.edu/ebl/)

Details of protocol for particular species:

[1]*Micromonas commoda*: Acclimated mid-exponential M. commoda cells grown in L1 medium at 21 oC were spun at 5000 x g for 10 min, the pellet was resuspended in Buffer SF (Lonza) premixed with carrier DNA (pUC19) and plasmid, and 3 x 107 cells were used per reaction. After applying the EW-113 pulse, 100 µl of ice-cold recovery buffer (10 mM HEPES-KOH, pH 7.5; 530 mM sorbitol; 4.7% [w/v] PEG 8000) was added to each well and incubated for 5 min at room temperature. Each reaction was then transferred into 2 ml L1, placed at 21 °C and light was increased stepwise over 72 h.

[2]*Perkinsus marinus*: In brief, a newly formulated transformation 3R buffer (200 mM Na2HPO4; 70 mM NaH2PO4; 15 mM KCl; 1.5 mM CaCl2; 150 mM HEPES-KOH, pH 7.3) was used to reduce the cost of electroporation. 5 x 107 cells were resuspended in 330 µl of fresh ATCC Medium 1886 and were mixed with 5.0 µg of linearized and circular [1:1] plasmid and 300 µl of glass beads (Sigma) in a 1.5 ml tube, vortexed for 30 s at maximum speed, and cells in 500 µl of culture medium were transferred to 6-well plates in a final volume of 3 ml.

[3]*Bodo saltans*: Square-wave electroporation (Nepa21) was used with a poring pulse of 250V (25 ms) and 5 transfer pulses of 60V (99 ms) in the presence of Cytomix buffer (120 mM KCl; 0.15 mM CaCl2; 10 mM KH2PO4; 2 mM EGTA; 5 mM MgCl2; 25 mM HEPES-KOH, pH 7.6).
